# All-*trans* retinoic acid enhances the anti-tumour effects of fimaporfin-based photodynamic therapy

**DOI:** 10.1101/2022.04.28.487643

**Authors:** Judith Jing Wen Wong, Susanne Lorenz, Pål Kristian Selbo

## Abstract

The vitamin A metabolite all-trans retinoic acid (ATRA; tretinoin) has anticancer potential. However, lack of clinical success has prevented its approval for solid tumours. Herein, we propose combining short-term low-dose ATRA preconditioning with fimaporfin-based photodynamic therapy (ATRA+PDT) for the improved treatment of solid cancers. Compared to monotherapies, ATRA+PDT induced synergistic cytotoxic responses including promotion of apoptosis in colon and breast carcinoma cell lines. Neither enhanced activity of alkaline phosphatase (ALP) nor increased expression of CD133 was detected after ATRA treatment indicating that ATRA+PDT cause cell death independent of differentiation. In the human colorectal adenocarcinoma cell line HT-29, the effect of ATRA+PDT on gene expression was evaluated by RNA sequencing (RNA-seq). We identified 1129 differentially expressed genes (DEGs) after ATRA+PDT compared to PDT. Ingenuity Pathway Analysis (IPA) predicted the unfolded protein response (UPR), interferon (IFN) signaling and retinoic acid-mediated apoptosis signaling as strongly activated canonical pathways after ATRA+PDT compared to PDT. A validation of the RNA-sec data by RT-qPCR revealed that ATRA+PDT elevated mRNA expression of early growth response 1 (EGR1) and strongly the stress-induced activating transcription factor 3 (ATF3), of which was confirmed on the protein level. In addition, ATRA+PDT abolished mRNA expression of regenerating islet-derived protein 4 (REG4). During the first 20 days post-ATRA+PDT, we obtained significant anti-tumour responses in HT-29 xenografts, including complete responses in 2/5 mice. In conclusion, ATRA+PDT represent a novel combination therapy for solid tumours that should be further tested in immunocompetent preclinical models.

## Introduction

ATRA is derived from vitamin A and plays a crucial role in biological processes, including embryonic development, immune responses, and vision, through binding and activation of the nuclear retinoic acid receptors RARα, RARβ, and RARγ [1]. Transcriptional activation by RARs induce differentiation, cell cycle arrest, anti-proliferative, and apoptosis-inducing effects [2], which make retinoids attractive in cancer therapy. As one of the first targeted cancer therapies, ATRA is successfully used in the differentiation and apoptosis induction therapy of acute promyelocytic leukaemia [1, 3] and is one of the first precision targeted therapies. Based on this, ATRA is currently being studied to prevent or treat carcinoma [4]. However, lack of effects in solid tumours has limited clinical translation of ATRA due to several factors: ATRA formulation and administration is challenging due to its low aqueous solubility. In addition, ATRA is oxidized when exposed to heat, UV light, or oxygen [4] and has a very short plasma half-life (∼45 min) [5]. Moreover, RARβ expression is frequently lost in solid tumours, or the RAR promoter is epigenetically silenced, resulting in ATRA resistance [6, 7]. Thus, there is a high need for new ATRA drug delivery or combinatorial strategies to overcome resistance and improve the survival of patients with solid tumours.

PDT is used clinically for the treatment of different solid cancers and is based on three non-toxic components: photosensitiser, light, and oxygen. Upon light activation of the photosensitiser, energy is transferred to molecular oxygen, resulting in generation of cytotoxic reactive oxygen species (ROS), of which singlet oxygen is the most common [8]. The use of differentiation-promoting agents, e.g., preconditioning with vitamin D, to enhance PDT efficacy was first demonstrated by Ortel *et al.* to improve 5-aminolevulinic acid (ALA)-based PDT. However, ATRA was only shown to slightly enhance the photosensitising compound PpIX production compared to ALA alone (non-significant) and no enhanced cytotoxic effects was demonstrated [9]. Currently, there is a lack of mechanistic evidence showing that ATRA preconditioning improves PDT of cancer.

In this study, we hypothesized that combining ATRA and PDT (ATRA+PDT) may exert stronger anti-carcinoma effects than either treatments alone. To address this, we aimed to investigate the potential anti-cancer effects of short-term low-dose ATRA combined with the endosome/lysosome targeting drug fimaporfin (TPCS_2a_) as the photosensitiser followed by controlled light exposure. Fimaporfin/TPCS_2a_ is used in the drug delivery technology photochemical internalization (PCI) [10], and has been tested clinically in combination with bleomycin [11] and vaccine peptide/protein-based antigens [12]. Recently, PCI in combination with gemcitabine was shown to be safe and effective in a Phase I study for the treatment of unresectable extrahepatic cholangiocarcinoma [13]. Here we show for the first time that combining ATRA+PDT results in synergistic cytotoxic responses in ATRA resistant colon and breast carcinoma cell lines compared to ATRA or PDT monotherapy. Mechanistically, we demonstrate that ATRA+PDT induce cell cycle arrest and augment cytotoxic responses including apoptotic cell death. Pathway analysis of the RNA-seq data confirmed apoptosis signaling and identified activation of the unfolded protein response (UPR). *In vivo,* one single treatment with ATRA+PDT induced significant anti-tumour effects during the first 20 days compared to no treatment in HT-29 tumour-bearing athymic nude mice, and ATRA+PDT was the only group that achieved complete responses.

## Materials and Methods

### Cell lines and culture conditions

The cell lines, HT-29, HCT116, SKBR3, and MDA-MB-231, were obtained from the American Type Culture Collections (ATCC, Manassas, VA, USA). The MC-38 (ENH204-FP) cell line was obtained from Kerafast (Boston, MA, USA). The HT-29, SKBR3 and HCT116 cells were cultivated in McCoy’s 5a medium (M9309, Sigma-Aldrich, St Louis, MO, USA), the MDA-MB-231 cells in RPMI-1640 (R8758, Sigma-Aldrich), and the MC-38 cells in DMEM (BE12-604F/U1, Lonza Group, Basel, Switzerland) supplemented with 10 mM HEPES (H0887, Sigma-Aldrich). All culture media were supplemented with 10% FBS (ThermoFisher Scientific, Waltham, MA, USA), 100 U/mL penicillin and 100 µg/mL streptomycin (Sigma-Aldrich). All cell lines were mycoplasma negative throughout the experimental period and maintained in a humidified incubator at 37 °C with 5% CO2. All cell lines were in all experiments used at low passage numbers (≤25).

### Drugs and chemicals

ATRA (Sigma-Aldrich, Saint-Louis, MO, USA) was dissolved in DMSO was stored at −80°C. The photosensitiser fimaporfin/TPCS_2a_ (in polysorbate 80, 2.8% mannitol, 50 mM Tris, pH 8.5) was provided by PCI Biotech AS (Oslo, Norway) and stored at 4 °C. All work with ATRA or TPCS_2a_ was performed under subdued light.

### *In vitro* ATRA+PDT treatment protocol

To evaluate the treatment effects of ATRA+PDT, 3000 (HCT116 and HT-29) or 8000 (SKBR3) cells/well were seeded in 96-well plates (Nunc). The cells were first incubated with 0.1 µM ATRA for 24 hours and further incubated with 0.1 µM ATRA and/or 0.4 µg/mL TPCS_2a_ for 18 hours. Before light exposure, the cells were washed twice with PBS, chased for 4 hours in fresh media, and illuminated with LumiSource (PCI Biotech, Oslo, Norway) [14]. Cell viability was evaluated 48 hours post-treatment. To evaluate ATRA+PDT response, MC-38 cells (3000 cells/well) were treated using 10 µM ATRA and 0.6 µg/mL TPCS_2a_. The MC-38 cells were first incubated for 24 hours with ATRA. The medium with ATRA was then replaced with new medium containing both ATRA and TPCS_2a_ and the cells were further incubated for 18 hours. For ATRA+PDT in MDA-MB-231 cells (3000 cells/well), 10 µM ATRA was co-incubated with 0.4 µg/mL TPCS_2a_ for 18 hours. Prior to light exposure, the cells were washed with PBS twice, chased for 4 hours in fresh media and illuminated. MC-38 and MDA-MB-231 viability was assessed 24 and 48 hours post-light, respectively.

Opposite to preconditioning the cells with ATRA before PDT we also aimed to assess ATRA incubation after PDT. HT-29 (pre-incubated with 0.4 µg/mL TPCS_2a_) and MC-38 (pre-incubated with 0.6 µg/mL TPCS_2a_) cells were incubated with 1 µM and 10 µM ATRA, respectively, for 48 hrs directly after PDT.

### Evaluation of treatment-induced cytotoxicity and effect on proliferation

The cell viability was assessed by the MTT assay as previously described [15]. Clonogenic assay was performed in 6-well plates (500 cells/well). The cells were allowed to attach and treated as described above. When sufficiently large colonies (>50 cells/colony) were formed in controls, the colonies were fixed and stained [14]. Real-time monitoring of cell proliferation was evaluated using the IncuCyte live-cell imaging system (Essens Bioscience, MI, USA). The cells were treated and immediately after light exposure placed in an incubator connected to the IncuCyte imaging system.

To evaluate the formation of potential phototoxic products generated by ATRA+PDT, 0.1 µM ATRA and 0.4 µg/ml TPCS_2a_ was first exposed to light in a cell-free system. Both ATRA and TPCS_2a_ was prepared in McCoy’s 5a culture medium, added in a 24-well plate (500 µL/well) and exposed to light at indicated time-points. HT-29 cells were subsequently incubated with the culture medium containing light-exposed ATRA and/or TPCS_2a_ in 96-well plates for 48 hours before viability was measured using the MTT assay.

### Detection of intracellular ROS

The cell permeant reagent 2’,7’-dichlorofluorescin diacetate (DCF-DA) and its downstream product 2’,7’-dichlorofluorescein (DCF) was used to quantitatively assess ROS by flow cytometry in live cell samples as previously described [16]. To evaluate generation of intracellular ROS post-treatments, 150 000 (HCT116 and HT-29) or 200 000 (SKBR3) cells/well were seeded in 6-well plates and treated as indicated. The cells were subjected to light exposure and immediately after treatment cells were detached using trypsin. The cells were washed with PBS and untreated CellTrace™ Violet-stained cells (ThermoFisher Scientific, 0.5 µM, 37 °C for 30 min) were added to all samples to serve as an internal control. The samples were incubated with 20 µM dichloro-dihydro-fluorescein diacetate (DCFH-DA) (Sigma-Aldrich) at 37 °C for 1 hour, washed and subsequently analysed on a flow cytometer described below.

### ALP assay

ALP activity in cells was measured using a fluorometric assay kit (ab83371, Abcam, Cambridge, UK) according to manufacturer’s instructions. In brief, 1.5×10^5^ cells/well (SKBR3) or 5×10^4^ cells/well (HCT116 and HT-29) were seeded in 12-well plates, allowed to attach and incubated with indicated ATRA concentrations for 42 hours. At the end of the incubation, the cells were washed with ice-cold PBS once and harvested using 100 µL assay buffer/well. A non-fluorescent substrate, 4-methylumbelliferyl phosphate disodium (MUP) salt, was added to the samples. MUP is converted to a fluorescent product (4-MU) by ALP, which was measured after 30 min reaction time at ambient temperature. A stop solution was added and fluorescence was measured using the Tecan Spark™ 10M microplate reader (Zürich, Switzerland). 4-MU was excited at 360 nm and fluorescence was detected at 440 nm. The fluorescence intensity was calculated relative to protein concentration in each sample as assessed by the DC™ Protein Assay (BioRad, Hercules, CA, USA) according to manufacturer’s protocol.

### Evaluation of CD133 expression

To assess the CD133 expression after ATRA treatment, 1.5×105 cells/well (HCT116 and HT-29) or 2×105 cells/well (SKBR3) were seeded in 6-well plates, allowed to attach and incubated with indicated ATRA concentrations for 42 hours. At the end of treatment, the cells were detached with trypsin, washed once and incubated with 0.5% BSA in PBS for 10 min on ice. The cells incubated with the primary antibody anti-CD133 (12.3 µg/mL, HB#7, Developmental Studies Hybdridoma Bank, Iowa City, IA, USA) and the secondary antibody rabbit anti-mouse-FITC (1:50, #F0232, Dako) as previously described [17]. The cells were subsequently analysed using a flow cytometer described below.

### Apoptosis and cell cycle analysis analysed by flow cytometry

The TUNEL assay was used to evaluate apoptosis. Acute lymphocytic leukaemia cells, Reh, treated with 4 Gy and harvested 24 hours post-irradiation were included as a positive control. Treated cells were harvested, washed, resuspended in ice-cold methanol, and stored at least 1 hour at −20 °C before staining with Tdt reaction mix (Roche Diagnostics, Risch-Rotkreuz, Switzerland) containing biotin-16-dUTP (Roche) [16]. The cells were then washed, and incubated in streptavidin-Cy5 (GE Healthcare, Chicago, IL, USA) [16]. Finally, the cells were resuspended in Hoechst 33258 (1.5 µg/mL, Sigma Aldrich), incubated overnight at 4 °C and subsequently analysed by flow cytometry using the LSRII flow cytometer (Becton Dickinson, Franklin Lakes, NJ, USA). Cell cycle analyses, based on Hoechst stain, were performed using the Watson model in FlowJo version 7.6.5 (Treestar).

### RNA extraction

Total cellular RNA was extracted post-treatment in HT-29 cells (1.5×105/well) plated in 6-well plates (Nunc) at indicated time-points. The GenElute Mammalian Total RNA Miniprep Kit (Sigma-Aldrich) was used according to manufacturer’s instructions. RNA purity and quantity was measured using the Nanodrop2000 spectrophotometer (ThermoFisher Scientific). All samples were stored at −80 °C until further analysis.

### RNA-seq and differential gene analysis

Total RNA was isolated from three biological replicates of HT-29 cells treated with PDT or ATRA+PDT. The samples were harvested 3 hours post-120 seconds blue light exposure (∼LD_50_). The RNA integrity number (RIN) was evaluated using the 2100 Bioanalyzer (Agilent Technologies, Santa Clara, CA, USA), where all the samples had a RIN=10. Total RNA (500 ng) was processed using a TruSeq Stranded mRNA kit (Illumina, San Diego, CA, USA) following the manufacturer’s recommended procedures. Final libraries were quality controlled using RT-qPCR and pooled. Pooled libraries were sequenced on the NextSeq500 instrument (Illumina) with a HighOutput flow cell, single read sequencing of 75 bp read length, and dual-indexes (IDT UDI’s 2×8 bp). For data analysis, FASTQ files were mapped using Star2 aligner (STAR v2.5.0b) and human reference hg19. Counting, normalization, and differential gene expression analysis were performed using Cufflinks2 (v2.2.1).

### Heat maps

Heat maps of DEGs were generated using the web tool ClustVis (https://biit.cs.ut.ee/clustvis/) [18]. Unit variance scaling was applied to rows, both rows and columns were clustered using Euclidean distance and average linkage.

### STRING and STITCH interaction networks

Protein and functional interaction network was constructed using the STRING v11 online tool (https://string-db.org/) [19]. The drug-target gene interaction network of retinoic acid was constructed using the STITCH online tool (http://stitch.embl.de/) [20].

### Gene set enrichment analyses

Broad Institute’s GSEA software 4.1.0 was used to determine whether an *a priori* defined set of genes is represented as DEGs in ATRA+PDT compared to PDT [21, 22]. The Hallmark Gene sets were obtained from the Molecular Signature Database (MSigDb v7.4) [23]. False discovery rate (FDR) of ≤ 25% and p-value ≤ 0.05 was considered significant. Further analysis was performed using the IPA software (Qiagen, Hilden, Germany), which identify canonical pathways and biological functions or diseases significantly associated with the identified DEGs. The IPA software was used to analyse upstream regulators that are predicted to be relevant for the biological mechanisms underlying the data based on information gathered in the literature [24]. An overlap of p-value ≤ 0.01 and an activation z-score ≥ |2| was considered significant. The z-score is used to deduce likely activation states based on significant pattern matches of up- or downregulated genes. A positive z-score predicts activation, and a negative z-score indicates inhibition.

### Western blot analysis

HT-29 cells (1.5 ×105/well) were plated in 6-well plates (Nunc), ATRA+PDT-treated, and harvested for Western blot analysis by first placing the plate on ice. The cells were then washed and detached in PBS using a cell scraper. The cells were centrifuged (1000 g, 5 min) at 4 °C and the pellets containing cells were kept at −80 °C. Total proteins were then extracted using RIPA-lysis buffer on ice. Additionally, the lysis samples were sonicated, and centrifuged (12000 g, 15 min) at 4 °C. The protein concentration was assessed using the DC Protein Assay (Bio-Rad Laboratories). The samples were stored at −80 °C until further analysis. Proteins were separated by SDS-PAGE and expression was detected using the following antibodies: ATF3 (sc-81189, Santa Cruz Biotechnology, Dallas, TX, USA), cleaved caspase-3 (#9664, Cell Signaling Technology, Danvers, MA, USA) and γ-tubulin (#5886, Cell Signaling Technology). HRP-linked rabbit (#7074, Cell Signaling Technology) and HRP-linked mouse (#7076S, Cell Signaling Technology) antibodies were used as secondary antibodies. SuperSignal™ West Dura Extended Duration Substrate (Sigma-Aldrich) was used according to the manufacturer’s instruction. Protein bands were detected using the ChemiDoc system (BioRad) with the ImageLab 4.1 software (Bio-Rad). Protein expression was quantified relative to the loading control.

### Animal studies

All animal procedures were performed according to protocols approved by the national animal research authority Mattilsynet (FOTS ID22020) and were conducted according to the regulations of the Federation of European Laboratory Animal Science Association (FELASA) Handling of and experiments with animals were therefore performed in compliance with EUs Directive 2010/63/EU on the protection of animals used for scientific purposes. The mice were on average 20–25 g (6–8 weeks old, n=5 per experimental group) and given a unique number by ear marking at the start of the experiments. HT-29 xenografts (2.5×10^6^ cells, 30 µL) were established intradermally in the flank of female HSD athymic Nude-Foxn1nu mice bred at the Department of Comparative Medicine at the Norwegian Radium Hospital, Oslo University Hospital. An intradermal tumour model was selected as it provides better control of intratumoral injections and improved light penetration for photosensitiser activation. When the tumours reached ∼70-150 mm^3^, the animals were randomized to different treatment groups. Two protocols were evaluated. In the first protocol (systemic), the ATRA+PDT group received five doses of ATRA prior to light treatment. 10 mg/kg ATRA (R2625, Sigma-Aldrich) in corn oil (C8267, Sigma-Aldrich) was delivered intraperitoneally once a day for five days. TPCS_2a_ (5 mg/kg) was delivered as a single dose intravenously three days prior to light exposure. In the second protocol (intratumoral), TPCS_2a_ (20 µg) and/or ATRA (0.3 µg) were dissolved in a solution containing polysorbate 80, 2.8% mannitol, 50 mM Tris, pH 8.5 and delivered as a single dose intratumorally (30 µL) one day prior to light exposure. In both the systemic and intratumoral ATRA+PDT protocols, the animal was anaesthetized with sevoflurane inhalation during light exposure, placed on a heating pad and covered in aluminium foil except for the tumour area. The tumours were illuminated using a diode laser (652 nm) equipped with an optical fibre with a microlens at the end (Medlight SA, Ecublens, Switzerland) (CeramOptec, Bonn, Germany). The mice received a light dose of 15 and 10 J/cm^2^, in the systemic and intratumoral protocol, respectively, using an irradiance of 90 mW/cm^2^. All mice that received TPCS_2a_ were kept in subdued light for a week post-illumination to avoid phototoxicity. The tumour volume and whole body weight of the animals were monitored 2-3 times a week as previously described [14]. Humane endpoints were set at weight loss ≥ 20% and/or tumour size ≥ 1000 mm^3^ or 90 days post-treatment. The animals were euthanized by cervical dislocation.

### Statistical Analysis and synergy calculations

Statistical analyses were performed using SigmaPlot 14.0 (Systat Software, San Jose, CA, USA). Statistical significance was set at p≤0.05. Survival analysis was performed by pairwise log-rank (Mantel-Cox) comparison using SPSS Statistics 26 (IBM, Armonk, NY, USA). The synergy was evaluated using the synergy/antagonism parameter difference in log (DL) and evaluated if the DL parameter is statistically significant from zero [25]. The DL parameter is based on the difference in the observed combined effect and the theoretical additive effect. A statistically significant positive DL value indicates synergism, and a negative value indicates antagonism. A DL value close to zero suggests additive effects. We evaluated the DL values based on the MTT data for HT-29 and SKBR3.

## Results

### ATRA+PDT enhances cytotoxic responses in breast and colon cancer cells

We evaluated the cytotoxic response of 0.01 to 100 µM ATRA monotherapy for 48 hours in SKBR3, HCT116, and HT-29 cells. For all cell lines, the viability was > 50% at ATRA concentrations < 50 µM (**Supplementary Fig. 1**). While the physiological concentration of ATRA is in the nanomolar range in human plasma [26], pharmacological doses of ATRA results in plasma concentrations in the micromolar range [27]. Thus, a therapeutic relevant concentration of 0.1 µM ATRA was selected for the subsequent experiments and combined with fimaporfin-PDT (ATRA+PDT). We examined the cytotoxic effects of ATRA+PDT in five different cancer cell lines; two human colon cancer cell lines: HCT116 (ATRA-resistant) and HT-29 (moderately ATRA-resistant) [28] and the MC-38 mouse colon cancer cell line. Two human breast cancer cell lines were also included: SKBR3 (ATRA-sensitive) [29] and MDA-MB-231 (ATRA-resistant) [30].

The cells were preconditioned with 0.1 µM ATRA and subjected to fimaporfin (TPCS2_a_)-PDT. In contrast to HCT116, enhanced cytotoxicity was observed in SKBR3 and HT-29 cells (**Fig. 1A**) and in MC-38 and MDA-MB-231 (**Supplementary Fig. 2**). The dual P13K and mTOR inhibitor NVP-BEZ235 augments ATRA efficacy in leukemic cells [31] and was therefore tested in combination with ATRA+PDT. However, no additional effect was found when combining 0.1 µM ATRA+PDT with NVP-BEZ235 at concentrations ranging from 5-100 nM in HT-29 and HCT116 (**data not shown**). Changing the treatment sequence where PDT was performed prior to ATRA incubation did not have any enhancement effect on the cell viability, indicating the importance of preconditioning with ATRA **(Supplementary Fig. 3)**.

**Fig. 1:**
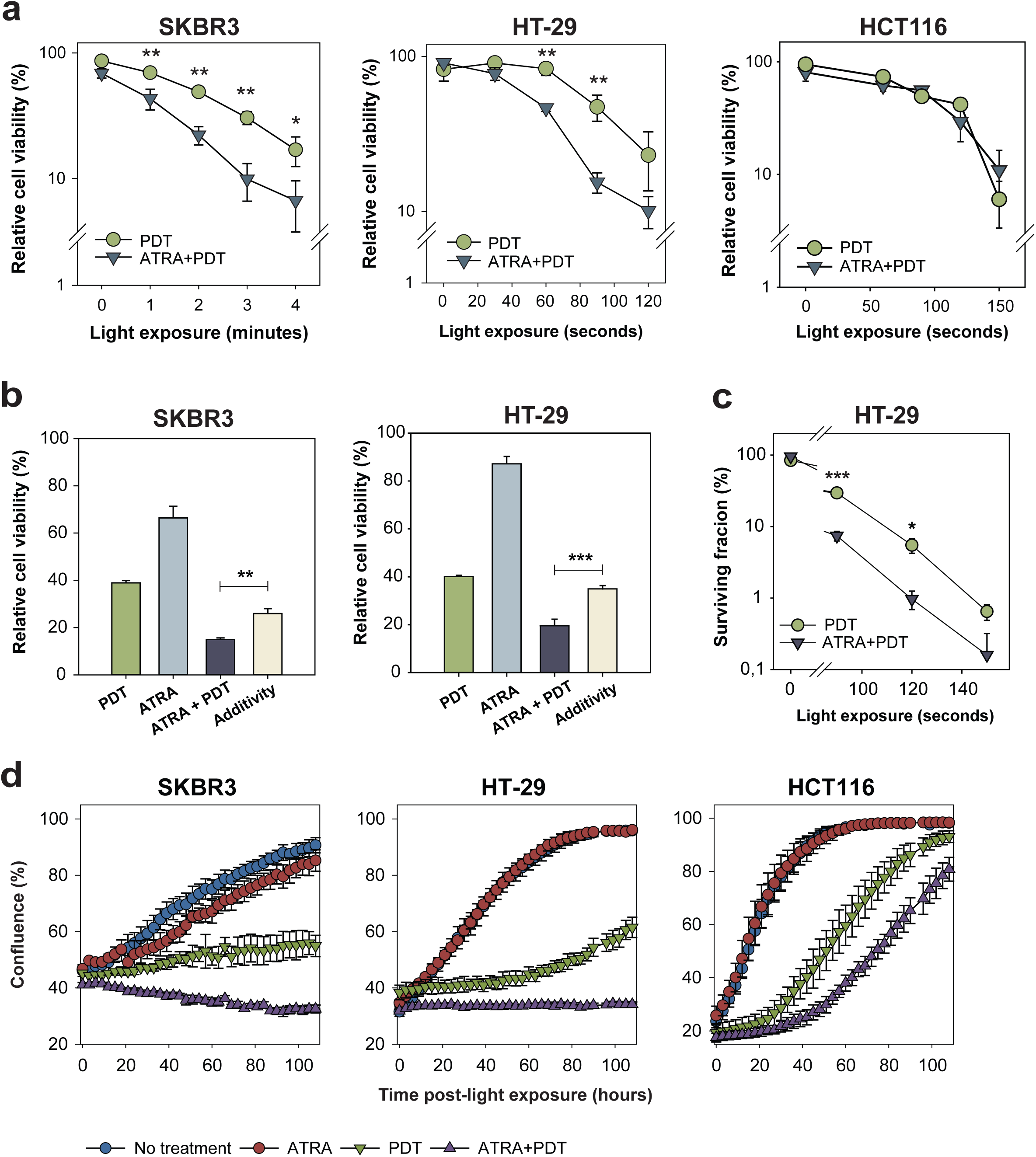
ATRA preconditioning enhances the cytotoxic efficacy of fimaporfin-PDT and strongly impairs proliferation in carcinoma cells. **(A)** Viability assessed by MTT 48 hours post-light exposure in SKBR3, HT-29, and HCT116 cells treated with fimaporfin-PDT or the combination ATRA+PDT. Representative data of at least three independent experiments. Data presented as mean of triplicates ± S.D. 60 seconds light exposure ≈ 0.58 J/cm^2^. **(B)** Synergistic cytotoxic effects induced by ATRA+PDT in HT-29 and SKBR3 cells. The observed effect of the combination in SKBR3 and HT-29 compared to a theoretical additive effect at ∼ LD_60_ of at least three independent experiments. Data presented as mean ± S.E. **(C)** Representative clonogenic assay of PDT compared to ATRA+PDT in HT-29 cells. **(D)** Proliferation of SKBR3, HT-29 and HCT116 up to 110 hours post-120 seconds light exposure (1.16 J/cm^2^). Representative experiment of three experiments, data presented as mean of triplicates ± S.E. Statistical significance calculated using Student’s two-tailed t-test (*** p ≤ 0.001, ** p ≤ 0.01, * p ≤ 0.05).

To further evaluate whether the cytotoxic effects of ATRA+PDT was synergistic in HT-29 and SKBR3, the observed effect was compared to the theoretical additive effect. A significant difference was found indicating synergy between ATRA and PDT and supported with synergy calculations (**Fig. 1B**). In both cell lines, significant DL values were found indicating synergistic cytotoxic effects; DLSKBR3 = 0.310±0.06 (p = 0.04, n = 3) at 3 min light exposure and DLHT-29 = 0.272±0.03 (p = 0.002, n = 4) at 120 seconds of light exposure. The clonogenic assay revealed that ATRA+PDT significantly induced a 4-fold and 6-fold reduction of the colony-forming ability of HT-29 following 90- and 120-seconds light exposure, respectively, whereas ATRA monotherapy failed to significantly affect survival (**Fig. 1C**). The impact of ATRA+PDT on proliferation was also evaluated (**Fig 1D**). Strikingly, ATRA+PDT blocked SKBR3 and HT-29 proliferation and attenuated proliferation of the ATRA-resistant HCT116 cells. ATRA monotherapy did not affect the proliferation of HT-29 and HCT116, in contrast to the SKBR3 cells, where the proliferation was slightly reduced.

### Light exposure of ATRA alone is not cytotoxic, and *ex vivo* light exposure of ATRA and fimaporfin does not lead to cytotoxic photoproducts

Retinoids absorb light in the UVA range (315-400 nm). Upon irradiation, retinoids may generate toxicity by forming photoreaction products or forming excited retinoids that directly or indirectly exert toxicity [32]. The lamp used in vitro emits blue light ranging from 400-500 nm, with the highest intensity around 435 nm. ATRA alone was therefore incubated in cells and exposed to blue light to evaluate the toxicity. Our results indicate that HT-29 and MDA-MB-231 cells incubated first with 0.1 µM and 10 µM ATRA, respectively, and then exposed to light, did not get any reduction of cell viability. **(Supplementary Fig. 4**). Furthermore, we also explored the formation of toxic photoproducts of ATRA alone or in the presence of TPCS_2a_/fimaporfin. TPCS_2a_ is an amphiphilic photosensitiser and is taken up to the cells by adsorptive endocytosis. The photosensitiser is, therefore, accumulated over time on the surface of the vesicle membranes. ATRA is a lipophilic molecule and can pass through the plasma membrane passively. An interaction between ATRA and TPCS_2a_ inside the cells is possible if ATRA co-localize in with TPCS_2a_ in the endo/lysosomal membranes during light exposure. To evaluate whether the potential interaction leads to formation of toxic products, ATRA and fimaporfin were exposed to different light doses without cells present and added to cells. We did not observe any significant decrease in cell viability, measured by the MTT assay, 48 hours after incubation start **(Supplementary Fig. 4**).

### ATRA+PDT induce cell cycle arrest and enhance apoptosis

The cell cycle was analysed using flow cytometry at 24 and 72 hours after light exposure (**Fig. 2A**). ATRA monotherapy did not induce any significant change in the cell cycle distribution. In the PDT- or ATRA+PDT-treated HCT116 cells, a slight non-significant increase in G1 was observed 24 hours post light-exposure. At 72 hours, a G2/M accumulation was observed in PDT- or ATRA+PDT-treated HCT116 cells compared to the NT and ATRA treatment (non-significant).

**Fig. 2:**
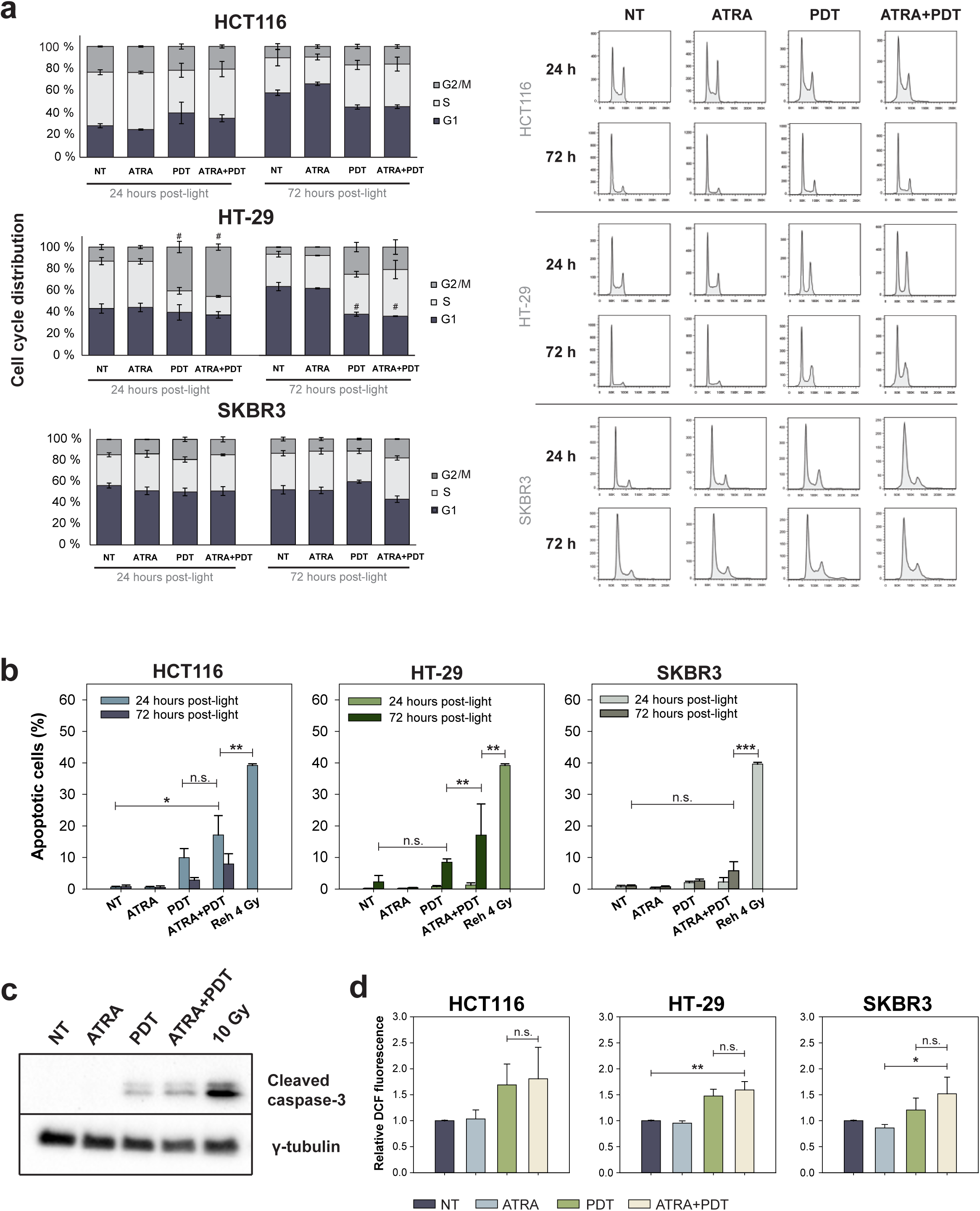
ATRA preconditioning increases PDT-induced cell cycle arrest and apoptosis. The cells were subjected to 120 seconds (SKBR3) and 90 seconds (HT-29 and HCT116) light exposure, which corresponds to ≈ 1.16 J/cm^2^ and 0.87 J/cm^2^, respectively. 4 Gy-treated Reh cells harvested 24 hours post-radiotherapy was included as a positive control. **(A)** Cell cycle analysis at 24- and 72-hours post-light exposure. Left panels, percent distribution in G2/M, S, and G1 phase after the different treatments. Data presented as mean of three independent experiments ± S.E. Right panels, representative cell cycle histograms. **(B)** Apoptosis was evaluated by the TUNEL assay at 24- and 72-hours post-treatment. Data presented as mean of three experiments ± S.E. **(C)** Western blot of cleaved caspase-3 in HT-29 cells harvested 72 hours post-treatment. Cells treated with 10 Gy were included as a positive control, and γ-tubulin was included as a loading control. Representative blot of three experiments. **(D)** Intracellular ROS was measured using the DCFH-DA assay immediately after light exposure. Data are normalized to non-treated samples and presented as mean of three experiments ± S.E. Statistical significance calculated with one-way ANOVA. (*** p ≤ 0.001, ** p ≤ 0.01 and * p ≤ 0.05, n.s.: not significant).

At 24 hours post-PDT or -ATRA+PDT, HT-29 cells significantly accumulated in G2/M, in line with previous PDT-based studies [16, 33]. Specifically, 36.9-43.7% of PDT- or ATRA+PDT-treated cells accumulated in G2/M compared to 12.6-12.9% in NT or ATRA-treated cells (**Fig. 2A**, p ≤ 0.001-0.002). A non-significant increase in G2/M accumulation in both PDT- and ATRA+PDT-treated HT-29 cells was also observed at 72 hours post-treatment. No statistically significant differences were found when comparing PDT- and ATRA+PDT-treated samples for HT-29 and HCT116. For SKBR3 cells, a slight increase in G2/M accumulation was observed 24 hours after PDT, whereas a G2/M accumulation for ATRA+PDT-treated cells was observed after 72-hour (non-significant).

As the anti-proliferation effect of ATRA+PDT increased with time (**Fig. 1D**), we decided to assess the apoptotic response (TUNEL assay) at both 24 and 72 hours after a light-dose inducing ∼ 50% reduction of cell viability (LD_50_). The apoptotic fraction 24 hours post-light was low for both SKBR3 and HT-29. In contrast, a higher (not significant) apoptotic fraction was observed in ATRA+PDT compared to PDT-treated HCT116 cells (**Fig. 2B**). At 72 hours post-ATRA+PDT, a 2-fold higher apoptotic response was detected in HT-29 (p ≤ 0.01) compared to PDT. A higher apoptotic response was also detected in HCT116 and SKBR3 post-ATRA+PDT, although not significant compared to PDT (**Fig. 2B**).

In line with the TUNEL data, no cleaved caspase-3 was detected 24 hours after light exposure of HT-29 (Supplementary data, Uncropped WBs). However, at 72 hours post-light, increased cleaved caspase-3 was observed in PDT- and ATRA+PDT-treated cells (**Fig. 2C**). A tendency (non-significant) of higher expression of cleaved caspase-3 was observed after ATRA+PDT compared to PDT across all three independent experiments. Altogether this indicates that increased cell death after ATRA+PDT is partly due to an enhanced apoptotic response.

### Low-dose ATRA neither elevate intracellular ROS levels nor alkaline phosphatase (ALP) activity and CD133 expression

ATRA induces ROS generation in leukaemia [34]. Compared to PDT, ROS levels post ATRA+PDT showed a tendency to increase (∼ 20%, non-significant) only in the SKBR3 cells. However, low-dose ATRA (0.1 µM) did not influence the ROS generation in all cell lines (**Fig. 2D**).

ALP is a marker of intestinal cell differentiation [35], and was used to evaluate treatment-induced differentiation. Cells were short-term incubated with ATRA concentrations up to 1 µM for 42 hours, harvested, and assayed. The basal ALP activity was high in SKBR3 and low in HCT116 and HT-29 (**Supplementary Fig. 5A**), and was not affected by ATRA treatment. Intriguingly, we observed a tendency (non-significant) of reduced ALP activity in ATRA-treated SKBR3 cells. The surface expression of the cancer stem cell and marker CD133 was evaluated as it has been associated to differentiation status [36]. The CD133 expression was not reduced after 42 hours incubation with 0.1 µM ATRA (**Supplementary Fig. 5B**).

### Differentially expressed genes (DEGs) after ATRA+PDT compared to PDT

As ATRA monotherapy did not have any effect on cell viability, we decided to perform DEGs analysis of RNA-seq data comparing only PDT with PDT+ATRA. By this, we were able to identify candidate genes and key pathways associated with the synergistic effect between ATRA and PDT in HT-29 cells. Based on Prasmickaite *et al*. [37], RNA was harvested 3 hours post-light exposure using a photochemical light dose corresponding to ∼ LD_50_. We identified 1129 genes that were significantly differentially expressed; 676 genes (59.9%) upregulated and 453 (40.1%) downregulated (**Supplementary Table 1**). Among the significant DEGs, 103 genes showed a log2-fold change of at least 1 (**Supplementary Table 2**) including, 56 upregulated (54.4%) and 47 downregulated (45.6%) of which a heat map was generated (**Fig. 3A**). The sequencing data are deposited at the European Nucleotide Archive https://www.ebi.ac.uk/ena/browser/view/PRJEB49953).

**Fig. 3:**
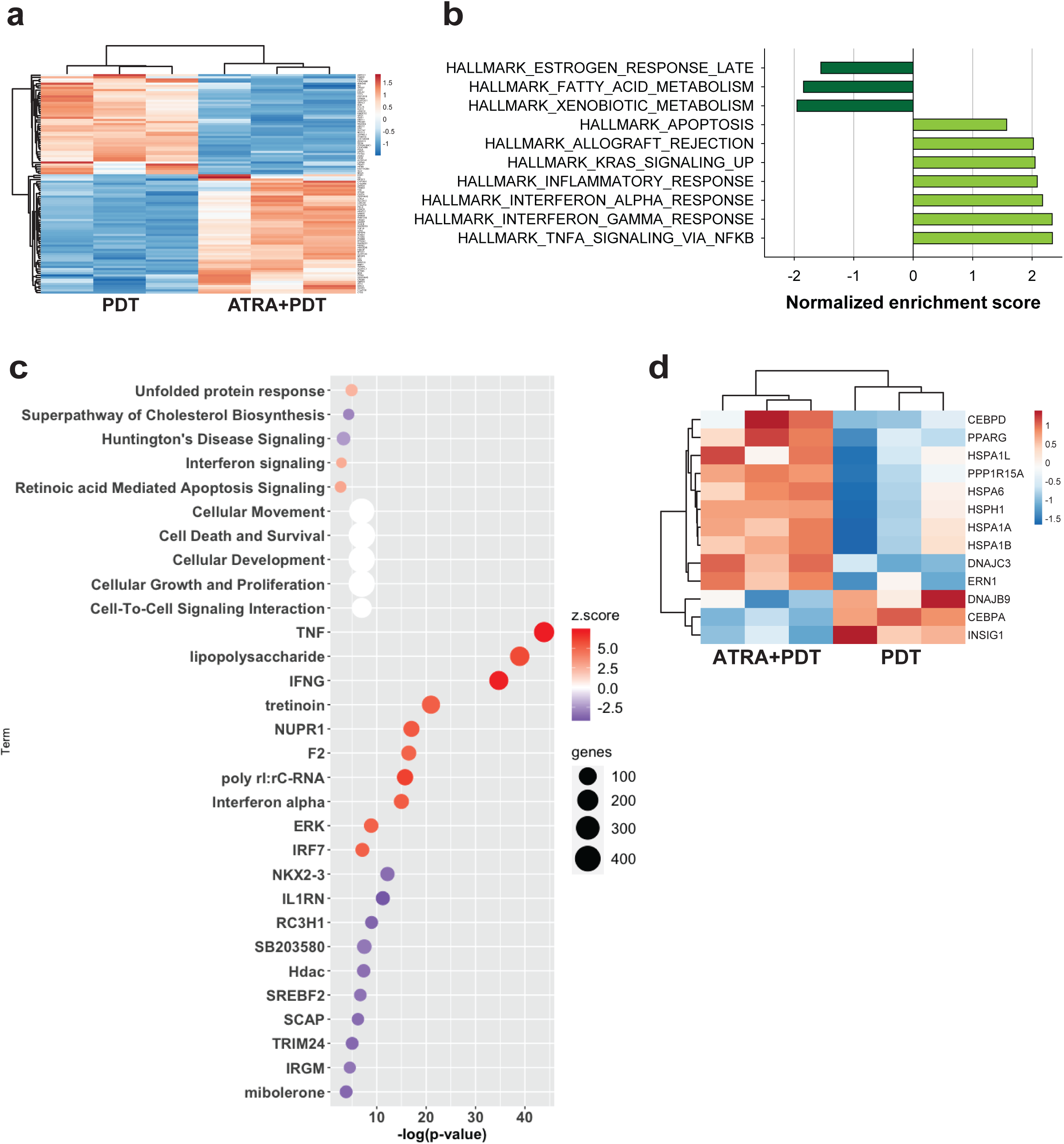
Differentially expressed genes (DEGs) and gene set enrichment analyses (GSEA) in ATRA+PDT-treated HT-29 cells. **(A)** Heat map of DEGs of fold change ≥ |2| at 3 hours post-light exposure in PDT and ATRA+PDT-treated samples. **(B)** All significantly enriched biological states or processes in ATRA+PDT compared to PDT determined by GSEA analysis using the Hallmark gene set. A p-value of ≤ 0.05 and an FDR of ≤ 25% were considered significant. **(C)** Visualization of IPA analysis of top five canonical pathways based on p-value, top five molecular and cellular functions, and top ten activated and inhibited based on z-value. A p-value of ≤ 0.01 and a predicted activation, z-score of ≥ |2| were considered significant. (**D)** Heat map of DEGs mapped to the unfolded protein response.

### Gene set enrichment analysis (GSEA)

To obtain specific, well-defined biological states or processes underlying ATRA+PDT, all significant DEGs were first submitted to the GSEA software using the MSigDB hallmark gene set collection. The analysis revealed ten significantly enriched gene sets (**Fig. 3B**). Apoptosis and upregulation of KRAS signaling were positively correlated to ATRA+PDT, whereas late oestrogen response, fatty acid metabolism, and xenobiotic metabolism were negatively correlated. Interestingly, a significant enrichment in immune-related gene sets were identified including allograft rejection, inflammation, IFN-α/ IFN-γ response and TNF-α signaling.

All significant DEGs were further analysed using the IPA software for gene set enrichment analysis to gain mechanistic insight into biological processes, pathways, molecular networks, and upstream regulators. 68 canonical pathways were significantly enriched, of which 12 pathways had a significant activation z-score (**Supplementary Table 3**). Five canonical pathways, including the UPR IFN signaling and retinoic acid-mediated apoptosis signaling, were predicted significantly activated, and seven were predicted to be significantly inhibited (**Fig. 3C).** Furthermore, top molecular and cellular functions identified by IPA were associated with cellular movement, cell death and survival, cellular development, cellular growth and proliferation, and cell-to-cell signaling interaction. As PDT is a strong inducer of cellular stress, we decided to further explore the UPR response with a separate heat map (**Fig. 3D**).

The upstream regulator analysis identified 1407 significant regulators (p ≤ 0.01), of which 229 were predicted as activated and 79 as inhibited (**Supplementary Table 4, Fig. 3C**). Of relevance, strong activations of the pro-inflammatory and immune-related regulators tumour necrosis factor (TNF), lipopolysaccharide (LPS), interferon-gamma (IFNG), poly rI:rC-RNA (a synthetic dsRNA analogue known as poly (I:C)), and interferon-alpha (IFNA) were detected.

### Validation of gene expression data

Reverse transcription quantitative real-time (RT-qPCR) was used to validate the RNA-seq data of six DEGs: *REG4, IGFBP6, ATF3, EGR1, LCN2,* and *ALDH1A1*. The RT-qPCR data revealed that the selected genes’ fold-change was comparable to the RNA-seq results (**Fig. 4A**)

**Fig. 4:**
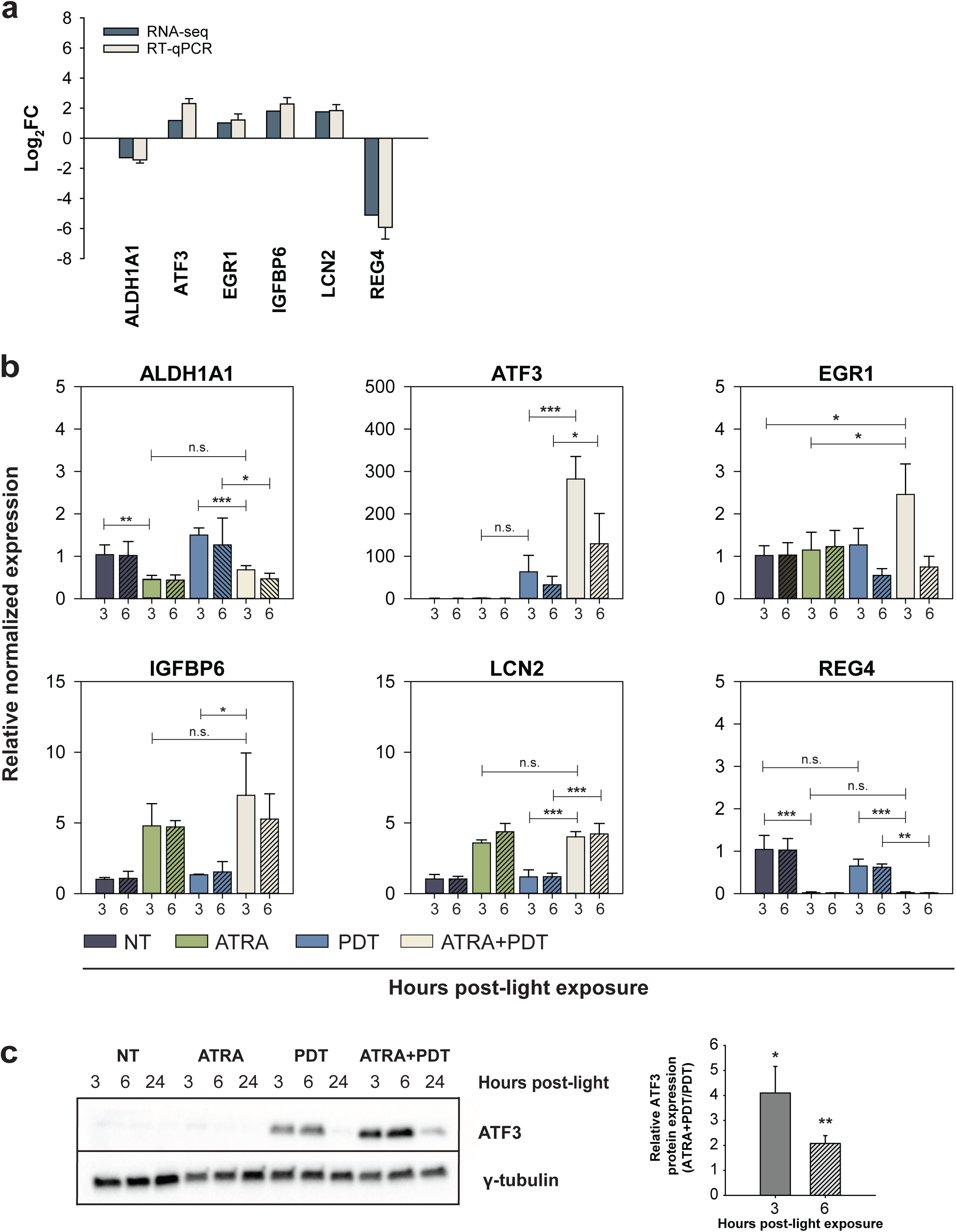
Post-RNA-seq validation of six selected DEGs by RT-qPCR and ATF3 protein expression by Western blotting. **(A)** Validation of mRNA expression of six selected DEGs (ALDH1A1, ATF3, EGR1, IGFBP6, LCN2 and REG4) by RT-qPCR of samples harvested 3 hours post-treatment. RT-qPCR data are presented as log-2-fold change (FC) (ratio of ATRA+PDT/PDT) and compared to the RNA-seq data. **(B)** Relative normalized expression of DEGs in samples obtained 3 or 6 hours post-ATRA+PDT treatment and compared to non-treated (NT), ATRA- and PDT-treated controls. The data are normalized to two reference genes (RPLP0 and GADPH) and presented as mean of three experiments ± S.E. Statistical significance calculated with one-way ANOVA. **(C)** Relative ATF3 signal in ATRA+PDT to PDT samples. The data were normalized to loading control. Data are presented as mean of four experiments ± S.E. One-sample t-test. Representative western blot of ATF3 at 3-, 6-, and 24-hours post-treatment. γ-tubulin was included as a loading control. (*** p ≤ 0.001, ** p ≤ 0.01 and * p ≤ 0.05, n.s.: not significant).

### ATRA+PDT rapidly induces ATF3 and EGR1 expression

ATRA+PDT-treated samples were evaluated using RT-qPCR at 3 and 6 hours post-treatment and compared with non-treated, PDT- and ATRA-treated cells (**Fig. 4B**). IGFBP6 and ALDH1A1 have previously been described in the context of ATRA [38, 39], whereas REG4, ATF3, and EGR1 have, to the best of our knowledge, not been associated with ATRA treatment. ATF3 and EGR1 were selected based on their association identified by the STRING network analysis (**Supplementary Fig. 6**). Both the ATF3 and EGR1 gene expressions were upregulated synergistically after ATRA+PDT. At 3 hours post-treatment, ATRA+PDT induced a 2.5-fold increase of the EGR1 expression, while no significant difference were detected in non-treated, PDT-, and ATRA-treated samples. Strikingly, a 282-fold increase of ATF3 expression was observed 3 hours post-ATRA+PDT, whereas PDT induced a 63-fold increase 3 hours post-light. These observations are in line with the literature, as ATF3 expression is rapidly upregulated by stress [40]. The mRNA expressions of ATF3 and, in particular, EGR1 were strongly downregulated 6 hours post-PDT or - ATRA+PDT **(Fig. 4B)**. As expected, no difference in expression of LCN2, ALDH1A1, or IGFBP6 was observed between ATRA and ATRA+PDT at 3 and 6 hours post-light. Of notice, the REG4 expression was entirely abolished in ATRA- or ATRA+PDT-treated cells, an observation that, to the best of our knowledge, has not been reported before **(Fig. 4B)**. The increased ATF3 and EGR1 expressions were identified as unique for the ATRA+PDT combination.

The ATF3 protein expression was validated by Western blotting of PDT- or ATRA+PDT- treated samples 3 and 6 hours post light exposure (**Fig. 4C**). At the 24 hour, the ATF3 expression was strongly downregulated, with slightly higher ATF3 expression in the ATRA+PDT samples compared to PDT. The basal level of ATF3 was low in the non-treated samples and after ATRA monotherapy (**Fig. 4C**).

### PDT+ATRA induces initial tumour growth delay and CRs in HT-29 xenografts

The anti-tumour activities of ATRA+PDT were first assessed in HT-29 xenografts in athymic nude mice (n=5 per experimental group) using a systemic protocol and a 652 nm laser dose of 15 J/cm^2^. The ATRA+PDT treatment had an initial strong anti-tumour effect; however, due to unexpected adverse effects including xeroderma and weight loss resulting in mortality the final group size (n=2) was too small to draw any conclusions (**Supplementary Fig. 7**). This effect was not expected as several reports delivered ATRA systematically and continuously [41, 42]. Also ATRA treatment alone induced a significant weight loss. To improve the safety profile, we decided to explore a focal ablation approach including intratumoral (i.t.) co-delivery of photosensitiser and ATRA combined with a reduced laser dose (10 J/cm^2^). When a primary lesion or its metastases are accessible, i.t. injection of anti-cancer drugs, in particular, immunostimulatory agents have been recommended [43]. In the second modified protocol, both ATRA and fimaporfin (TPCS_2a_) were injected i.t. as a single dose one day (18-24 hours) before light exposure. As anticipated, compared to systemic delivery, i.t. caused only mild adverse effects including, oedema and erythema in tumour areas after PDT or ATRA+PDT **(Fig. 5A)**. All ATRA+PDT-treated animals (5/5) had an initial tumour growth delay compared to the other treatment groups (**Fig. 5B, C and Supplementary Fig. 8A**). By comparing the mean tumour size in each treatment group at 5, 10, and 20 days post-treatment, ATRA+PDT significantly inhibited tumour growth up to 20 days post-treatment compared to non-treated (NT). The mean tumour volume in the ATRA+PDT group was up to 3-fold smaller compared to the other treatment groups (not significant) (**Fig. 5B**). Although not reflected in the tumour measurements, visual examination of the PDT- or ATRA+PDT-treated tumours indicated they were flatter or had a necrotic centre (**Fig. 5A**). The tumour volume measurements are based on the assumption that tumours are spherical. As the PDT- and ATRA+PDT-treated tumours are not spherical (but flat with a necrotic core), these groups’ tumour measurements were most likely overestimated. During laser illumination, fluorescence was observed in tumours injected with fimaporfin (TPCS_2a_) alone or in combination with ATRA (ATRA+PDT) (**Fig. 5D**). In the PDT group, the fluorescence was limited to the tumour, indicating that most of the photosensitiser is confined in the tumour. Intriguingly, for mice treated with ATRA+PDT, we additionally detected fluorescence in the tumour margins (**Fig. 5D**).

**Fig. 5:**
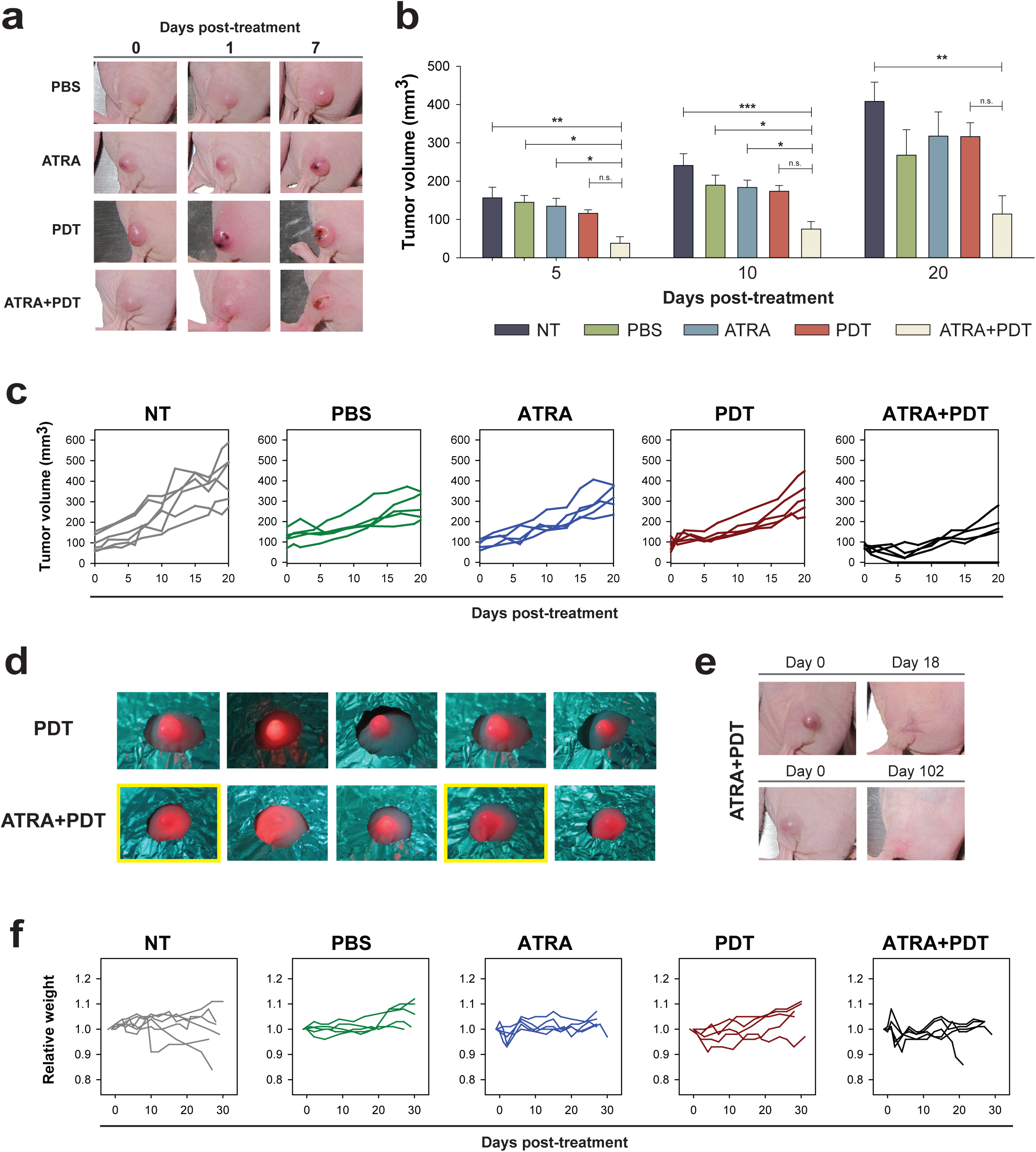
Treatment response in HT-29 xenografts following i.t. delivery of ATRA, fimaporfin, and photochemical treatment. **(A)** Representative images of one animal in each treatment group at day 0 (before light exposure), 1 and 7 post-treatment. **(B)** Tumour volume (mm^3^) of each treatment group at day 5, 10, and 20 post-treatment. Data presented as mean ± S.E. Statistical significance calculated with one-way ANOVA. (*** p ≤ 0.001, ** p ≤ 0.01 and * p ≤ 0.05, n.s.: not significant)**. (C)** Tumour growth of individual animals in each treatment group up to 20 days post-treatment. Each line represents one animal. **(D)** Images of fimaporfin fluorescence in the tumours during light exposure in the PDT- and ATRA+PDT-group. The yellow frames indicate animals that achieved complete responses. **(E)** Images of the two mice that were cured in the ATRA+PDT group before light exposure (day 0) and post-treatment at a later time-point (day 18 and 102). **(F)** Relative weight of each individual animal, relative to day -1, in each treatment group. Each line represents one animal. For all treatment groups, n = 5, except for NT where n = 6.

Two out of five (40%) ATRA+PDT-treated animals achieved complete responses (CR) after only one treatment (**Fig. 5E**). However, one of the cured animals was euthanized on day 21 due to weight loss, not associated with the treatment as one animal in the non-treated control group was also euthanized due to weight loss (**Fig. 5F**). None of the mice in any of the other treatment groups achieved CR. The animals were followed up to at least 90 days post-treatment. Surprisingly, in the ATRA group, 3/5 animals reached day 90 post-treatment. The ATRA-treated tumours grew similar to the controls up to 20 days post-treatment. However, tumour growth stagnated over time (**Supplementary Fig. 8B**), and surprisingly, the best overall survival was obtained with ATRA monotherapy (p = 0.012) compared to NT (**Supplementary Fig. 8C**). The mean time to reach the endpoint (day 90, weight loss ≥ 20% and/or tumour size ≥ 1000 mm^3^) was longest in the ATRA group (76.8 ± 8.1 days).

As the impact of the needle trauma and injection of fluids on the tumour microenvironment was unknown, a sham procedure including an i.t. injection of PBS was included in addition to the NT control group. No animals in the NT group reached day 90. However, 2/5 animals in the PBS treatment group reached day 90, of which one animal was at endpoint (tumour volume >1000 mm^3^). The fact that PBS-treated animals also survived up to day 90 indicate that i.t. injection of fluids itself affects tumour growth. One of five animals in the PDT group (tumour volume >1000 mm^3^) and in the ATRA+PDT group (CR) also reached day 90 (**Supplementary Fig. 8D**).

## Discussion

ATRA has not proven successful in the treatment of solid tumours due to intrinsic or acquired drug resistance. It is therefore suggested that ATRA needs to be combined with other therapies to improve therapeutic outcomes [4]. In this study, we present for the first time therapeutic and mechanistic evidence demonstrating that ATRA preconditioning combined with PDT (ATRA+PDT) is a promising anti-carcinoma strategy. Different experiments were conducted to identify the possible mechanism behind the enhanced efficacy of ATRA+PDT, including assessment of differentiation (ALP and CD133 expression), ROS-generation, mTOR inhibition, cell cycle distribution, and cytotoxic products of ATRA and fimaporfin, of which none were identified as the mechanism. Thus, in an effort to provide a mechanistic rationale for how ATRA preconditioning is able to make carcinoma cells vulnerable and enhance the cytotoxic efficacy of PDT, we decided to determine the DEGs levels between the two experimental conditions ATRA+PDT and PDT. DEG analysis of the RNA-seq data revealed a complex transcriptome profile post-ATRA+PDT. The DEG analysis indicated that EGR1 and ATF3 were significantly upregulated in ATRA+PDT-treated cells. In line with our observation, the stress response transcription regulator ATF3 has previously shown to be induced after PDT [37, 44, 45], and implicated in the UPR [46]. ATF3 was therefore selected for further analysis and validation on the mRNA and protein level. Strikingly, the RT-qPCR results indicated a 282- and 63-fold increase in ATF3 mRNA in the ATRA+PDT- and PDT- treated cells, respectively. Increased ATF3 protein expression after ATRA+PDT was also confirmed. Strikingly, REG4 expression was reduced by a factor of ∼35X by ATRA+PDT compared to PDT. REG4 is overexpressed in colorectal carcinomas and is predictive of poor prognosis and drug resistance [47]. REG4 has also shown to protect colon crypt stem cell and colorectal cancer cells from radiation-induced apoptosis [48] and, recently, it was demonstrated that REG4 interacts with CD44 and improve the stemness of colorectal cancer cells [49]. Altogether, this suggests that the ATRA-PDT-induced blockage of REG4 expression may explain in part the enhanced apoptotic response in the HT-29 cells.

The bioinformatics analysis revealed pathways and upstream regulators supporting our observation related to enhanced cytotoxicity of ATRA+PDT, including apoptosis and the UPR as one of the top pathways. Misfolded protein or the formation of protein aggregates due to ROS-induced proteotoxic stress is a primary response to PDT independent of the cell type and PDT strategy [50]. Upon endoplasmic reticulum (ER) stress, UPR is activated to reduce accumulation and aggregation of misfolded or unfolded proteins playing an important role in homeostasis and cell survival. However, if the ER stress is excessive and sustained, apoptosis is induced [51]. Indeed, our observation of ATRA+PDT-enhanced apoptotic cell death by the TUNEL assay was supported by both GSEA and IPA. Western blot analysis revealed increased cleaved caspase-3 post-ATRA+PDT, but not significantly elevated compared to PDT, which was confirmed by the DEG-analysis. However, compared to PDT monotherapy, DEG-analysis revealed significant upregulation of caspase-4, -7, and -9 post-ATRA+PDT. Furthermore, TNF-α, a potent pro-apoptotic trigger that may also induce necrosis [52], was identified as an activated upstream regulator after ATRA+PDT.

The downstream effects of the p38 kinase inhibitor SB203580 were predicted to be significantly inhibited suggesting that p38 kinase is involved in stress-induced death signalling after ATRA+PDT. This is consistent with previous observations; Weyergang *et al*. demonstrated that SB203580 in combination with TPPS_2a_-based PDT resulted in increased cell viability [25], while similar effects were shown by Olsen *et al.* where SB203580 was combined with fimaporfin-PDT [16].

HDAC was identified as one of the upstream regulators with a negative z-score indicating HDAC inhibition (HDACi) despite that the class II deacetylase HDAC9 was 2.2-fold upregulated by ATRA+PDT. HDAC9 negatively r**e**gulates the function of TRIM29 by non-histone deacetylation of the protein [53], and thereby alters its ability to associate with p53 and consequently inhibits its cell proliferation-promoting activity. Interestingly, our RNA-seq data reveal that TRIM29 is 30% downregulated post-ATRA+PDT (p = 5.00E-05), which may strengthen the concept of the HDACi effect of ATRA+PDT.

The ATRA+PDT combination strategy was further evaluated *in vivo* by using the HT-29 xenograft model in athymic nude mice. High morbidity (significant weight loss and xeroderma) was observed when ATRA was given systemically. In an effort to enhance the safety profile and anti-tumour efficacy, ATRA and TPCS_2a_ were co-administered i.t. This technique was recently suggested by Marabelle *et al.* as a strategy to deliver high drug (immunostimulatory agents) concentration in the tumour [43].

All ATRA+PDT-treated animals had smaller tumours than control groups during the first 20 days post laser exposure and 2 of 5 ATRA+PDT-treated mice achieved CR after only one treatment. In the ATRA+PDT group, fimaporfin fluorescence was observed in the tumours as well as in the tumour margins during light exposure, as opposed to the PDT treatment group, where the fluorescence was confined to the tumours only. Although the exact mechanism of this ATRA effect is not clear, the enhanced TPCS_2a_ distribution in the tumour margin may contribute to the improved overall PDT response and explain the initial tumour growth inhibition in the ATRA+PDT group. In addition to direct cell killing, PDT with PCI-based photosensitisers also targets the tumour vasculature [14, 54, 55]. Thus, light activation of fimaporfin in the tumour periphery of the ATRA+PDT-treated animals may induce vascular shutdown, which limits the oxygen and nutrition supply to the tumour, but this remains to be tested experimentally. Despite the CRs obtained after ATRA+PDT, ATRA monotherapy surprisingly gave the best overall survival, of which 60% of the mice were still alive at day 90.

A limitation of the present study is that we do not have any validation on the protein level both on EGR1, REG4 and the UPR in general. In addition, the xenograft model is immune deficient and hence limits the potential anti-immune effects of ATRA+PDT. Thus, more research needs to be conducted regarding the role of inflammatory and immune regulators after ATRA+PDT in syngeneic *in vivo* models.

In conclusion, our study shows that low-dose short term ATRA preconditioning is a promising strategy to improve the efficacy of fimaporfin-PDT in carcinomas independent on a differentiation effect. DEG analysis predicted the unfolded protein response as the most significantly upregulated pathway and enhanced activation of apoptosis as a major cell death mechanism. To fully exploit the potential of the ATRA+PDT combinatorial approach, preclinical studies of ATRA in combination with laser-activation of the clinical relevant photosensitiser fimaporfin in immunocompetent murine models are warranted.

## Statements and Declarations

## Acknowledgements

We are grateful to Drs. Trond Stokke and Idun Dale Rein at the Flow Cytometry Core Facility (OUS) for advice on cell cycle analysis. We also thank Ane Sager Longva for excellent technical assistance and advice on RNA extraction, RT-qPCR and Western blotting, Miriam Rosvold From for assistance on the MDA-MB-231 cell experiments and Ane Sofie Fremstedal for processing the tumour measurement data. We thank Dr. Anders Høgset/PCI Biotech AS for providing us with the photosensitiser fimaporfin.

## Authors’ contributions

**Judith Jing Wen Wong:** Writing - Original Draft, Investigation, Formal analysis, Validation, Methodology, Data Curation, Visualization. **Susanne Lorenz:** Investigation, Formal analysis, Data Curation, Writing - Review & Editing. **Pål Kristian Selbo:** Conceptualization, Supervision, Project administration, Funding acquisition, Resources, Methodology, Writing - Review & Editing

## Data Availability

The RNA-seq datasets generated and analysed during the current study are available in the EMBL Nucleotide Sequence Database: European Nucleotide Archive (ENA) repository, https://www.ebi.ac.uk/ena/browser/view/PRJEB49953. The other datasets generated, used and/or analysed during the current study are available from the corresponding author on reasonable request.

## Ethics approval

All animal procedures were performed according to protocols approved by the national animal research authority Mattilsynet (FOTS ID22020) and were conducted according to the regulations of the Federation of European Laboratory Animal Science Association (FELASA) Handling of and experiments with animals were therefore performed in compliance with EUs Directive 2010/63/EU on the protection of animals used for scientific purposes.

## Consent to participate

Not applicable

## Consent to publish

Not Applicable

## Abbreviations

ALDH1A1: aldehyde dehydrogenase 1 family member A1
ALP: alkaline phosphatase
ATF3: activating transcription factor 3
ATRA: all-trans retinoic acid (tretinoin)
ATRA+PDT: ATRA preconditioning with fimaporfin-based photodynamic therapy
DEG: differentially expressed gene
EGR1: early growth response 1
GSEA: gene set enrichment analysis
HDAC9: histone deacetylase 9
IFN: interferon
IGFBP6: insulin-like growth factor-binding protein 6
IPA: Ingenuity pathway analysis
LCN2: lipocalin 2
LD50: light-dose inducing 50% reduction of cell viability
LPS: lipopolysaccharide
PCI: photochemical internalisation
PDT: photodynamic therapy
RAR: retinoic acid receptor
REG4: regenerating islet-derived protein 4
RNA-seq: RNA sequencing
ROS: reactive oxygen species
UPR: unfolded protein response
TNF: tumour necrosis factor
TPCS_2a_: disulfonated tetraphenyl chlorin (fimaporfin)
TUNEL: Terminal deoxynucleotidyl transferase (TdT) dUTP nick-end labelling

