## Supplementary figure legends for "All-*trans* retinoic acid enhances the anti-tumour effects of fimaporfin-based photodynamic therapy"

**Supplementary Fig. 1. ATRA alone is sub-toxic at physiological/therapeutic relevant concentrations in HT-29, HCT116, and SKBR3 cells.** Relative cell viability in cells incubated with increasing ATRA concentrations (0–100 μM) for 48 hours. Data presented as mean of three experiments ± S.E.

**Supplementary Fig. 2. ATRA pre-conditioning enhances fimaporfin-PDT in MDA-MB-231 and MC-38 cells.** Left panel, relative cell viability of MDA-MB-231 cells after 18 hrs co-incubation with 10 μM ATRA and 0.4 μg/mL $TPCS_{2a}$ prior to light exposure (PDT). Viability (MTT) was assessed 48 hours after PDT. Representative experiment of three independent experiments, data presented as mean of triplicates ± S.D Right panel, relative cell viability of MC-38 cells after 42 hrs pre-incubation of 1 μM ATRA including 18 hrs co-incubation with 0.6 μg/mL TPCS2a prior to light exposure (PDT). Viability (MTT) was assessed 24 hours after PDT. Representative experiment of three independent experiments, data presented as mean of triplicates ± S.D. * p<0.05, *** p<0.001

**Supplementary Fig. 3. ATRA do not enhance fimaporfin-PDT when given after light.** ATRA incubation for 48 hours after PDT does not induce enhanced cytotoxicity. HT-29 (left panel) and MC-38 cells (right panel) were incubated with 1 μM and 10 μM ATRA, respectively. Representative experiment of three independent experiments, data presented as mean of triplicates ± S.D.

**Supplementary Fig. 4. ATRA+light is non-toxic and no phototoxic products are generated after light exposure of ATRA and fimaporfin ($TPCS_{2a}$) in a cell-free system.**

**(A)** ATRA alone in HT-29 (0.1 μM) and MDA-MB-231 (10 μM) cells exposed to increasing light exposure. Viability evaluated 48 hours post-light exposure. (**B)** 0.1 μM ATRA and 0.4 μg/mL fimaporfin ($TPCS_{2a}$) exposed to light ex vivo and subsequently incubated for 48 hours in HT-29 cells. Representative experiment of at least two experiments, data presented as mean of triplicates ± S.D.

**Supplementary Fig. 5. Sub-lethal ATRA concentrations does not significantly affect the differential marker alkaline phosphatase and the stem cell marker CD133.**

**(A)** Alkaline phosphatase expression per mg protein and (**B),** relative CD133 expression in HCT116, SKBR3 and HT-29 incubated with 0.1–1 μM ATRA for 42 hours. Data presented as mean of three experiments ± S.E.

**Supplementary Fig. 6. Drug-protein and protein-protein interaction network of retinoic acid and DEGs (ATRA+PDT versus ATRA) of at least 1 $log_2$-fold change.**

**(A)** The drug-gene and protein-protein interaction network from the STITCH database of retinoic acid. The block represents the drug and circle represents gene. Associations are indicated with a line. Stronger associations are represented by thicker lines. (**B)** Protein-protein network from the STRING database. Each circle represents a protein. Association between the proteins is indicated with a line.

**Supplementary Fig. 7. Treatment response following systemic ATRA and fimaporfin-PDT (ATRA+PDT) in HT-29 -tumor bearing mice.**

**(A)** Relative body weight in the ATRA-treatment group. The animals received ATRA i.p. once a day for five days. The weight of each animal was normalized to the weight at the day -4. **(B)** ATRA+PDT group received five doses of ATRA prior to light treatment. 10 mg/kg ATRA was delivered intraperitoneally once a day for five days. TPCS2a (5 mg/kg) was delivered as a single dose intravenously three days prior to laser exposure with a light dose of 15 $J/cm^2$ using an irradiance of 90 $mW/cm^2$. The Kaplan-Meier survival curve shows the treatment response and the table shows the estimated time to reach endpoint in the different treatment groups. Note that ATRA+PDT group is only 2 animals as 4 animals had severe toxic effects after systemic delivery of ATRA. Consequently, it was decided to co-deliver ATRA and fimaporfin/TPCS2a by intratumoral injections in the further experiments (Figure 5 + Supplementary Fig.8). **(C)** Mean tumor volume ($mm^3$), and **(D)** waterfall plots of each treatment group at 5, 10, and 20 days post-treatment. Each bar indicate an individual animal and percentage change of tumor size compared to day 0. NT, ATRA and PDT: 6 mice per group, except for ATRA+PDT where $n = 2$. Data presented as mean ± S.E.

**Supplementary Fig. 8. Survival and tumor response following intratumoral injection of ATRA and fimaporfin, and photochemical treatment.**

**(A)** Waterfall plots of each treatment group at 5, 10 and 20 days post-treatment. Each bar indicate an individual animal. The tumor size at indicated time-point relative to tumor size at treatment start (day 0). **(B)** Tumor volume ($mm^3$) of individual animal in each treatment group up to 90 days post-treatment. Each line represents one animal. **(C)** Survival curve of treatment response and table of estimated mean time (days) to reach endpoint for each treatment groups. For all treatment groups $n = 5$, except for NT where $n = 6$. Survival evaluated using log-rank (Mantel-Cox) * $p \leq 0.05$. **(D)** Tumor size ($mm^3$) of animals in each group that reached day 90.
