## Supplementary figures and images for "All-*trans* retinoic acid enhances the anti-tumour effects of fimaporfin-based photodynamic therapy"

### Supplementary Fig.1

Supplementary Figure 1

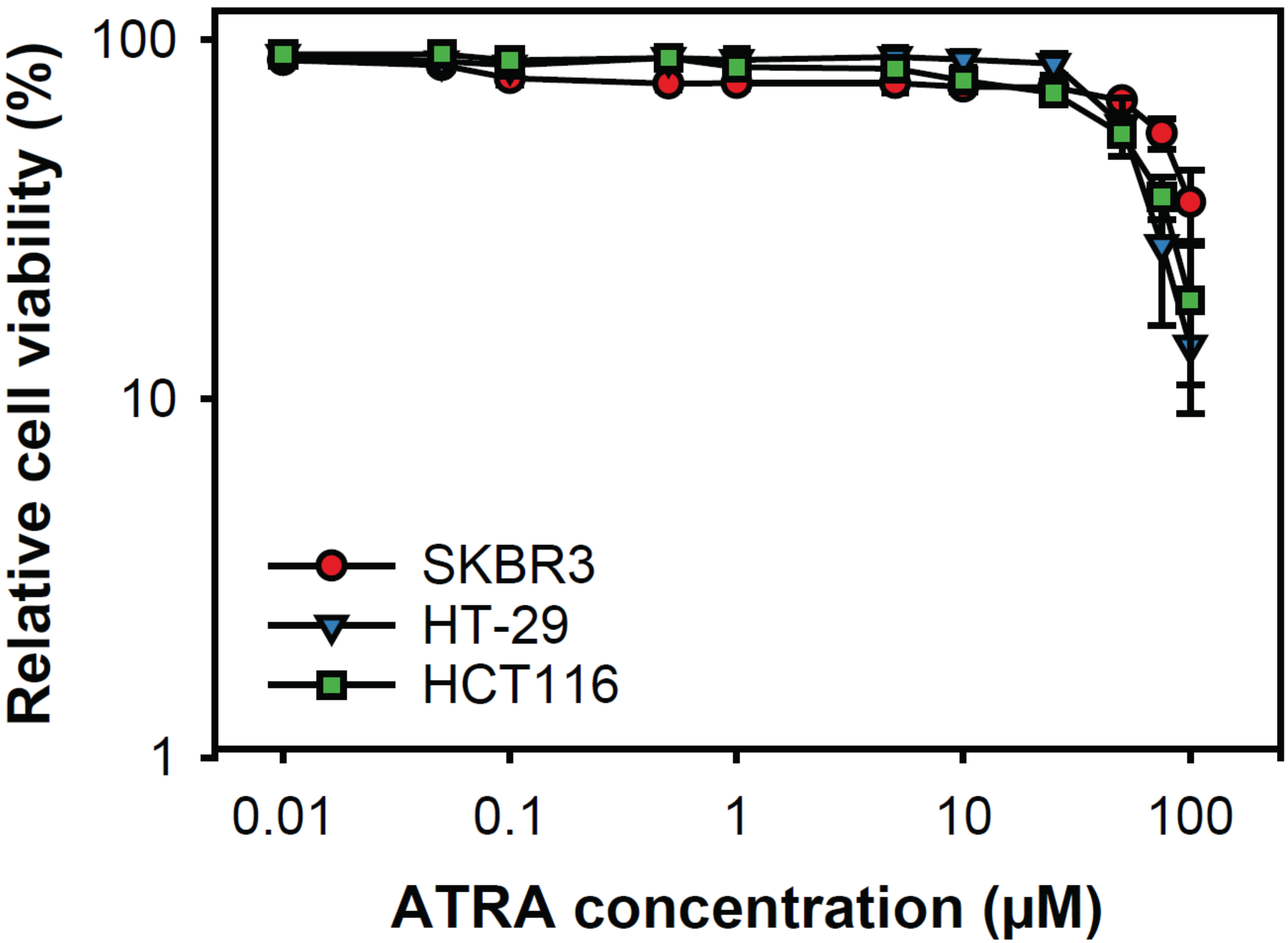

### Supplementary Fig.2

Supplementary Figure 2

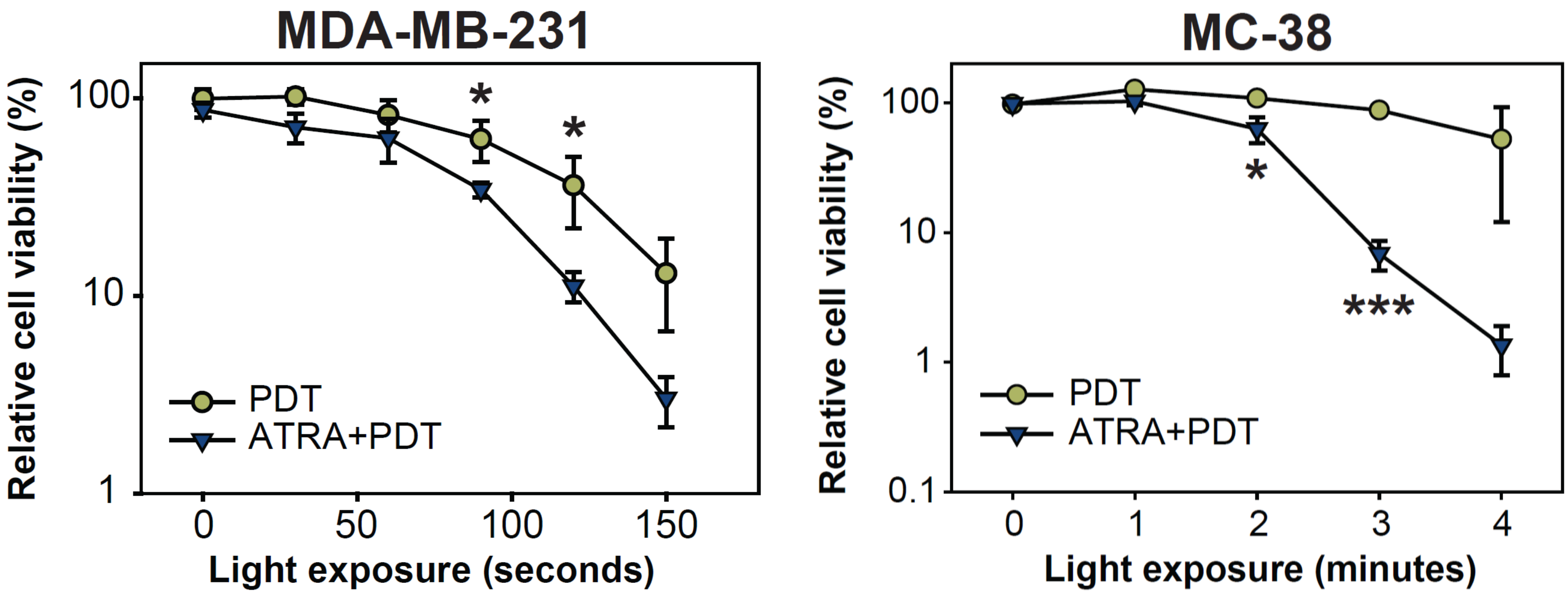

### Supplementary Fig.3

Supplementary Figure 3

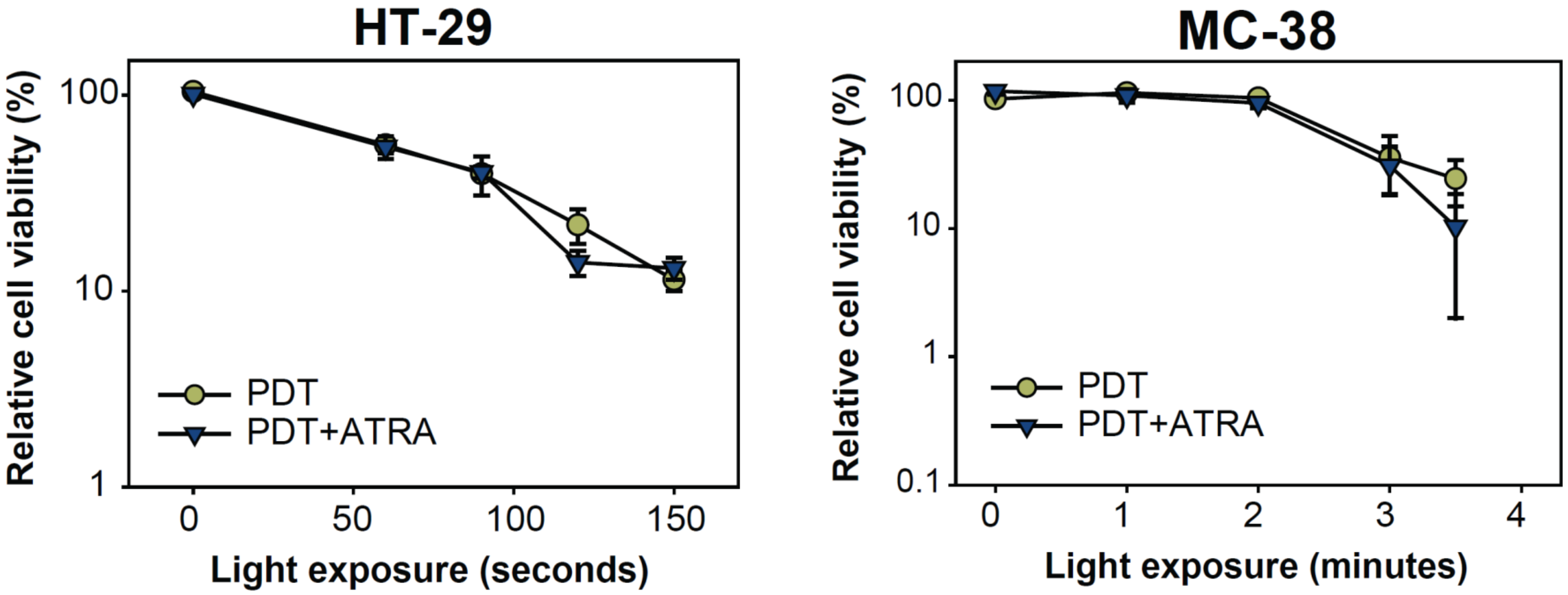

### Supplementary Fig.4

Supplementary Figure 4

A

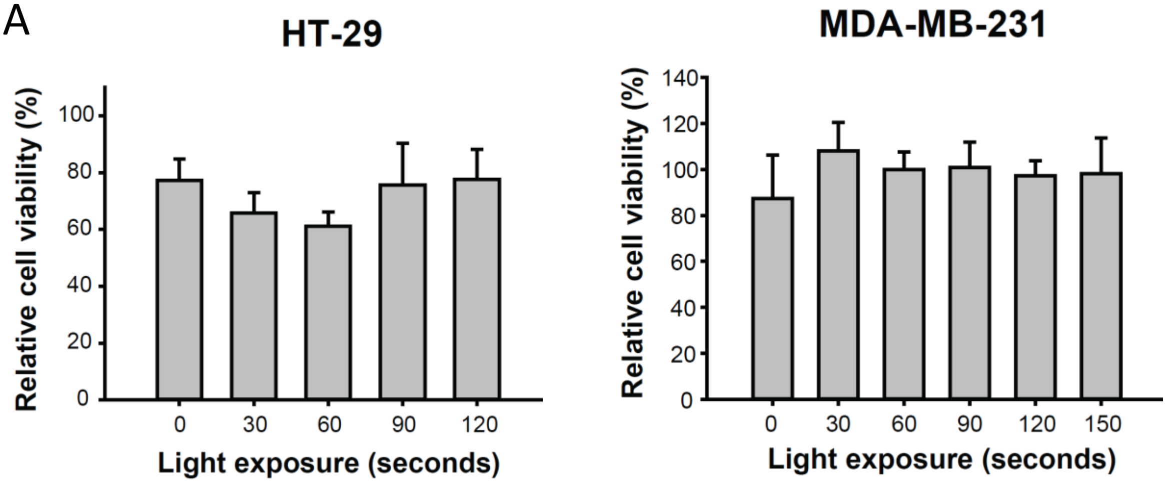


B

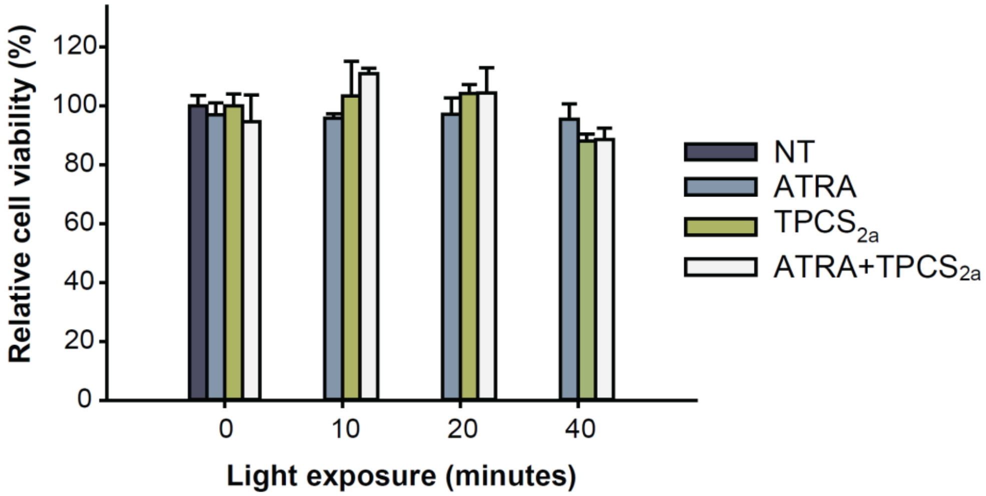

### Supplementary Fig.5

Supplementary Figure 5

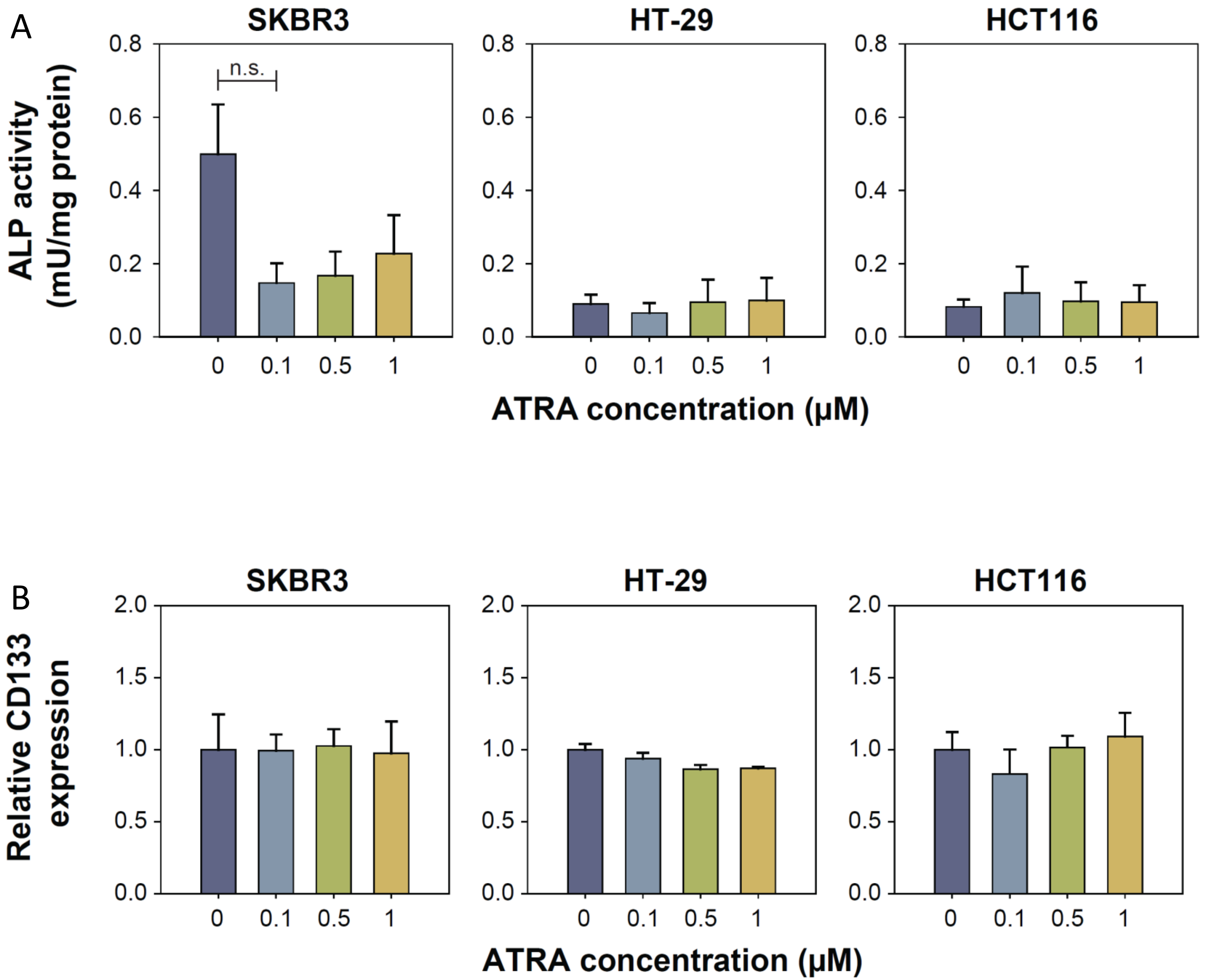

### Supplementary Fig.6

Supplementary Figure 6

A

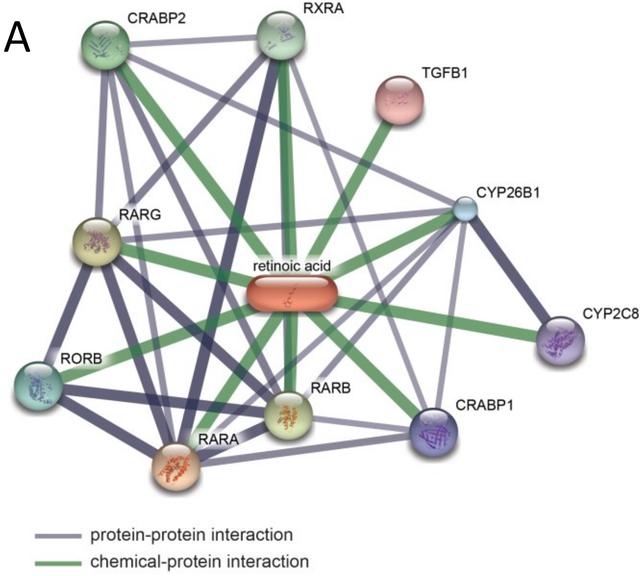


B

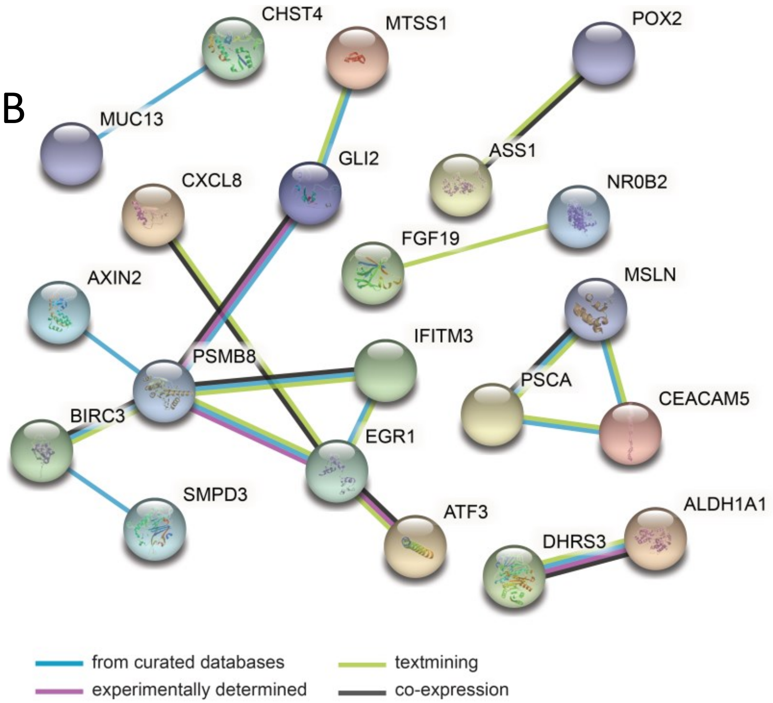
