## Supplementary Fig.7 for "All-*trans* retinoic acid enhances the anti-tumour effects of fimaporfin-based photodynamic therapy"

Supplementary Figure 7

**a**

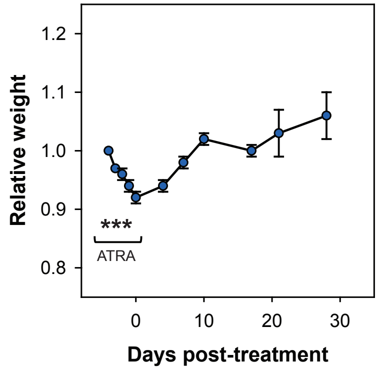


**b**

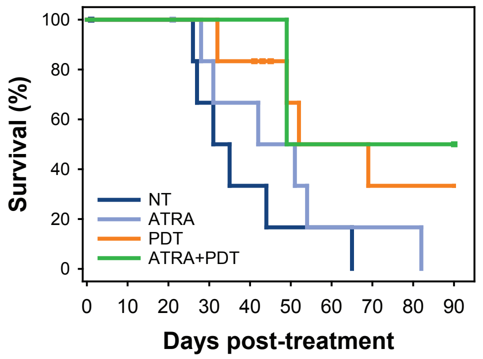


| Treatment group | Mean time to reach endpoint (days) | S.E. | No. of animals |
|---|---|---|---|
| NT | 38.0 | 6.0 | 6 |
| ATRA | 48.0 | 8.0 | 6 |
| PDT | 63.7 | 8.8 | 6 |
| ATRA+PDT | 69.5 | 14.5 | 2 |

**c**

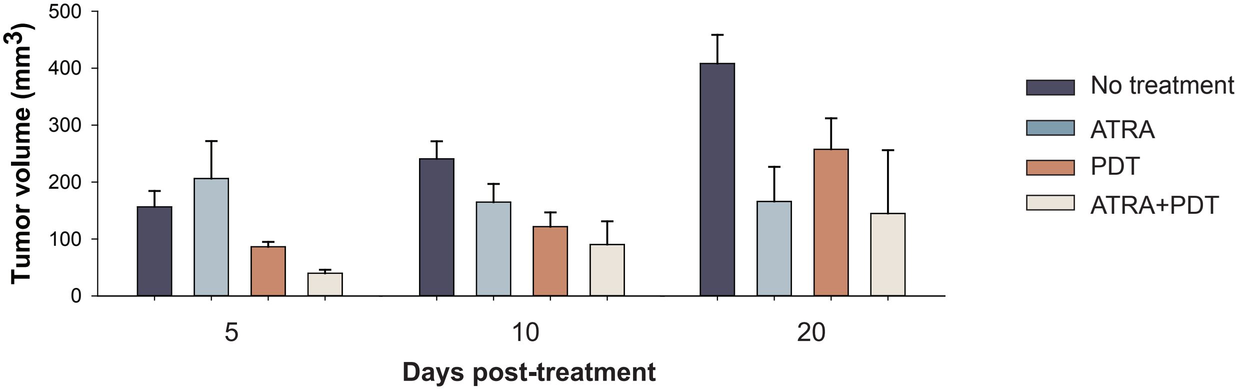


**d**

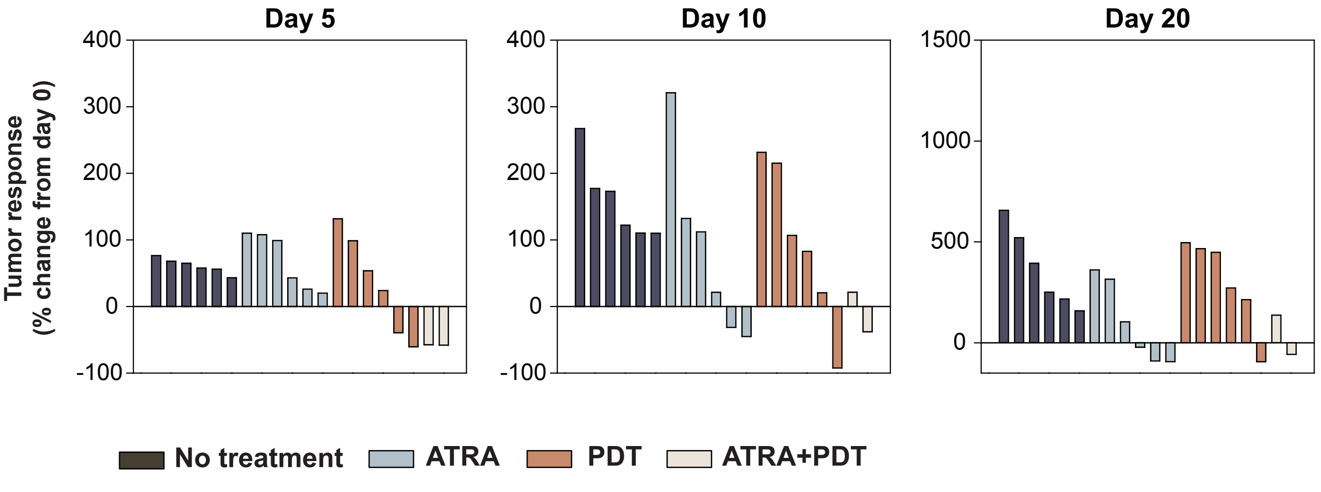
