## Supplementary Fig.8 for "All-*trans* retinoic acid enhances the anti-tumour effects of fimaporfin-based photodynamic therapy"

Supplementary Figure 8

a

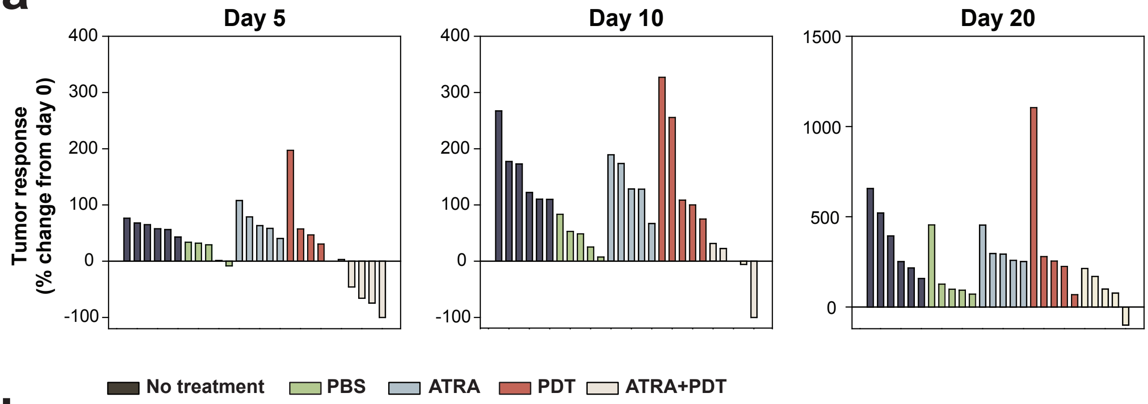


b

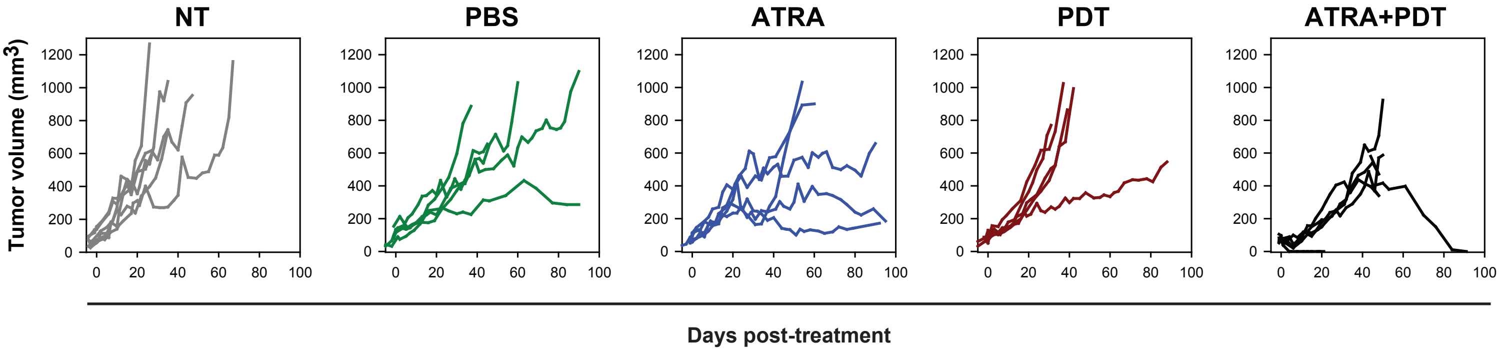


c

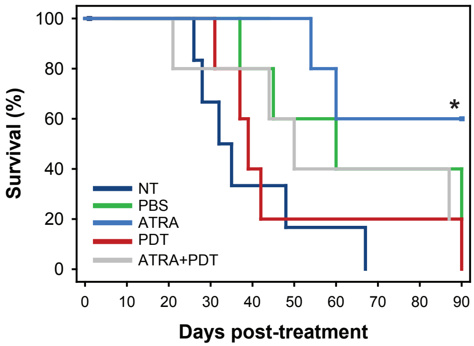


| Treatment group | Mean time to reach endpoint (days) | S.E. | No. of animals |
|---|---|---|---|
| NT | 38.0 | 6.0 | 6 |
| PBS | 64.4 | 11.1 | 5 |
| ATRA | 76.8 | 8.1 | 5 |
| PDT | 47.8 | 10.7 | 5 |
| ATRA+PDT | 58.4 | 13.2 | 5 |

d

**Tumor size (mm³) at day 90**

| PBS | ATRA | PDT | ATRA+PDT |
|---|---|---|---|
| 1098.1 | 658.0 | 1022.4 | 0 |
| 285.9 | 259.9 | | |
| | 172.2 | | |
