## Supplementary Table 1 for "All-*trans* retinoic acid enhances the anti-tumour effects of fimaporfin-based photodynamic therapy"

**Table S1: Significant differentially expressed genes (DEGs) after ATRA+PDT compared to PDT.**

| # | gene | locus | sample_1 | sample_2 | value_1 | value_2 | log2(fold_change) | p_value | q_value |
|---|---|---|---|---|---|---|---|---|---|
| 1 | AADAC | chr3:151347319-151546276 | PDT | ATRA+PDT | 4.18233 | 2.51685 | -0.732689 | 0.0008 | 0.0114972 |
| 2 | AAK1 | chr2:69685126-69870977 | PDT | ATRA+PDT | 1.64626 | 2.29391 | 0.478624 | 0.00005 | 0.00102476 |
| 3 | ABCA2 | chr9:139901685-139931234 | PDT | ATRA+PDT | 12.3197 | 8.26399 | -0.576054 | 0.00005 | 0.00102476 |
| 4 | ABCB10 | chr1:229652328-229694442 | PDT | ATRA+PDT | 8.39398 | 6.2916 | -0.415928 | 0.00055 | 0.0085238 |
| 5 | ABCC1 | chr16:16043433-16236930 | PDT | ATRA+PDT | 21.4222 | 26.0988 | 0.284878 | 0.00325 | 0.0342581 |
| 6 | ABCC2 | chr10:101542462-101611662 | PDT | ATRA+PDT | 1.13309 | 0.335678 | -1.75511 | 0.00005 | 0.00102476 |
| 7 | ABHD6 | chr3:58223258-58280461 | PDT | ATRA+PDT | 2.07661 | 1.27837 | -0.699919 | 0.0012 | 0.0156263 |
| 8 | ABLIM1 | chr10:116190868-116444414 | PDT | ATRA+PDT | 54.6826 | 41.5563 | -0.396015 | 0.0001 | 0.00194145 |
| 9 | ABP1 | chr7:150549572-150558379 | PDT | ATRA+PDT | 2.27635 | 1.40021 | -0.701078 | 0.00135 | 0.0171 |
| 10 | ABTB2 | chr11:34172533-34379555 | PDT | ATRA+PDT | 20.8252 | 27.426 | 0.397214 | 0.0001 | 0.00194145 |
| 11 | ACAA1 | chr3:38080695-38178733 | PDT | ATRA+PDT | 33.7981 | 27.0265 | -0.32257 | 0.0023 | 0.0255827 |
| 12 | ACAT1 | chr11:107992257-108018891 | PDT | ATRA+PDT | 25.2783 | 17.6737 | -0.516291 | 0.00005 | 0.00102476 |
| 13 | ACCN2 | chr12:50451486-50477394 | PDT | ATRA+PDT | 3.70639 | 2.03942 | -0.861857 | 0.00005 | 0.00102476 |
| 14 | ACER2 | chr9:19408924-19452500 | PDT | ATRA+PDT | 8.75569 | 14.5626 | 0.733978 | 0.00005 | 0.00102476 |
| 15 | ACO1 | chr9:32384600-32450832 | PDT | ATRA+PDT | 14.6191 | 11.3469 | -0.365548 | 0.0012 | 0.0156263 |
| 16 | ACOT11 | chr1:55013806-55100417 | PDT | ATRA+PDT | 5.78464 | 4.43867 | -0.382102 | 0.00225 | 0.0252369 |
| 17 | ACSL1 | chr4:185676748-185747215 | PDT | ATRA+PDT | 8.09953 | 4.95421 | -0.709184 | 0.00005 | 0.00102476 |
| 18 | ACSS2 | chr20:33462765-33515769 | PDT | ATRA+PDT | 2.4964 | 1.62324 | -0.620967 | 0.0009 | 0.0125788 |
| 19 | ACVR1 | chr2:158592957-158732374 | PDT | ATRA+PDT | 24.0242 | 29.2492 | 0.283907 | 0.00515 | 0.0488381 |
| 20 | ADAM8 | chr10:135075919-135090407 | PDT | ATRA+PDT | 2.6359 | 1.80406 | -0.547053 | 0.00115 | 0.0151042 |
| 21 | ADAM9 | chr8:38854504-38962779 | PDT | ATRA+PDT | 54.3784 | 68.1282 | 0.32522 | 0.00095 | 0.0131061 |
| 22 | ADAMTS6 | chr5:64444562-64777704 | PDT | ATRA+PDT | 1.65403 | 2.38699 | 0.529205 | 0.00005 | 0.00102476 |
| 23 | AGMAT | chr1:15853351-15911605 | PDT | ATRA+PDT | 12.0281 | 8.43434 | -0.51206 | 0.00045 | 0.00719326 |
| 24 | AGR2 | chr7:16832263-16844738 | PDT | ATRA+PDT | 472.679 | 277.218 | -0.769839 | 0.00005 | 0.00102476 |
| 25 | AGR3 | chr7:16899029-16921613 | PDT | ATRA+PDT | 21.2253 | 13.6572 | -0.636122 | 0.00005 | 0.00102476 |
| 26 | AIFM3 | chr22:21319417-21335649 | PDT | ATRA+PDT | 1.47668 | 0.587014 | -1.33089 | 0.00005 | 0.00102476 |
| 27 | AJUBA | chr14:23440409-23451848 | PDT | ATRA+PDT | 20.9896 | 25.772 | 0.296128 | 0.00435 | 0.0428104 |
| 28 | AK2 | chr1:33473540-33502512 | PDT | ATRA+PDT | 74.9825 | 61.4369 | -0.287448 | 0.0044 | 0.0431831 |
| 29 | AK4 | chr1:65613231-65697828 | PDT | ATRA+PDT | 5.16637 | 2.81494 | -0.876049 | 0.00005 | 0.00102476 |
| 30 | AKAP12 | chr6:151561133-151679694 | PDT | ATRA+PDT | 5.55638 | 17.5063 | 1.65566 | 0.00005 | 0.00102476 |
| 31 | AKR1B10 | chr7:134212343-134226166 | PDT | ATRA+PDT | 1046.65 | 831.293 | -0.332352 | 0.0019 | 0.0221245 |
| 32 | AKR1C1 | chr10:5005453-5020158 | PDT | ATRA+PDT | 50.1133 | 24.6981 | -1.02079 | 0.00005 | 0.00102476 |
| 33 | AKR1C2 | chr10:5029967-5060225 | PDT | ATRA+PDT | 18.5005 | 10.5262 | -0.813584 | 0.00005 | 0.00102476 |
| 34 | AKR1C3 | chr10:5090957-5149878 | PDT | ATRA+PDT | 113.964 | 81.5445 | -0.482918 | 0.00005 | 0.00102476 |
| 35 | AKR7A3 | chr1:19609056-19615280 | PDT | ATRA+PDT | 18.7555 | 34.5418 | 0.881027 | 0.00005 | 0.00102476 |
| 36 | AKR7L | chr1:19592475-19600568 | PDT | ATRA+PDT | 2.7002 | 4.73611 | 0.810636 | 0.00005 | 0.00102476 |
| 37 | ALCAM | chr3:105085556-105295757 | PDT | ATRA+PDT | 10.8802 | 14.7247 | 0.43654 | 0.00005 | 0.00102476 |
| 38 | ALDH1A1 | chr9:75515577-75568233 | PDT | ATRA+PDT | 405.218 | 165.602 | -1.29098 | 0.00005 | 0.00102476 |
| 39 | ALDH1A3 | chr15:101420008-101456830 | PDT | ATRA+PDT | 17.4142 | 37.3478 | 1.10076 | 0.00005 | 0.00102476 |
| 40 | ALDH1B1 | chr9:38392660-38398662 | PDT | ATRA+PDT | 10.8202 | 7.59703 | -0.510217 | 0.00005 | 0.00102476 |
| 41 | ALDH1L1 | chr3:125822407-125899485 | PDT | ATRA+PDT | 0.337508 | 1.1976 | 1.82715 | 0.00005 | 0.00102476 |
| 42 | ALDH2 | chr12:112204690-112247789 | PDT | ATRA+PDT | 32.9702 | 40.9484 | 0.312646 | 0.0022 | 0.024728 |
| 43 | ALDH7A1 | chr5:125877532-125931082 | PDT | ATRA+PDT | 7.3038 | 9.72537 | 0.413106 | 0.00015 | 0.00281494 |
| 44 | ALG2 | chr9:101978706-101984246 | PDT | ATRA+PDT | 17.2807 | 13.2892 | -0.378913 | 0.0004 | 0.00650107 |
| 45 | ALPK3 | chr15:85359910-85416713 | PDT | ATRA+PDT | 14.89 | 19.9106 | 0.419193 | 0.00005 | 0.00102476 |
| 46 | AMBP | chr9:116822407-116840752 | PDT | ATRA+PDT | 2.98429 | 1.4942 | -0.998018 | 0.0004 | 0.00650107 |
| 47 | AMBRA1 | chr11:46417963-46612914 | PDT | ATRA+PDT | 7.53966 | 9.82411 | 0.381827 | 0.0004 | 0.00650107 |
| 48 | AMD1 | chr6:111195986-111216913 | PDT | ATRA+PDT | 73.7935 | 59.5988 | -0.308209 | 0.0019 | 0.0221245 |
| 49 | AMN1 | chr12:31824070-31882108 | PDT | ATRA+PDT | 4.18126 | 6.13014 | 0.551983 | 0.00005 | 0.00102476 |
| 50 | AMOTL2 | chr3:134074189-134093406 | PDT | ATRA+PDT | 7.07425 | 12.6607 | 0.839711 | 0.00005 | 0.00102476 |
| 51 | ANGPT1 | chr8:108261709-108510254 | PDT | ATRA+PDT | 2.09696 | 2.80515 | 0.419779 | 0.00185 | 0.0217319 |
| 52 | ANKMY1 | chr2:241418838-241497405 | PDT | ATRA+PDT | 4.63985 | 6.0158 | 0.374679 | 0.0048 | 0.0461751 |
| 53 | ANKRD13B | chr17:27920526-27948441 | PDT | ATRA+PDT | 6.37219 | 4.92921 | -0.370435 | 0.0041 | 0.040954 |
| 54 | ANKS3 | chr16:4746510-4799397 | PDT | ATRA+PDT | 2.8895 | 3.81208 | 0.399756 | 0.00505 | 0.0482325 |
| 55 | ANKS4B | chr16:21245015-21263750 | PDT | ATRA+PDT | 1.89722 | 0.765152 | -1.31007 | 0.00005 | 0.00102476 |
| 56 | ANKS6 | chr9:101494290-101558794 | PDT | ATRA+PDT | 7.94934 | 6.24206 | -0.348812 | 0.0015 | 0.0184954 |
| 57 | ANXA1 | chr9:75766780-75785307 | PDT | ATRA+PDT | 666.622 | 827.567 | 0.312008 | 0.0031 | 0.0328065 |
| 58 | ANXA10 | chr4:169013687-169108893 | PDT | ATRA+PDT | 116.272 | 212.071 | 0.867047 | 0.00005 | 0.00102476 |
| 59 | ANXA11 | chr10:81914879-81965328 | PDT | ATRA+PDT | 91.7455 | 111.849 | 0.285847 | 0.004 | 0.0401429 |
| 60 | ANXA13 | chr8:124693033-124749647 | PDT | ATRA+PDT | 5.31172 | 1.27258 | -2.06142 | 0.00005 | 0.00102476 |
| 61 | APOBEC3B | chr22:39353526-39388783 | PDT | ATRA+PDT | 4.55594 | 8.63943 | 0.923187 | 0.00005 | 0.00102476 |
| 62 | APOL1 | chr22:36649116-36663577 | PDT | ATRA+PDT | 4.95905 | 9.82204 | 0.98596 | 0.00005 | 0.00102476 |
| 63 | APOL2 | chr22:36622254-36636000 | PDT | ATRA+PDT | 4.72196 | 7.00791 | 0.5696 | 0.00005 | 0.00102476 |
| 64 | APOL6 | chr22:36044423-36064456 | PDT | ATRA+PDT | 2.85787 | 5.44977 | 0.931257 | 0.00005 | 0.00102476 |
| 65 | APRT | chr16:88875876-88878342 | PDT | ATRA+PDT | 200.504 | 160.185 | -0.323894 | 0.0011 | 0.0146092 |
| 66 | AQP5 | chr12:50355278-50359461 | PDT | ATRA+PDT | 12.7712 | 9.55428 | -0.41867 | 0.00085 | 0.0119898 |
| 67 | ARHGAP29 | chr1:94634462-94703307 | PDT | ATRA+PDT | 73.0217 | 93.0992 | 0.350444 | 0.0007 | 0.0103814 |
| 68 | ARHGDIB | chr12:15094949-15114562 | PDT | ATRA+PDT | 20.5853 | 43.954 | 1.09438 | 0.00005 | 0.00102476 |
| 69 | ARHGEF40 | chr14:21538526-21558036 | PDT | ATRA+PDT | 1.39385 | 0.992707 | -0.489635 | 0.0036 | 0.0371409 |
| 70 | ARID5B | chr10:63661012-63856707 | PDT | ATRA+PDT | 1.53658 | 1.16869 | -0.394839 | 0.00465 | 0.0450978 |
| 71 | ARL14 | chr3:160394947-160396235 | PDT | ATRA+PDT | 32.1274 | 57.9683 | 0.851458 | 0.00005 | 0.00102476 |
| 72 | ARL4A | chr7:12726451-12730558 | PDT | ATRA+PDT | 13.1366 | 10.3091 | -0.349671 | 0.00155 | 0.0189586 |
| 73 | ARL6IP4 | chr12:123464879-123467460 | PDT | ATRA+PDT | 50.1261 | 40.2597 | -0.316227 | 0.0019 | 0.0221245 |
| 74 | ARL8A | chr1:202102531-202113871 | PDT | ATRA+PDT | 10.0644 | 14.1927 | 0.495885 | 0.0001 | 0.00194145 |
| 75 | ARRDC3 | chr5:90664540-90716532 | PDT | ATRA+PDT | 5.84636 | 7.96361 | 0.445885 | 0.00035 | 0.00584867 |
| 76 | ARRDC4 | chr15:98503932-98517068 | PDT | ATRA+PDT | 9.12536 | 7.30596 | -0.320808 | 0.00475 | 0.045818 |
| 77 | ARSJ | chr4:114821439-114900878 | PDT | ATRA+PDT | 11.1443 | 8.63381 | -0.368237 | 0.0005 | 0.00788626 |
| 78 | ASB13 | chr10:5680819-5708558 | PDT | ATRA+PDT | 9.28322 | 6.93377 | -0.420986 | 0.00065 | 0.00978942 |
| 79 | ASMTL | chrY:1469423-1522655 | PDT | ATRA+PDT | 11.024 | 7.71413 | -0.515076 | 0.00005 | 0.00102476 |
| 80 | ASS1 | chr9:133320093-133376661 | PDT | ATRA+PDT | 5.71336 | 2.75499 | -1.05229 | 0.00005 | 0.00102476 |
| 81 | ATAD3B | chr1:1407163-1431582 | PDT | ATRA+PDT | 24.6905 | 19.5282 | -0.338397 | 0.00155 | 0.0189586 |
| 82 | ATF3 | chr1:212738675-212794119 | PDT | ATRA+PDT | 97.0536 | 219.94 | 1.18025 | 0.00005 | 0.00102476 |
| 83 | ATHL1 | chr11:289137-295688 | PDT | ATRA+PDT | 14.4644 | 24.4115 | 0.755047 | 0.00005 | 0.00102476 |
| 84 | ATP1B1 | chr1:169075946-169337186 | PDT | ATRA+PDT | 182.748 | 105.694 | -0.789958 | 0.00005 | 0.00102476 |
| 85 | ATP1B3 | chr3:141595469-141645382 | PDT | ATRA+PDT | 52.6144 | 42.1756 | -0.319049 | 0.00135 | 0.0171 |
| 86 | ATP2A3 | chr17:3827162-3867758 | PDT | ATRA+PDT | 3.1096 | 2.14415 | -0.536323 | 0.0001 | 0.00194145 |
| 87 | ATP2B1 | chr12:89981825-90049844 | PDT | ATRA+PDT | 28.9522 | 22.6681 | -0.353007 | 0.0002 | 0.00358322 |
| 88 | ATP5G1 | chr17:46970147-46973232 | PDT | ATRA+PDT | 188.142 | 138.532 | -0.441608 | 0.00005 | 0.00102476 |
| 89 | ATP5G3 | chr2:176040985-176046490 | PDT | ATRA+PDT | 57.8546 | 46.8294 | -0.305018 | 0.00235 | 0.0259765 |
| 90 | ATP6V1B1 | chr2:71162997-71192561 | PDT | ATRA+PDT | 3.24369 | 1.8939 | -0.776278 | 0.0006 | 0.00910057 |
| 91 | ATP9A | chr20:50213313-50384908 | PDT | ATRA+PDT | 27.2854 | 43.0077 | 0.656466 | 0.00005 | 0.00102476 |
| 92 | ATXN2 | chr12:111890017-112037480 | PDT | ATRA+PDT | 11.3362 | 14.1905 | 0.323992 | 0.002 | 0.0229388 |
| 93 | AURKA | chr20:54944444-54967351 | PDT | ATRA+PDT | 72.5295 | 58.7749 | -0.303368 | 0.0017 | 0.0202822 |
| 94 | AXIN2 | chr17:63524682-63557740 | PDT | ATRA+PDT | 14.4559 | 4.59396 | -1.65385 | 0.00005 | 0.00102476 |
| 95 | AZGP1 | chr7:99564349-99573735 | PDT | ATRA+PDT | 9.89214 | 6.2095 | -0.671806 | 0.00005 | 0.00102476 |
| 96 | B2M | chr15:45003684-45010357 | PDT | ATRA+PDT | 459.14 | 652.756 | 0.507611 | 0.00005 | 0.00102476 |
| 97 | B3GNT5 | chr3:182895830-183145855 | PDT | ATRA+PDT | 26.8585 | 37.6283 | 0.486439 | 0.00005 | 0.00102476 |
| 98 | B7H6 | chr11:17373308-17398868 | PDT | ATRA+PDT | 9.70913 | 6.97765 | -0.4766 | 0.00005 | 0.00102476 |
| 99 | BACE2 | chr21:42539727-42648524 | PDT | ATRA+PDT | 13.7798 | 10.5462 | -0.385838 | 0.0008 | 0.0114972 |
| 100 | BAG3 | chr10:121410881-121437329 | PDT | ATRA+PDT | 199.728 | 279.991 | 0.487348 | 0.00005 | 0.00102476 |
| 101 | BAIAP2L1 | chr7:97910978-98030427 | PDT | ATRA+PDT | 54.0082 | 71.1708 | 0.398108 | 0.0002 | 0.00358322 |
| 102 | BAIAP2L2 | chr22:38480895-38506676 | PDT | ATRA+PDT | 4.25175 | 2.3843 | -0.834492 | 0.00005 | 0.00102476 |

**Table S1: Significant differentially expressed genes (DEGs) after ATRA+PDT compared to PDT.**

| # | gene | locus | sample_1 | sample_2 | value_1 | value_2 | log2(fold_change) | p_value | q_value |
|---|---|---|---|---|---|---|---|---|---|
| 103 | BAZ2B | chr2:160175489-160473059 | PDT | ATRA+PDT | 7.00159 | 9.66999 | 0.465831 | 0.00005 | 0.00102476 |
| 104 | BBC3 | chr19:47724078-47736023 | PDT | ATRA+PDT | 40.8982 | 50.9416 | 0.316808 | 0.0023 | 0.0255827 |
| 105 | BBS4 | chr15:72978519-73030817 | PDT | ATRA+PDT | 6.72558 | 8.77988 | 0.384542 | 0.00195 | 0.0225592 |
| 106 | BCAM | chr19:45312337-45324678 | PDT | ATRA+PDT | 36.6104 | 49.3033 | 0.429432 | 0.00005 | 0.00102476 |
| 107 | BCAS1 | chr20:52560078-52687304 | PDT | ATRA+PDT | 17.0823 | 27.5657 | 0.690368 | 0.00005 | 0.00102476 |
| 108 | BCAS3 | chr17:58755171-59470199 | PDT | ATRA+PDT | 1.34242 | 2.28822 | 0.769385 | 0.00005 | 0.00102476 |
| 109 | BCDIN3D | chr12:50222325-50236912 | PDT | ATRA+PDT | 1.17038 | 1.78497 | 0.608924 | 0.00245 | 0.0267496 |
| 110 | BCL10 | chr1:85731459-85743771 | PDT | ATRA+PDT | 21.8727 | 35.4025 | 0.694719 | 0.00005 | 0.00102476 |
| 111 | BCL3 | chr19:45251977-45263301 | PDT | ATRA+PDT | 10.2934 | 17.5884 | 0.77291 | 0.00005 | 0.00102476 |
| 112 | BCL6 | chr3:187416046-187463513 | PDT | ATRA+PDT | 6.68803 | 8.88905 | 0.410447 | 0.0007 | 0.0103814 |
| 113 | BGN | chrX:152760346-152775004 | PDT | ATRA+PDT | 16.5827 | 6.73104 | -1.30078 | 0.00005 | 0.00102476 |
| 114 | BIRC3 | chr11:102188180-102210135 | PDT | ATRA+PDT | 1.25282 | 2.58 | 1.04219 | 0.00005 | 0.00102476 |
| 115 | BMP4 | chr14:54416454-54423554 | PDT | ATRA+PDT | 124.779 | 91.4887 | -0.447709 | 0.00005 | 0.00102476 |
| 116 | BOLA3 | chr2:74362527-74375039 | PDT | ATRA+PDT | 48.6335 | 37.0183 | -0.393712 | 0.00285 | 0.0305545 |
| 117 | BRPF3 | chr6:36164549-36200567 | PDT | ATRA+PDT | 19.2699 | 24.3925 | 0.340091 | 0.00075 | 0.0109256 |
| 118 | BTBD10 | chr11:13409555-13484838 | PDT | ATRA+PDT | 29.8294 | 39.4629 | 0.403762 | 0.00005 | 0.00102476 |
| 119 | BTG1 | chr12:92378751-92539673 | PDT | ATRA+PDT | 22.097 | 27.4426 | 0.312566 | 0.00175 | 0.0207397 |
| 120 | BTN2A1 | chr6:26458152-26482737 | PDT | ATRA+PDT | 11.2454 | 14.5205 | 0.368763 | 0.00115 | 0.0151042 |
| 121 | BTN3A1 | chr6:26402464-26415444 | PDT | ATRA+PDT | 0.773794 | 1.7837 | 1.20485 | 0.00005 | 0.00102476 |
| 122 | BTN3A2 | chr6:26365397-26378548 | PDT | ATRA+PDT | 1.8277 | 2.68976 | 0.557449 | 0.00045 | 0.00719326 |
| 123 | BTNL9 | chr5:180467224-180488523 | PDT | ATRA+PDT | 6.38227 | 4.6704 | -0.450523 | 0.0004 | 0.00650107 |
| 124 | C10orf10 | chr10:45455218-45490172 | PDT | ATRA+PDT | 3.78595 | 9.13062 | 1.27006 | 0.00005 | 0.00102476 |
| 125 | C10orf125 | chr10:135168657-135171529 | PDT | ATRA+PDT | 15.3108 | 9.96173 | -0.620081 | 0.001 | 0.0135165 |
| 126 | C10orf47 | chr10:11865396-11936709 | PDT | ATRA+PDT | 13.1369 | 20.2372 | 0.623382 | 0.00005 | 0.00102476 |
| 127 | C10orf54 | chr10:73156690-73575704 | PDT | ATRA+PDT | 1.13266 | 2.95064 | 1.38132 | 0.00005 | 0.00102476 |
| 128 | C11orf68 | chr11:65684282-65686531 | PDT | ATRA+PDT | 15.6446 | 20.922 | 0.419358 | 0.0005 | 0.00788626 |
| 129 | C11orf75 | chr11:93211638-93276546 | PDT | ATRA+PDT | 10.0852 | 7.34606 | -0.457202 | 0.0041 | 0.040954 |
| 130 | C12orf35 | chr12:32112352-32146043 | PDT | ATRA+PDT | 24.3656 | 43.641 | 0.840838 | 0.00005 | 0.00102476 |
| 131 | C12orf36 | chr12:13523604-13529679 | PDT | ATRA+PDT | 1.77168 | 1.09353 | -0.696119 | 0.00075 | 0.0109256 |
| 132 | C14orf43 | chr14:74181824-74253896 | PDT | ATRA+PDT | 3.43389 | 4.65663 | 0.439443 | 0.0001 | 0.00194145 |
| 133 | C15orf52 | chr15:40623652-40633168 | PDT | ATRA+PDT | 7.17081 | 4.87871 | -0.555636 | 0.00005 | 0.00102476 |
| 134 | C16orf52 | chr16:22019455-22095972 | PDT | ATRA+PDT | 6.24092 | 8.43014 | 0.433798 | 0.0001 | 0.00194145 |
| 135 | C17orf28 | chr17:72946838-72968900 | PDT | ATRA+PDT | 6.1793 | 4.19248 | -0.559639 | 0.00005 | 0.00102476 |
| 136 | C19orf66 | chr19:10196805-10213425 | PDT | ATRA+PDT | 1.11955 | 2.17595 | 0.958731 | 0.00045 | 0.00719326 |
| 137 | C1orf116 | chr1:207191865-207206101 | PDT | ATRA+PDT | 17.3812 | 23.2522 | 0.419839 | 0.0002 | 0.00358322 |
| 138 | C1orf130 | chr1:24882566-24935818 | PDT | ATRA+PDT | 1.04809 | 2.20534 | 1.07323 | 0.00005 | 0.00102476 |
| 139 | C20orf96 | chr20:251503-271419 | PDT | ATRA+PDT | 1.39756 | 2.5256 | 0.853715 | 0.00135 | 0.0171 |
| 140 | C22orf46 | chr22:42086546-42094140 | PDT | ATRA+PDT | 2.23184 | 3.21502 | 0.526595 | 0.00005 | 0.00102476 |
| 141 | C2orf42 | chr2:70377016-70418151 | PDT | ATRA+PDT | 2.01778 | 2.86738 | 0.506964 | 0.00425 | 0.0420979 |
| 142 | C2orf49 | chr2:105954012-105961984 | PDT | ATRA+PDT | 12.3893 | 16.0922 | 0.377273 | 0.00375 | 0.0383182 |
| 143 | C2orf72 | chr2:231902280-231914427 | PDT | ATRA+PDT | 1.31922 | 0.840518 | -0.65033 | 0.00305 | 0.0323415 |
| 144 | C5orf41 | chr5:172483354-172566291 | PDT | ATRA+PDT | 7.08993 | 9.68801 | 0.45043 | 0.00005 | 0.00102476 |
| 145 | C6orf1 | chr6:34214156-34216904 | PDT | ATRA+PDT | 4.87146 | 7.2764 | 0.578871 | 0.003 | 0.0318746 |
| 146 | C7orf46 | chr7:23719748-23742269 | PDT | ATRA+PDT | 14.7916 | 19.7694 | 0.418488 | 0.00075 | 0.0109256 |
| 147 | C8orf84 | chr8:73976777-74005507 | PDT | ATRA+PDT | 6.66596 | 4.13785 | -0.687932 | 0.00005 | 0.00102476 |
| 148 | C9orf152 | chr9:112961845-112970413 | PDT | ATRA+PDT | 5.62529 | 3.32061 | -0.760478 | 0.00005 | 0.00102476 |
| 149 | CA9 | chr9:35673914-35681154 | PDT | ATRA+PDT | 34.2291 | 25.0317 | -0.451469 | 0.0001 | 0.00194145 |
| 150 | CALB2 | chr16:71392615-71424342 | PDT | ATRA+PDT | 89.8282 | 123.84 | 0.46324 | 0.00005 | 0.00102476 |
| 151 | CAMKK1 | chr17:3763616-3796337 | PDT | ATRA+PDT | 5.71221 | 4.31302 | -0.405349 | 0.00185 | 0.0217319 |
| 152 | CAPN5 | chr11:76777991-76837198 | PDT | ATRA+PDT | 4.82162 | 3.23871 | -0.574099 | 0.00005 | 0.00102476 |
| 153 | CAPN9 | chr1:230883129-230937749 | PDT | ATRA+PDT | 1.45467 | 0.70276 | -1.04958 | 0.00145 | 0.0179618 |
| 154 | CARD10 | chr22:37886399-37915210 | PDT | ATRA+PDT | 5.04292 | 6.86843 | 0.445722 | 0.0002 | 0.00358322 |
| 155 | CARD6 | chr5:40841409-40855456 | PDT | ATRA+PDT | 0.883466 | 1.61116 | 0.866857 | 0.00005 | 0.00102476 |
| 156 | CASP4 | chr11:104813593-104839325 | PDT | ATRA+PDT | 48.7865 | 59.6521 | 0.290091 | 0.00425 | 0.0420979 |
| 157 | CASP7 | chr10:115438934-115490664 | PDT | ATRA+PDT | 20.5481 | 27.0151 | 0.394762 | 0.0001 | 0.00194145 |
| 158 | CASP9 | chr1:15818768-15851285 | PDT | ATRA+PDT | 3.65432 | 5.51936 | 0.594897 | 0.00005 | 0.00102476 |
| 159 | CAV1 | chr7:116164838-116201239 | PDT | ATRA+PDT | 10.8401 | 6.42212 | -0.755251 | 0.00005 | 0.00102476 |
| 160 | CAV2 | chr7:116139654-116148595 | PDT | ATRA+PDT | 19.9623 | 13.9617 | -0.515804 | 0.00005 | 0.00102476 |
| 161 | CBLB | chr3:105377108-105587887 | PDT | ATRA+PDT | 2.45668 | 3.47512 | 0.500354 | 0.00015 | 0.00281494 |
| 162 | CBX7 | chr22:39526778-39548538 | PDT | ATRA+PDT | 1.32238 | 2.05347 | 0.634922 | 0.00025 | 0.00439786 |
| 163 | CCDC112 | chr5:114602884-114632458 | PDT | ATRA+PDT | 8.91704 | 11.5731 | 0.376144 | 0.0016 | 0.0194588 |
| 164 | CCDC117 | chr22:29168661-29185283 | PDT | ATRA+PDT | 32.13 | 43.7731 | 0.446124 | 0.00005 | 0.00102476 |
| 165 | CCDC28A | chr6:139046347-139114456 | PDT | ATRA+PDT | 13.8248 | 17.8438 | 0.368169 | 0.0033 | 0.0346484 |
| 166 | CCDC34 | chr11:27360060-27384795 | PDT | ATRA+PDT | 21.2716 | 16.3716 | -0.377735 | 0.00165 | 0.0198632 |
| 167 | CCDC68 | chr18:52568739-52626739 | PDT | ATRA+PDT | 3.26671 | 2.34128 | -0.480542 | 0.0003 | 0.00511725 |
| 168 | CCDC86 | chr11:60609428-60623444 | PDT | ATRA+PDT | 48.8511 | 37.736 | -0.372449 | 0.0002 | 0.00358322 |
| 169 | CCDC88C | chr14:91737666-91884188 | PDT | ATRA+PDT | 4.44669 | 5.82279 | 0.38898 | 0.0003 | 0.00511725 |
| 170 | CCL15 | chr17:34310691-34329084 | PDT | ATRA+PDT | 15.6012 | 4.77259 | -1.70881 | 0.00005 | 0.00102476 |
| 171 | CCNA2 | chr4:122722471-122745088 | PDT | ATRA+PDT | 58.14 | 44.5609 | -0.383754 | 0.00105 | 0.0140676 |
| 172 | CCNDBP1 | chr15:43477465-43489375 | PDT | ATRA+PDT | 8.45993 | 12.1829 | 0.526144 | 0.00005 | 0.00102476 |
| 173 | CCNF | chr16:2479394-2508859 | PDT | ATRA+PDT | 12.2325 | 8.17091 | -0.582151 | 0.00005 | 0.00102476 |
| 174 | CCNG2 | chr4:78078356-78091213 | PDT | ATRA+PDT | 11.3243 | 15.0479 | 0.410131 | 0.0001 | 0.00194145 |
| 175 | CCNJL | chr5:159678670-159739573 | PDT | ATRA+PDT | 3.33525 | 4.28638 | 0.361966 | 0.00435 | 0.0428104 |
| 176 | CD44 | chr11:35160416-35253949 | PDT | ATRA+PDT | 98.6039 | 74.61 | -0.402276 | 0.00005 | 0.00102476 |
| 177 | CD55 | chr1:207494816-207534311 | PDT | ATRA+PDT | 362.706 | 446.974 | 0.30139 | 0.00335 | 0.0351043 |
| 178 | CD58 | chr1:117057155-117113715 | PDT | ATRA+PDT | 27.0231 | 20.3912 | -0.406248 | 0.00095 | 0.0131061 |
| 179 | CD68 | chr17:7482804-7485429 | PDT | ATRA+PDT | 6.98815 | 9.08423 | 0.378455 | 0.00265 | 0.0286404 |
| 180 | CDC14B | chr9:99262394-99382112 | PDT | ATRA+PDT | 10.515 | 14.1561 | 0.428976 | 0.00005 | 0.00102476 |
| 181 | CDC25A | chr3:48198667-48229801 | PDT | ATRA+PDT | 19.8524 | 15.7683 | -0.332287 | 0.00125 | 0.0161202 |
| 182 | CDC42BPG | chr11:64591661-64612041 | PDT | ATRA+PDT | 2.57052 | 3.90719 | 0.604068 | 0.00005 | 0.00102476 |
| 183 | CDC42EP1 | chr22:37956470-37965410 | PDT | ATRA+PDT | 29.3322 | 41.0904 | 0.486314 | 0.00005 | 0.00102476 |
| 184 | CDC42EP2 | chr11:65082288-65089900 | PDT | ATRA+PDT | 29.0189 | 40.3226 | 0.474595 | 0.00005 | 0.00102476 |
| 185 | CDHR5 | chr11:616564-625067 | PDT | ATRA+PDT | 3.37929 | 2.60197 | -0.377115 | 0.0046 | 0.0446535 |
| 186 | CDK6 | chr7:92234234-92465941 | PDT | ATRA+PDT | 16.7786 | 20.8185 | 0.311246 | 0.001 | 0.0135165 |
| 187 | CDKN2AIP | chr4:184365788-184369049 | PDT | ATRA+PDT | 11.8498 | 14.9428 | 0.334588 | 0.0029 | 0.0310283 |
| 188 | CDKN2B | chr9:21994789-22121093 | PDT | ATRA+PDT | 9.39304 | 18.1238 | 0.94822 | 0.00005 | 0.00102476 |
| 189 | CEACAM1 | chr19:43011457-43032661 | PDT | ATRA+PDT | 18.4867 | 22.8958 | 0.308597 | 0.0025 | 0.027212 |
| 190 | CEACAM5 | chr19:42212529-42234437 | PDT | ATRA+PDT | 12.0597 | 5.06265 | -1.25223 | 0.00005 | 0.00102476 |
| 191 | CEACAM6 | chr19:42259397-42276113 | PDT | ATRA+PDT | 47.7562 | 22.2689 | -1.10066 | 0.00005 | 0.00102476 |
| 192 | CEBPA | chr19:33790839-33793430 | PDT | ATRA+PDT | 7.40735 | 5.55553 | -0.415033 | 0.0014 | 0.0175049 |
| 193 | CEBPD | chr8:48649475-48650726 | PDT | ATRA+PDT | 6.30402 | 9.0375 | 0.519651 | 0.0003 | 0.00511725 |
| 194 | CELSR1 | chr22:46756730-46933067 | PDT | ATRA+PDT | 0.901511 | 1.32008 | 0.550213 | 0.00005 | 0.00102476 |
| 195 | CEP120 | chr5:122680578-122759286 | PDT | ATRA+PDT | 5.71684 | 7.40525 | 0.373331 | 0.00065 | 0.00978942 |
| 196 | CEP128 | chr14:80962820-81405884 | PDT | ATRA+PDT | 1.26871 | 1.81083 | 0.513293 | 0.0017 | 0.0202822 |
| 197 | CEP135 | chr4:56815036-56899527 | PDT | ATRA+PDT | 4.08846 | 5.34231 | 0.385909 | 0.0008 | 0.0114972 |
| 198 | CEP170 | chr1:243287729-243418708 | PDT | ATRA+PDT | 5.2673 | 6.54725 | 0.313827 | 0.00355 | 0.0368028 |
| 199 | CEP41 | chr7:130036374-130080854 | PDT | ATRA+PDT | 2.04751 | 3.43399 | 0.746012 | 0.00005 | 0.00102476 |
| 200 | CEP70 | chr3:138213186-138313129 | PDT | ATRA+PDT | 3.35163 | 4.6257 | 0.464808 | 0.0007 | 0.0103814 |
| 201 | CEP95 | chr17:62503157-62534062 | PDT | ATRA+PDT | 20.5436 | 25.4745 | 0.310362 | 0.0025 | 0.027212 |
| 202 | CFTR | chr7:117120016-117308718 | PDT | ATRA+PDT | 14.818 | 11.4256 | -0.375083 | 0.00065 | 0.00978942 |
| 203 | CGREF1 | chr2:27309610-27341995 | PDT | ATRA+PDT | 12.4164 | 7.361 | -0.754276 | 0.00005 | 0.00102476 |
| 204 | CHAC1 | chr15:41245635-41248717 | PDT | ATRA+PDT | 39.802 | 55.5993 | 0.482225 | 0.00005 | 0.00102476 |

**Table S1: Significant differentially expressed genes (DEGs) after ATRA+PDT compared to PDT.**

| # | gene | locus | sample_1 | sample_2 | value_1 | value_2 | log2(fold_change) | p_value | q_value |
|---|---|---|---|---|---|---|---|---|---|
| 205 | CHD2 | chr15:93443550-93571237 | PDT | ATRA+PDT | 32.4564 | 43.5391 | 0.423808 | 0.00005 | 0.00102476 |
| 206 | CHORDC1 | chr11:89933596-89956532 | PDT | ATRA+PDT | 94.5056 | 117.498 | 0.314168 | 0.00135 | 0.0171 |
| 207 | CHPF | chr2:220403668-220408487 | PDT | ATRA+PDT | 4.02569 | 3.01637 | -0.416425 | 0.0033 | 0.0346484 |
| 208 | CHPT1 | chr12:102091416-102133250 | PDT | ATRA+PDT | 15.0998 | 8.0349 | -0.910182 | 0.00005 | 0.00102476 |
| 209 | CHST4 | chr16:71560022-71572493 | PDT | ATRA+PDT | 0.355725 | 1.21076 | 1.76709 | 0.00015 | 0.00281494 |
| 210 | CIB1 | chr15:90773476-90777279 | PDT | ATRA+PDT | 112.778 | 140.187 | 0.313869 | 0.00245 | 0.0267496 |
| 211 | CIRBP | chr19:1267469-1274809 | PDT | ATRA+PDT | 41.7846 | 53.1301 | 0.346557 | 0.00195 | 0.0225592 |
| 212 | CKAP2L | chr2:113495443-113522254 | PDT | ATRA+PDT | 9.43968 | 7.47605 | -0.336462 | 0.0036 | 0.0371409 |
| 213 | CKB | chr14:103985994-103989196 | PDT | ATRA+PDT | 72.3156 | 50.3455 | -0.522445 | 0.00005 | 0.00102476 |
| 214 | CLCF1 | chr11:67085309-67159158 | PDT | ATRA+PDT | 33.6729 | 52.7771 | 0.648324 | 0.00005 | 0.00102476 |
| 215 | CLDN1 | chr3:190023489-190040235 | PDT | ATRA+PDT | 15.3463 | 21.516 | 0.487519 | 0.00005 | 0.00102476 |
| 216 | CLDN12 | chr7:90032647-90045268 | PDT | ATRA+PDT | 29.5243 | 42.4496 | 0.523846 | 0.00005 | 0.00102476 |
| 217 | CLDN2 | chrX:106143292-106174091 | PDT | ATRA+PDT | 19.9892 | 11.7522 | -0.766294 | 0.00005 | 0.00102476 |
| 218 | CLIC5 | chr6:45866189-46048085 | PDT | ATRA+PDT | 1.02119 | 1.59094 | 0.639622 | 0.00235 | 0.0259765 |
| 219 | CLIP2 | chr7:73703804-73820273 | PDT | ATRA+PDT | 2.2237 | 1.393 | -0.674764 | 0.00005 | 0.00102476 |
| 220 | CLK1 | chr2:201717731-201729467 | PDT | ATRA+PDT | 103.345 | 137.873 | 0.415875 | 0.00005 | 0.00102476 |
| 221 | CLK4 | chr5:178029664-178054054 | PDT | ATRA+PDT | 6.81798 | 8.81923 | 0.371308 | 0.00275 | 0.0296013 |
| 222 | CLMN | chr14:95648275-95786245 | PDT | ATRA+PDT | 2.9246 | 3.76489 | 0.364365 | 0.00055 | 0.0085238 |
| 223 | CLN8 | chr8:1711869-1734736 | PDT | ATRA+PDT | 2.529 | 1.89329 | -0.417668 | 0.00105 | 0.0140676 |
| 224 | CLP1 | chr11:57425215-57429337 | PDT | ATRA+PDT | 13.7435 | 9.95338 | -0.465492 | 0.0003 | 0.00511725 |
| 225 | CLRN3 | chr10:129676113-129691211 | PDT | ATRA+PDT | 7.69706 | 5.49485 | -0.486227 | 0.004 | 0.0401429 |
| 226 | CLU | chr8:27454433-27472328 | PDT | ATRA+PDT | 4.61759 | 10.0572 | 1.12302 | 0.00005 | 0.00102476 |
| 227 | CMTM3 | chr16:66637934-66647795 | PDT | ATRA+PDT | 6.29977 | 9.54492 | 0.599434 | 0.00005 | 0.00102476 |
| 228 | CMTM7 | chr3:32433162-32496333 | PDT | ATRA+PDT | 17.8085 | 23.9263 | 0.426029 | 0.00045 | 0.00719326 |
| 229 | CNGA1 | chr4:47937993-48014961 | PDT | ATRA+PDT | 0.664166 | 1.77801 | 1.42065 | 0.00005 | 0.00102476 |
| 230 | CNKSR1 | chr1:26503980-26516375 | PDT | ATRA+PDT | 8.17194 | 10.6539 | 0.382633 | 0.00145 | 0.0179618 |
| 231 | CNTRL | chr9:123850573-123939886 | PDT | ATRA+PDT | 5.08383 | 6.99577 | 0.460567 | 0.0001 | 0.00194145 |
| 232 | COL17A1 | chr10:105791045-105845638 | PDT | ATRA+PDT | 13.9053 | 9.03744 | -0.621652 | 0.00005 | 0.00102476 |
| 233 | COL18A1 | chr21:46825096-46933634 | PDT | ATRA+PDT | 2.93481 | 2.10741 | -0.477797 | 0.00015 | 0.00281494 |
| 234 | COTL1 | chr16:84599203-84651669 | PDT | ATRA+PDT | 110.534 | 141.466 | 0.355972 | 0.0002 | 0.00358322 |
| 235 | CPT1A | chr11:68522087-68609399 | PDT | ATRA+PDT | 47.0072 | 30.818 | -0.60911 | 0.00005 | 0.00102476 |
| 236 | CRABP2 | chr1:156669399-156675608 | PDT | ATRA+PDT | 154.472 | 108.962 | -0.50352 | 0.00005 | 0.00102476 |
| 237 | CREB5 | chr7:28338939-28865511 | PDT | ATRA+PDT | 3.6883 | 8.02894 | 1.12225 | 0.00005 | 0.00102476 |
| 238 | CRELD1 | chr3:9975523-9987097 | PDT | ATRA+PDT | 4.03205 | 2.67534 | -0.59179 | 0.0005 | 0.00788626 |
| 239 | CRISPLD2 | chr16:84853586-84943116 | PDT | ATRA+PDT | 0.782842 | 1.32989 | 0.764514 | 0.0003 | 0.00511725 |
| 240 | CSPG4 | chr15:75966662-76005189 | PDT | ATRA+PDT | 3.78188 | 1.95068 | -0.955129 | 0.00005 | 0.00102476 |
| 241 | CSRNP1 | chr3:39183341-39195102 | PDT | ATRA+PDT | 4.48779 | 7.52922 | 0.746495 | 0.00005 | 0.00102476 |
| 242 | CTSE | chr1:206317458-206332104 | PDT | ATRA+PDT | 8.26341 | 5.46912 | -0.595429 | 0.00005 | 0.00102476 |
| 243 | CTSS | chr1:150702671-150738433 | PDT | ATRA+PDT | 0.43546 | 1.76088 | 2.01569 | 0.00005 | 0.00102476 |
| 244 | CTTNBP2NL | chr1:112938799-113003786 | PDT | ATRA+PDT | 30.9598 | 43.1174 | 0.477873 | 0.00005 | 0.00102476 |
| 245 | CXCL1 | chr4:74735108-74737019 | PDT | ATRA+PDT | 5.62294 | 11.2404 | 0.999296 | 0.00005 | 0.00102476 |
| 246 | CXCL16 | chr17:4634722-4643223 | PDT | ATRA+PDT | 3.56652 | 6.74273 | 0.918813 | 0.00005 | 0.00102476 |
| 247 | CXCL2 | chr4:74962753-74964997 | PDT | ATRA+PDT | 1.67488 | 2.92825 | 0.805984 | 0.00395 | 0.0398659 |
| 248 | CXCL3 | chr4:74902311-74904490 | PDT | ATRA+PDT | 3.00394 | 5.38244 | 0.841402 | 0.00015 | 0.00281494 |
| 249 | CXCR4 | chr2:136871918-136875725 | PDT | ATRA+PDT | 17.1971 | 12.0182 | -0.51694 | 0.0001 | 0.00194145 |
| 250 | CXXC5 | chr5:139028300-139062680 | PDT | ATRA+PDT | 19.0937 | 13.8853 | -0.459533 | 0.0002 | 0.00358322 |
| 251 | CYB5RL | chr1:54638026-54665746 | PDT | ATRA+PDT | 2.05215 | 1.21173 | -0.760073 | 0.00025 | 0.00439786 |
| 252 | CYP1A1 | chr15:75011882-75017877 | PDT | ATRA+PDT | 6.05798 | 9.12181 | 0.590483 | 0.00005 | 0.00102476 |
| 253 | CYP24A1 | chr20:52769987-52790516 | PDT | ATRA+PDT | 0.491874 | 2.60664 | 2.40583 | 0.00005 | 0.00102476 |
| 254 | CYP2B6 | chr19:41497203-41524301 | PDT | ATRA+PDT | 0.928489 | 1.77284 | 0.933107 | 0.00015 | 0.00281494 |
| 255 | CYP2S1 | chr19:41699114-41713444 | PDT | ATRA+PDT | 23.4328 | 31.7265 | 0.437162 | 0.0001 | 0.00194145 |
| 256 | CYP3A5 | chr7:99245812-99277621 | PDT | ATRA+PDT | 65.7934 | 91.3335 | 0.473202 | 0.00005 | 0.00102476 |
| 257 | CYR61 | chr1:86046443-86049648 | PDT | ATRA+PDT | 4.62284 | 7.36744 | 0.672385 | 0.00005 | 0.00102476 |
| 258 | DAPK1 | chr9:90112755-90323549 | PDT | ATRA+PDT | 9.61867 | 13.932 | 0.53449 | 0.00005 | 0.00102476 |
| 259 | DAPP1 | chr4:100737980-100791346 | PDT | ATRA+PDT | 1.78116 | 2.99943 | 0.751871 | 0.00005 | 0.00102476 |
| 260 | DDC | chr7:50526133-50633154 | PDT | ATRA+PDT | 11.9731 | 4.23146 | -1.50057 | 0.00005 | 0.00102476 |
| 261 | DDIT4 | chr10:74033676-74035797 | PDT | ATRA+PDT | 177.845 | 221.999 | 0.319935 | 0.00115 | 0.0151042 |
| 262 | DDX60L | chr4:169277885-169401638 | PDT | ATRA+PDT | 1.0759 | 1.9709 | 0.873306 | 0.00005 | 0.00102476 |
| 263 | DEDD2 | chr19:42702751-42721813 | PDT | ATRA+PDT | 127.921 | 162.117 | 0.341777 | 0.0006 | 0.00910057 |
| 264 | DEGS2 | chr14:100612752-100626012 | PDT | ATRA+PDT | 3.86987 | 2.3815 | -0.700417 | 0.0022 | 0.024728 |
| 265 | DENND3 | chr8:142138719-142205900 | PDT | ATRA+PDT | 7.69182 | 10.605 | 0.463353 | 0.00005 | 0.00102476 |
| 266 | DEPDC1B | chr5:59892738-59995993 | PDT | ATRA+PDT | 17.2863 | 22.3816 | 0.372682 | 0.0003 | 0.00511725 |
| 267 | DEXI | chr16:11022747-11036257 | PDT | ATRA+PDT | 8.73179 | 6.4536 | -0.436174 | 0.00065 | 0.00978942 |
| 268 | DGAT2 | chr11:75479777-75512581 | PDT | ATRA+PDT | 5.80363 | 3.82 | -0.603383 | 0.00005 | 0.00102476 |
| 269 | DHODH | chr16:72042642-72059316 | PDT | ATRA+PDT | 8.02902 | 6.1418 | -0.386561 | 0.00275 | 0.0296013 |
| 270 | DHRS3 | chr1:12627938-12677820 | PDT | ATRA+PDT | 7.01205 | 21.5381 | 1.61899 | 0.00005 | 0.00102476 |
| 271 | DIP2C | chr10:320129-735608 | PDT | ATRA+PDT | 1.47699 | 2.23676 | 0.598746 | 0.00005 | 0.00102476 |
| 272 | DKK1 | chr10:54074040-54077417 | PDT | ATRA+PDT | 67.0006 | 99.9039 | 0.576367 | 0.00005 | 0.00102476 |
| 273 | DMTF1 | chr7:86781676-86849031 | PDT | ATRA+PDT | 23.6529 | 30.6114 | 0.372051 | 0.00125 | 0.0161202 |
| 274 | DNAJA1 | chr9:33025208-33039062 | PDT | ATRA+PDT | 581.316 | 721.099 | 0.310875 | 0.003 | 0.0318746 |
| 275 | DNAJA4 | chr15:78556486-78574538 | PDT | ATRA+PDT | 57.131 | 87.063 | 0.607786 | 0.00005 | 0.00102476 |
| 276 | DNAJB1 | chr19:14625581-14629201 | PDT | ATRA+PDT | 531.15 | 857.804 | 0.691529 | 0.00005 | 0.00102476 |
| 277 | DNAJB4 | chr1:78470635-78482995 | PDT | ATRA+PDT | 72.7418 | 98.3063 | 0.4345 | 0.00005 | 0.00102476 |
| 278 | DNAJB9 | chr7:108210188-108215294 | PDT | ATRA+PDT | 37.4037 | 26.737 | -0.484344 | 0.00005 | 0.00102476 |
| 279 | DNAJC22 | chr12:49741040-49745685 | PDT | ATRA+PDT | 6.21878 | 4.60419 | -0.433685 | 0.0018 | 0.0212615 |
| 280 | DNAJC3 | chr13:96329392-96447243 | PDT | ATRA+PDT | 19.6839 | 15.3869 | -0.355314 | 0.0004 | 0.00650107 |
| 281 | DNMT3A | chr2:25455829-25565459 | PDT | ATRA+PDT | 1.09346 | 1.85713 | 0.764178 | 0.00005 | 0.00102476 |
| 282 | DOCK5 | chr8:25042286-25270619 | PDT | ATRA+PDT | 5.07166 | 6.63786 | 0.38826 | 0.0002 | 0.00358322 |
| 283 | DPEP1 | chr16:89679715-89704839 | PDT | ATRA+PDT | 6.81875 | 2.92142 | -1.22284 | 0.00005 | 0.00102476 |
| 284 | DPP4 | chr2:162848754-162931052 | PDT | ATRA+PDT | 8.24666 | 4.60386 | -0.840966 | 0.00005 | 0.00102476 |
| 285 | DPP9 | chr19:4675243-4723855 | PDT | ATRA+PDT | 18.5075 | 23.507 | 0.344986 | 0.0006 | 0.00910057 |
| 286 | DST | chr6:56322784-56507694 | PDT | ATRA+PDT | 13.8114 | 10.3563 | -0.415349 | 0.00005 | 0.00102476 |
| 287 | DTX3L | chr3:122246759-122294049 | PDT | ATRA+PDT | 14.324 | 21.944 | 0.615392 | 0.00005 | 0.00102476 |
| 288 | DUSP1 | chr5:172195092-172198203 | PDT | ATRA+PDT | 37.0726 | 70.0712 | 0.918467 | 0.00005 | 0.00102476 |
| 289 | DUSP16 | chr12:12626215-12715448 | PDT | ATRA+PDT | 21.2594 | 25.9134 | 0.285595 | 0.0035 | 0.0363197 |
| 290 | DUSP2 | chr2:96808907-96811179 | PDT | ATRA+PDT | 12.1886 | 26.538 | 1.12252 | 0.00005 | 0.00102476 |
| 291 | DUSP5 | chr10:112257624-112271302 | PDT | ATRA+PDT | 39.574 | 55.5091 | 0.488171 | 0.00005 | 0.00102476 |
| 292 | DVL1 | chr1:1270657-1284492 | PDT | ATRA+PDT | 31.0137 | 24.4283 | -0.344352 | 0.00085 | 0.0119898 |
| 293 | DYRK2 | chr12:68042511-68056444 | PDT | ATRA+PDT | 11.899 | 8.84758 | -0.427486 | 0.00005 | 0.00102476 |
| 294 | E2F7 | chr12:77415025-77459360 | PDT | ATRA+PDT | 13.4159 | 17.4383 | 0.378318 | 0.0001 | 0.00194145 |
| 295 | ECE2 | chr3:183967444-184010819 | PDT | ATRA+PDT | 17.6741 | 10.8401 | -0.705252 | 0.00005 | 0.00102476 |
| 296 | ECM1 | chr1:150480486-150486265 | PDT | ATRA+PDT | 2.36948 | 1.44101 | -0.717492 | 0.0014 | 0.0175049 |
| 297 | EDN1 | chr6:12290528-12297427 | PDT | ATRA+PDT | 14.2647 | 22.7092 | 0.670833 | 0.00005 | 0.00102476 |
| 298 | EEF1A2 | chr20:62119365-62130505 | PDT | ATRA+PDT | 29.2765 | 20.1294 | -0.540436 | 0.00005 | 0.00102476 |
| 299 | EEPD1 | chr7:36192835-36341152 | PDT | ATRA+PDT | 6.13859 | 7.7133 | 0.32944 | 0.00375 | 0.0383182 |
| 300 | EFCAB4A | chr11:827584-831991 | PDT | ATRA+PDT | 11.8512 | 8.80399 | -0.428809 | 0.0006 | 0.00910057 |
| 301 | EFNA1 | chr1:155100348-155107386 | PDT | ATRA+PDT | 23.3266 | 18.0179 | -0.372544 | 0.0014 | 0.0175049 |
| 302 | EGLN1 | chr1:231499496-231560790 | PDT | ATRA+PDT | 4.00706 | 5.21901 | 0.381232 | 0.001 | 0.0135165 |
| 303 | EGR1 | chr5:137801180-137805004 | PDT | ATRA+PDT | 4.82157 | 9.81097 | 1.02489 | 0.00005 | 0.00102476 |
| 304 | EHF | chr11:34642587-34684834 | PDT | ATRA+PDT | 92.6037 | 123.486 | 0.415203 | 0.00005 | 0.00102476 |
| 305 | EIF2AK2 | chr2:37332283-37384190 | PDT | ATRA+PDT | 10.5599 | 13.6644 | 0.37183 | 0.0003 | 0.00511725 |
| 306 | EIF2C4 | chr1:36273827-36323490 | PDT | ATRA+PDT | 3.13563 | 4.12885 | 0.396986 | 0.00045 | 0.00719326 |

**Table S1: Significant differentially expressed genes (DEGs) after ATRA+PDT compared to PDT.**

| # | gene | locus | sample_1 | sample_2 | value_1 | value_2 | log2(fold_change) | p_value | q_value |
|---|---|---|---|---|---|---|---|---|---|
| 307 | EIF4A2 | chr3:186501360-186524484 | PDT | ATRA+PDT | 163.926 | 206.699 | 0.334491 | 0.00135 | 0.0171 |
| 308 | EIF5A2 | chr3:170606203-170626426 | PDT | ATRA+PDT | 3.07978 | 2.33192 | -0.401311 | 0.00215 | 0.0243712 |
| 309 | ELAC2 | chr17:12692828-12921381 | PDT | ATRA+PDT | 18.7339 | 15.1593 | -0.305453 | 0.00335 | 0.0351043 |
| 310 | ELF3 | chr1:201979689-201986315 | PDT | ATRA+PDT | 37.099 | 66.1482 | 0.834321 | 0.00005 | 0.00102476 |
| 311 | ELOVL6 | chr4:110970228-111119820 | PDT | ATRA+PDT | 17.373 | 14.134 | -0.297673 | 0.0044 | 0.0431831 |
| 312 | EMP1 | chr12:13349601-13369708 | PDT | ATRA+PDT | 67.9701 | 92.6699 | 0.447201 | 0.00005 | 0.00102476 |
| 313 | ENDOD1 | chr11:94822973-94865815 | PDT | ATRA+PDT | 11.7833 | 7.78629 | -0.59773 | 0.00005 | 0.00102476 |
| 314 | ENG | chr9:130577290-130617047 | PDT | ATRA+PDT | 4.39529 | 2.73046 | -0.686818 | 0.00005 | 0.00102476 |
| 315 | ENTPD5 | chr14:74433180-74486026 | PDT | ATRA+PDT | 4.50015 | 3.24469 | -0.471895 | 0.00265 | 0.0286404 |
| 316 | EPB41L1 | chr20:34700347-34820721 | PDT | ATRA+PDT | 22.7706 | 34.5998 | 0.603593 | 0.00005 | 0.00102476 |
| 317 | EPB49 | chr8:21911080-21940036 | PDT | ATRA+PDT | 3.52815 | 2.54822 | -0.469422 | 0.0015 | 0.0184954 |
| 318 | EPC1 | chr10:32557858-32636113 | PDT | ATRA+PDT | 12.8604 | 17.367 | 0.433417 | 0.0001 | 0.00194145 |
| 319 | EPHA2 | chr1:16450831-16482582 | PDT | ATRA+PDT | 40.8386 | 66.5434 | 0.704364 | 0.00005 | 0.00102476 |
| 320 | EPHB6 | chr7:142552791-142568847 | PDT | ATRA+PDT | 7.61521 | 5.03569 | -0.596695 | 0.00005 | 0.00102476 |
| 321 | EPN3 | chr17:48610047-48621111 | PDT | ATRA+PDT | 2.3054 | 1.62994 | -0.500193 | 0.00135 | 0.0171 |
| 322 | ERAP2 | chr5:96211643-96255406 | PDT | ATRA+PDT | 11.845 | 23.0945 | 0.963269 | 0.00005 | 0.00102476 |
| 323 | ERCC6L | chrX:71401525-71483814 | PDT | ATRA+PDT | 8.34576 | 6.30818 | -0.403821 | 0.00425 | 0.0420979 |
| 324 | EREG | chr4:75230859-75254477 | PDT | ATRA+PDT | 109.653 | 161.325 | 0.557018 | 0.00005 | 0.00102476 |
| 325 | ERN1 | chr17:62120389-62207502 | PDT | ATRA+PDT | 4.52517 | 6.46555 | 0.514798 | 0.00005 | 0.00102476 |
| 326 | ERRFI1 | chr1:8071778-8086393 | PDT | ATRA+PDT | 345.906 | 470.603 | 0.444132 | 0.00005 | 0.00102476 |
| 327 | ERVMER34-1 | chr4:53609683-53617807 | PDT | ATRA+PDT | 3.02422 | 2.13684 | -0.501086 | 0.00135 | 0.0171 |
| 328 | ESAM | chr11:124623018-124632223 | PDT | ATRA+PDT | 29.0409 | 36.6252 | 0.334753 | 0.00175 | 0.0207397 |
| 329 | ETS1 | chr11:128328655-128457453 | PDT | ATRA+PDT | 14.9045 | 20.5587 | 0.464007 | 0.00005 | 0.00102476 |
| 330 | ETV1 | chr7:13930855-14031050 | PDT | ATRA+PDT | 0.749919 | 1.22532 | 0.70835 | 0.00005 | 0.00102476 |
| 331 | ETV6 | chr12:11802787-12048325 | PDT | ATRA+PDT | 7.4987 | 10.5491 | 0.492402 | 0.00005 | 0.00102476 |
| 332 | EXOC8 | chr1:231468481-231473578 | PDT | ATRA+PDT | 7.75358 | 6.02451 | -0.36402 | 0.00085 | 0.0119898 |
| 333 | EXPH5 | chr11:108376157-108464374 | PDT | ATRA+PDT | 6.8025 | 8.9817 | 0.400923 | 0.00005 | 0.00102476 |
| 334 | F2R | chr5:76011867-76031595 | PDT | ATRA+PDT | 5.86953 | 9.46569 | 0.689462 | 0.00005 | 0.00102476 |
| 335 | F2RL1 | chr5:76114832-76131140 | PDT | ATRA+PDT | 49.4055 | 65.413 | 0.404907 | 0.00005 | 0.00102476 |
| 336 | F5 | chr1:169481191-169555769 | PDT | ATRA+PDT | 10.6716 | 6.61326 | -0.690341 | 0.00005 | 0.00102476 |
| 337 | FAAH | chr1:46859938-46879520 | PDT | ATRA+PDT | 5.50656 | 7.94572 | 0.529027 | 0.00005 | 0.00102476 |
| 338 | FAM102B | chr1:109102970-109181949 | PDT | ATRA+PDT | 6.13952 | 8.4052 | 0.453155 | 0.0001 | 0.00194145 |
| 339 | FAM105A | chr5:14581890-14616287 | PDT | ATRA+PDT | 2.59782 | 1.64428 | -0.659843 | 0.00005 | 0.00102476 |
| 340 | FAM107B | chr10:14560558-14816896 | PDT | ATRA+PDT | 59.2022 | 73.3432 | 0.309013 | 0.0012 | 0.0156263 |
| 341 | FAM108C1 | chr15:80987651-81047962 | PDT | ATRA+PDT | 32.8262 | 26.161 | -0.327426 | 0.00155 | 0.0189586 |
| 342 | FAM111B | chr11:58874657-58894888 | PDT | ATRA+PDT | 31.7401 | 24.7476 | -0.359017 | 0.0006 | 0.00910057 |
| 343 | FAM123B | chrX:63404996-63425624 | PDT | ATRA+PDT | 1.69093 | 1.18022 | -0.518764 | 0.00005 | 0.00102476 |
| 344 | FAM127B | chrX:134184962-134186221 | PDT | ATRA+PDT | 16.058 | 21.8891 | 0.446926 | 0.004 | 0.0401429 |
| 345 | FAM131B | chr7:143050492-143059840 | PDT | ATRA+PDT | 0.409476 | 3.47658 | 3.08582 | 0.00005 | 0.00102476 |
| 346 | FAM135A | chr6:71123106-71270877 | PDT | ATRA+PDT | 18.3524 | 13.9109 | -0.399752 | 0.00015 | 0.00281494 |
| 347 | FAM13A | chr4:89630939-89978323 | PDT | ATRA+PDT | 4.12024 | 7.15203 | 0.795624 | 0.00005 | 0.00102476 |
| 348 | FAM149A | chr4:187065994-187093817 | PDT | ATRA+PDT | 8.48109 | 16.2294 | 0.936285 | 0.00005 | 0.00102476 |
| 349 | FAM149B1 | chr10:74927876-75001939 | PDT | ATRA+PDT | 3.59557 | 4.85356 | 0.432823 | 0.0011 | 0.0146092 |
| 350 | FAM162A | chr3:122103022-122128961 | PDT | ATRA+PDT | 82.1723 | 61.1637 | -0.425977 | 0.0001 | 0.00194145 |
| 351 | FAM174B | chr15:93160678-93199031 | PDT | ATRA+PDT | 7.42036 | 5.03265 | -0.560172 | 0.00005 | 0.00102476 |
| 352 | FAM198B | chr4:159045731-159094202 | PDT | ATRA+PDT | 12.9282 | 7.67835 | -0.751657 | 0.00005 | 0.00102476 |
| 353 | FAM210A | chr18:13663345-13726591 | PDT | ATRA+PDT | 15.116 | 11.8696 | -0.3488 | 0.0012 | 0.0156263 |
| 354 | FAM3D | chr3:58619669-58652561 | PDT | ATRA+PDT | 8.77298 | 4.24277 | -1.04806 | 0.00005 | 0.00102476 |
| 355 | FAM63B | chr15:59063392-59149734 | PDT | ATRA+PDT | 2.30668 | 3.07012 | 0.412473 | 0.00105 | 0.0140676 |
| 356 | FAM81A | chr15:59730371-59815751 | PDT | ATRA+PDT | 5.00287 | 3.87761 | -0.367588 | 0.0046 | 0.0446535 |
| 357 | FAM84B | chr8:127564682-127570711 | PDT | ATRA+PDT | 127.905 | 156.797 | 0.293824 | 0.00435 | 0.0428104 |
| 358 | FAM86EP | chr4:3943668-3957148 | PDT | ATRA+PDT | 1.55741 | 2.30187 | 0.563655 | 0.00065 | 0.00978942 |
| 359 | FAM98A | chr2:33808728-33824362 | PDT | ATRA+PDT | 27.1483 | 20.8592 | -0.380176 | 0.0003 | 0.00511725 |
| 360 | FANCF | chr11:22644078-22647387 | PDT | ATRA+PDT | 4.17794 | 5.47064 | 0.388919 | 0.0024 | 0.0263926 |
| 361 | FAS | chr10:90694830-90775542 | PDT | ATRA+PDT | 2.10634 | 1.40791 | -0.581187 | 0.0034 | 0.0355585 |
| 362 | FBXL19-AS1 | chr16:30930639-30934590 | PDT | ATRA+PDT | 1.41865 | 2.01877 | 0.508954 | 0.0021 | 0.0238805 |
| 363 | FBXO32 | chr8:124510126-124553493 | PDT | ATRA+PDT | 4.9402 | 6.10708 | 0.305914 | 0.00455 | 0.0443293 |
| 364 | FEM1C | chr5:114856607-114880591 | PDT | ATRA+PDT | 30.0869 | 38.8026 | 0.36702 | 0.0002 | 0.00358322 |
| 365 | FER | chr5:108083522-108523373 | PDT | ATRA+PDT | 2.62255 | 3.97463 | 0.59985 | 0.00005 | 0.00102476 |
| 366 | FEZ2 | chr2:36779403-36825332 | PDT | ATRA+PDT | 21.8097 | 26.8895 | 0.302071 | 0.0049 | 0.0470101 |
| 367 | FGD3 | chr9:95709600-95798518 | PDT | ATRA+PDT | 1.54982 | 2.16782 | 0.484138 | 0.00495 | 0.0474046 |
| 368 | FGF19 | chr11:69513005-69519106 | PDT | ATRA+PDT | 6.27328 | 25.0306 | 1.9964 | 0.00005 | 0.00102476 |
| 369 | FGFR2 | chr10:123237843-123357972 | PDT | ATRA+PDT | 8.45316 | 13.0471 | 0.626161 | 0.00005 | 0.00102476 |
| 370 | FHL2 | chr2:105977282-106055230 | PDT | ATRA+PDT | 60.6957 | 86.9494 | 0.518583 | 0.00005 | 0.00102476 |
| 371 | FHL3 | chr1:38462441-38471187 | PDT | ATRA+PDT | 12.2879 | 16.2252 | 0.400995 | 0.00185 | 0.0217319 |
| 372 | FIGNL1 | chr7:50511831-50518088 | PDT | ATRA+PDT | 13.0673 | 10.5123 | -0.313882 | 0.00345 | 0.0359757 |
| 373 | FLNB | chr3:57994126-58157982 | PDT | ATRA+PDT | 89.7689 | 121.2 | 0.433102 | 0.00005 | 0.00102476 |
| 374 | FLOT1 | chr6_ssto_hap7:2027817-204277 | PDT | ATRA+PDT | 49.3978 | 62.6809 | 0.343578 | 0.001 | 0.0135165 |
| 375 | FLRT3 | chr20:13976145-16033841 | PDT | ATRA+PDT | 1.48851 | 2.13722 | 0.521865 | 0.0014 | 0.0175049 |
| 376 | FOS | chr14:75745480-75748937 | PDT | ATRA+PDT | 9.11561 | 13.3146 | 0.546601 | 0.00005 | 0.00102476 |
| 377 | FOSB | chr19:45971252-45978437 | PDT | ATRA+PDT | 2.84898 | 5.04937 | 0.825659 | 0.00005 | 0.00102476 |
| 378 | FOSL1 | chr11:65659691-65667997 | PDT | ATRA+PDT | 152.947 | 219.618 | 0.521971 | 0.00005 | 0.00102476 |
| 379 | FOXL1 | chr16:86612114-86615304 | PDT | ATRA+PDT | 1.38773 | 2.05807 | 0.568566 | 0.0021 | 0.0238805 |
| 380 | FOXN2 | chr2:48541794-48606434 | PDT | ATRA+PDT | 13.7006 | 17.568 | 0.358703 | 0.00015 | 0.00281494 |
| 381 | FRMD4B | chr3:69219145-69435430 | PDT | ATRA+PDT | 2.05717 | 3.05091 | 0.568578 | 0.00005 | 0.00102476 |
| 382 | FRRS1 | chr1:100111430-100231349 | PDT | ATRA+PDT | 9.3853 | 14.3505 | 0.612622 | 0.00005 | 0.00102476 |
| 383 | FRYL | chr4:48499379-48782316 | PDT | ATRA+PDT | 8.78851 | 10.6727 | 0.280238 | 0.00455 | 0.0443293 |
| 384 | FSTL3 | chr19:676388-683392 | PDT | ATRA+PDT | 1.03427 | 1.77411 | 0.778476 | 0.00085 | 0.0119898 |
| 385 | FTSJD1 | chr16:71316202-71323509 | PDT | ATRA+PDT | 37.7123 | 52.1353 | 0.467224 | 0.00005 | 0.00102476 |
| 386 | FUT4 | chr11:94277016-94283064 | PDT | ATRA+PDT | 18.9388 | 15.3605 | -0.302118 | 0.0023 | 0.0255827 |
| 387 | FUT8 | chr14:65877309-66210839 | PDT | ATRA+PDT | 1.64239 | 1.10566 | -0.570888 | 0.0011 | 0.0146092 |
| 388 | FXYD3 | chr19:35606731-35615228 | PDT | ATRA+PDT | 17.5797 | 8.62688 | -1.027 | 0.00005 | 0.00102476 |
| 389 | FYN | chr6:111981534-112194655 | PDT | ATRA+PDT | 5.51577 | 3.87692 | -0.50865 | 0.0001 | 0.00194145 |
| 390 | GAB1 | chr4:144257982-144395718 | PDT | ATRA+PDT | 6.95646 | 8.89793 | 0.355116 | 0.00095 | 0.0131061 |
| 391 | GAB2 | chr11:77926335-78128868 | PDT | ATRA+PDT | 1.8574 | 2.53823 | 0.45054 | 0.00075 | 0.0109256 |
| 392 | GABRP | chr5:170210722-170241050 | PDT | ATRA+PDT | 1.85944 | 4.52118 | 1.28183 | 0.00005 | 0.00102476 |
| 393 | GADD45A | chr1:68150859-68154021 | PDT | ATRA+PDT | 32.4531 | 53.294 | 0.715616 | 0.00005 | 0.00102476 |
| 394 | GADD45B | chr19:2476122-2478257 | PDT | ATRA+PDT | 3.93333 | 7.29827 | 0.891804 | 0.00005 | 0.00102476 |
| 395 | GALM | chr2:38893051-38961909 | PDT | ATRA+PDT | 7.12751 | 9.34785 | 0.391235 | 0.00205 | 0.0234618 |
| 396 | GALNT6 | chr12:51745832-51785200 | PDT | ATRA+PDT | 1.32323 | 1.87155 | 0.500173 | 0.002 | 0.0229388 |
| 397 | GATA2 | chr3:128198264-128212030 | PDT | ATRA+PDT | 2.09529 | 3.76078 | 0.843884 | 0.00005 | 0.00102476 |
| 398 | GATA6 | chr18:19749415-19782227 | PDT | ATRA+PDT | 12.7737 | 16.4298 | 0.363143 | 0.00055 | 0.0085238 |
| 399 | GBP3 | chr1:89472359-89488549 | PDT | ATRA+PDT | 5.81741 | 10.962 | 0.914066 | 0.00005 | 0.00102476 |
| 400 | GCNT2 | chr6:10521567-10629601 | PDT | ATRA+PDT | 4.33216 | 5.83905 | 0.430648 | 0.00035 | 0.00584867 |
| 401 | GDF15 | chr19:18496967-18499986 | PDT | ATRA+PDT | 395.98 | 673.908 | 0.767123 | 0.00005 | 0.00102476 |
| 402 | GFOD1 | chr6:13363586-13487869 | PDT | ATRA+PDT | 4.02601 | 2.65386 | -0.601256 | 0.00035 | 0.00584867 |
| 403 | GJB2 | chr13:20761603-20767114 | PDT | ATRA+PDT | 13.3763 | 10.3672 | -0.367655 | 0.0019 | 0.0221245 |
| 404 | GLB1L2 | chr11:134201767-134246218 | PDT | ATRA+PDT | 40.5745 | 33.0921 | -0.294089 | 0.0033 | 0.0346484 |
| 405 | GLCCI1 | chr7:8008373-8128709 | PDT | ATRA+PDT | 5.11203 | 7.22144 | 0.498392 | 0.00005 | 0.00102476 |
| 406 | GLI1 | chr12:57853917-57873633 | PDT | ATRA+PDT | 1.47099 | 0.87076 | -0.756439 | 0.00085 | 0.0119898 |
| 407 | GLI2 | chr2:121554866-121750229 | PDT | ATRA+PDT | 1.73488 | 0.852615 | -1.02487 | 0.00005 | 0.00102476 |
| 408 | GLIS2 | chr16:4382224-4389598 | PDT | ATRA+PDT | 6.53027 | 8.92753 | 0.451118 | 0.0003 | 0.00511725 |

**Table S1: Significant differentially expressed genes (DEGs) after ATRA+PDT compared to PDT.**

| # | gene | locus | sample_1 | sample_2 | value_1 | value_2 | log2(fold_change) | p_value | q_value |
|---|---|---|---|---|---|---|---|---|---|
| 409 | GLIS3 | chr9:3824127-4300035 | PDT | ATRA+PDT | 2.9426 | 4.17559 | 0.50489 | 0.00005 | 0.00102476 |
| 410 | GLTPD1 | chr1:1260142-1264276 | PDT | ATRA+PDT | 6.41022 | 4.68666 | -0.451815 | 0.0008 | 0.0114972 |
| 411 | GMEB2 | chr20:62218954-62258381 | PDT | ATRA+PDT | 17.8984 | 22.2288 | 0.3126 | 0.00245 | 0.0267496 |
| 412 | GNA14 | chr9:80037994-80263232 | PDT | ATRA+PDT | 1.83269 | 1.07864 | -0.764748 | 0.0013 | 0.0166643 |
| 413 | GNAI1 | chr7:79764139-79848725 | PDT | ATRA+PDT | 18.3225 | 14.5946 | -0.328176 | 0.00165 | 0.0198632 |
| 414 | GNAZ | chr22:23401592-23484241 | PDT | ATRA+PDT | 1.1314 | 0.568899 | -0.991869 | 0.00085 | 0.0119898 |
| 415 | GNE | chr9:36214438-36277053 | PDT | ATRA+PDT | 7.61007 | 5.82517 | -0.385611 | 0.0004 | 0.00650107 |
| 416 | GNPAT | chr1:231359508-231413719 | PDT | ATRA+PDT | 22.1603 | 31.3272 | 0.499438 | 0.00005 | 0.00102476 |
| 417 | GOLGA8A | chr15:34671269-34729667 | PDT | ATRA+PDT | 5.68254 | 4.37806 | -0.376245 | 0.00225 | 0.0252369 |
| 418 | GOLPH3L | chr1:150618700-150669672 | PDT | ATRA+PDT | 19.8668 | 28.0126 | 0.495718 | 0.00005 | 0.00102476 |
| 419 | GOLT1A | chr1:204167287-204183220 | PDT | ATRA+PDT | 7.49469 | 11.4978 | 0.61742 | 0.0003 | 0.00511725 |
| 420 | GPC4 | chrX:132435063-132549205 | PDT | ATRA+PDT | 2.61393 | 3.85772 | 0.561528 | 0.00005 | 0.00102476 |
| 421 | GPCPD1 | chr20:5525079-5591672 | PDT | ATRA+PDT | 2.764 | 3.95326 | 0.516285 | 0.00005 | 0.00102476 |
| 422 | GPD1 | chr12:50497800-50505095 | PDT | ATRA+PDT | 2.28955 | 1.01978 | -1.1668 | 0.00005 | 0.00102476 |
| 423 | GPR110 | chr6:46967812-47010082 | PDT | ATRA+PDT | 31.3419 | 51.5418 | 0.717651 | 0.00005 | 0.00102476 |
| 424 | GPR114 | chr16:57576600-57611100 | PDT | ATRA+PDT | 3.23369 | 2.00971 | -0.686193 | 0.00005 | 0.00102476 |
| 425 | GPR158 | chr10:25447000-25891157 | PDT | ATRA+PDT | 0.87409 | 1.31409 | 0.588213 | 0.00025 | 0.00439786 |
| 426 | GPR160 | chr3:169755734-169803183 | PDT | ATRA+PDT | 7.7991 | 10.3693 | 0.410936 | 0.0013 | 0.0166643 |
| 427 | GPR35 | chr2:241544824-241570676 | PDT | ATRA+PDT | 25.9273 | 20.6714 | -0.326836 | 0.002 | 0.0229388 |
| 428 | GPR87 | chr3:150804675-151176497 | PDT | ATRA+PDT | 4.55177 | 8.63195 | 0.92326 | 0.00005 | 0.00102476 |
| 429 | GPRC5A | chr12:13043955-13066600 | PDT | ATRA+PDT | 195.824 | 291.91 | 0.575964 | 0.00005 | 0.00102476 |
| 430 | GRB7 | chr17:37894161-37903538 | PDT | ATRA+PDT | 26.9977 | 40.8461 | 0.597359 | 0.00005 | 0.00102476 |
| 431 | GRHL3 | chr1:24645811-24741587 | PDT | ATRA+PDT | 1.75026 | 0.776258 | -1.17296 | 0.002 | 0.0229388 |
| 432 | GRIN2D | chr19:48898131-48948188 | PDT | ATRA+PDT | 3.05194 | 5.32452 | 0.802924 | 0.00005 | 0.00102476 |
| 433 | GRK5 | chr10:120967196-121215131 | PDT | ATRA+PDT | 5.1872 | 3.94614 | -0.394514 | 0.00235 | 0.0259765 |
| 434 | GSDMB | chr17:38060847-38074903 | PDT | ATRA+PDT | 2.28629 | 3.67046 | 0.682951 | 0.00165 | 0.0198632 |
| 435 | GSDMD | chr8:144635556-144645231 | PDT | ATRA+PDT | 7.49063 | 11.6451 | 0.636561 | 0.00005 | 0.00102476 |
| 436 | GSN | chr9:124030379-124095120 | PDT | ATRA+PDT | 41.6084 | 30.6138 | -0.442694 | 0.00005 | 0.00102476 |
| 437 | GSTA4 | chr6:52842745-52860178 | PDT | ATRA+PDT | 7.19953 | 4.1013 | -0.811822 | 0.00005 | 0.00102476 |
| 438 | GTPBP2 | chr6:43543877-43596936 | PDT | ATRA+PDT | 48.7001 | 66.9444 | 0.459038 | 0.00005 | 0.00102476 |
| 439 | GUK1 | chr1:228327784-228336655 | PDT | ATRA+PDT | 92.4045 | 73.8656 | -0.32306 | 0.00165 | 0.0198632 |
| 440 | GYG1 | chr3:148709194-148745456 | PDT | ATRA+PDT | 27.1253 | 21.9331 | -0.306532 | 0.0038 | 0.0386074 |
| 441 | H1F0 | chr22:38201113-38203443 | PDT | ATRA+PDT | 89.3849 | 124.932 | 0.483036 | 0.00005 | 0.00102476 |
| 442 | H1FX | chr3:129033613-129043412 | PDT | ATRA+PDT | 45.3794 | 59.1282 | 0.38181 | 0.00025 | 0.00439786 |
| 443 | HAGHL | chr16:777265-779715 | PDT | ATRA+PDT | 4.11839 | 2.28962 | -0.846973 | 0.00045 | 0.00719326 |
| 444 | HBEGF | chr5:139712427-139726188 | PDT | ATRA+PDT | 20.527 | 41.5863 | 1.01859 | 0.00005 | 0.00102476 |
| 445 | HCAR1 | chr12:123212152-123215129 | PDT | ATRA+PDT | 5.34507 | 7.07409 | 0.404334 | 0.0014 | 0.0175049 |
| 446 | HDAC9 | chr7:18126571-19036992 | PDT | ATRA+PDT | 1.44155 | 3.20797 | 1.15404 | 0.00005 | 0.00102476 |
| 447 | HEPH | chrX:65382432-65487230 | PDT | ATRA+PDT | 3.62214 | 1.79148 | -1.01569 | 0.00005 | 0.00102476 |
| 448 | HERC6 | chr4:89299890-89364249 | PDT | ATRA+PDT | 0.683215 | 1.25848 | 0.881267 | 0.00015 | 0.00281494 |
| 449 | HES6 | chr2:239146907-239148681 | PDT | ATRA+PDT | 5.73401 | 2.55042 | -1.16881 | 0.00005 | 0.00102476 |
| 450 | HGD | chr3:120347014-120401418 | PDT | ATRA+PDT | 10.2884 | 5.52027 | -0.898213 | 0.00005 | 0.00102476 |
| 451 | HHLA2 | chr3:108021331-108097126 | PDT | ATRA+PDT | 4.62643 | 1.14286 | 0.36638 | 0.00005 | 0.00102476 |
| 452 | HIPK2 | chr7:139246315-139477693 | PDT | ATRA+PDT | 5.89243 | 7.59602 | 0.36638 | 0.0003 | 0.00511725 |
| 453 | HIST1H2BN | chr6:27806439-27806888 | PDT | ATRA+PDT | 1.30759 | 0 | -inf | 0.00145 | 0.0179618 |
| 454 | HIVEP2 | chr6:143072603-143266338 | PDT | ATRA+PDT | 6.94549 | 9.85117 | 0.504219 | 0.00005 | 0.00102476 |
| 455 | HJURP | chr2:234745485-234763212 | PDT | ATRA+PDT | 19.6228 | 15.0074 | -0.386854 | 0.00025 | 0.00439786 |
| 456 | HK1 | chr10:71029755-71161637 | PDT | ATRA+PDT | 39.6626 | 31.8751 | -0.315349 | 0.001 | 0.0135165 |
| 457 | HLA-B | chr6_ssto_hap7:2655407-265874 | PDT | ATRA+PDT | 2.29347 | 4.53191 | 0.98259 | 0.00005 | 0.00102476 |
| 458 | HLA-C | chr6_ssto_hap7:2570004-257338 | PDT | ATRA+PDT | 28.2747 | 48.785 | 0.786925 | 0.00005 | 0.00102476 |
| 459 | HLA-E | chr6_ssto_hap7:1789472-179427 | PDT | ATRA+PDT | 30.4013 | 38.3495 | 0.335076 | 0.00095 | 0.0131061 |
| 460 | HLA-J | chr6_ssto_hap7:1298114-135940 | PDT | ATRA+PDT | 1.6197 | 2.77771 | 0.778169 | 0.00445 | 0.0435537 |
| 461 | HMGA2 | chr12:66151800-66360071 | PDT | ATRA+PDT | 6.8733 | 4.23936 | -0.697159 | 0.00005 | 0.00102476 |
| 462 | HMGCR | chr5:74632992-74657926 | PDT | ATRA+PDT | 86.0972 | 66.7796 | -0.366559 | 0.00035 | 0.00584867 |
| 463 | HMGCS2 | chr1:120290618-120311555 | PDT | ATRA+PDT | 16.1869 | 0.803481 | -4.33242 | 0.00005 | 0.00102476 |
| 464 | HNF4A | chr20:42984440-43060030 | PDT | ATRA+PDT | 16.8354 | 9.53061 | -0.820854 | 0.00005 | 0.00102476 |
| 465 | HNF4G | chr8:76452202-76479061 | PDT | ATRA+PDT | 13.4329 | 19.4032 | 0.530523 | 0.00005 | 0.00102476 |
| 466 | HOXA3 | chr7:27145808-27166639 | PDT | ATRA+PDT | 2.22132 | 3.42141 | 0.623174 | 0.00005 | 0.00102476 |
| 467 | HOXA5 | chr7:27179982-27196296 | PDT | ATRA+PDT | 9.4125 | 12.6717 | 0.428962 | 0.00405 | 0.0405304 |
| 468 | HPDL | chr1:45792544-45794346 | PDT | ATRA+PDT | 7.45459 | 5.14936 | -0.533734 | 0.00005 | 0.00102476 |
| 469 | HPGD | chr4:175411327-175444049 | PDT | ATRA+PDT | 10.8983 | 5.70085 | -0.934858 | 0.00005 | 0.00102476 |
| 470 | HRH1 | chr3:11178778-11304939 | PDT | ATRA+PDT | 5.2823 | 8.67514 | 0.71572 | 0.00005 | 0.00102476 |
| 471 | HSPA1A | chr6_qbl_hap6:3076937-307936 | PDT | ATRA+PDT | 612.758 | 897.988 | 0.551379 | 0.00005 | 0.00102476 |
| 472 | HSPA1B | chr6_qbl_hap6:3089162-309168 | PDT | ATRA+PDT | 745.494 | 1083.52 | 0.539457 | 0.00005 | 0.00102476 |
| 473 | HSPA1L | chr6_qbl_hap6:3071043-307648 | PDT | ATRA+PDT | 2.68991 | 3.82212 | 0.506814 | 0.00095 | 0.0131061 |
| 474 | HSPA4L | chr4:128703452-128754526 | PDT | ATRA+PDT | 45.3591 | 59.7923 | 0.398568 | 0.00005 | 0.00102476 |
| 475 | HSPA6 | chr1:161494035-161496687 | PDT | ATRA+PDT | 159.467 | 281.821 | 0.821519 | 0.00005 | 0.00102476 |
| 476 | HSPA7 | chr1:161575848-161578341 | PDT | ATRA+PDT | 3.44082 | 6.33443 | 0.880463 | 0.00005 | 0.00102476 |
| 477 | HSPB1 | chr7:75931874-75933614 | PDT | ATRA+PDT | 102.047 | 199.387 | 0.96634 | 0.00005 | 0.00102476 |
| 478 | HSPG2 | chr1:22138757-22263750 | PDT | ATRA+PDT | 13.7059 | 21.6875 | 0.66207 | 0.00005 | 0.00102476 |
| 479 | HSPH1 | chr13:31710762-31736117 | PDT | ATRA+PDT | 462.619 | 655.509 | 0.50279 | 0.00005 | 0.00102476 |
| 480 | HTR1D | chr1:23518387-23521222 | PDT | ATRA+PDT | 2.95125 | 1.97948 | -0.576204 | 0.0004 | 0.00650107 |
| 481 | ID2 | chr2:8822112-8824583 | PDT | ATRA+PDT | 5.50856 | 2.75203 | -1.00118 | 0.00005 | 0.00102476 |
| 482 | IDI1 | chr10:1064846-1095061 | PDT | ATRA+PDT | 129.123 | 102.959 | -0.326683 | 0.00055 | 0.0085238 |
| 483 | IER5L | chr9:131937830-131940540 | PDT | ATRA+PDT | 1.89291 | 2.85555 | 0.593163 | 0.0008 | 0.0114972 |
| 484 | IFFO2 | chr1:19230773-19282826 | PDT | ATRA+PDT | 55.4573 | 77.6592 | 0.485779 | 0.00005 | 0.00102476 |
| 485 | IFI35 | chr17:41158741-41166476 | PDT | ATRA+PDT | 3.36478 | 6.42601 | 0.933414 | 0.00005 | 0.00102476 |
| 486 | IFI44 | chr1:79115476-79129763 | PDT | ATRA+PDT | 1.76976 | 3.04735 | 0.783999 | 0.00055 | 0.0085238 |
| 487 | IFIH1 | chr2:163123588-163175039 | PDT | ATRA+PDT | 1.31458 | 1.92628 | 0.551214 | 0.0023 | 0.0255827 |
| 488 | IFIT3 | chr10:91087601-91100725 | PDT | ATRA+PDT | 3.12177 | 4.27384 | 0.453168 | 0.00175 | 0.0207397 |
| 489 | IFITM2 | chr11:308106-309410 | PDT | ATRA+PDT | 16.2241 | 28.8464 | 0.830253 | 0.00005 | 0.00102476 |
| 490 | IFITM3 | chr11:319672-320914 | PDT | ATRA+PDT | 3.17628 | 9.52404 | 1.58423 | 0.00005 | 0.00102476 |
| 491 | IFNGR2 | chr21:34775201-34852316 | PDT | ATRA+PDT | 14.2102 | 18.7851 | 0.402665 | 0.00165 | 0.0198632 |
| 492 | IGF2 | chr11:2150341-2182439 | PDT | ATRA+PDT | 6.58505 | 4.4414 | -0.568179 | 0.0004 | 0.00650107 |
| 493 | IGFBP2 | chr2:217498126-217529158 | PDT | ATRA+PDT | 9.34685 | 6.63453 | -0.494486 | 0.0005 | 0.00788626 |
| 494 | IGFBP6 | chr12:53491435-53496128 | PDT | ATRA+PDT | 31.6079 | 110.977 | 1.81191 | 0.00005 | 0.00102476 |
| 495 | IGSF1 | chrX:130407482-130423403 | PDT | ATRA+PDT | 3.04443 | 1.18424 | -1.36221 | 0.00005 | 0.00102476 |
| 496 | IGSF9 | chr1:159896828-159915386 | PDT | ATRA+PDT | 3.79281 | 2.55118 | -0.572102 | 0.00005 | 0.00102476 |
| 497 | IL15RA | chr10:5994333-6020150 | PDT | ATRA+PDT | 1.29552 | 3.53252 | 1.44716 | 0.00005 | 0.00102476 |
| 498 | IL1R2 | chr2:102608305-102644884 | PDT | ATRA+PDT | 6.14538 | 4.29709 | -0.516143 | 0.0019 | 0.0221245 |
| 499 | IL1RN | chr2:113875469-113891593 | PDT | ATRA+PDT | 13.7209 | 19.7421 | 0.524894 | 0.00005 | 0.00102476 |
| 500 | IL8 | chr4:74606222-74609433 | PDT | ATRA+PDT | 14.3248 | 29.5956 | 1.04686 | 0.00005 | 0.00102476 |
| 501 | IMPDH1 | chr7:128032330-128050036 | PDT | ATRA+PDT | 58.24 | 47.5302 | -0.293165 | 0.0032 | 0.0337978 |
| 502 | INCENP | chr11:61891444-61920635 | PDT | ATRA+PDT | 17.7265 | 14.0139 | -0.339053 | 0.00125 | 0.0161202 |
| 503 | INPPL1 | chr11:71935881-71955220 | PDT | ATRA+PDT | 7.33072 | 9.52766 | 0.378168 | 0.0006 | 0.00910057 |
| 504 | INSIG1 | chr7:155089485-155101945 | PDT | ATRA+PDT | 92.3001 | 69.1934 | -0.415699 | 0.00005 | 0.00102476 |
| 505 | INTU | chr4:128554086-128637934 | PDT | ATRA+PDT | 1.52329 | 2.25255 | 0.564362 | 0.00135 | 0.0171 |
| 506 | IQGAP2 | chr5:75699148-76003957 | PDT | ATRA+PDT | 6.99109 | 5.53759 | -0.336261 | 0.00185 | 0.0217319 |
| 507 | IRF1 | chr5:131817300-131826465 | PDT | ATRA+PDT | 6.32604 | 9.99935 | 0.660531 | 0.00005 | 0.00102476 |
| 508 | IRF7 | chr11:612554-615999 | PDT | ATRA+PDT | 9.12755 | 11.8924 | 0.381736 | 0.00245 | 0.0267496 |
| 509 | IRS2 | chr13:110406183-110438914 | PDT | ATRA+PDT | 3.94161 | 5.33563 | 0.436872 | 0.00005 | 0.00102476 |
| 510 | ITGB6 | chr2:160958232-161056590 | PDT | ATRA+PDT | 9.28624 | 14.4733 | 0.640229 | 0.00005 | 0.00102476 |

**Table S1: Significant differentially expressed genes (DEGs) after ATRA+PDT compared to PDT.**

| # | gene | locus | sample_1 | sample_2 | value_1 | value_2 | log2(fold_change) | p_value | q_value |
|---|---|---|---|---|---|---|---|---|---|
| 511 | ITM2B | chr13:48807273-48836232 | PDT | ATRA+PDT | 45.3908 | 56.6793 | 0.320422 | 0.0016 | 0.0194588 |
| 512 | ITPKA | chr15:41786055-41795757 | PDT | ATRA+PDT | 0.556708 | 1.31432 | 1.23932 | 0.00425 | 0.0420979 |
| 513 | ITPR2 | chr12:26488284-26986131 | PDT | ATRA+PDT | 2.36664 | 1.77543 | -0.414676 | 0.00045 | 0.00719326 |
| 514 | ITPRIP | chr10:106071898-106093663 | PDT | ATRA+PDT | 3.14414 | 4.07388 | 0.373739 | 0.004 | 0.0401429 |
| 515 | JAG2 | chr14:105608075-105635161 | PDT | ATRA+PDT | 3.13358 | 2.14484 | -0.546942 | 0.00005 | 0.00102476 |
| 516 | JDP2 | chr14:75894508-75939404 | PDT | ATRA+PDT | 2.75811 | 4.48053 | 0.699991 | 0.00005 | 0.00102476 |
| 517 | JMJD1C | chr10:64926987-65226322 | PDT | ATRA+PDT | 28.1971 | 38.0703 | 0.433119 | 0.00005 | 0.00102476 |
| 518 | JMJD5 | chr16:27214806-27233089 | PDT | ATRA+PDT | 1.04725 | 1.86261 | 0.830721 | 0.0008 | 0.0114972 |
| 519 | JUN | chr1:59246462-59249785 | PDT | ATRA+PDT | 62.6204 | 87.3252 | 0.479765 | 0.00005 | 0.00102476 |
| 520 | JUND | chr19:18390562-18392432 | PDT | ATRA+PDT | 50.5425 | 72.2088 | 0.514676 | 0.00005 | 0.00102476 |
| 521 | KANK1 | chr9:504702-746103 | PDT | ATRA+PDT | 4.76117 | 6.11556 | 0.361169 | 0.00265 | 0.0286404 |
| 522 | KANSL3 | chr2:97258906-97304115 | PDT | ATRA+PDT | 17.7835 | 22.5732 | 0.344067 | 0.0005 | 0.00788626 |
| 523 | KCNE3 | chr11:74165885-74178600 | PDT | ATRA+PDT | 3.59091 | 7.78606 | 1.11655 | 0.00005 | 0.00102476 |
| 524 | KCNK5 | chr6:39156746-39197251 | PDT | ATRA+PDT | 10.4488 | 7.78294 | -0.424949 | 0.0002 | 0.00358322 |
| 525 | KDM2B | chr12:121866899-122018920 | PDT | ATRA+PDT | 6.14892 | 4.89176 | -0.329979 | 0.00325 | 0.0342581 |
| 526 | KDM6B | chr17:7743234-7758118 | PDT | ATRA+PDT | 8.33288 | 10.5209 | 0.336372 | 0.00095 | 0.0131061 |
| 527 | KIAA0040 | chr1:175126122-175162229 | PDT | ATRA+PDT | 9.05332 | 11.1212 | 0.296789 | 0.00515 | 0.0488381 |
| 528 | KIAA0182 | chr16:85645028-85709812 | PDT | ATRA+PDT | 25.1433 | 31.1985 | 0.311303 | 0.0018 | 0.0212615 |
| 529 | KIAA0430 | chr16:15688225-15737023 | PDT | ATRA+PDT | 6.13795 | 7.74811 | 0.336087 | 0.001 | 0.0135165 |
| 530 | KIAA0664L3 | chr16:31711933-31718745 | PDT | ATRA+PDT | 1.59993 | 2.86062 | 0.838318 | 0.0007 | 0.0103814 |
| 531 | KIAA0907 | chr1:155882835-155904188 | PDT | ATRA+PDT | 14.384 | 10.9516 | -0.393324 | 0.00055 | 0.0085238 |
| 532 | KIAA1109 | chr4:123091757-123283914 | PDT | ATRA+PDT | 6.77937 | 8.55588 | 0.335765 | 0.00075 | 0.0109256 |
| 533 | KIAA1161 | chr9:34368906-34376894 | PDT | ATRA+PDT | 13.347 | 9.93011 | -0.426637 | 0.00005 | 0.00102476 |
| 534 | KIAA1199 | chr15:81071711-81243999 | PDT | ATRA+PDT | 18.2246 | 26.0793 | 0.517021 | 0.00005 | 0.00102476 |
| 535 | KIAA1211 | chr4:57036360-57196890 | PDT | ATRA+PDT | 2.71753 | 1.96603 | -0.467014 | 0.00025 | 0.00439786 |
| 536 | KIAA1244 | chr6:138483052-138665800 | PDT | ATRA+PDT | 4.25439 | 5.27576 | 0.310426 | 0.0027 | 0.0291218 |
| 537 | KIAA1462 | chr10:30301728-30348488 | PDT | ATRA+PDT | 2.75442 | 2.14844 | -0.358459 | 0.0027 | 0.0291218 |
| 538 | KIF13B | chr8:28924794-29120610 | PDT | ATRA+PDT | 3.97397 | 4.9588 | 0.319411 | 0.0035 | 0.0363197 |
| 539 | KITLG | chr12:88886569-88974250 | PDT | ATRA+PDT | 62.7278 | 48.1623 | -0.381202 | 0.0002 | 0.00358322 |
| 540 | KLF12 | chr13:74260148-74708066 | PDT | ATRA+PDT | 4.1927 | 5.99807 | 0.516618 | 0.00005 | 0.00102476 |
| 541 | KLF4 | chr9:110247132-110252047 | PDT | ATRA+PDT | 20.167 | 15.3591 | -0.392903 | 0.00035 | 0.00584867 |
| 542 | KLF6 | chr10:3818187-3827473 | PDT | ATRA+PDT | 116.265 | 182.985 | 0.65431 | 0.00005 | 0.00102476 |
| 543 | KLF9 | chr9:72999512-73029573 | PDT | ATRA+PDT | 1.03209 | 1.46042 | 0.500816 | 0.0024 | 0.0263926 |
| 544 | KLHL24 | chr3:183353410-183402304 | PDT | ATRA+PDT | 9.81722 | 12.0617 | 0.297047 | 0.0038 | 0.0386074 |
| 545 | KLHL25 | chr15:86302558-86338189 | PDT | ATRA+PDT | 5.03188 | 6.50075 | 0.369509 | 0.0031 | 0.0328065 |
| 546 | KLHL5 | chr4:39046450-39127853 | PDT | ATRA+PDT | 2.10193 | 3.84225 | 0.870241 | 0.00005 | 0.00102476 |
| 547 | KLK10 | chr19:51515999-51523431 | PDT | ATRA+PDT | 52.0493 | 83.349 | 0.679285 | 0.00005 | 0.00102476 |
| 548 | KLK12 | chr19:51532347-51538148 | PDT | ATRA+PDT | 2.7591 | 0 | -inf | 0.00005 | 0.00102476 |
| 549 | KLK6 | chr19:51461886-51472929 | PDT | ATRA+PDT | 250.781 | 468.7 | 0.902237 | 0.00005 | 0.00102476 |
| 550 | KRT13 | chr17:39657232-39661865 | PDT | ATRA+PDT | 10.4704 | 14.8408 | 0.503253 | 0.0002 | 0.00358322 |
| 551 | KRT20 | chr17:39032140-39041495 | PDT | ATRA+PDT | 285.041 | 226.863 | -0.329347 | 0.00115 | 0.0151042 |
| 552 | KRT7 | chr12:52626953-52642709 | PDT | ATRA+PDT | 2.09329 | 3.50519 | 0.743723 | 0.0006 | 0.00910057 |
| 553 | KRT80 | chr12:52562779-52585784 | PDT | ATRA+PDT | 37.3471 | 58.4232 | 0.645543 | 0.00005 | 0.00102476 |
| 554 | LAMA5 | chr20:60884120-60942368 | PDT | ATRA+PDT | 13.9332 | 16.9153 | 0.279795 | 0.0043 | 0.0424749 |
| 555 | LAMC2 | chr1:183155173-183214262 | PDT | ATRA+PDT | 11.3733 | 14.2343 | 0.323717 | 0.0011 | 0.0146092 |
| 556 | LAT2 | chr7:73624086-73644164 | PDT | ATRA+PDT | 2.73752 | 1.68019 | -0.704247 | 0.0008 | 0.0114972 |
| 557 | LATS2 | chr13:21547175-21635722 | PDT | ATRA+PDT | 7.21982 | 9.1952 | 0.348918 | 0.0011 | 0.0146092 |
| 558 | LCN2 | chr9:130911731-130915734 | PDT | ATRA+PDT | 67.9208 | 229.774 | 1.75829 | 0.00005 | 0.00102476 |
| 559 | LDHA | chr11:18415935-18429765 | PDT | ATRA+PDT | 520.471 | 387.074 | -0.427208 | 0.00005 | 0.00102476 |
| 560 | LEPREL1 | chr3:189674516-189840226 | PDT | ATRA+PDT | 3.14966 | 0.868411 | -1.85875 | 0.00005 | 0.00102476 |
| 561 | LEPREL4 | chr17:39958204-39968451 | PDT | ATRA+PDT | 5.03458 | 3.84597 | -0.388524 | 0.0035 | 0.0363197 |
| 562 | LGALS3 | chr14:55595934-55612148 | PDT | ATRA+PDT | 93.1758 | 66.449 | -0.487708 | 0.00005 | 0.00102476 |
| 563 | LGALS3BP | chr17:76967334-76976061 | PDT | ATRA+PDT | 169.931 | 209.535 | 0.302242 | 0.0023 | 0.0255827 |
| 564 | LGALS4 | chr19:39292310-39303740 | PDT | ATRA+PDT | 473.603 | 343.759 | -0.46228 | 0.00005 | 0.00102476 |
| 565 | LGALS9 | chr17:25958173-25976586 | PDT | ATRA+PDT | 8.81966 | 19.451 | 1.14105 | 0.00005 | 0.00102476 |
| 566 | LGMN | chr14:93170151-93215047 | PDT | ATRA+PDT | 27.1521 | 41.8574 | 0.624417 | 0.00005 | 0.00102476 |
| 567 | LGR4 | chr11:27387507-27494334 | PDT | ATRA+PDT | 36.4515 | 29.9764 | -0.282149 | 0.00385 | 0.0390041 |
| 568 | LIF | chr22:30636441-30642796 | PDT | ATRA+PDT | 6.03009 | 9.60605 | 0.671763 | 0.00005 | 0.00102476 |
| 569 | LIMA1 | chr12:50569562-50677353 | PDT | ATRA+PDT | 71.8704 | 98.5638 | 0.45566 | 0.00005 | 0.00102476 |
| 570 | LIMS2 | chr2:128395995-128439360 | PDT | ATRA+PDT | 17.001 | 10.3959 | -0.709613 | 0.00005 | 0.00102476 |
| 571 | LINC00263 | chr10:102133332-102148111 | PDT | ATRA+PDT | 9.00983 | 12.3297 | 0.452564 | 0.0023 | 0.0255827 |
| 572 | LINC00346 | chr13:111516333-111522655 | PDT | ATRA+PDT | 1.45147 | 2.37626 | 0.711183 | 0.00005 | 0.00102476 |
| 573 | LIPG | chr18:47088426-47119278 | PDT | ATRA+PDT | 29.3519 | 41.6539 | 0.504996 | 0.00005 | 0.00102476 |
| 574 | LMO7 | chr13:76194569-76434006 | PDT | ATRA+PDT | 63.2636 | 78.1874 | 0.305562 | 0.0023 | 0.0255827 |
| 575 | LOC10028925 | chr17:10698229-10707416 | PDT | ATRA+PDT | 8.25004 | 10.9374 | 0.406794 | 0.0017 | 0.0202822 |
| 576 | LOC10049917 | chr4:83814604-83841284 | PDT | ATRA+PDT | 52.2529 | 65.9451 | 0.335755 | 0.0008 | 0.0114972 |
| 577 | LOC10050654 | chr5:40825364-40829244 | PDT | ATRA+PDT | 21.4239 | 26.6867 | 0.316904 | 0.00195 | 0.0225592 |
| 578 | LOC10085993 | chr5:180256953-180262726 | PDT | ATRA+PDT | 5.83042 | 7.81612 | 0.422853 | 0.0035 | 0.0363197 |
| 579 | LOC146336 | chr16:1114081-1131454 | PDT | ATRA+PDT | 3.05651 | 0.610603 | -2.32358 | 0.00005 | 0.00102476 |
| 580 | LOC388796 | chr20:37049238-37064018 | PDT | ATRA+PDT | 52.6232 | 68.5071 | 0.380555 | 0.0005 | 0.00788626 |
| 581 | LOC400027 | chr12:46119502-46121704 | PDT | ATRA+PDT | 2.69541 | 3.95591 | 0.553504 | 0.00105 | 0.0140676 |
| 582 | LOC439990 | chr10:81967465-81979413 | PDT | ATRA+PDT | 2.51556 | 4.04375 | 0.684811 | 0.00005 | 0.00102476 |
| 583 | LOC541471 | chr2:112124590-112252692 | PDT | ATRA+PDT | 10.7023 | 15.7466 | 0.557115 | 0.00395 | 0.0398659 |
| 584 | LOC550112 | chr4:68566995-68588222 | PDT | ATRA+PDT | 1.86914 | 2.90238 | 0.63486 | 0.001 | 0.0135165 |
| 585 | LOC729966 | chr19:18360759-18366229 | PDT | ATRA+PDT | 16.137 | 21.0502 | 0.383457 | 0.00525 | 0.0496541 |
| 586 | LOC730102 | chr1:177975274-178007142 | PDT | ATRA+PDT | 4.1893 | 3.06842 | -0.449213 | 0.00075 | 0.0109256 |
| 587 | LOXL4 | chr10:100007442-100028007 | PDT | ATRA+PDT | 1.62446 | 4.43023 | 1.44742 | 0.00005 | 0.00102476 |
| 588 | LPCAT1 | chr5:1461541-1524076 | PDT | ATRA+PDT | 23.2019 | 16.2627 | -0.512683 | 0.00005 | 0.00102476 |
| 589 | LPP | chr3:187868993-188608460 | PDT | ATRA+PDT | 3.2364 | 4.52808 | 0.484508 | 0.00005 | 0.00102476 |
| 590 | LRIF1 | chr1:111489811-111506566 | PDT | ATRA+PDT | 35.0027 | 44.2119 | 0.336968 | 0.00185 | 0.0217319 |
| 591 | LRIG1 | chr3:66119284-66550845 | PDT | ATRA+PDT | 5.95481 | 4.12301 | -0.530357 | 0.00055 | 0.0085238 |
| 592 | LRP8 | chr1:53692563-53793821 | PDT | ATRA+PDT | 42.4996 | 30.5249 | -0.477462 | 0.00005 | 0.00102476 |
| 593 | LRRC31 | chr3:169557028-169587660 | PDT | ATRA+PDT | 1.39178 | 0.656509 | -1.08404 | 0.0006 | 0.00910057 |
| 594 | LRRCC1 | chr8:86019322-86058314 | PDT | ATRA+PDT | 3.93226 | 5.094 | 0.373438 | 0.004 | 0.0401429 |
| 595 | LRRK1 | chr15:101459459-101610317 | PDT | ATRA+PDT | 8.13602 | 9.94322 | 0.289389 | 0.0038 | 0.0386074 |
| 596 | LSS | chr21:47608359-47648738 | PDT | ATRA+PDT | 15.5509 | 11.379 | -0.450621 | 0.00005 | 0.00102476 |
| 597 | LTBP3 | chr11:65292547-65325699 | PDT | ATRA+PDT | 4.15846 | 7.24278 | 0.800493 | 0.00055 | 0.0085238 |
| 598 | LTBP4 | chr19:41099071-41135725 | PDT | ATRA+PDT | 4.87091 | 3.07992 | -0.661299 | 0.00005 | 0.00102476 |
| 599 | LYSMD4 | chr15:100267611-100273626 | PDT | ATRA+PDT | 3.71944 | 4.81908 | 0.373671 | 0.0039 | 0.0394358 |
| 600 | MACC1 | chr7:20174278-20257013 | PDT | ATRA+PDT | 3.37263 | 4.86252 | 0.527829 | 0.00005 | 0.00102476 |
| 601 | MACF1 | chr1:39547088-39952810 | PDT | ATRA+PDT | 8.65205 | 11.2662 | 0.380886 | 0.0006 | 0.00910057 |
| 602 | MACROD1 | chr11:63766029-63933585 | PDT | ATRA+PDT | 12.4501 | 8.82427 | -0.496606 | 0.0003 | 0.00511725 |
| 603 | MAFF | chr22:38597938-38612517 | PDT | ATRA+PDT | 54.3501 | 77.4675 | 0.511308 | 0.00005 | 0.00102476 |
| 604 | MALAT1 | chr11:65265232-65273939 | PDT | ATRA+PDT | 48.4011 | 35.4144 | -0.450706 | 0.00005 | 0.00102476 |
| 605 | MANSC1 | chr12:12482217-12503169 | PDT | ATRA+PDT | 16.582 | 12.9391 | -0.357878 | 0.00175 | 0.0207397 |
| 606 | MAOA | chrX:43515408-43606068 | PDT | ATRA+PDT | 6.1329 | 4.33142 | -0.50173 | 0.00005 | 0.00102476 |
| 607 | MAOB | chrX:43625856-43741721 | PDT | ATRA+PDT | 2.99967 | 4.34618 | 0.534944 | 0.0002 | 0.00358322 |
| 608 | MAP1LC3B | chr16:87425800-87438380 | PDT | ATRA+PDT | 43.6199 | 60.1844 | 0.464405 | 0.00005 | 0.00102476 |
| 609 | MAP3K14 | chr17:43325302-43394414 | PDT | ATRA+PDT | 4.90305 | 6.19553 | 0.337549 | 0.005 | 0.0477977 |
| 610 | MAP4K4 | chr2:102314164-102511152 | PDT | ATRA+PDT | 13.2854 | 22.7571 | 0.776472 | 0.00005 | 0.00102476 |
| 611 | MAPRE3 | chr2:27193524-27250087 | PDT | ATRA+PDT | 13.7126 | 8.25953 | -0.731375 | 0.00005 | 0.00102476 |
| 612 | MARS2 | chr2:198570027-198573114 | PDT | ATRA+PDT | 5.86098 | 4.26192 | -0.459639 | 0.00025 | 0.00439786 |

**Table S1: Significant differentially expressed genes (DEGs) after ATRA+PDT compared to PDT.**

| # | gene | locus | sample_1 | sample_2 | value_1 | value_2 | log2(fold_change) | p_value | q_value |
|---|---|---|---|---|---|---|---|---|---|
| 613 | MAT2A | chr2:85766100-85788657 | PDT | ATRA+PDT | 59.464 | 47.5279 | -0.323243 | 0.00245 | 0.0267496 |
| 614 | MB21D1 | chr6:74134855-74162043 | PDT | ATRA+PDT | 29.1977 | 39.2172 | 0.425631 | 0.0001 | 0.00194145 |
| 615 | MBD6 | chr12:57916658-57923931 | PDT | ATRA+PDT | 18.5689 | 22.9335 | 0.304567 | 0.0028 | 0.0301092 |
| 616 | MBOAT1 | chr6:20100934-20212670 | PDT | ATRA+PDT | 1.39007 | 2.26188 | 0.702363 | 0.00005 | 0.00102476 |
| 617 | MBOAT7 | chr19:54677105-54693733 | PDT | ATRA+PDT | 32.6092 | 40.3291 | 0.306542 | 0.0019 | 0.0221245 |
| 618 | MCAM | chr11:119179233-119187840 | PDT | ATRA+PDT | 4.99053 | 3.26224 | -0.613332 | 0.00005 | 0.00102476 |
| 619 | MCL1 | chr1:150547026-150552214 | PDT | ATRA+PDT | 143.54 | 179.524 | 0.322729 | 0.0017 | 0.0202822 |
| 620 | MDK | chr11:46402617-46405375 | PDT | ATRA+PDT | 4.9587 | 26.8511 | 2.43695 | 0.00005 | 0.00102476 |
| 621 | ME1 | chr6:83920109-84140938 | PDT | ATRA+PDT | 53.5323 | 44.352 | -0.271411 | 0.00445 | 0.0435537 |
| 622 | MED15 | chr22:20861885-20941919 | PDT | ATRA+PDT | 45.8395 | 56.1238 | 0.292021 | 0.0034 | 0.0355585 |
| 623 | MEGF6 | chr1:3404505-3528059 | PDT | ATRA+PDT | 0.819526 | 1.67993 | 1.03554 | 0.00005 | 0.00102476 |
| 624 | MEIS2 | chr15:37183221-37393500 | PDT | ATRA+PDT | 7.09086 | 10.394 | 0.551714 | 0.00005 | 0.00102476 |
| 625 | METTL23 | chr17:74722911-74729962 | PDT | ATRA+PDT | 31.6363 | 40.056 | 0.340438 | 0.00445 | 0.0435537 |
| 626 | METTL7B | chr12:56075329-56106089 | PDT | ATRA+PDT | 10.9456 | 5.80892 | -0.914008 | 0.00005 | 0.00102476 |
| 627 | MEX3A | chr1:156041803-156051789 | PDT | ATRA+PDT | 6.24003 | 8.4898 | 0.444176 | 0.00005 | 0.00102476 |
| 628 | MGAT1 | chr5:180217540-180237137 | PDT | ATRA+PDT | 32.4596 | 24.78 | -0.38947 | 0.0003 | 0.00511725 |
| 629 | MGC45800 | chr4:183060158-183065668 | PDT | ATRA+PDT | 1.85735 | 1.25974 | -0.560126 | 0.00045 | 0.00719326 |
| 630 | MIA3 | chr1:222791443-222841351 | PDT | ATRA+PDT | 8.65341 | 10.5993 | 0.292628 | 0.0045 | 0.0439625 |
| 631 | MIB2 | chr1:1550794-1565990 | PDT | ATRA+PDT | 1.57156 | 0.890457 | -0.819578 | 0.0005 | 0.00788626 |
| 632 | MICALL2 | chr7:1473994-1499109 | PDT | ATRA+PDT | 9.61778 | 7.67023 | -0.326434 | 0.00475 | 0.045818 |
| 633 | MID1 | chrX:10413349-10851809 | PDT | ATRA+PDT | 1.16894 | 1.93127 | 0.72435 | 0.00005 | 0.00102476 |
| 634 | MMP15 | chr16:58059281-58080804 | PDT | ATRA+PDT | 10.7939 | 7.95429 | -0.440406 | 0.00005 | 0.00102476 |
| 635 | MMP7 | chr11:102391238-102401478 | PDT | ATRA+PDT | 71.2114 | 148.966 | 1.0648 | 0.00005 | 0.00102476 |
| 636 | MMRN2 | chr10:88695297-88717425 | PDT | ATRA+PDT | 2.00218 | 1.12145 | -0.836208 | 0.00005 | 0.00102476 |
| 637 | MOB3B | chr9:27325206-27529850 | PDT | ATRA+PDT | 4.24961 | 2.94053 | -0.531254 | 0.00005 | 0.00102476 |
| 638 | MPP6 | chr7:24613084-24727498 | PDT | ATRA+PDT | 10.2552 | 7.66042 | -0.420862 | 0.00085 | 0.0119898 |
| 639 | MRPL18 | chr6:160211491-160219461 | PDT | ATRA+PDT | 167.637 | 222.897 | 0.411037 | 0.00005 | 0.00102476 |
| 640 | MRPL2 | chr6:43021766-43027242 | PDT | ATRA+PDT | 35.858 | 28.0635 | -0.353601 | 0.0014 | 0.0175049 |
| 641 | MRPL34 | chr19:17416476-17417652 | PDT | ATRA+PDT | 8.90721 | 6.14515 | -0.535525 | 0.00155 | 0.0189586 |
| 642 | MRPL4 | chr19:10362639-10370736 | PDT | ATRA+PDT | 34.4222 | 26.3991 | -0.382847 | 0.00075 | 0.0109256 |
| 643 | MRPS2 | chr9:138392482-138396519 | PDT | ATRA+PDT | 63.5914 | 50.2259 | -0.3404 | 0.0009 | 0.0125788 |
| 644 | MRPS34 | chr16:1821895-1823140 | PDT | ATRA+PDT | 61.5177 | 47.7933 | -0.364193 | 0.0009 | 0.0125788 |
| 645 | MSL3P1 | chr2:234774089-234777055 | PDT | ATRA+PDT | 11.855 | 8.87928 | -0.416976 | 0.0009 | 0.0125788 |
| 646 | MSLN | chr16:810764-818865 | PDT | ATRA+PDT | 2.81002 | 1.28159 | -1.13265 | 0.00005 | 0.00102476 |
| 647 | MSMO1 | chr4:166248817-166264314 | PDT | ATRA+PDT | 57.2861 | 42.5788 | -0.428049 | 0.00005 | 0.00102476 |
| 648 | MT1E | chr16:56659584-56661024 | PDT | ATRA+PDT | 599.264 | 810.957 | 0.436434 | 0.00005 | 0.00102476 |
| 649 | MT1X | chr16:56716381-56718108 | PDT | ATRA+PDT | 183.778 | 269.734 | 0.553572 | 0.00005 | 0.00102476 |
| 650 | MT2A | chr16:56642477-56643409 | PDT | ATRA+PDT | 727.536 | 1063.81 | 0.548144 | 0.00005 | 0.00102476 |
| 651 | MTMR11 | chr1:149900542-149908791 | PDT | ATRA+PDT | 9.21467 | 7.00351 | -0.395854 | 0.00145 | 0.0179618 |
| 652 | MTSS1 | chr8:125563027-125740730 | PDT | ATRA+PDT | 1.69547 | 5.32383 | 1.65078 | 0.00005 | 0.00102476 |
| 653 | MTUS1 | chr8:17501302-17658426 | PDT | ATRA+PDT | 11.0399 | 7.61404 | -0.535987 | 0.00005 | 0.00102476 |
| 654 | MUC13 | chr3:124624288-124653595 | PDT | ATRA+PDT | 28.9957 | 13.5277 | -1.09993 | 0.00005 | 0.00102476 |
| 655 | MUC2 | chr11:1074874-1104417 | PDT | ATRA+PDT | 2.51755 | 1.38985 | -0.857089 | 0.00005 | 0.00102476 |
| 656 | MUC20 | chr3:195447752-195464540 | PDT | ATRA+PDT | 2.92427 | 4.21721 | 0.528214 | 0.00005 | 0.00102476 |
| 657 | MXD1 | chr2:70142172-70170076 | PDT | ATRA+PDT | 41.2028 | 57.6778 | 0.485271 | 0.00005 | 0.00102476 |
| 658 | MXD4 | chr4:2249159-2263739 | PDT | ATRA+PDT | 4.39778 | 3.34473 | -0.394888 | 0.00165 | 0.0198632 |
| 659 | MYADM | chr19:54369610-54379689 | PDT | ATRA+PDT | 40.4218 | 57.1709 | 0.500148 | 0.00005 | 0.00102476 |
| 660 | MYB | chr6:135502452-135540311 | PDT | ATRA+PDT | 8.49932 | 10.5838 | 0.316437 | 0.00495 | 0.0474046 |
| 661 | MYEOV | chr11:69061621-69064754 | PDT | ATRA+PDT | 36.4184 | 58.2045 | 0.676462 | 0.00005 | 0.00102476 |
| 662 | MYO1B | chr2:192110106-192290115 | PDT | ATRA+PDT | 14.5609 | 20.5701 | 0.498451 | 0.00005 | 0.00102476 |
| 663 | MYOF | chr10:95066185-95242074 | PDT | ATRA+PDT | 26.7778 | 33.3691 | 0.317473 | 0.00125 | 0.0161202 |
| 664 | MYOM3 | chr1:24382530-24438665 | PDT | ATRA+PDT | 5.16202 | 3.00742 | -0.77941 | 0.00005 | 0.00102476 |
| 665 | MYZAP | chr15:57884101-58009755 | PDT | ATRA+PDT | 9.78209 | 15.8685 | 0.697952 | 0.00125 | 0.0161202 |
| 666 | NAA16 | chr13:41885340-41951166 | PDT | ATRA+PDT | 12.5809 | 16.0165 | 0.348331 | 0.00135 | 0.0171 |
| 667 | NAA20 | chr20:19997933-20014273 | PDT | ATRA+PDT | 154.599 | 122.571 | -0.33491 | 0.00035 | 0.00584867 |
| 668 | NAAA | chr4:76831807-76862166 | PDT | ATRA+PDT | 3.09494 | 2.00349 | -0.627393 | 0.003 | 0.0318746 |
| 669 | NARFL | chr16:779768-790997 | PDT | ATRA+PDT | 7.36565 | 5.68953 | -0.372504 | 0.00365 | 0.0375479 |
| 670 | NCOA3 | chr20:46130600-46285621 | PDT | ATRA+PDT | 9.3187 | 12.5625 | 0.430927 | 0.00005 | 0.00102476 |
| 671 | NCOA7 | chr6:126102306-126253176 | PDT | ATRA+PDT | 7.94673 | 10.0077 | 0.332673 | 0.001 | 0.0135165 |
| 672 | NDRG2 | chr14:21484921-21493935 | PDT | ATRA+PDT | 2.02451 | 1.04624 | -0.952352 | 0.0002 | 0.00358322 |
| 673 | NEDD4L | chr18:55711609-56068772 | PDT | ATRA+PDT | 15.5074 | 20.3359 | 0.391068 | 0.00005 | 0.00102476 |
| 674 | NEDD9 | chr6:11183530-11382581 | PDT | ATRA+PDT | 13.5981 | 21.1826 | 0.639473 | 0.00005 | 0.00102476 |
| 675 | NELF | chr9:140342022-140353786 | PDT | ATRA+PDT | 21.2559 | 16.8792 | -0.332613 | 0.00085 | 0.0119898 |
| 676 | NEO1 | chr15:73344824-73597547 | PDT | ATRA+PDT | 4.66325 | 6.55559 | 0.49139 | 0.00005 | 0.00102476 |
| 677 | NFKBIZ | chr3:101498028-101579869 | PDT | ATRA+PDT | 11.8167 | 16.851 | 0.512008 | 0.00005 | 0.00102476 |
| 678 | NFYA | chr6:41040706-41108573 | PDT | ATRA+PDT | 22.5346 | 27.3822 | 0.281099 | 0.0038 | 0.0386074 |
| 679 | NHS | chrX:17393542-17754113 | PDT | ATRA+PDT | 4.98575 | 7.55745 | 0.60009 | 0.00005 | 0.00102476 |
| 680 | NIPAL1 | chr4:48018790-48039080 | PDT | ATRA+PDT | 12.6601 | 17.6904 | 0.482675 | 0.00005 | 0.00102476 |
| 681 | NME3 | chr16:1820320-1821710 | PDT | ATRA+PDT | 8.76519 | 5.55801 | -0.657217 | 0.00055 | 0.0085238 |
| 682 | NME4 | chr16:447191-450754 | PDT | ATRA+PDT | 8.29672 | 5.15854 | -0.685579 | 0.0001 | 0.00194145 |
| 683 | NMI | chr2:152126981-152146430 | PDT | ATRA+PDT | 14.5752 | 23.5218 | 0.690489 | 0.00005 | 0.00102476 |
| 684 | NMUR2 | chr5:151771101-151784840 | PDT | ATRA+PDT | 38.8992 | 51.4381 | 0.403097 | 0.00005 | 0.00102476 |
| 685 | NOSTRIN | chr2:169643048-169721849 | PDT | ATRA+PDT | 4.83264 | 3.53551 | -0.450891 | 0.00255 | 0.0276998 |
| 686 | NOX1 | chrX:100098312-100129334 | PDT | ATRA+PDT | 2.19042 | 1.46503 | -0.580285 | 0.00365 | 0.0375479 |
| 687 | NPAS2 | chr2:101436612-101613287 | PDT | ATRA+PDT | 4.74617 | 6.4326 | 0.438638 | 0.00085 | 0.0119898 |
| 688 | NPC1 | chr18:21083461-21166581 | PDT | ATRA+PDT | 30.5634 | 38.4756 | 0.332142 | 0.001 | 0.0135165 |
| 689 | NPDC1 | chr9:139933908-139940676 | PDT | ATRA+PDT | 15.0162 | 8.51311 | -0.818761 | 0.00005 | 0.00102476 |
| 690 | NPNT | chr4:106816596-106892828 | PDT | ATRA+PDT | 25.931 | 19.6757 | -0.398259 | 0.00005 | 0.00102476 |
| 691 | NQO2 | chr6:3000066-3019994 | PDT | ATRA+PDT | 19.8193 | 12.8652 | -0.623437 | 0.00005 | 0.00102476 |
| 692 | NR0B2 | chr1:27237974-27240567 | PDT | ATRA+PDT | 1.38854 | 2.94617 | 1.08527 | 0.00055 | 0.0085238 |
| 693 | NR1D1 | chr17:38218445-38256973 | PDT | ATRA+PDT | 15.3103 | 24.2233 | 0.661888 | 0.00005 | 0.00102476 |
| 694 | NR1D2 | chr3:23986750-24022109 | PDT | ATRA+PDT | 25.6726 | 31.1112 | 0.277209 | 0.00475 | 0.045818 |
| 695 | NR5A2 | chr1:199996769-200146550 | PDT | ATRA+PDT | 1.27497 | 2.3158 | 0.861049 | 0.00005 | 0.00102476 |
| 696 | NRIP1 | chr21:16333555-16437126 | PDT | ATRA+PDT | 9.31825 | 11.6883 | 0.326936 | 0.0012 | 0.0156263 |
| 697 | NRP1 | chr10:33466418-33623833 | PDT | ATRA+PDT | 4.46624 | 6.98659 | 0.645528 | 0.00015 | 0.00281494 |
| 698 | NRSN2 | chr20:327369-335512 | PDT | ATRA+PDT | 2.17623 | 1.18721 | -0.874258 | 0.0003 | 0.00511725 |
| 699 | NT5E | chr6:86159301-86205509 | PDT | ATRA+PDT | 19.0717 | 13.5303 | -0.495233 | 0.00005 | 0.00102476 |
| 700 | NTN4 | chr12:96051582-96184536 | PDT | ATRA+PDT | 35.7761 | 60.9329 | 0.768226 | 0.00005 | 0.00102476 |
| 701 | NUCB2 | chr11:17298285-17353070 | PDT | ATRA+PDT | 10.6956 | 7.62311 | -0.488564 | 0.0001 | 0.00194145 |
| 702 | NUDC | chr1:27248223-27272887 | PDT | ATRA+PDT | 212.793 | 262.453 | 0.302609 | 0.0017 | 0.0202822 |
| 703 | NUDT19 | chr19:33182866-33204702 | PDT | ATRA+PDT | 9.55354 | 6.97424 | -0.453999 | 0.00015 | 0.00281494 |
| 704 | OAS3 | chr12:113376248-113411054 | PDT | ATRA+PDT | 4.98403 | 6.86002 | 0.460901 | 0.00005 | 0.00102476 |
| 705 | OBFC2A | chr2:192542797-192553248 | PDT | ATRA+PDT | 7.33341 | 10.0349 | 0.452475 | 0.00005 | 0.00102476 |
| 706 | ONECUT2 | chr18:55102916-55158530 | PDT | ATRA+PDT | 5.40996 | 6.58805 | 0.284233 | 0.00415 | 0.041376 |
| 707 | OR7E14P | chr11:17073432-17074591 | PDT | ATRA+PDT | 6.80136 | 9.1838 | 0.433267 | 0.00385 | 0.0390041 |
| 708 | OSBPL5 | chr11:3108345-3186582 | PDT | ATRA+PDT | 10.7194 | 7.38127 | -0.53829 | 0.00005 | 0.00102476 |
| 709 | OTUD7B | chr1:149912231-149982686 | PDT | ATRA+PDT | 5.74496 | 7.20416 | 0.326533 | 0.0022 | 0.024728 |
| 710 | OVOL1 | chr11:65554504-65564690 | PDT | ATRA+PDT | 8.87289 | 11.53 | 0.377916 | 0.00075 | 0.0109256 |
| 711 | PADI1 | chr1:17531620-17572501 | PDT | ATRA+PDT | 2.82304 | 2.17378 | -0.377045 | 0.0051 | 0.048623 |
| 712 | PADI2 | chr1:17393255-17445948 | PDT | ATRA+PDT | 1.47832 | 0.896892 | -0.720949 | 0.0006 | 0.00910057 |
| 713 | PAG1 | chr8:81880045-82024303 | PDT | ATRA+PDT | 1.66817 | 0.914237 | -0.867625 | 0.00005 | 0.00102476 |
| 714 | PAPSS1 | chr4:108534821-108641419 | PDT | ATRA+PDT | 17.4956 | 22.2315 | 0.345608 | 0.00115 | 0.0151042 |

#### Table S1: Significant differentially expressed genes (DEGs) after ATRA+PDT compared to PDT.

| # | gene | locus | sample_1 | sample_2 | value_1 | value_2 | log2(fold_change) | p_value | q_value |
|---|---|---|---|---|---|---|---|---|---|
| 715 | PAQR5 | chr15:69591293-69699976 | PDT | ATRA+PDT | 29.5384 | 20.7262 | -0.511137 | 0.00005 | 0.00102476 |
| 716 | PARD6B | chr20:49348080-49370278 | PDT | ATRA+PDT | 10.5288 | 13.243 | 0.330893 | 0.0016 | 0.0194588 |
| 717 | PARP10 | chr8:145051319-145060635 | PDT | ATRA+PDT | 0.401217 | 1.33341 | 1.73267 | 0.00005 | 0.00102476 |
| 718 | PARP12 | chr7:139723548-139763521 | PDT | ATRA+PDT | 4.93932 | 7.8736 | 0.672709 | 0.00005 | 0.00102476 |
| 719 | PARP14 | chr3:122399671-122449687 | PDT | ATRA+PDT | 8.13795 | 15.8278 | 0.959724 | 0.00005 | 0.00102476 |
| 720 | PARP8 | chr5:49961732-50142356 | PDT | ATRA+PDT | 1.65131 | 2.28313 | 0.467401 | 0.0002 | 0.00358322 |
| 721 | PATZ1 | chr22:31721789-31742249 | PDT | ATRA+PDT | 6.64987 | 8.72047 | 0.391079 | 0.001 | 0.0135165 |
| 722 | PCK2 | chr14:24563482-24573339 | PDT | ATRA+PDT | 15.5475 | 19.8844 | 0.354956 | 0.00145 | 0.0179618 |
| 723 | PCSK5 | chr9:78505559-78977255 | PDT | ATRA+PDT | 7.01257 | 10.5902 | 0.59471 | 0.00005 | 0.00102476 |
| 724 | PCSK6 | chr15:101844132-102030187 | PDT | ATRA+PDT | 17.9251 | 14.5859 | -0.297413 | 0.0046 | 0.0446535 |
| 725 | PDE12 | chr3:57541980-57547768 | PDT | ATRA+PDT | 7.58494 | 5.83222 | -0.379092 | 0.00125 | 0.0161202 |
| 726 | PDE9A | chr21:44073861-44195618 | PDT | ATRA+PDT | 2.65823 | 4.41355 | 0.731474 | 0.0004 | 0.00650107 |
| 727 | PDIA5 | chr3:122785855-122880953 | PDT | ATRA+PDT | 14.8652 | 11.5385 | -0.36549 | 0.0017 | 0.0202822 |
| 728 | PDK4 | chr7:95212808-95225925 | PDT | ATRA+PDT | 1.91366 | 2.81879 | 0.55874 | 0.00015 | 0.00281494 |
| 729 | PDLIM1 | chr10:96997329-97050781 | PDT | ATRA+PDT | 72.8709 | 101.499 | 0.478051 | 0.00005 | 0.00102476 |
| 730 | PDLIM7 | chr5:176910394-176924602 | PDT | ATRA+PDT | 11.7486 | 8.96007 | -0.390911 | 0.00415 | 0.041376 |
| 731 | PELI2 | chr14:56585092-56768031 | PDT | ATRA+PDT | 1.87101 | 3.7817 | 1.01522 | 0.00005 | 0.00102476 |
| 732 | PEX11A | chr15:90226286-90233958 | PDT | ATRA+PDT | 11.7028 | 7.50754 | -0.640445 | 0.00005 | 0.00102476 |
| 733 | PFKM | chr12:48499655-48540187 | PDT | ATRA+PDT | 33.1889 | 26.1917 | -0.341586 | 0.0005 | 0.00788626 |
| 734 | PGM1 | chr1:64058946-64125916 | PDT | ATRA+PDT | 8.02227 | 5.37656 | -0.577327 | 0.00005 | 0.00102476 |
| 735 | PGM2L1 | chr11:74041360-74109502 | PDT | ATRA+PDT | 0.714012 | 1.42471 | 0.996651 | 0.00005 | 0.00102476 |
| 736 | PGPEP1 | chr19:18451407-18480763 | PDT | ATRA+PDT | 2.68912 | 4.16227 | 0.630238 | 0.00005 | 0.00102476 |
| 737 | PHF21A | chr11:45950869-46142985 | PDT | ATRA+PDT | 1.81217 | 2.57575 | 0.507269 | 0.00005 | 0.00102476 |
| 738 | PHLDA1 | chr12:76419226-76425556 | PDT | ATRA+PDT | 28.3072 | 49.6068 | 0.80937 | 0.00005 | 0.00102476 |
| 739 | PHLPP1 | chr18:60382671-60647676 | PDT | ATRA+PDT | 3.19185 | 4.34827 | 0.446049 | 0.0001 | 0.00194145 |
| 740 | PI3 | chr20:43803539-43805185 | PDT | ATRA+PDT | 11.7493 | 6.65439 | -0.82019 | 0.001 | 0.0135165 |
| 741 | PIK3R1 | chr5:67511583-67597649 | PDT | ATRA+PDT | 17.1353 | 13.837 | -0.308442 | 0.0019 | 0.0221245 |
| 742 | PIM1 | chr6:37137921-37143204 | PDT | ATRA+PDT | 11.4269 | 18.507 | 0.695638 | 0.00005 | 0.00102476 |
| 743 | PIP5K1A | chr1:151171020-151222007 | PDT | ATRA+PDT | 33.9012 | 42.7141 | 0.333379 | 0.00085 | 0.0119898 |
| 744 | PITPNC1 | chr17:65373923-65689647 | PDT | ATRA+PDT | 31.2742 | 43.5841 | 0.478831 | 0.00005 | 0.00102476 |
| 745 | PITPNM3 | chr17:6354582-6459877 | PDT | ATRA+PDT | 1.93026 | 1.34623 | -0.519863 | 0.00005 | 0.00102476 |
| 746 | PJA1 | chrX:68380580-68385365 | PDT | ATRA+PDT | 3.37515 | 5.14484 | 0.608175 | 0.00005 | 0.00102476 |
| 747 | PKIB | chr6:122793061-123047518 | PDT | ATRA+PDT | 25.5099 | 12.9605 | -0.976938 | 0.00005 | 0.00102476 |
| 748 | PKN1 | chr19:14544165-14582679 | PDT | ATRA+PDT | 8.80354 | 16.3822 | 0.895976 | 0.00005 | 0.00102476 |
| 749 | PKNOX1 | chr21:44394642-44453688 | PDT | ATRA+PDT | 11.0656 | 13.5507 | 0.292295 | 0.0048 | 0.0461751 |
| 750 | PLA2G16 | chr11:63341943-63381941 | PDT | ATRA+PDT | 137.417 | 169.418 | 0.302024 | 0.0024 | 0.0263926 |
| 751 | PLAC8 | chr4:84011210-84035911 | PDT | ATRA+PDT | 2.40008 | 5.49895 | 1.19608 | 0.00005 | 0.00102476 |
| 752 | PLCXD1 | chrY:142990-170022 | PDT | ATRA+PDT | 7.37322 | 4.9257 | -0.581968 | 0.00005 | 0.00102476 |
| 753 | PLD1 | chr3:171318194-171528284 | PDT | ATRA+PDT | 1.88317 | 1.23716 | -0.606141 | 0.00005 | 0.00102476 |
| 754 | PLD2 | chr17:4710395-4726727 | PDT | ATRA+PDT | 2.13226 | 3.16203 | 0.56847 | 0.0001 | 0.00194145 |
| 755 | PLEKHA2 | chr8:38758752-38831430 | PDT | ATRA+PDT | 1.47715 | 2.04398 | 0.468562 | 0.00045 | 0.00719326 |
| 756 | PLEKHG5 | chr1:6521213-6580069 | PDT | ATRA+PDT | 2.24091 | 1.63582 | -0.454065 | 0.0014 | 0.0175049 |
| 757 | PLK3 | chr1:45266035-45272957 | PDT | ATRA+PDT | 12.6041 | 17.9976 | 0.513909 | 0.00005 | 0.00102476 |
| 758 | PLSCR1 | chr3:146232966-146262628 | PDT | ATRA+PDT | 22.742 | 30.7511 | 0.435277 | 0.00005 | 0.00102476 |
| 759 | PLXNB2 | chr22:50713407-50746001 | PDT | ATRA+PDT | 36.1477 | 43.869 | 0.279295 | 0.0043 | 0.0424749 |
| 760 | PLXND1 | chr3:129274055-129325582 | PDT | ATRA+PDT | 4.77635 | 3.82235 | -0.32145 | 0.0049 | 0.0470101 |
| 761 | PMAIP1 | chr18:57567191-57571538 | PDT | ATRA+PDT | 27.4374 | 39.4407 | 0.523542 | 0.00005 | 0.00102476 |
| 762 | PML | chr15:74287013-74340155 | PDT | ATRA+PDT | 2.86592 | 4.01457 | 0.486249 | 0.0006 | 0.00910057 |
| 763 | PMPCA | chr9:139305115-139318213 | PDT | ATRA+PDT | 61.022 | 49.0771 | -0.314278 | 0.0015 | 0.0184954 |
| 764 | PMS2CL | chr7:6774935-6791232 | PDT | ATRA+PDT | 1.72805 | 3.29832 | 0.932582 | 0.00015 | 0.00281494 |
| 765 | PNPLA2 | chr11:818900-825571 | PDT | ATRA+PDT | 40.2315 | 51.0456 | 0.343462 | 0.0007 | 0.0103814 |
| 766 | POLI | chr18:51795848-51824604 | PDT | ATRA+PDT | 0.696334 | 1.13724 | 0.70768 | 0.0004 | 0.00650107 |
| 767 | POLR2L | chr11:839720-842529 | PDT | ATRA+PDT | 123.563 | 94.9511 | -0.37999 | 0.00005 | 0.00102476 |
| 768 | PPAP2B | chr1:56960418-57045257 | PDT | ATRA+PDT | 2.69017 | 3.70388 | 0.461337 | 0.0006 | 0.00910057 |
| 769 | PPARG | chr3:12329348-12475855 | PDT | ATRA+PDT | 63.8493 | 81.2108 | 0.347002 | 0.00035 | 0.00584867 |
| 770 | PPFIBP2 | chr11:7534995-7674996 | PDT | ATRA+PDT | 3.94301 | 2.67874 | -0.557743 | 0.0002 | 0.00358322 |
| 771 | PPIF | chr10:81107219-81115089 | PDT | ATRA+PDT | 91.7911 | 72.915 | -0.33214 | 0.0004 | 0.00650107 |
| 772 | PPL | chr16:4932507-4987136 | PDT | ATRA+PDT | 16.0735 | 20.51 | 0.351644 | 0.00035 | 0.00584867 |
| 773 | PPP1R10 | chr6_ssto_hap7:1900460-191729 | PDT | ATRA+PDT | 31.3853 | 42.0273 | 0.421239 | 0.00005 | 0.00102476 |
| 774 | PPP1R15A | chr19:49375648-49379319 | PDT | ATRA+PDT | 282.868 | 421 | 0.57369 | 0.00005 | 0.00102476 |
| 775 | PPP1R1B | chr17:37783176-37792878 | PDT | ATRA+PDT | 8.15642 | 2.98407 | -1.45065 | 0.00005 | 0.00102476 |
| 776 | PPP1R21 | chr2:48667907-48742531 | PDT | ATRA+PDT | 4.05097 | 5.46955 | 0.433155 | 0.00075 | 0.0109256 |
| 777 | PPP1R3B | chr8:8993763-9009152 | PDT | ATRA+PDT | 3.24328 | 2.51084 | -0.369287 | 0.00365 | 0.0375479 |
| 778 | PPP2R5B | chr11:64692142-64703360 | PDT | ATRA+PDT | 1.19621 | 1.8247 | 0.609196 | 0.0052 | 0.0492248 |
| 779 | PPP4R1L | chr20:56807832-56884495 | PDT | ATRA+PDT | 3.49219 | 4.75011 | 0.443829 | 0.00165 | 0.0198632 |
| 780 | PRAP1 | chr10:135160843-135166187 | PDT | ATRA+PDT | 3.20396 | 1.43399 | -1.15982 | 0.00515 | 0.0488381 |
| 781 | PRDM8 | chr4:81106423-81125482 | PDT | ATRA+PDT | 1.23561 | 1.83677 | 0.57195 | 0.0016 | 0.0194588 |
| 782 | PRDX4 | chrX:23685644-23704514 | PDT | ATRA+PDT | 57.3371 | 45.153 | -0.344648 | 0.00155 | 0.0189586 |
| 783 | PRIC285 | chr20:62189438-62205592 | PDT | ATRA+PDT | 3.42848 | 5.91896 | 0.787776 | 0.00005 | 0.00102476 |
| 784 | PRKAR2A | chr3:48788092-48885270 | PDT | ATRA+PDT | 16.3822 | 12.1706 | -0.42873 | 0.0003 | 0.00511725 |
| 785 | PRKCA | chr17:64298925-64806862 | PDT | ATRA+PDT | 5.41637 | 6.90526 | 0.35037 | 0.00075 | 0.0109256 |
| 786 | PRKD2 | chr19:47150868-47220384 | PDT | ATRA+PDT | 33.1834 | 42.336 | 0.351424 | 0.00055 | 0.0085238 |
| 787 | PROCR | chr20:33759773-33765165 | PDT | ATRA+PDT | 45.1156 | 30.1563 | -0.581167 | 0.00005 | 0.00102476 |
| 788 | PRODH | chr22:18900286-18924066 | PDT | ATRA+PDT | 1.66635 | 0.459819 | -1.85756 | 0.0001 | 0.00194145 |
| 789 | PROS1 | chr3:93591880-93692934 | PDT | ATRA+PDT | 4.99684 | 3.49117 | -0.517305 | 0.00025 | 0.00439786 |
| 790 | PRR15L | chr17:46029333-46035110 | PDT | ATRA+PDT | 3.83366 | 2.37869 | -0.688556 | 0.00145 | 0.0179618 |
| 791 | PRRG4 | chr11:32851480-32879669 | PDT | ATRA+PDT | 9.22956 | 11.4977 | 0.317005 | 0.00255 | 0.0276998 |
| 792 | PSCA | chr8:143751725-143764145 | PDT | ATRA+PDT | 1.7825 | 0.615034 | -1.53516 | 0.002 | 0.0229388 |
| 793 | PSMB10 | chr16:67968406-67970780 | PDT | ATRA+PDT | 18.4903 | 29.683 | 0.682866 | 0.00005 | 0.00102476 |
| 794 | PSMB8 | chr6_ssto_hap7:4239266-425252 | PDT | ATRA+PDT | 1.01457 | 4.79809 | 2.24159 | 0.00005 | 0.00102476 |
| 795 | PSME1 | chr14:24605377-24610797 | PDT | ATRA+PDT | 97.3296 | 148.202 | 0.606612 | 0.00005 | 0.00102476 |
| 796 | PSME2 | chr14:24612573-24615855 | PDT | ATRA+PDT | 82.8818 | 113.648 | 0.455451 | 0.00005 | 0.00102476 |
| 797 | PSPH | chr7:56078743-56119268 | PDT | ATRA+PDT | 23.4394 | 28.9095 | 0.302606 | 0.0037 | 0.0379161 |
| 798 | PTGS2 | chr1:186640943-186649559 | PDT | ATRA+PDT | 3.23432 | 5.40639 | 0.741203 | 0.00005 | 0.00102476 |
| 799 | PTPN21 | chr14:88932121-89021123 | PDT | ATRA+PDT | 3.89511 | 5.04068 | 0.371953 | 0.0014 | 0.0175049 |
| 800 | PTPN9 | chr15:75759461-75871625 | PDT | ATRA+PDT | 15.0615 | 18.4692 | 0.294255 | 0.0047 | 0.0455001 |
| 801 | PTPRB | chr12:70910631-71031220 | PDT | ATRA+PDT | 2.99581 | 3.86343 | 0.366936 | 0.0007 | 0.0103814 |
| 802 | PTPRH | chr19:55692614-55720874 | PDT | ATRA+PDT | 24.2026 | 47.4306 | 0.970657 | 0.00005 | 0.00102476 |
| 803 | PXMP2 | chr12:133264191-133281577 | PDT | ATRA+PDT | 30.9691 | 22.2824 | -0.474925 | 0.00025 | 0.00439786 |
| 804 | PYGB | chr20:25228705-25371618 | PDT | ATRA+PDT | 220.407 | 173.113 | -0.348456 | 0.00145 | 0.0179618 |
| 805 | QPCT | chr2:37571752-37600465 | PDT | ATRA+PDT | 28.6729 | 21.0747 | -0.444174 | 0.00005 | 0.00102476 |
| 806 | QSOX1 | chr1:180123967-180169859 | PDT | ATRA+PDT | 33.2027 | 24.2491 | -0.453372 | 0.0001 | 0.00194145 |
| 807 | RAB27B | chr18:52495707-52562747 | PDT | ATRA+PDT | 4.65883 | 3.32112 | -0.4883 | 0.00005 | 0.00102476 |
| 808 | RAB30 | chr11:82692477-82782884 | PDT | ATRA+PDT | 1.93354 | 3.05249 | 0.658741 | 0.0046 | 0.0446535 |
| 809 | RAB3D | chr19:11406823-11450344 | PDT | ATRA+PDT | 5.91284 | 3.44236 | -0.780454 | 0.00005 | 0.00102476 |
| 810 | RAB40B | chr17:80614942-80656598 | PDT | ATRA+PDT | 10.2786 | 7.73325 | -0.410498 | 0.00135 | 0.0171 |
| 811 | RAB40C | chr16:639356-679273 | PDT | ATRA+PDT | 9.25892 | 7.21951 | -0.358942 | 0.00405 | 0.0405304 |
| 812 | RAET1K | chr6:150319154-150326280 | PDT | ATRA+PDT | 0.984729 | 1.75706 | 0.83536 | 0.0044 | 0.0431831 |
| 813 | RALGAPA2 | chr20:20370271-20693266 | PDT | ATRA+PDT | 8.27806 | 10.3653 | 0.324402 | 0.00175 | 0.0207397 |
| 814 | RANBP10 | chr16:67757004-67840555 | PDT | ATRA+PDT | 4.63285 | 5.81037 | 0.32673 | 0.00455 | 0.0443293 |
| 815 | RAP1GAP | chr1:21922707-21995856 | PDT | ATRA+PDT | 32.5006 | 21.8294 | -0.574195 | 0.00005 | 0.00102476 |
| 816 | RAP2B | chr3:152880028-152886263 | PDT | ATRA+PDT | 15.2802 | 19.9637 | 0.385712 | 0.0002 | 0.00358322 |

#### Table S1: Significant differentially expressed genes (DEGs) after ATRA+PDT compared to PDT.

| # | gene | locus | sample_1 | sample_2 | value_1 | value_2 | log2(fold_change) | p_value | q_value |
|---|---|---|---|---|---|---|---|---|---|
| 817 | RARG | chr12:53604349-53626040 | PDT | ATRA+PDT | 37.369 | 57.0455 | 0.610268 | 0.00005 | 0.00102476 |
| 818 | RARRES1 | chr3:158414896-158450275 | PDT | ATRA+PDT | 2.68665 | 16.1452 | 2.58723 | 0.00005 | 0.00102476 |
| 819 | RARRES3 | chr11:63014620-63330855 | PDT | ATRA+PDT | 7.65704 | 32.1209 | 2.06866 | 0.00005 | 0.00102476 |
| 820 | RASA1 | chr5:86564069-86687743 | PDT | ATRA+PDT | 32.5508 | 47.4891 | 0.544902 | 0.00005 | 0.00102476 |
| 821 | RASSF6 | chr4:74438861-74486340 | PDT | ATRA+PDT | 4.25043 | 5.66768 | 0.415152 | 0.0007 | 0.0103814 |
| 822 | RBM48 | chr7:92158086-92166823 | PDT | ATRA+PDT | 9.83332 | 12.7478 | 0.374499 | 0.00345 | 0.0359757 |
| 823 | RBPMS | chr8:30239634-30429734 | PDT | ATRA+PDT | 5.87097 | 8.64539 | 0.558333 | 0.00015 | 0.00281494 |
| 824 | RBPMS2 | chr15:65032094-65067770 | PDT | ATRA+PDT | 3.70713 | 2.624 | -0.498537 | 0.0038 | 0.0386074 |
| 825 | RCAN1 | chr21:35888783-35987382 | PDT | ATRA+PDT | 4.20594 | 6.83073 | 0.69961 | 0.00005 | 0.00102476 |
| 826 | RCAN3 | chr1:24822822-24863510 | PDT | ATRA+PDT | 12.2376 | 18.93 | 0.629347 | 0.00005 | 0.00102476 |
| 827 | RCBTB1 | chr13:50106081-50159719 | PDT | ATRA+PDT | 8.17925 | 11.2587 | 0.461001 | 0.00005 | 0.00102476 |
| 828 | RCN1 | chr11:32112476-32127272 | PDT | ATRA+PDT | 16.5146 | 24.007 | 0.539712 | 0.00005 | 0.00102476 |
| 829 | REG4 | chr1:120336640-120354203 | PDT | ATRA+PDT | 70.8796 | 2.04563 | -5.11475 | 0.00005 | 0.00102476 |
| 830 | REL | chr2:61108751-61150178 | PDT | ATRA+PDT | 4.36843 | 5.64892 | 0.37086 | 0.0039 | 0.0394358 |
| 831 | REN | chr1:204123943-204135465 | PDT | ATRA+PDT | 0.636161 | 2.66058 | 2.06428 | 0.00005 | 0.00102476 |
| 832 | REPS2 | chrX:16964813-17171403 | PDT | ATRA+PDT | 2.24299 | 2.91552 | 0.378333 | 0.00235 | 0.0259765 |
| 833 | REXO2 | chr11:114310107-114321000 | PDT | ATRA+PDT | 94.0016 | 77.1618 | -0.284798 | 0.00515 | 0.0488381 |
| 834 | RG9MTD1 | chr3:101280711-101285089 | PDT | ATRA+PDT | 57.3518 | 42.9105 | -0.418507 | 0.00005 | 0.00102476 |
| 835 | RGS2 | chr1:192778168-192781407 | PDT | ATRA+PDT | 28.7673 | 45.9333 | 0.67511 | 0.00005 | 0.00102476 |
| 836 | RHBDD2 | chr7:75508316-75518244 | PDT | ATRA+PDT | 24.1826 | 30.4657 | 0.333219 | 0.00235 | 0.0259765 |
| 837 | RHBDD3 | chr22:29655843-29663914 | PDT | ATRA+PDT | 8.435 | 10.8511 | 0.363385 | 0.0043 | 0.0424749 |
| 838 | RHOB | chr2:20646834-20649201 | PDT | ATRA+PDT | 80.1921 | 101.159 | 0.33509 | 0.00055 | 0.0085238 |
| 839 | RHOD | chr11:66824288-66839488 | PDT | ATRA+PDT | 10.1643 | 7.44284 | -0.44958 | 0.0025 | 0.027212 |
| 840 | RHOU | chr1:228780393-228882416 | PDT | ATRA+PDT | 5.61287 | 2.90298 | -0.951202 | 0.00005 | 0.00102476 |
| 841 | RNASE4 | chr14:21152335-21168758 | PDT | ATRA+PDT | 2.37664 | 0.837993 | -1.50391 | 0.00085 | 0.0119898 |
| 842 | RND3 | chr2:151324706-151344209 | PDT | ATRA+PDT | 49.8429 | 67.3551 | 0.434401 | 0.00005 | 0.00102476 |
| 843 | RNF126 | chr19:647525-663233 | PDT | ATRA+PDT | 18.2231 | 13.7825 | -0.402933 | 0.0006 | 0.00910057 |
| 844 | RNF128 | chrX:105937067-106040246 | PDT | ATRA+PDT | 24.7876 | 18.2125 | -0.444685 | 0.00005 | 0.00102476 |
| 845 | RNF19A | chr8:101269287-101322327 | PDT | ATRA+PDT | 11.9951 | 14.7277 | 0.296081 | 0.00435 | 0.0428104 |
| 846 | RNF213 | chr17:78234659-78411884 | PDT | ATRA+PDT | 3.77922 | 5.10192 | 0.432953 | 0.0004 | 0.00650107 |
| 847 | RNF24 | chr20:3912068-3996216 | PDT | ATRA+PDT | 8.05694 | 5.00173 | -0.687806 | 0.00005 | 0.00102476 |
| 848 | ROR1 | chr1:64239689-64644707 | PDT | ATRA+PDT | 10.0918 | 17.2802 | 0.77594 | 0.00005 | 0.00102476 |
| 849 | RPH3AL | chr17:62179-202633 | PDT | ATRA+PDT | 5.19426 | 6.82169 | 0.393211 | 0.0021 | 0.0238805 |
| 850 | RPS6KB2 | chr11:67195934-67202879 | PDT | ATRA+PDT | 32.1139 | 25.1005 | -0.355484 | 0.0008 | 0.0114972 |
| 851 | RPUSD1 | chr16:834973-838383 | PDT | ATRA+PDT | 16.0041 | 11.7535 | -0.445345 | 0.0002 | 0.00358322 |
| 852 | RTKN | chr2:74652987-74669060 | PDT | ATRA+PDT | 17.4717 | 13.5318 | -0.368662 | 0.00115 | 0.0151042 |
| 853 | RUNX2 | chr6:44796469-45518819 | PDT | ATRA+PDT | 0.807263 | 1.52946 | 0.921912 | 0.0022 | 0.024728 |
| 854 | S100A10 | chr1:151955385-151966714 | PDT | ATRA+PDT | 722.813 | 500.148 | -0.531269 | 0.00005 | 0.00102476 |
| 855 | S100A14 | chr1:153586731-153588808 | PDT | ATRA+PDT | 76.7108 | 35.2653 | -1.12118 | 0.00005 | 0.00102476 |
| 856 | S100A16 | chr1:153579366-153585514 | PDT | ATRA+PDT | 239.074 | 168.412 | -0.505466 | 0.00005 | 0.00102476 |
| 857 | S100A2 | chr1:153533584-153538306 | PDT | ATRA+PDT | 18.6421 | 14.2378 | -0.388835 | 0.002 | 0.0229388 |
| 858 | S100A4 | chr1:153516094-153518282 | PDT | ATRA+PDT | 326.579 | 215.965 | -0.596635 | 0.00005 | 0.00102476 |
| 859 | SAMD12 | chr8:119201694-119738306 | PDT | ATRA+PDT | 7.72496 | 10.9211 | 0.499518 | 0.00005 | 0.00102476 |
| 860 | SAMD4A | chr14:55034329-55260033 | PDT | ATRA+PDT | 9.68298 | 12.3034 | 0.345532 | 0.0007 | 0.0103814 |
| 861 | SAMD9 | chr7:92728825-92747336 | PDT | ATRA+PDT | 8.37085 | 23.382 | 1.48196 | 0.00005 | 0.00102476 |
| 862 | SAPCD2 | chr9:139956578-139965028 | PDT | ATRA+PDT | 32.3841 | 26.2546 | -0.302715 | 0.0022 | 0.024728 |
| 863 | SAT1 | chrX:23801274-23804327 | PDT | ATRA+PDT | 77.3754 | 104.301 | 0.4308 | 0.00005 | 0.00102476 |
| 864 | SBNO2 | chr19:1107632-1174282 | PDT | ATRA+PDT | 7.70866 | 10.4313 | 0.436367 | 0.00005 | 0.00102476 |
| 865 | SC5DL | chr11:121163387-121184119 | PDT | ATRA+PDT | 25.2215 | 19.8586 | -0.344895 | 0.00025 | 0.00439786 |
| 866 | SCARB1 | chr12:125262173-125348519 | PDT | ATRA+PDT | 40.9505 | 31.1269 | -0.395722 | 0.00005 | 0.00102476 |
| 867 | SCEL | chr13:78109808-78219398 | PDT | ATRA+PDT | 9.97633 | 19.3169 | 0.953286 | 0.00005 | 0.00102476 |
| 868 | SCO1 | chr17:10583648-10600885 | PDT | ATRA+PDT | 21.1395 | 16.1884 | -0.384976 | 0.0006 | 0.00910057 |
| 869 | SDC4 | chr20:43953928-43977064 | PDT | ATRA+PDT | 97.778 | 124.698 | 0.350852 | 0.0002 | 0.00358322 |
| 870 | SDCCAG8 | chr1:243419306-244006886 | PDT | ATRA+PDT | 4.41229 | 5.76522 | 0.385848 | 0.0036 | 0.0371409 |
| 871 | SDF2L1 | chr22:21996541-21998588 | PDT | ATRA+PDT | 13.9605 | 10.0903 | -0.468384 | 0.0018 | 0.0212615 |
| 872 | SEC11C | chr18:56807124-56826063 | PDT | ATRA+PDT | 36.4528 | 28.2563 | -0.36746 | 0.0033 | 0.0346484 |
| 873 | SEC14L2 | chr22:30792929-30821291 | PDT | ATRA+PDT | 2.02921 | 3.7383 | 0.881462 | 0.00005 | 0.00102476 |
| 874 | SEC24D | chr4:119643977-119757326 | PDT | ATRA+PDT | 10.2168 | 7.21767 | -0.501332 | 0.00005 | 0.00102476 |
| 875 | SEC61A1 | chr3:127771211-127790526 | PDT | ATRA+PDT | 128.699 | 104.128 | -0.30565 | 0.00235 | 0.0259765 |
| 876 | SEC61B | chr9:101984569-101992901 | PDT | ATRA+PDT | 162.163 | 128.308 | -0.337826 | 0.00125 | 0.0161202 |
| 877 | SEC61G | chr7:54819939-54826939 | PDT | ATRA+PDT | 301.576 | 241.039 | -0.323257 | 0.0022 | 0.024728 |
| 878 | SEMA3A | chr7:83587658-83824217 | PDT | ATRA+PDT | 5.20312 | 7.70564 | 0.566537 | 0.00005 | 0.00102476 |
| 879 | SEMA3B | chr3:50305039-50314572 | PDT | ATRA+PDT | 13.5343 | 16.9983 | 0.328766 | 0.0022 | 0.024728 |
| 880 | SEMA3E | chr7:82993221-83278479 | PDT | ATRA+PDT | 1.42158 | 0.875195 | -0.699815 | 0.00005 | 0.00102476 |
| 881 | SEMA4B | chr15:90728151-90772892 | PDT | ATRA+PDT | 58.9744 | 90.4247 | 0.616629 | 0.00005 | 0.00102476 |
| 882 | SEMA7A | chr15:74701629-74726299 | PDT | ATRA+PDT | 5.13759 | 8.93069 | 0.797679 | 0.00005 | 0.00102476 |
| 883 | SERINC5 | chr5:79407049-79551898 | PDT | ATRA+PDT | 34.2876 | 27.0545 | -0.341818 | 0.00035 | 0.00584867 |
| 884 | SERPINA1 | chr14:94843083-94857029 | PDT | ATRA+PDT | 52.1035 | 35.7555 | -0.543216 | 0.00005 | 0.00102476 |
| 885 | SERPINA3 | chr14:95078713-95090390 | PDT | ATRA+PDT | 0.68675 | 2.43736 | 1.82746 | 0.0001 | 0.00194145 |
| 886 | SERPINA5 | chr14:95047730-95059457 | PDT | ATRA+PDT | 0.689563 | 1.79855 | 1.38308 | 0.00005 | 0.00102476 |
| 887 | SERPINE2 | chr2:224839764-224904036 | PDT | ATRA+PDT | 12.5507 | 8.2725 | -0.601378 | 0.00005 | 0.00102476 |
| 888 | SERPINH1 | chr11:75273100-75283849 | PDT | ATRA+PDT | 64.3244 | 101.369 | 0.656182 | 0.00005 | 0.00102476 |
| 889 | SERTAD2 | chr2:64858754-64881046 | PDT | ATRA+PDT | 33.6969 | 42.7119 | 0.342025 | 0.0004 | 0.00650107 |
| 890 | SESN2 | chr1:28585962-28609002 | PDT | ATRA+PDT | 31.424 | 51.1889 | 0.703964 | 0.00005 | 0.00102476 |
| 891 | SEZ6L2 | chr16:29882479-29910585 | PDT | ATRA+PDT | 3.51491 | 4.63187 | 0.398107 | 0.002 | 0.0229388 |
| 892 | SFTA2 | chr6_qbl_hap6:2192097-219292 | PDT | ATRA+PDT | 18.8994 | 11.8409 | -0.674553 | 0.0021 | 0.0238805 |
| 893 | SGK1 | chr6:134490383-134639196 | PDT | ATRA+PDT | 13.7474 | 10.7069 | -0.360616 | 0.0014 | 0.0175049 |
| 894 | SH2B3 | chr12:111843751-111889427 | PDT | ATRA+PDT | 3.18564 | 4.18057 | 0.392119 | 0.00115 | 0.0151042 |
| 895 | SH2D3A | chr19:6752172-6767523 | PDT | ATRA+PDT | 14.9455 | 19.5506 | 0.3875 | 0.00045 | 0.00719326 |
| 896 | SH3BGRL2 | chr6:80340999-80413369 | PDT | ATRA+PDT | 22.0365 | 15.048 | -0.550319 | 0.00005 | 0.00102476 |
| 897 | SH3RF2 | chr5:145316125-145442879 | PDT | ATRA+PDT | 8.67702 | 6.11854 | -0.504012 | 0.00005 | 0.00102476 |
| 898 | SHCBP1 | chr16:46614467-46655311 | PDT | ATRA+PDT | 9.33162 | 7.17744 | -0.378658 | 0.001 | 0.0135165 |
| 899 | SHROOM1 | chr5:132157832-132166590 | PDT | ATRA+PDT | 6.71456 | 9.36809 | 0.480463 | 0.00005 | 0.00102476 |
| 900 | SIGMAR1 | chr9:34634718-34637768 | PDT | ATRA+PDT | 97.8308 | 78.3167 | -0.320969 | 0.00115 | 0.0151042 |
| 901 | SIPA1L1 | chr14:71996041-72206120 | PDT | ATRA+PDT | 12.1664 | 15.2208 | 0.323133 | 0.0013 | 0.0166643 |
| 902 | SIPA1L2 | chr1:232533711-232651243 | PDT | ATRA+PDT | 6.69335 | 4.95696 | -0.433271 | 0.0002 | 0.00358322 |
| 903 | SIX4 | chr14:61176255-61190852 | PDT | ATRA+PDT | 1.17064 | 1.96518 | 0.747368 | 0.00005 | 0.00102476 |
| 904 | SKAP1 | chr17:46210801-46507594 | PDT | ATRA+PDT | 3.27684 | 4.95002 | 0.595128 | 0.00085 | 0.0119898 |
| 905 | SKAP2 | chr7:26706687-26904341 | PDT | ATRA+PDT | 19.9257 | 29.4178 | 0.562056 | 0.00005 | 0.00102476 |
| 906 | SLC12A4 | chr16:67973786-68002597 | PDT | ATRA+PDT | 3.92501 | 6.3475 | 0.693492 | 0.00005 | 0.00102476 |
| 907 | SLC16A1 | chr1:113454469-113498975 | PDT | ATRA+PDT | 49.6228 | 38.2808 | -0.374383 | 0.0002 | 0.00358322 |
| 908 | SLC16A3 | chr17:80186281-80197375 | PDT | ATRA+PDT | 14.5481 | 18.1504 | 0.319175 | 0.00425 | 0.0420979 |
| 909 | SLC17A9 | chr20:61583998-61599949 | PDT | ATRA+PDT | 37.6118 | 30.334 | -0.310248 | 0.00245 | 0.0267496 |
| 910 | SLC22A18 | chr11:2909326-2946476 | PDT | ATRA+PDT | 18.2722 | 13.6659 | -0.419075 | 0.00095 | 0.0131061 |
| 911 | SLC22A23 | chr6:3269206-3456793 | PDT | ATRA+PDT | 1.72442 | 0.932151 | -0.887479 | 0.00005 | 0.00102476 |
| 912 | SLC22A5 | chr5:131630144-131731306 | PDT | ATRA+PDT | 1.53996 | 2.33011 | 0.59751 | 0.00075 | 0.0109256 |
| 913 | SLC23A3 | chr2:220026180-220034817 | PDT | ATRA+PDT | 2.36227 | 1.29953 | -0.862185 | 0.0004 | 0.00650107 |
| 914 | SLC24A1 | chr15:65914269-65948598 | PDT | ATRA+PDT | 0.97534 | 1.44064 | 0.562731 | 0.0007 | 0.0103814 |
| 915 | SLC25A28 | chr10:101370274-101380221 | PDT | ATRA+PDT | 13.5257 | 10.0388 | -0.430107 | 0.0009 | 0.0125788 |
| 916 | SLC25A39 | chr17:42396992-42402217 | PDT | ATRA+PDT | 119.986 | 95.6822 | -0.326543 | 0.0007 | 0.0103814 |
| 917 | SLC26A6 | chr3:48663155-48672926 | PDT | ATRA+PDT | 4.11243 | 3.02687 | -0.442163 | 0.0014 | 0.0175049 |
| 918 | SLC28A3 | chr9:86890764-86983413 | PDT | ATRA+PDT | 1.35204 | 1.95367 | 0.531051 | 0.00075 | 0.0109256 |

### Table S1: Significant differentially expressed genes (DEGs) after ATRA+PDT compared to PDT.

| # | gene | locus | sample_1 | sample_2 | value_1 | value_2 | log2(fold_change) | p_value | q_value |
|---|---|---|---|---|---|---|---|---|---|
| 919 | SLC2A12 | chr6:134308718-134373789 | PDT | ATRA+PDT | 0.608026 | 1.3385 | 1.13841 | 0.00005 | 0.00102476 |
| 920 | SLC30A1 | chr1:211748380-211752099 | PDT | ATRA+PDT | 50.3641 | 62.676 | 0.315518 | 0.00175 | 0.0207397 |
| 921 | SLC35E4 | chr22:31031792-31043862 | PDT | ATRA+PDT | 4.46778 | 3.03284 | -0.558887 | 0.00015 | 0.00281494 |
| 922 | SLC38A5 | chrX:48316926-48328644 | PDT | ATRA+PDT | 13.129 | 7.8112 | -0.749135 | 0.00005 | 0.00102476 |
| 923 | SLC39A10 | chr2:196521531-196602426 | PDT | ATRA+PDT | 6.11085 | 4.30522 | -0.505288 | 0.00005 | 0.00102476 |
| 924 | SLC39A8 | chr4:103172197-103266655 | PDT | ATRA+PDT | 9.63548 | 5.401 | -0.83513 | 0.00005 | 0.00102476 |
| 925 | SLC4A4 | chr4:72053002-72437804 | PDT | ATRA+PDT | 2.03784 | 3.17406 | 0.639291 | 0.00005 | 0.00102476 |
| 926 | SLC7A7 | chr14:23242431-23289020 | PDT | ATRA+PDT | 4.55966 | 0.977627 | -2.22157 | 0.00005 | 0.00102476 |
| 927 | SLC9A7 | chrX:46466372-46618472 | PDT | ATRA+PDT | 4.74234 | 7.44846 | 0.651342 | 0.00005 | 0.00102476 |
| 928 | SLCO4A1 | chr20:61273796-61303647 | PDT | ATRA+PDT | 26.2854 | 20.3592 | -0.368578 | 0.0003 | 0.00511725 |
| 929 | SLFN5 | chr17:33570085-33594761 | PDT | ATRA+PDT | 2.51404 | 3.53672 | 0.492403 | 0.0002 | 0.00358322 |
| 930 | SMAD3 | chr15:67358194-67487533 | PDT | ATRA+PDT | 59.5952 | 83.9935 | 0.49508 | 0.00005 | 0.00102476 |
| 931 | SMARCA1 | chrX:128580477-128657460 | PDT | ATRA+PDT | 9.95072 | 12.3985 | 0.317295 | 0.0026 | 0.0281856 |
| 932 | SMCHD1 | chr18:2655885-2805015 | PDT | ATRA+PDT | 21.4244 | 26.7571 | 0.320665 | 0.0007 | 0.0103814 |
| 933 | SMPD3 | chr16:68392229-68482409 | PDT | ATRA+PDT | 2.86222 | 6.1399 | 1.10108 | 0.00005 | 0.00102476 |
| 934 | SMURF1 | chr7:98625057-98741743 | PDT | ATRA+PDT | 44.3285 | 53.9284 | 0.282811 | 0.00475 | 0.045818 |
| 935 | SNAP23 | chr15:42787503-42825259 | PDT | ATRA+PDT | 40.841 | 50.8188 | 0.315344 | 0.00155 | 0.0189586 |
| 936 | SNCG | chr10:88718287-88723017 | PDT | ATRA+PDT | 97.5439 | 65.8508 | -0.566852 | 0.00005 | 0.00102476 |
| 937 | SNTB1 | chr8:121547984-121824309 | PDT | ATRA+PDT | 32.41 | 25.4212 | -0.350408 | 0.0004 | 0.00650107 |
| 938 | SORD | chr15:45315301-45367287 | PDT | ATRA+PDT | 55.6577 | 45.7389 | -0.283161 | 0.00425 | 0.0420979 |
| 939 | SORL1 | chr11:121322911-121504471 | PDT | ATRA+PDT | 14.4875 | 11.7749 | -0.299093 | 0.0024 | 0.0263926 |
| 940 | SOSTDC1 | chr7:16501105-16505474 | PDT | ATRA+PDT | 1.20435 | 0.404531 | -1.57393 | 0.0015 | 0.0184954 |
| 941 | SOX12 | chr20:306238-310867 | PDT | ATRA+PDT | 1.2375 | 0.834728 | -0.568053 | 0.00315 | 0.0333027 |
| 942 | SOX4 | chr6:21593971-21598849 | PDT | ATRA+PDT | 29.6801 | 40.2276 | 0.438686 | 0.00005 | 0.00102476 |
| 943 | SP100 | chr2:231280870-231410317 | PDT | ATRA+PDT | 15.6481 | 20.7536 | 0.407377 | 0.00035 | 0.00584867 |
| 944 | SP110 | chr2:231033644-231090444 | PDT | ATRA+PDT | 3.29061 | 4.76151 | 0.533064 | 0.0002 | 0.00358322 |
| 945 | SP140L | chr2:231191893-231268445 | PDT | ATRA+PDT | 4.7695 | 7.90811 | 0.729495 | 0.00005 | 0.00102476 |
| 946 | SPCS3 | chr4:177241089-177253396 | PDT | ATRA+PDT | 26.9071 | 20.9101 | -0.363784 | 0.00015 | 0.00281494 |
| 947 | SPDEF | chr6:34505578-34524110 | PDT | ATRA+PDT | 16.5058 | 8.19596 | -1.00999 | 0.00005 | 0.00102476 |
| 948 | SPHK2 | chr19:49122547-49133663 | PDT | ATRA+PDT | 7.70829 | 9.8477 | 0.353375 | 0.00385 | 0.0390041 |
| 949 | SPINK1 | chr5:147204142-147211260 | PDT | ATRA+PDT | 106.5 | 158.725 | 0.575683 | 0.00005 | 0.00102476 |
| 950 | SPINK4 | chr9:33240195-33248565 | PDT | ATRA+PDT | 5.51131 | 0 | -inf | 0.00005 | 0.00102476 |
| 951 | SQLE | chr8:126010719-126034525 | PDT | ATRA+PDT | 278.875 | 208.842 | -0.417209 | 0.00005 | 0.00102476 |
| 952 | SRD5A3 | chr4:56212387-56251747 | PDT | ATRA+PDT | 1.50107 | 2.06191 | 0.457987 | 0.0035 | 0.0363197 |
| 953 | SRI | chr7:87834431-87856308 | PDT | ATRA+PDT | 43.9469 | 59.8828 | 0.446382 | 0.00005 | 0.00102476 |
| 954 | SRPRB | chr3:133502876-133540336 | PDT | ATRA+PDT | 34.7247 | 28.4416 | -0.287961 | 0.00425 | 0.0420979 |
| 955 | SSFA2 | chr2:182756471-182795464 | PDT | ATRA+PDT | 79.3028 | 63.3638 | -0.323715 | 0.0009 | 0.0125788 |
| 956 | SSR1 | chr6:7281287-7313541 | PDT | ATRA+PDT | 18.5387 | 15.3097 | -0.276093 | 0.00515 | 0.0488381 |
| 957 | ST3GAL2 | chr16:70413337-70472991 | PDT | ATRA+PDT | 2.67229 | 1.86208 | -0.521164 | 0.00005 | 0.00102476 |
| 958 | ST3GAL4 | chr11:126225539-126284536 | PDT | ATRA+PDT | 83.6796 | 64.7482 | -0.370035 | 0.0001 | 0.00194145 |
| 959 | ST6GALNAC1 | chr17:74620844-74639894 | PDT | ATRA+PDT | 1.73672 | 0.466286 | -1.89708 | 0.00005 | 0.00102476 |
| 960 | ST6GALNAC4 | chr9:130670164-130679305 | PDT | ATRA+PDT | 9.82944 | 6.35071 | -0.630191 | 0.00005 | 0.00102476 |
| 961 | STAG3L4 | chr7:66767624-66786513 | PDT | ATRA+PDT | 4.26266 | 6.58727 | 0.627925 | 0.00005 | 0.00102476 |
| 962 | STARD10 | chr11:72465773-72504750 | PDT | ATRA+PDT | 89.0213 | 63.1855 | -0.494558 | 0.00005 | 0.00102476 |
| 963 | STEAP1 | chr7:89783688-89794141 | PDT | ATRA+PDT | 4.94591 | 9.43399 | 0.931631 | 0.00005 | 0.00102476 |
| 964 | STEAP2 | chr7:89840999-89866992 | PDT | ATRA+PDT | 2.36953 | 4.0594 | 0.776666 | 0.0003 | 0.00511725 |
| 965 | STEAP3 | chr2:119981383-120023227 | PDT | ATRA+PDT | 8.3467 | 6.32523 | -0.400089 | 0.00035 | 0.00584867 |
| 966 | STEAP4 | chr7:87905743-87936228 | PDT | ATRA+PDT | 1.91198 | 2.73698 | 0.517518 | 0.00005 | 0.00102476 |
| 967 | STK17A | chr7:43622691-43666978 | PDT | ATRA+PDT | 11.1558 | 15.412 | 0.466257 | 0.00005 | 0.00102476 |
| 968 | STK32C | chr10:134020995-134121477 | PDT | ATRA+PDT | 4.70709 | 3.51555 | -0.421086 | 0.0037 | 0.0379161 |
| 969 | STK39 | chr2:168810529-169104105 | PDT | ATRA+PDT | 62.3222 | 86.5812 | 0.474308 | 0.00005 | 0.00102476 |
| 970 | STMN1 | chr1:26210676-26233368 | PDT | ATRA+PDT | 170.312 | 210.954 | 0.308745 | 0.00145 | 0.0179618 |
| 971 | STOX1 | chr10:70587293-70655209 | PDT | ATRA+PDT | 0.733568 | 1.32757 | 0.855784 | 0.0024 | 0.0263926 |
| 972 | STRA6 | chr15:74471807-74502046 | PDT | ATRA+PDT | 0.553158 | 1.47588 | 1.41581 | 0.00005 | 0.00102476 |
| 973 | STX1A | chr7:73113534-73134017 | PDT | ATRA+PDT | 24.2662 | 30.793 | 0.343655 | 0.00085 | 0.0119898 |
| 974 | SUCLA2 | chr13:48516790-48575462 | PDT | ATRA+PDT | 29.5879 | 23.5821 | -0.327312 | 0.00205 | 0.0234618 |
| 975 | SULT2B1 | chr19:49055428-49102684 | PDT | ATRA+PDT | 30.4892 | 18.3777 | -0.730343 | 0.00005 | 0.00102476 |
| 976 | SUV420H2 | chr19:55851220-55859489 | PDT | ATRA+PDT | 4.81079 | 6.42984 | 0.418508 | 0.0012 | 0.0156263 |
| 977 | SVIL | chr10:29746276-30024730 | PDT | ATRA+PDT | 4.67726 | 6.41431 | 0.45563 | 0.00005 | 0.00102476 |
| 978 | SWAP70 | chr11:9685627-9774507 | PDT | ATRA+PDT | 10.2031 | 13.3352 | 0.386231 | 0.0004 | 0.00650107 |
| 979 | SYT7 | chr11:61281187-61348344 | PDT | ATRA+PDT | 2.19574 | 1.45466 | -0.59402 | 0.00005 | 0.00102476 |
| 980 | SYTL2 | chr11:85405264-85522178 | PDT | ATRA+PDT | 6.81333 | 9.03548 | 0.407242 | 0.00035 | 0.00584867 |
| 981 | SYTL5 | chrX:37865834-37988073 | PDT | ATRA+PDT | 1.21634 | 0.704342 | -0.788197 | 0.0003 | 0.00511725 |
| 982 | TAF1A | chr1:222731243-222763275 | PDT | ATRA+PDT | 11.8473 | 16.7531 | 0.499865 | 0.00005 | 0.00102476 |
| 983 | TAF1D | chr11:93469095-93474703 | PDT | ATRA+PDT | 74.4728 | 92.6756 | 0.315475 | 0.00195 | 0.0225592 |
| 984 | TAP2 | chr6_ssto_hap7:4220532-423731 | PDT | ATRA+PDT | 8.33497 | 15.5791 | 0.902366 | 0.00005 | 0.00102476 |
| 985 | TAPBP | chr6_qbl_hap6:4499711-451441( | PDT | ATRA+PDT | 17.4531 | 27.3118 | 0.646043 | 0.00005 | 0.00102476 |
| 986 | TAPBPL | chr12:6561176-6579843 | PDT | ATRA+PDT | 0.508899 | 1.72817 | 1.7638 | 0.00035 | 0.00584867 |
| 987 | TBC1D1 | chr4:37892704-38140796 | PDT | ATRA+PDT | 10.7113 | 14.0503 | 0.391464 | 0.00025 | 0.00439786 |
| 988 | TBC1D2 | chr9:100961279-101018003 | PDT | ATRA+PDT | 9.12037 | 6.95428 | -0.391191 | 0.0011 | 0.0146092 |
| 989 | TBC1D4 | chr13:75858808-76056250 | PDT | ATRA+PDT | 15.7906 | 12.6676 | -0.317923 | 0.00105 | 0.0140676 |
| 990 | TBC1D9 | chr4:141541935-141677471 | PDT | ATRA+PDT | 4.7026 | 6.63498 | 0.496634 | 0.00005 | 0.00102476 |
| 991 | TBL1X | chrX:9431334-9687780 | PDT | ATRA+PDT | 9.27934 | 5.89633 | -0.654204 | 0.00005 | 0.00102476 |
| 992 | TBL3 | chr16:2022063-2028751 | PDT | ATRA+PDT | 21.0564 | 16.4915 | -0.35254 | 0.0013 | 0.0166643 |
| 993 | TBXAS1 | chr7:139478046-139720125 | PDT | ATRA+PDT | 7.8039 | 11.8389 | 0.601267 | 0.00005 | 0.00102476 |
| 994 | TCEAL1 | chrX:102883647-102885876 | PDT | ATRA+PDT | 1.55868 | 2.89536 | 0.893418 | 0.00215 | 0.0243712 |
| 995 | TCN2 | chr22:31003069-31023047 | PDT | ATRA+PDT | 4.27892 | 2.4986 | -0.776127 | 0.00005 | 0.00102476 |
| 996 | TCP11L2 | chr12:106696580-106740792 | PDT | ATRA+PDT | 7.1208 | 10.0693 | 0.499859 | 0.0002 | 0.00358322 |
| 997 | TCTN2 | chr12:124155659-124192950 | PDT | ATRA+PDT | 2.0913 | 3.06846 | 0.553113 | 0.00085 | 0.0119898 |
| 998 | TEAD2 | chr19:49838676-49865714 | PDT | ATRA+PDT | 13.329 | 26.0582 | 0.967171 | 0.00005 | 0.00102476 |
| 999 | TFF3 | chr21:43731776-43735706 | PDT | ATRA+PDT | 6.41603 | 0.37653 | -4.09084 | 0.0026 | 0.0281856 |
| 1000 | TFPI | chr2:188328957-188419219 | PDT | ATRA+PDT | 74.811 | 105.917 | 0.501613 | 0.00005 | 0.00102476 |
| 1001 | TGFB2 | chr1:218517537-218617961 | PDT | ATRA+PDT | 2.41102 | 4.22929 | 0.810768 | 0.00005 | 0.00102476 |
| 1002 | TGFBR2 | chr3:30647993-30735633 | PDT | ATRA+PDT | 27.0082 | 18.9255 | -0.513065 | 0.00005 | 0.00102476 |
| 1003 | TGM2 | chr20:36756863-36793700 | PDT | ATRA+PDT | 10.6596 | 17.9277 | 0.75004 | 0.00005 | 0.00102476 |
| 1004 | TIA1 | chr2:70436575-70475779 | PDT | ATRA+PDT | 14.983 | 19.6966 | 0.394623 | 0.00015 | 0.00281494 |
| 1005 | TINAGL1 | chr1:32042085-32053287 | PDT | ATRA+PDT | 77.1995 | 103.768 | 0.426698 | 0.00005 | 0.00102476 |
| 1006 | TIPARP | chr3:156390959-156424557 | PDT | ATRA+PDT | 74.905 | 118.938 | 0.667071 | 0.00005 | 0.00102476 |
| 1007 | TJP1 | chr15:29992356-30114706 | PDT | ATRA+PDT | 49.6946 | 60.8908 | 0.293135 | 0.00285 | 0.0305545 |
| 1008 | TJP2 | chr9:71736179-71870124 | PDT | ATRA+PDT | 79.8813 | 100.228 | 0.327349 | 0.00095 | 0.0131061 |
| 1009 | TJP3 | chr19:3728373-3750682 | PDT | ATRA+PDT | 4.72629 | 7.85532 | 0.732962 | 0.00005 | 0.00102476 |
| 1010 | TK2 | chr16:66541905-66584315 | PDT | ATRA+PDT | 2.36491 | 3.59582 | 0.604535 | 0.00005 | 0.00102476 |
| 1011 | TM4SF4 | chr3:149192367-149221181 | PDT | ATRA+PDT | 312.173 | 251.628 | -0.311053 | 0.0014 | 0.0175049 |
| 1012 | TMC5 | chr16:19422056-19510434 | PDT | ATRA+PDT | 18.8617 | 28.4905 | 0.595021 | 0.00005 | 0.00102476 |
| 1013 | TMED1 | chr19:10943113-10946983 | PDT | ATRA+PDT | 11.0032 | 8.3463 | -0.398713 | 0.0017 | 0.0202822 |
| 1014 | TMED3 | chr15:79603490-79615189 | PDT | ATRA+PDT | 88.2767 | 67.8847 | -0.378947 | 0.0002 | 0.00358322 |
| 1015 | TMEM141 | chr9:139685776-139687769 | PDT | ATRA+PDT | 114.982 | 93.1895 | -0.303171 | 0.00285 | 0.0305545 |
| 1016 | TMEM147 | chr19:36024313-36038429 | PDT | ATRA+PDT | 60.3456 | 47.624 | -0.341561 | 0.00305 | 0.0323415 |
| 1017 | TMEM43 | chr3:14166439-14185180 | PDT | ATRA+PDT | 21.6461 | 16.6741 | -0.376502 | 0.00015 | 0.00281494 |
| 1018 | TMEM57 | chr1:25757387-25826698 | PDT | ATRA+PDT | 13.4423 | 10.9402 | -0.297135 | 0.0052 | 0.0492248 |
| 1019 | TMPRSS4 | chr11:117886486-117990556 | PDT | ATRA+PDT | 12.7967 | 16.5614 | 0.372054 | 0.00075 | 0.0109256 |
| 1020 | TNFAIP2 | chr14:103592663-103603776 | PDT | ATRA+PDT | 11.2927 | 19.0707 | 0.755967 | 0.00005 | 0.00102476 |

**Table S1: Significant differentially expressed genes (DEGs) after ATRA+PDT compared to PDT.**

| # | gene | locus | sample_1 | sample_2 | value_1 | value_2 | log2(fold_change) | p_value | q_value |
|---|---|---|---|---|---|---|---|---|---|
| 1021 | TNFRSF12A | chr16:3070312-3072383 | PDT | ATRA+PDT | 260.635 | 317.793 | 0.286056 | 0.003 | 0.0318746 |
| 1022 | TNK2 | chr3:195590235-195635880 | PDT | ATRA+PDT | 17.786 | 22.1477 | 0.316414 | 0.0023 | 0.0255827 |
| 1023 | TOB1 | chr17:48939586-48945732 | PDT | ATRA+PDT | 47.7377 | 35.9599 | -0.40874 | 0.00005 | 0.00102476 |
| 1024 | TP53I11 | chr11:44785975-44972608 | PDT | ATRA+PDT | 26.3408 | 41.2581 | 0.647376 | 0.00005 | 0.00102476 |
| 1025 | TP53INP2 | chr20:33292147-33301237 | PDT | ATRA+PDT | 2.06393 | 2.71965 | 0.398031 | 0.0038 | 0.0386074 |
| 1026 | TPPP | chr5:659976-693510 | PDT | ATRA+PDT | 4.12975 | 2.4613 | -0.746634 | 0.00005 | 0.00102476 |
| 1027 | TRAF4 | chr17:27071022-27077976 | PDT | ATRA+PDT | 44.8849 | 57.8295 | 0.365577 | 0.0002 | 0.00358322 |
| 1028 | TRAPPC6A | chr19:45666185-45681485 | PDT | ATRA+PDT | 16.4438 | 10.1838 | -0.691265 | 0.00005 | 0.00102476 |
| 1029 | TRIM21 | chr11:4406126-4414926 | PDT | ATRA+PDT | 4.84589 | 7.53851 | 0.637519 | 0.00005 | 0.00102476 |
| 1030 | TRIM25 | chr17:54965269-54991409 | PDT | ATRA+PDT | 26.2805 | 33.0008 | 0.32851 | 0.00105 | 0.0140676 |
| 1031 | TRIM29 | chr11:119981993-120008863 | PDT | ATRA+PDT | 25.5675 | 18.1135 | -0.497249 | 0.00005 | 0.00102476 |
| 1032 | TRIM3 | chr11:6469842-6495689 | PDT | ATRA+PDT | 2.8097 | 1.83959 | -0.611033 | 0.0004 | 0.00650107 |
| 1033 | TRIM31 | chr6_ssto_hap7:1401126-141131 | PDT | ATRA+PDT | 23.1248 | 76.6326 | 1.72852 | 0.00005 | 0.00102476 |
| 1034 | TRIM38 | chr6:25963070-25985352 | PDT | ATRA+PDT | 2.49065 | 4.52013 | 0.859841 | 0.00005 | 0.00102476 |
| 1035 | TRIM56 | chr7:100728785-100733889 | PDT | ATRA+PDT | 8.56345 | 11.6197 | 0.440307 | 0.00005 | 0.00102476 |
| 1036 | TRIM8 | chr10:104404251-104418076 | PDT | ATRA+PDT | 18.8275 | 23.8045 | 0.338397 | 0.00125 | 0.0161202 |
| 1037 | TRIP6 | chr7:100464949-100471076 | PDT | ATRA+PDT | 4.77133 | 6.54758 | 0.456571 | 0.0013 | 0.0166643 |
| 1038 | TRMT6 | chr20:5918485-5931173 | PDT | ATRA+PDT | 25.4935 | 20.6174 | -0.306271 | 0.0037 | 0.0379161 |
| 1039 | TRNP1 | chr1:27320194-27327377 | PDT | ATRA+PDT | 94.1771 | 116.26 | 0.303907 | 0.00195 | 0.0225592 |
| 1040 | TRPS1 | chr8:116420723-116681228 | PDT | ATRA+PDT | 7.50335 | 10.0167 | 0.416798 | 0.00005 | 0.00102476 |
| 1041 | TSC22D1 | chr13:45006278-45154568 | PDT | ATRA+PDT | 68.9779 | 53.2376 | -0.373689 | 0.0001 | 0.00194145 |
| 1042 | TSC22D2 | chr3:150126787-150177615 | PDT | ATRA+PDT | 22.2714 | 29.6708 | 0.413853 | 0.00005 | 0.00102476 |
| 1043 | TSC22D3 | chrX:106956451-107019017 | PDT | ATRA+PDT | 10.6117 | 20.1811 | 0.927348 | 0.00005 | 0.00102476 |
| 1044 | TSFM | chr12:58176527-58209852 | PDT | ATRA+PDT | 30.5172 | 24.5388 | -0.314557 | 0.00515 | 0.0488381 |
| 1045 | TSPAN13 | chr7:16793350-16824161 | PDT | ATRA+PDT | 57.6264 | 41.0611 | -0.488958 | 0.00005 | 0.00102476 |
| 1046 | TSPAN4 | chr11:842823-867116 | PDT | ATRA+PDT | 9.36181 | 5.15974 | -0.85949 | 0.00005 | 0.00102476 |
| 1047 | TSPAN6 | chrX:99883794-99891794 | PDT | ATRA+PDT | 15.0487 | 18.9021 | 0.328911 | 0.0036 | 0.0371409 |
| 1048 | TSPAN8 | chr12:71518876-71551779 | PDT | ATRA+PDT | 409.565 | 337.399 | -0.279638 | 0.005 | 0.0477977 |
| 1049 | TST | chr22:37406905-37415491 | PDT | ATRA+PDT | 32.4685 | 25.9519 | -0.3232 | 0.00485 | 0.0466141 |
| 1050 | TSTA3 | chr8:144694787-144699732 | PDT | ATRA+PDT | 50.2639 | 37.6337 | -0.417498 | 0.00005 | 0.00102476 |
| 1051 | TTC39A | chr1:51752929-51810785 | PDT | ATRA+PDT | 4.78511 | 3.21035 | -0.575822 | 0.0001 | 0.00194145 |
| 1052 | TUBA4A | chr2:220110191-220136910 | PDT | ATRA+PDT | 79.3541 | 57.7594 | -0.45825 | 0.00005 | 0.00102476 |
| 1053 | TUBB2A | chr6:3153901-3157783 | PDT | ATRA+PDT | 9.30193 | 7.19573 | -0.370388 | 0.0036 | 0.0371409 |
| 1054 | TUBGCP6 | chr22:50656117-50683400 | PDT | ATRA+PDT | 3.25603 | 2.5207 | -0.369291 | 0.00375 | 0.0383182 |
| 1055 | TUFT1 | chr1:151512780-151556059 | PDT | ATRA+PDT | 18.614 | 25.7644 | 0.46899 | 0.00005 | 0.00102476 |
| 1056 | TXK | chr4:48068409-48136273 | PDT | ATRA+PDT | 0.361822 | 1.3053 | 1.85103 | 0.00005 | 0.00102476 |
| 1057 | TXNDC5 | chr6:7727010-8102828 | PDT | ATRA+PDT | 130.123 | 104.162 | -0.321045 | 0.0047 | 0.0455001 |
| 1058 | TXNIP | chr1:145438461-145442628 | PDT | ATRA+PDT | 51.5463 | 70.4953 | 0.451659 | 0.00005 | 0.00102476 |
| 1059 | UBC | chr12:125396191-125399587 | PDT | ATRA+PDT | 831.594 | 1063.52 | 0.354898 | 0.0019 | 0.0221245 |
| 1060 | UBXN11 | chr1:26608772-26647014 | PDT | ATRA+PDT | 8.10727 | 10.8199 | 0.416395 | 0.00215 | 0.0243712 |
| 1061 | UCA1 | chr19:15939756-15946230 | PDT | ATRA+PDT | 10.7972 | 17.4755 | 0.694673 | 0.00005 | 0.00102476 |
| 1062 | UGCG | chr9:114659205-114695433 | PDT | ATRA+PDT | 47.1696 | 59.2197 | 0.328219 | 0.0011 | 0.0146092 |
| 1063 | UGT1A10 | chr2:234526290-234681951 | PDT | ATRA+PDT | 160.905 | 214.324 | 0.413587 | 0.00155 | 0.0189586 |
| 1064 | UGT8 | chr4:115519610-115598202 | PDT | ATRA+PDT | 21.5381 | 17.0898 | -0.333753 | 0.0008 | 0.0114972 |
| 1065 | UNC93B1 | chr11:67758574-67771593 | PDT | ATRA+PDT | 10.2936 | 13.4578 | 0.386694 | 0.00095 | 0.0131061 |
| 1066 | UPK3B | chr7:76139744-76157199 | PDT | ATRA+PDT | 12.5212 | 7.33083 | -0.772327 | 0.00005 | 0.00102476 |
| 1067 | UQCRFS1 | chr19:29698166-29704136 | PDT | ATRA+PDT | 84.513 | 67.3338 | -0.327843 | 0.001 | 0.0135165 |
| 1068 | UQCRHL | chr1:16133656-16134194 | PDT | ATRA+PDT | 29.9609 | 22.5527 | -0.409778 | 0.004 | 0.0401429 |
| 1069 | URB2 | chr1:229761980-229795946 | PDT | ATRA+PDT | 12.5981 | 9.88058 | -0.350534 | 0.0006 | 0.00910057 |
| 1070 | USH1C | chr11:17515441-17565963 | PDT | ATRA+PDT | 22.9992 | 16.7359 | -0.458641 | 0.00005 | 0.00102476 |
| 1071 | USMG5 | chr10:105127723-105156270 | PDT | ATRA+PDT | 401.557 | 316.853 | -0.34179 | 0.003 | 0.0318746 |
| 1072 | USP11 | chrX:47092313-47107727 | PDT | ATRA+PDT | 12.1379 | 15.491 | 0.351919 | 0.001 | 0.0135165 |
| 1073 | USP53 | chr4:120133781-120216673 | PDT | ATRA+PDT | 30.8233 | 40.165 | 0.381915 | 0.0002 | 0.00358322 |
| 1074 | VAMP5 | chr2:85811530-85820511 | PDT | ATRA+PDT | 3.04146 | 7.92904 | 1.38238 | 0.00005 | 0.00102476 |
| 1075 | VDAC1 | chr5:133307565-133340824 | PDT | ATRA+PDT | 108.093 | 85.0046 | -0.346664 | 0.0004 | 0.00650107 |
| 1076 | VDR | chr12:48235319-48298814 | PDT | ATRA+PDT | 6.92137 | 4.76545 | -0.538444 | 0.00005 | 0.00102476 |
| 1077 | VEZF1 | chr17:56048909-56065615 | PDT | ATRA+PDT | 20.2986 | 25.483 | 0.328156 | 0.00095 | 0.0131061 |
| 1078 | VIL1 | chr2:219283837-219314248 | PDT | ATRA+PDT | 31.2981 | 16.9876 | -0.881593 | 0.00005 | 0.00102476 |
| 1079 | VMA21 | chrX:150565656-150577836 | PDT | ATRA+PDT | 18.8352 | 15.2752 | -0.302238 | 0.0022 | 0.024728 |
| 1080 | VSIG10L | chr19:51834794-51845378 | PDT | ATRA+PDT | 7.17934 | 10.1251 | 0.49601 | 0.00005 | 0.00102476 |
| 1081 | VWA1 | chr1:1370902-1378262 | PDT | ATRA+PDT | 11.894 | 8.45271 | -0.492746 | 0.00005 | 0.00102476 |
| 1082 | VWA2 | chr10:115999017-116049751 | PDT | ATRA+PDT | 1.92412 | 3.07645 | 0.677069 | 0.00005 | 0.00102476 |
| 1083 | WARS | chr14:100800124-100842680 | PDT | ATRA+PDT | 48.7871 | 59.4294 | 0.284679 | 0.00405 | 0.0405304 |
| 1084 | WBP5 | chrX:102611379-102613397 | PDT | ATRA+PDT | 13.9992 | 18.3838 | 0.393092 | 0.00165 | 0.0198632 |
| 1085 | WDR19 | chr4:39184023-39287430 | PDT | ATRA+PDT | 2.8049 | 3.69424 | 0.397329 | 0.00245 | 0.0267496 |
| 1086 | WFDC2 | chr20:44098393-44110172 | PDT | ATRA+PDT | 2.28187 | 6.53873 | 1.51879 | 0.0021 | 0.0238805 |
| 1087 | WHAMM | chr15:83477972-83503613 | PDT | ATRA+PDT | 5.53372 | 6.93922 | 0.326523 | 0.0048 | 0.0461751 |
| 1088 | WNK4 | chr17:40932648-40949084 | PDT | ATRA+PDT | 1.35766 | 0.74025 | -0.875036 | 0.00005 | 0.00102476 |
| 1089 | WNT11 | chr11:75897369-75917574 | PDT | ATRA+PDT | 4.1994 | 1.77853 | -1.2395 | 0.00005 | 0.00102476 |
| 1090 | WWC3 | chrX:9983794-10112518 | PDT | ATRA+PDT | 1.36101 | 2.01822 | 0.568405 | 0.00005 | 0.00102476 |
| 1091 | XPA | chr9:100437190-100459691 | PDT | ATRA+PDT | 7.37596 | 5.13513 | -0.522429 | 0.00045 | 0.00719326 |
| 1092 | ZAK | chr2:173940564-174146764 | PDT | ATRA+PDT | 21.2333 | 15.5073 | -0.453377 | 0.00005 | 0.00102476 |
| 1093 | ZBTB10 | chr8:81398447-81434610 | PDT | ATRA+PDT | 1.3336 | 2.32843 | 0.804026 | 0.00005 | 0.00102476 |
| 1094 | ZBTB4 | chr17:7362684-7387568 | PDT | ATRA+PDT | 2.95671 | 4.5059 | 0.60782 | 0.00005 | 0.00102476 |
| 1095 | ZC3H12A | chr1:37940118-37949978 | PDT | ATRA+PDT | 14.3521 | 18.3946 | 0.358025 | 0.00105 | 0.0140676 |
| 1096 | ZC3H6 | chr2:113033177-113097640 | PDT | ATRA+PDT | 1.08125 | 1.53073 | 0.501519 | 0.00015 | 0.00281494 |
| 1097 | ZC3HAV1 | chr7:138728265-138794465 | PDT | ATRA+PDT | 26.4315 | 36.6715 | 0.472401 | 0.00005 | 0.00102476 |
| 1098 | ZC3HAV1L | chr7:138710451-138720775 | PDT | ATRA+PDT | 3.21045 | 1.95516 | -0.715491 | 0.001 | 0.0135165 |
| 1099 | ZDHHC23 | chr3:113666747-113681827 | PDT | ATRA+PDT | 6.00029 | 4.23378 | -0.503085 | 0.00005 | 0.00102476 |
| 1100 | ZFAND2A | chr7:1192542-1199855 | PDT | ATRA+PDT | 98.4655 | 139.074 | 0.498159 | 0.00005 | 0.00102476 |
| 1101 | ZFHX3 | chr16:72816785-73092534 | PDT | ATRA+PDT | 2.83608 | 3.50976 | 0.307476 | 0.0051 | 0.048623 |
| 1102 | ZFP36 | chr19:39897486-39900045 | PDT | ATRA+PDT | 69.397 | 85.3345 | 0.298256 | 0.0029 | 0.0310283 |
| 1103 | ZFP36L1 | chr14:69254371-69262960 | PDT | ATRA+PDT | 30.635 | 39.7444 | 0.375571 | 0.00005 | 0.00102476 |
| 1104 | ZHX2 | chr8:123793900-123986755 | PDT | ATRA+PDT | 4.98183 | 7.87996 | 0.661512 | 0.00005 | 0.00102476 |
| 1105 | ZNF143 | chr11:9482511-9550071 | PDT | ATRA+PDT | 10.1289 | 12.9935 | 0.359311 | 0.00165 | 0.0198632 |
| 1106 | ZNF184 | chr6:27418520-27440897 | PDT | ATRA+PDT | 4.11238 | 5.61524 | 0.449374 | 0.0009 | 0.0125788 |
| 1107 | ZNF189 | chr9:104161162-104172942 | PDT | ATRA+PDT | 4.06617 | 5.74544 | 0.498748 | 0.0001 | 0.00194145 |
| 1108 | ZNF195 | chr11:3379156-3400452 | PDT | ATRA+PDT | 19.9948 | 25.1975 | 0.333654 | 0.00115 | 0.0151042 |
| 1109 | ZNF200 | chr16:3272324-3285457 | PDT | ATRA+PDT | 4.65756 | 3.54515 | -0.393726 | 0.0017 | 0.0202822 |
| 1110 | ZNF212 | chr7:148936741-148952700 | PDT | ATRA+PDT | 14.9627 | 18.5768 | 0.31213 | 0.00345 | 0.0359757 |
| 1111 | ZNF213 | chr16:3185056-3192805 | PDT | ATRA+PDT | 2.20527 | 2.92962 | 0.409759 | 0.0037 | 0.0379161 |
| 1112 | ZNF292 | chr6:87865268-87973406 | PDT | ATRA+PDT | 5.00352 | 6.18856 | 0.306661 | 0.003 | 0.0318746 |
| 1113 | ZNF334 | chr20:45129706-45142194 | PDT | ATRA+PDT | 3.43199 | 4.67598 | 0.446221 | 0.0016 | 0.0194588 |
| 1114 | ZNF473 | chr19:50529211-50552031 | PDT | ATRA+PDT | 28.2838 | 35.599 | 0.33186 | 0.001 | 0.0135165 |
| 1115 | ZNF488 | chr10:48355088-48373866 | PDT | ATRA+PDT | 3.47533 | 6.63871 | 0.933755 | 0.00005 | 0.00102476 |
| 1116 | ZNF513 | chr2:27600097-27603611 | PDT | ATRA+PDT | 10.3136 | 13.352 | 0.372507 | 0.002 | 0.0229388 |
| 1117 | ZNF516 | chr18:74069636-74207146 | PDT | ATRA+PDT | 2.15709 | 3.05855 | 0.503761 | 0.00005 | 0.00102476 |
| 1118 | ZNF552 | chr19:58318449-58326281 | PDT | ATRA+PDT | 8.4639 | 12.3848 | 0.549174 | 0.00005 | 0.00102476 |
| 1119 | ZNF609 | chr15:64791618-64978266 | PDT | ATRA+PDT | 26.6047 | 35.7902 | 0.427885 | 0.00005 | 0.00102476 |
| 1120 | ZNF627 | chr19:11708234-11729974 | PDT | ATRA+PDT | 2.83543 | 3.88525 | 0.454441 | 0.0012 | 0.0156263 |
| 1121 | ZNF641 | chr12:48733792-48745029 | PDT | ATRA+PDT | 2.01609 | 2.61535 | 0.375441 | 0.0045 | 0.0439625 |
| 1122 | ZNF674 | chrX:46357159-46404892 | PDT | ATRA+PDT | 1.61444 | 2.52034 | 0.642585 | 0.00005 | 0.00102476 |

**Table S1: Significant differentially expressed genes (DEGs) after ATRA+PDT compared to PDT.**

| # | gene | locus | sample_1 | sample_2 | value_1 | value_2 | log2(fold_change) | p_value | q_value |
|---|---|---|---|---|---|---|---|---|---|
| 1123 | ZNF76 | chr6:35227509-35263760 | PDT | ATRA+PDT | 7.93857 | 10.2398 | 0.367238 | 0.00195 | 0.0225592 |
| 1124 | ZNF764 | chr16:30565084-30569642 | PDT | ATRA+PDT | 1.23965 | 1.95514 | 0.657332 | 0.0021 | 0.0238805 |
| 1125 | ZNF776 | chr19:58258163-58269527 | PDT | ATRA+PDT | 8.13752 | 10.4735 | 0.364081 | 0.0005 | 0.00788626 |
| 1126 | ZNF823 | chr19:11832079-11849760 | PDT | ATRA+PDT | 4.46785 | 6.29258 | 0.494072 | 0.0003 | 0.00511725 |
| 1127 | ZNFX1-AS1 | chr20:47862438-47905795 | PDT | ATRA+PDT | 399.818 | 534.27 | 0.418226 | 0.00295 | 0.0315316 |
| 1128 | ZSWIM6 | chr5:60628099-60841999 | PDT | ATRA+PDT | 9.50869 | 12.4155 | 0.384823 | 0.00005 | 0.00102476 |
| 1129 | ZYX | chr7:143078359-143220540 | PDT | ATRA+PDT | 77.6904 | 119.543 | 0.621723 | 0.00005 | 0.00102476 |
