## Supplementary Table 2 for "All-*trans* retinoic acid enhances the anti-tumour effects of fimaporfin-based photodynamic therapy"

Table S2: Significant DEGs with a |log2(fold change)| of ≥1 after ATRA+PDT compared to PDT.

| gene | locus | sample_1 | sample_2 | value_1 | value_2 | log2(fold_change) | p_value | q_value |
|---|---|---|---|---|---|---|---|---|
| Up-regulated genes | | | | | | | | |
| FAM131B | chr7:143050492-143059840 | PDT | ATRA+PDT | 0.409476 | 3.47658 | 3.08582 | 0.00005 | 0.00102476 |
| RARRES1 | chr3:158414896-158450275 | PDT | ATRA+PDT | 2.68665 | 16.1452 | 2.58723 | 0.00005 | 0.00102476 |
| MDK | chr11:46402617-46405375 | PDT | ATRA+PDT | 4.9587 | 26.8511 | 2.43695 | 0.00005 | 0.00102476 |
| CYP24A1 | chr20:52769987-52790516 | PDT | ATRA+PDT | 0.491874 | 2.60664 | 2.40583 | 0.00005 | 0.00102476 |
| PSMB8 | chr6_ssto_hap7:4239266-4252521 | PDT | ATRA+PDT | 1.01457 | 4.79809 | 2.24159 | 0.00005 | 0.00102476 |
| RARRES3 | chr11:63014620-63330855 | PDT | ATRA+PDT | 7.65704 | 32.1209 | 2.06866 | 0.00005 | 0.00102476 |
| REN | chr1:204123943-204135465 | PDT | ATRA+PDT | 0.636161 | 2.66058 | 2.06428 | 0.00005 | 0.00102476 |
| CTSS | chr1:150702671-150738433 | PDT | ATRA+PDT | 0.43546 | 1.76088 | 2.01569 | 0.00005 | 0.00102476 |
| FGF19 | chr11:69513005-69519106 | PDT | ATRA+PDT | 6.27328 | 25.0306 | 1.9964 | 0.00005 | 0.00102476 |
| TXK | chr4:48068409-48136273 | PDT | ATRA+PDT | 0.361822 | 1.3053 | 1.85103 | 0.00005 | 0.00102476 |
| SERPINA3 | chr14:95078713-95090390 | PDT | ATRA+PDT | 0.68675 | 2.43736 | 1.82746 | 0.0001 | 0.00194145 |
| ALDH1L1 | chr3:125822407-125899485 | PDT | ATRA+PDT | 0.337508 | 1.1976 | 1.82715 | 0.00005 | 0.00102476 |
| IGFBP6 | chr12:53491435-53496128 | PDT | ATRA+PDT | 31.6079 | 110.977 | 1.81191 | 0.00005 | 0.00102476 |
| CHST4 | chr16:71560022-71572493 | PDT | ATRA+PDT | 0.355725 | 1.21076 | 1.76709 | 0.00015 | 0.00281494 |
| TAPBPL | chr12:6561176-6579843 | PDT | ATRA+PDT | 0.508899 | 1.72817 | 1.7638 | 0.00035 | 0.00584867 |
| LCN2 | chr9:130911731-130915734 | PDT | ATRA+PDT | 67.9208 | 229.774 | 1.75829 | 0.00005 | 0.00102476 |
| PARP10 | chr8:145051319-145060635 | PDT | ATRA+PDT | 0.401217 | 1.33341 | 1.73267 | 0.00005 | 0.00102476 |
| TRIM31 | chr6_ssto_hap7:1401126-1411311 | PDT | ATRA+PDT | 23.1248 | 76.6326 | 1.72852 | 0.00005 | 0.00102476 |
| AKAP12 | chr6:151561133-151679694 | PDT | ATRA+PDT | 5.55638 | 17.5063 | 1.65566 | 0.00005 | 0.00102476 |
| MTSS1 | chr8:125563027-125740730 | PDT | ATRA+PDT | 1.69547 | 5.32383 | 1.65078 | 0.00005 | 0.00102476 |
| DHRS3 | chr1:12627938-12677820 | PDT | ATRA+PDT | 7.01205 | 21.5381 | 1.61899 | 0.00005 | 0.00102476 |
| IFITM3 | chr11:319672-320914 | PDT | ATRA+PDT | 3.17628 | 9.52404 | 1.58423 | 0.00005 | 0.00102476 |
| WFDC2 | chr20:44098393-44110172 | PDT | ATRA+PDT | 2.28187 | 6.53873 | 1.51879 | 0.0021 | 0.0238805 |
| SAMD9 | chr7:92728825-92747336 | PDT | ATRA+PDT | 8.37085 | 23.382 | 1.48196 | 0.00005 | 0.00102476 |
| LOXL4 | chr10:100007442-100028007 | PDT | ATRA+PDT | 1.62446 | 4.43023 | 1.44742 | 0.00005 | 0.00102476 |
| IL15RA | chr10:5994333-6020150 | PDT | ATRA+PDT | 1.29552 | 3.53252 | 1.44716 | 0.00005 | 0.00102476 |
| CNGA1 | chr4:47937993-48014961 | PDT | ATRA+PDT | 0.664166 | 1.77801 | 1.42065 | 0.00005 | 0.00102476 |
| STRA6 | chr15:74471807-74502046 | PDT | ATRA+PDT | 0.553158 | 1.47588 | 1.41581 | 0.00005 | 0.00102476 |
| SERPINA5 | chr14:95047730-95059457 | PDT | ATRA+PDT | 0.689563 | 1.79855 | 1.38308 | 0.00005 | 0.00102476 |
| VAMP5 | chr2:85811530-85820511 | PDT | ATRA+PDT | 3.04146 | 7.92904 | 1.38238 | 0.00005 | 0.00102476 |
| C10orf54 | chr10:73156690-73575704 | PDT | ATRA+PDT | 1.13266 | 2.95064 | 1.38132 | 0.00005 | 0.00102476 |
| GABRP | chr5:170210722-170241050 | PDT | ATRA+PDT | 1.85944 | 4.52118 | 1.28183 | 0.00005 | 0.00102476 |
| C10orf10 | chr10:45455218-45490172 | PDT | ATRA+PDT | 3.78595 | 9.13062 | 1.27006 | 0.00005 | 0.00102476 |
| ITPKA | chr15:41786055-41795757 | PDT | ATRA+PDT | 0.556708 | 1.31432 | 1.23932 | 0.00425 | 0.0420979 |
| BTN3A1 | chr6:26402464-26415444 | PDT | ATRA+PDT | 0.773794 | 1.7837 | 1.20485 | 0.00005 | 0.00102476 |
| PLAC8 | chr4:84011210-84035911 | PDT | ATRA+PDT | 2.40008 | 5.49895 | 1.19608 | 0.00005 | 0.00102476 |
| ATF3 | chr1:212738675-212794119 | PDT | ATRA+PDT | 97.0536 | 219.94 | 1.18025 | 0.00005 | 0.00102476 |
| HDAC9 | chr7:18126571-19036992 | PDT | ATRA+PDT | 1.44155 | 3.20797 | 1.15404 | 0.00005 | 0.00102476 |
| LGALS9 | chr17:25958173-25976586 | PDT | ATRA+PDT | 8.81966 | 19.451 | 1.14105 | 0.00005 | 0.00102476 |
| SLC2A12 | chr6:134308718-134373789 | PDT | ATRA+PDT | 0.608026 | 1.3385 | 1.13841 | 0.00005 | 0.00102476 |
| CLU | chr8:27454433-27472328 | PDT | ATRA+PDT | 4.61759 | 10.0572 | 1.12302 | 0.00005 | 0.00102476 |
| DUSP2 | chr2:96808907-96811179 | PDT | ATRA+PDT | 12.1886 | 26.538 | 1.12252 | 0.00005 | 0.00102476 |
| CREB5 | chr7:28338939-28865511 | PDT | ATRA+PDT | 3.6883 | 8.02894 | 1.12225 | 0.00005 | 0.00102476 |
| KCNE3 | chr11:74165885-74178600 | PDT | ATRA+PDT | 3.59091 | 7.78606 | 1.11655 | 0.00005 | 0.00102476 |
| SMPD3 | chr16:68392229-68482409 | PDT | ATRA+PDT | 2.86222 | 6.1399 | 1.10108 | 0.00005 | 0.00102476 |
| ALDH1A3 | chr15:101420008-101456830 | PDT | ATRA+PDT | 17.4142 | 37.3478 | 1.10076 | 0.00005 | 0.00102476 |
| ARHGDIB | chr12:15094949-15114562 | PDT | ATRA+PDT | 20.5853 | 43.954 | 1.09438 | 0.00005 | 0.00102476 |
| NR0B2 | chr1:27237974-27240567 | PDT | ATRA+PDT | 1.38854 | 2.94617 | 1.08527 | 0.00055 | 0.0085238 |
| C1orf130 | chr1:24882566-24935818 | PDT | ATRA+PDT | 1.04809 | 2.20534 | 1.07323 | 0.00005 | 0.00102476 |
| MMP7 | chr11:102391238-102401478 | PDT | ATRA+PDT | 71.2114 | 148.966 | 1.0648 | 0.00005 | 0.00102476 |
| IL8 | chr4:74606222-74609433 | PDT | ATRA+PDT | 14.3248 | 29.5956 | 1.04686 | 0.00005 | 0.00102476 |
| BIRC3 | chr11:102188180-102210135 | PDT | ATRA+PDT | 1.25282 | 2.58 | 1.04219 | 0.00005 | 0.00102476 |
| MEGF6 | chr1:3404505-3528059 | PDT | ATRA+PDT | 0.819526 | 1.67993 | 1.03554 | 0.00005 | 0.00102476 |
| EGR1 | chr5:137801180-137805004 | PDT | ATRA+PDT | 4.82157 | 9.81097 | 1.02489 | 0.00005 | 0.00102476 |
| HBEGF | chr5:139712427-139726188 | PDT | ATRA+PDT | 20.527 | 41.5863 | 1.01859 | 0.00005 | 0.00102476 |
| PELI2 | chr14:56585092-56768031 | PDT | ATRA+PDT | 1.87101 | 3.7817 | 1.01522 | 0.00005 | 0.00102476 |
| Down-regulated genes | | | | | | | | |
| ID2 | chr2:8822112-8824583 | PDT | ATRA+PDT | 5.50856 | 2.75203 | -1.00118 | 0.00005 | 0.00102476 |
| SPDEF | chr6:34505578-34524110 | PDT | ATRA+PDT | 16.5058 | 8.19596 | -1.00999 | 0.00005 | 0.00102476 |
| HEPH | chrX:65382432-65487230 | PDT | ATRA+PDT | 3.62214 | 1.79148 | -1.01569 | 0.00005 | 0.00102476 |
| AKR1C1 | chr10:5005453-5020158 | PDT | ATRA+PDT | 50.1133 | 24.6981 | -1.02079 | 0.00005 | 0.00102476 |
| GLI2 | chr2:121554866-121750229 | PDT | ATRA+PDT | 1.73488 | 0.852615 | -1.02487 | 0.00005 | 0.00102476 |
| FXYD3 | chr19:35606731-35615228 | PDT | ATRA+PDT | 17.5797 | 8.62688 | -1.027 | 0.00005 | 0.00102476 |
| FAM3D | chr3:58619669-58652561 | PDT | ATRA+PDT | 8.77298 | 4.24277 | -1.04806 | 0.00005 | 0.00102476 |
| CAPN9 | chr1:230883129-230937749 | PDT | ATRA+PDT | 1.45467 | 0.70276 | -1.04958 | 0.00145 | 0.0179618 |
| ASS1 | chr9:133320093-133376661 | PDT | ATRA+PDT | 5.71336 | 2.75499 | -1.05229 | 0.00005 | 0.00102476 |
| LRRC31 | chr3:169557028-169587660 | PDT | ATRA+PDT | 1.39178 | 0.656509 | -1.08404 | 0.0006 | 0.00910057 |
| MUC13 | chr3:124624288-124653595 | PDT | ATRA+PDT | 28.9957 | 13.5277 | -1.09993 | 0.00005 | 0.00102476 |
| CEACAM6 | chr19:42259397-42276113 | PDT | ATRA+PDT | 47.7562 | 22.2689 | -1.10066 | 0.00005 | 0.00102476 |
| S100A14 | chr1:153586731-153588808 | PDT | ATRA+PDT | 76.7108 | 35.2653 | -1.12118 | 0.00005 | 0.00102476 |
| MSLN | chr16:810764-818865 | PDT | ATRA+PDT | 2.81002 | 1.28159 | -1.13265 | 0.00005 | 0.00102476 |
| PRAP1 | chr10:135160843-135166187 | PDT | ATRA+PDT | 3.20396 | 1.43399 | -1.15982 | 0.00515 | 0.0488381 |
| GPD1 | chr12:50497800-50505095 | PDT | ATRA+PDT | 2.28955 | 1.01978 | -1.1668 | 0.00005 | 0.00102476 |
| HES6 | chr2:239146907-239148681 | PDT | ATRA+PDT | 5.73401 | 2.55042 | -1.16881 | 0.00005 | 0.00102476 |
| GRHL3 | chr1:24645811-24741587 | PDT | ATRA+PDT | 1.75026 | 0.776258 | -1.17296 | 0.002 | 0.0229388 |
| DPEP1 | chr16:89679715-89704839 | PDT | ATRA+PDT | 6.81875 | 2.92142 | -1.22284 | 0.00005 | 0.00102476 |
| WNT11 | chr11:75897369-75917574 | PDT | ATRA+PDT | 4.1994 | 1.77853 | -1.2395 | 0.00005 | 0.00102476 |
| CEACAM5 | chr19:42212529-42234437 | PDT | ATRA+PDT | 12.0597 | 5.06265 | -1.25223 | 0.00005 | 0.00102476 |
| ALDH1A1 | chr9:75515577-75568233 | PDT | ATRA+PDT | 405.218 | 165.602 | -1.29098 | 0.00005 | 0.00102476 |
| BGN | chrX:152760346-152775004 | PDT | ATRA+PDT | 16.5827 | 6.73104 | -1.30078 | 0.00005 | 0.00102476 |
| ANKS4B | chr16:21245015-21263750 | PDT | ATRA+PDT | 1.89722 | 0.765152 | -1.31007 | 0.00005 | 0.00102476 |
| AIFM3 | chr22:21319417-21335649 | PDT | ATRA+PDT | 1.47668 | 0.587014 | -1.33089 | 0.00005 | 0.00102476 |
| IGSF1 | chrX:130407482-130423403 | PDT | ATRA+PDT | 3.04443 | 1.18424 | -1.36221 | 0.00005 | 0.00102476 |
| PPP1R1B | chr17:37783176-37792878 | PDT | ATRA+PDT | 8.15642 | 2.98407 | -1.45065 | 0.00005 | 0.00102476 |
| DDC | chr7:50526133-50633154 | PDT | ATRA+PDT | 11.9731 | 4.23146 | -1.50057 | 0.00005 | 0.00102476 |
| RNASE4 | chr14:21152335-21168758 | PDT | ATRA+PDT | 2.37664 | 0.837993 | -1.50391 | 0.00085 | 0.0119898 |
| PSCA | chr8:143751725-143764145 | PDT | ATRA+PDT | 1.7825 | 0.615034 | -1.53516 | 0.002 | 0.0229388 |
| SOSTDC1 | chr7:16501105-16505474 | PDT | ATRA+PDT | 1.20435 | 0.404531 | -1.57393 | 0.0015 | 0.0184954 |
| AXIN2 | chr17:63524682-63557740 | PDT | ATRA+PDT | 14.4559 | 4.59396 | -1.65385 | 0.00005 | 0.00102476 |
| CCL15 | chr17:34310691-34329084 | PDT | ATRA+PDT | 15.6012 | 4.77259 | -1.70881 | 0.00005 | 0.00102476 |
| ABCC2 | chr10:101542462-101611662 | PDT | ATRA+PDT | 1.13309 | 0.335678 | -1.75511 | 0.00005 | 0.00102476 |
| PRODH | chr22:18900286-18924066 | PDT | ATRA+PDT | 1.66635 | 0.459819 | -1.85756 | 0.0001 | 0.00194145 |
| LEPREL1 | chr3:189674516-189840226 | PDT | ATRA+PDT | 3.14966 | 0.868411 | -1.85875 | 0.00005 | 0.00102476 |
| ST6GALNAC1 | chr17:74620844-74639894 | PDT | ATRA+PDT | 1.73672 | 0.466286 | -1.89708 | 0.00005 | 0.00102476 |
| HHLA2 | chr3:108021331-108097126 | PDT | ATRA+PDT | 4.62643 | 1.14286 | -2.01725 | 0.00005 | 0.00102476 |
| ANXA13 | chr8:124693033-124749647 | PDT | ATRA+PDT | 5.31172 | 1.27258 | -2.06142 | 0.00005 | 0.00102476 |
| SLC7A7 | chr14:23242431-23289020 | PDT | ATRA+PDT | 4.55966 | 0.977627 | -2.22157 | 0.00005 | 0.00102476 |
| LOC146336 | chr16:1114081-1131454 | PDT | ATRA+PDT | 3.05651 | 0.610603 | -2.32358 | 0.00005 | 0.00102476 |
| TFF3 | chr21:43731776-43735706 | PDT | ATRA+PDT | 6.41603 | 0.37653 | -4.09084 | 0.0026 | 0.0281856 |
| HMGCS2 | chr1:120290618-120311555 | PDT | ATRA+PDT | 16.1869 | 0.803481 | -4.33242 | 0.00005 | 0.00102476 |
| REG4 | chr1:120336640-120354203 | PDT | ATRA+PDT | 70.8796 | 2.04563 | -5.11475 | 0.00005 | 0.00102476 |
| HIST1H2BN | chr6:27806439-27806888 | PDT | ATRA+PDT | 1.30759 | 0 | -inf | 0.00145 | 0.0179618 |
| KLK12 | chr19:51532347-51538148 | PDT | ATRA+PDT | 2.7591 | 0 | -inf | 0.00005 | 0.00102476 |
| SPINK4 | chr9:33240195-33248565 | PDT | ATRA+PDT | 5.51131 | 0 | -inf | 0.00005 | 0.00102476 |
