## Supplementary Table 3 for "All-*trans* retinoic acid enhances the anti-tumour effects of fimaporfin-based photodynamic therapy"

Table S3: Pathways identification after ATRA+PDT compared to PDT.

| Ingenuity Canonical Pathways | -log(p-value) | Ratio | z-score | Molecules |
|---|---|---|---|---|
| Aryl Hydrocarbon Receptor Signaling | 5.89 | 0.154 | 1.155 | ALDH1A1,ALDH1A3,ALDH1B1,ALDH1L1,ALDH2,ALDH7A1,CCNA2,CDK6,CYP1A1,FAS,FOS,GSTA4,HSPB1,JUN,NCOA3,NCOA7,NQO2,NR0B2,NRIP1,RARG,TGFB2,TGM2 |
| Tryptophan Degradation X (Mammalian, via Tryptamine) | 5.88 | 0.36 | 0 | AKR1B10,ALDH1A1,ALDH1A3,ALDH1B1,ALDH2,ALDH7A1,DDC,MAOA,MAOB |
| Putrescine Degradation III | 5.5 | 0.381 | 0.707 | ALDH1A1,ALDH1A3,ALDH1B1,ALDH2,ALDH7A1,MAOA,MAOB,SAT1 |
| Unfolded protein response | 4.95 | 0.214 | 2.333 | CEBPA,CEBPD,DNAJB9,DNAJC3,ERN1,HSPA1A/HSPA1B,HSPA1L,HSPA6,HSPH1,INSIG1,PPARG,PPP1R15A |
| Oxidative Ethanol Degradation III | 4.76 | 0.368 | -0.378 | ACSL1,ACSS2,ALDH1A1,ALDH1A3,ALDH1B1,ALDH2,ALDH7A1 |
| Superpathway of Cholesterol Biosynthesis | 4.32 | 0.276 | -2.828 | ACAT1,HMGCR,HMGCS2,IDI1,LSS,MSMO1,SC5D,SQLE |
| Ethanol Degradation IV | 4.15 | 0.304 | -0.378 | ACSL1,ACSS2,ALDH1A1,ALDH1A3,ALDH1B1,ALDH2,ALDH7A1 |
| VDR/RXR Activation | 4.09 | 0.167 | -0.378 | CEBPA,CYP24A1,GADD45A,IGFBP6,KLF4,KLK6,MXD1,NCOA3,PRKCA,RUNX2,SEMA3B,TGFB2,VDR |
| Histamine Degradation | 4.03 | 0.353 | 0 | ALDH1A1,ALDH1A3,ALDH1B1,ALDH2,ALDH7A1,AOC1 |
| Tumor Microenvironment Pathway | 3.95 | 0.119 | 0.218 | CD44,CSPG4,CXCL8,CXCR4,FAS,FGF19,FOS,HLA-B,HLA-C,HLA-E,IGF2,JUN,LGALS9,MAP3K14,MMP15,MMP7,PIK3R1,PTGS2,RAP2B,SLC16A1,TGFB2 |
| Senescence Pathway | 3.81 | 0.102 | 1.512 | ACVR1,ATF3,CAPN5,CAPN9,CBX7,CDC25A,CDK6,CDKN2B,CGAS,CXCL8,DMTF1,E2F7,ELF3,ETS1,GADD45A,GADD45B,HIPK2,ITPR2,JUN,PDK4,PIK3R1,PML,PPP2R5B,RAP2B,SMAD3,TGFB2,TGFBR2,ZFP36L1 |
| Aldosterone Signaling in Epithelial Cells | 3.67 | 0.12 | -0.816 | ASIC1,DNAJA1,DNAJB1,DNAJB4,DNAJB9,DNAJC22,DNAJC3,DUSP1,HSPA1A/HSPA1B,HSPA1L,HSPA4L,HSPA6,HSPB1,HSPH1,ITPR2,PIK3R1,PIP5K1A,PRKCA,SGK1 |
| Molecular Mechanisms of Cancer | 3.63 | 0.09 | #NUM! | AURKA,BBC3,BIRC3,BMP4,CASP7,CASP9,CDC25A,CDK6,CDKN2B,DVL1,E2F7,FAS,FOS,FYN,GAB1,GAB2,GLI1,GNA14,GNAI1,GNAZ,HIPK2,JUN,PIK3R1,PMAIP1,PRKAR2A,PRKCA,RAP2B,RASA1,RHOB,RHOD,RHOU,RND3,SMAD3,TGFB2,TGFBR2,WNT11 |
| PXR/RXR Activation | 3.61 | 0.169 | #NUM! | ABCC2,ALDH1A1,CPT1A,CYP2B6,CYP3A5,HMGCS2,HNF4A,NR0B2,PCK2,PRKAR2A,UGT1A10 (includes others) |
| Fatty Acid α-oxidation | 3.58 | 0.3 | 0.816 | ALDH1A1,ALDH1A3,ALDH1B1,ALDH2,ALDH7A1,PTGS2 |
| Protein Ubiquitination Pathway | 3.5 | 0.0989 | #NUM! | B2M,BIRC3,DNAJA1,DNAJB1,DNAJB4,DNAJB9,DNAJC22,DNAJC3,HLA-B,HLA-C,HLA-E,HSPA1A/HSPA1B,HSPA1L,HSPA4L,HSPA6,HSPB1,HSPH1,NEDD4L,PSMB10,PSMB8,PSME1,PSME2,SMURF1,TAP2,UBC,USP11,USP53 |
| Sumoylation Pathway | 3.39 | 0.136 | 0 | ARHGDIB,CEBPA,DNMT3A,ETS1,FAS,FOS,JUN,MYB,PML,RHOB,RHOD,RHOU,RND3,SP100 |
| Death Receptor Signaling | 3.36 | 0.141 | 1.941 | ARHGDIB,BIRC3,CASP7,CASP9,FAS,HSPB1,MAP3K14,MAP4K4,PARP10,PARP12,PARP14,PARP8,TIPARP |
| Dopamine Degradation | 3.35 | 0.233 | 0.378 | ALDH1A1,ALDH1A3,ALDH1B1,ALDH2,ALDH7A1,MAOA,MAOB |
| Huntington's Disease Signaling | 3.27 | 0.1 | -2.324 | ATP5MC1,CAPN5,CAPN9,CASP4,CASP7,CASP9,CREB5,DNAJB1,GNA14,HDAC9,HSPA1A/HSPA1B,HSPA1L,HSPA6,JUN,PIK3R1,POLR2L,PRKCA,PSME1,PSME2,RASA1,SGK1,STX1A,TGM2,UBC |
| LPS/IL-1 Mediated Inhibition of RXR Function | 3.27 | 0.102 | 0.333 | ABCC2,ACSL1,ALDH1A1,ALDH1A3,ALDH1B1,ALDH1L1,ALDH2,ALDH7A1,CHST4,CPT1A,CYP2B6,CYP3A5,GSTA4,HMGCS2,IL1R2,IL1RN,JUN,MAOA,MAOB,NR0B2,NR5A2,SCARB1,SULT2B1 |
| TGF-β Signaling | 3.18 | 0.135 | 0.905 | ACVR1,BMP4,FOS,HNF4A,IRF7,JUN,RAP2B,RUNX2,SMAD3,SMURF1,TGFB2,TGFBR2,VDR |
| Ethanol Degradation II | 3.17 | 0.219 | -0.378 | ACSL1,ACSS2,ALDH1A1,ALDH1A3,ALDH1B1,ALDH2,ALDH7A1 |
| Caveolar-mediated Endocytosis Signaling | 3.16 | 0.151 | #NUM! | B2M,CAV1,CD55,FLNB,FLOT1,FYN,HLA-B,HLA-C,HLA-E,ITGB6,PRKCA |
| Endothelin-1 Signaling | 3.13 | 0.106 | 1.606 | CASP4,CASP7,CASP9,ECE2,EDN1,FOS,GAB1,GNA14,GNAI1,GNAZ,ITPR2,JUN,PIK3R1,PLAAT3,PLAAT4,PLD1,PLD2,PRKCA,PTGS2,RAP2B |
| NRF2-mediated Oxidative Stress Response | 3.1 | 0.106 | 0.632 | ABCC1,ABCC2,AKR7A3,DNAJA1,DNAJA4,DNAJB1,DNAJB4,DNAJB9,DNAJC3,FOS,FOSL1,GSTA4,JUN,JUND,MAFF,NQO2,PIK3R1,PRKCA,RAP2B,SCARB1 |
| UVA-Induced MAPK Signaling | 3.09 | 0.133 | 0 | CASP9,FOS,JUN,PARP10,PARP12,PARP14,PARP8,PIK3R1,PRKCA,RAP2B,RPS6KB2,SMPD3,TIPARP |
| Xenobiotic Metabolism PXR Signaling Pathway | 3.02 | 0.104 | 0.447 | ABCC2,ALDH1A1,ALDH1A3,ALDH1B1,ALDH1L1,ALDH2,ALDH7A1,CHST4,CYP2B6,CYP3A5,GSTA4,MAOA,MAOB,NRIP1,PPP1R10,PRKAR2A,PRKCA,SULT2B1,UGT1A10 (includes others),UGT8 |
| Cytotoxic T Lymphocyte-mediated Apoptosis of Target Cells | 3 | 0.206 | #NUM! | B2M,CASP7,CASP9,FAS,HLA-B,HLA-C,HLA-E |
| Noradrenaline and Adrenaline Degradation | 2.92 | 0.2 | 0.378 | ALDH1A1,ALDH1A3,ALDH1B1,ALDH2,ALDH7A1,MAOA,MAOB |
| Interferon Signaling | 2.85 | 0.194 | 2.646 | IFI35,IFIT3,IFITM2,IFITM3,IFNGR2,IRF1,PSMB8 |
| Xenobiotic Metabolism CAR Signaling Pathway | 2.72 | 0.101 | 1.147 | ABCC1,ABCC2,ALDH1A1,ALDH1A3,ALDH1B1,ALDH1L1,ALDH2,ALDH7A1,CHST4,CYP1A1,CYP2B6,CYP3A5,GSTA4,NRIP1,PPP2R5B,PRKCA,SULT2B1,UGT1A10 (includes others),UGT8 |
| Stearate Biosynthesis I (Animals) | 2.67 | 0.163 | 0.378 | ACOT11,ACSL1,ELOVL6,GNPAT,HNF4A,MBOAT7,SRD5A3,TBXAS1 |
| Retinoic acid Mediated Apoptosis Signaling | 2.67 | 0.15 | 2.828 | CASP9,CRABP2,IRF1,PARP10,PARP12,PARP14,PARP8,RARG,TIPARP |
| Antigen Presentation Pathway | 2.64 | 0.179 | #NUM! | B2M,HLA-B,HLA-C,HLA-E,PSMB8,TAP2,TAPBP |
| Xenobiotic Metabolism AHR Signaling Pathway | 2.61 | 0.129 | 0.905 | ALDH1A1,ALDH1A3,ALDH1B1,ALDH1L1,ALDH2,ALDH7A1,CYP1A1,GSTA4,NQO2,NRIP1,UGT1A10 (includes others) |
| p53 Signaling | 2.6 | 0.122 | 1.265 | BBC3,FAS,GADD45A,GADD45B,HDAC9,HIPK2,JUN,PIK3R1,PMAIP1,PML,SERPINE2,TRIM29 |
| Cholesterol Biosynthesis I | 2.58 | 0.308 | -2 | LSS,MSMO1,SC5D,SQLE |
| Cholesterol Biosynthesis II (via 24,25-dihydrolanosterol) | 2.58 | 0.308 | -2 | LSS,MSMO1,SC5D,SQLE |
| Cholesterol Biosynthesis III (via Desmosterol) | 2.58 | 0.308 | -2 | LSS,MSMO1,SC5D,SQLE |
| Hepatic Fibrosis Signaling Pathway | 2.57 | 0.082 | 0.18 | ACVR1,COL18A1,CREB5,CXCL8,DVL1,EDN1,FOS,GLI1,GLI2,GLIS2,GNAI1,IL1R2,IL1RN,IRS2,JUN,KLF9,NOX1,PIK3R1,PPARG,PRKAR2A,PRKCA,RAP2B,RHOB,RHOD,RHOU,RND3,RPS6KB2,SMAD3,TGFB2,TGFBR2,WNT11 |
| Axonal Guidance Signaling | 2.52 | 0.0769 | #NUM! | ABLIM1,ADAM8,ADAM9,ADAMTS6,BMP4,CXCR4,ECE2,EFNA1,EPHA2,EPHB6,ERAP2,FYN,GLI1,GLI2,GLIS2,GNA14,GNAI1,GNAZ,MMP15,MMP7,NRP1,NTN4,PIK3R1,PLXNB2,PLXND1,PRKAR2A,PRKCA,RAP2B,RASA1,RHOD,SEMA3A,SEMA3B,SEMA3E,SEMA4B,SEMA7A,TUBA4A,TUBB2A,WNT11 |
| Virus Entry via Endocytic Pathways | 2.45 | 0.118 | #NUM! | B2M,CAV1,CD55,FLNB,FYN,HLA-B,HLA-C,HLA-E,ITGB6,PIK3R1,PRKCA,RAP2B |
| Mevalonate Pathway I | 2.44 | 0.286 | -2 | ACAT1,HMGCR,HMGCS2,IDI1 |
| IL-8 Signaling | 2.44 | 0.095 | 1.606 | ANGPT1,CXCL1,CXCL8,FOS,GNAI1,HBEGF,JUN,MAP4K4,NOX1,PIK3R1,PLD1,PLD2,PRKCA,PTGS2,RAP2B,RHOB,RHOD,RHOU,RND3 |
| Hepatic Cholestasis | 2.42 | 0.0968 | #NUM! | ABCC1,ABCC2,CLCF1,CXCL8,CYP3A5,FGF19,HNF4A,IL1R2,IL1RN,JUN,LIF,MAP3K14,NR0B2,NR5A2,PRKAR2A,PRKCA,TGFB2,TJP2 |
| Granulocyte Adhesion and Diapedesis | 2.39 | 0.0983 | #NUM! | CCL15,CLDN1,CLDN12,CLDN2,CXCL1,CXCL16,CXCL2,CXCL3,CXCL8,CXCR4,GNAI1,HRH1,IL1R2,IL1RN,MMP15,MMP7,SDC4 |
| Polyamine Regulation in Colon Cancer | 2.39 | 0.217 | #NUM! | MXD1,PPARG,PSME1,PSME2,SAT1 |
| IGF-1 Signaling | 2.38 | 0.115 | -0.333 | CASP9,CCN1,FOS,IGFBP2,IGFBP6,IRS2,JUN,PIK3R1,PRKAR2A,RAP2B,RASA1,RPS6KB2 |
| Role of PKR in Interferon Induction and Antiviral Response | 2.36 | 0.11 | 1.387 | ATF3,CASP9,DNAJC3,EIF2AK2,FAS,FOS,HSPA1A/HSPA1B,HSPA1L,HSPA6,IFIH1,IFNGR2,IRF1,JUN |
| HIF1α Signaling | 2.32 | 0.0927 | -1.147 | CCNG2,EDN1,EGLN1,HK1,HSPA1A/HSPA1B,HSPA1L,HSPA6,IGF2,JUN,LDHA,MMP15,MMP7,NOX1,PIK3R1,PRKCA,RAP2B,RPS6KB2,SAT1,TGFB2 |
| Glucocorticoid Receptor Signaling | 2.27 | 0.0758 | #NUM! | ANXA1,ATP5MC1,CAV1,CEBPA,CXCL3,CXCL8,DUSP1,FOS,HSPA1A/HSPA1B,HSPA1L,HSPA6,IL15RA,IL1R2,IL1RN,JUN,KRT13,KRT20,KRT7,KRT80,MAP3K14,NCOA3,NRIP1,PCK2,PDK4,PIK3R1,POLR2L,PPARG,PTGS2,RAP2B,SGK1,SMAD3,TGFB2,TGFBR2,TSC22D3,UQCRFS1 |
| Coagulation System | 2.23 | 0.171 | 0 | F2R,F5,PROS1,SERPINA1,SERPINA5,TFPI |
| RAR Activation | 2.23 | 0.0928 | #NUM! | AKR1B10,AKR1C3,ALDH1A1,ALDH1A3,CRABP2,DHRS3,DUSP1,FOS,JUN,NRIP1,PIK3R1,PML,PRKAR2A,PRKCA,RARG,REL,SMAD3,TGFB2 |
| Xenobiotic Metabolism Signaling | 2.21 | 0.0836 | #NUM! | ABCC2,ALDH1A1,ALDH1A3,ALDH1B1,ALDH1L1,ALDH2,ALDH7A1,CHST4,CYP1A1,CYP2B6,CYP3A5,GSTA4,MAOA,MAOB,MAP3K14,NQO2,NRIP1,PIK3R1,PPP2R5B,PRKCA,RAP2B,SULT2B1,UGT1A10 (includes others),UGT8 |
| Anandamide Degradation | 2.18 | 0.667 | #NUM! | FAAH,NAAA |
| Systemic Lupus Erythematosus In B Cell Signaling Pathway | 2.15 | 0.0836 | 3.545 | BCL10,CLCF1,CXCL8,FOS,FYN,GAB1,IFIH1,IFIT3,IFNGR2,INPPL1,IRF7,JUN,LIF,MAP3K14,MAP4K4,MCL1,PAG1,PIK3R1,PLAAT4,PRKCA,RAP2B,TGFB2,TRAF4 |
| Salvage Pathways of Pyrimidine Ribonucleotides | 2.14 | 0.112 | 0 | AK4,APOBEC3B,CDK6,DAPK1,EIF2AK2,GRK5,NME3,NME4,PIM1,PKN1,SGK1 |
| FXR/RXR Activation | 2.12 | 0.103 | #NUM! | ABCC2,AMBP,APOL1,CLU,FGF19,HNF4A,IL1RN,NR0B2,NR5A2,PCK2,PPARG,SCARB1,SERPINA1 |
| Semaphorin Signaling in Neurons | 2.12 | 0.133 | #NUM! | FYN,NRP1,RHOB,RHOD,RHOU,RND3,SEMA3A,SEMA7A |
| MSP-RON Signaling In Macrophages Pathway | 2.09 | 0.106 | 1.155 | CREB5,FOS,GAB2,IFNGR2,JUN,KLK10,KLK6,NFKBIZ,PIK3R1,PTGS2,RAP2B,SBNO2 |
| PDGF Signaling | 2.09 | 0.116 | 1.897 | CAV1,EIF2AK2,FOS,INPPL1,JUN,PIK3R1,PRKCA,RAP2B,RASA1,SPHK2 |
| Germ Cell-Sertoli Cell Junction Signaling | 2.08 | 0.0936 | #NUM! | EPN3,FER,GSN,MAP3K14,PIK3R1,RAP2B,RHOB,RHOD,RHOU,RND3,TGFB2,TGFBR2,TJP1,TUBA4A,TUBB2A,ZYX |
| Apoptosis Signaling | 2.07 | 0.11 | -1.667 | BIRC3,CAPN5,CAPN9,CASP7,CASP9,FAS,MAP3K14,MAP4K4,MCL1,PRKCA,RAP2B |
| Superpathway of Geranylgeranyldiphosphate Biosynthesis I (via Mevalonate) | 2.03 | 0.222 | -2 | ACAT1,HMGCR,HMGCS2,IDI1 |
| Wnt/β-catenin Signaling | 2.03 | 0.0925 | -1.291 | ACVR1,AXIN2,CD44,DKK1,DVL1,JUN,MMP7,NR5A2,PPP2R5B,RARG,SOX12,SOX4,TGFB2,TGFBR2,UBC,WNT11 |
| Regulation Of The Epithelial Mesenchymal Transition By Growth Factors Pathway | 2.03 | 0.0904 | 2 | EGR1,ETS1,FGF19,FGFR2,FOS,GAB1,HMGA2,ID2,JUN,LATS2,PARD6B,PIK3R1,RAP2B,SMAD3,SMURF1,TGFB2,TGFBR2 |
| Osteoarthritis Pathway | 2 | 0.0864 | 0.775 | CASP4,CASP7,CASP9,CREB5,CXCL8,DDIT4,DKK1,ELF3,GLI1,GLI2,GLIS2,IL1R2,PPARG,PTGS2,RUNX2,SDC4,SLC39A8,SMAD3,TGFBR2 |
| Prostanoid Biosynthesis | 1.99 | 0.3 | #NUM! | CYP2S1,PTGS2,TBXAS1 |
| GDNF Family Ligand-Receptor Interactions | 1.98 | 0.118 | 1.633 | FOS,GAB1,IRS2,ITPR2,JUN,PDLIM7,PIK3R1,RAP2B,RASA1 |
| PPARα/RXRα Activation | 1.97 | 0.089 | -1.5 | ACAA1,ACVR1,BCL3,GNA14,GPD1,IL1R2,JUN,MAP3K14,MAP4K4,NCOA3,NR0B2,PRKAR2A,PRKCA,RAP2B,SMAD3,TGFB2,TGFBR2 |
| Regulation of the Epithelial-Mesenchymal Transition Pathway | 1.95 | 0.0885 | #NUM! | DVL1,EGR1,ETS1,FGF19,FGFR2,GAB1,HMGA2,ID2,JAG2,PARD6B,PIK3R1,RAP2B,SMAD3,SMURF1,TGFB2,TGFBR2,WNT11 |
| PPAR Signaling | 1.92 | 0.105 | -1.508 | FOS,IL1R2,IL1RN,JUN,MAP3K14,MAP4K4,NR0B2,NRIP1,PPARG,PTGS2,RAP2B |
| Melatonin Degradation II | 1.89 | 0.5 | #NUM! | MAOA,MAOB |
| Acetate Conversion to Acetyl-CoA | 1.89 | 0.5 | #NUM! | ACSL1,ACSS2 |
| Integrin Signaling | 1.84 | 0.0845 | -0.277 | CAPN5,CAPN9,CAV1,FYN,GRB7,GSN,ITGB6,NEDD9,PIK3R1,RAP2B,RHOB,RHOD,RHOU,RND3,TNK2,TSPAN4,TSPAN6,ZYX |
| Serotonin Degradation | 1.84 | 0.119 | 0.707 | ALDH1A1,ALDH1A3,ALDH1B1,ALDH2,ALDH7A1,MAOA,MAOB,UGT1A10 (includes others) |
| Estrogen Biosynthesis | 1.84 | 0.143 | 1.342 | AKR1C1/AKR1C2,AKR1C3,CYP1A1,CYP2B6,CYP2S1,CYP3A5 |
| CXCR4 Signaling | 1.83 | 0.0898 | 0 | CXCR4,EGR1,FOS,GNA14,GNAI1,GNAZ,ITPR2,JUN,PIK3R1,PRKCA,RAP2B,RHOB,RHOD,RHOU,RND3 |
| Pancreatic Adenocarcinoma Signaling | 1.81 | 0.101 | 0 | CASP9,CDKN2B,E2F7,HBEGF,PIK3R1,PLD1,PLD2,PTGS2,SMAD3,TGFB2,TGFBR2 |
| Endoplasmic Reticulum Stress Pathway | 1.79 | 0.19 | 2 | CASP7,CASP9,DNAJC3,ERN1 |
| Retinol Biosynthesis | 1.75 | 0.136 | 0.447 | AADAC,AKR1B10,AKR1C3,DHRS3,LIPG,PNPLA2 |
| B Cell Receptor Signaling | 1.75 | 0.086 | 1.5 | BCL10,BCL6,CARD10,CREB5,DAPP1,EGR1,ETS1,GAB1,GAB2,INPPL1,JUN,MAP3K14,PAG1,PIK3R1,RAP2B,RPS6KB2 |
| Protein Kinase A Signaling | 1.74 | 0.0725 | -1.177 | AKAP12,CDC14B,CDC25A,CNGA1,CREB5,DUSP1,DUSP16,DUSP2,DUSP5,FLNB,GNAI1,H1-0,H1-10,ITPR2,PDE12,PDE9A,PPP1R10,PPP1R1B,PRKAR2A,PRKCA,PTGS2,PTPN21,PTPN9,PTPRB,PTPRH,PYGB,SMAD3,TGFB2,TGFBR2 |
| TR/RXR Activation | 1.72 | 0.107 | #NUM! | AKR1C1/AKR1C2,AKR1C3,BCL3,KLF9,ME1,NCOA3,PIK3R1,SCARB1,SLC16A3 |
| Production of Nitric Oxide and Reactive Oxygen Species in Macrophages | 1.7 | 0.0847 | 1 | APOL1,CLU,FOS,IFNGR2,IRF1,JUN,MAP3K14,PIK3R1,PPP1R10,PPP2R5B,PRKCA,RHOB,RHOD,RHOU,RND3,SERPINA1 |
| HIPPO signaling | 1.69 | 0.106 | -1.134 | AJUBA,CD44,LATS2,PPP1R10,PPP2R5B,RASSF6,SMAD3,TEAD2,TJP2 |
| Protein Citrullination | 1.68 | 0.4 | #NUM! | PADI1,PADI2 |
| Creatine-phosphate Biosynthesis | 1.68 | 0.4 | #NUM! | CKB,MAP4K4 |
| Retinoate Biosynthesis I | 1.66 | 0.147 | -0.447 | AKR1B10,AKR1C3,ALDH1A1,ALDH1A3,DHRS3 |
| PFKFB4 Signaling Pathway | 1.66 | 0.13 | 0 | CREB5,HK1,NCOA3,PFKM,PRKAR2A,TGFB2 |
| ERK5 Signaling | 1.66 | 0.111 | 1.414 | CREB5,FOS,FOSL1,GAB1,LIF,RAP2B,RPS6KB2,SGK1 |

Table S3: Pathways identification after ATRA+PDT compared to PDT.

| Ingenuity Canonical Pathways | -log(p-value) | Ratio | z-score | Molecules |
|---|---|---|---|---|
| Glioma Invasiveness Signaling | 1.63 | 0.11 | 0 | CD44,F2R,PIK3R1,RAP2B,RHOB,RHOD,RHOU,RND3 |
| Role of Tissue Factor in Cancer | 1.63 | 0.0948 | #NUM! | CCN1,CXCL1,CXCL8,EGR1,F2RL1,FYN,GNA14,HBEGF,PIK3R1,PRKCA,RAP2B |
| Agranulocyte Adhesion and Diapedesis | 1.62 | 0.0829 | #NUM! | CCL15,CLDN1,CLDN12,CLDN2,CXCL1,CXCL16,CXCL2,CXCL3,CXCL8,CXCR4,GNAI1,HRH1,IL1RN,MMP15,MMP7,SDC4 |
| TWEAK Signaling | 1.61 | 0.143 | 1.342 | BIRC3,CASP7,CASP9,MAP3K14,TNFRSF12A |
| Sphingosine-1-phosphate Signaling | 1.6 | 0.094 | -1.265 | ACER2,CASP4,CASP7,CASP9,GNAI1,PIK3R1,RHOB,RHOD,RHOU,RND3,SMPD3 |
| HER-2 Signaling in Breast Cancer | 1.6 | 0.0825 | 1.291 | CASP7,CASP9,ELF3,ETS1,FOS,FYN,HBEGF,ITGB6,JUN,PARD6B,PIK3R1,PRKCA,PTGS2,RAP2B,RPS6KB2,SMAD3 |
| Estrogen Receptor Signaling | 1.57 | 0.0732 | -1.043 | ATP5MC1,CAV1,CREB5,FBXO32,FOS,GNA14,GNAI1,GNAZ,IGF2,JUN,MDK,MED15,MMP15,MMP7,NCOA3,NR0B2,NRIP1,PIK3R1,PRKAR2A,PRKCA,RAP2B,RPS6KB2,RUNX2,UQCRFS1 |
| Role of IL-17A in Psoriasis | 1.57 | 0.214 | #NUM! | CXCL1,CXCL3,CXCL8 |
| Acyl-CoA Hydrolysis | 1.57 | 0.214 | #NUM! | ACOT11,GNPAT,HNF4A |
| Phenylalanine Degradation IV (Mammalian, via Side Chain) | 1.57 | 0.214 | #NUM! | ALDH2,MAOA,MAOB |
| IL-17A Signaling in Fibroblasts | 1.56 | 0.139 | #NUM! | CEBPD,FOS,JUN,LCN2,NFKBIZ |
| Cholecystokinin/Gastrin-mediated Signaling | 1.55 | 0.0924 | 1.508 | FOS,IL1RN,ITPR2,JUN,PRKCA,PTGS2,RAP2B,RHOB,RHOD,RHOU,RND3 |
| Toll-like Receptor Signaling | 1.54 | 0.105 | 1.633 | EIF2AK2,FOS,IL1RN,JUN,MAP3K14,MAP4K4,TRAF4,UBC |
| Regulation of IL-2 Expression in Activated and Anergic T Lymphocytes | 1.54 | 0.1 | #NUM! | BCL10,FOS,FYN,JUN,RAP2B,SMAD3,TGFB2,TGFBR2,TOB1 |
| Bupropion Degradation | 1.53 | 0.16 | 2 | CYP1A1,CYP2B6,CYP2S1,CYP3A5 |
| PD-1, PD-L1 cancer immunotherapy pathway | 1.51 | 0.0943 | 0 | B2M,CBLB,HLA-B,HLA-C,HLA-E,IFNGR2,LATS2,PIK3R1,SMAD3,TGFB2 |
| Thrombopoietin Signaling | 1.51 | 0.111 | 1.89 | FOS,GAB2,IRS2,JUN,PIK3R1,PRKCA,RAP2B |
| T Cell Receptor Signaling | 1.51 | 0.0943 | #NUM! | BCL10,FOS,FYN,JUN,PAG1,PIK3R1,PTPRH,RAP2B,RASA1,TXK |
| Myc Mediated Apoptosis Signaling | 1.5 | 0.12 | 0 | BBC3,CASP9,FAS,MCL1,PMAIP1,PRKAR2A |
| TNFR1 Signaling | 1.5 | 0.12 | 1.633 | BIRC3,CASP7,CASP9,FOS,JUN,MAP3K14 |
| Tight Junction Signaling | 1.5 | 0.0833 | #NUM! | CEBPA,CLDN1,CLDN12,CLDN2,FOS,JUN,PPP2R5B,PRKAR2A,SMURF1,TGFB2,TGFBR2,TJP1,TJP2,TJP3 |
| Neuroinflammation Signaling Pathway | 1.49 | 0.0733 | 1.789 | ACVR1,B2M,BACE2,BIRC3,CALB2,CREB5,CXCL8,FAS,FOS,GABRP,GRIN2D,HLA-B,HLA-C,HLA-E,IFNGR2,IRF7,JUN,NOX1,PIK3R1,PTGS2,TGFB2,TGFBR2 |
| Choline Biosynthesis III | 1.49 | 0.2 | #NUM! | CHPT1,PLD1,PLD2 |
| PI3K Signaling in B Lymphocytes | 1.47 | 0.087 | 1.265 | ATF3,BCL10,CARD10,DAPP1,FOS,FYN,IRS2,ITPR2,JUN,PIK3R1,PLEKHA2,RAP2B |
| IL-17A Signaling in Gastric Cells | 1.47 | 0.154 | 2 | CXCL1,CXCL8,FOS,JUN |
| Superpathway of Melatonin Degradation | 1.45 | 0.108 | 1.89 | CYP1A1,CYP2B6,CYP2S1,CYP3A5,MAOA,MAOB,UGT1A10 (includes others) |
| Antiproliferative Role of TOB in T Cell Signaling | 1.42 | 0.128 | 0.447 | CCNA2,SMAD3,TGFB2,TGFBR2,TOB1 |
| Pyridoxal 5'-phosphate Salvage Pathway | 1.42 | 0.106 | 0.816 | CDK6,DAPK1,EIF2AK2,GRK5,PIM1,PKN1,SGK1 |
| Ephrin Receptor Signaling | 1.41 | 0.0794 | -0.707 | ANGPT1,CREB5,CXCR4,EFNA1,EPHA2,EPHB6,FYN,GNA14,GNAI1,GNAZ,GRIN2D,MAP3K14,MAP4K4,RAP2B,RASA1 |
| Extrinsic Prothrombin Activation Pathway | 1.41 | 0.188 | #NUM! | F5,PROS1,TFPI |
| IL-6 Signaling | 1.4 | 0.0873 | 2.111 | CXCL8,FOS,HSPB1,IL1R2,IL1RN,JUN,MAP3K14,MAP4K4,MCL1,PIK3R1,RAP2B |
| Cyclins and Cell Cycle Regulation | 1.39 | 0.0988 | -1.134 | CCNA2,CDC25A,CDK6,CDKN2B,E2F7,HDAC9,PPP2R5B,TGFB2 |
| Cell Cycle: G1/S Checkpoint Regulation | 1.39 | 0.104 | 0 | CDC25A,CDK6,CDKN2B,E2F7,HDAC9,SMAD3,TGFB2 |
| Prolactin Signaling | 1.39 | 0.0988 | 1.134 | FOS,FYN,IRF1,JUN,NMI,PIK3R1,PRKCA,RAP2B |
| eNOS Signaling | 1.37 | 0.0818 | -0.707 | AQP5,CASP9,CAV1,CCNA2,CNGA1,HSPA1A/HSPA1B,HSPA1L,HSPA6,ITPR2,NOSTRIN,PIK3R1,PRKAR2A,PRKCA |
| Regulation Of The Epithelial Mesenchymal Transition In Development Pathway | 1.32 | 0.0952 | -2.121 | AXIN2,DVL1,GLI1,GLI2,GLIS2,JAG2,S100A4,WNT11 |
| EGF Signaling | 1.32 | 0.109 | 0 | FOS,ITPR2,JUN,PIK3R1,PRKCA,RASA1 |
| Sonic Hedgehog Signaling | 1.32 | 0.138 | 0 | GLI1,GLI2,GLIS2,PRKAR2A |
| Sorbitol Degradation I | 1.32 | 1 | #NUM! | SORD |
| Lanosterol Biosynthesis | 1.32 | 1 | #NUM! | LSS |
| Intrinsic Prothrombin Activation Pathway | 1.31 | 0.119 | 0.447 | COL18A1,F5,KLK10,KLK6,PROS1 |
| AMPK Signaling | 1.29 | 0.0751 | -0.277 | AK2,AK4,CCNA2,CFTR,CPT1A,CREB5,HMGCR,HNF4A,IRS2,PCK2,PFKM,PHLPP1,PIK3R1,PPP2R5B,PRKAR2A,TBC1D1 |
| Neuroprotective Role of THOP1 in Alzheimer's Disease | 1.28 | 0.0862 | 0.447 | DPP4,ECE2,HLA-B,HLA-C,HLA-E,KLK10,KLK6,NFYA,PRKAR2A,SERPINA3 |
| FAT10 Signaling Pathway | 1.28 | 0.167 | #NUM! | MAP1LC3B,PSME1,PSME2 |
| Systemic Lupus Erythematosus In T Cell Signaling Pathway | 1.27 | 0.0689 | -1.043 | B2M,BCL6,CASP4,CASP7,CASP9,CD44,CREB5,FAS,FOS,GADD45A,GNAI1,HLA-B,HLA-C,HLA-E,JUN,PIK3R1,PPP2R5B,RAP2B,RHOB,RHOD,RHOU,RND3,RPS6KB2 |
| BAG2 Signaling Pathway | 1.27 | 0.116 | 0.447 | HSPA1A/HSPA1B,HSPA1L,HSPA6,PSME1,PSME2 |
| TNFR2 Signaling | 1.27 | 0.133 | 1 | BIRC3,FOS,JUN,MAP3K14 |
| Salvage Pathways of Pyrimidine Deoxyribonucleotides | 1.27 | 0.25 | #NUM! | APOBEC3B,TK2 |
| HMGB1 Signaling | 1.26 | 0.0788 | 1.732 | CLCF1,CXCL8,FOS,IFNGR2,JUN,LIF,PIK3R1,RAP2B,RHOB,RHOD,RHOU,RND3,TGFB2 |
| IL-12 Signaling and Production in Macrophages | 1.26 | 0.0827 | #NUM! | APOL1,CLU,FOS,IRF1,JUN,PIK3R1,PPARG,PRKCA,REL,SERPINA1,TGFB2 |
| Triacylglycerol Biosynthesis | 1.23 | 0.114 | -0.447 | DGAT2,ELOVL6,LPCAT1,MBOAT7,PLPP3 |
| Acetone Degradation I (to Methylglyoxal) | 1.23 | 0.129 | 2 | CYP1A1,CYP2B6,CYP2S1,CYP3A5 |
| Chronic Myeloid Leukemia Signaling | 1.23 | 0.0874 | #NUM! | CDK6,E2F7,GAB2,HDAC9,PIK3R1,RAP2B,SMAD3,TGFB2,TGFBR2 |
| Ceramide Signaling | 1.22 | 0.0909 | 2.449 | CNKSR1,FOS,JUN,PIK3R1,PPP2R5B,RAP2B,SMPD3,SPHK2 |
| Methylglyoxal Degradation III | 1.22 | 0.158 | #NUM! | AKR1B10,AKR1C1/AKR1C2,AKR1C3 |
| Coronavirus Replication Pathway | 1.2 | 0.111 | -1.342 | BAG3,IFITM2,IFITM3,TUBA4A,TUBB2A |
| Colorectal Cancer Metastasis Signaling | 1.2 | 0.0711 | -0.258 | CASP9,DVL1,FOS,JUN,MMP15,MMP7,PIK3R1,PRKAR2A,PTGS2,RAP2B,RHOB,RHOD,RHOU,RND3,SMAD3,TGFB2,TGFBR2,WNT11 |
| ERK/MAPK Signaling | 1.2 | 0.0743 | 0 | CREB5,DUSP1,DUSP2,ELF3,ETS1,FOS,FYN,HSPB1,PIK3R1,PPARG,PPP1R10,PPP2R5B,PRKAR2A,PRKCA,RAP2B |
| HGF Signaling | 1.2 | 0.0833 | 1.89 | ELF3,ETS1,FOS,GAB1,JUN,MAP3K14,PIK3R1,PRKCA,PTGS2,RAP2B |
| Hepatic Fibrosis / Hepatic Stellate Cell Activation | 1.19 | 0.0753 | #NUM! | COL17A1,COL18A1,CXCL3,CXCL8,EDN1,FAS,FGFR2,IFNGR2,IGF2,IL1R2,KLF6,SMAD3,TGFB2,TGFBR2 |
| IL-15 Production | 1.18 | 0.0826 | 2.53 | CLK1,CLK4,EPHA2,FER,FGFR2,FYN,IRF1,ROR1,TNK2,TXK |
| SPINK1 Pancreatic Cancer Pathway | 1.17 | 0.1 | -0.816 | F2RL1,KLK10,KLK6,SMAD3,SPINK1,TGFBR2 |
| PCP pathway | 1.17 | 0.1 | -0.447 | CELSR1,DVL1,JUN,JUND,LGR4,WNT11 |
| FAT10 Cancer Signaling Pathway | 1.17 | 0.109 | 0.447 | ACVR1,CXCR4,SMAD3,TGFB2,TGFBR2 |
| Sirtuin Signaling Pathway | 1.16 | 0.0687 | 1.069 | ACSS2,ATP5MC1,CPT1A,CXCL8,GADD45A,GADD45B,H1-0,H1-10,HMGCS2,JUN,LDHA,MAP1LC3B,PCK2,PFKM,PPARG,PPIF,TUBA4A,UQCRFS1,VDAC1,XPA |
| The Visual Cycle | 1.16 | 0.15 | #NUM! | AKR1B10,AKR1C3,DHRS3 |
| Insulin Receptor Signaling | 1.15 | 0.0791 | -0.302 | ASIC1,FYN,GAB1,INPPL1,IRS2,PIK3R1,PPP1R10,PRKAR2A,RAP2B,RPS6KB2,SGK1 |
| Induction of Apoptosis by HIV1 | 1.15 | 0.0984 | 0 | BBC3,BIRC3,CASP9,CXCR4,FAS,MAP3K14 |
| Tec Kinase Signaling | 1.13 | 0.0751 | 0 | FAS,FOS,FYN,GNA14,GNAI1,GNAZ,PIK3R1,PRKCA,RHOB,RHOD,RHOU,RND3,TXK |
| ILK Signaling | 1.13 | 0.0737 | 0.832 | CREB5,FLNB,FOS,IRS2,ITGB6,JUN,LIMS2,PIK3R1,PPP2R5B,PTGS2,RHOB,RHOD,RHOU,RND3 |
| nNOS Signaling in Neurons | 1.13 | 0.106 | #NUM! | CAPN5,CAPN9,GRIN2D,PFKM,PRKCA |
| Gαq Signaling | 1.12 | 0.0764 | -0.577 | GNA14,HRH1,ITPR2,PIK3R1,PLD1,PLD2,PRKCA,RGS2,RHOB,RHOD,RHOU,RND3 |
| Dopamine Receptor Signaling | 1.12 | 0.0909 | #NUM! | DDC,MAOA,MAOB,PPP1R10,PPP1R1B,PPP2R5B,PRKAR2A |
| Inhibition of Angiogenesis by TSP1 | 1.11 | 0.118 | #NUM! | FYN,HSPG2,JUN,TGFBR2 |
| T Cell Exhaustion Signaling Pathway | 1.1 | 0.0743 | 0.707 | ACVR1,BCL6,FOS,HLA-B,HLA-C,HLA-E,JUN,LGALS9,PIK3R1,PPP2R5B,RAP2B,SMAD3,TGFBR2 |
| Graft-versus-Host Disease Signaling | 1.1 | 0.104 | #NUM! | FAS,HLA-B,HLA-C,HLA-E,IL1RN |
| Calcium Transport I | 1.1 | 0.2 | #NUM! | ATP2A3,ATP2B1 |
| Ferroptosis Signaling Pathway | 1.09 | 0.0794 | 0 | CHAC1,DPP4,EMP1,HMGCR,HSPB1,NOX1,RAP2B,SAT1,SLC39A8,STEAP3 |
| Leukocyte Extravasation Signaling | 1.09 | 0.0725 | 0 | CD44,CLDN1,CLDN12,CLDN2,CXCR4,FER,GNAI1,MMP15,MMP7,NOX1,PIK3R1,PRKCA,RAP1GAP,TXK |
| Fcγ Receptor-mediated Phagocytosis in Macrophages and Monocytes | 1.09 | 0.0851 | 0 | FYN,GAB2,PIK3R1,PIP5K1A,PLD1,PLD2,PRKCA,RPS6KB2 |
| Cardiac Hypertrophy Signaling (Enhanced) | 1.08 | 0.0624 | 0.73 | ACVR1,ATP2A3,CLCF1,CXCL8,DVL1,EDN1,FGF19,FGFR2,GNA14,GNAI1,HDAC9,HSPB1,IL15RA,IL1R2,ITPR2,JUN,LIF,MAP3K14,PDE12,PDE9A,PIK3R1,PKN1,PRKAR2A,PRKCA,PTGS2,RAP2B,RCAN1,RPS6KB2,TGFB2,TGFBR2,WNT11 |
| Thyroid Cancer Signaling | 1.07 | 0.0886 | 1.134 | CXCL8,CXCR4,FOS,IRS2,JUN,PIK3R1,RAP2B |
| Sertoli Cell-Sertoli Cell Junction Signaling | 1.07 | 0.0722 | #NUM! | CLDN1,CLDN12,CLDN2,EPN3,JUN,MAP3K14,MPP6,PRKAR2A,RAP2B,TJP1,TJP2,TJP3,TUBA4A,TUBB2A |
| Atherosclerosis Signaling | 1.07 | 0.0787 | #NUM! | APOL1,CLU,COL18A1,CXCL8,CXCR4,IL1RN,PLAAT3,PLAAT4,SERPINA1,TNFRSF12A |
| Pyrimidine Deoxyribonucleotides De Novo Biosynthesis I | 1.06 | 0.136 | #NUM! | AK4,NME3,NME4 |
| Corticotropin Releasing Hormone Signaling | 1.05 | 0.0759 | 0 | CREB5,FOS,GLI1,GLI2,GNAI1,ITPR2,JUN,JUND,PRKAR2A,PRKCA,PTGS2 |
| Chemokine Signaling | 1.05 | 0.0875 | 0.378 | CXCR4,FOS,GNAI1,JUN,NOX1,PRKCA,RAP2B |
| Renal Cell Carcinoma Signaling | 1.05 | 0.0875 | 1.633 | ETS1,FOS,GAB1,JUN,PIK3R1,RAP2B,UBC |
| Triacylglycerol Degradation | 1.04 | 0.1 | 0.447 | AADAC,ABHD6,FAAH,LIPG,PNPLA2 |
| Amyotrophic Lateral Sclerosis Signaling | 1.03 | 0.0825 | 0 | BIRC3,CAPN5,CAPN9,CASP7,CASP9,GRIN2D,PIK3R1,RNF19A |
| Epoxysqualene Biosynthesis | 1.03 | 0.5 | #NUM! | SQLE |
| Spermine Biosynthesis | 1.03 | 0.5 | #NUM! | AMD1 |

Table S3: Pathways identification after ATRA+PDT compared to PDT.

| Ingenuity Canonical Pathways | -log(p-value) | Ratio | z-score | Molecules |
|---|---|---|---|---|
| Choline Degradation I | 1.03 | 0.5 | #NUM! | ALDH7A1 |
| Spermidine Biosynthesis I | 1.03 | 0.5 | #NUM! | AMD1 |
| Sulfate Activation for Sulfonation | 1.03 | 0.5 | #NUM! | PAPSS1 |
| GDP-L-fucose Biosynthesis I (from GDP-D-mannose) | 1.03 | 0.5 | #NUM! | TSTA3 |
| Ketogenesis | 1.02 | 0.182 | #NUM! | ACAT1,HMGCS2 |
| GDP-glucose Biosynthesis | 1.02 | 0.182 | #NUM! | HK1,PGM1 |
| Eicosanoid Signaling | 1.02 | 0.0909 | #NUM! | AKR1C3,DPEP1,PLAAT3,PLAAT4,PTGS2,TBXAS1 |
| Superpathway of Inositol Phosphate Compounds | 1.01 | 0.0704 | 1.604 | CDC25A,DUSP1,DUSP16,DUSP2,DUSP5,INPPL1,ITPKA,PIK3R1,PIP5K1A,PPFIBP2,PPP1R1B,PPP2R5B,PTPRH,RASA1 |
| UVC-Induced MAPK Signaling | 1.01 | 0.098 | 2.236 | FOS,JUN,PRKCA,RAP2B,SMPD3 |
| Apelin Endothelial Signaling Pathway | 0.996 | 0.0783 | 1 | ANGPT1,FOS,GNAI1,JUN,PIK3R1,PRKCA,RAP2B,RPS6KB2,SMAD3 |
| Signaling by Rho Family GTPases | 0.987 | 0.0672 | 0.535 | CDC42EP1,CDC42EP2,FOS,GNA14,GNAI1,GNAZ,JUN,NOX1,PIK3R1,PIP5K1A,PKN1,PLD1,RHOB,RHOD,RHOU,RND3,STMN1 |
| 3-phosphoinositide Biosynthesis | 0.987 | 0.0723 | 1.155 | CDC25A,DUSP1,DUSP16,DUSP2,DUSP5,PIK3R1,PIP5K1A,PPFIBP2,PPP1R1B,PPP2R5B,PTPRH,RASA1 |
| Role of Osteoblasts, Osteoclasts and Chondrocytes in Rheumatoid Arthritis | 0.987 | 0.0688 | #NUM! | BIRC3,BMP4,CASP9,DKK1,DVL1,FOS,GSN,IL1R2,IL1RN,JUN,MAP3K14,PIK3R1,RUNX2,SMURF1,WNT11 |
| Tumoricidal Function of Hepatic Natural Killer Cells | 0.975 | 0.125 | #NUM! | CASP7,CASP9,FAS |
| Coronavirus Pathogenesis Pathway | 0.971 | 0.0733 | 0.905 | CASP9,CXCL8,E2F7,EEF1A2,FOS,IRF7,JUN,PTGS2,SMAD3,TGFBR2,TRIM25 |
| Lymphotoxin β Receptor Signaling | 0.963 | 0.0943 | 1.342 | CASP9,CXCL1,MAP3K14,PIK3R1,TRAF4 |
| Glucose and Glucose-1-phosphate Degradation | 0.959 | 0.167 | #NUM! | HK1,PGM1 |
| Glycogen Degradation II | 0.959 | 0.167 | #NUM! | PGM1,PYGB |
| SPINK1 General Cancer Pathway | 0.947 | 0.087 | -0.816 | MT1E,MT1X,MT2A,PIK3R1,RAP2B,SPINK1 |
| p38 MAPK Signaling | 0.943 | 0.0763 | 0 | CREB5,DUSP1,FAS,HSPB1,IL1R2,IL1RN,RPS6KB2,TGFB2,TGFBR2 |
| Renin-Angiotensin Signaling | 0.943 | 0.0763 | 0.333 | FOS,ITPR2,JUN,NOX1,PIK3R1,PRKAR2A,PRKCA,RAP2B,REN |
| BMP signaling pathway | 0.943 | 0.0824 | 0.378 | BMP4,JUN,PRKAR2A,RAP2B,RUNX2,SMURF1,SOSTDC1 |
| Airway Pathology in Chronic Obstructive Pulmonary Disease | 0.943 | 0.0763 | #NUM! | AMBP,CLCF1,CXCL1,CXCL3,CXCL8,FGF19,LCN2,LIF,TGFB2 |
| IL-17 Signaling | 0.936 | 0.0695 | 3.051 | CLCF1,CXCL1,CXCL3,CXCL8,FOS,JUN,LCN2,LIF,MAP3K14,PIK3R1,PTGS2,RAP2B,TGFB2 |
| Sperm Motility | 0.924 | 0.0673 | 0.816 | CLK1,CLK4,CNGA1,EPHA2,FER,FGFR2,FYN,ITPR2,PLAAT3,PLAAT4,PRKAR2A,PRKCA,ROR1,TNK2,TXK |
| Role of Hypercytokinemia/hyperchemokinemia in the Pathogenesis of Influenza | 0.924 | 0.0814 | 2.646 | CXCL3,CXCL8,EIF2AK2,IFIT3,IL1RN,IRF7,OAS3 |
| IL-10 Signaling | 0.924 | 0.0857 | #NUM! | FOS,IL1R2,IL1RN,JUN,MAP3K14,MAP4K4 |
| Role of IL-17A in Arthritis | 0.91 | 0.0909 | #NUM! | CXCL1,CXCL3,CXCL8,PIK3R1,PTGS2 |
| Growth Hormone Signaling | 0.903 | 0.0845 | -0.447 | CEBPA,FOS,IGF2,PIK3R1,PRKCA,RPS6KB2 |
| Small Cell Lung Cancer Signaling | 0.903 | 0.0845 | 0 | CASP9,CDK6,CDKN2B,PIK3R1,PTGS2,TRAF4 |
| Estrogen-mediated S-phase Entry | 0.896 | 0.115 | #NUM! | CCNA2,CDC25A,E2F7 |
| Bile Acid Biosynthesis, Neutral Pathway | 0.896 | 0.154 | #NUM! | AKR1C1/AKR1C2,AKR1C3 |
| Apelin Liver Signaling Pathway | 0.896 | 0.115 | #NUM! | COL18A1,EDN1,FAS |
| Thrombin Signaling | 0.893 | 0.0673 | 0.577 | F2R,GATA2,GATA6,GNA14,GNAI1,GNAZ,ITPR2,PIK3R1,PRKCA,RAP2B,RHOB,RHOD,RHOU,RND3 |
| Insulin Secretion Signaling Pathway | 0.889 | 0.0656 | -2 | AGO4,CREB5,EIF4A2,FYN,GNA14,ITPR2,PIK3R1,PRKAR2A,PRKCA,RPS6KB2,SEC11C,SEC61A1,SEC61B,SEC61G,SPCS3,SSR1 |
| Erythropoietin Signaling Pathway | 0.889 | 0.0694 | 0 | BIRC3,CLCF1,CXCL8,FOS,IRS2,ITPR2,JUN,LIF,PIK3R1,PRKCA,RAP2B,TGFB2 |
| GNRH Signaling | 0.889 | 0.0694 | 1.155 | CREB5,EGR1,FOS,GNA14,GNAI1,HBEGF,ITPR2,JUN,MAP3K14,PRKAR2A,PRKCA,RAP2B |
| Basal Cell Carcinoma Signaling | 0.883 | 0.0833 | -1 | BMP4,DVL1,GLI1,GLI2,GLIS2,WNT11 |
| 3-phosphoinositide Degradation | 0.883 | 0.0705 | 1.508 | CDC25A,DUSP1,DUSP16,DUSP2,DUSP5,INPPL1,PPFIBP2,PPP1R1B,PPP2R5B,PTPRH,RASA1 |
| G Beta Gamma Signaling | 0.879 | 0.0738 | -1 | CAV1,CAV2,GNA14,GNAI1,GNAZ,HBEGF,PRKAR2A,PRKCA,RAP2B |
| Ovarian Cancer Signaling | 0.879 | 0.0719 | 0 | CD44,DVL1,EDN1,MMP7,PIK3R1,PRKAR2A,PTGS2,RAP2B,RPS6KB2,WNT11 |
| Neuregulin Signaling | 0.879 | 0.0762 | 0.707 | EREG,ERRFI1,GRB7,HBEGF,PIK3R1,PRKCA,RAP2B,RPS6KB2 |
| Acute Myeloid Leukemia Signaling | 0.87 | 0.0787 | -0.816 | CEBPA,KITLG,PIK3R1,PIM1,PML,RAP2B,RPS6KB2 |
| CTLA4 Signaling in Cytotoxic T Lymphocytes | 0.87 | 0.0787 | #NUM! | B2M,FYN,HLA-B,HLA-C,HLA-E,PIK3R1,PPP2R5B |
| Nicotine Degradation III | 0.863 | 0.0877 | 2.236 | CYP1A1,CYP2B6,CYP2S1,CYP3A5,UGT1A10 (includes others) |
| Methylglyoxal Degradation I | 0.863 | 0.333 | #NUM! | HAGHL |
| Proline Degradation | 0.863 | 0.333 | #NUM! | LOC102724788/PRODH |
| Thiosulfate Disproportionation III (Rhodanese) | 0.863 | 0.333 | #NUM! | TST |
| Glycerol-3-phosphate Shuttle | 0.863 | 0.333 | #NUM! | GPD1 |
| Adenine and Adenosine Salvage I | 0.863 | 0.333 | #NUM! | APRT |
| S-adenosyl-L-methionine Biosynthesis | 0.863 | 0.333 | #NUM! | MAT2A |
| Non-Small Cell Lung Cancer Signaling | 0.86 | 0.0822 | -0.447 | CASP9,CDK6,ITPR2,PIK3R1,PRKCA,RAP2B |
| Cdc42 Signaling | 0.851 | 0.0682 | 1.633 | B2M,CDC42EP2,EXOC8,FGD3,FOS,HLA-B,HLA-C,HLA-E,IQGAP2,JUN,RASA1,TNK2 |
| Glycogen Degradation III | 0.845 | 0.143 | #NUM! | PGM1,PYGB |
| Urate Biosynthesis/Inosine 5'-phosphate Degradation | 0.845 | 0.143 | #NUM! | IMPDH1,NT5E |
| Regulation of Cellular Mechanics by Calpain Protease | 0.842 | 0.0811 | 0 | CAPN5,CAPN9,CCNA2,CDK6,CNGA1,RAP2B |
| D-myo-inositol (1,4,5,6)-Tetrakisphosphate Biosynthesis | 0.839 | 0.0704 | 1.265 | CDC25A,DUSP1,DUSP16,DUSP2,DUSP5,PPFIBP2,PPP1R1B,PPP2R5B,PTPRH,RASA1 |
| D-myo-inositol (3,4,5,6)-tetrakisphosphate Biosynthesis | 0.839 | 0.0704 | 1.265 | CDC25A,DUSP1,DUSP16,DUSP2,DUSP5,PPFIBP2,PPP1R1B,PPP2R5B,PTPRH,RASA1 |
| Cancer Drug Resistance By Drug Efflux | 0.839 | 0.0862 | #NUM! | ABCC1,ABCC2,PIK3R1,PTGS2,RAP2B |
| CDK5 Signaling | 0.83 | 0.0741 | 0.707 | EGR1,FOSB,LAMA5,PPP1R10,PPP1R1B,PPP2R5B,PRKAR2A,RAP2B |
| Serotonin Receptor Signaling | 0.824 | 0.093 | #NUM! | DDC,HTR1D,MAOA,MAOB |
| Angiopoietin Signaling | 0.821 | 0.08 | 1 | ANGPT1,CASP9,GRB7,PIK3R1,RAP2B,RASA1 |
| Estrogen-Dependent Breast Cancer Signaling | 0.821 | 0.08 | 1.342 | AKR1C1/AKR1C2,CREB5,FOS,JUN,PIK3R1,RAP2B |
| Role of MAPK Signaling in Inhibiting the Pathogenesis of Influenza | 0.821 | 0.08 | 2.449 | CXCL8,EIF2AK2,JUN,PLAAT3,PLAAT4,PTGS2 |
| IL-1 Signaling | 0.815 | 0.0761 | 1 | FOS,GNA14,GNAI1,GNAZ,JUN,MAP3K14,PRKAR2A |
| NF-κB Signaling | 0.815 | 0.067 | 1.155 | BCL10,BMP4,CARD10,EIF2AK2,FGFR2,IL1R2,IL1RN,MAP3K14,MAP4K4,PIK3R1,RAP2B,TGFBR2 |
| Neurotrophin/TRK Signaling | 0.804 | 0.0789 | 1.633 | CREB5,FOS,GAB1,JUN,PIK3R1,RAP2B |
| Melatonin Degradation I | 0.796 | 0.0833 | 2.236 | CYP1A1,CYP2B6,CYP2S1,CYP3A5,UGT1A10 (includes others) |
| Superpathway of Citrulline Metabolism | 0.796 | 0.133 | #NUM! | ASS1,LOC102724788/PRODH |
| Dopamine-DARPP32 Feedback in cAMP Signaling | 0.793 | 0.0675 | 0.632 | ATP2A3,CAMKK1,CREB5,GNAI1,GRIN2D,ITPR2,PPP1R10,PPP1R1B,PPP2R5B,PRKAR2A,PRKCA |
| Role of NFAT in Regulation of the Immune Response | 0.79 | 0.0663 | -0.905 | FOS,FYN,GNA14,GNAI1,GNAZ,HLA-B,ITPR2,JUN,PIK3R1,RAP2B,RCAN1,RCAN3 |
| Type I Diabetes Mellitus Signaling | 0.785 | 0.0721 | 1.342 | CASP9,FAS,HLA-B,HLA-C,HLA-E,IFNGR2,IRF1,MAP3K14 |
| ErbB Signaling | 0.783 | 0.0745 | 1.89 | EREG,FOS,HBEGF,JUN,PIK3R1,PRKCA,RAP2B |
| iNOS Signaling | 0.775 | 0.0889 | #NUM! | FOS,IFNGR2,IRF1,JUN |
| Glioblastoma Multiforme Signaling | 0.77 | 0.0667 | -1 | CDK6,E2F7,IGF2,ITPR2,PIK3R1,RAP2B,RHOB,RHOD,RHOU,RND3,WNT11 |
| IL-7 Signaling Pathway | 0.767 | 0.0769 | -1 | BCL6,CDC25A,FYN,JUN,MCL1,PIK3R1 |
| PI3K/AKT Signaling | 0.757 | 0.0652 | 0 | GAB1,GAB2,GDF15,IL15RA,IL1R2,INPPL1,MCL1,PIK3R1,PPP2R5B,PTGS2,RAP2B,RPS6KB2 |
| IL-3 Signaling | 0.75 | 0.0759 | 1.633 | FOS,GAB2,JUN,PIK3R1,PRKCA,RAP2B |
| Communication between Innate and Adaptive Immune Cells | 0.75 | 0.0729 | #NUM! | B2M,CCL15,CXCL8,HLA-B,HLA-C,HLA-E,IL1RN |
| Glutaryl-CoA Degradation | 0.75 | 0.125 | #NUM! | ACAT1,CYP2S1 |
| Androgen Biosynthesis | 0.75 | 0.125 | #NUM! | AKR1C3,SRD5A3 |
| Spermine and Spermidine Degradation I | 0.747 | 0.25 | #NUM! | SAT1 |
| Catecholamine Biosynthesis | 0.747 | 0.25 | #NUM! | DDC |
| Phenylethylamine Degradation I | 0.747 | 0.25 | #NUM! | ALDH2 |
| Cardiac Hypertrophy Signaling | 0.745 | 0.0625 | 0.535 | GNA14,GNAI1,GNAZ,HSPB1,JUN,MAP3K14,PIK3R1,PRKAR2A,RAP2B,RHOB,RHOD,RHOU,RND3,TGFB2,TGFBR2 |
| NGF Signaling | 0.74 | 0.0702 | 1.134 | CREB5,GAB1,MAP3K14,PIK3R1,RAP2B,RPS6KB2,SMPD3,TRAF4 |
| Ephrin A Signaling | 0.726 | 0.0851 | #NUM! | EFNA1,EPHA2,FYN,PIK3R1 |
| Phagosome Maturation | 0.721 | 0.0662 | #NUM! | ATP6V1B1,B2M,CTSE,CTSS,HLA-B,HLA-C,HLA-E,NOX1,TUBA4A,TUBB2A |
| MSP-RON Signaling In Cancer Cells Pathway | 0.71 | 0.0672 | 2.333 | CREB5,ELF3,ETS1,FOS,JUN,KLK10,KLK6,PIK3R1,RAP2B |
| Mitochondrial L-carnitine Shuttle Pathway | 0.71 | 0.118 | #NUM! | ACSL1,CPT1A |
| Epithelial Adherens Junction Signaling | 0.71 | 0.0658 | #NUM! | ACVR1,EPN3,FER,LMO7,RAP2B,TGFB2,TGFBR2,TUBA4A,TUBB2A,ZYX |
| Adipogenesis pathway | 0.71 | 0.0672 | #NUM! | BMP4,CEBPA,CEBPD,FGFR2,HDAC9,NR1D2,PPARG,SMAD3,TXNIP |

| Ingenuity Canonical Pathways | -log(p-value) | Ratio | z-score | Molecules |
|---|---|---|---|---|
| Airway Inflammation in Asthma | 0.706 | 0.0938 | #NUM! | CXCL8,IFNGR2,TGFB2 |
| PEDF Signaling | 0.699 | 0.0732 | 0.816 | CASP7,FAS,PIK3R1,PNPLA2,PPARG,RAP2B |
| Mitochondrial Dysfunction | 0.699 | 0.0643 | #NUM! | ACO1,ATP5MC1,ATP5MC3,BACE2,CASP9,CPT1A,DHODH,MAOA,MAOB,UQCRFS1,VDAC1 |
| CD40 Signaling | 0.697 | 0.0769 | 1 | FOS,JUN,MAP3K14,PIK3R1,PTGS2 |
| Phospholipases | 0.697 | 0.0769 | 1.342 | LIPG,PLAAT3,PLAAT4,PLD1,PLD2 |
| Nicotine Degradation II | 0.697 | 0.0769 | 2.236 | CYP1A1,CYP2B6,CYP2S1,CYP3A5,UGT1A10 (includes others) |
| Human Embryonic Stem Cell Pluripotency | 0.697 | 0.0667 | #NUM! | ACVR1,BMP4,DVL1,FGFR2,PIK3R1,SMAD3,TGFB2,TGFBR2,WNT11 |
| Role of Pattern Recognition Receptors in Recognition of Bacteria and Viruses | 0.686 | 0.0649 | 1.342 | CLCF1,CXCL8,EIF2AK2,IFIH1,IRF7,LIF,OAS3,PIK3R1,PRKCA,TGFB2 |
| Androgen Signaling | 0.684 | 0.0662 | 0 | DNAJB1,GNA14,GNAI1,GNAZ,JUN,POLR2L,PRKAR2A,PRKCA,SMAD3 |
| LPS-stimulated MAPK Signaling | 0.682 | 0.0723 | 1.633 | FOS,JUN,MAP3K14,PIK3R1,PRKCA,RAP2B |
| Autoimmune Thyroid Disease Signaling | 0.682 | 0.0816 | #NUM! | FAS,HLA-B,HLA-C,HLA-E |
| mTOR Signaling | 0.678 | 0.0619 | -0.302 | DDIT4,EIF4A2,PIK3R1,PLD1,PLD2,PPP2R5B,PRKCA,RAP2B,RHOB,RHOD,RHOU,RND3,RPS6KB2 |
| Circadian Rhythm Signaling | 0.678 | 0.0909 | #NUM! | CREB5,GRIN2D,NR1D1 |
| IL-9 Signaling | 0.678 | 0.0909 | #NUM! | BCL3,IRS2,PIK3R1 |
| Pyrimidine Ribonucleotides Interconversion | 0.678 | 0.0909 | #NUM! | AK4,NME3,NME4 |
| Role of NANOG in Mammalian Embryonic Stem Cell Pluripotency | 0.672 | 0.0672 | 1 | BMP4,DVL1,GAB1,GATA6,LIF,PIK3R1,RAP2B,WNT11 |
| 1D-myo-inositol Hexakisphosphate Biosynthesis II (Mammalian) | 0.672 | 0.111 | #NUM! | INPPL1,ITPKA |
| D-myo-inositol (1,3,4)-trisphosphate Biosynthesis | 0.672 | 0.111 | #NUM! | INPPL1,ITPKA |
| Iron homeostasis signaling pathway | 0.672 | 0.0657 | #NUM! | ABCB10,ACO1,ATP6V1B1,BMP4,CIAO3,GDF15,HEPH,SMAD3,STEAP3 |
| FGF Signaling | 0.666 | 0.0714 | 1.633 | CREB5,FGF19,FGFR2,GAB1,PIK3R1,PRKCA |
| Serine Biosynthesis | 0.662 | 0.2 | #NUM! | PSPH |
| CMP-N-acetylneuraminate Biosynthesis I (Eukaryotes) | 0.662 | 0.2 | #NUM! | GNE |
| Lysine Degradation II | 0.662 | 0.2 | #NUM! | ALDH7A1 |
| Lysine Degradation V | 0.662 | 0.2 | #NUM! | ALDH7A1 |
| Trans, trans-farnesyl Diphosphate Biosynthesis | 0.662 | 0.2 | #NUM! | IDI1 |
| Citrulline-Nitric Oxide Cycle | 0.662 | 0.2 | #NUM! | ASS1 |
| Galactose Degradation I (Leloir Pathway) | 0.662 | 0.2 | #NUM! | GALM |
| Tyrosine Degradation I | 0.662 | 0.2 | #NUM! | HGD |
| Amyloid Processing | 0.662 | 0.08 | #NUM! | BACE2,CAPN5,CAPN9,PRKAR2A |
| D-myo-inositol-5-phosphate Metabolism | 0.652 | 0.0637 | 1.265 | CDC25A,DUSP1,DUSP16,DUSP2,DUSP5,PPFIBP2,PPP1R1B,PPP2R5B,PTPRH,RASA1 |
| Semaphorin Neuronal Repulsive Signaling Pathway | 0.65 | 0.0647 | -0.333 | CD44,CSPG4,FYN,NRP1,PIK3R1,PLXND1,PRKAR2A,SEMA3A,SEMA3E |
| Rac Signaling | 0.648 | 0.0661 | -0.707 | CD44,IQGAP2,JUN,NOX1,PIK3R1,PIP5K1A,PLD1,RAP2B |
| LXR/RXR Activation | 0.648 | 0.0661 | -0.378 | AMBP,APOL1,CLU,HMGCR,IL1R2,IL1RN,PTGS2,SERPINA1 |
| Hereditary Breast Cancer Signaling | 0.636 | 0.0643 | #NUM! | CDK6,FANCF,GADD45A,GADD45B,HDAC9,PIK3R1,POLR2L,RAP2B,UBC |
| Granzyme A Signaling | 0.636 | 0.105 | #NUM! | H1-0,H1-10 |
| GADD45 Signaling | 0.636 | 0.105 | #NUM! | GADD45A,GADD45B |
| Purine Nucleotides Degradation II (Aerobic) | 0.636 | 0.105 | #NUM! | IMPDH1,NT5E |
| Inhibition of ARE-Mediated mRNA Degradation Pathway | 0.635 | 0.0656 | -1.414 | AGO4,PPP2R5B,PRKAR2A,PSME1,PSME2,TIA1,ZFP36,ZFP36L1 |
| Pyrimidine Ribonucleotides De Novo Biosynthesis | 0.629 | 0.0857 | #NUM! | AK4,NME3,NME4 |
| Natural Killer Cell Signaling | 0.623 | 0.0609 | 0 | B2M,COL18A1,FYN,HLA-B,HLA-C,HLA-E,HSPA1A/HSPA1B,HSPA1L,HSPA6,MAP3K14,PIK3R1,RAP2B |
| RhoA Signaling | 0.622 | 0.065 | 0.378 | CDC42EP1,CDC42EP2,PIP5K1A,PKN1,PLD1,PLEKHG5,RND3,RTKN |
| UVB-Induced MAPK Signaling | 0.622 | 0.0769 | 1 | FOS,JUN,PIK3R1,PRKCA |
| Gap Junction Signaling | 0.613 | 0.0606 | #NUM! | CAV1,GJB2,GNAI1,ITPR2,PIK3R1,PRKAR2A,PRKCA,RAP2B,TJP1,TJP2,TUBA4A,TUBB2A |
| G-Protein Coupled Receptor Signaling | 0.611 | 0.0584 | #NUM! | CREB5,DUSP1,FYN,GNA14,GNAI1,HRH1,HTR1D,PDE12,PDE9A,PIK3R1,PRKAR2A,PRKCA,RAP1GAP,RAP2B,RASA1,RGS2 |
| Xenobiotic Metabolism General Signaling Pathway | 0.604 | 0.0629 | -0.333 | ABCA2,GSTA4,MAP3K14,NQO2,PIK3R1,PRKCA,RAP2B,UGT1A10 (includes others),UGT8 |
| Endocannabinoid Cancer Inhibition Pathway | 0.604 | 0.0629 | 1.667 | ATF3,CASP4,CASP7,CASP9,CREB5,GNAI1,PIK3R1,PRKAR2A,SMPD3 |
| CD27 Signaling in Lymphocytes | 0.604 | 0.0755 | #NUM! | CASP9,FOS,JUN,MAP3K14 |
| GP6 Signaling Pathway | 0.599 | 0.064 | -0.707 | COL17A1,COL18A1,FYN,LAMA5,LAMC2,NOX1,PIK3R1,PRKCA |
| RANK Signaling in Osteoclasts | 0.593 | 0.0674 | 0.816 | BIRC3,FOS,GSN,JUN,MAP3K14,PIK3R1 |
| Arginine Biosynthesis IV | 0.592 | 0.167 | #NUM! | ASS1 |
| Pyruvate Fermentation to Lactate | 0.592 | 0.167 | #NUM! | LDHA |
| Urea Cycle | 0.592 | 0.167 | #NUM! | ASS1 |
| Chondroitin and Dermatan Biosynthesis | 0.592 | 0.167 | #NUM! | CHPF |
| Ceramide Biosynthesis | 0.592 | 0.167 | #NUM! | DEGS2 |
| Serotonin and Melatonin Biosynthesis | 0.592 | 0.167 | #NUM! | DDC |
| Glycerol Degradation I | 0.592 | 0.167 | #NUM! | GPD1 |
| Ceramide Degradation | 0.592 | 0.167 | #NUM! | ACER2 |
| Zymosterol Biosynthesis | 0.592 | 0.167 | #NUM! | MSMO1 |
| Glycogen Biosynthesis II (from UDP-D-Glucose) | 0.592 | 0.167 | #NUM! | GYG1 |
| Cell Cycle Regulation by BTG Family Proteins | 0.583 | 0.0811 | #NUM! | BTG1,E2F7,PPP2R5B |
| Ephrin B Signaling | 0.58 | 0.0694 | -2.236 | CXCR4,EPHB6,GNA14,GNAI1,GNAZ |
| P2Y Purigenic Receptor Signaling Pathway | 0.575 | 0.063 | 1.414 | CREB5,FOS,GNAI1,JUN,PIK3R1,PRKAR2A,PRKCA,RAP2B |
| Docosahexaenoic Acid (DHA) Signaling | 0.562 | 0.0789 | #NUM! | CASP9,PIK3R1,PNPLA2 |
| Glioma Signaling | 0.558 | 0.0636 | 0 | CDK6,CDKN2B,E2F7,IGF2,PIK3R1,PRKCA,RAP2B |
| Role of MAPK Signaling in Promoting the Pathogenesis of Influenza | 0.558 | 0.0636 | 1.89 | ATP6V1B1,JUN,PLAAT3,PLAAT4,PRKCA,PTGS2,RAP2B |
| Synaptic Long Term Potentiation | 0.554 | 0.062 | 0 | CREB5,GNA14,GRIN2D,ITPR2,PPP1R10,PRKAR2A,PRKCA,RAP2B |
| p70S6K Signaling | 0.554 | 0.062 | 0 | F2R,F2RL1,GNAI1,PIK3R1,PLD1,PPP2R5B,PRKCA,RAP2B |
| CDP-diacylglycerol Biosynthesis I | 0.545 | 0.0909 | #NUM! | LPCAT1,MBOAT7 |
| Inhibition of Matrix Metalloproteases | 0.541 | 0.0769 | #NUM! | HSPG2,MMP15,MMP7 |
| IL-15 Signaling | 0.536 | 0.0667 | 0.447 | CXCL8,IL15RA,PIK3R1,RAP2B,RPS6KB2 |
| Trehalose Degradation II (Trehalase) | 0.536 | 0.143 | #NUM! | HK1 |
| Superpathway of Serine and Glycine Biosynthesis I | 0.536 | 0.143 | #NUM! | PSPH |
| Phosphatidylcholine Biosynthesis I | 0.536 | 0.143 | #NUM! | CHPT1 |
| Factors Promoting Cardiogenesis in Vertebrates | 0.532 | 0.06 | -0.333 | ACVR1,BMP4,CREB5,DKK1,DVL1,PRKCA,TGFB2,TGFBR2,WNT11 |
| Melanocyte Development and Pigmentation Signaling | 0.527 | 0.0638 | -0.816 | CREB5,KITLG,PIK3R1,PRKAR2A,RAP2B,RPS6KB2 |
| RhoGDI Signaling | 0.524 | 0.0582 | 0 | ARHGDIB,CD44,GNA14,GNAI1,GNAZ,PIP5K1A,PRKCA,RHOB,RHOD,RHOU,RND3 |
| Opioid Signaling Pathway | 0.524 | 0.0567 | 0.277 | CREB5,FOS,FOSB,FYN,GNAI1,GRIN2D,GRK5,ITPR2,PLD2,PRKAR2A,PRKCA,RAP2B,RPS6KB2,SIGMAR1 |
| Relaxin Signaling | 0.521 | 0.0596 | 0 | FOS,GNA14,GNAI1,GNAZ,JUN,PDE12,PDE9A,PIK3R1,PRKAR2A |
| NF-κB Activation by Viruses | 0.521 | 0.0658 | 1.342 | EIF2AK2,MAP3K14,PIK3R1,PRKCA,RAP2B |
| April Mediated Signaling | 0.521 | 0.075 | #NUM! | FOS,JUN,MAP3K14 |
| Differential Regulation of Cytokine Production in Intestinal Epithelial Cells by IL-17A | 0.519 | 0.087 | #NUM! | CXCL1,LCN2 |
| Superpathway of D-myo-inositol (1,4,5)-trisphosphate Metabolism | 0.519 | 0.087 | #NUM! | INPPL1,ITPKA |
| Tryptophan Degradation III (Eukaryotic) | 0.519 | 0.087 | #NUM! | ACAT1,CYP2S1 |
| α-Adrenergic Signaling | 0.516 | 0.0632 | -0.447 | GNAI1,ITPR2,PRKAR2A,PRKCA,PYGB,RAP2B |
| Th1 and Th2 Activation Pathway | 0.514 | 0.0585 | #NUM! | ACVR1,CXCR4,HLA-B,IFNGR2,IRF1,JAG2,JUN,LGALS9,PIK3R1,TGFBR2 |
| B Cell Activating Factor Signaling | 0.503 | 0.0732 | #NUM! | FOS,JUN,MAP3K14 |
| Phosphatidylglycerol Biosynthesis II (Non-plastidic) | 0.493 | 0.0833 | #NUM! | LPCAT1,MBOAT7 |
| TCA Cycle II (Eukaryotic) | 0.493 | 0.0833 | #NUM! | ACO1,SUCLA2 |
| ATM Signaling | 0.492 | 0.0619 | 0 | CDC25A,CREB5,GADD45A,GADD45B,JUN,PPP2R5B |
| STAT3 Pathway | 0.492 | 0.0593 | 0.447 | CDC25A,FGFR2,IL15RA,IL1R2,PIM1,RAP2B,TGFB2,TGFBR2 |
| Bladder Cancer Signaling | 0.492 | 0.0619 | #NUM! | CXCL8,DAPK1,FGF19,MMP15,MMP7,RAP2B |
| Maturity Onset Diabetes of Young (MODY) Signaling | 0.488 | 0.0667 | #NUM! | APOL1,APOL2,APOL6,HNF4A |

Table S3: Pathways identification after ATRA+PDT compared to PDT.

| Ingenuity Canonical Pathways | -log(p-value) | Ratio | z-score | Molecules |
|---|---|---|---|---|
| Sphingosine and Sphingosine-1-phosphate Metabolism | 0.488 | 0.125 | #NUM! | ACER2 |
| Sphingomyelin Metabolism | 0.488 | 0.125 | #NUM! | SMPD3 |
| Breast Cancer Regulation by Stathmin1 | 0.485 | 0.0525 | 3.772 | ADGRF1,ADGRG5,AURKA,CDK6,CELSR1,CREB5,E2F7,F2R,F2RL1,GPR158,GPR160,GPR35,GPR87,GPRC5A,HCAR1,HRH1,HTR1D,JUN,LGR4,NMUR2,PIK3R1,PPP1R10,PPP2R5B,PRKAR2A,PRKCA,RAP2B,SPDEF,STMN1,TGFB2,TUBA4A,TUBB2A |
| MIF Regulation of Innate Immunity | 0.484 | 0.0714 | #NUM! | FOS,JUN,PTGS2 |
| BEX2 Signaling Pathway | 0.483 | 0.0633 | -0.447 | BBC3,ITGB6,JUN,PPP2R5B,SWAP70 |
| Role of NFAT in Cardiac Hypertrophy | 0.479 | 0.0561 | -0.302 | GNAI1,HDAC9,ITPR2,LIF,PIK3R1,PRKAR2A,PRKCA,RAP2B,RCAN1,RCAN3,TGFB2,TGFBR2 |
| IL-2 Signaling | 0.474 | 0.0656 | #NUM! | FOS,JUN,PIK3R1,RAP2B |
| Nitric Oxide Signaling in the Cardiovascular System | 0.47 | 0.0606 | 0 | ATP2A3,CAV1,ITPR2,PIK3R1,PRKAR2A,PRKCA |
| Role of MAPK Signaling in the Pathogenesis of Influenza | 0.47 | 0.0625 | #NUM! | PLAAT3,PLAAT4,PRKCA,PTGS2,RAP2B |
| Role of Macrophages, Fibroblasts and Endothelial Cells in Rheumatoid Arthritis | 0.469 | 0.0541 | #NUM! | CEBPA,CEBPD,CREB5,CXCL8,DKK1,DVL1,F2RL1,FOS,IL1R2,IL1RN,JUN,MAP3K14,PIK3R1,PRKCA,RAP2B,TRAF4,WNT11 |
| Necroptosis Signaling Pathway | 0.467 | 0.0573 | -0.333 | BIRC3,CAPN5,CAPN9,DAPK1,EIF2AK2,FAS,PPIF,UBC,VDAC1 |
| Role of IL-17F in Allergic Inflammatory Airway Diseases | 0.467 | 0.0698 | #NUM! | CREB5,CXCL1,CXCL8 |
| Wnt/Ca+ pathway | 0.46 | 0.0645 | 1 | CREB5,DVL1,PRKCA,ROR1 |
| Actin Nucleation by ARP-WASP Complex | 0.458 | 0.0617 | 0.447 | RAP2B,RHOB,RHOD,RHOU,RND3 |
| Role of RIG1-like Receptors in Antiviral Innate Immunity | 0.451 | 0.0682 | #NUM! | IFIH1,IRF7,TRIM25 |
| Apelin Pancreas Signaling Pathway | 0.451 | 0.0682 | #NUM! | ERN1,PIK3R1,PRKAR2A |
| Neuropathic Pain Signaling In Dorsal Horn Neurons | 0.449 | 0.0594 | 0 | FOS,GRIN2D,ITPR2,PIK3R1,PRKAR2A,PRKCA |
| Sucrose Degradation V (Mammalian) | 0.446 | 0.111 | #NUM! | GALM |
| Citrulline Biosynthesis | 0.446 | 0.111 | #NUM! | LOC102724788/PRODH |
| CD28 Signaling in T Helper Cells | 0.441 | 0.0579 | 0 | BCL10,FOS,FYN,HLA-B,ITPR2,JUN,PIK3R1 |
| Acute Phase Response Signaling | 0.439 | 0.0556 | 2.333 | AMBP,CRABP2,FOS,IL1RN,JUN,MAP3K14,PIK3R1,RAP2B,SERPINA1,SERPINA3 |
| SAPK/JNK Signaling | 0.438 | 0.0588 | 0.816 | GAB1,GADD45A,JUN,MAP4K4,PIK3R1,RAP2B |
| Role of PI3K/AKT Signaling in the Pathogenesis of Influenza | 0.433 | 0.0625 | #NUM! | CASP9,GNAI1,PIK3R1,PLAC8 |
| Cardiac β-adrenergic Signaling | 0.428 | 0.0563 | -1.134 | AKAP12,ATP2A3,PDE12,PDE9A,PKIB,PPP1R10,PPP2R5B,PRKAR2A |
| Mouse Embryonic Stem Cell Pluripotency | 0.427 | 0.0583 | -0.816 | BMP4,DVL1,ID2,LIF,PIK3R1,RAP2B |
| Regulation of Actin-based Motility by Rho | 0.427 | 0.0583 | 0.447 | GSN,PIP5K1A,RHOB,RHOD,RHOU,RND3 |
| VEGF Family Ligand-Receptor Interactions | 0.424 | 0.0595 | 1 | FOS,NRP1,PIK3R1,PRKCA,RAP2B |
| IL-4 Signaling | 0.412 | 0.0588 | #NUM! | HLA-B,INPPL1,PIK3R1,RAP2B,RPS6KB2 |
| Ketolysis | 0.41 | 0.1 | #NUM! | ACAT1 |
| Dendritic Cell Maturation | 0.409 | 0.0543 | 1 | B2M,CD58,COL18A1,CREB5,HLA-B,HLA-C,HLA-E,IL1RN,MAP3K14,PIK3R1 |
| Calcium-induced T Lymphocyte Apoptosis | 0.408 | 0.0606 | 0 | ATP2A3,HLA-B,ITPR2,PRKCA |
| Allograft Rejection Signaling | 0.401 | 0.0581 | #NUM! | B2M,FAS,HLA-B,HLA-C,HLA-E |
| Calcium Signaling | 0.396 | 0.0534 | -0.707 | ATP2A3,ATP2B1,CAMKK1,CREB5,GRIN2D,HDAC9,ITPR2,PRKAR2A,RAP2B,RCAN1,RCAN3 |
| Chondroitin Sulfate Biosynthesis (Late Stages) | 0.39 | 0.0625 | #NUM! | CHPF,CHST4,SULT2B1 |
| Telomerase Signaling | 0.389 | 0.0561 | 0.447 | ELF3,ETS1,HDAC9,PIK3R1,PPP2R5B,RAP2B |
| 14-3-3-mediated Signaling | 0.388 | 0.0551 | 1.342 | FOS,JUN,PIK3R1,PRKCA,RAP2B,TUBA4A,TUBB2A |
| Remodeling of Epithelial Adherens Junctions | 0.384 | 0.0588 | #NUM! | MAPRE3,TUBA4A,TUBB2A,ZYX |
| Endocannabinoid Neuronal Synapse Pathway | 0.38 | 0.0547 | 0.447 | ABHD6,FAAH,GNA14,GNAI1,GRIN2D,PRKAR2A,PTGS2 |
| Purine Nucleotides De Novo Biosynthesis II | 0.379 | 0.0909 | #NUM! | IMPDH1 |
| cAMP-mediated signaling | 0.378 | 0.0524 | -0.302 | AKAP12,CNGA1,CREB5,DUSP1,GNAI1,HTR1D,PDE12,PDE9A,PKIB,PRKAR2A,RAP1GAP,RGS2 |
| Hematopoiesis from Pluripotent Stem Cells | 0.377 | 0.0612 | #NUM! | CXCL8,KITLG,LIF |
| Cell Cycle: G2/M DNA Damage Checkpoint Regulation | 0.377 | 0.0612 | #NUM! | AURKA,GADD45A,HIPK2 |
| Role of JAK1 and JAK3 in γc Cytokine Signaling | 0.373 | 0.058 | #NUM! | IL15RA,IRS2,PIK3R1,RAP2B |
| White Adipose Tissue Browning Pathway | 0.372 | 0.0543 | 0.816 | CREB5,FGFR2,LDHA,PPARG,PRKAR2A,RARG,SLC16A1 |
| Crosstalk between Dendritic Cells and Natural Killer Cells | 0.372 | 0.0562 | #NUM! | FAS,HLA-B,HLA-C,HLA-E,IL15RA |
| OX40 Signaling Pathway | 0.362 | 0.0556 | #NUM! | B2M,HLA-B,HLA-C,HLA-E,JUN |
| GM-CSF Signaling | 0.362 | 0.0571 | #NUM! | ETS1,PIK3R1,PIM1,RAP2B |
| Assembly of RNA Polymerase I Complex | 0.351 | 0.0833 | #NUM! | TAF1A |
| Hematopoiesis from Multipotent Stem Cells | 0.351 | 0.0833 | #NUM! | KITLG |
| 4-1BB Signaling in T Lymphocytes | 0.338 | 0.0625 | #NUM! | JUN,MAP3K14 |
| GPCR-Mediated Integration of Enteroendocrine Signaling Exemplified by an L Cell | 0.331 | 0.0548 | 0 | GNA14,GNAI1,ITPR2,PRKAR2A |
| T Helper Cell Differentiation | 0.331 | 0.0548 | #NUM! | BCL6,HLA-B,IFNGR2,TGFBR2 |
| PKCθ Signaling in T Lymphocytes | 0.33 | 0.0516 | 1.414 | BCL10,FOS,FYN,HLA-B,JUN,MAP3K14,PIK3R1,RAP2B |
| Fatty Acid Activation | 0.326 | 0.0769 | #NUM! | ACSL1 |
| Guanosine Nucleotides Degradation III | 0.326 | 0.0769 | #NUM! | NT5E |
| UDP-N-acetyl-D-galactosamine Biosynthesis II | 0.326 | 0.0769 | #NUM! | HK1 |
| CCR5 Signaling in Macrophages | 0.325 | 0.0532 | 1 | FAS,FOS,GNAI1,JUN,PRKCA |
| Fatty Acid β-oxidation I | 0.323 | 0.0606 | #NUM! | ACAA1,ACSL1 |
| Endocannabinoid Developing Neuron Pathway | 0.321 | 0.0522 | 0.447 | CREB5,GNAI1,PIK3R1,PRKAR2A,RAP1GAP,RAP2B |
| Hypoxia Signaling in the Cardiovascular System | 0.321 | 0.0541 | #NUM! | CREB5,EDN1,JUN,LDHA |
| Th2 Pathway | 0.319 | 0.0515 | -1.342 | ACVR1,CXCR4,HLA-B,JAG2,JUN,PIK3R1,TGFBR2 |
| PTEN Signaling | 0.319 | 0.0515 | 0.816 | CASP9,FGFR2,INPPL1,PIK3R1,RAP2B,RPS6KB2,TGFBR2 |
| Transcriptional Regulatory Network in Embryonic Stem Cells | 0.316 | 0.0556 | #NUM! | GATA6,HNF4A,ZFHX3 |
| fMLP Signaling in Neutrophils | 0.314 | 0.0517 | -0.816 | GNAI1,ITPR2,NOX1,PIK3R1,PRKCA,RAP2B |
| Role of JAK2 in Hormone-like Cytokine Signaling | 0.31 | 0.0588 | #NUM! | IRS2,SH2B3 |
| Fc Epsilon RI Signaling | 0.307 | 0.0513 | 0 | FYN,GAB1,INPPL1,PIK3R1,PRKCA,RAP2B |
| CREB Signaling in Neurons | 0.304 | 0.0487 | 1.961 | ADGRF1,ADGRG5,CELSR1,CREB5,F2R,F2RL1,FGFR2,GNA14,GNAI1,GNAZ,GPR158,GPR160,GPR35,GPR87,GPRC5A,GRIN2D,HCAR1,HRH1,HTR1D,ITPR2,LGR4,NMUR2,PIK3R1,POLR2L,PRKAR2A,PRKCA,RAP2B,TGFB2,TGFBR2 |
| Leukotriene Biosynthesis | 0.303 | 0.0714 | #NUM! | DPEP1 |
| Colanic Acid Building Blocks Biosynthesis | 0.303 | 0.0714 | #NUM! | TSTA3 |
| Macropinocytosis Signaling | 0.303 | 0.0526 | #NUM! | ITGB6,PIK3R1,PRKCA,RAP2B |
| Nucleotide Excision Repair Pathway | 0.296 | 0.0571 | #NUM! | POLR2L,XPA |
| Chondroitin Sulfate Biosynthesis | 0.294 | 0.0536 | #NUM! | CHPF,CHST4,SULT2B1 |
| Phospholipase C Signaling | 0.29 | 0.0489 | 0.632 | CREB5,FYN,HDAC9,ITPR2,PLD1,PLD2,PRKCA,RAP2B,RHOB,RHOD,RHOU,RND3,TGM2 |
| CNTF Signaling | 0.284 | 0.0526 | #NUM! | PIK3R1,RAP2B,RPS6KB2 |
| Role of CHK Proteins in Cell Cycle Checkpoint Control | 0.284 | 0.0526 | #NUM! | CDC25A,E2F7,PPP2R5B |
| Type II Diabetes Mellitus Signaling | 0.279 | 0.0493 | -0.447 | ACSL1,IRS2,MAP3K14,PIK3R1,PPARG,PRKCA,SMPD3 |
| MSP-RON Signaling Pathway | 0.275 | 0.0517 | #NUM! | KLK10,KLK6,PIK3R1 |
| FLT3 Signaling in Hematopoietic Progenitor Cells | 0.268 | 0.05 | 1 | CREB5,GAB2,PIK3R1,RAP2B |
| JAK/Stat Signaling | 0.268 | 0.05 | 1 | FOS,JUN,PIK3R1,RAP2B |
| Dermatan Sulfate Biosynthesis | 0.265 | 0.0508 | #NUM! | CHPF,CHST4,SULT2B1 |
| Granzyme B Signaling | 0.264 | 0.0625 | #NUM! | CASP9 |
| Chondroitin Sulfate Degradation (Metazoa) | 0.264 | 0.0625 | #NUM! | CEMIP |
| Adenosine Nucleotides Degradation II | 0.264 | 0.0625 | #NUM! | NT5E |
| Parkinson's Signaling | 0.264 | 0.0625 | #NUM! | CASP9 |
| Endometrial Cancer Signaling | 0.256 | 0.05 | #NUM! | CASP9,PIK3R1,RAP2B |
| Gαi Signaling | 0.253 | 0.048 | 0.447 | CAV1,GNAI1,HTR1D,PRKAR2A,RAP1GAP,RAP2B |
| FAK Signaling | 0.249 | 0.0481 | #NUM! | CAPN5,CAPN9,FYN,PIK3R1,RAP2B |
| Autophagy | 0.247 | 0.0492 | #NUM! | CTSE,CTSS,MAP1LC3B |
| Isoleucine Degradation I | 0.246 | 0.0588 | #NUM! | ACAT1 |
| γ-linolenate Biosynthesis II (Animals) | 0.246 | 0.0588 | #NUM! | ACSL1 |
| Dermatan Sulfate Degradation (Metazoa) | 0.246 | 0.0588 | #NUM! | CEMIP |
| D-myo-inositol (1,4,5)-trisphosphate Degradation | 0.246 | 0.0588 | #NUM! | INPPL1 |

Table S3: Pathways identification after ATRA+PDT compared to PDT.

| Ingenuity Canonical Pathways | -log(p-value) | Ratio | z-score | Molecules |
|---|---|---|---|---|
| Differential Regulation of Cytokine Production in Macrophages and T Helper Cells by | 0.231 | 0.0556 | #NUM! | CXCL1 |
| Cardiomyocyte Differentiation via BMP Receptors | 0.203 | 0.05 | #NUM! | BMP4 |
| HOTAIR Regulatory Pathway | 0 | 0.0437 | -1.89 | AGO4,CD44,IRF1,MMP15,MMP7,PIK3R1,WNT11 |
| Synaptogenesis Signaling Pathway | 0 | 0.0449 | -1.069 | CREB5,EFNA1,EPHA2,EPHB6,FYN,GRIN2D,LRP8,PIK3R1,PRKAR2A,RAP2B,RPS6KB2,SNCG,STX1A,SYT7 |
| Antioxidant Action of Vitamin C | 0 | 0.0459 | -1 | PLAAT3,PLAAT4,PLD1,PLD2,SLC23A3 |
| Apelin Cardiomyocyte Signaling Pathway | 0 | 0.0404 | -1 | ATP2A3,GNAI1,PIK3R1,PRKCA |
| Reelin Signaling in Neurons | 0 | 0.041 | -0.447 | FYN,GRIN2D,LRP8,PDK4,PIK3R1 |
| GPCR-Mediated Nutrient Sensing in Enteroendocrine Cells | 0 | 0.0446 | -0.447 | GNA14,GNAI1,ITPR2,PRKAR2A,PRKCA |
| Synaptic Long Term Depression | 0 | 0.0476 | -0.333 | GNA14,GNAI1,GNAZ,ITPR2,PLAAT3,PLAAT4,PPP2R5B,PRKCA,RAP2B |
| EIF2 Signaling | 0 | 0.0357 | 0 | AGO4,ATF3,EIF2AK2,EIF4A2,PIK3R1,PPP1R15A,RAP2B,WARS1 |
| Th1 Pathway | 0 | 0.0413 | 0 | HLA-B,IFNGR2,IRF1,LGALS9,PIK3R1 |
| Adrenomedullin signaling pathway | 0 | 0.0406 | 0 | FOS,GNA14,IL1RN,ITPR2,PIK3R1,PPARG,PRKAR2A,RAP2B |
| Gα12/13 Signaling | 0 | 0.0458 | 0.816 | F2R,F2RL1,JUN,PIK3R1,RAP2B,RASA1 |
| Actin Cytoskeleton Signaling | 0 | 0.0352 | 1.89 | F2R,FGD3,FGF19,GSN,IQGAP2,PIK3R1,PIP5K1A,RAP2B |
| Role of BRCA1 in DNA Damage Response | 0 | 0.0375 | #NUM! | E2F7,FANCF,GADD45A |
| Activation of IRF by Cytosolic Pattern Recognition Receptors | 0 | 0.0476 | #NUM! | IFIH1,IRF7,JUN |
| Clathrin-mediated Endocytosis Signaling | 0 | 0.0466 | #NUM! | AAK1,APOL1,CLU,F2R,FGF19,ITGB6,PIK3R1,SERPINA1,UBC |
| TREM1 Signaling | 0 | 0.04 | #NUM! | CXCL3,CXCL8,LAT2 |
| FcγRIIB Signaling in B Lymphocytes | 0 | 0.0267 | #NUM! | PIK3R1,RAP2B |
| CCR3 Signaling in Eosinophils | 0 | 0.0403 | #NUM! | GNAI1,ITPR2,PIK3R1,PRKCA,RAP2B |
| Oncostatin M Signaling | 0 | 0.0465 | #NUM! | MT2A,RAP2B |
| Role of Cytokines in Mediating Communication between Immune Cells | 0 | 0.037 | #NUM! | CXCL8,IL1RN |
| Mechanisms of Viral Exit from Host Cells | 0 | 0.0244 | #NUM! | PRKCA |
| Melatonin Signaling | 0 | 0.0417 | #NUM! | GNAI1,PRKAR2A,PRKCA |
| Cellular Effects of Sildenafil (Viagra) | 0 | 0.0153 | #NUM! | ITPR2,PRKAR2A |
| Agrin Interactions at Neuromuscular Junction | 0 | 0.0429 | #NUM! | DVL1,JUN,RAP2B |
| iCOS-iCOSL Signaling in T Helper Cells | 0 | 0.045 | #NUM! | GAB2,HLA-B,ITPR2,PIK3R1,PLEKHA2 |
| Lipid Antigen Presentation by CD1 | 0 | 0.0385 | #NUM! | B2M |
| Mitotic Roles of Polo-Like Kinase | 0 | 0.0455 | #NUM! | CDC25A,PLK3,PPP2R5B |
| DNA Methylation and Transcriptional Repression Signaling | 0 | 0.0286 | #NUM! | DNMT3A |
| Antiproliferative Role of Somatostatin Receptor 2 | 0 | 0.026 | #NUM! | PIK3R1,RAP2B |
| Role of Oct4 in Mammalian Embryonic Stem Cell Pluripotency | 0 | 0.0435 | #NUM! | CCNF,NR5A2 |
| Melanoma Signaling | 0 | 0.04 | #NUM! | PIK3R1,RAP2B |
| Prostate Cancer Signaling | 0 | 0.044 | #NUM! | CASP9,CREB5,PIK3R1,RAP2B |
| Primary Immunodeficiency Signaling | 0 | 0.02 | #NUM! | TAP2 |
| Systemic Lupus Erythematosus Signaling | 0 | 0.0349 | #NUM! | FOS,HLA-B,HLA-C,HLA-E,IL1RN,JUN,PIK3R1,RAP2B |
| PAK Signaling | 0 | 0.0189 | #NUM! | PIK3R1,RAP2B |
| Altered T Cell and B Cell Signaling in Rheumatoid Arthritis | 0 | 0.0444 | #NUM! | FAS,HLA-B,IL1RN,MAP3K14 |
| Regulation of eIF4 and p70S6K Signaling | 0 | 0.0301 | #NUM! | AGO4,EIF4A2,PIK3R1,PPP2R5B,RAP2B |
| Leptin Signaling in Obesity | 0 | 0.027 | #NUM! | PIK3R1,PRKAR2A |
| B Cell Development | 0 | 0.0278 | #NUM! | HLA-B |
| Role of Wnt/GSK-3β Signaling in the Pathogenesis of Influenza | 0 | 0.0385 | #NUM! | DVL1,NCOA3,WNT11 |
| Nur77 Signaling in T Lymphocytes | 0 | 0.0441 | #NUM! | CASP9,HDAC9,PRKCA |
| MIF-mediated Glucocorticoid Regulation | 0 | 0.0294 | #NUM! | PTGS2 |
| Cell Cycle Control of Chromosomal Replication | 0 | 0.0179 | #NUM! | CDK6 |
| Assembly of RNA Polymerase II Complex | 0 | 0.02 | #NUM! | POLR2L |
| IL-17A Signaling in Airway Cells | 0 | 0.0462 | #NUM! | CXCL1,CXCL3,PIK3R1 |
| Role of JAK1, JAK2 and TYK2 in Interferon Signaling | 0 | 0.0417 | #NUM! | IFNGR2 |
| Paxillin Signaling | 0 | 0.0278 | #NUM! | ITGB6,PIK3R1,RAP2B |
| nNOS Signaling in Skeletal Muscle Cells | 0 | 0.0244 | #NUM! | SNTB1 |
| ErbB2-ErbB3 Signaling | 0 | 0.0462 | #NUM! | JUN,PIK3R1,RAP2B |
| ErbB4 Signaling | 0 | 0.0448 | #NUM! | PIK3R1,PRKCA,RAP2B |
| Netrin Signaling | 0 | 0.0308 | #NUM! | ABLIM1,PRKAR2A |
| Heparan Sulfate Biosynthesis | 0 | 0.038 | #NUM! | AADAC,CHST4,SULT2B1 |
| Thyroid Hormone Metabolism II (via Conjugation and/or Degradation) | 0 | 0.0263 | #NUM! | UGT1A10 (includes others) |
| Heparan Sulfate Biosynthesis (Late Stages) | 0 | 0.0417 | #NUM! | AADAC,CHST4,SULT2B1 |
| tRNA Splicing | 0 | 0.0455 | #NUM! | PDE12,PDE9A |
| NAD Salvage Pathway II | 0 | 0.0385 | #NUM! | NT5E |
| D-myo-inositol (1,4,5)-Trisphosphate Biosynthesis | 0 | 0.04 | #NUM! | PIP5K1A |
| Glutathione-mediated Detoxification | 0 | 0.0312 | #NUM! | GSTA4 |
| tRNA Charging | 0 | 0.0256 | #NUM! | WARS1 |
| Dermatan Sulfate Biosynthesis (Late Stages) | 0 | 0.0435 | #NUM! | CHST4,SULT2B1 |
| Glycolysis I | 0 | 0.0385 | #NUM! | PFKM |
| Gluconeogenesis I | 0 | 0.0385 | #NUM! | ME1 |
| Methionine Degradation I (to Homocysteine) | 0 | 0.0455 | #NUM! | MAT2A |
| Superpathway of Methionine Degradation | 0 | 0.027 | #NUM! | MAT2A |
| Gαs Signaling | 0 | 0.0374 | #NUM! | CNGA1,CREB5,PRKAR2A,RGS2 |
| Cysteine Biosynthesis III (mammalia) | 0 | 0.0417 | #NUM! | MAT2A |
| Role of p14/p19ARF in Tumor Suppression | 0 | 0.0345 | #NUM! | PIK3R1 |
| Vitamin-C Transport | 0 | 0.0417 | #NUM! | GJB2 |
| Oxidative Phosphorylation | 0 | 0.0275 | #NUM! | ATP5MC1,ATP5MC3,UQCRFS1 |
| GABA Receptor Signaling | 0 | 0.0211 | #NUM! | GABRP,UBC |
| Phototransduction Pathway | 0 | 0.0385 | #NUM! | CNGA1,PRKAR2A |
| Glutamate Receptor Signaling | 0 | 0.0175 | #NUM! | GRIN2D |
| Notch Signaling | 0 | 0.027 | #NUM! | JAG2 |
| VEGF Signaling | 0 | 0.0303 | #NUM! | PIK3R1,PRKCA,RAP2B |
| Gustation Pathway | 0 | 0.0323 | #NUM! | ASIC1,ITPR2,PDE12,PDE9A,PRKAR2A |
| Phagosome Formation | 0 | 0.0451 | #NUM! | PIK3R1,PRKCA,RHOB,RHOD,RHOU,RND3 |
| NER Pathway | 0 | 0.0194 | #NUM! | POLR2L,XPA |
| Apelin Cardiac Fibroblast Signaling Pathway | 0 | 0.0435 | #NUM! | TGFB2 |
| Apelin Adipocyte Signaling Pathway | 0 | 0.0366 | #NUM! | GNAI1,NOX1,PRKAR2A |
| IL-23 Signaling Pathway | 0 | 0.0227 | #NUM! | PIK3R1 |
| Kinetochore Metaphase Signaling Pathway | 0 | 0.0297 | #NUM! | INCENP,PPP1R10,PPP2R5B |
| Complement System | 0 | 0.027 | #NUM! | CD55 |
