## Supplementary Table 4 for "All-*trans* retinoic acid enhances the anti-tumour effects of fimaporfin-based photodynamic therapy"

Supplementary Table 4. Upstream regulator analysis based gene set enrichment analysis (GSEA) of differentially expressed genes (DEGs) after ATRA+PDT compared to PDT

© 2000-2021 QIAGEN. All rights reserved.

| Upstream Regulator | Expr Log Ratio | Molecule Type | Predicted Activation State | Activation z-score | p-value of overlap | Target Molecules in Dataset | Mechanistic Network |
|---|---|---|---|---|---|---|---|
| TNF | | cytokine | Activated | 7.478 | 1.36E-44 | K2,ELF3,EMP1,ENG,ENTPD5,EPHA2,EREG,ERN1,ETS1,ETV6,F2RL1,FAS,FBXO32,FGFR2,FOS,FOSB,FOSL1,FUT4,FYN,GAB1,GADD45A,GADD45B,GASK1B,GATA2,GBP3,GDF15 | 498 (19) |
| IFNG | | cytokine | Activated | 7.045 | 1.95E-35 | R158,GSDMD,HBEGF,HDAC9,HERC6,HLA-B,HLA-C,HLA-E,HMGCR,HSPA1A/HSPA1B,HSPA1L,HSPB1,HSPG2,ID2,IDI1,IFI35,IFI44,IFIH1,IFIT3,IFITM2,IFITM3,IFNGR2,IL15RA,IL1RN | 510 (19) |
| poly rI:rC-RNA | | biologic drug | Activated | 6.103 | 1.85E-16 | DF15,GLI1,GNPAT,HBEGF,HERC6,HLA-B,HLA-C,HLA-E,HLA-J,IFI44,IFIH1,IFIT3,IFITM3,IGFBP6,IL15RA,IQGAP2,IRF1,IRF7,JUN,KLF4,KLF6,LGALS9,LIF,LIPG,MYB,NAAA,NCOA7,NE | 493 (19) |
| lipopolysaccharide | | chemical drug | Activated | 5.841 | 1.16E-39 | TV6,F2R,F2RL1,FAAH,FAS,FBXO32,FEM1C,FOS,FOSB,FOXL1,FUT8,FYN,GAB1,GADD45A,GADD45B,GATA2,GBP3,GJB2,GLCCI1,GLIS2,GNPAT,GPC4,GPD1,GPR35,GSN,H1-10,H | 630 (20) |
| NUPR1 | | transcription regulator | Activated | 5.347 | 9.61E-18 | 2,GADD45A,GCNT2,GDF15,GNE,GPCPD1,GSTA4,HBEGF,HJURP,IRS2,KLF4,KLF6,LIF,LRP8,MAT2A,MT1X,MXD1,NAAA,NR1D1,NR1D2,NRSN2,PHLDA1,PIK3R1,PIM1,PLK3,PMPCA | 424 (7) |
| Interferon alpha | | group | Activated | 5.255 | 1.01E-15 | SP3,GLI1,HERC6,HLA-B,HLA-C,HSPA1A/HSPA1B,IFI35,IFI44,IFIH1,IFIT3,IFITM2,IFITM3,IL15RA,IL1RN,IRF1,IRF7,LGALS9,LIF,MCL1,NABP1,NMI,NT5E,OAS3,PARP10,PARP12,PARI | 368 (16) |
| tretinoin | | chemical - endogenous mammalian | Activated | 5.134 | 1.04E-21 | HOXA3,HOXA5,HSPB1,ID2,IFI35,IFI44,IFIH1,IFIT3,IGF2,IGFBP2,IGFBP6,IRF1,ITM2B,ITPR2,JUN,KANSL3,KITLG,KLF4,KLF9,KRT20,KRT7,LATS2,LCN2,LGALS3BP,LGALS9,MAOB,M | 550 (20) |
| IRF7 | 0.382 | transcription regulator | Activated | 5.114 | 8.36E-08 | DNAJA1,GBP3,IFI35,IFI44,IFIH1,IFIT3,IFITM2,IFITM3,IL15RA,IRF1,IRF7,MCL1,NMI,OAS3,PARP12,PARP14,PLAC8,PLSCR1,PMAIP1,PSMB10,PSMB8,PSME1,PSME2,TAP2,TRIM21,Z | 220 (13) |
| ERK | | group | Activated | 5.038 | 1.35E-09 | CXCL8,CYP1A1,DUSP1,EDN1,EGR1,EIF2AK2,EPHA2,EREG,ETS1,FAS,FOS,FOSL1,GDF15,HBEGF,JUN,LCN2,LIF,MAFF,MAP4K4,MCAM,MCL1,NFKBIZ,PDK4,PMAIP1,PTGS2,RUN | 483 (21) |
| F2 | | peptidase | Activated | 4.98 | 3.34E-17 | CL1,CXCL3,CXCL8,CXCR4,CYP2B6,DST,DUSP1,E2F7,EDN1,EGR1,EREG,ETS1,F2R,F2RL1,FOS,FOSB,FOSL1,HBEGF,HDAC9,IL15RA,JUN,KLF6,MTSS1,NCOA7,PGM2L1,PMAIP1, | 442 (20) |
| IFNL1 | | cytokine | Activated | 4.927 | 2.84E-14 | 6,ATF3,CHAC1,CXCL8,DDX60L,EIF2AK2,F2R,HERC6,HLA-B,HLA-C,IFI35,IFI44,IFIH1,IFIT3,IFITM2,IFITM3,LGALS3BP,OAS3,PLAAT4,PLSCR1,PML,SAMD9,SHFL,SP100,SP110,ZC3 | 330 (13) |
| tetradecanoylphorbol acetate | | chemical drug | Activated | 4.881 | 5.76E-23 | TA2,GDF15,GNAZ,GPR35,GPR87,GPRC5A,GRK5,GSDMB,H1-10,HBEGF,HJURP,HMGA2,HNF4A,HOXA5,HRH1,HTR1D,ID2,IFI35,IFNGR2,IGF2,IGFBP2,IGFBP6,IL1RN,IRF1,IRF7,JC | 510 (19) |
| OSM | | cytokine | Activated | 4.768 | 1.1E-14 | 1,DHRS3,DNAJC3,ECM1,EGR1,EPHB6,FOS,FOSB,FOSL1,GAB1,HBEGF,HLA-B,HLA-C,ID2,IFI35,IGFBP6,IL15RA,IL1R2,IRF1,IRF7,JMJD1C,JUN,LCN2,LIF,MAOA,MT1X,MT2A,NEDD | 450 (21) |
| STAT1 | | transcription regulator | Activated | 4.711 | 9.63E-15 | P3A5,DPP4,EDN1,EGR1,EIF2AK2,FAS,FOS,GBP3,HERC6,HLA-E,IFI35,IFI44,IFIH1,IFIT3,IFITM2,IFITM3,IL15RA,IRF1,IRF7,JUN,KLF4,LCN2,NOX1,OAS3,PIM1,PLSCR1,PMAIP1,PPAR | 483 (20) |
| IFNA2 | | cytokine | Activated | 4.582 | 7.31E-14 | 2AK2,FAS,HBEGF,HERC6,HLA-B,HLA-C,HLA-E,HSPA6,IFI35,IFI44,IFIH1,IFIT3,IFITM2,IFITM3,IRF1,IRF7,LGALS3BP,MT1X,MT2A,NPDC1,NT5E,OAS3,PARP12,PLAAT4,PLSCR1,PM | 455 (20) |
| IL1B | | cytokine | Activated | 4.561 | 3.58E-27 | ADD45B,GBP3,GDF15,GNPAT,GRB7,H1-10,HBEGF,HLA-E,HNF4A,HSPA1A/HSPA1B,HSPA1L,HSPB1,HSPG2,ID2,IER5L,IFIT3,IGFBP6,IL15RA,IL1R2,IL1RN,IRF1,IRF7,IRS2,JUN,JUN | 498 (16) |
| NFkB (complex) | | complex | Activated | 4.545 | 4.06E-17 | P2,DUSP5,E2F7,EDN1,EFNA1,EGR1,EHF,ELF3,ERAP2,ETS1,F2RL1,FAS,FOS,GADD45A,GADD45B,GDF15,GLI2,GNPAT,GRK5,HDAC9,HNF4A,HSPA1L,HSPB1,IGFBP2,IL15RA,IL1F | 463 (17) |
| TGM2 | 0.75 | enzyme | Activated | 4.276 | 0.00000237 | RC3,CDK6,CEACAM1,CSRNP1,CXCL8,DDX60L,HLA-B,IFI35,IFIT3,LGALS9,MPP6,MT2A,OAS3,PARP14,PHLDA1,PIM1,PLAAT4,PLSCR1,RNF213,RUNX2,SEMA7A,SLC16A1,SLFN5,S | 435 (11) |
| PDGF BB | | complex | Activated | 4.175 | 8.88E-15 | 1,DUSP5,EDN1,EGR1,EREG,ETS1,FHL2,FOS,FOSB,FOSL1,GADD45A,GDF15,HBEGF,HLA-E,HMGCR,IGF2,JUN,KLF6,LGALS3,LIF,MAT2A,MCL1,PDLIM1,PHLDA1,PIM1,PPARG,PP | 503 (24) |
| GPER1 | | G-protein coupled receptor | Activated | 4.156 | 3.03E-10 | ATF3,CCN1,CEBPD,DDIT4,DUSP1,DUSP5,EDN1,EGR1,ERRFI1,FOS,FOSB,ITPRIP,JUN,MT1X,MT2A,PPP1R15A,TUFT1,ZFP36 | 593 (24) |
| 2-(4-amino-1-isopropyl-1H-pyrazolo[3,4-d]pyrimidin-3-yl)-1H-indol-5-ol | | chemical reagent | Activated | 4.038 | 5.07E-09 | BC3,BBS4,BCL6,BTG1,CCNF,CD44,CDC25A,CPT1A,CXCL8,DDIT4,EDN1,EGR1,FOSL1,HBEGF,HMGCR,IL1RN,KLF6,LIF,NABP1,PIM1,PML,PPIF,PPP1R15A,PTGS2,SDC4,TCP11L2 | 418 (17) |
| thapsigargin | | chemical toxicant | Activated | 4.028 | 3.12E-16 | CPT1A,CXCL2,CXCL3,CXCL8,DDIT4,DNAJB9,EDN1,EGR1,ERN1,ERRFI1,ETS1,FOS,GADD45A,GADD45B,HMGCR,HMGCS2,HSPA1A/HSPA1B,INSIG1,JUN,KLF4,MCL1,NCOA3,NED | 493 (24) |
| EIF2AK2 | 0.372 | kinase | Activated | 3.995 | 5.14E-11 | ATF3,CASP9,CEBPD,CXCL8,EGR1,EIF2AK2,ERAP2,FAS,FOS,IFI35,IFITM2,IRF1,IRS2,JUN,LGALS3BP,NEDD9,NMI,OAS3,PARP12,PLSCR1,PPP1R15A,PSMB10,ZC3HAV1,ZNF292 | 335 (16) |
| prexasertib | | chemical drug | Activated | 3.962 | 8.33E-14 | ATF3,BCL3,BCL6,BIRC3,CDKN2B,CSRNP1,DUSP1,EGR1,FOS,GADD45B,GDF15,H1-0,IFIH1,IFIT3,IRF7,KLF4,PHLDA1,PMAIP1,PPP1R15A,PTGS2,SGK1,TXNIP,ZFP36 | 398 (7) |
| IFNAR1 | | transmembrane receptor | Activated | 3.921 | 0.000000559 | ATF3,B2M,CGAS,CLIC5,CXCL2,CXCL3,EIF2AK2,HMGCR,IDI1,IFI44,IFIH1,IFIT3,IRF1,IRF7,KLK6,LIF,OAS3,PARP12,PTGS2,SQLE,TAPBPL,TGM2,TNFAIP2,TRIM21 | 318 (16) |
| Ifnar | | group | Activated | 3.919 | 0.0000163 | APOBEC3B,B2M,EIF2AK2,HLA-E,IFI35,IFIH1,IFIT3,IFITM3,IRF1,IRF7,PSMB8,RNF213,TAP2,TAPBP,TRIM21,UNC93B1 | 339 (16) |
| PRL | | cytokine | Activated | 3.912 | 3.59E-12 | L,DVL1,ECM1,EGR1,EIF2AK2,FOS,HERC6,ID2,IFI35,IFI44,IFIH1,IFIT3,IGF2,IRF1,IRF7,JUN,OAS3,PARP10,PARP12,PARP14,PDIA5,PDK4,PIM1,PLSCR1,PSME1,PSME2,RNASE4,S | 385 (21) |
| IRF1 | 0.661 | transcription regulator | Activated | 3.902 | 2E-12 | CFTR,CTSS,CXCL16,CXCL2,CXCL3,CXCL8,DST,EIF2AK2,GATA6,IFI35,IFI44,IFIH1,IFIT3,IFITM3,IRF1,IRF7,MAP4K4,MYB,OAS3,PLAAT3,PLAAT4,PML,PSMB10,PSMB8,PSME1,PSM | 438 (16) |
| 4-hydroxytamoxifen | | chemical drug | Activated | 3.841 | 3.37E-10 | ,CEBPA,CEBPD,CLU,CYP1A1,DDIT4,DUSP1,DUSP5,EDN1,EGR1,ERRFI1,FOS,FOSB,ITPRIP,JUN,KRT13,MT1X,MT2A,NCOA3,NEDD9,PDK4,PMAIP1,PPP1R15A,PTPRH,SGK1,SLC | 505 (22) |
| cigarette smoke | | chemical toxicant | Activated | 3.824 | 0.00000465 | DC25A,CKB,CXCL2,CXCL3,CXCL8,CYP1A1,CYP2B6,CYP3A5,DDIT4,EGR1,ERN1,FOS,FOSL1,JUN,KITLG,KLF4,MAFF,MAP1LC3B,MAP4K4,OAS3,PPARG,PRKCA,PTGS2,PTPRB,R | 497 (21) |
| VEGFA | | growth factor | Activated | 3.803 | 0.000000962 | TP5MC3,BMP4,CAV1,CCN1,CD55,CD68,CEACAM1,CTSS,CXCL2,CXCL8,CXCR4,DUSP5,EDN1,EGR1,ETS1,FOS,FOSB,HBEGF,HK1,MCL1,MMP15,NRP1,PIM1,PLPP3,PNPLA2,PRK | 441 (22) |
| Salmonella enterica serotype abortus equi lipopolysaccharide | | chemical toxicant | Activated | 3.686 | 8.2E-12 | XCL1,CXCL2,CXCL3,DNAJB4,DUSP2,E2F7,EDN1,EGR1,EREG,FOSL1,GADD45A,GADD45B,GBP3,GJB2,GPRC5A,ID2,IL15RA,IRF1,KANK1,MAFF,NEDD4L,NFKBIZ,PHLDA1,PLAC8, | 483 (18) |
| salmonella minnesota R595 lipopolysaccharides | | chemical - endogenous non-mammalian | Activated | 3.671 | 0.00000002 | ACSL1,ADAM9,BCL3,CASP4,CEBPD,CXCL1,CXCL2,CXCL3,CXCL8,EGR1,FOS,FOSB,FOSL1,GDF15,IRF1,ITPR2,JUN,MAFF,MT1X,NFKBIZ,PTGS2,SLFN5,TSC22D1 | 472 (16) |
| PRKCD | | kinase | Activated | 3.598 | 9.4E-12 | CL8,DENND3,ERCC6L,FOS,FOSL1,GDF15,GLI1,GPR87,GPRC5A,HJURP,HRH1,IL1RN,JCAD,JUN,KLF6,KRT20,LIF,LIPG,MAP1LC3B,MCL1,NOX1,NR0B2,PDLIM7,PLSCR1,PPP1R15 | 454 (19) |
| TLR3 | | transmembrane receptor | Activated | 3.534 | 0.000000805 | CXCL3,CXCL8,DUSP1,EDN1,EIF2AK2,FAS,FOS,GADD45B,GBP3,IFI44,IFIH1,IFIT3,IL1R2,IL1RN,IRF1,IRF7,LCN2,LIPG,NFKBIZ,NMI,PHLDA1,PIK3R1,PMAIP1,PROCR,PTGS2,RHBD | 282 (17) |
| MAPK3 | | kinase | Activated | 3.513 | 0.0000622 | BMP4,CDKN2B,CXCL8,CYP24A1,EGR1,FOS,FOSB,FOSL1,JUN,JUND,MCL1,PTGS2,RGS2,SMAD3 | 501 (26) |
| IRF5 | | transcription regulator | Activated | 3.45 | 0.00119 | CXCL2,CXCR4,IFI44,IFIH1,IFIT3,IFITM3,IRF7,PARP12,PLSCR1,PMAIP1,PTGS2,SP110 | 493 (20) |
| TLR4 | | transmembrane receptor | Activated | 3.431 | 0.0000495 | S,CEBPD,CXCL2,CXCL3,CXCL8,CYP3A5,DNMT3A,DUSP2,EDN1,EFNA1,EREG,FAS,FBXO32,HBEGF,IFIT3,IFITM3,IL15RA,IRF1,IRF7,LCN2,NFKBIZ,NMI,PML,PPARG,PTGS2,REL,S | 441 (15) |
| carbamazepine | | chemical drug | Activated | 3.407 | 0.00000918 | ATF3,CYP2B6,CYP3A5,DNAJB1,EIF4A2,GDF15,JUND,KLF9,MAP1LC3B,MCL1,PSPH,TNFRSF12A | 173 (7) |
| IL6 | | cytokine | Activated | 3.381 | 1.1E-13 | ,DUSP16,DVL1,EGR1,EREG,ERN1,ERRFI1,FAS,FOS,FOSL1,FRRS1,GADD45A,GADD45B,GBP3,GLI1,GSTA4,HPGD,ID2,IFITM3,IGF2,IGFBP6,IL1RN,IRF1,JUN,JUND,LCN2,LGR4,LI | 511 (21) |
| carrageenan | | chemical drug | Activated | 3.378 | 0.00000605 | BCL10,CXCL2,CXCL3,CXCL8,EGR1,F2RL1,FOS,FOSB,FOSL1,JUN,JUND,PTGS2 | 346 (17) |
| CD40LG | | cytokine | Activated | 3.377 | 4.08E-13 | DUSP1,DUSP2,DUSP5,EGR1,EIF2AK2,FAS,FOS,FOSB,GADD45A,HSPA1A/HSPA1B,ID2,IFI44,IFIT3,IRF1,JUN,JUND,KDM6B,MCL1,MT1E,MT2A,NCOA3,PIM1,PMAIP1,PML,PSMB10 | 478 (19) |
| carbonyl cyanide m-chlorophenyl hydrazone | | chemical toxicant | Activated | 3.357 | 3.86E-08 | AADAC,CLDN1,CXCL8,DDIT4,DUSP5,ERRFI1,GDF15,LCN2,MAP1LC3B,NFKBIZ,PHLDA1,PTGS2,RND3,SPINK1,ZFP36L1 | 439 (19) |
| C11orf95-RELA | | fusion gene/product | Activated | 3.357 | 3.53E-09 | BCL3,BIRC3,CASP4,CD44,CXCL2,CXCL3,DUSP2,IL15RA,IL1RN,IRF1,IRF7,PTGS2,SDC4,TAPBP,TGM2 | |
| C5 | | cytokine | Activated | 3.352 | 0.000407 | ATF3,BCL6,CD55,CXCL1,CXCL2,CXCL3,CXCL8,EGR1,EIF2AK2,ERN1,GDF15,IFNGR2,PLK3,PPP1R15A,TNFAIP2,ZFP36 | 456 (16) |
| F7 | | peptidase | Activated | 3.334 | 3.93E-10 | CASP7,CCN1,CLDN1,CXCL2,CXCL8,EGR1,FOS,FOSL1,GADD45A,HBEGF,LIF,MMP7,PTGS2,RND3,ZFP36 | 437 (18) |
| trovafloxacin | | chemical drug | Activated | 3.302 | 0.000438 | B2M,BIRC3,CEBPA,CXCL2,CXCL3,ERRFI1,FOS,IRF1,NEDD4L,PHLDA1,TGM2 | 459 (13) |
| TLR7 | | transmembrane receptor | Activated | 3.23 | 0.0111 | ATF3,CD44,CREB5,CXCL1,CXCL2,CXCL3,CXCL8,DKK1,FGF19,FOSL1,HIVEP2,IFI35,IFI44,IFIT3,IRF1,IRF7,OAS3,TRIM38,ZC3H12A | |
| SAMSN1 | | other | Activated | 3.207 | 0.000133 | CXCL2,EDN1,HBEGF,IFIT3,IL15RA,IRF1,IRF7,NFKBIZ,NMI,PML,PTGS2,TRIM21,TSC22D1,ZC3H12A | 214 (7) |
| tributyrin | | chemical drug | Activated | 3.201 | 1.67E-15 | AKAP12,ATF3,ATP2A3,AZGP1,BIRC3,C1orf116,CCN1,CD55,CLDN1,CXCL3,CXCL8,CXCR4,DAPP1,DUSP1,EDN1,ETS1,HBEGF,ID2,PTGS2,SAMD4A,SLC30A1,TUFT1,UGCG,USP53 | 321 (7) |
| PPRC1 | | transcription regulator | Activated | 3.201 | 4.29E-08 | AADAC,CLDN1,CXCL8,DDIT4,DUSP5,ERRFI1,GDF15,LCN2,NFKBIZ,PHLDA1,PTGS2,RND3,SPINK1,ZFP36L1 | |
| 25-hydroxycholesterol | | chemical reagent | Activated | 3.185 | 0.000000424 | ACSL1,ATF3,CHAC1,CXCL8,FOS,FOSB,FOSL1,HMGCR,INSIG1,JUN,JUND,MAFF,PTGS2,SCARB1,SULT2B1 | 432 (21) |
| TLR9 | | transmembrane receptor | Activated | 3.164 | 0.000445 | ARRDC4,ATF3,CCNA2,CXCL2,CXCL3,CXCL8,DUSP1,EDN1,EGR1,FAS,GADD45B,IFI35,IFIT3,IRF1,IRF7,LCN2,LIPG,MCL1,NFKBIZ,NR0B2,OAS3,PHLDA1,PROCR,PTGS2,UGCG | 435 (18) |
| IRF3 | | transcription regulator | Activated | 3.145 | 0.00000967 | C2,ADAM9,APOBEC3B,B2M,BIRC3,CD58,CXCL1,CXCL8,EIF2AK2,FAS,IFI44,IFIH1,IFIT3,IFITM3,IRF1,IRF7,LCN2,OAS3,PARP12,PARP14,PLAC8,PMAIP1,PML,RASA1,SORL1,ZC3H | 247 (14) |
| ELAVL1 | | other | Activated | 3.145 | 0.000028 | ANXA11,ATF3,ATP1B3,B2M,CASP9,CLDN1,CTSS,DNMT3A,DUSP1,FOS,IFI44,IFIH1,IFITM3,IRF1,LGALS3BP,MAT2A,PTGS2,REN,SAT1,SLC7A7,TIA1,TJP1,TSC22D3,WARS1,ZFP36 | 449 (16) |
| APP | | other | Activated | 3.128 | 8.69E-10 | HERC6,HIPK2,HK1,HLA-E,HMGCR,HSPA1A/HSPA1B,HSPB1,HSPG2,IFI35,IFIH1,IGF2,IGFBP2,IRF1,IRF7,IRS2,JUN,JUND,KCNE3,LDHA,LGALS3,LIF,MAFF,MDK,NCOA3,NR1D1,NRI | 504 (20) |
| Tlr | | group | Activated | 3.12 | 0.00484 | BCL6,CASP4,CXCL3,CXCL8,DUSP1,HBEGF,ID2,IRF1,IRF7,LCN2,PTGS2,SEMA3A,ZC3H12A | 387 (16) |
| PML | 0.486 | transcription regulator | Activated | 3.12 | 4.51E-10 | CXCL1,CXCR4,DNAJA4,DNAJB4,DNAJB9,ELOVL6,FAS,GADD45A,HERC6,HMGCR,HSPB1,ID2,IDI1,IFI35,IFI44,IFIH1,IFIT3,IRF7,MAFF,ME1,OAS3,PDK4,PLSCR1,PML,PRKCA,PSM | 468 (20) |
| methylprednisolone | | chemical drug | Activated | 3.117 | 4.21E-11 | GPD1,HSPA1A/HSPA1B,HSPH1,IL1RN,INSIG1,IRF1,IRF7,JUN,KLF9,LGR4,MAOB,MAP1LC3B,ME1,MGAT1,MYO1B,NR0B2,PEX11A,PIK3R1,PLAAT3,PPL,PSME2,REL,RGS2,RND3,S | 473 (22) |
| MAPK8 | | kinase | Activated | 3.104 | 0.000000272 | BIRC3,BMP4,CASP9,CCNA2,CXCL16,CXCL8,CYP1A1,DUSP1,EGR1,ETS1,FOS,FOSL1,GADD45A,HMGCR,JUN,JUND,KLF6,LIF,MAOB,MMP7,NRP1,PTGS2,SESN2,TSC22D3,VDR,Z | 434 (18) |
| resiquimod | | chemical drug | Activated | 3.098 | 2.61E-08 | CXCL1,CXCL2,CXCL3,CXCL8,DNAJB9,DNAJC3,EIF2AK2,ELF3,F2R,FAS,GDF15,HIVEP2,HSPB1,IFIT3,IFITM3,IRF7,JUN,LIF,MCL1,NFKBIZ,OAS3,PLSCR1,PMAIP1,PML,PTGS2,REL, | 453 (18) |
| STING1 | | other | Activated | 3.077 | 0.0000476 | CALB2,CD68,CGAS,CXCL2,CXCL8,IFI44,IFIT3,IFITM3,IRF7,LCN2,LIF,PPP1R15A | 370 (19) |
| CpG ODN 2006 | | chemical reagent | Activated | 3.072 | 0.0232 | BIRC3,CXCL8,IFI35,IFI44,IFIH1,IFIT3,IL15RA,IRF7,NFKBIZ,SP100 | |
| gentamicin | | chemical drug | Activated | 3.061 | 1.15E-09 | EP1,DPP4,EGR1,FOS,GADD45A,GDF15,GJB2,GUK1,HIVEP2,HNF4A,HSPA1A/HSPA1B,HSPB1,IRF1,IRF7,JUN,KLF6,LCN2,LGALS3,MYO1B,NPNT,NR1D1,NR1D2,NRIP1,NUDC,PPI | 484 (19) |
| ATF4 | | transcription regulator | Activated | 3.055 | 5.22E-12 | CXCL1,CXCL2,CXCL8,DDIT4,FYN,GADD45A,GDF15,HNF4A,IGFBP2,JUN,KLF4,KLF9,LGALS3,MAP1LC3B,MCL1,MT2A,PCK2,PMAIP1,PPARG,PPP1R15A,PSPH,PTGS2,SIGMAR1,S | 380 (14) |
| MKNK1 | | kinase | Activated | 3.051 | 0.00143 | BAG3,CCN1,ETS1,FLNB,FOS,FOSB,GNAI1,GPC4,HSPA4L,LCN2,PLXNB2,SERPINH1,TXNIP | |
| leukotriene D4 | | chemical - endogenous mammalian | Activated | 3.05 | 4.86E-09 | ARID5B,ATF3,CCN1,CXCL2,CXCL8,DUSP1,EGR1,HBEGF,KLF4,MCL1,NEDD9,PDK4,PTGS2,RCAN1,RGS2,ZFP36 | 501 (23) |
| FOXO3 | | transcription regulator | Activated | 3.05 | 8.2E-14 | T1A,CXCL8,CXCR4,DDIT4,DEPP1,EGR1,FBXO32,FGFR2,FLNB,FOS,FOSB,FYN,GADD45A,GADD45B,GATA2,HMGCR,HMGCS2,IFIH1,IRS2,LCN2,MAP1LC3B,MCAM,MXD1,MXD4,N | 570 (28) |
| TICAM1 | | other | Activated | 2.992 | 0.000000545 | ACSL1,APOBEC3B,BCL3,CASP4,CEBPD,CXCL2,CXCL3,CXCL8,DUSP1,EGR1,FAS,GDF15,IFIT3,IL15RA,IRF1,IRF7,ITPR2,MAFF,NFKBIZ,PTGS2,RHOB,TSC22D1,ZC3HAV1 | 462 (22) |
| camptothecin | | chemical drug | Activated | 2.991 | 2.59E-16 | GRK5,GSE1,HBEGF,HJURP,HLA-B,HLA-C,HLA-E,IL15RA,INSIG1,IQGAP2,IRF1,JUN,LGALS3,LOC102724788/PRODH,MAFF,MAP4K4,MBOAT7,MCL1,MYB,NRP1,PDLIM7,PHF21A,PI | 439 (14) |
| mycophenolic acid | | chemical drug | Activated | 2.991 | 2.08E-14 | P12,ATF3,AZGP1,BIRC3,C1orf116,CCN1,CD55,CDK6,CLDN1,CLU,CXCL3,CXCL8,CXCR4,DAPP1,DUSP1,EDN1,ETS1,HBEGF,ID2,PTGS2,RUNX2,SAMD4A,SLC30A1,TUFT1,UGCG,U | 454 (22) |
| Cdk | | group | Activated | 2.99 | 0.00000253 | ATP5MC1,CXCL2,CXCL8,DDIT4,DUSP1,GADD45A,NEDD9,PHLDA1,RGS2,TXNIP | 387 (12) |
| FOXL2 | | transcription regulator | Activated | 2.982 | 0.000031 | ATF3,CXCL2,CXCL3,FAS,FOS,LIF,MAFF,MYB,NR5A2,PPP1R15A,PTGS2,RGS2,SERPINE2,SOX4 | 226 (9) |
| IL27 | | cytokine | Activated | 2.958 | 0.0000308 | APOL1,B2M,CEBPD,CXCL8,DEPP1,EGR1,EIF2AK2,FOS,HLA-B,HLA-C,HLA-E,HPGD,ID2,IL15RA,IL1RN,IRF1,MCL1,PIM1,PLAAT4,PTGS2 | 516 (19) |
| CD24 | | other | Activated | 2.935 | 3.86E-15 | I,CFTR,CHAC1,CXCL8,DEPDC1B,DUSP5,ECM1,EPHB6,FOS,FRYL,GDF15,JMJD1C,JUN,LAT2,LPP,MCAM,MTUS1,NCOA3,NRIP1,PPP1R15A,RARRES1,RASA1,SLC30A1,SLC4A4,T | 410 (18) |
| PTGS2 | 0.741 | enzyme | Activated | 2.935 | 0.0000159 | ANGPT1,ANXA1,AXIN2,BBC3,BIRC3,CD44,CD68,CLU,CXCL3,CXCL8,CXCR4,DUSP1,EGR1,FOS,HBEGF,MCL1,MMP7,PPARG,PTGS2,PYGB,RUNX2,SEMA7A,TGM2,TNFRSF12A | 474 (22) |
| Jnk | | group | Activated | 2.91 | 1.92E-09 | CXCL3,CXCL8,CXCR4,DKK1,DUSP1,EDN1,EGR1,ETS1,F2RL1,FAS,FBXO32,FOS,FOSB,GADD45A,GDF15,HNF4A,JUN,JUND,KLF6,MCL1,PLSCR1,PTGS2,RARG,SESN2,STRA6,TC | 455 (20) |
| cholesterol | | chemical - endogenous mammalian | Activated | 2.892 | 4.9E-09 | V1,CAV2,CD68,CPT1A,CXCL2,CXCL3,CXCL8,CYP3A5,DDC,GLI2,GNAI1,HBEGF,HES6,HMGCR,IDI1,IL1RN,INSIG1,KITLG,LGALS3,MAP1LC3B,ME1,MSMO1,NR0B2,PPARG,PTGS2,S | 438 (18) |
| oblimersen | | biologic drug | Activated | 2.887 | 0.0000112 | ATF3,CEBPD,CXCL8,DUSP1,GADD45A,GADD45B,IFI35,IFI44,IFIT3,IRF7,MAFF,PRKCA | |
| dalfampridine | | chemical drug | Activated | 2.887 | 0.0000286 | ATF3,CCN1,DUSP1,EGR1,FOS,FOSB,GADD45B,JUN,KLF4,MAFF,PTGS2,TIPARP | 400 (19) |
| bicuculline | | chemical - endogenous non-mammalian | Activated | 2.884 | 0.000000453 | ATF3,CCN1,DNMT3A,DUSP1,EGR1,FOS,FOSB,GADD45B,JUN,KLF4,MAFF,PTGS2,TGFB2,TIPARP | 472 (22) |
| 3-deazaneplanocin | | chemical drug | Activated | 2.884 | 0.000056 | FAS,FBXO32,KLF6,KRT7,LYSMD4,MAP3K14,PJA1,PPP1R15A,PRKCA,SOX12,TSC22D3,USP53 | |
| JUN | 0.48 | transcription regulator | Activated | 2.852 | 6.19E-13 | XCL8,DKK1,DMTF1,DUSP1,DUSP5,E2F7,EDN1,EREG,FAS,FOS,FOSL1,GADD45A,HK1,HLA-B,HPGD,ID2,IGFBP2,IGFBP6,IRS2,JUN,LGALS3,LIF,MAOB,MAT2A,MMP7,MT2A,MYB,N | 595 (24) |
| IFN Beta | | group | Activated | 2.838 | 0.0000176 | ATF3,CASP4,CEBPD,CGAS,CXCL1,CXCL8,DPP4,DUSP1,EIF2AK2,EREG,HLA-B,IFI35,IFI44,IFIH1,IFIT3,IL15RA,IL1RN,INSIG1,IRF1,IRF7,OAS3,PRKCA,TAPBP | 480 (18) |
| aldesleukin | | biologic drug | Activated | 2.828 | 0.00213 | B2M,HLA-B,HLA-C,IFI35,IFNGR2,IRF1,ITM2B,TAPBP | |
| MYD88 | | other | Activated | 2.826 | 0.000000097 | ASP4,CCN1,CD44,CEBPD,CXCL1,CXCL2,CXCL3,CXCL8,DNMT3A,DUSP1,EGR1,FANCF,FBXO32,FOS,GDF15,IL1RN,IRF1,IRF7,ITPR2,JUN,MAFF,MMP7,MUC2,NFKBIZ,NOX1,PTGS | 454 (17) |
| palmitic acid | | chemical - endogenous mammalian | Activated | 2.823 | 4.27E-09 | CD68,CGAS,CXCL2,CXCL3,CXCL8,DGAT2,ELOVL6,ERN1,F2RL1,FBXO32,FOS,GDF15,HNF4A,IL1RN,IRS2,KLF4,LCN2,MAP1LC3B,MCL1,NOX1,NR0B2,PDK4,PFKM,PNPLA2,PPAR | 459 (22) |
| kainic acid | | chemical toxicant | Activated | 2.822 | 0.00000101 | ATF3,CALB2,CSRNP1,CTSE,DUSP1,EGR1,ERRFI1,FGFR2,FOS,FOSB,FOSL1,GADD45B,HBEGF,JUN,JUND,KLK6,LCN2,MMP7,PPP1R15A,PTGS2,RGS2,SLC16A1,TIPARP,TJP1 | 356 (23) |
| TLR7/8 | | group | Activated | 2.813 | 0.000731 | EDN1,EREG,NFKBIZ,PMAIP1,PPP1R15A,PTGS2,REPS2,VDR | 458 (12) |
| IL3 | | cytokine | Activated | 2.791 | 0.0000822 | B,BIRC3,CASP9,CPT1A,CXCL8,CXCR4,EGR1,EIF2AK2,F2R,FAS,FOS,GADD45A,GADD45B,GATA2,HBEGF,HK1,HNF4A,JUN,KLF9,LATS2,LCN2,LIF,MCL1,MID1,NEDD9,NRP1,PIM1 | 467 (21) |
| hydrogen peroxide | | chemical - endogenous mammalian | Activated | 2.783 | 6.25E-15 | 16,DUSP2,EDN1,EGR1,ENG,EPHA2,ETS1,F2RL1,FAS,FBXO32,FOS,FOSL1,GADD45A,GADD45B,GDF15,GLI1,HBEGF,HIPK2,HMGCR,HMGCS2,INSIG1,JUN,KITLG,KLF4,LCN2,LGI | 490 (18) |
| methyl methanesulfonate | | chemical toxicant | Activated | 2.78 | 0.000000889 | ATF3,BTG1,EGR1,GADD45A,GADD45B,JUN,KLF4,LIF,MCL1,NFYA,NMI,PLXNB2 | 454 (16) |
| bromodeoxyuridine | | chemical drug | Activated | 2.771 | 0.00404 | CXCL8,IFI44,IFIH1,IFIT3,IRF1,IRF7,PML,SP110 | 290 (10) |
| stallimycin | | biologic drug | Activated | 2.771 | 0.00161 | CXCL8,IFI44,IFIH1,IFIT3,IRF1,IRF7,PML,SP110 | 290 (10) |
| calcimycin | | chemical reagent | Activated | 2.768 | 0.0000501 | AMD1,ATF3,CXCL8,DDIT4,EGR1,FOS,FOSB,FOSL1,JUN,LCN2,MUC2,PPARG,PTGS2,RCAN1,REN,S100A2,SDF2L1 | 449 (18) |
| peptidoglycan | | chemical - endogenous non-mammalian | Activated | 2.762 | 0.000101 | CXCL1,CXCL16,CXCL2,CXCL3,CXCL8,CXCR4,DUSP1,EGR1,FAS,FOS,IL1RN,IRF1,MCL1,PTGS2,VDR,VEZF1 | 398 (16) |
| anisomycin | | chemical - endogenous non-mammalian | Activated | 2.755 | 0.00218 | ATF3,CXCL8,CYP1A1,DUSP1,EGR1,FOS,IRF1,JUN,MCL1 | 415 (19) |
| E. coli lipopolysaccharide | | chemical - endogenous non-mammalian | Activated | 2.755 | 0.0281 | CCN1,CEBPA,CEBPD,CXCL1,CXCL2,CXCL3,CXCL8,PTGS2 | |
| TRAF3IP2 | | enzyme | Activated | 2.735 | 0.0026 | CEBPD,CXCL16,CXCL2,CXCL3,LCN2,NFKBIZ,STEAP4,TNFRSF12A,TRAF4,ZC3H12A | 445 (18) |
| formaldehyde | | chemical - endogenous mammalian | Activated | 2.731 | 0.0000351 | CYP1A1,EGR1,FOS,FOSB,FOSL1,JUN,JUND,PTGS2,SGK1 | 435 (23) |
| DOCK8 | | other | Activated | 2.714 | 0.000499 | EDN1,IFIT3,IL15RA,IRF1,IRF7,NFKBIZ,NMI,PTGS2,REL,TRIM21,TSC22D1 | |

Supplementary Table 4. Upstream regulator analysis based gene set enrichment analysis (GSEA) of differentially expressed genes (DEGs) after ATRA+PDT compared to PDT

| Upstream Regulator | Expr Log Ratio | Molecule Type | Predicted Activation State | Activation z-score | p-value of overlap | Target Molecules in Dataset | Mechanistic Network |
|---|---|---|---|---|---|---|---|
| SASH1 | | other | Activated | 2.714 | 0.00064 | EDN1,IFIT3,IL15RA,IRF1,IRF7,NFKBIZ,NMI,PML,PTGS2,TRIM21,TSC22D1 | 311 (7) |
| RARG | 0.61 | ligand-dependent nuclear receptor | Activated | 2.708 | 0.000431 | CRABP2,DHRS3,DUSP1,EGR1,FOS,HOXA3,HOXA5,IGFBP6,JUN,KLF4,KRT20,SLC17A9,STRA6 | 499 (25) |
| ID2 | -1.001 | transcription regulator | Activated | 2.704 | 0.000679 | AXIN2,BCL3,BCL6,BIRC3,CCNG2,CD44,CXCR4,DUSP1,FAS,GADD45B,IFNGR2,JDP2,KLF6,LRIG1,MAP3K14,MYB,PIK3R1,RAP1GAP,REL,SOX4,SPDEF,TRAF4,WNT11 | 236 (9) |
| cocaine | | chemical drug | Activated | 2.701 | 0.000574 | CPT1A,CXCL2,CXCL8,DNMT3A,DUSP1,DUSP5,EGR1,FHL2,FOS,FOSB,FOSL1,GADD45A,GADD45B,IRS2,JUN,LAMC2,MCL1,PLCXD1,PPP1R1B,SIGMAR1,TIPARP | 486 (22) |
| CG | | complex | Activated | 2.697 | 4.17E-18 | FAM3D,FAS,FOS,FYN,GASK1B,GATA6,GJB2,HMGA2,HMGCR,HPGD,IFIT3,IGFBP2,IL1R2,IRS2,JUN,KLF4,LGALS3BP,LGR4,LIF,MCAM,MCL1,NR0B2,NR5A2,NRIP1,NRP1,NUCB2,P | 594 (23) |
| bortezomib | | chemical drug | Activated | 2.685 | 0.000000228 | CAV1,CEBPD,CXCL2,CXCL8,CXCR4,DKK1,DNAJA1,DNAJB1,DNAJB4,ERN1,FANCF,FAS,GADD45A,GADD45B,GDF15,GLI1,HSPA1A/HSPA1B,KLF9,LIF,MCL1,PKIB,PMAIP1,PPP1 | 515 (20) |
| GnRH analog | | biologic drug | Activated | 2.683 | 0.0964 | ABCC1,ADAM8,ANXA11,ATF3,BCL3,CIB1,DUSP1,DVL1,FAS,FOS,GAB1,JUN,MARF1,MIA3,NCOA3,PKNOX1,PLAAT3,SKAP2,SRI,SSR1,TNK2,TXK | |
| PRKCA | 0.35 | kinase | Activated | 2.66 | 0.00000287 | ARHGDIB,CAV1,CD55,CXCL2,CXCL3,EGR1,ETS1,FOS,GDF15,HSPA1A/HSPA1B,ID2,IFIT3,INSIG1,JUN,PLD1,PML,PTGS2,RUNX2,SP100,TGFB2 | 430 (18) |
| TP53 | | transcription regulator | Activated | 2.659 | 2.32E-30 | HA2,F2R,F5,FAS,FHL2,FIGNL1,FOS,FOSL1,FSTL3,FXYD3,FYN,GADD45A,GADD45B,GATA6,GDF15,GLI2,GNA14,GNAI1,GPD1,GPR160,GPR87,GSN,HBEGF,HDAC9,HJURP,HLA-B, | 523 (20) |
| CHUK | | kinase | Activated | 2.653 | 0.000000057 | BPD,CHST4,CLDN1,CLU,COL18A1,CXCL2,CXCL3,CXCL8,EREG,FAS,FOS,GADD45B,GBP3,HES6,IFI35,IGFBP6,IL1RN,ITM2B,LCN2,MAT2A,MXD1,OVOL1,PLSCR1,PTGS2,RCAN1, | 435 (14) |
| collagenase | | group | Activated | 2.646 | 0.000763 | CEBPD,CXCL2,CXCL3,EGR1,FOSL1,IRF1,JUN | 441 (13) |
| GNRH | | group | Activated | 2.609 | 0.000514 | ATF3,DUSP1,EGR1,FOS,JUN,PTGS2,SEMA3A,SGK1 | 461 (21) |
| diclofenac | | chemical drug | Activated | 2.608 | 1.04E-08 | ATF3,DNAJB1,EIF4A2,FOS,GDF15,HPGD,JUN,JUND,KLF4,KLF9,LDHA,LGALS3,MAP1LC3B,MCL1,PPP1R15A,PSPH,PTGS2,SLC16A1,TNFRSF12A,WARS1 | 483 (21) |
| CREBBP | | transcription regulator | Activated | 2.601 | 5.15E-08 | R4,CYP24A1,DDC,DHRS3,DNAJB4,DUSP1,EGR1,ESAM,FOS,FOSB,FRMD4B,GLCCI1,GLI1,GRIN2D,HLA-B,HMGCS2,IRF1,JUN,JUND,LGALS3BP,NPAS2,OAS3,PPFIBP2,PTGS2,RE | 446 (23) |
| PDGFB | | growth factor | Activated | 2.599 | 0.0517 | BBC3,CXCL8,EGR1,FOS,IL1RN,PHLDA1,PTGS2 | |
| Mapk | | group | Activated | 2.585 | 0.0169 | CXCL1,CXCL8,EGR1,FOS,HPGD,JUN,MMP7,NOX1,PTGS2,RUNX2,SNCG | |
| quinolinic acid | | chemical - endogenous mammalian | Activated | 2.578 | 0.00846 | EGR1,FOS,FOSB,HSPA1A/HSPA1B,JUN,JUND,PTGS2 | 422 (21) |
| 9,10-dimethyl-1,2-benzanthracene | | chemical toxicant | Activated | 2.573 | 0.000748 | CXCL2,CXCL3,CYP1A1,EPHA2,GADD45A,HBEGF,JUN,PTGS2,REL | 455 (16) |
| MAP3K1 | | kinase | Activated | 2.572 | 0.0000512 | ATF3,BIRC3,CXCL8,DUSP1,EGR1,FAS,FOS,HSPB1,JUN,MAOB,PTGS2 | 400 (16) |
| uric acid | | chemical - endogenous mammalian | Activated | 2.57 | 0.16 | ANXA1,CXCL2,CXCL3,CXCL8,EIF2AK2,IFIT3,IRF7,PTGS2 | |
| F2RL1 | 0.405 | G-protein coupled receptor | Activated | 2.569 | 0.000000342 | CASP7,CAV1,CCN1,CXCL1,CXCL2,CXCL8,F2RL1,FAS,FOS,HBEGF,KITLG,LIF,PTGS2,RARG,TXNIP | 463 (20) |
| PRKCE | | kinase | Activated | 2.564 | 0.00143 | BIRC3,CAV1,CCN1,CD55,CXCL8,EGR1,FOS,JUN,PIM1,PTGS2,RGS2 | 454 (24) |
| cytokine | | group | Activated | 2.56 | 0.00000635 | BCL3,CCN1,CLU,CXCL1,CXCL3,CXCL8,DUSP1,EDN1,EFNA1,EGR1,FAS,FOS,IGFBP2,IRF1,JUN,LIF,LIPG,PLAAT4,PTGS2,S100A4,SGK1,TGFB2 | 376 (14) |
| MEF2C | | transcription regulator | Activated | 2.555 | 0.00612 | ATF3,CKB,CXCL2,CXCR4,FOS,FOSB,HDAC9,JUN,JUND,PTGS2,PTPRB,RUNX2,S100A4,ZFP36 | 382 (12) |
| RAF1 | | kinase | Activated | 2.537 | 9.14E-16 | P1,CDKN2B,CLDN1,CLK1,CXCL3,DHRS3,DUSP2,DUSP5,EGR1,EMP1,FAM13A,FAS,FOS,HBEGF,HMGA2,HSPB1,JUN,LDHA,LIF,MAOB,MXD1,PCSK6,PHLDA1,PLD1,PPP1R10,PTG | 459 (19) |
| TGFA | | growth factor | Activated | 2.527 | 0.000168 | BIRC3,CXCL2,CXCL8,EREG,ERRFI1,FOS,GJB2,LAMC2,LCN2,MUC2,PTGS2,S100A10,SERPINA1 | 521 (27) |
| 5-O-mycolyl-beta-araf-(1->2)-5-O-mycolyl-alpha-araf-(1->1')-glycerol | | chemical - endogenous non-mammalian | Activated | 2.524 | 2.19E-08 | ACSL1,ATF3,BCL3,CD44,CXCL1,CXCL2,CXCL3,CXCL8,DUSP1,HBEGF,IFIT3,NFKBIZ,NT5E,PCSK5,PTGS2,RCAN1,RND3,SDC4,SERPINE2 | 437 (14) |
| sulindac sulfide | | chemical drug | Activated | 2.521 | 0.00000746 | ATF3,BCAM,CYP1A1,EGR1,GADD45A,GDF15,INSIG1,LDHA,MMP7,MSMO1,PIK3R1,PTGS2,RAP1GAP,TUBA4A | 467 (20) |
| IL1 | | group | Activated | 2.519 | 0.000012 | ,CEBPD,CPT1A,CXCL1,CXCL2,CXCL3,CXCL8,CYP1A1,CYP2B6,DDC,DUSP1,EDN1,EGR1,ELF3,FAS,FOS,GDF15,HMGCR,IL1R2,IL1RN,IRF1,JUN,KITLG,KLF6,LIF,MMP7,NOX1,PLD | 493 (19) |
| MAP2K1 | | kinase | Activated | 2.511 | 2.31E-13 | CD44,CDKN2B,CEBPA,CLDN2,CXCL3,CXCL8,DKK1,DPP4,DUSP1,DUSP5,F2R,FOS,FOSL1,GLI2,JUN,JUND,LGALS3,MAOB,NUDT19,PIM1,PTGS2,RAP1GAP,RAP2B,RUNX2,SCARE | 503 (21) |
| Am 580 | | chemical reagent | Activated | 2.503 | 0.000256 | CPT1A,DUSP1,HMGCR,IRF1,LSS,PPARG,RARG,SQLE,STRA6,TGM2 | 481 (23) |
| Ifn | | group | Activated | 2.502 | 0.00000756 | B2M,BCL6,CD44,CD58,EDN1,EIF2AK2,FAS,FOS,IFIH1,IFITM3,IL15RA,IRF1,IRF7,LCN2,OAS3,PLAAT4,PML,SP100,TRIM21 | 308 (15) |
| IL1A | | cytokine | Activated | 2.502 | 5.65E-13 | XCL1,CXCL2,CXCL3,CXCL8,CYP1A1,CYP2B6,CYP3A5,DPP4,F2RL1,FAS,FOS,FOSB,FOSL1,HSPG2,IFNGR2,IL1R2,IL1RN,IRF1,JUN,KITLG,LCN2,LDHA,LGALS9,LIF,MCAM,MT2A,N | 445 (15) |
| IKBKG | | kinase | Activated | 2.502 | 0.0000405 | BBC3,BIRC3,CEBPD,CLU,CXCL2,CXCL3,CXCL8,EREG,FOS,GBP3,IGFBP6,IL1RN,IRF7,LCN2,PPARG,RCAN1,SERPINE2,SGK1,TNFAIP2 | 449 (16) |
| ELK1 | | transcription regulator | Activated | 2.501 | 0.000000424 | EGR1,FOS,FOSB,FOSL1,FUT4,ITGB6,JUN,MCL1,PRKCA,PSMB10,PSMB8,PTGS2,RUNX2,TIPARP,ZFP36 | 519 (23) |
| MIF | | cytokine | Activated | 2.5 | 0.00152 | CD44,CXCL16,CXCL2,CXCL3,CXCL8,CXCR4,DGAT2,DUSP1,F2R,F2RL1,FOS,IL15RA,JUN,MAP1LC3B,PTGS2 | 472 (21) |
| IL33 | | cytokine | Activated | 2.48 | 0.00956 | CAT1,ALDH1A3,BCL3,BIRC3,CXCL16,CXCL2,CXCL3,CXCL8,DGAT2,DUSP2,FAS,GADD45B,IL1RN,LAMA5,MDK,NABP1,NFKBIZ,NT5E,REL,SCARB1,TNFAIP2,ZC3H12A,ZFHX3,ZH | 377 (13) |
| GnRH-A | | chemical reagent | Activated | 2.479 | 5.06E-08 | AKR1B10,ATF3,CD68,EGR1,FER,FOS,FOSB,FOSL1,GADD45B,GCNT2,JUN,KLF4,KLF6,NFKBIZ,PPP1R15A,RGS2,ZFP36 | 409 (20) |
| TNFSF11 | | cytokine | Activated | 2.475 | 1.58E-08 | BRF,CXCL3,CXCL8,CYP1A1,DUSP1,DUSP16,ETS1,FAS,FHL2,FOS,FOSL1,FRRS1,GADD45B,HIPK2,IL15RA,IL1RN,JDP2,JUN,KDM8,NFKBIZ,PAG1,PIM1,PLD1,PTGS2,RCAN1,RHO | 464 (20) |
| Ap1 | | complex | Activated | 2.469 | 0.000382 | ATF3,CCNA2,CD44,CLU,CXCL8,EDN1,ETS1,F2RL1,FAS,FOS,FOSL1,HPGD,JUN,KLF9,MMP7,MT2A,PTGS2,S100A4,SNCG,VDR | 443 (18) |
| dopamine | | chemical - endogenous mammalian | Activated | 2.469 | 0.0931 | ATF3,CSRNP1,CXCL8,DUSP2,EGR1,FOS,FOSB,GATA2,JUN,KLF4,PPP1R1B | |
| GH1 | | growth factor | Activated | 2.468 | 0.0000559 | ABCC1,ACVR1,ATF3,CEBPA,CEBPD,CLU,CXCR4,EGR1,FOS,GADD45A,GATA2,ID2,JUN,MYB,UQCRHL,VEZF1,ZFP36 | 553 (26) |
| MAP4K4 | 0.776 | kinase | Activated | 2.468 | 0.000656 | CBLB,CEBPA,CHPT1,DHODH,IMPDH1,PEX11A,PFKM,PGM1,PPARG,PRKAR2A,PXMP2,RPS6KB2,ST3GAL2,ST3GAL4,UQCRFS1 | 96 (3) |
| tunicamycin | | chemical - endogenous non-mammalian | Activated | 2.456 | 4.03E-12 | ,DNAJB9,DNAJC3,EGR1,ERN1,ETS1,GADD45A,GADD45B,KLF4,MCL1,MUC2,NCOA3,NEDD4L,PCK2,PHLDA1,PMAIP1,PPARG,PPP1R15A,PTGS2,S100A4,SDF2L1,SEC24D,SEC61 | 390 (16) |
| Ifn gamma | | complex | Activated | 2.453 | 0.000265 | CXCL16,CXCL8,EDN1,EIF2AK2,FAS,LGALS9,LIF,MAP4K4,PMAIP1,PML,PPARG,PSME1,PSME2,TGFB2 | 491 (19) |
| Collagen type II | | complex | Activated | 2.449 | 0.0196 | BMP4,CXCL1,CXCL2,CXCL3,IL1RN,MMP7 | |
| OSCAR | | other | Activated | 2.449 | 0.00565 | CXCL1,CXCL2,CXCL3,CXCL8,IL1RN,MMP7 | 356 (16) |
| MARK2 | | kinase | Activated | 2.449 | 0.00485 | CD44,CXCL2,CXCL3,IRF1,NFKBIZ,PTGS2 | 303 (9) |
| TRADD | | other | Activated | 2.449 | 0.00203 | CXCL1,CXCL2,CXCL3,CXCL8,IRF1,TNFAIP2 | 362 (16) |
| GNA13 | | enzyme | Activated | 2.44 | 0.0000423 | CXCL1,CXCL2,CXCL3,CXCL8,EGR1,FOS,PTGS2 | 432 (19) |
| MASTL | | kinase | Activated | 2.433 | 0.024 | ACVR1,ARHGDIB,CDK6,ESAM,LPP,SH2B3 | |
| MEF2D | | transcription regulator | Activated | 2.433 | 0.0785 | FOS,FOSB,HDAC9,JUN,JUND,TGFB2,ZFP36 | |
| CREB1 | | transcription regulator | Activated | 2.432 | 3.8E-11 | 1,ETV6,FGF19,FGFR2,FOS,FOSB,GADD45A,GADD45B,HMGCR,HPGD,IDI1,IRF7,IRS2,JUN,KLF4,LCN2,LSS,MAT2A,MCAM,MCL1,MSMO1,MTSS1,MUC2,MYO1B,NEO1,NRP1,PGM2 | 514 (23) |
| UXT | | transcription regulator | Activated | 2.429 | 0.00414 | CCNA2,CXCL8,F5,HPGD,SEC24D,SORD | |
| MEIS1 | | transcription regulator | Activated | 2.425 | 0.00112 | BAZ2B,FOS,HOXA3,HOXA5,MYB,RUNX2,TSC22D2 | |
| Nfat (family) | | group | Activated | 2.42 | 0.0000401 | ATF3,CCN1,CXCL8,F2RL1,FAS,HBEGF,ITPR2,KLF6,NCOA7,PGM2L1,PTGS2,RCAN1,RNF128,SAMD4A,SDC4,TGFB2 | 439 (19) |
| EDN1 | 0.671 | cytokine | Activated | 2.413 | 0.000971 | ANXA1,CCN1,CDC25A,CXCL8,CXCR4,EDN1,EGR1,EREG,ERRFI1,FOS,FOSB,FOSL1,HBEGF,ITPR2,JUN,MCAM,MMP7,PRKCA,PTGS2,REN,TGM2 | 463 (24) |
| IL36A | | cytokine | Activated | 2.412 | 0.00655 | CXCL1,CXCL2,CXCL3,CXCL8,LCN2,NFKBIZ | 421 (18) |
| JAK1 | | kinase | Activated | 2.412 | 0.0551 | BCL3,EIF2AK2,FOS,HLA-C,IRF1,IRF7,LIF | |
| N-acetylsphingosine | | chemical reagent | Activated | 2.408 | 0.024 | CXCL2,CXCL8,FOS,JUN,PTGS2,UGCG | |
| SB 216763 | | chemical - kinase inhibitor | Activated | 2.408 | 0.0346 | CCN1,CXCL8,FOSB,PTGS2,RGS2,RND3 | |
| 4-methylnitrosoamino-1-(3-pyridinyl)-1-butanone | | chemical toxicant | Activated | 2.405 | 0.0142 | ANXA1,FOS,JUN,PTGS2,SERPINA1,TBXAS1 | |
| doxorubicin | | chemical drug | Activated | 2.401 | 1.01E-12 | SP5,EDN1,EGR1,ELF3,ERN1,FAS,FOS,GADD45A,GDF15,HBEGF,HIPK2,HSPH1,ID2,IGFBP6,IRF1,IRF7,JUN,LAMC2,LCN2,LGALS3,LGALS3BP,LIMA1,LOC102724788/PRODH,MCL1 | 510 (18) |
| IL17A | | cytokine | Activated | 2.4 | 0.00000017 | CXCL1,CXCL2,CXCL3,CXCL8,CXCR4,ELF3,EREG,FAS,FOS,FOSL1,GADD45A,HBEGF,IL1RN,ITPR2,JUN,LCN2,LIF,MCL1,NFKBIZ,NME3,NRP1,PI3,PLXNB2,PPARG,PTGS2,SGK1, | 469 (16) |
| nelfinavir | | chemical drug | Activated | 2.397 | 0.00013 | ATF3,ATP2A3,CEBPA,CHAC1,JDP2,KLF4,PCK2,PPARG,PPP1R15A,PSPH,SESN2,TSC22D3,ZNF674 | |
| isotretinoin | | biologic drug | Activated | 2.395 | 3.31E-10 | ALDH1A3,CALB2,CRABP2,CXCL8,ELF3,HOXA5,HPGD,IFIT3,IRF1,LCN2,PLAAT4,PRKCA,PSMB10,PTGS2,RARG,RARRES1,S100A2,SERPINA3,TMPRSS4,TNFAIP2 | 547 (22) |
| RPS6KA5 | | kinase | Activated | 2.395 | 0.000393 | CDKN2B,DUSP1,FOS,FOSL1,JUN,PTGS2 | 422 (18) |
| TNFSF13B | | cytokine | Activated | 2.393 | 0.0484 | BCL6,CTSS,CXCL8,DUSP5,JUN,MAP3K14,MCL1 | |
| GRP | | growth factor | Activated | 2.388 | 0.00246 | CXCL8,FOS,FOSB,JUN,PRKCA,PTGS2 | 438 (23) |
| IFNB1 | | cytokine | Activated | 2.382 | 0.000000466 | 2,CXCL3,CXCL8,CYP1A1,CYP3A5,EIF2AK2,F2R,FOS,GBP3,GLIS2,GPC4,HMGCR,IDI1,IFIH1,IFIT3,IRF1,IRF7,NMI,PARP12,PARP14,PDK4,PLAAT4,PMAIP1,PML,PTGS2,REL,RNAS | 318 (15) |
| TRAF6 | | enzyme | Activated | 2.373 | 0.00234 | BIRC3,CXCL1,CXCL16,CXCL2,CXCL8,MAP1LC3B,MAT2A,MCL1,PAG1,SERPINA3 | 402 (18) |
| F2R | 0.689 | G-protein coupled receptor | Activated | 2.373 | 2.28E-08 | ANGPT1,CASP4,CCN1,CD44,CD55,CXCL3,CXCL8,DUSP1,EGR1,F2R,FGFR2,FOS,HMGA2,MCAM,PTGS2,S100A4,TFPI,TGFB2,TGM2,TJP1,TJP3 | 495 (22) |
| amphetamine | | chemical drug | Activated | 2.36 | 0.000071 | ATF3,BMP4,DDC,EGR1,FGF19,FOS,FOSB,FOSL1,JUN,MAOB,PTPN21,RGS2,SGK1,TGFBR2 | 357 (22) |
| ABT-737 | | chemical drug | Activated | 2.359 | 0.000393 | CXCL3,CXCL8,GADD45A,IRF1,MCL1,PMAIP1 | 439 (16) |
| uranyl nitrate | | chemical toxicant | Activated | 2.355 | 0.0414 | ATP5MD,CYP2B6,CYP3A5,ENTPD5,ID2,IFIT3,LGALS3,NUDT19,STARD10,UBC | |
| FOXO4 | | transcription regulator | Activated | 2.343 | 0.00000642 | ANGPT1,BCL6,CAV1,CCNG2,CDKN2B,ELOVL6,FLNB,FYN,GADD45A,GADD45B,HMGCR,IDI1,MAP1LC3B,OVOL1,PTPRB,RUNX2,SGK1,TXNIP | 380 (12) |
| salirasib | | chemical drug | Activated | 2.339 | 0.00115 | AK4,ATF3,CA9,CYP2B6,LDHA,PGM1,TGM2,TJP1,TJP2 | |
| etoposide | | chemical drug | Activated | 2.338 | 0.0000257 | ATF3,BBC3,BIRC3,CDC25A,CEBPA,CXCL3,CXCL8,CYP3A5,DDIT4,DUSP1,FAS,GADD45A,GDF15,HK1,ID2,IRF1,IRF7,JUN,LIMA1,MCL1,PFKM,REL,S100A2,SGK1,SVIL | 445 (17) |
| dimethyl sulfoxide | | chemical drug | Activated | 2.337 | 0.0000192 | ABCC2,ANXA1,BMP4,CXCL8,CYP1A1,CYP2B6,CYP3A5,DNAJB4,FOS,FUT8,IRS2,JUN,JUND,LCN2,LOXL4,PLD1,PLD2,RUNX2,SMPD3 | 471 (23) |
| PTPRJ | | phosphatase | Activated | 2.333 | 0.0000728 | CCNA2,CEBPD,CXCL2,CXCL3,EDN1,FOS,HBEGF,NFKBIZ,PTGS2,TSC22D1,ZC3H12A | 455 (22) |
| ARHGAP21 | | other | Activated | 2.333 | 0.000197 | CXCL2,IFIT3,IL15RA,IRF1,NFKBIZ,NMI,PTGS2,TRIM21,TSC22D1 | |
| E. coli B5 lipopolysaccharide | | chemical - endogenous non-mammalian | Activated | 2.332 | 0.0000891 | M,BCL3,BCL6,BIRC3,CD44,CDK6,CEBPA,CXCL2,CXCL3,CXCL8,CXCR4,CYP3A5,DUSP1,ERRFI1,FOS,IL15RA,IL1R2,IRF1,LIF,MAT2A,NEDD4L,PHLDA1,PTGS2,RHOB,TGFB2,TGF | 342 (15) |
| TNFSF10 | | cytokine | Activated | 2.321 | 3.5E-09 | ANGPT1,BIRC3,CASP9,CXCL8,CXCR4,EIF2AK2,FOS,HLA-C,IFITM2,IL1RN,JUN,MCL1,PMAIP1,PPP1R15A,PRAP1,PSME2,PTGS2,RCAN1,SERPINH1,SP100,TNFAIP2,TNK2 | 472 (22) |
| ionomycin | | chemical reagent | Activated | 2.32 | 0.0000764 | ATP2A3,CCNG2,CXCL8,DNAJC3,DUSP1,DUSP2,DUSP5,EGR1,EIF2AK2,ETS1,FAS,FOS,FOSB,ITPR2,LDHA,NRP1,PIM1,PPARG,PPP1R15A,PTGS2,RCAN1,RGS2,SGK1 | 467 (19) |
| reactive oxygen species | | chemical toxicant | Activated | 2.314 | 0.00141 | ABCC1,ADAM9,CXCL8,CXCR4,DUSP2,EDN1,ETS1,FAS,GADD45A,HBEGF,JUN,KLF6,MCL1,PMAIP1,PTGS2 | 467 (20) |
| JAK2 | | kinase | Activated | 2.311 | 0.000235 | ACER2,AKAP12,AKR1C3,BBC3,CDC25A,CXCL8,EGR1,F5,FOS,GCNT2,IRF1,LAT2,LCN2,NFKBIZ,PTGS2,SDC4,SERPINH1,TGM2,TSC22D1 | 419 (21) |
| PAF1 | | other | Activated | 2.309 | 0.0000031 | ARL4A,BIRC3,IFI44,IFIT3,IFITM3,KLF4,MXD1,NFKBIZ,OAS3,SDC4,ZC3HAV1,ZFP36 | |
| FN1 | | enzyme | Activated | 2.294 | 0.000121 | BIRC3,CASP7,CCN1,CDK6,CXCL1,CXCL2,CXCL3,CXCL8,EPB41L1,FOS,HK1,HLA-B,JUND,KRT7,MAT2A,MUC2,RUNX2,SDC4,SNAP23,TGFBR2,TGM2,TNFAIP2,VDAC1,ZYX | 475 (16) |
| AGN194204 | | chemical drug | Activated | 2.287 | 3.19E-13 | COTL1,CPT1A,CRABP2,DUSP1,EMP1,F2RL1,GADD45A,GBP3,GNAI1,GPRC5A,IDI1,IFITM3,IL1R2,IRF1,IRS2,KLF6,KRT80,LSS,NRIP1,PDK4,PHLDA1,PIM1,PLAC8,PPARG,PPL,PR | 269 (11) |
| FCGR2A | | transmembrane receptor | Activated | 2.287 | 0.0003 | CXCL2,CXCL3,CXCL8,F2RL1,FAS,IFIT3,IFITM3,IRF7,OAS3,PTGS2 | 418 (15) |
| IKBKB | | kinase | Activated | 2.272 | 3.07E-10 | CXCL16,CXCL2,CXCL3,CXCL8,CXCR4,EDN1,EGLN1,EGR1,EREG,ETS1,FAS,FOS,FYN,GADD45A,GBP3,GLI1,GRK5,IGFBP2,IGFBP6,IL1RN,IRF1,LCN2,MAT2A,MUC2,PTGS2,RCA | 443 (14) |
| MAPK9 | | kinase | Activated | 2.263 | 1.95E-09 | CAV1,CXCL8,CYP1A1,EDN1,EGR1,FAS,FOS,FOSL1,GADD45A,GADD45B,HMGCR,IGFBP2,IL15RA,JUN,JUND,KLF6,LGALS3BP,LIF,LMO7,NMI,PARP14,PPARG,PPP1R15A,SERPIN | 509 (19) |
| CD40 | | transmembrane receptor | Activated | 2.263 | 0.0000423 | BCL6,BIRC3,CCNA2,CD44,CDK6,CXCL8,DAPP1,DUSP1,ERN1,FAS,FOS,IL15RA,IL1RN,IRF1,JUN,KLF4,KLF9,LGALS3,MAP3K14,PIM1,PSMB10,PTGS2,REL,TAP2,TAPBP,TRAF4 | 341 (17) |
| SMARCA4 | | transcription regulator | Activated | 2.259 | 5.46E-17 | EG,FAS,FGFR2,FHL2,FLNB,FOS,GADD45A,GADD45B,GLI1,GPR158,H1-10,HEPH,HLA-B,HLA-C,HLA-E,HLA-J,HMGA2,HNF4A,HPGD,IFITM2,IFITM3,IL15RA,IRF1,JUN,KDM6B,LAMC | 507 (21) |
| deoxycholate | | chemical - endogenous mammalian | Activated | 2.257 | 0.000748 | CAV1,CXCL8,CYP2B6,EGR1,FGF19,GJB2,HPGD,MCL1,PTGS2 | 454 (20) |
| RUNX2 | 0.922 | transcription regulator | Activated | 2.254 | 0.021 | AXIN2,CDHR5,CEBPA,CEBPD,CXCL8,EDN1,FGFR2,GADD45B,LGALS3,RUNX2,SVIL,UGCG | |
| CHD1 | | enzyme | Activated | 2.236 | 0.00106 | CXCL1,CXCL2,CXCL3,LIF,PTGS2 | |
| CYP51A1 | | enzyme | Activated | 2.236 | 0.000259 | HMGCR,LSS,MSMO1,SC5D,SQLE | |
| SPIB | | transcription regulator | Activated | 2.236 | 0.331 | ATF3,EGR1,HSPA1A/HSPA1B,KDM6B,KLF6,PRDX4 | |
| ribavirin | | chemical drug | Activated | 2.236 | 0.0158 | APOBEC3B,CXCL8,EIF2AK2,IFIT3,IRF1,IRF7 | |
| D-galactosamine | | chemical - endogenous mammalian | Activated | 2.236 | 0.0177 | CXCL2,FOS,GDF15,JUN,JUND,MCL1 | |
| ST3-Hel2A-2 | | chemical reagent | Activated | 2.236 | 0.000101 | ATF3,BCL6,FOS,GADD45A,PHLDA1 | 261 (8) |

Supplementary Table 4. Upstream regulator analysis based gene set enrichment analysis (GSEA) of differentially expressed genes (DEGs) after ATRA+PDT compared to PDT

| Upstream Regulator | Expr Log Ratio | Molecule Type | Predicted Activation State | Activation z-score | p-value of overlap | Target Molecules in Dataset | Mechanistic Network |
|---|---|---|---|---|---|---|---|
| GNRH1 | | other | Activated | 2.229 | 0.154 | EGR1,FOS,FOSB,JUN,PTGS2 | |
| 1-methyl-4-phenyl-1,2,3,6-tetrahydropyridine | | chemical toxicant | Activated | 2.228 | 0.0139 | CASP9,CLDN1,EGR1,FOS,FOSB,HSPB1,JUN,LIF,PTGS2 | |
| NFATC3 | | transcription regulator | Activated | 2.226 | 0.00234 | ANGPT1,ATF3,DDIT4,EDN1,FOS,FOSB,JUN,MUC2,PPARG,PTGS2 | 317 (12) |
| Ni2+ | | chemical reagent | Activated | 2.224 | 0.00546 | BIRC3,CXCL1,CXCL2,CXCL3,CXCL8 | 441 (16) |
| NfkB1-RelA | | complex | Activated | 2.222 | 0.0446 | CXCL1,CXCL3,CXCL8,LCN2,PTGS2 | |
| PTEN | | phosphatase | Activated | 2.222 | 8.58E-11 | N1,ELOVL6,ERRFI1,ETS1,FAS,FGF19,FGFR2,FOS,GPR160,GTPBP2,HMGCR,HSPA1A/HSPA1B,IDI1,IFNGR2,IGF2,IGFBP2,IGFBP6,INPPL1,IRS2,JAG2,JUND,KLF6,KLHL24,LGALS | 510 (18) |
| EP300 | | transcription regulator | Activated | 2.222 | 8.24E-12 | CTSE,CXCL3,CXCL8,CXCR4,CYP1A1,DDC,DHRS3,DNAJB4,DUSP1,EDN1,EGR1,FOS,FRMD4B,GLCCI1,HES6,HLA-B,HMGCR,HMGCS2,ID2,IGFBP6,IL15RA,JUN,LAMC2,LDHA,LGA | 512 (23) |
| HOXA5 | 0.429 | transcription regulator | Activated | 2.219 | 0.00546 | CXCL8,EGR1,GADD45B,RUNX2,SAT1 | |
| PBX3 | | transcription regulator | Activated | 2.219 | 0.00367 | BAZ2B,HOXA3,HOXA5,RUNX2,TSC22D2 | |
| SLC27A2 | | transporter | Activated | 2.216 | 0.0579 | ACAA1,ACSL1,ALDH1A1,ME1,PEX11A | |
| Pam3-Cys-Ser-Lys4 | | chemical reagent | Activated | 2.214 | 0.000353 | CASP4,CD55,CD58,CLDN1,CXCL2,CXCL3,CXCL8,IRF1,IRF7,JUN,KDM6B,MCL1,NFKBIZ,PARP14,PTGS2,REL,RGS2,SLC16A3,TGFB2 | 354 (15) |
| STAT | | group | Activated | 2.213 | 0.0368 | CEBPD,EIF2AK2,FOS,IRF1,SERPINA3 | |
| NPPB | | other | Activated | 2.213 | 0.0212 | HMGCR,IDI1,LSS,MSMO1,SCARB1 | |
| imiquimod | | chemical drug | Activated | 2.208 | 0.00119 | BCL10,CXCL2,CXCL8,FOS,FOSL1,GLI1,IFI35,IL1RN,IRF7,LCN2,PTGS2,RNF213,XPA | 328 (20) |
| IFT88 | | other | Activated | 2.208 | 0.000188 | ACAT1,CEBPA,GLI1,HMGCR,IDI1,LSS,PPARG,RUNX2 | 354 (12) |
| MALP-2s | | chemical reagent | Activated | 2.207 | 0.0107 | CXCL2,CXCL3,IL15RA,MCL1,PTGS2 | |
| MAP3K8 | | kinase | Activated | 2.207 | 0.00056 | ARRDC3,BIRC3,CEBPD,CLIC5,CXCL2,CXCL3,CXCL8,FAM107B,FLNB,FOS,GAB1,GPR160,JUN,PTGS2,SLCO4A1,TAPBPL,TXNIP,ZC3H6 | 357 (16) |
| USF1 | | transcription regulator | Activated | 2.204 | 0.000561 | AKAP12,B2M,CCNG2,CEACAM1,CEBPA,CPT1A,CTSS,DUSP1,FBXO32,PTGS2,REN,TXNIP | 203 (7) |
| Pdgf (complex) | | complex | Activated | 2.203 | 0.000117 | CAV1,CD44,CDK6,CDKN2B,CXCL3,EDN1,EGR1,ERRFI1,FOS,FOSB,IRS2,JUN,NOX1,PPARG,PTGS2 | 427 (19) |
| phenylephrine | | chemical drug | Activated | 2.202 | 0.0000358 | ANGPT1,DUSP1,EDN1,EGR1,FOS,ITPR2,JUN,PDK4,PTGS2,RCAN1,REN,RGS2,STEAP3,UCA1 | 477 (25) |
| adenosine | | chemical - endogenous mammalian | Activated | 2.2 | 0.0268 | CXCL8,CXCR4,DPP4,FOS,PMAIP1,PTGS2,TXNIP | |
| taurocholic acid | | chemical - endogenous mammalian | Activated | 2.2 | 0.0144 | ATF3,EGR1,GADD45B,NR0B2,PTGS2 | |
| bile acid | | chemical - endogenous mammalian | Activated | 2.199 | 0.00565 | CAV1,FGF19,HNF4A,MUC2,NR0B2,PTGS2 | 506 (22) |
| PRKD1 | | kinase | Activated | 2.193 | 0.17 | AJUBA,CXCL2,CXCL3,ERN1,PTGS2 | |
| USP19 | | peptidase | Activated | 2.19 | 0.00546 | BIRC3,CBLB,DDIT4,FBXO32,LCN2 | |
| CD36 | | transmembrane receptor | Activated | 2.189 | 0.0538 | CXCL1,CXCL3,CXCL8,ERN1,FAS,IGFBP6,LIF,PNPLA2,PPARG | |
| ITGB3 | | transmembrane receptor | Activated | 2.186 | 0.128 | CEACAM1,FOS,PTGS2,RUNX2,SMAD3,TGFBR2 | |
| VitaminD3-VDR-RXR | | complex | Activated | 2.178 | 0.00161 | CYP24A1,GADD45A,IGFBP6,KLF4,KLK6,MXD1,SEMA3B,TGFB2 | |
| ozone | | chemical toxicant | Activated | 2.172 | 0.00919 | CXCL2,CXCL3,CXCL8,FOS,JUN | 421 (17) |
| collagen type i (family) | | group | Activated | 2.169 | 0.0107 | ABCC1,CLRN3,HNF4A,IL1RN,LGALS4 | |
| silicon dioxide | | chemical drug | Activated | 2.169 | 0.00106 | CXCL8,EGR1,FOS,FOSL1,JUN | 415 (20) |
| FOXO1 | | transcription regulator | Activated | 2.166 | 1.21E-08 | N1,EGR1,ELOVL6,FAS,FBXO32,FLNB,FOS,FOSB,FYN,GADD45A,GADD45B,GATA2,GPD1,HMGCR,IRS2,ITGB6,JUN,KLF4,MAP1LC3B,ME1,MRPL34,MRPS2,NPC1,OVOL1,PCK2,P | 465 (20) |
| TEAD1 | | transcription regulator | Activated | 2.164 | 0.000723 | AMOTL2,ARID5B,CCN1,CDK6,EDN1,LATS2,MSLN,NFKBIZ,PTGS2,SLFN5,TRNP1 | 357 (12) |
| ABCB6 | | transporter | Activated | 2.158 | 0.0579 | CYP1A1,CYP2B6,CYP3A5,HK1,SLC25A39 | |
| AMPK | | complex | Activated | 2.158 | 0.0354 | CLDN1,CXCL8,CYP2B6,EGR1,FBXO32,IL1RN,NR0B2,PMAIP1,PPARG,PTGS2,RUNX2 | |
| IL6R | | transmembrane receptor | Activated | 2.156 | 0.00314 | ABCC1,CXCL2,CXCL3,CXCL8,IGF2,IRF1,LCN2,MCL1,PTGS2,SERPINA3,TGFBR2 | 438 (21) |
| TSH | | complex | Activated | 2.156 | 0.104 | CTSS,CXCL8,EGR1,FOS,GNAI1,IL1RN | |
| carbon tetrachloride | | chemical toxicant | Activated | 2.152 | 0.0000015 | S1,ATF3,CASP7,CYP1A1,DUSP1,FAAH,FOS,GADD45A,GADD45B,GDF15,HSPB1,IGFBP6,JUN,JUND,ME1,NDRG2,NR0B2,PPARG,PPP1R15A,PTGS2,SMAD3,SORD,SPHK2,TGFB | 510 (19) |
| PTH | | other | Activated | 2.142 | 0.0000529 | ANGPT1,CXCL2,CXCR4,CYP24A1,DUSP1,ESAM,FOS,FOSB,JUN,LIMA1,NDRG2,PDK4,PPP1R1B,PTGS2,RAP1GAP,RGS2,RUNX2,SDC4,SEC61B,SMAD3,TJP2,VDR | 449 (22) |
| IFI16 | | transcription regulator | Activated | 2.138 | 0.0394 | CDC25A,CXCL8,CYP1A1,EDN1,GADD45A,ID2,IL1RN,PIM1 | |
| LCK | | kinase | Activated | 2.137 | 0.00917 | ANXA1,CCNA2,EGR1,FOS,GADD45B,IRF1,JUN,PPARG,SCARB1 | 466 (23) |
| NORAD | | other | Activated | 2.121 | 0.000731 | CCN1,FSTL3,GADD45B,JUN,RHOB,SEC14L2,SGK1,TNFRSF12A | 204 (7) |
| TEAD3 | | transcription regulator | Activated | 2.121 | 0.00358 | AMOTL2,ARID5B,CCN1,EDN1,LATS2,NFKBIZ,SLFN5,TRNP1 | |
| TNFSF12 | | cytokine | Activated | 2.12 | 0.0158 | ADAM8,CD68,CXCL2,CXCL3,CXCL8,FBXO32,HES6,IL1R2,JAG2,MAP3K14,TNFRSF12A | |
| TREM1 | | transmembrane receptor | Activated | 2.115 | 1.81E-12 | CL3,CXCL8,CXCR4,E2F7,EDN1,EGR1,FOSL1,GADD45B,GPRC5A,HBEGF,IL15RA,IRF1,KANK1,LIF,MAFF,MT1E,NEDD4L,NPC1,NT5E,PHLDA1,PLAC8,PLCXD1,PLD1,PLPP3,PPAR | 444 (19) |
| l-asparaginase | | biologic drug | Activated | 2.111 | 0.0271 | AURKA,CCNA2,E2F7,ERCC6L,HMGCR,IDI1,INCENP,INSIG1,LIPG,MSMO1,SQLE | |
| tosedostat | | chemical drug | Activated | 2.111 | 8.62E-08 | ASS1,ATF3,CCNG2,CHAC1,CXCL8,DDIT4,FYN,GADD45A,PPP1R15A,SESN2,WARS1 | |
| IL12 (family) | | group | Activated | 2.109 | 0.0863 | ATF3,BCL3,BCL6,CD44,DDIT4,GMEB2,IRF1,LIF,NFKBIZ,PIM1,TRIM29 | |
| MAP2K4 | | kinase | Activated | 2.104 | 0.00000353 | ATF3,BIRC3,CD44,CXCL8,ERRFI1,FAS,FOS,GADD45A,JUN,PPARG,PTGS2,TUBB2A,VDR | 426 (16) |
| Lh | | complex | Activated | 2.104 | 0.000000251 | P9,CD55,COL18A1,CXCL8,CXCR4,DAPK1,DHRS3,DUSP1,EREG,FOS,GRK5,MAP1LC3B,MAP4K4,MCL1,MSMO1,PMAIP1,PPP2R5B,PRKAR2A,PTGS2,PTPN21,RHOB,RUNX2,SGK1 | 472 (22) |
| RAS | | group | Activated | 2.101 | 0.00000673 | AURKA,CAV1,CDKN2B,CEBPA,CXCL8,CYP24A1,EDN1,EGR1,FOS,FOSL1,IRF1,JUN,LIF,MAOB,NOX1,PLD1,PML,PTGS2,SERPINE2,TGM2 | 394 (23) |
| CXCL12 | | cytokine | Activated | 2.096 | 0.0107 | AKR1C3,BCL3,CD44,CXCL8,CXCR4,CYP3A5,DPP4,EGR1,FAS,FOS,FYN,HLA-B,IFNGR2,JMJD1C,JUN,MAPRE3,PTGS2,RUNX2 | |
| EIF2AK3 | | kinase | Activated | 2.081 | 0.0000975 | ATF3,BIRC3,CA9,DNAJB9,DNAJC3,EGR1,ERN1,FOSL1,GADD45A,JUN,KLF4,MAP1LC3B,PCK2,PPP1R15A,SEC24D,SEC61A1,TXNIP,WARS1,ZFP36 | 420 (23) |
| sphingosine-1-phosphate | | chemical - endogenous mammalian | Activated | 2.078 | 0.000169 | ANGPT1,CD44,CXCL2,CXCL8,EGR1,ERN1,ETS1,FAS,FOS,GATA6,JUN,NRP1,PTGS2,RUNX2 | 443 (18) |
| 1,4-bis[2-(3,5-dichloropyridyloxy)]benzene | | chemical toxicant | Activated | 2.076 | 0.0000196 | ABCC2,CCNA2,CDK6,CPT1A,CYP2B6,CYP3A5,FAM107B,GADD45B,HMGCS2,INSIG1,MAFF,NEDD9,PNPLA2,RARRES1,SAT1,STEAP4,TNFAIP2,TUBA4A | 241 (7) |
| NFATC2 | | transcription regulator | Activated | 2.07 | 0.0000901 | AQP5,CASP4,CCNA2,CCNF,CXCL2,CXCL3,EDN1,FAS,IFIT3,IRF1,IRF7,JUN,KDM6B,LDHA,NFKBIZ,NMI,PLD1,PML,PPARG,PTGS2,RCAN1,REL,RGS2,TSC22D1 | 496 (19) |
| IL18 | | cytokine | Activated | 2.067 | 0.0029 | ATF3,BCL6,CD44,CXCL16,CXCL3,CXCL8,DDIT4,FAS,GADD45B,IRF1,JUN,LIF,MMP15,NFKBIZ,PTGS2,SMAD3,TGFBR2,TRIM29,TXK | 438 (17) |
| EPO | | cytokine | Activated | 2.061 | 0.00000917 | BTG1,CCNG2,CXCR4,EDN1,EGR1,FAS,FOS,FOSB,GATA2,GDF15,ID2,IGF2,JUN,KLHL24,MCL1,MYB,PIM1,PRKCA,PTGS2,PTPRB,REN,RNF128,SLC24A1,SNAP23,ST3GAL4,TJP1, | 510 (22) |
| TGFB1 | | growth factor | Activated | 2.054 | 7.82E-33 | L1,FSTL3,FUT8,FYN,GADD45A,GADD45B,GALM,GDF15,GJB2,GLI1,GLI2,GNA14,GNAZ,GNPAT,GPR158,GPRC5A,GSE1,GSN,GYG1,H1-10,HBEGF,HDAC9,HK1,HMGA2,HNF4A,HP | 629 (22) |
| MRTFB | | transcription regulator | Activated | 2.052 | 0.0000283 | AMOTL2,ARID5B,CCN1,CXCR4,DNAJB4,EDN1,EGR1,ETV6,F2R,GADD45A,HPGD,LCN2,NFKBIZ,PTGS2,RAB27B,SCARB1,SGK1,SLFN5,TBXAS1,TGFB2,TRNP1,TST | 314 (14) |
| Z-LLL-CHO | | chemical - protease inhibitor | Activated | 2.049 | 0.00000268 | F3,AURKA,BIRC3,CEBPA,CLU,CXCL2,CXCL8,CYP1A1,DUSP1,ETS1,FAS,FOS,GLI1,GNAI1,HSPB1,IRF7,IRS2,ITPR2,JUN,MAT2A,MCL1,MUC2,MYB,PLK3,PMAIP1,PPARG,PTGS2,R | 471 (19) |
| hyaluronic acid | | chemical - endogenous mammalian | Activated | 2.041 | 0.0242 | BCL3,CD44,CXCL2,CXCL3,CXCL8,CXCR4,FOS,JUN,PPARG,PTGS2,TGFB2 | |
| ACSL4 | | enzyme | Activated | 2.041 | 0.0000728 | CALB2,CCDC28A,CDKN2B,DOCK5,NT5E,PPARG,PTGS2,RHOU,SAT1,STK39,TFPI | |
| TAC1 | | other | Activated | 2.028 | 0.0109 | CCN1,CXCL3,CXCL8,CXCR4,FOS,FOSB,FOXL1,IL1RN,KITLG,PTGS2 | |
| Igm | | complex | Activated | 2.008 | 0.000438 | CCNA2,CD44,CXCL8,DUSP2,EGR1,FAS,MCL1,PIM1,POLI,UGCG,ZFP36L1 | 473 (22) |
| PARPBP | | other | Activated | 2 | 0.0207 | AMD1,KRT7,REN,TST | |
| motexafin gadolinium | | chemical drug | Activated | 2 | 0.000593 | MT1E,MT1X,MT2A,SLC30A1 | |
| RNY3 | | other | Activated | 2 | 0.041 | IFI44,IFIT3,IFITM3,OAS3 | |
| Smad2/3 | | group | Activated | 2 | 0.187 | CDKN2B,FSTL3,MXD1,RUNX2,TGM2 | |
| PROM1 | | other | Activated | 2 | 0.00507 | IDI1,INSIG1,LSS,SQLE | 181 (7) |
| RNF138 | | enzyme | Activated | 2 | 0.0279 | BIRC3,CXCL1,GADD45B,PLK3 | |
| IFNK | | cytokine | Activated | 2 | 0.0241 | EIF2AK2,IFIH1,IRF1,PLAAT4 | |
| IFNAR2 | | transmembrane receptor | Activated | 2 | 0.0218 | IFI44,IFIH1,PSMB10,PSMB8,PSME2,TGM2 | |
| CYBB | | enzyme | Activated | 2 | 0.098 | CXCL3,DUSP1,PPARG,PTGS2 | |
| IL12B | | cytokine | Activated | 2 | 0.328 | CEBPD,CXCL8,DUSP5,ERN1 | |
| LTB4R | | G-protein coupled receptor | Activated | 2 | 0.0571 | CXCL2,CXCL3,CXCL8,PTGS2 | |
| DGAT1 | | enzyme | Activated | 2 | 0.000387 | CPT1A,DGAT2,PDK4,PNPLA2,PPARG | 428 (19) |
| BTC | | growth factor | Activated | 2 | 0.00507 | CXCL8,EREG,IRS2,PTGS2 | 472 (23) |
| BAG1 | | other | Activated | 2 | 0.00135 | FOS,GADD45A,JUN,MT2A,NEDD4L,PTGS2 | 481 (22) |
| acetaldehyde | | chemical - endogenous mammalian | Activated | 2 | 0.0176 | FOS,JUN,KLF9,PPARG | |
| leukotriene C4 | | chemical - endogenous mammalian | | 1.997 | 0.00818 | CXCL8,EGR1,FOS,PTGS2 | 454 (19) |
| SYVN1 | | transporter | | 1.997 | 6.47E-09 | ,ATP1B3,CD44,CSPG4,DUSP1,EPHA2,ERN1,FAS,GPRC5A,HLA-C,HSPB1,IFI44,IFITM2,LGALS3BP,MCAM,MYOF,PLD2,PLPP3,PNPLA2,SLC16A1,SLC30A1,SLC39A10,SLCO4A1,S | |
| TCR | | complex | | 1.994 | 0.00000901 | XCR4,EGR1,F5,FAS,FOS,FYN,GADD45B,IFI44,IFIH1,IFIT3,IRF1,IRF7,JUN,KCNK5,MARF1,MCL1,NRP1,PIK3R1,PIM1,PMAIP1,REL,RNF128,SEMA3A,SERPINA3,SOX12,TNFRSF12 | 378 (18) |
| CpG ODN 1668 | | chemical reagent | | 1.991 | 0.317 | CXCL3,CXCL8,IRF7,PTGS2 | |
| RELA | | transcription regulator | | 1.988 | 1.49E-11 | 2B6,DMTF1,DUSP1,EDN1,EGLN1,EGR1,EHF,ELF3,ERAP2,FAS,FOS,FOSB,GDF15,GLI2,GNPAT,GRK5,HLA-B,HNF4A,IFNGR2,IGFBP2,IL15RA,IL1RN,IRF1,IRF7,JUN,MAT2A,MMP7 | 485 (17) |
| oltipraz | | chemical drug | | 1.987 | 0.0632 | AKR7A3,CYP1A1,MAP3K14,UGT1A10 (includes others) | |
| CSF2RB | | transmembrane receptor | | 1.987 | 0.00507 | FOS,JUN,PIM1,SNAP23 | 456 (22) |
| TXN | | enzyme | | 1.987 | 0.218 | BCL6,CXCL8,CYP1A1,FOS | |
| EPHB1 | | kinase | | 1.987 | 0.00144 | EGR1,FOS,JUN,PTGS2 | 460 (23) |
| ELOVL5 | | enzyme | | 1.987 | 0.0461 | ELOVL6,HMGCS2,PDK4,PNPLA2 | |
| CFB | | peptidase | | 1.987 | 0.0632 | ACO1,CXCL2,CXCL3,TGFB2 | |
| 1-chloro-2-(2,2,2-trichloro-1-(4-chlorophenyl)ethyl)benzene | | chemical toxicant | | 1.987 | 0.00287 | CYP2B6,CYP3A5,GADD45A,GADD45B | |
| NAMPT | | cytokine | | 1.984 | 0.00521 | CXCL1,CXCL2,CXCL8,EPHA2,JUN,TFPI,TGFB2 | 451 (18) |
| ERVW-1 | | other | | 1.982 | 0.0148 | CCNA2,CXCL1,CXCL8,LATS2 | |
| NQO1 | | enzyme | | 1.982 | 0.114 | BIRC3,CEBPA,CXCR4,PTGS2 | |
| HOXB13 | | transcription regulator | | 1.982 | 0.0514 | CEP170,NIPAL1,SEMA3A,SPINK1 | |
| IFNE | | cytokine | | 1.982 | 0.158 | IFIH1,IFITM3,PLAAT4,PTGS2 | |
| HNRNPK | | other | | 1.982 | 0.029 | KDM6B,LPP,PPARG,PTGS2,RASA1,TCN2 | |
| CRP | | other | | 1.982 | 0.0579 | CXCL8,EGR1,IL1RN,PRKCA,PSME2 | |
| 2-mercaptoacetate | | chemical drug | | 1.982 | 0.00367 | CD68,CXCL2,CXCL3,PPARG,S100A10 | 285 (12) |
| Zn2+ | | chemical - endogenous mammalian | | 1.982 | 0.0207 | FOS,MT2A,SLC30A1,SLC39A8 | |
| SMAD3 | 0.495 | transcription regulator | | 1.981 | 0.00000802 | KN2B,CRISPLD2,CXCL3,CYP1A1,DAPK1,EDN1,EGR1,ENG,EREG,FOS,FSTL3,GADD45B,GLI1,GLI2,HBEGF,HNF4A,ID2,JUN,JUND,MUC2,MXD1,PCSK5,PTGS2,RHOB,RUNX2,SM | 519 (20) |

Supplementary Table 4. Upstream regulator analysis based gene set enrichment analysis (GSEA) of differentially expressed genes (DEGs) after ATRA+PDT compared to PDT

| Upstream Regulator | Expr Log Ratio | Molecule Type | Predicted Activation State | Activation z-score | p-value of overlap | Target Molecules in Dataset | Mechanistic Network |
|---|---|---|---|---|---|---|---|
| arsenic trioxide | | chemical drug | | 1.981 | 0.00000131 | CASP7,CASP9,CD44,CDK6,CXCL8,DEDD2,DNMT3A,FAS,GLI1,GLI2,HMGCR,IDI1,IRF1,LOC102724788/PRODH,LSS,MCL1,ME1,MED15,MSMO1,MT1E,MT1X,MT2A,NQO2,PIM1,PML, | 459 (19) |
| CHRM1 | | G-protein coupled receptor | | 1.981 | 0.00287 | CCN1,EGR1,FOS,JUN | 384 (18) |
| cyclooxygenase | | group | | 1.98 | 0.0148 | FOS,LIF,PTGS2,RGS2 | |
| ALOX12 | | enzyme | | 1.98 | 0.00207 | CXCL3,FOSL1,JUN,TGFBR2 | 446 (22) |
| CSF2RA | | transmembrane receptor | | 1.98 | 0.00287 | FOS,JUN,PIM1,SNAP23 | 369 (16) |
| WNT3 | | other | | 1.98 | 0.00387 | GLI1,GLI2,PTGS2,RUNX2 | 346 (12) |
| phorbol esters | | chemical - other | | 1.978 | 0.000000367 | ANXA1,B2M,CD44,CXCL8,CYP24A1,DUSP1,DUSP2,EGR1,FGFR2,FOS,HOXA5,JUN,LIF,PRKCA,PSCA,PTGS2,REL,VDR | 411 (20) |
| heparin | | chemical - endogenous mammalian | | 1.977 | 0.336 | DUSP1,FOS,PROCR,RUNX2,TFPI | |
| NLRC5 | | transcription regulator | | 1.976 | 0.0176 | B2M,HLA-B,HLA-C,HLA-E | |
| NUP98-DDX10 | | fusion gene/product | | 1.976 | 0.000000341 | ANGPT1,CDK6,EGR1,GADD45B,HOXA3,HOXA5,MCL1,MYB,NDRG2,PTGS2,REN,SOX4,ZFHX3 | |
| capsaicin | | chemical drug | | 1.975 | 0.000254 | AQP5,CXCL3,CXCL8,CYP3A5,FOS,GDF15,KITLG,PPARG,PTGS2,SGK1,TGFB2 | 472 (18) |
| CGAS | 0.426 | enzyme | | 1.974 | 0.041 | CXCL8,IFI44,IFIT3,IRF7 | |
| P2RY2 | | G-protein coupled receptor | | 1.974 | 0.0319 | CXCL8,JUN,PTGS2,SESN2 | |
| TLR5 | | transmembrane receptor | | 1.973 | 0.114 | ATF3,CXCL2,CXCL3,CXCL8 | |
| anandamide | | chemical - endogenous mammalian | | 1.972 | 0.131 | BBC3,FOS,PPARG,PTGS2 | |
| EBI3 | | cytokine | | 1.97 | 0.019 | B2M,CTSE,FOS,HLA-B,HLA-C,IRF1,PIM1 | |
| Pam3-Cys | | chemical toxicant | | 1.97 | 0.187 | CXCL2,CXCL3,CXCL8,IL1RN | |
| CDK9 | | kinase | | 1.969 | 0.0174 | CXCL2,CXCL8,CXCR4,FOS,GATA2,PDK4,PPARG | |
| peroxynitrite | | chemical toxicant | | 1.969 | 0.0123 | CXCL8,FOS,PTGS2,RGS2 | |
| KLF4 | -0.393 | transcription regulator | | 1.966 | 0.00000017 | RABP2,CXCL8,CYP1A1,DUSP1,DUSP5,DYRK2,EPHA2,ETV1,GADD45A,GBP3,GJB2,GRHL3,HBEGF,HSPB1,IFITM3,KLF4,KLHL24,KRT13,KRT7,MUC2,NRP1,OVOL1,PPL,RUNX2,S | 491 (23) |
| IL1R1 | | transmembrane receptor | | 1.965 | 0.00864 | ABCC1,CXCL1,CXCL2,CXCL3,CXCL8,EGR1,FOS,PTGS2 | 419 (17) |
| MAP3K3 | | kinase | | 1.964 | 0.00546 | CXCL8,FOS,JUN,STEAP4,TGFB2 | 371 (19) |
| TBX5 | | transcription regulator | | 1.964 | 0.191 | CXCL2,CXCL8,CXCR4,PTGS2,PTPRB,S100A4 | |
| finasteride | | chemical drug | | 1.964 | 0.0148 | CLU,EDN1,FOS,TGFB2 | |
| TERT | | enzyme | | 1.964 | 0.000268 | ALDH1A3,ATF3,B2M,CAV1,CXCL8,EIF5A2,EREG,ETS1,F2R,FUT4,GATA6,GSN,HLA-B,IQGAP2,LIPG,PSCA,PTGS2,SMAD3 | 520 (24) |
| thyroid hormone | | chemical - endogenous mammalian | | 1.964 | 0.02 | AK4,CCNA2,CDKN2B,CPT1A,DDC,DNAJC3,EGR1,FOS,HMGCR,IGFBP2,KLF9,ME1,PML,S100A10,SERPINE2,UQCRHL | |
| ISGF3 | | complex | | 1.961 | 0.00287 | EIF2AK2,IFIH1,IRF1,IRF7 | |
| IPMK | | kinase | | 1.961 | 0.0279 | CCN1,FOS,FOSB,JUN | |
| EHD2 | | other | | 1.961 | 0.000343 | CAV1,CAV2,PNPLA2,PPARG | |
| ALB | | transporter | | 1.961 | 0.168 | CXCL8,EDN1,ERN1,PTGS2 | |
| di(2-ethylhexyl) phthalate | | chemical toxicant | | 1.961 | 0.000468 | CEACAM1,CXCL1,CXCL8,DPP4,EGR1,FOS,FOSB,JUN,PEX11A,ZFP36 | 419 (16) |
| EWSR1 | | other | | 1.96 | 0.0831 | F2RL1,FOS,KCNE3,MCL1 | |
| bisphenol A | | chemical - endogenous mammalian | | 1.956 | 0.0112 | AQP5,CCN1,CLU,EGR1,ELAC2,EREG,FOS,HSPA1A/HSPA1B,LCN2,MAOA,MPP6,PPARG,PPP1R10,PTGS2,RBPMS,SGK1,SRD5A3 | |
| Ras homolog | | group | | 1.953 | 0.0176 | EGR1,FOS,PTGS2,TJP1 | |
| SMAD1 | | transcription regulator | | 1.953 | 0.0264 | CDKN2B,CXCL2,CXCL3,GADD45B,IRF1,RUNX2 | |
| N-methyl-D-aspartate | | chemical drug | | 1.952 | 0.373 | EGR1,FOS,LIF,PTGS2 | |
| MAP2K7 | | kinase | | 1.951 | 0.0017 | ATF3,BIRC3,CXCL8,FAS,HSPA1L,LMO7,MAOB,RHOB,VDR | 426 (17) |
| PTK2B | | kinase | | 1.951 | 0.0123 | CXCL8,FOS,JUN,NEDD9 | |
| AMH | | growth factor | | 1.95 | 0.041 | ACVR1,CEACAM1,CXCL2,IRF1 | |
| cisplatin | | chemical drug | | 1.949 | 1.42E-18 | DD45A,GDF15,GPR87,HDAC9,HIPK2,HJURP,HK1,HMGCR,HSPA4L,ID2,IFIH1,JUN,JUND,KCNK5,KLF12,LAMC2,LATS2,LCN2,LGALS3,LRIG1,MAOA,MAP1LC3B,MCAM,MCL1,MEGF | 551 (17) |
| E. coli B4 lipopolysaccharide | | chemical toxicant | | 1.945 | 0.000072 | NT5,CASP7,CASP9,CLDN1,CXCL2,CXCL3,CXCL8,CYP3A5,F5,FAS,FOSB,GNA14,GNAI1,IFI35,IFIT3,IFITM3,IL1RN,LCN2,MCL1,NOX1,PIM1,PPP1R1B,PROS1,PSMB8,PTGS2,SCAR | 348 (18) |
| tert-butyl-hydroquinone | | chemical reagent | | 1.944 | 0.000169 | ABCC1,AKR1C1/AKR1C2,ATF3,CYP1A1,DYRK2,FOS,FOSL1,GADD45A,JUN,ME1,PPARG,SESN2,UGT1A10 (includes others),ZNF184 | 513 (22) |
| CAMK4 | | kinase | | 1.944 | 0.106 | ATF3,FOS,FOSB,JUN | |
| EIF2AK4 | | kinase | | 1.941 | 0.0016 | ATF3,CHAC1,EGR1,PCK2,PPARG,PPP1R15A,PSPH | |
| imipramine blue | | chemical drug | | 1.941 | 0.000928 | AURKA,CASP4,CCNA2,CCNF,CDC25A,GADD45A,STMN1 | |
| Ca2+ | | chemical - endogenous mammalian | | 1.941 | 5.54E-08 | ,CXCR4,CYP1A1,DUSP1,EDN1,EGR1,ELF3,FAAH,FGFR2,FOS,FOSL1,GRK5,HBEGF,HMGCR,IRF1,IRS2,ITPR2,JUN,JUND,KLF4,MCAM,MXD1,MYB,NR1D1,PADI1,PPARG,PTGS2,R | 428 (21) |
| TRH | | other | | 1.94 | 0.106 | DUSP1,F2R,FOS,JUN | |
| ceramide | | chemical - endogenous mammalian | | 1.94 | 0.041 | CXCL8,FOS,JUN,PTGS2 | |
| LIF | 0.672 | cytokine | | 1.938 | 0.0025 | CCNA2,CEBPD,CXCL2,CXCL8,EGR1,ELF3,EREG,FOS,GATA6,GPCPD1,HBEGF,IRF1,JUN,NMI,PIM1,PLD1,PPARG,PTGS2,RHOU,SERPINA3,SNAP23 | 515 (24) |
| TEAD4 | | transcription regulator | | 1.938 | 0.0000927 | ARID5B,BMP4,CAV2,CCN1,CDK6,CEBPA,DGAT2,EDN1,FGFR2,JUN,NFKBIZ,PPARG,RCAN1,SLFN5,TRNP1 | 248 (7) |
| doxycycline | | chemical drug | | 1.937 | 0.0385 | CEBPD,CHAC1,CXCL3,HSPA1A/HSPA1B,MMP7,MXD4,PCK2,PLK3,PTGS2 | |
| PLCE1 | | enzyme | | 1.936 | 0.00387 | CXCL2,CXCL3,CXCL8,PTGS2 | 320 (20) |
| Endothelin | | group | | 1.934 | 0.0101 | EDN1,FOS,JUN,PTGS2 | |
| NFKBIZ | 0.512 | transcription regulator | | 1.932 | 0.041 | CEBPD,CXCL3,CXCL8,LCN2 | |
| methapyrilene | | chemical drug | | 1.929 | 0.0406 | ACAA1,AMBP,ASS1,ATF3,CPT1A,CTSS,CXCL3,GADD45A,HMGCR,HMGCS2,MAOB,STMN1 | |
| green tea polyphenol | | chemical drug | | 1.914 | 0.00134 | ANXA1,CAV1,FOS,JUN,PIK3R1,PPARG,PTGS2 | 466 (22) |
| ATF2 | | transcription regulator | | 1.914 | 0.00000284 | ATF3,CCNA2,CXCL8,DUSP1,DUSP5,EGR1,FOS,GADD45A,JUN,MAT2A,MCL1,MEIS2,MT2A,PPARG,PTGS2,RHOB,TGFB2 | 470 (15) |
| crocidolite asbestos | | chemical toxicant | | 1.912 | 0.0000448 | CD44,CXCL2,FOS,FOSL1,GADD45A,JUN,JUND,LIF,ZYX | 509 (21) |
| ROCK | | group | | 1.912 | 0.0158 | CXCL8,ETS1,FOS,LPP,PPARG,RUNX2,TJP1 | |
| JAK1/2 | | group | | 1.912 | 0.0586 | EIF2AK2,FAM81A,IRF1,IRF7,ITPKA,LMO7,PLAC8 | |
| laminaran | | chemical drug | | 1.912 | 0.00347 | CLDN1,CXCL1,CXCL8,EREG,MT2A,PTGS2,SLC39A8 | |
| Fibrinogen | | complex | | 1.911 | 0.0176 | BCL3,CXCL1,CXCL3,CXCL8 | |
| IRF8 | | transcription regulator | | 1.903 | 0.0000184 | ARID5B,ATF3,B2M,BCL6,CEBPA,CTSS,CXCL16,EGR1,ETV6,FANCF,FAS,FOS,IFIT3,KLF4,KLF6,LIF,MAP4K4,PML,TRIM21,ZFHX3 | 482 (20) |
| CCL2 | | cytokine | | 1.901 | 0.00623 | CEBPA,CEBPD,CXCL3,CXCL8,FOS,LCN2,PPARG,PTGS2,RNF128,SCARB1,ZC3H12A | 440 (18) |
| HNF1B | | transcription regulator | | 1.898 | 2.36E-08 | MBP,AXIN2,BGN,CD44,CLDN1,CXCL1,DST,DUSP1,GATA6,HMGA2,HNF4A,HNF4G,ITGB6,MMP7,NR5A2,RNASE4,S100A2,S100A4,SERPINA1,SERPINH1,TM4SF4,UGT1A10 (includ | 405 (10) |
| NFYA | 0.281 | transcription regulator | | 1.897 | 0.0000196 | BBC3,CCNA2,CYP1A1,EGR1,FGFR2,GADD45A,GADD45B,IRF1,JUN,JUND,PMAIP1,PPP1R15A,RHOB,TGFBR2,TXNIP | 440 (13) |
| lactacystin | | chemical - protease inhibitor | | 1.89 | 0.000000502 | 2,ARHGDIB,ATF3,ATP2A3,BAG3,CXCL8,CXCR4,CYP1A1,CYP24A1,DUSP1,FAS,FOSL1,GNAI1,HLA-B,HLA-C,HSPB1,IRS2,ITPRID2,NEDD9,PMAIP1,PRKCA,PTGS2,RUNX2,SAT1,S | 552 (21) |
| CX3CL1 | | cytokine | | 1.89 | 0.0172 | CRABP2,ETV1,GSN,GYG1,ID2,IFIT3,IMPDH1,IRF7,LGALS3BP,MEIS2,PPARG,SOX4,TGFB2 | |
| APEX1 | | enzyme | | 1.89 | 0.000763 | ABCB10,ABCC1,EGR1,FOS,FOSB,PTGS2,RGS2 | 482 (22) |
| HGF | | growth factor | | 1.888 | 7.44E-13 | ,EDN1,EGR1,EHF,ELF3,EMP1,ERRFI1,ETS1,FOS,FOSB,FOSL1,FOXN2,GAB1,GRK5,HBEGF,HK1,IL15RA,INPPL1,INSIG1,KITLG,LAMC2,LCN2,LDHA,LGMN,LRP8,MAT2A,MCL1,MIC | 502 (23) |
| IL12 (complex) | | complex | | 1.885 | 0.00724 | BCL3,CD44,CEACAM1,CXCL8,FAAH,FAS,FOS,GADD45B,GCNT2,IFIH1,IFNGR2,IRF1,JUN,PIM1,PLAC8,PTGS2,SMAD3,TGFBR2,TXK | 377 (17) |
| DDX58 | | enzyme | | 1.875 | 0.000000145 | CASP4,CXCL8,EIF2AK2,GBP3,IFI35,IFI44,IFIH1,IFIT3,IRF1,IRF7,KLF4,LTBP3,MAT2A,NMI,PTGS2,TRIM25,ZC3HAV1 | 323 (15) |
| lysophosphatidic acid | | chemical - other | | 1.873 | 0.00000591 | BCL6,CCN1,CXCL1,CXCL2,CXCL3,CXCL8,DUSP1,EDN1,EGR1,FOS,JUN,LIF,LIPG,PPARG,PPP1R3B,PTGS2,SCARB1 | 445 (20) |
| TGFB2 | 0.811 | growth factor | | 1.873 | 0.000189 | ALDH1A1,BMP4,CYP24A1,EDN1,EGR1,FAS,FOS,HBEGF,HSPG2,LIPG,PDK4,SMAD3,TGFB2,TGFBR2,VDR,WNT11 | 541 (23) |
| INHBA | | growth factor | | 1.873 | 0.0283 | ACVR1,AKR1C3,CDC25A,CDKN2B,CXCL8,CXCR4,EREG,FSTL3,IRF1,KLF4,MCL1,PCSK5,PCSK6,PGM2L1,SOX4,ZBTB10 | |
| Notch | | group | | 1.871 | 0.00614 | ABLIM1,CDKN2B,CXXC5,DUSP1,GADD45B,ID2,MCL1,MEX3A,PATZ1,SKAP2,SMAD3,SMURF1,SORL1,TRAF4,TSPAN13 | 210 (7) |
| carbamylcholine | | chemical drug | | 1.87 | 0.0126 | AQP5,EGR1,FOS,ITPR2,JUN,RGS2 | |
| EGF | | growth factor | | 1.868 | 6.48E-19 | FOS,FOSB,FOSL1,GAB2,GADD45A,GATA2,GDF15,GRB7,GSTA4,HBEGF,HNF4A,HPGD,ID2,IDI1,IGF2,IGFBP2,IL1R2,JAG2,JUN,JUND,LCN2,LDHA,MAOA,MCL1,MUC2,NRP1,PDK4, | 616 (21) |
| IL15 | | cytokine | | 1.864 | 1.92E-08 | ,CXCR4,DPP4,DUSP5,EDN1,EGLN1,ETS1,FAS,FYN,GALM,GPRC5A,HK1,IFI35,IL15RA,IRF1,JUN,LDHA,LIF,MCL1,MSLN,NME4,PGM1,PIK3R1,PIM1,PLD1,PMAIP1,PRDX4,PSMB10, | 509 (19) |
| RARB | | ligand-dependent nuclear receptor | | 1.855 | 0.0000444 | CASP4,CASP7,CDKN2B,CXCL2,CXCL8,DHRS3,EGR1,ETS1,GADD45A,HLA-C,HOXA3,HOXA5,IGFBP6,JUN,KLF4,KRT20,MMP7,PTGS2,SERPINA1,SLC17A9,STRA6,TJP1 | 539 (25) |
| SRC | | kinase | | 1.849 | 0.00305 | ACAT1,CAV1,CXCL8,CYP1A1,DUSP1,F2R,FOS,FOSL1,JUN,MCL1,PRKCA,PTGS2,SCARB1,TJP1 | 458 (23) |
| 5-azacytidine | | chemical drug | | 1.847 | 3.38E-10 | C25A,CDKN2B,CLCF1,CLDN1,DKK1,DUSP1,EPHA2,FANCF,FOSL1,FYN,GADD45A,GDF15,HBEGF,IRF7,IRS2,KLF4,KLF6,MAFF,MCL1,MUC2,MYO1B,PMAIP1,PTGS2,RHOB,STK17 | 419 (16) |
| EIF4E | | translation regulator | | 1.846 | 0.0148 | B2M,CEBPA,CXCL16,DEDD2,EREG,FOSL1,IL15RA,INSIG1,LAT2,LGALS3BP,MCL1,NRP1,PIM1,REPS2,SVIL,TAPBP,TRIP6,VSIR | |
| calcitriol | | chemical drug | | 1.846 | 2.61E-15 | 1,EDN1,EGR1,EHF,ERRFI1,F2RL1,FAS,FIGNL1,FOS,GADD45A,GATA6,GRK5,GSDMB,HBEGF,HIPK2,HPGD,ID2,IL1RN,INCENP,IRF7,JUN,KLF4,KLK10,KLK6,LCN2,LDHA,MAOA,M | 589 (22) |
| IL2 | | cytokine | | 1.842 | 1.49E-11 | 6,DAPK1,DHODH,DPP4,DUSP2,DUSP5,EDN1,EMP1,ETS1,FAS,FGFR2,FOS,FYN,GADD45B,GDF15,HSPA1A/HSPA1B,IDI1,IFNGR2,IL1R2,IRF1,ITM2B,JUN,JUND,KLF6,LIF,LTBP3, | 483 (18) |
| NRG1 | | growth factor | | 1.833 | 0.00000404 | ,CAV1,CCN1,CDC42EP1,CXCL8,CXCR4,DUSP1,EDN1,EGR1,ELF3,EPHA2,ERRFI1,FOS,FOSB,GATA2,GRB7,HMGCR,IDI1,IGF2,JUN,JUND,MCL1,PTGS2,SGK1,SHCBP1,SOX4,STN | 475 (18) |
| zymosan | | chemical - endogenous non-mammalian | | 1.831 | 0.0743 | CXCL2,CXCL3,CXCL8,DEPP1,LGALS9,LIF,PTGS2 | |
| IL17F | | cytokine | | 1.828 | 0.00485 | CXCL1,CXCL2,CXCL3,CXCL8,CXCR4,PTGS2 | 362 (15) |
| CSF3 | | cytokine | | 1.823 | 0.00000209 | IRC3,CD44,CDK6,CEBPA,CEBPD,CXCL2,CXCL3,CXCL8,CXCR4,EDN1,EGR1,ERAP2,ETS1,FOS,GADD45A,GADD45B,IL1RN,JUN,KITLG,LCN2,MCL1,PIM1,PLSCR1,PPARG,PRKCA | 447 (17) |
| nitric oxide | | chemical - endogenous mammalian | | 1.814 | 0.000881 | ATF3,BBC3,CASP7,CD44,CXCL8,DUSP1,EDN1,EGR1,FAS,FOS,GADD45A,GSTA4,HSPG2,JUN,PMAIP1,PTGS2,SGK1,ZFHX3 | 461 (19) |
| norepinephrine | | chemical - endogenous mammalian | | 1.799 | 0.00106 | CXCL8,CXCR4,DUSP1,EDN1,EGR1,ELOVL6,FOS,GDF15,GPD1,GRK5,IRS2,MAT2A,MCAM,PPARG,PTGS2,PTPN21,SGK1,TBC1D1,TINAGL1 | 457 (22) |
| PD173074 | | chemical reagent | | 1.796 | 0.000215 | CEBPA,DEPP1,EFNA1,ELF3,GABRP,GATA2,ITGB6,KRT7,MXD1,OVOL1,PPARG,TGM2,UPK3B | |
| PRKCB | | kinase | | 1.793 | 0.0682 | EGR1,FOS,IRS2,JUN,PTGS2,TGFBR2 | |
| cytarabine | | chemical drug | | 1.793 | 0.00523 | BBC3,CD55,CDC25A,FAS,GADD45A,ITPR2,MCL1,SESN2,TGFB2 | 481 (18) |
| RAC1 | | enzyme | | 1.782 | 0.00484 | ATF3,BCL6,CDC25A,CXCL8,DUSP1,FAS,FOS,FOSL1,GSN,HSPB1,JUN,MCL1,PTGS2 | 490 (21) |
| Pkc(s) | | group | | 1.78 | 2.97E-12 | L8,CXCR4,CYP1A1,DNAJC3,DUSP1,EDN1,EGR1,ERN1,FAS,FOS,FOSL1,GADD45A,GADD45B,GRHL3,HMGCR,HPGD,ID2,IGF2,JUN,KLF6,MAOB,NR1D1,OVOL1,PHLDA1,PPARG,P | 478 (20) |
| STAT3 | | transcription regulator | | 1.778 | 2.31E-14 | ,EIF2AK2,FAS,FLRT3,FOS,FUT4,GADD45A,GBP3,HERC6,HNF4A,ID2,IFI35,IFI44,IFIH1,IFIT3,IFITM2,IFITM3,IL1RN,IRF1,IRF7,ITGB6,KLF4,LDHA,LGR4,LIF,MCL1,MMP7,MYB,NFKBI | 469 (21) |
| calcipotriene | | chemical drug | | 1.777 | 0.00108 | CXCL8,FOS,FOSL1,JUN,PRKCA,PTGS2 | 444 (23) |
| progesterone | | chemical - endogenous mammalian | | 1.771 | 1.18E-16 | FI1,F2R,F2RL1,FAAH,FGFR2,FOS,GATA2,GJB2,HBEGF,HMGCR,IGFBP2,IRS2,JUN,KLF9,KRT7,LCN2,LDHA,LGALS3,LIF,LMO7,MAOA,MDK,ME1,MMP7,MT2A,MTSS1,MYB,NDRG2, | 484 (26) |
| nitrofurantoin | | chemical drug | | 1.763 | 0.000189 | ABCC2,ACAA1,AMBP,AMD1,ASS1,CASP7,CLU,CPT1A,CTSS,CXCL3,FAS,GADD45A,HMGCR,HMGCS2,HNF4A,LGALS3,S100A10,SERPINA1,STMN1,TNFRSF12A | |
| NFKB1 | | transcription regulator | | 1.76 | 0.0000012 | ,CCN1,CD44,CEBPA,CFTR,CXCL2,CXCL3,CXCL8,DUSP1,E2F7,EGR1,EHF,ELF3,FAS,FOS,FOSB,GADD45A,GNPAT,GRK5,IFNGR2,IL1RN,IRF1,MAT2A,MCL1,MMP7,MYB,NMI,PI3, | 475 (15) |
| KMT2A | | transcription regulator | | 1.757 | 0.0167 | ABTB2,BAG3,EGR1,ETS1,HOXA5,KLF6,MEIS2,PRDM8,SCARB1,ZFAND2A | |

### Supplementary Table 4. Upstream regulator analysis based gene set enrichment analysis (GSEA) of differentially expressed genes (DEGs) after ATRA+PDT compared to PDT

| Upstream Regulator | Expr Log Ratio | Molecule Type | Predicted Activation State | Activation z-score | p-value of overlap | Target Molecules in Dataset | Mechanistic Network |
|---|---|---|---|---|---|---|---|
| 3-methylcholanthrene | | chemical toxicant | | 1.756 | 0.0515 | CCN1,CYP1A1,FOS,ME1,NEDD9,PTGS2 | |
| KITLG | -0.381 | growth factor | | 1.751 | 0.0000266 | NA2,CD68,CDKN2B,CEBPA,CLU,CXCL2,CXCL3,CXCL8,CXCR4,EGR1,F2R,FOS,GNAZ,ID2,IL1RN,KLHL24,MYB,PIM1,PLSCR1,PRKCA,PTGS2,RNF128,SNAP23,ST3GAL4,TSC22D3 | 410 (19) |
| PARP9 | | enzyme | | 1.746 | 0.0000132 | BCL6,HLA-E,IFI44,IFIT3,IRF1,IRF7,SP110 | 198 (7) |
| MAPKAPK2 | | kinase | | 1.741 | 0.0227 | CEBPD,CXCL2,CXCL3,CXCL8,HSPB1,TSC22D1,ZFP36 | |
| NFKB2 | | transcription regulator | | 1.741 | 0.000628 | BIRC3,CTSS,CXCL8,CXCR4,DMTF1,FOS,MAT2A,MCL1,MYB,PLPP3,PTGS2,TNFAIP2 | 410 (15) |
| PADI2 | -0.721 | enzyme | | 1.732 | 0.0101 | CXCL2,CXCL3,CXCL8,PTGS2 | |
| NEDD9 | 0.639 | other | | 1.732 | 0.00119 | AURKA,CA9,CD44,DDIT4,ELF3,ERRFI1,FOS,GDF15,NAAA,PLAC8,PPP1R3B,TXNIP | 398 (12) |
| 6,7-dinitroquinoxaline-2,3-dione | | chemical reagent | | 1.732 | 0.0000795 | EGR1,FOS,FOSB,JUN | 307 (11) |
| tert-butyl hydroperoxide | | chemical toxicant | | 1.732 | 0.000293 | BBC3,CASP9,CLU,CXCL8,FAS,PTGS2 | 442 (21) |
| L-glutamic acid | | chemical - endogenous mammalian | | 1.728 | 0.000154 | B2M,BAG3,CASP9,CLDN1,CYP2B6,EGR1,FOS,FOSB,GADD45B,JUN,NOX1,PTGS2,RUNX2,SORL1,TAP2,TJP1 | 395 (22) |
| folic acid | | chemical - endogenous mammalian | | 1.718 | 0.0393 | ABCC1,ANGPT1,CLU,COL18A1,DKK1,IGF2,JUN | |
| H1-2 | | other | | 1.718 | 0.00651 | AXIN2,GADD45A,HSPA1A/HSPA1B,PMAIP1 | |
| CCND1 | | transcription regulator | | 1.714 | 0.0000615 | 2B,E2F7,EIF5A2,ENDOD1,EPHA2,EREG,ERRFI1,GPRC5A,HJURP,HSPA1A/HSPA1B,HSPB1,ITGB6,KDM6B,KLHL24,MACF1,MAFF,MEIS2,NCOA3,NT5E,PPARG,RCN1,RGS2,SEC6 | 436 (12) |
| aplidine | | biologic drug | | 1.709 | 0.000000352 | DUSP1,FOS,FOSB,FOSL1,JUN,JUND | 441 (19) |
| P38 MAPK | | group | | 1.707 | 4.86E-10 | A,CEBPD,CXCL1,CXCL2,CXCL3,CXCL8,CXCR4,CYP1A1,DDC,DUSP1,EDN1,EGR1,ETS1,FAS,FBXO32,FOS,IL1RN,IRF7,JUN,KLF4,MAP4K4,MCL1,MUC2,NOX1,PMAIP1,PML,PPAR | 482 (18) |
| SCARB1 | -0.396 | transporter | | 1.706 | 0.00846 | CD68,CXCL2,DDIT4,FBXO32,HMGCR,IL1RN,PTGS2 | 408 (21) |
| 6-hydroxydopamine | | chemical toxicant | | 1.706 | 0.000595 | ACO1,BBC3,CASP9,CEACAM1,DDIT4,FOS,FOSB,HEPH,HSPA1A/HSPA1B,JUN,NOX1,PTGS2,UBC | 509 (25) |
| PRMT1 | | enzyme | | 1.703 | 0.00427 | CLDN1,CXCL3,IGFBP6,KLF4,KLHL24,MT2A,PDK4,PPARG,SPINK1 | 355 (13) |
| Muscarinic cholinergic receptor | | group | | 1.702 | 0.000178 | CCN1,FOS,JUN,RGS2 | 496 (20) |
| MMP1 | | peptidase | | 1.702 | 0.00115 | ANGPT1,CXCL8,EGR1,FGFR2,FOS,JUN,PTGS2,RUNX2,S100A4 | 469 (24) |
| SMAD4 | | transcription regulator | | 1.701 | 0.00000628 | MP4,CCNG2,CDKN2B,DAPK1,EDN1,ENG,EREG,FOS,FSTL3,GADD45A,GADD45B,GLI1,HMGA2,HNF4A,ID2,JAG2,LOC102724788/PRODH,MUC2,OVOL1,PTGS2,RUNX2,SERPINE2, | 500 (18) |
| arsenite | | chemical toxicant | | 1.694 | 0.000000856 | ABCC1,CAV1,CXCL3,CXCL8,CYP1A1,CYP2B6,DUSP1,EGR1,EREG,FOS,GADD45A,HSPA1A/HSPA1B,HSPA4L,HSPB1,IRF1,JUN,MAFF,MCL1,PTGS2,TGFB2 | 461 (20) |
| ID3 | | transcription regulator | | 1.689 | 0.000801 | BCL3,BCL6,CCNG2,CD44,CXCL1,CXCL8,CXCR4,DUSP1,ELOVL6,GADD45B,IFNGR2,JDP2,KLF6,MAP3K14,MYB,PIK3R1,RAP1GAP,REL,SEMA7A,SOX4,TRAF4 | 329 (13) |
| THPO | | cytokine | | 1.687 | 0.000147 | AURKA,BIRC3,CCNA2,CD44,CEBPA,CXCL1,FOS,GATA2,IRF1,KLF4,NFKBIZ,PIM1,REL,SH2B3,VDR | 476 (21) |
| cis-urocanic acid | | chemical drug | | 1.687 | 0.00000044 | AKR1C1/AKR1C2,CD55,CLDN1,CXCL8,GADD45B,HSPA1A/HSPA1B,PMAIP1,PPIF,PTGS2 | 351 (10) |
| MRTFA | | transcription regulator | | 1.684 | 0.000728 | AMOTL2,CCN1,CDC42EP2,CXCR4,DNAJB4,EDN1,EGR1,ETV6,F2R,FOS,GADD45A,HPGD,LCN2,PTGS2,RAB27B,SCARB1,SGK1,TBXAS1,TGFB2,TST | 400 (24) |
| EGR1 | 1.025 | transcription regulator | | 1.684 | 1.28E-11 | CLU,CXCL2,CXCL3,CXCL8,EGR1,EREG,FAS,FOSL1,FYN,GADD45A,GADD45B,GDF15,GLI1,HBEGF,HMGCR,HPGD,IGF2,JUN,JUND,MAOB,MAP1LC3B,ME1,MXD1,MYB,PNPLA2,PP | 568 (21) |
| KDM3A | | transcription regulator | | 1.677 | 0.000865 | AJUBA,ALCAM,CCN1,EDN1,ETS1,FSTL3,GDF15,MCAM | |
| ethanol | | chemical - endogenous mammalian | | 1.674 | 1.52E-08 | CR4,CYP2B6,CYP3A5,DKK1,DUSP1,EDN1,EGR1,FAAH,FAS,FOS,FOSB,GADD45A,GNAI1,HMGCR,IFIT3,IL1R2,IL1RN,IRF7,JUN,KLF6,KLHL24,LSS,MAT2A,NMI,PLK3,PNPLA2,PPA | 514 (19) |
| LDL | | complex | | 1.674 | 2.35E-08 | ,CXCL2,CXCL3,CXCL8,DUSP1,DUSP2,EGR1,FAS,FOS,FOSB,GNAI1,HMGCR,HSPA6,IGFBP2,IL1RN,INSIG1,IRF7,ITPR2,JUN,MCL1,NOX1,NPC1,NR0B2,NRP1,PMAIP1,PPARG,PT | 465 (18) |
| ELF4 | | transcription regulator | | 1.673 | 0.00863 | CXCL2,CXCL8,DUSP1,DUSP5,KLF4,RCAN1 | 260 (7) |
| IFN type 1 | | group | | 1.671 | 0.00316 | BCL6,CGAS,CXCL8,EIF2AK2,IFIH1,IRF1,PLAAT4,PML | 321 (16) |
| NR3C2 | | ligand-dependent nuclear receptor | | 1.671 | 0.000034 | ATP1B1,BMP4,CXCL8,CXCR4,DDC,DDIT4,EDN1,EGR1,KLF9,LCN2,NDRG2,PRDX4,PROCR,PTGS2,RGS2,SERPINA3,SGK1,TGM2,TSC22D3,TUBA4A | 359 (21) |
| hydrocortisone | | chemical - endogenous mammalian | | 1.668 | 0.00149 | ACO1,ALDH1L1,CEBPA,CEBPD,CIAO3,CXCL8,CXCR4,FAS,GADD45B,IGFBP6,PPARG,PTGS2,STX1A,TNFAIP2,TP53I11,TSC22D3,ZNF143 | 456 (21) |
| H1-6 | | other | | 1.667 | 0.0248 | B2M,BCL3,DEGS2,DUSP16,FOS,IGF2,MMP15,PHLDA1,ZNF143 | |
| JAG1 | | growth factor | | 1.667 | 0.00131 | CDKN2B,DUSP1,DUSP2,DUSP5,GATA2,GLI2,RNF128,RUNX2,SMAD3 | 480 (20) |
| H1f1 | | other | | 1.667 | 0.0248 | B2M,BCL3,DEGS2,DUSP16,FOS,IGF2,MMP15,PHLDA1,ZNF143 | |
| ELANE | | peptidase | | 1.666 | 0.00457 | CEBPD,CXCL2,CXCL8,F2RL1,FOS,MMP7,SERPINA1 | 392 (19) |
| F3 | | transmembrane receptor | | 1.664 | 0.000000163 | ANGPT1,CASP7,CCN1,CXCL8,DUSP2,EGR1,FOS,FUT8,HMGA2,MDK,MMP7,MSLN,PLPP3,SEMA3A,TGFBR2,TGM2 | 352 (19) |
| steroid | | chemical - endogenous mammalian | | 1.664 | 0.00221 | ANXA1,CCN1,CEBPA,CYP2B6,CYP3A5,FOS,KLK10 | 521 (23) |
| mir-183 | | microRNA | | 1.663 | 0.000185 | DDX60L,EGR1,HERC6,KLF4,NMI,SAMD9,SERPINE2,SLC30A1,SLFN5,SP110 | 202 (7) |
| STAT2 | | transcription regulator | | 1.654 | 0.00014 | CAV1,CXCL8,GLI1,IFI35,IFIT3,IFITM2,IRF1,IRF7,NOX1,PSMB8,WARS1 | 440 (22) |
| let-7 | | microRNA | | 1.637 | 0.000169 | H1B1,AURKA,CCNA2,CCNF,CD44,CDC25A,CDK6,CEBPD,CXCL8,DNAJB9,FOSL1,GAB2,HMGA2,ID2,IRS2,LRIG1,MGAT1,PDE12,PTGS2,RCAN1,S100A4,SH2B3,SIGMAR1,SSR1,ZF | 336 (19) |
| CCAR2 | | peptidase | | 1.633 | 0.00863 | CDK6,CREB5,MED15,NR1D1,PRKCA,VDAC1 | |
| JAK | | group | | 1.633 | 0.00457 | EIF2AK2,FBXO32,HES6,IFIH1,IFIT3,IFITM3,PTGS2 | 322 (12) |
| NSUN6 | | enzyme | | 1.633 | 0.0112 | AMOTL2,CCN1,SGK1,STMN1,TGFB2,TGM2 | |
| TANK | | other | | 1.633 | 0.0000566 | CXCL2,CXCL3,CXCL8,DNMT3A,NFKBIZ,PLAAT4,PTGS2,TSC22D1,ZC3H12A | 361 (16) |
| TEAD2 | 0.967 | transcription regulator | | 1.633 | 0.0117 | ARID5B,CCN1,CCNF,EDN1,NFKBIZ,SLFN5,TRNP1 | |
| inosine | | chemical - endogenous mammalian | | 1.622 | 0.013 | ALCAM,ANXA1,CD68,DPP4,IFITM3,IRF1,LGALS3BP | |
| okadaic acid | | chemical toxicant | | 1.62 | 4.96E-09 | ANXA1,BCL6,CCNG2,CRABP2,CXCL2,CXCL8,CYP24A1,CYP2B6,EGR1,FOS,FOSB,HNF4A,IL1RN,JUN,JUND,KLF4,MCL1,PIM1,PRKCA,PTGS2,RASA1 | 484 (24) |
| EGR3 | | transcription regulator | | 1.616 | 0.00244 | BCL6,CBLB,JUN,LMO7,LTBP3,MYB,PLXND1,ZNF292 | |
| CREM | | transcription regulator | | 1.604 | 0.0000798 | ATF3,CRABP2,CSRNP1,CXCL8,DUSP1,EGR1,ERRFI1,FOS,GADD45B,HMGCR,IDI1,IRS2,LSS,MCL1,MSMO1,PPP1R15A,REN,RHOB,SLC16A1,TIPARP | 527 (20) |
| triamterene | | chemical drug | | 1.604 | 0.00000302 | ANXA1,BIRC3,CD44,CLU,ELF3,LAMC2,LCN2,LGALS3,PSMB10,PSMB8,RCN1,S100A10,TNFRSF12A,TSPAN8 | |
| MAP2K1/2 | | group | | 1.6 | 0.000444 | ATF3,CXCL3,DUSP1,EGR1,FOS,FOSL1,HSPB1,JUN,MCL1,NOX1,PHLDA1,SCARB1 | 471 (23) |
| GAST | | other | | 1.599 | 0.00215 | ABCC2,DNAJA1,EGR1,FOS,HBEGF,JUN,MMP7,MUC2,PIK3R1,PPARG,PTGS2 | 434 (21) |
| arachidonic acid | | chemical - endogenous mammalian | | 1.591 | 0.000051 | CLU,DNAJB9,EDN1,EGR1,ELOVL6,FAS,FOS,FOSL1,GLI1,JUN,KLF6,PLD1,PNPLA2,PPARG,PTGS2,RGS2 | 458 (21) |
| NFATC1 | | transcription regulator | | 1.588 | 0.0352 | CYP1A1,EDN1,FAS,GLI1,HNF4A,ITPR2,PAG1,PPARG,PTGS2,RCAN1 | |
| tripterine | | chemical - endogenous non-mammalian | | 1.587 | 0.0000554 | BAG3,DNAJB1,DNAJB9,HRH1,HSPA1A/HSPA1B,HSPA1L,HSPA4L,HSPH1 | 63 (3) |
| ECSIT | | transcription regulator | | 1.584 | 0.000249 | BCL3,CD44,CXCL8,IL1RN,IRF7,PIM1,PTGS2 | 393 (16) |
| isoproterenol | | chemical drug | | 1.584 | 0.00352 | AQP5,ATF3,AURKA,CAV1,CD68,CYP1A1,DHRS3,EDN1,FOS,FOSL1,GRK5,JUN,LGALS3,LIF,PNPLA2,PTGS2,REN | 488 (22) |
| SPHK1 | | kinase | | 1.577 | 0.0264 | BIRC3,CXCL8,EGR1,ERN1,JUN,PTGS2 | |
| DKK1 | 0.576 | growth factor | | 1.572 | 0.0208 | AXIN2,BMP4,CD44,F2RL1,HNF4A,PPARG,RUNX2,WNT11 | |
| methylnitronitrosoguanidine | | chemical toxicant | | 1.569 | 0.0000065 | BBC3,BTG1,CEACAM1,CEBPA,EGR1,FAS,GADD45A,GDF15,IRF1 | 451 (18) |
| 5-hydroxytryptamine | | chemical - endogenous mammalian | | 1.566 | 0.00484 | ANGPT1,ASIC1,CASP9,CXCL8,EGR1,F2R,FOS,HBEGF,NUCB2,PKN1,PTGS2,RUNX2,TUBA4A | 489 (24) |
| R5020 | | chemical reagent | | 1.563 | 7.06E-08 | BCL6,BIRC3,CEBPA,DKK1,DUSP5,HBEGF,IFIT3,IRS2,LAMC2,LGALS3BP,MT1X,MT2A,PDK4,PPP1R1B,RARG,RGS2,SGK1,TIPARP,TXNIP | 436 (25) |
| Ige | | complex | | 1.561 | 2.44E-10 | L2,CXCL3,CXCL8,DUSP2,EDN1,EGR1,ERRFI1,F2R,FAS,FHL2,FYN,GADD45B,GDF15,GSN,HBEGF,LAMA5,LIF,NABP1,NOX1,NRP1,PIP5K1A,PLK3,PPARG,PPIF,PPP1R3B,PRKCA, | 488 (18) |
| trinitrobenzenesulfonic acid | | chemical reagent | | 1.556 | 0.00066 | ABCC2,ADAM8,CD44,CD68,CLDN1,CXCL2,CXCL3,FOS,IL1R2,LCN2,NR0B2,PTGS2,TNFRSF12A | 414 (15) |
| TIMP3 | | other | | 1.554 | 0.0000291 | BMP4,ENG,FAS,FOS,ITGB6,JUN,KLF4,LAMC2,LRP8,TGFBR2,WNT11 | 541 (24) |
| SOX4 | 0.439 | transcription regulator | | 1.552 | 0.00756 | ALDH1B1,B3GNT5,BBC3,CGAS,CHAC1,EMP1,FOS,GRB7,H1-10,HIPK2,IGF2,KANK1,LMO7,MEX3A,RARRES1,RNASE4,SERPINE2,TSPAN6 | |
| ATM | | kinase | | 1.552 | 0.000814 | BBC3,CLU,CXCL8,DUSP1,FAS,GADD45A,GADD45B,IRF1,JUN,PSME2,TGM2 | 420 (17) |
| delta-9-tetrahydrocannabinol | | chemical drug | | 1.551 | 0.042 | ALCAM,CASP7,CEBPA,CXCR4,FOS,FOSB,TAP2,TRIM38 | |
| JUNB | | transcription regulator | | 1.551 | 0.00000707 | ATF3,BCL3,CASP4,CAV1,CD44,CD68,CLU,CXCL16,CXCL2,CXCL3,DMTF1,DUSP1,E2F7,FOSL1,HPGD,ID2,LCN2,PIM1,PNPLA2,RUNX2,SGK1,SWAP70 | 491 (17) |
| NCOA2 | | transcription regulator | | 1.548 | 0.000000618 | ALDH1B1,BCAS1,CEBPA,CYP1A1,CYP2B6,EGR1,ELOVL6,HMGCR,IDI1,IRF1,IRF7,KLF4,LIPG,MYB,NR1D1,OAS3,PARD6B,PPARG,PPP1R3B,PTGS2,RARG,TSC22D3,VDR | 591 (25) |
| phorbol 12,13-dibutyrate | | chemical - endogenous non-mammalian | | 1.545 | 0.00754 | CCNG2,CD55,CXCL8,FOS,PRKCA,PTGS2 | 465 (23) |
| SPI1 | | transcription regulator | | 1.543 | 0.00000279 | 25A,CDK6,CEBPA,CTSE,CTSS,DMTF1,FOS,GATA2,ID2,IFI44,IFIT3,IFITM3,IL1R2,IL1RN,IRF7,JDP2,JUN,KLF4,MCL1,MT1E,MT1X,MT2A,MYB,PARP12,PML,PRDX4,PSMB10,PSMB8 | 354 (17) |
| TGIF1 | | transcription regulator | | 1.539 | 0.000778 | CXCL1,CXCL2,CXCL3,CXCL8,STRA6 | 396 (15) |
| PSEN1 | | peptidase | | 1.539 | 0.00218 | S,CXCL2,DUSP1,EGR1,EIF5A2,EMP1,F2R,FOS,GADD45B,HK1,HLA-E,IRF7,LRP8,PIK3R1,PMAIP1,PPARG,PTGS2,SCARB1,SRD5A3,STMN1,TGFBR2,TIPARP,TPPP,TUBA4A,TUBE | 491 (22) |
| sodium arsenite | | chemical drug | | 1.531 | 0.00000933 | ABCC1,ABCC2,CDK6,CEBPA,CXCL8,DDC,EGR1,FOS,GADD45A,GADD45B,JUN,MAT2A,PPARG | 430 (19) |
| Il3 | | cytokine | | 1.528 | 0.00917 | CD68,EGR1,FAS,FOS,LCN2,MCL1,MYB,PIM1,SNAP23 | 533 (23) |
| mitomycin C | | chemical drug | | 1.518 | 0.00000641 | ATF3,CDC25A,CXCL8,EGR1,FAS,GADD45A,HBEGF,JUN,MALAT1,MCAM,PMAIP1,PPP1R15A,SESN2 | 415 (16) |
| AVP | | other | | 1.516 | 0.029 | ATF3,EGR1,FOS,GATA2,PRKCA,RUNX2 | |
| colchicine | | chemical drug | | 1.511 | 0.00131 | ABCC2,ATF3,CYP2B6,FAS,FOS,JUN,JUND,MCL1,PTGS2 | 501 (25) |
| CXCR4 | -0.517 | G-protein coupled receptor | | 1.506 | 0.0105 | CD44,CXCL1,CXCL8,CXCR4,DPP4,EGR1,ID2,RUNX2 | |
| corticosterone | | chemical - endogenous mammalian | | 1.505 | 0.0000308 | ANXA1,ATP1B1,ATP2B1,CAV1,CPT1A,CRISPLD2,DDIT4,EDN1,FOS,JUN,KLF9,MAOA,PDK4,PNPLA2,PPARG,PTGS2,SCARB1,SGK1,TGM2,TSC22D3 | 384 (21) |
| CEBPD | 0.52 | transcription regulator | | 1.502 | 0.0113 | CEBPD,CLU,CXCL1,CXCL3,CXCL8,CXCR4,CYP1A1,FOS,NOX1,PPARG,PTGS2 | |
| dexamethasone phosphate | | chemical drug | | 1.492 | 0.00000106 | BCL6,CXCR4,DDIT4,DUSP1,ERRFI1,GPD1,PDK4,SGK1,TIPARP | 218 (7) |
| S1PR3 | | G-protein coupled receptor | | 1.491 | 0.000778 | CD44,ETS1,FOS,JUN,PTGS2 | 470 (22) |
| N-acetylmuramyl-L-alanyl-D-isoglutamine | | chemical - endogenous non-mammalian | | 1.488 | 0.0291 | AXIN2,CLDN1,CXCL2,CXCL3,CXCL8,ETS1,PTGS2 | |
| fenretinide | | chemical drug | | 1.483 | 0.000444 | ACER2,ATF3,BBC3,ELF3,FYN,JUN,PMAIP1,PPP1R15A,PRKAR2A,PTGS2,RARG,XPA | 453 (20) |
| IL22 | | cytokine | | 1.481 | 0.00121 | BCL3,CLDN1,CLDN2,CXCL1,CXCL2,CXCL3,CXCL8,HBEGF,HLA-B,HSPB1,LCN2,MCL1,MUC2,PI3,PTGS2,SERPINA3 | 432 (17) |
| thioacetamide | | chemical toxicant | | 1.478 | 3.53E-12 | KB,CLIP2,CLU,CYP1A1,ELF3,EMP1,F2R,FOS,GPD1,GYG1,HK1,IFITM2,JUN,LAMC2,LCN2,LGALS3,LGALS3BP,LRP8,PCSK6,PLPP3,PSMB10,PSMB8,PTGS2,RCN1,S100A10,SLC22 | 515 (14) |
| vinblastine | | chemical drug | | 1.474 | 0.00351 | CHST4,CYP1A1,FOS,MCL1,PMAIP1,PTGS2 | 408 (19) |
| 2,4,5,2',4',5'-hexachlorobiphenyl | | chemical toxicant | | 1.468 | 0.00307 | AKAP12,CAMKK1,CYP2B6,CYP3A5,ELAC2,ELOVL6,GADD45A,GADD45B,HMGCR,MPP6,PPP1R10,RBPMS,SMPD3 | |
| Cdc42 | | enzyme | | 1.461 | 0.0177 | CEBPA,CTSS,FAS,GATA2,LIF,STMN1 | |
| LOC105372576 | | other | | 1.461 | 0.0239 | CDK6,DOCK5,EDN1,FOS,TNFRSF12A | |
| NADPH oxidase | | complex | | 1.455 | 0.000672 | CXCL1,EDN1,FOS,JUN,NOX1,PTGS2 | 426 (21) |
| amitriptyline | | chemical drug | | 1.455 | 0.00656 | CYP2B6,CYP3A5,DEPP1,LSS,SERPINA3 | |
| BMP4 | -0.448 | growth factor | | 1.454 | 0.000638 | BMP4,CCN1,CEBPA,CXXC5,DEPP1,EFNA1,EIF4A2,ELF3,FGFR2,GABRP,GATA2,HMGA2,ID2,ITGB6,JUN,KLF4,KRT7,MXD1,OVOL1,PPARG,RUNX2,SCARB1,TGM2 | 544 (22) |
| STAT4 | | transcription regulator | | 1.448 | 0.00364 | BCL3,CAPN5,CXCL2,CXCL3,ERRFI1,GCNT2,HES6,IFIH1,IRF1,JAG2,KDM6B,KLF9,LTBP3,MUC2,PLAC8,RNF128,S100A4,SAT1,STK32C,VDAC1,VSIR | 394 (18) |
| ADRB3 | | G-protein coupled receptor | | 1.446 | 0.00367 | EGR1,FOS,JUN,PTGS2,SPHK2 | 485 (22) |
| 5-fluorouracil | | chemical drug | | 1.442 | 0.000000184 | DF15,GSN,HBEGF,HSPA4L,JAG2,JUN,LDHA,LIF,LOC102724788/PRODH,MAT2A,MCL1,MXD1,MXD4,PLK3,POLR2L,PPP1R15A,PROCR,PSME1,PSME2,PTGS2,RAP2B,RHOB,SAT1, | 496 (19) |
| TNFRSF1A | | transmembrane receptor | | 1.439 | 0.00684 | ABCC1,AQP5,BIRC3,CD44,CD68,CXCL1,CXCL2,CXCL3,CXCL8,CYP3A5,EFNA1,JUN,LATS2,LCN2,MMP7,PIM1 | 412 (15) |

Supplementary Table 4. Upstream regulator analysis based gene set enrichment analysis (GSEA) of differentially expressed genes (DEGs) after ATRA+PDT compared to PDT

| Upstream Regulator | Expr Log Ratio | Molecule Type | Predicted Activation State | Activation z-score | p-value of overlap | Target Molecules in Dataset | Mechanistic Network |
|---|---|---|---|---|---|---|---|
| SMARCB1 | | transcription regulator | | 1.435 | 0.000000946 | CSL1,ATP1B1,AURKA,BAG3,BTG1,CCNA2,CD44,CEACAM1,COL18A1,CXCR4,DAPK1,EIF2AK2,FAS,FOS,GADD45A,GJB2,GSN,HBEGF,IGFBP2,IL15RA,NEO1,OAS3,PLXNB2,PPAI | 508 (18) |
| MED12 | | transcription regulator | | 1.432 | 0.00000373 | BMP4,CREB5,EREG,GDF15,GLI1,KITLG,LIF,TGFBR2 | 250 (7) |
| diallyl disulfide | | chemical - endogenous non-mammalian | | 1.432 | 0.00167 | ATF3,CASP9,CD55,CYP2B6,GDF15,NT5E | |
| BMP6 | | growth factor | | 1.431 | 0.0000117 | AQP5,BMP4,CD44,ERRFI1,FOSL1,FRMD4B,HBEGF,ID2,KLF4,PCSK6,PTGS2,SCEL,SEMA3A,SLC39A10,SMARCA1,SORD,TNFRSF12A | 426 (16) |
| choline fenofibrate | | chemical drug | | 1.418 | 0.000319 | ACSL1,CAV1,CPT1A,NPC1,NR1D1,PCSK5,UGT1A10 (includes others) | 249 (10) |
| MVP | | other | | 1.418 | 0.000387 | CXCL2,CXCL3,CXCL8,IRF7,JUN | 532 (25) |
| CDK4/6 | | group | | 1.414 | 0.0225 | CDK6,EPHA2,ERRFI1,HSPB1,MACF1,NCOA3,RCN1,SEC61B | |
| mir-96 | | microRNA | | 1.414 | 0.000865 | DDX60L,HERC6,NMI,RPH3AL,SAMD9,SLC30A1,SLFN5,SP110 | |
| IFIH1 | 0.551 | enzyme | | 1.414 | 0.00186 | CXCL2,CXCL8,EGR1,FAS,IRF7,PHLDA1,RND3,ZC3HAV1 | 274 (17) |
| DPP-23 | | chemical reagent | | 1.414 | 0.00000151 | DNAJA1,DNAJB4,DNAJB9,ERN1,HSPA1A/HSPA1B,HSPA6,PADI1,PPP1R15A | |
| Vegf | | group | | 1.411 | 1.28E-10 | I,EGR1,EHF,EMP1,ETS1,FAS,FOSB,FOSL1,FOXN2,GAB1,GASK1B,GJB2,GRK5,HBEGF,HID1,HMGCS2,IL15RA,INSIG1,IRF1,JUN,KITLG,LCN2,LRIG1,LRP8,MAOB,MCL1,MID1,NOS | 444 (22) |
| RUVBL1 | | transcription regulator | | 1.408 | 0.024 | AXIN2,ETS1,GADD45B,METTL23,PLAC8,SESN2 | |
| indican | | chemical - endogenous mammalian | | 1.406 | 0.0268 | CA9,CXCR4,PTGS2,REN,SMAD3 | |
| metyrapone | | chemical drug | | 1.406 | 0.00183 | CXCR4,CYP1A1,CYP3A5,FOS,SCARB1 | 439 (18) |
| bromobenzene | | chemical toxicant | | 1.4 | 0.000138 | ABCC1,ABCC2,AKR7A3,CYP2B6,FOS,GADD45A,HSPB1,JUN,JUND,MSMO1,PPP1R15A,TSC22D1 | |
| POU5F1 | | transcription regulator | | 1.399 | 0.0000198 | CARD6,CASP7,CCNF,CRABP2,DAPK1,DUSP1,DUSP5,EPHA2,FAS,FUT4,GADD45A,GATA2,GATA6,GBP3,GRHL3,IER5L,KDM2B,KLF4,LDHA,MCL1,NRP1,OVOL1,PCSK6,PFKM,PM | 351 (11) |
| NR1I3 | | ligand-dependent nuclear receptor | | 1.397 | 0.0000221 | ABCC2,ALDH1A1,CYP2B6,CYP3A5,FAM107B,GADD45B,HK1,HNF4A,INSIG1,MAFF,NEDD9,RARRES1,SAT1,STEAP4,SULT2B1,TNFAIP2,TUBA4A | 407 (7) |
| benzo(a)pyrene | | chemical toxicant | | 1.395 | 0.000000148 | CASP9,CCN1,CYP1A1,ELF3,ENTPD5,EREG,FOS,FYN,GNA14,GNAI1,HSPA1A/HSPA1B,HSPA4L,HSPB1,IGFBP6,JUN,MAFF,ME1,NMI,PDLIM1,PPIF,PRKCA,PTGS2,RAP1GAP,SERI | 453 (18) |
| HIF1A | | transcription regulator | | 1.39 | 2.69E-11 | A1,CYP2S1,DDIT4,EDN1,EGLN1,ENG,EREG,ETS1,FAM13A,FAM162A,FHL2,FHL3,FOS,FYN,GADD45B,GATA6,HES6,HSPB1,ID2,IGF2,IGFBP2,IRS2,JUN,KITLG,LDHA,MAFF,MCL1, | 504 (16) |
| YAP1 | | transcription regulator | | 1.39 | 0.0000927 | ,CCNA2,CCNF,CDK6,CDKN2B,CPT1A,CXCL8,EDN1,EGR1,FBXO32,FOS,GAB1,GADD45A,GDF15,HMGCR,HMGCS2,ID2,IGF2,LATS2,LDHA,MSLN,PMAIP1,PPP1R3B,PTGS2,SGK1, | 393 (18) |
| PGF | | growth factor | | 1.389 | 0.000134 | ATF3,CASP7,CXCL8,CXCR4,EDN1,EGR1,EREG,FOSB,MAP4K4 | 483 (23) |
| gentamicin C | | chemical drug | | 1.387 | 0.0000112 | ABCC2,ANXA1,BIRC3,CD44,CLU,ELF3,LAMC2,LCN2,LGALS3,PSMB10,PSMB8,S100A10,TNFRSF12A | |
| Raf | | group | | 1.387 | 3.72E-09 | EDN1,EGR1,ETV1,FOS,FOSL1,GDF15,HMGA2,ID2,JUN,MAFF,MANSC1,MCL1,MUC2,PLPP3,PMAIP1,RND3,SH2B3,STK17A | 421 (20) |
| deoxycorticosterone acetate/potassium chloride/sodium chloride | | chemical reagent | | 1.387 | 0.00656 | CXCL2,HNF4A,PTGS2,REN,VDR | 191 (7) |
| MSTN | | growth factor | | 1.38 | 0.0208 | CXCL1,FBXO32,FOS,IGF2,MAP1LC3B,RUNX2,SERPINH1,SMAD3 | |
| Fcer1 | | complex | | 1.378 | 0.034 | CXCL3,CXCL8,FOS,FYN,IL1RN,JUN,LIF | |
| Tnf (family) | | group | | 1.375 | 0.000014 | ATF3,BBC3,BIRC3,CIRBP,CLDN2,CXCL1,CXCL3,CXCL8,CYP1A1,CYP2B6,CYP3A5,DKK1,EIF2AK2,FBXO32,JAG2,JUN,LCN2,LGALS9,NT5E,PTGS2,RUNX2 | 443 (20) |
| CRH | | cytokine | | 1.374 | 0.00699 | CCNA2,CXCL8,EDN1,FOS,HPGD,IL1RN,SCARB1,SGK1,VDR | 462 (20) |
| VCAN | | other | | 1.361 | 0.000205 | ASS1,CLU,CRISPLD2,CXCL2,CXCL3,CXCL8,FAS,GLIS3,IFI44,IL1RN,LCN2,LGALS3,LOXL4,MSLN,OAS3,PARP14,PCSK5,RGS2,RNASE4,SMAD3 | 467 (18) |
| alitretinoin | | chemical drug | | 1.35 | 0.0247 | CASP4,CASP7,CYP24A1,CYP2B6,DUSP1,FOS,HNF4A,IRF1,NR0B2,PMAIP1,PPARG,RARG,SMAD3,STRA6 | |
| RELB | | transcription regulator | | 1.35 | 0.0128 | B2M,BIRC3,CXCL8,DMTF1,MAT2A,MYB,PLPP3,PTGS2,TNFAIP2 | |
| chenodeoxycholic acid | | chemical - endogenous mammalian | | 1.348 | 0.00000515 | ABCC2,CAV1,CPT1A,CXCL8,CYP3A5,EGR1,FGF19,HNF4A,HPGD,MUC2,NR0B2,NR5A2,PCK2,PDK4,PTGS2,VDR | 537 (24) |
| TLR2 | | transmembrane receptor | | 1.344 | 0.0283 | ATF3,CEBPD,CXCL2,CXCL3,CXCL8,DUSP1,FBXO32,GADD45A,GBP3,IL15RA,IL1RN,IRF1,MUC2,PTGS2,TRIM38,VDR | |
| C3 | | peptidase | | 1.343 | 0.0475 | CASP4,CD55,CXCL16,CXCL2,CXCL3,CXCL8,JUN,LCN2 | |
| UCP1 | | transporter | | 1.343 | 0.0151 | AMD1,CD68,ERN1,FBXO32,GADD45A,GDF15,IFIT3,NUDT19,OAS3,PCK2,PSPH,RAB40B,TPPP,UBC,USH1C | |
| (+)-catechin | | chemical drug | | 1.342 | 0.00557 | CASP7,CASP9,JUN,RUNX2,SLC16A1 | 367 (7) |
| CRNDE | | other | | 1.342 | 0.0299 | DUSP5,EPHA2,GAB1,GADD45B,HSPA6 | |
| Collagen(s) | | complex | | 1.342 | 0.0218 | BCL3,CCN1,CD55,CD68,PPP1R15A,SERPINH1 | |
| Betacatenin/TCF | | complex | | 1.342 | 0.000557 | CD44,JUN,MMP7,NR5A2,RUNX2 | |
| EIF3E | | other | | 1.342 | 0.0446 | ANGPT1,CCNF,CMTM7,COL18A1,CXCL1 | |
| SOCS6 | | other | | 1.342 | 0.00295 | CEBPD,CXCL2,CXCL3,HBEGF,PTGS2 | 369 (14) |
| BAX | | transporter | | 1.342 | 0.0443 | CTSS,DNAJB9,ETV1,IFITM2,IRF7,TGM2 | |
| nonylphenol | | chemical toxicant | | 1.342 | 0.0488 | CYP3A5,EREG,FAS,HSPA1A/HSPA1B,PTGS2 | |
| NCD-38 | | chemical reagent | | 1.342 | 0.00234 | ATF3,ERN1,FOSB,MYB,PPP1R15A | 341 (11) |
| (1S,2R)-NCL-1 | | chemical reagent | | 1.342 | 0.00234 | ATF3,ERN1,FOSB,MYB,PPP1R15A | 313 (9) |
| PROC | | peptidase | | 1.337 | 0.00047 | ANGPT1,CLU,CXCL2,CXCL3,CXCL8,F5,HK1,PROCR,TJP1 | 495 (21) |
| nitroprusside | | chemical drug | | 1.333 | 0.00404 | CAV1,CXCL3,EGR1,FAS,FOS,IFNGR2,PTGS2,SLC12A4 | 434 (19) |
| TNFSF14 | | cytokine | | 1.328 | 0.0239 | ADAM8,BIRC3,CXCL1,CXCL3,F2RL1 | |
| PDX1 | | transcription regulator | | 1.318 | 0.0000828 | ARRDC4,ATF3,ATP2A3,BBC3,CD44,CKB,CXCL2,CXCR4,DUSP5,EFNA1,EGR1,HK1,HNF4A,ID2,INSIG1,IRS2,JUN,KLF6,MAOB,NR5A2,PMAIP1,TSPAN8,TXNIP | 503 (21) |
| cycloheximide | | chemical reagent | | 1.315 | 5.09E-18 | P1,EDN1,EGR1,FAS,FOS,FOSB,GADD45A,GADD45B,GNAI1,HBEGF,HMGA2,HNF4A,IGF2,IGFBP2,IGFBP6,IL1R2,IL1RN,IRF7,IRS2,JUN,JUND,KDM6B,LCN2,LDHA,MCL1,MMP7,MX | 488 (17) |
| TIRAP | | other | | 1.311 | 0.0188 | CXCL2,CXCL3,CXCL8,IRF1,PTGS2 | |
| NFAT5 | | transcription regulator | | 1.305 | 0.00141 | CXCL16,CXCL3,DDIT4,DKK1,EDN1,EREG,IFIT3,IL1RN,IRF1,LCN2,LIF,MUC2,S100A4,SGK1,TNFAIP2 | 453 (19) |
| BCR (complex) | | complex | | 1.297 | 0.0000329 | BCL6,CCNA2,CCNG2,CD44,CDC25A,CDK6,CXCR4,EGR1,FAS,FOS,FOSB,IGSF1,KLF4,KLF9,MCL1,PIM1,PTGS2,REL,S100A10 | 366 (18) |
| Ins1 | | other | | 1.289 | 0.00315 | BPD,COL18A1,CYP1A1,DGAT2,EDN1,EGR1,ELOVL6,ERRFI1,FOS,GNPAT,GPD1,HMGCR,IGF2,IGFBP2,IRS2,JUN,LDHA,LGALS3BP,PIK3R1,PPARG,PRKCA,PTGS2,REN,RUNX2,U | 448 (20) |
| histamine | | chemical - endogenous mammalian | | 1.285 | 0.000274 | CLDN1,CXCL2,CXCL8,CXCR4,FOS,GDF15,HBEGF,HRH1,JUN,MUC2,PTGS2,SEMA3A,TJP1 | 440 (19) |
| CCR5 | | G-protein coupled receptor | | 1.28 | 0.0158 | CASP9,CXCL2,CXCL3,FAS,FOS,SERPINA1 | |
| zinc | | chemical drug | | 1.276 | 0.0202 | AKR1B10,FAS,HNF4A,IDI1,IGF2,MT2A,PTGS2,SLC30A1,TGM2 | |
| IKBKE | | kinase | | 1.275 | 0.00000933 | BIRC3,CXCL2,CXCL8,GLI1,IFIH1,IFIT3,IRF7,LDHA,MAT2A,PDK4,PGM1,PSPH,PTGS2,SLC16A1,SLC16A3,ZC3HAV1 | 219 (13) |
| E. coli serotype 0127B8 lipopolysaccharide | | chemical - endogenous non-mammalian | | 1.272 | 0.00012 | LDH1L1,ANXA1,CALB2,CAV1,CXCL16,CXCL2,CXCR4,DNAJC3,F2R,F2RL1,GJB2,GNA14,GPR158,IRF1,LAMC2,LCN2,LGALS3,LIF,PCK2,PPP1R1B,PTGS2,SEMA7A,SMAD3,TGFBR | 468 (18) |
| Map3k7 | | kinase | | 1.269 | 0.00168 | BMP4,CXCL3,EDN1,HBEGF,IL15RA,JUN,NFKBIZ,PTGS2,TSC22D1,WNT11 | 441 (22) |
| vancomycin | | biologic drug | | 1.261 | 0.0000032 | NXA1,BIRC3,BMP4,CD44,CLU,CXCL2,DNAJB4,ELF3,FOS,LAMC2,LCN2,LGALS3,MYO1B,NPNT,NRIP1,PSMB10,PSMB8,RCN1,S100A10,SLCO4A1,SMAD3,SOX4,TJP1,TNFRSF12A | |
| TCF7L2 | | transcription regulator | | 1.26 | 5.56E-10 | O32,FGFR2,FOS,GAB1,GLI1,GSN,HIPK2,HMGCS2,ID2,IGFBP2,IRS2,JUN,LAMC2,LGALS4,MAP4K4,MCAM,ME1,MMP7,MUC2,MYB,NPC1,OTUD7B,PAPSS1,PPARG,PTGS2,RCBTB | 485 (25) |
| hexachlorobenzene | | chemical toxicant | | 1.257 | 0.00000287 | ANXA1,CD44,CLU,ELF3,LAMC2,LCN2,LGALS3,PSMB10,PSMB8,RCN1,S100A10,TNFRSF12A,TSPAN8 | |
| homocysteine | | chemical - endogenous mammalian | | 1.251 | 0.00344 | ATF3,CASP9,CLU,CXCL8,FYN,GADD45A,HMGCR,IDI1,PHLDA1,PMAIP1,PTGS2 | 497 (22) |
| Tgf beta | | group | | 1.248 | 0.0000827 | KN2B,CLU,CXCL2,CXCL3,CXCL8,CXCR4,EDN1,ENG,FOS,FSTL3,GADD45B,GLI2,HSPG2,IL1RN,JUN,KDM6B,LIF,MMP15,MMP7,MXD1,NEDD9,PLPP3,PTGS2,RHOB,RUNX2,SEC14I | 524 (22) |
| prostaglandin E2 | | chemical - endogenous mammalian | | 1.243 | 6.33E-11 | D,CXCL1,CXCL3,CXCL8,CXCR4,DUSP1,EGR1,EREG,ETS1,F2R,F2RL1,FAS,FOS,FOSB,FOSL1,FSTL3,IL15RA,IL1R2,IL1RN,IRF1,JAG2,JUN,KLF4,MCL1,MUC2,NEDD9,NR5A2,NRP | 489 (19) |
| aspirin | | chemical drug | | 1.242 | 0.00000188 | AKAP12,CAV1,CDKN2B,CXCL1,CXCL8,CXCR4,DPP4,EFNA1,EMP1,ENG,F2RL1,GDF15,GLI1,HBEGF,HPGD,IGFBP2,KITLG,MCAM,MCL1,NPC1,PMAIP1,PTGS2,SCARB1,SOX4,TJP | 530 (23) |
| POR | | enzyme | | 1.237 | 0.00000284 | LDH1A1,CPT1A,CYP1A1,CYP2B6,ELOVL6,ENTPD5,EREG,GADD45B,HMGCR,HSPH1,IDI1,INSIG1,JDP2,LSS,MSMO1,NR0B2,PDK4,PPARG,RAB30,SC5D,SDC4,SERPINA3,SLC22 | |
| fluticasone propionate | | chemical drug | | 1.237 | 3.38E-10 | L2,CXCL8,DDIT4,EMP1,GADD45B,GJB2,HPGD,IL1R2,IL1RN,IRS2,KLF9,MMP7,MT1X,MT2A,PDK4,PHLDA1,PLAAT4,PLAC8,RARRES1,RCAN1,SDC4,SERPINA1,TCN2,TFPI,TGM2,T | 322 (13) |
| EREG | 0.557 | growth factor | | 1.235 | 0.0101 | CXCL3,EREG,HBEGF,PTGS2 | |
| deoxynivalenol | | chemical toxicant | | 1.231 | 0.00246 | CLDN1,CLDN2,CXCL8,DNAJC3,ERN1,TJP1 | |
| forskolin | | chemical toxicant | | 1.23 | 2.71E-17 | EG,ERRFI1,FAS,FGFR2,FOS,FOSB,FOSL1,FRRS1,GAB2,GADD45A,GADD45B,GLI1,GNA14,GRK5,HOXA5,HPGD,HSPA1A/HSPA1B,HTR1D,ID2,IGF2,IGFBP6,IRS2,ITPR2,JUN,JUN | 508 (20) |
| Mek | | group | | 1.229 | 1.68E-11 | V1,FOS,FOSL1,GCNT2,GDF15,HMGA2,HRH1,HSPB1,ID2,IRF1,ITGB6,JUN,LAMC2,LRP8,MAFF,MANSC1,MCL1,MSLN,MUC2,NOX1,NPC1,NT5E,PHLDA1,PLPP3,PMAIP1,PTGS2,RN | 496 (23) |
| GC-GCR dimer | | complex | | 1.227 | 0.000503 | ANXA1,DUSP1,IL1R2,IL1RN,PDK4,PIK3R1,TSC22D3 | |
| RUNX1 | | transcription regulator | | 1.225 | 0.00746 | ALCAM,BBC3,CDKN2B,CEBPA,ENG,FAS,GATA2,HMGA2,ID2,IFITM3,IRF7,KRT80,MYB,OAS3,PIK3R1,PLXND1,PMAIP1,PPP1R15A,RUNX2,SEC61A1,TJP1,UGCG | 490 (22) |
| CCL5 | | cytokine | | 1.224 | 0.0000231 | ALCAM,CD44,CXCL2,CXCL3,CXCL8,DUSP1,EMP1,F2R,F2RL1,FOS,PPIF,SGK1,SQLE,ZFP36 | 494 (21) |
| CASR | | G-protein coupled receptor | | 1.222 | 0.0000405 | ABCC2,AGR2,CASP9,CD44,CYP2B6,EDN1,EGR1,GADD45A,GATA6,INCENP,NEDD9,PPARG,PTGS2,SERPINA1,SLC7A7,TGFB2,TXNIP,VDR,ZYX | 476 (25) |
| bee venom | | chemical - endogenous non-mammalian | | 1.222 | 0.00000513 | CXCL8,DNAJB1,FEM1C,FOSB,GADD45A,GPR35,HSPA4L,HSPB1,IRF1,JUN,KLF4,PPP1R15A,PTGS2,RCAN1,RGS2,TNFAIP2,TSC22D1 | 385 (14) |
| erlotinib | | chemical drug | | 1.221 | 0.0205 | BBC3,CXCL1,CXCL3,CXCL8,EGR1,FAS,IGFBP6,LIF,NOX1,PML | |
| MAVS | | other | | 1.219 | 0.00699 | CGAS,CXCL2,CXCL8,IFIT3,IFITM3,IRF7,MAT2A,PARP12,ZC3HAV1 | 219 (16) |
| GLI3 | | transcription regulator | | 1.214 | 0.0000646 | BMP4,DUSP1,DUSP2,DUSP5,EGR1,FGF19,FOSB,GLI1,GLI2,IRF1,JAG2,NFKBIZ,RUNX2,SMAD3,STMN1 | |
| hydrochloric acid | | chemical - endogenous mammalian | | 1.213 | 0.0107 | CXCL2,CXCL3,CXCL8,FOS,JUN | |
| RHOA | | enzyme | | 1.213 | 0.0342 | CDKN2B,CXCL8,FOS,GDF15,JUN,RHOB,RND3,RUNX2,SCARB1 | |
| clopidogrel | | chemical drug | | 1.212 | 0.000138 | ATF3,CHAC1,CHD2,DHRS3,DUSP2,HOXA3,HOXA5,LAMA5,PCK2,PMAIP1,PPP1R15A,SESN2 | |
| MAC | | complex | | 1.211 | 0.000249 | AKR1C3,CXCL2,EGR1,IRF1,ITPRIP,PTGS2,ZFP36 | 409 (17) |
| Ccl2 | | cytokine | | 1.208 | 0.0478 | CXCL2,CXCL3,ETS1,KLF4,PPARG,SDC4 | |
| HSF1 | | transcription regulator | | 1.203 | 3.91E-09 | LU,COL18A1,DEDD2,DNAJA1,DNAJA4,DNAJB1,EIF4A2,HSPA1A/HSPA1B,HSPA4L,HSPB1,HSPH1,JUN,LDHA,LIF,LRIF1,MAT2A,MBOAT7,MCL1,MRPL18,PLAAT4,PPARG,RHBDD | 435 (12) |
| ESRRG | | ligand-dependent nuclear receptor | | 1.2 | 0.00917 | ACO1,ATP5MC3,FBXO32,KCNE3,KLK6,LDHA,NR0B2,NRP1,PDK4 | 182 (7) |
| streptozocin | | chemical drug | | 1.2 | 0.00239 | ATF3,ATP1B1,CD68,CEBPA,CLU,CPT1A,DNMT3A,ELOVL6,FAS,FBXO32,FOS,HNF4A,HSPG2,IRS2,LGALS3,MAP1LC3B,NT5E,PDK4,PPARG,PTGS2,RUNX2,TXNIP,UBC,VDAC1 | 529 (24) |
| TRPS1 | 0.417 | transcription regulator | | 1.195 | 0.0117 | ACSL1,CCNA2,DKK1,RUNX2,SMPD3,TGFB2,VDR | |
| C5AR1 | | G-protein coupled receptor | | 1.195 | 0.0315 | CD55,CXCL2,CXCL3,CXCL8,ERN1,TGFB2,TGFBR2 | |
| IFNA1/IFNA13 | | cytokine | | 1.193 | 0.00193 | CYP1A1,CYP3A5,EIF2AK2,IFIH1,IRF1,IRF7,PLAAT4,PLSCR1,PTGS2 | 361 (18) |
| Pdgf Ab | | complex | | 1.192 | 0.0101 | ANGPT1,EREG,FOS,HBEGF | |
| hemoglobin | | complex | | 1.188 | 0.0101 | CXCL8,EDN1,ME1,REN | |
| bryostatin 1 | | chemical drug | | 1.179 | 0.00565 | BIRC3,CD44,CXCL8,CXCR4,IFNGR2,PRKCA | 484 (21) |
| DETA-NONOate | | chemical reagent | | 1.178 | 0.000000219 | ACO1,BGN,BIRC3,CA9,CAV1,CXCL3,CXCL8,DUSP1,FAS,FOS,HMGCR,IRF7,PTGS2,TXNIP | 446 (20) |
| dinoprost | | chemical - endogenous mammalian | | 1.17 | 0.00426 | AKR1C1/AKR1C2,EDN1,FOS,HMGCS2,HSPA1A/HSPA1B,ID2,JUN,NOX1,PTGS2,RUNX2 | 431 (22) |
| TBK1 | | kinase | | 1.166 | 0.0000443 | APOBEC3B,CXCL2,CXCL3,CXCL8,EDN1,IFI44,IFIT3,IL15RA,IRF7,NFKBIZ,NMI,OAS3,PTGS2,TSC22D1,ZC3H12A | 278 (18) |
| KDM5B | | transcription regulator | | 1.163 | 0.00000293 | AURKA,CAV1,CLCF1,CLDN1,EGR1,EPB41L1,FGFR2,GADD45A,GJB2,IGFBP2,INSIG1,LGALS3BP,MCAM,MIA3,MT1E,MT1X,NEDD9,SAT1,SWAP70,TGFB2,TUBB2A | |
| VDR | -0.538 | transcription regulator | | 1.159 | 0.0000335 | ,CD44,CXCL8,CYP24A1,CYP2B6,EDN1,EGR1,GADD45A,GATA6,GLI1,GPD1,IL1RN,INCENP,JUN,NEDD9,NR0B2,PLD1,PNPLA2,PPARG,PTGS2,REN,SERPINA1,SLC7A7,TGFB2,TX | 411 (22) |
| SLC13A1 | | transporter | | 1.155 | 0.00176 | AKR1C3,ALDH1A1,ANKS4B,ELOVL6,HSPB1,IGFBP2,INSIG1,KLF9,MPP6,SDF2L1,SLC16A1,ZNF292 | |
| FASLG | | cytokine | | 1.154 | 0.00399 | CD44,CXCL8,EGR1,FAS,FOS,RUNX2,TGFB2 | 439 (18) |

Supplementary Table 4. Upstream regulator analysis based gene set enrichment analysis (GSEA) of differentially expressed genes (DEGs) after ATRA+PDT compared to PDT

| Upstream Regulator | Expr Log Ratio | Molecule Type | Predicted Activation State | Activation z-score | p-value of overlap | Target Molecules in Dataset | Mechanistic Network |
|---|---|---|---|---|---|---|---|
| RHOB | 0.335 | enzyme | | 1.154 | 0.00651 | CXCL8,PTGS2,RHOB,TGFBR2 | 483 (21) |
| proteasome inhibitor PSI | | chemical - protease inhibitor | | 1.151 | 0.0000933 | CD55,CXCL2,CYP1A1,FAS,PRKCA,RUNX2,SMAD3,TJP1 | 496 (17) |
| enalapril | | biologic drug | | 1.15 | 0.0174 | ABCC2,CLU,LCN2,LGALS3,REN,TGFBR2,TNFRSF12A | |
| tamoxifen | | chemical drug | | 1.144 | 8.51E-10 | 6,CEBPD,CLU,CXCL8,CXCR4,CYP2B6,CYP3A5,DEPP1,DNAJA1,F2R,FHL2,FLOT1,FOS,GDF15,GJB2,HMGCR,IGF2,IL1RN,INPPL1,IRF1,JUN,KLHL24,KRT80,LIF,LSS,MAFF,NR0B2,I | 548 (21) |
| 8-bromo-cAMP | | chemical reagent | | 1.135 | 1.65E-11 | P2B6,DKK1,DUSP1,EGR1,ETS1,F2R,FAS,FLNB,FOS,FOSL1,FUT8,FYN,GATA6,GLI1,GNAI1,GSN,HPGD,ID2,IGF2,IQGAP2,ITPR2,JUN,KITLG,LGMN,LIF,MAOA,MAOB,MAT2A,MMP7 | 562 (21) |
| LLGL2 | | other | | 1.134 | 0.000503 | AMOTL2,CCN1,DUSP1,SGK1,SHCBP1,STMN1,TGFB2 | |
| vincristine | | chemical drug | | 1.134 | 0.00186 | EGR1,FAS,JUN,MCL1,PMAIP1,PRKCA,PTGS2,TUBB2A | 435 (20) |
| TP53COR1 | | other | | 1.131 | 0.0148 | BBC3,BIRC3,FAS,PMAIP1 | |
| PF4 | | cytokine | | 1.129 | 0.00388 | BIRC3,CEBPA,CXCL3,CXCL8,GATA2,IRF1,KLF4,NFKBIZ,REL,VDR | 458 (16) |
| fenamic acid | | chemical reagent | | 1.127 | 0.000000965 | ABCC2,ANXA1,BIRC3,CD44,CLU,ELF3,LAMC2,LCN2,LGALS3,PSMB10,PSMB8,RCN1,S100A10,TNFRSF12A,TSPAN8 | |
| IL5 | | cytokine | | 1.125 | 1.13E-13 | ,EGR1,FAM162A,FAS,GADD45A,HBEGF,HMGCR,HSPA6,HSPH1,IDI1,IL1R2,IRF7,ITGB6,LMO7,MAT2A,MYADM,NABP1,PDLIM1,PGM1,PIM1,PKIB,PPIF,PRDX4,QSOX1,RAP1GAP,R | 516 (18) |
| mir-133 | | microRNA | | 1.124 | 0.0208 | AK2,CASP9,CXCL3,ITPR2,MALAT1,PPARG,TBL1X,TRPS1 | |
| mir-142 | | microRNA | | 1.119 | 0.0461 | CXCL3,HMGA2,MYB,TGFBR2 | |
| ATF1 | | transcription regulator | | 1.119 | 0.000163 | EGR1,FOS,FOSL1,HLA-B,JUN,NOX1,PTGS2,REN,TGFBR2 | 269 (12) |
| PELP1 | | other | | 1.117 | 0.000117 | BCAS3,CXCL1,CXCL8,CXXC5,FOS,FYN,GJB2,IFI44,MYADM,P3H2,PI3,PPARG,PTGS2,SNCG,SYT7,TCN2,TGM2 | 449 (18) |
| Pkg | | group | | 1.109 | 0.0045 | EDN1,EGR1,FOS,FOSB,JUN | 349 (21) |
| YWHAZ | | enzyme | | 1.109 | 0.000111 | CXCL2,FOSL1,MCL1,MMP7,PPARG,SMAD3,TSC22D3 | 335 (12) |
| cholic acid | | chemical - endogenous mammalian | | 1.108 | 0.0118 | ABCC2,CIRBP,CYP2B6,CYP3A5,HMGCR,MDK,MYB,NR0B2,SQLE | |
| Ngf | | group | | 1.107 | 0.00168 | BBC3,CLU,DUSP1,EGR1,FOS,JUN,MYB,PRKCA,SLC12A4,TGFB2 | 466 (23) |
| JUND | 0.515 | transcription regulator | | 1.104 | 0.00613 | BCL3,CLU,CXCL16,FOSL1,HPGD,JUND,MMP7,RUNX2,SAT1,SOX4 | 434 (17) |
| WNT5A | | cytokine | | 1.103 | 0.0115 | ARHGDIB,AXIN2,CXCL1,CXCL2,CXCL8,CXCR4,EREG,ID2,IRF1,MMP7,NR0B2,PTGS2,RUNX2 | |
| arotinoid acid | | chemical toxicant | | 1.103 | 0.0363 | DUSP1,FOS,RARG,RARRES1 | |
| pregnenolone carbonitrile | | chemical drug | | 1.094 | 0.000514 | ABCC2,CPT1A,CYP24A1,CYP2B6,CYP3A5,HMGCS2,INSIG1,UGT1A10 (includes others) | 261 (5) |
| captopril | | chemical drug | | 1.093 | 3.73E-08 | ABCC2,ANXA1,BGN,CD44,CLU,IGFBP6,IL1RN,LAMC2,LCN2,LGALS3,MAOA,PSMB8,PTGS2,REN,S100A10,TGFB2,TNFRSF12A,TSPAN8 | 445 (21) |
| betamethasone | | chemical drug | | 1.091 | 0.0241 | CAV1,CRISPLD2,CYP3A5,TGM2 | |
| TGFB3 | | growth factor | | 1.086 | 0.0148 | CCN1,CDKN2B,ENG,ETS1,F2RL1,FOS,FOSB,JUN,JUND,SERPINE2,TJP1,TSPAN13 | |
| CAMP | | other | | 1.085 | 0.00387 | CXCL1,CXCL3,CXCL8,CXCR4,HBEGF,IL1R2,IL1RN,MCL1,MUC2,PTGS2,TNFAIP2,XPA | 428 (19) |
| bleomycin | | biologic drug | | 1.08 | 0.0000347 | ANGPT1,BMP4,CD68,CEBPA,CTSE,CTSS,CXCL1,CXCL2,CXCR4,E2F7,F2R,FAS,FOSL1,GAB1,GAB2,HBEGF,KITLG,LGALS3,MID1,S100A4,SDC4,TGFB2,TGFBR2,TINAGL1 | 531 (20) |
| NGF | | growth factor | | 1.078 | 0.00517 | ACER2,AKR1B10,ASIC1,ATF3,AXIN2,BBC3,CAV1,DUSP1,EGR1,ERRFI1,FAS,FOS,FOSL1,FRRS1,GADD45A,IGF2,JUN,MACROD1,PPARG,REN,TNFRSF12A | 484 (23) |
| PDLIM2 | | other | | 1.069 | 0.00019 | CEMIP,CHPF,CRISPLD2,CXCL8,F2RL1,HBEGF,IFI44,IFIT3,LGALS3BP,MYEOV,OAS3,PHLDA1,RNF128,TXNIP | 341 (14) |
| GATA1 | | transcription regulator | | 1.069 | 0.0328 | ABCA2,ABCB10,CDK6,CEBPA,DMTN,EGR1,ETS1,FOSB,GATA2,HIPK2,JDP2,MYB,NCOA7,PIM1,PRDM8,REL,RUNX2,SLC24A1,TFF3,TGM2,TRIM29,VDR | |
| lomustine | | chemical drug | | 1.069 | 0.00000249 | ABCC2,ANXA1,BIRC3,CD44,CLU,ELF3,LAMC2,LCN2,LGALS3,PSMB8,RCN1,S100A10,TNFRSF12A,TSPAN8 | |
| phytohemagglutinin | | chemical drug | | 1.068 | 0.0000447 | :D44,CGAS,COL18A1,CXCL8,CXCR4,ENG,FAS,FOS,GAB2,GATA2,GNAI1,GPD1,HPGD,IFNGR2,IL15RA,IL1R2,KRT13,MAOA,MEIS2,MT1E,NRIP1,NT5E,PLD1,PRKCA,RASA1,SERPI | 533 (24) |
| CCAR1 | | transcription regulator | | 1.067 | 0.0000795 | CEBPA,CEBPD,PPARG,RGS2 | 217 (7) |
| EHHADH | | enzyme | | 1.067 | 0.00507 | ACAA1,ACSL1,PDK4,PEX11A | 170 (7) |
| KLB | | enzyme | | 1.067 | 0.0000574 | EGR1,FGF19,FOS,HMGCR,NR0B2 | 143 (6) |
| buthionine sulfoximine | | chemical drug | | 1.067 | 0.00387 | CXCR4,FOS,HBEGF,PTGS2 | 468 (21) |
| FGF1 | | growth factor | | 1.066 | 0.00353 | AKR1B10,AXIN2,CCN1,CD68,CXCL1,EDN1,EGR1,FOS,JUN,PTGS2,RHOU,TGFB2,TGM2,TNFRSF12A | 472 (23) |
| PRKAA1 | | kinase | | 1.06 | 4.06E-09 | ,BIRC3,BTG1,CAMKK1,CLU,CXCL8,CYP2B6,CYP3A5,ERN1,FGFR2,GALM,GJB2,ID2,IFI44,LDHA,NDRG2,PPP1R3B,PSMB8,RGS2,RHOU,RND3,SLC16A1,SMAD3,SORL1,TGM2,TJP | 513 (23) |
| ZBED6 | | transcription regulator | | 1.06 | 0.0239 | DDIT4,IGF2,KITLG,MAP3K14,SGK1 | |
| IL24 | | cytokine | | 1.059 | 0.000552 | BCL6,CXCL8,EIF2AK2,FAS,GADD45A,HBEGF,MCL1,PPP1R15A,PTGS2 | 503 (18) |
| HOXA10 | | transcription regulator | | 1.058 | 0.00872 | ALDH1A1,ARL4A,CDKN2B,DKK1,DPP4,HMGCR,HOXA5,ID2,KLF9,LCN2,ME1,MEIS2,NDRG2,PIK3R1,PROS1,RNASE4,SAT1 | |
| HMGA1 | | transcription regulator | | 1.055 | 0.0000041 | ARL4A,B2M,CAV1,CAV2,CCN1,CD44,EGR1,FOS,GSN,HMGCR,IDI1,INSIG1,KITLG,MGAT1,PPARG,PTGS2,RHOB,SDC4,SERPINH1,SOX4,XPA,ZYX | 519 (22) |
| SMPD1 | | enzyme | | 1.054 | 0.013 | CLCF1,CXCL1,CXCL2,CXCL8,CXCR4,MAP1LC3B,UGCG | |
| GNAS | | enzyme | | 1.053 | 0.0227 | CEBPA,FOS,GLI1,GLI2,MUC2,NRP1,PPARG | |
| Creb | | group | | 1.052 | 0.00000315 | 3PD,CLMN,CPT1A,CXCL8,DUSP1,EGR1,FAS,FLNB,FOS,FOSB,FOSL1,HPGD,IRS2,JUN,KDM2B,LTBP3,MALAT1,MAT2A,MCAM,MCL1,MYB,NEDD9,PPARG,PRKCA,PTGS2,RGS2,S | 522 (22) |
| TP73 | | transcription regulator | | 1.048 | 3.27E-14 | /ITF1,ECM1,EDN1,EGR1,ENG,EPHA2,EREG,FAS,FGFR2,GAB2,GADD45A,GATA2,GRK5,HBEGF,HSPA1A/HSPA1B,ID2,JAG2,LDHA,LOC102724788/PRODH,MDK,NRP1,PMAIP1,PM | 470 (19) |
| APC | | enzyme | | 1.048 | 0.00000559 | AXIN2,B2M,CD44,CD55,CLDN1,CLU,CXCL2,CXCL3,DPP4,EDN1,GLI1,GLI2,ID2,KLF4,MMP7,MUC2,MXD1,PDK4,PRKCA,PTGS2,SGK1,TFF3,TJP1,TNFRSF12A | 489 (22) |
| quercetin | | chemical drug | | 1.047 | 0.00076 | ABCC2,ARRDC4,CASP9,CAV1,CXCL8,CYP1A1,EDN1,EGLN1,EGR1,FOS,HMGCS2,JUN,LCN2,MCL1,NEDD9,PLD1,PTGS2 | 437 (23) |
| ERK1/2 | | group | | 1.046 | 0.000000156 | )44,CDC25A,CEBPA,CXCL1,CXCL3,CXCL8,CXCR4,DAPK1,DKK1,DUSP1,EDN1,EGR1,ETS1,FGFR2,FOS,FOSB,FOSL1,GRK5,HBEGF,HPGD,IRS2,JUN,JUND,NOX1,PSMB10,PSMB8 | 495 (20) |
| fluoxetine | | chemical drug | | 1.045 | 0.0079 | DEPP1,EGR1,FOS,FOSB,GNAI1,GNAZ,LSS,PPP1R1B,S100A10,SERPINA3 | 330 (20) |
| CEBPB | | transcription regulator | | 1.041 | 0.00000464 | CXCL3,CXCL8,CXCR4,CYP1A1,CYP24A1,DAPK1,DGAT2,FAS,FBXO32,FGFR2,FHL2,FOS,GADD45A,IFIT3,IFITM3,IFNGR2,IL1RN,IRF7,IRS2,JUN,LCN2,LGMN,NFKBIZ,NRP1,PDK4,I | 540 (24) |
| methamphetamine | | chemical drug | | 1.033 | 0.0433 | ABCC1,EGR1,FAS,FOS,FOSB,JUN,MAOA,SIGMAR1,UGT8 | |
| PLA2G6 | | enzyme | | 1.026 | 0.0045 | NOX1,PPARG,PTGS2,RGS2,RUNX2 | 518 (25) |
| MTOR | | kinase | | 1.023 | 0.0000454 | .1,CXCL2,CXCL8,CXCR4,DDIT4,DUSP1,EFNA1,FBXO32,HK1,HMGCS2,HSPB1,IGF2,IRF1,IRS2,LDHA,MAP1LC3B,MCL1,PDK4,PHLPP1,PIM1,PLD2,PPARG,PRKAR2A,PYGB,RUNX2 | 489 (24) |
| phenacetin | | chemical drug | | 1.022 | 5.93E-08 | ABCC2,ANXA1,BIRC3,CD44,CLU,ELF3,LAMC2,LCN2,LGALS3,PSMB8,S100A10,TNFRSF12A,TSPAN8 | 290 (7) |
| LEPR | | transmembrane receptor | | 1.022 | 0.0000303 | ADAM8,BTG1,CD68,EDN1,FOS,GADD45B,HMGCR,ID2,IL1R2,IL1RN,INPPL1,IRS2,LCN2,LGMN,LIF,MAP4K4,MMP7,PHLPP1,PPARG,PSMB8,SCARB1,TGFBR2,TSC22D3 | 566 (20) |
| NDRG1 | | kinase | | 1.002 | 1.23E-08 | ATF3,CASP4,CAV1,CEACAM5,CXCL1,CXCL2,CXCL3,CXCL8,ERRFI1,FOS,JUN,MAP1LC3B,PML,SMAD3 | 468 (22) |
| 3-deoxy-2-octulosonic acid(2)-lipid A | | chemical - endogenous non-mammalian | | 1 | 0.00818 | HMGCR,LSS,PTGS2,SQLE | 396 (7) |
| kawain | | chemical - endogenous non-mammalian | | 1 | 0.0148 | CEBPA,CYP3A5,FAS,PPARG | |
| PKNOX2 | | transcription regulator | | 1 | 0.0101 | BBC3,EGR1,FAS,GADD45A | |
| SUZ12 | | enzyme | | 1 | 0.0163 | CALB2,CDC25A,FBXO32,GATA6,HNF4A,INSIG1,JUND,KLF4,LAMC2,LGALS3,MMP7,SERTAD2,TNFAIP2 | |
| nilvadipine | | chemical drug | | 1 | 0.041 | AMBP,CASP9,GABRP,MAP4K4 | |
| ACVR1C | | kinase | | 1 | 0.0207 | CCNG2,CDKN2B,FOS,KLF4 | |
| ARRB1 | | transcription regulator | | 1 | 0.00565 | CXCL8,EGR1,IRS2,PTGS2,SCARB1,TSC22D3 | 460 (26) |
| DDX17 | | enzyme | | 1 | 0.00818 | FOSL1,JUN,LCN2,S100A4 | |
| TFPI2 | | other | | 1 | 0.0241 | ATF3,CXCL2,GADD45B,HTR1D | |
| acetyl-L-carnitine | | chemical - endogenous mammalian | | 1 | 0.0123 | BBC3,CPT1A,GADD45A,PPARG | |
| 7-ketocholesterol | | chemical - endogenous mammalian | | 1 | 0.00234 | CXCL8,CYP1A1,EGR1,GNAI1,SCARB1 | 341 (12) |
| NOS2 | | enzyme | | 0.997 | 0.000377 | ACO1,AZGP1,CD44,CXCL2,CXCL3,CXCL8,EDN1,FAS,GPD1,IFIT3,IL1RN,IRS2,KLF6,KRT13,LCN2,LGALS3BP,PLSCR1,PTGS2,SERPINA3,SERPINH1,TNK2 | 461 (18) |
| LIPE | | enzyme | | 0.992 | 0.000000169 | ACSL1,ACSS2,ALDH1A1,ANGPT1,ATF3,BCL6,CEBPA,CLK1,EDN1,EGR1,FOS,HSPA1A/HSPA1B,ID2,KITLG,NRIP1,PDK4,PPARG,PTGS2,RUNX2,SCARB1,SDC4,SUCLA2 | 477 (22) |
| ATP | | chemical - endogenous mammalian | | 0.981 | 0.0000155 | CLDN1,CXCL2,CXCL3,CXCL8,CXCR4,DUSP1,EIF2AK2,FOS,HBEGF,HK1,IL1RN,LCN2,NRP1,PLD2,PTGS2,REN,SAT1,TXNIP | 414 (23) |
| FGF2 | | growth factor | | 0.981 | 0.00000371 | C25A,CXCL2,CXCR4,DKK1,EGR1,EREG,ERRFI1,ETS1,ETV1,FAS,FGFR2,FOS,FOSL1,FRRS1,GADD45A,GLI1,HBEGF,ID2,IGF2,IGFBP2,JUN,KDM2B,MACROD1,MCAM,PPARG,PT( | 504 (24) |
| vorinostat | | chemical drug | | 0.975 | 0.0000593 | RKA,BBC3,BIRC3,CARD6,CEBPA,CLDN2,CLU,CXCL8,DUSP1,EDN1,EGR1,FAS,HMGCR,HPGD,IDI1,IRF1,JUN,KLF4,LIPG,LSS,MCL1,MMP15,MYB,PMAIP1,PTGS2,RGS2,RUNX2,SEI | 436 (20) |
| geldanamycin | | chemical drug | | 0.975 | 0.00781 | XA1,AURKA,BGN,CDKN2AIP,CEBPD,CHORDC1,CLU,CXCL8,CYP1A1,DNAJB1,EIF2AK2,FYN,HSPA1A/HSPA1B,IRF1,IRS2,JUN,LIF,NUDC,PTGS2,QSOX1,SMARCA1,TPPP,TRIM3,Z | 424 (19) |
| NCOA3 | 0.431 | transcription regulator | | 0.973 | 0.00000129 | CCN1,CCNA2,CDC25A,CEBPA,CYP1A1,CYP2B6,FGFR2,GJB2,IRF1,LDHA,MUC2,NCOA3,PARD6B,PPARG,PTGS2,SLC16A3,TGFB2 | 551 (23) |
| RET | | kinase | | 0.97 | 0.000345 | ANXA1,BBC3,CLU,CXCL8,CXCR4,DNAJA1,DNAJC3,EGR1,ETV1,FOS,HSPA1A/HSPA1B,HSPA1L,HSPH1,MMP7,PMAIP1,RNF19A,TGFB2,WNT11 | 516 (22) |
| NRAS | | enzyme | | 0.968 | 0.00000124 | 38,CDKN2B,CEACAM1,CRABP2,CTSE,CXCL2,CXXC5,DUSP2,EGR1,ETS1,IFI35,IFIH1,IL1RN,LCN2,MCL1,P3H4,PMAIP1,PSMB10,PSMB8,PTGS2,RBPMS,RHOB,SBNO2,SERPINE2, | 504 (22) |
| MAPK14 | | kinase | | 0.967 | 0.0000247 | AXIN2,CCNA2,CEBPD,CXCL8,DKK1,DNMT3A,DUSP1,EFNA1,EREG,FAS,FOS,GADD45A,HMGCR,IGF2,JUN,JUND,KITLG,MAOB,NRP1,PTGS2,SGK1,TGM2,VDR,ZFP36 | 470 (18) |
| 15(S)-HETE | | chemical - endogenous mammalian | | 0.961 | 0.0000132 | EGR1,FOSB,FOSL1,HMGCR,JUN,PPARG,PTGS2 | 424 (24) |
| KLF11 | | transcription regulator | | 0.961 | 0.00207 | APOL6,CAV1,CXCR4,EDN1,ENG,ITGB6,MAOB,PCK2,PDK4,PPARG,SERPINA1,SERPINH1,SMAD3,TGFB2,TGFBR2 | 253 (7) |
| CSF1 | | cytokine | | 0.96 | 0.0000952 | CDC25A,CEBPA,CXCL2,CXCL3,DUSP1,DUSP5,EGR1,F2R,F2RL1,FAS,FHL2,FOS,GDF15,HMGCR,IDI1,IRF7,JDP2,JUN,LDHA,LGR4,LSS,MSMO1,PPARG,SC5D,SERPINE2,SLC16A3 | 454 (19) |
| IRF6 | | transcription regulator | | 0.959 | 0.000185 | APOBEC3B,CXCL8,GRHL3,KLF4,LCN2,OVOL1,PPARG,PTGS2,SCARB1,TJP1 | 396 (20) |
| Ncoa-Nr1i3-Rxra | | complex | | 0.958 | 0.0207 | ABCC1,ABCC2,CYP2B6,CYP3A5 | |
| cyclopiazonic acid | | chemical - endogenous non-mammalian | | 0.957 | 0.00651 | CXCL2,EGR1,FAS,FOS | 485 (24) |
| VHL | | transcription regulator | | 0.956 | 0.0181 | ALDH1A1,AURKA,BBC3,CA9,CAV1,CXCR4,IFITM3,NEDD9,PMAIP1,SEC61G,SLC16A3,TMED3,ZFP36L1 | |
| vitamin K3 | | chemical drug | | 0.954 | 0.000928 | ABCC1,DYRK2,EGR1,FAS,FOS,KLF6,MAP1LC3B | 435 (20) |
| azoxymethane | | chemical toxicant | | 0.947 | 0.015 | BBC3,BMP4,CXCL2,CXCL3,FANCF,PRKCA,PTGS2,REN | |
| triadimefon | | chemical toxicant | | 0.946 | 0.0346 | ALDH1A1,CYP24A1,CYP2B6,CYP3A5,SQLE,TST | |
| indomethacin | | chemical drug | | 0.941 | 4.86E-08 | RC3,CD44,CD55,CLDN1,CLU,CPT1A,CXCR4,CYP2B6,DDC,ELF3,FOS,GDF15,IFITM3,LAMC2,LCN2,LGALS3,LIMA1,PIM1,PPARG,PRKCA,PSMB10,PSMB8,PTGS2,RCN1,REN,S100A | 480 (18) |
| GDF2 | | growth factor | | 0.935 | 0.00176 | CAV1,CXCL8,CXCR4,EDN1,ENG,FOS,GADD45B,GATA2,ID2,NRP1,SEMA3B,SERPINE2 | 361 (14) |
| NFKBIA | | transcription regulator | | 0.933 | 3.11E-10 | 3,CXCL8,CXCR4,ERAP2,EREG,FAS,FHL2,FOS,GADD45A,GADD45B,GBP3,GRK5,IGF2,IGFBP6,IL15RA,IL1RN,IRF1,JDP2,JUN,LAMA5,LCN2,LIF,LIMA1,MMP15,MUC2,MXD4,NDRG2 | 412 (18) |
| ALOX15 | | enzyme | | 0.933 | 0.00485 | AXIN2,CXCL2,FOSL1,JUN,PPARG,RGS2 | 480 (25) |
| BRCA1 | | transcription regulator | | 0.929 | 0.000000763 | PT1,AQP5,CARD10,COL18A1,CYP1A1,DDIT4,EGR1,FAS,FHL2,FXYD3,GADD45A,GADD45B,GDF15,HMGA2,HSPB1,IFIT3,IRF7,NEDD9,PHLDA1,PLSCR1,S100A2,SAT1,SERPINE2,S | 434 (16) |
| BMPER | | other | | 0.927 | 0.0279 | ACVR1,BMP4,HNF4A,RUNX2 | |
| FOXC2 | | transcription regulator | | 0.913 | 0.00953 | CASP9,CEBPA,CXCR4,DKK1,IRS2,PPARG,PTGS2,RUNX2 | 452 (23) |
| ethionine | | chemical toxicant | | 0.905 | 0.00000176 | ANXA1,CD44,CLU,LAMC2,LCN2,LGALS3,PSMB10,PSMB8,S100A10,TNFRSF12A,TSPAN8 | |
| EGLN | | group | | 0.904 | 0.00257 | AK4,CA9,EFNA1,EGLN1,FEM1C,FSTL3,FYN,HLA-B,LCN2,LDHA,PEX11A,PGM1,PPP2R5B,RNASE4,SLC16A3,UPK3B,ZNF292 | 330 (7) |
| polyamines | | chemical - other | | 0.9 | 0.00387 | CD44,CEBPA,PPARG,SAT1 | 309 (7) |
| CPT1B | | enzyme | | 0.898 | 0.0347 | ACSL1,CBLB,DGAT2,FLOT1,HK1,INSIG1,IRS2,PCK2,PDK4,PIK3R1,PPP1R3B,PRKAR2A,PYGB,RPS6KB2,SCARB1,UQCRFS1,UQCRHL | |
| HBEGF | 1.019 | growth factor | | 0.895 | 0.00139 | CXCR4,EGR1,EREG,LDHA,NRP1,PDK4,PTGS2,SLC16A3 | 495 (20) |
| potassium chloride | | chemical drug | | 0.894 | 0.0169 | ATF3,ATP1B1,ATP2B1,CASP7,EGR1,FGFR2,FOS,JUN,PIK3R1,PTGS2,TXNIP | |

Supplementary Table 4. Upstream regulator analysis based gene set enrichment analysis (GSEA) of differentially expressed genes (DEGs) after ATRA+PDT compared to PDT

| Upstream Regulator | Expr Log Ratio | Molecule Type | Predicted Activation State | Activation z-score | p-value of overlap | Target Molecules in Dataset | Mechanistic Network |
|---|---|---|---|---|---|---|---|
| trichostatin A | | chemical drug | | 0.89 | 8.53E-14 | ,FUT8,GADD45A,GATA6,GLI2,GSN,HDAC9,HMGA2,HMGCR,HMGCS2,ID2,IGSF1,IGSF9,INPPL1,IRF1,JUND,KLF4,LGMN,LIMA1,LRP8,MAP1LC3B,MAP4K4,MAT2A,MICALL2,MMP7, | 499 (20) |
| NR5A2 | 0.861 | ligand-dependent nuclear receptor | | 0.889 | 0.000371 | ABCC2,CEBPD,CXCL16,FGF19,FOS,HMGCR,HNF4A,IL1RN,JUN,MMP7,NR0B2,NR5A2,PLK3,SCARB1,TGFB2,TNFRSF12A | 527 (26) |
| BMP15 | | growth factor | | 0.882 | 0.0241 | EREG,ID2,PCSK6,PTGS2 | |
| HRAS | | enzyme | | 0.871 | 1.08E-13 | 1,DPP4,DUSP1,EGR1,EIF5A2,ETS1,F2R,FAS,FOS,FOSB,FOSL1,FSTL3,GJB2,GNA14,GSN,HLA-B,HSPB1,ID2,IFITM2,IGF2,IQGAP2,JUN,KLF6,LATS2,LIPG,LPP,MAOB,MCAM,MSM | 484 (19) |
| 17-alpha-ethinylestradiol | | chemical drug | | 0.87 | 0.0000233 | ABCC2,ARRDC3,BCL3,CD68,CKB,CTSS,CXCL3,CYP2B6,FOS,GDF15,JUN,KRT7,LCN2,NR0B2,PLK3,SCARB1,SERPINH1,SGK1,TFF3,TGM2,VEZF1 | 618 (21) |
| diethylnitrosamine | | chemical toxicant | | 0.863 | 0.0228 | HNF4A,IGFBP2,JUN,LCN2,MCL1,PHLDA1,PLSCR1,PRKCA,QPCT,SESN2,TM4SF4 | |
| AREG | | growth factor | | 0.859 | 0.00119 | CCNA2,CCNF,CXCR4,EGR1,ELF3,EREG,FOS,HJURP,IFIT3,JUN,MMP15,PTGS2 | 433 (24) |
| TAS1R3 | | G-protein coupled receptor | | 0.849 | 0.00144 | CEBPA,DDIT4,PPARG,SLC16A1 | 85 (3) |
| salmeterol | | chemical drug | | 0.849 | 0.00387 | CFTR,DDIT4,DUSP1,RGS2 | 217 (7) |
| farnesyl pyrophosphate | | chemical - endogenous mammalian | | 0.849 | 0.00651 | LIPG,PTGS2,SCARB1,SMAD3 | 375 (7) |
| ELF3 | 0.834 | transcription regulator | | 0.837 | 0.00485 | ANGPT1,EHF,ELF3,PPARG,PTGS2,TGFBR2 | 359 (17) |
| topotecan | | chemical drug | | 0.837 | 4.69E-08 | F3,AURKA,BCL3,BTG1,CDC25A,CLP1,DUSP5,EGR1,ENG,FAS,FOSB,GDF15,HLA-E,HSPA1A/HSPA1B,HSPA1L,JMJD1C,KIAA0040,MAT2A,MCL1,MYB,PLK3,PMAIP1,PTGS2,SESN | 404 (12) |
| cadmium | | chemical toxicant | | 0.835 | 0.00103 | ACAA1,B2M,CASP9,CHPT1,CYP1A1,JUN,LGALS3,PTGS2,SGK1,SLC30A1,SLC39A8 | 406 (19) |
| ETV4 | | transcription regulator | | 0.834 | 0.000288 | CAV1,CXCL8,CXCR4,HPGD,MMP7,MYB,PTGS2,TGFBR2 | 342 (7) |
| dehydroisoandrosterone | | chemical - endogenous mammalian | | 0.832 | 0.00613 | B2M,BMP4,CPT1A,CYP1A1,CYP2B6,CYP3A5,ECM1,LGALS3,PPARG,TAP2 | 542 (14) |
| ziritaxestat | | chemical drug | | 0.832 | 0.000000734 | AKR1C1/AKR1C2,AKR1C3,ALDH1A3,APOL6,CA9,CEMIP,CREBRF,DDIT4,GPRC5A,IFIT3,KLHL24,OAS3,SGK1 | |
| BIRC2 | | enzyme | | 0.831 | 0.00183 | BIRC3,CCNA2,CXCL1,CXCL8,MAP3K14 | 423 (17) |
| PRKN | | enzyme | | 0.831 | 0.0406 | AXIN2,CDK6,MAOA,MAOB,MCL1 | |
| IGF2 | -0.568 | growth factor | | 0.83 | 0.00000303 | L4A,CCNG2,CXCR4,DUSP1,EGR1,ELOVL6,EREG,GADD45A,GATA6,HBEGF,IFITM2,IGF2,IGFBP2,IGFBP6,KLF9,PHLDA1,PIK3R1,PIM1,PTGS2,RHOB,SAT1,SEZ6L2,SKAP2,SLC30 | 482 (20) |
| GSK3B | | kinase | | 0.828 | 0.0198 | BIRC3,CXCL8,DNMT3A,FOS,GLI1,IRS2,LCN2,MCL1,MYB,NFKBIZ,PTGS2,SLC16A1 | |
| MET | | kinase | | 0.823 | 0.0000107 | AURKA,CD44,CLDN1,CXCL1,CXCL8,EDN1,FBXO32,FOS,HBEGF,HSPB1,IL15RA,JAG2,JUN,KLF4,NFKBIZ,ONECUT2,PIM1,PTGS2,TRIM21,TSC22D1 | 398 (20) |
| eicosapentenoic acid | | chemical drug | | 0.822 | 0.000296 | ACAT1,CD44,CEBPD,DNAJB9,FOS,IL1R2,JUN,KRT20,PDK4,PNPLA2,PPARG,PTGS2,TGFB2,TGFBR2 | 418 (23) |
| ketamine | | chemical drug | | 0.821 | 0.00367 | CXCL8,FOS,HK1,KLF6,PTGS2 | 443 (20) |
| ciprofibrate | | chemical drug | | 0.819 | 0.0000308 | ACAA1,CEACAM1,DPP4,IGFBP2,LCN2,LGALS3BP,MYB,NPC1,PDK4,PEX11A,PHLDA1,PLSCR1,PPARG,PXMP2,QPCT,SCARB1,SERPINA1,SLC16A1,SLC4A4,TM4SF4 | 412 (19) |
| asoprisnil | | chemical drug | | 0.816 | 0.00754 | DKK1,DPP4,IGFBP2,RARRES1,SGK1,TGFBR2 | |
| SELPLG | | other | | 0.816 | 0.0177 | CLK1,CXCL2,CXCL8,CXCR4,FOS,IL1R2 | |
| LCAT | | enzyme | | 0.816 | 0.000518 | AXIN2,CEBPA,CLU,HMGCR,PPARG,SCARB1 | 183 (7) |
| AIP | | transcription regulator | | 0.816 | 0.00946 | CYP1A1,DUSP1,F2RL1,GPRC5A,HTR1D,PDE9A,RGS2 | |
| CSF2 | | cytokine | | 0.813 | 1.66E-09 | L8,CXCR4,DUSP2,E2F7,EDN1,EGR1,ERCC6L,F2R,F2RL1,FAS,FIGNL1,FOS,FOSL1,GDF15,HBEGF,HJURP,HMGCR,HSPH1,ID2,IFITM3,IL1RN,JUN,LDHA,MAT2A,MCL1,MYB,NOX1 | 465 (16) |
| AHR | | ligand-dependent nuclear receptor | | 0.809 | 0.000000984 | ,CYP2B6,CYP2S1,DNAJB1,EDN1,EGLN1,FAS,FGFR2,FOS,FOSL1,GADD45A,HDAC9,HMGCR,HSPH1,IRF1,JUN,LDHA,LTBP3,MT2A,NEDD9,NT5E,PJA1,PPARG,PPP1R15A,PTGS2 | 573 (24) |
| GDF9 | | growth factor | | 0.808 | 0.0368 | HSPG2,ID2,KITLG,NRIP1,PTGS2 | |
| PRKAA | | group | | 0.808 | 0.0264 | CLDN1,CPT1A,DUSP1,DUSP2,MCL1,PMAIP1 | |
| PIK3R1 | -0.308 | kinase | | 0.807 | 0.000071 | CD44,CXCL8,DNAJB9,DNAJC3,ENG,ERN1,FOS,IRF7,LDHA,PIK3R1,PPARG,PTGS2,TGFB2,TXNIP | 526 (22) |
| IL7 | | cytokine | | 0.804 | 0.00614 | CBLB,CDK6,CEACAM1,CXCL1,CXCL2,CXCL3,CXCR4,DPP4,DUSP5,FAS,LIF,MCL1,PIM1,PMAIP1,RUNX2 | 388 (19) |
| beta-naphthoflavone | | chemical toxicant | | 0.802 | 0.00316 | AKR1C1/AKR1C2,AKR7A3,CCN1,CYP1A1,CYP2S1,FOS,ME1,UGT1A10 (includes others) | 468 (20) |
| MMP9 | | peptidase | | 0.8 | 0.000114 | CD44,CD68,COL18A1,CXCL2,CXCR4,FOS,GADD45A,HPGD,JAG2,SDC4,SERPINA1,SERPINE2,TJP1 | 510 (19) |
| MYBL2 | | transcription regulator | | 0.798 | 0.00946 | AURKA,BIRC3,CCNA2,CCNG2,CEBPA,CLU,GATA2 | |
| CTNNBIP1 | | other | | 0.795 | 0.00207 | AXIN2,HMGCR,IDI1,NEDD9 | 290 (7) |
| KMT2D | | transcription regulator | | 0.794 | 0.00214 | CD55,CSPG4,DKK1,DUSP2,GJB2,GRHL3,IGSF1,IRF1,JAG2,KLF4,MYOM3,NT5E,PPARG,SCARB1,SH3BGRL2,SIX4,SORD,TCEAL1,ZNF184,ZNF641 | 257 (7) |
| triamcinolone acetonide | | chemical drug | | 0.786 | 6.97E-10 | 4,DUSP1,EGR1,FGFR2,FOS,FUT4,GNAZ,HIPK2,INSIG1,IRF7,JUN,KLF9,MT1E,MT1X,MT2A,MXD4,PDIA5,PSPH,RGS2,SERPINA3,SLC16A1,SOX4,TBC1D1,TGFB2,TRIM21,TSC22D | 324 (13) |
| AKT1 | | kinase | | 0.781 | 2.59E-08 | IRC3,CD44,CEBPD,CLU,CXCL2,CXCL8,CXCR4,EGR1,FAS,FBXO32,FOS,FOSB,GDF15,HMGCR,IGF2,IRS2,JUND,LCN2,LSS,MCL1,ME1,MSMO1,MT2A,MYB,PEX11A,PHLDA1,PIK3 | 517 (22) |
| GATA3 | | transcription regulator | | 0.778 | 0.0108 | BCAS3,CEBPA,CRABP2,CXCR4,ELOVL6,ETS1,FOS,GRHL3,KLF4,LOC102724788/PRODH,NCOA3,NR5A2,OVOL1,PPARG,PPL,RCAN1,SP110,TJP1,TJP2,VSIR,WNT11 | |
| Akt | | group | | 0.778 | 1.09E-09 | SL1,GATA2,GCNT2,GLI1,GLI2,HPGD,HSPG2,IGF2,IGFBP2,JUN,LAMC2,LCN2,MCAM,MCL1,MSLN,MT1X,MT2A,MYB,NPC1,NR0B2,NT5E,PDK4,PMAIP1,PPARG,PPP1R1B,PTGS2,F | 613 (22) |
| RGS2 | 0.675 | enzyme | | 0.775 | 0.0000927 | CEBPD,EFNA1,ERRFI1,FXYD3,HBEGF,PDLIM1,PLXND1,PPARG,RCAN1,SCARB1,SDF2L1,SLC16A1,SLC16A3,SLCO4A1,UGCG | 381 (12) |
| Pka | | complex | | 0.768 | 0.000225 | CAV1,CD44,DUSP1,EDN1,EGR1,FOS,HMGCR,HNF4A,IRF1,JUN,KLF6,NOX1,PLD1,PPP1R1B,PTGS2,REN,RGS2,SGK1 | 372 (22) |
| butyric acid | | chemical - endogenous mammalian | | 0.766 | 8.92E-12 | APK1,DUSP1,FOS,FYN,GADD45A,GATA2,GATA6,GPRC5A,GSN,HDAC9,HK1,HSPG2,IFIT3,IFITM2,IFNGR2,IGF2,IRF1,ITPR2,KLF4,KRT13,MCL1,MMP7,MT2A,MUC2,MXD1,MYB,NF | 521 (21) |
| ABL1 | | kinase | | 0.765 | 0.0446 | CAV1,CCNA2,FAS,GADD45A,TGFB2 | |
| WR 1065 | | chemical drug | | 0.765 | 0.00167 | CASP4,FAS,GADD45A,GDF15,S100A4,ZFP36L1 | 456 (17) |
| corticosteroid | | chemical drug | | 0.762 | 0.0302 | CXCL8,DUSP1,HIPK2,IL1R2,IL1RN,MAFF,PARP14,PPARG | |
| MAPKAPK3 | | kinase | | 0.762 | 0.0101 | CXCL2,CXCL3,HSPB1,ZFP36 | |
| vinorelbine | | chemical drug | | 0.762 | 0.0101 | B2M,HLA-E,LGALS3,PMAIP1 | |
| SRF | | transcription regulator | | 0.761 | 7.81E-08 | EGR1,ETS1,ETV1,FHL2,FOS,FOSB,FOSL1,GADD45A,GSN,HES6,HSPG2,JUN,LCN2,MALAT1,MAP3K14,MAP4K4,MCL1,MINDY2,MYB,MYO1B,NCOA7,PDLIM7,PGM1,PTGS2,RAB2 | 331 (19) |
| REL | 0.371 | transcription regulator | | 0.756 | 0.00333 | ACSS2,B2M,BBC3,BCL3,COTL1,CXCL8,DMTF1,FAS,FOS,FOSL1,FRMD4B,GADD45B,HBEGF,JUN,PARP14,PFKM,PLAAT3,PMAIP1,PPP1R15A,RAB40C,REL,SH2B3 | 474 (20) |
| KIF3A | | enzyme | | 0.751 | 0.00367 | ACAT1,GLI1,GLI2,HMGCR,IDI1 | 285 (7) |
| MST1 | | growth factor | | 0.748 | 0.00135 | FAS,FOS,IL1RN,NR0B2,PTGS2,SOX4 | 419 (19) |
| ARNT | | transcription regulator | | 0.744 | 0.0000277 | ALDH7A1,ATP5MC1,CA9,CAV1,CLDN1,CXCR4,CYP1A1,CYP2S1,DEGS2,EDN1,ENG,HES6,HNF4A,ID2,IRS2,LDHA,PCK2,TUBA4A | 441 (11) |
| CDH1 | | other | | 0.742 | 0.000872 | BMP4,CD44,FOSL1,GLI1,GLI2,JUN,KRT13,KRT7,LCN2,MCAM,PIK3R1,S100A4 | 446 (20) |
| CUX1 | | transcription regulator | | 0.728 | 0.0138 | ARHGDIB,CCNA2,CXCL1,CXCL2,CXCL3,CXCL8,HLA-B,LIMA1 | |
| zoledronic acid | | chemical drug | | 0.728 | 0.00592 | CCN1,CXCL8,CXCR4,DKK1,PPARG,PTGS2,RUNX2 | 384 (21) |
| letrozole | | chemical drug | | 0.728 | 0.0125 | AGR2,DKK1,FOSL1,LIF,NCOA3 | |
| decitabine | | chemical drug | | 0.723 | 3.8E-17 | FAS,FLRT3,FRYL,GADD45B,GATA6,GDF15,GLI2,GSN,HBEGF,HLA-B,HLA-C,HLA-E,HMGA2,HMGCS2,HNF4A,HPGD,HSPA1A/HSPA1B,IFI35,IGFBP2,IGSF9,INPPL1,IRF7,KLF4,KLF | 549 (18) |
| ketoconazole | | chemical drug | | 0.721 | 0.00246 | ABCC2,CXCL8,CYP1A1,CYP2B6,HMGCR,VDR | 602 (24) |
| seocalcitol | | chemical drug | | 0.718 | 0.00092 | CEACAM1,CYP24A1,FOSL1,GADD45A,MCL1,PDLIM7,PTGS2,SGK1,TSPAN6,VDR | 451 (24) |
| CXCL8 | 1.047 | cytokine | | 0.718 | 0.000148 | ANXA1,CD44,COL18A1,CXCL2,CXCL3,CXCL8,FAS,HSPG2,LGALS3,NOX1,PLD2,PTGS2,ZFP36 | 443 (19) |
| SULT1E1 | | enzyme | | 0.714 | 0.0045 | CD68,ELOVL6,LCN2,PTGS2,SCARB1 | 237 (7) |
| E2F3 | | transcription regulator | | 0.713 | 0.0000485 | R1C1/AKR1C2,AKR1C3,BAIAP2L1,CASP7,CAV2,CCNA2,CDC25A,COL18A1,CXCL1,EDN1,ELF3,FGFR2,IGF2,KRT80,MALAT1,MT1X,MYB,RBPMS,SERPINH1,TNFAIP2,WFDC2,ZNF | |
| KAT2A | | enzyme | | 0.707 | 0.00288 | AKAP12,CDC25A,DNAJB9,FBXO32,JUN,LDHA,NUDC,PIK3R1,SLC16A1,TAF1D | 386 (14) |
| ZEB1 | | transcription regulator | | 0.694 | 0.0156 | ALDH1A1,CCN1,CCNG2,CDKN2B,EDN1,LAMC2,LPCAT1,NRP1,PTGS2,S100A10 | |
| GAB2 | 0.451 | other | | 0.692 | 0.0239 | ANGPT1,FOS,GSN,MCL1,MUC2 | |
| KT5823 | | chemical - kinase inhibitor | | 0.692 | 0.000778 | AQP5,EDN1,FOS,S100A10,SLC12A4 | 426 (17) |
| IRAK4 | | kinase | | 0.689 | 2.23E-11 | ATF3,BIRC3,CSRNP1,CXCL16,CXCL2,CXCL3,CXCL8,EDN1,EGR1,ERRFI1,FOS,FOSB,IRF1,IRF7,JUN,LIF,NFKBIZ,PHLDA1,PLSCR1,PPARG,ZC3H12A,ZFP36 | 494 (16) |
| FGF19 | 1.996 | growth factor | | 0.689 | 0.000000258 | ACSL1,ALDH1A1,CTSE,CXCL2,EGR1,FOS,GADD45A,HMGCR,IL1RN,LCN2,MCAM,NR0B2,RGS2,SCARB1,SERPINA1,SERPINA3,SGK1,SQLE | 413 (12) |
| WT1 | | transcription regulator | | 0.687 | 0.000000019 | EFNA1,EGR1,EREG,ETS1,FOSB,GSN,HBEGF,HSPG2,IDI1,IGF2,ITM2B,JUND,KLF4,LGALS3,LSS,MCL1,MSLN,MT1X,NQO2,PCK2,PHLDA1,PTPN9,RAB40C,SMAD3,SMIM29,SQLE, | 376 (9) |
| FOXP3 | | transcription regulator | | 0.686 | 0.000634 | ANXA1,BBC3,CXCR4,EGR1,FOS,ID2,IFNGR2,JUN,LATS2,LDHA,LGALS3,MGAT1,NRP1,NT5E,PPARG,SDC4,SLC16A3,SLCO4A1,SWAP70,TGM2 | 462 (17) |
| PRKAR2B | | kinase | | 0.686 | 0.0461 | BIRC3,EGR1,FOS,PPARG | |
| phenobarbital | | chemical drug | | 0.682 | 0.0248 | ABCC2,CYP1A1,CYP2B6,CYP3A5,GADD45B,HNF4A,INSIG1,SAT1,VEZF1 | |
| 10E,12Z-octadecadienoic acid | | chemical - endogenous mammalian | | 0.68 | 0.000326 | ACSS2,ADAM8,ATF3,CAV1,CEBPA,CLIC5,CPT1A,CREB5,CXCL8,DGAT2,GDF15,IL1RN,ITPKA,ME1,PNPLA2,PPARG,PPP1R15A,PPP1R3B,PTGS2 | 368 (22) |
| oxaliplatin | | chemical drug | | 0.677 | 0.0138 | BBC3,CASP9,FAS,FXYD3,GDF15,MMP7,PMAIP1,SAT1 | |
| cholecalciferol | | chemical - endogenous mammalian | | 0.676 | 0.00101 | CEACAM6,CEBPA,CEBPD,CYP24A1,EDN1,FOS,JUN,ME1,NFYA,NR0B2,PPARG,PTPRB,RUNX2,TGFBR2,VDR | 425 (21) |
| PLG | | peptidase | | 0.67 | 0.0323 | CAV1,CCN1,CYP2S1,EDN1,FYN,PIP5K1A,PTGS2,VDAC1 | |
| FXR ligand-FXR-Retinoic acid-RXRα | | complex | | 0.667 | 0.00656 | ABCC2,FGF19,NR0B2,NR5A2,SCARB1 | |
| trans-hydroxytamoxifen | | chemical drug | | 0.661 | 4.44E-09 | ,BTG1,CAV1,CCNA2,CLU,CXCL8,DDIT4,DUSP1,EFNA1,EGR1,GNE,GPRC5A,KLF6,KRT7,LDHA,LOXL4,MXD4,MYO1B,RAB30,RAP1GAP,RND3,SLC22A5,SMAD3,SNCG,SP100,TGF | 466 (13) |
| methyl-beta-cyclodextrin | | chemical drug | | 0.651 | 0.00135 | CAV1,CXCL1,CXCL8,HMGCR,PPARG,SCARB1 | 490 (26) |
| MAPK10 | | kinase | | 0.649 | 0.00000516 | BBC3,CASP9,CD44,CXCL8,FOS,JUN,PTGS2 | 500 (19) |
| SAA1 | | transporter | | 0.648 | 0.0368 | BGN,CEBPA,CXCL8,IL1RN,PPARG | |
| ANGPT2 | | growth factor | | 0.644 | 0.000174 | AXIN2,CD68,COL18A1,DNAJB4,EDN1,ETS1,FAS,GATA6,HSPA1A/HSPA1B,KDM6B,MMP7,NOX1,PDIA5,PHLDA1,PROS1,PTGS2,RCAN1,RHOB,STK17A,TCN2,TGFBR2,TNFRSF12 | 494 (22) |
| mitoxantrone | | chemical drug | | 0.64 | 0.0126 | BBC3,CXCL8,FAS,IRF7,MCL1,PMAIP1 | |
| imipramine | | chemical drug | | 0.64 | 0.00203 | DEPP1,EGR1,FOS,LSS,MAP1LC3B,SERPINA3 | 479 (24) |
| SIRT6 | | enzyme | | 0.633 | 0.000809 | BIRC3,CD68,CXCL8,ELOVL6,HMGCR,IRS2,LSS,NPNT,PLPP3,PTGS2 | 523 (23) |
| BTNL2 | | transmembrane receptor | | 0.632 | 0.00352 | CYP2S1,DAPP1,GBP3,GJB2,IFITM3,MXD1,NCOA7,NT5E,PHLPP1,S100A4 | |
| DSCAM | | other | | 0.632 | 0.0179 | CEBPA,GSN,KLF4,KLF9,MICALL2,NTN4,PLD1,SEMA3E,SEMA7A,TUBA4A | |
| mir-204 | | microRNA | | 0.632 | 0.00000353 | ACVR1,CXCL1,CXCL2,CXCL8,DNMT3A,ID2,MCL1,PIK3R1,RUNX2,SMAD3,SPDEF,STX1A,TGFBR2 | 251 (7) |
| 3,3'-diindolylmethane | | chemical drug | | 0.623 | 0.000438 | ATF3,BBC3,CDC25A,CDK6,FAS,GDF15,HLA-B,HLA-C,MAP1LC3B,PTGS2,TAP2 | 514 (18) |
| DRD2 | | G-protein coupled receptor | | 0.621 | 0.0158 | FOS,FOSB,ITGB6,JUN,PNPLA2,TGFBR2 | |
| nifedipine | | chemical drug | | 0.62 | 0.0346 | ATP2B1,CYP2B6,CYP3A5,EDN1,FOS,HMGCR,PTGS2,SGK1 | |
| E2F1 | | transcription regulator | | 0.619 | 0.000185 | NA2,CCNF,CD44,CDC25A,CEBPA,CKB,CRABP2,DUSP1,EGR1,EIF2AK2,FAS,FGFR2,FOS,IGF2,IRS2,JAG2,MAP3K14,MCL1,MYB,NCOA3,NMI,NRIP1,NRP1,NUDC,PDK4,PIK3R1,PK | 475 (19) |
| BCR-ABL1 | | fusion gene/product | | 0.613 | 0.00488 | AURKA,CAV1,CEBPA,CXCR4,FYN,HK1,LCN2,LDHA,MXD4,MYB,PIM1 | 519 (23) |
| DDIT3 | | transcription regulator | | 0.611 | 0.0000013 | ATF3,BBC3,CASP4,CEBPA,CEBPD,CXCL8,DNAJA1,LCN2,LTBP3,MCL1,NPAS2,PCK2,PPARG,PPP1R15A,STX1A,TGFB2,TGM2,TSC22D3,TXNIP,WARS1 | 476 (15) |
| acetaminophen | | chemical drug | | 0.608 | 0.0182 | ABCC1,ABCC2,ACO1,AK4,CXCL2,CXCL3,CYP3A5,FAS,GADD45A,HMGCR,HMGCS2,JUN,MCL1,ME1,NDRG2,PTGS2 | |
| Laminin (complex) | | complex | | 0.602 | 0.00457 | BIRC3,ID2,IGF2,PTGS2,RUNX2,SDC4,TGFB2 | 477 (22) |
| PRKCI | | kinase | | 0.6 | 0.0144 | CXCL2,CXCL3,ELF3,HK1,PTGS2 | |
| NOTCH1 | | transcription regulator | | 0.586 | 5.04E-08 | CCN1,CD44,CD68,CEBPA,CEBPD,CXCL1,CXCL8,CXCR4,DKK1,DNAJA1,FOS,FOSB,FOSL1,GATA2,GLI1,GLI2,HSPH1,ID2,IGFBP2,IRF1,MCAM,MUC2,PPARG,PTGS2,REL,REN,RH | 502 (23) |

Supplementary Table 4. Upstream regulator analysis based gene set enrichment analysis (GSEA) of differentially expressed genes (DEGs) after ATRA+PDT compared to PDT

| Upstream Regulator | Expr Log Ratio | Molecule Type | Predicted Activation State | Activation z-score | p-value of overlap | Target Molecules in Dataset | Mechanistic Network |
|---|---|---|---|---|---|---|---|
| IL32 | | cytokine | | 0.58 | 0.00635 | BBC3,CXCL1,CXCL3,CXCL8,EIF2AK2,FAS,MCL1,PTGS2 | 424 (17) |
| retinoid | | chemical drug | | 0.579 | 0.00102 | CEACAM1,DUSP1,EDN1,HBEGF,LIMA1,MUC2,PLAAT4,PTGS2 | 478 (24) |
| miR-145-5p (and other miRNAs w/seed UCCAGUU) | | mature microRNA | | 0.578 | 0.00578 | ACSS2,CCNA2,KLF4,KRT7,PADI1,PARP8,RASA1,RTKN,SWAP70 | |
| TET2 | | enzyme | | 0.577 | 0.0224 | ADAM8,BCL3,CEBPA,CYP2S1,GRK5,IFIT3,IFITM3,KCNE3,MAFF,MTSS1,RND3,SH2B3,SORL1 | |
| BHLHE40 | | transcription regulator | | 0.577 | 0.0131 | ANXA1,ATP1B1,CRABP2,CXCL3,ECM1,IFITM2,IL1R2,MYB,PHLDA1,PLAC8,RND3,TXK | |
| PRDX1 | | enzyme | | 0.577 | 0.00387 | ECM1,EGLN1,PTGS2,REL | 217 (7) |
| lipoarabinomannan | | chemical - endogenous non-mammalian | | 0.577 | 0.0241 | CXCL2,CXCL8,CXCR4,IRF1 | |
| CCN2 | | growth factor | | 0.574 | 0.0371 | CD44,EGR1,FLNB,IGF2,JUN,KITLG,LTBP3,MIA3,SDC4,SOX4 | |
| mir-223 | | microRNA | | 0.573 | 0.0113 | ALCAM,CAMKK1,EMP1,GASK1B,ID2,IL1RN,MSMO1,MT1E,MUC13,MYOF,NDRG2,SCARB1,STMN1,TOB1 | |
| NR3C1 | | ligand-dependent nuclear receptor | | 0.57 | 6.41E-15 | SP1,DUSP16,EDN1,EFNA1,EGR1,EMP1,ERN1,ERRFI1,F2R,F2RL1,FOS,GAB1,GADD45A,GADD45B,GLCCI1,HGD,HMGCS2,HSPA6,IFIH1,IL15RA,IRF1,ITGB6,JUN,JUND,KLF9,LIF,I | 562 (21) |
| SHH | | peptidase | | 0.57 | 0.00274 | AMD1,ANGPT1,AQP5,BMP4,CAV1,CCNG2,CDC25A,DDC,FGF19,GATA2,GLI1,GLI2,IGF2,PPARG,RUNX2,SOSTDC1,TGFB2,TJP1,TSC22D3 | 463 (22) |
| fish oils | | chemical drug | | 0.57 | 0.000131 | ACSL1,CEBPA,CPT1A,CYP1A1,CYP2B6,EDN1,NRIP1,NUCB2,PIK3R1,PTGS2 | 362 (10) |
| Tcf7 | | transcription regulator | | 0.566 | 0.0000362 | 6,ARRDC4,ECM1,FAAH,FYN,ID2,IL15RA,KLF6,LRIG1,MBOAT1,MSMO1,MXD1,NCOA7,NR5A2,NTN4,OSBPL5,PHLDA1,PPARG,RARG,REL,SC5D,SH3BGRL2,SOX12,SQLE,TBL1X,T | |
| INSIG1 | -0.416 | other | | 0.566 | 0.000000547 | ACSS2,CD68,CXCL2,CXCL3,CXCR4,ELOVL6,HMGCR,HMGCS2,IDI1,IRS2,LCN2,LIPG,LSS,NFKBIZ,PLD1,PPARG,SCARB1,SQLE,STEAP2 | 343 (11) |
| SQSTM1 | | transcription regulator | | 0.563 | 0.000866 | CXCL2,CYP2B6,EDN1,HBEGF,IL15RA,IRF1,JUN,MAP1LC3B,TSC22D1 | 422 (16) |
| UCHL1 | | peptidase | | 0.563 | 0.0346 | CGAS,HDAC9,HERC6,ITPRIP,LPP,PFKM,PSME1,SLC39A8 | |
| WNT3A | | cytokine | | 0.557 | 9.21E-09 | IN2,BBC3,BGN,BMP4,CCN1,CD44,CD68,CEBPA,CYP24A1,DDIT4,DKK1,DVL1,EDN1,ENG,FHL2,GLI1,IRS2,JDP2,JUN,KITLG,KLF4,LCN2,MCL1,MT1X,NR5A2,PLD1,PPARG,PTGS2,I | 461 (22) |
| 4-nitroquinoline-1-oxide | | chemical toxicant | | 0.555 | 0.000778 | AURKA,CCNA2,CDK6,EGR1,PTGS2 | 367 (7) |
| POMC | | other | | 0.547 | 0.0263 | CXCR4,DUSP1,FOS,FOSL1,IGF2,JUN,PCSK5,REN,SCARB1,SMAD3 | |
| EGFR | | kinase | | 0.539 | 8.93E-13 | F2RL1,FOS,FOSL1,GADD45A,GCNT2,HBEGF,HMGA2,HSPB1,IGF2,IQGAP2,IRF1,ITGB6,JUN,JUND,LAMC2,LCN2,LRP8,MACF1,MCL1,MGAT1,NOX1,NT5E,NUDC,PTGS2,RHOB,RU | 530 (22) |
| plicamycin | | chemical drug | | 0.537 | 0.000000412 | BBC3,CA9,CDC25A,CXCL1,FYN,GJB2,GPD1,HMGCR,MAOA,MUC2,PMAIP1,PPP1R1B,ROR1,SEMA7A,SERPINE2,STARD10,STX1A,TGFBR2,UBC,VDR | 481 (19) |
| EHF | 0.415 | transcription regulator | | 0.535 | 0.000238 | BMP4,CRABP2,CYP1A1,EHF,EREG,GATA2,GRHL3,IL1RN,KLK6,SCEL,SERPINA3,SPDEF,TFPI,ZFP36L1 | |
| phenylbutazone | | chemical drug | | 0.532 | 0.00000528 | ABCC2,ANXA1,CD44,CLU,LAMC2,LCN2,LGALS3,PSMB8,PTGS2,RCN1,S100A10,TNFRSF12A,TSPAN8 | 328 (11) |
| CDK8 | | kinase | | 0.528 | 0.00295 | CD44,CXCL1,CXCL2,CXCL8,CXCR4 | 368 (11) |
| ADM | | other | | 0.526 | 0.0443 | CXCL2,EDN1,FOS,PPARG,PTGS2,REN | |
| CIP2A | | other | | 0.525 | 0.0000893 | CDKN2B,GADD45A,PDK4,PDLIM7,PTPRB,RHOD,S100A16,SAT1,SERPINE2,SLC16A3,SLC22A18,TUBA4A | 349 (8) |
| deferoxamine | | chemical drug | | 0.508 | 0.000000102 | 3A5,DUSP1,DUSP16,DUSP5,ECM1,EGLN1,ERRFI1,FOS,GADD45A,GADD45B,GDF15,HSPA1A/HSPA1B,IGFBP2,JAG2,JUN,LCN2,LDHA,MAP1LC3B,ME1,PMAIP1,PPARG,PRKCA,S | 452 (18) |
| TFAP4 | | transcription regulator | | 0.508 | 0.0000133 | CASP9,CCNG2,CD44,CLDN1,EGR1,ETV1,FOS,GDF15,LIF,STEAP3 | 368 (7) |
| AR | | ligand-dependent nuclear receptor | | 0.507 | 1.56E-12 | H1,IFIH1,IGF2,KITLG,LAMA5,LRIG1,LRRK1,MAOA,MSMO1,MT1X,MYB,MYO1B,NPC1,NR5A2,PDE9A,PDIA5,PMAIP1,PRKCA,PROS1,PSCA,RARRES1,REN,RHOB,SCEL,SEC61B,SE | 547 (26) |
| NR1I2 | | ligand-dependent nuclear receptor | | 0.504 | 0.00000158 | 10,ABCC2,ALDH1A1,CEBPA,CPT1A,CYP1A1,CYP24A1,CYP2B6,CYP3A5,EGR1,ELOVL6,ENTPD5,FGF19,HMGCS2,HNF4A,INSIG1,JUN,LCN2,MYB,NR0B2,PPARG,SCARB1,TCN2, | 651 (24) |
| SP1 | | transcription regulator | | 0.494 | 7.54E-16 | R1,EIF2AK2,ENG,ENTPD5,EREG,F2R,FAS,FOS,FOSL1,FYN,GDF15,HBEGF,HMGCR,HNF4A,IFITM3,IGF2,IRF1,JUN,KLF4,KLF6,LTBP3,MAOA,MAOB,MAT2A,MCL1,MUC2,NCOA3,I | 537 (20) |
| EGR2 | | transcription regulator | | 0.493 | 0.00102 | ABCC1,ACVR1,ASS1,BCL6,CBLB,EGR1,FOS,HMGCR,ID2,IGF2,IL1R2,JUN,KITLG,LTBP3,MYB,NCMAP,SLC16A1,SQLE | 476 (20) |
| KAT2B | | transcription regulator | | 0.491 | 0.0346 | B2M,FOSB,GLI1,HLA-B,PTGS2,TGFBR2 | |
| 3-methyladenine | | chemical toxicant | | 0.489 | 0.00221 | AGR2,CXCL2,CXCL8,MAP1LC3B,PTGS2,RCAN1,RUNX2 | 401 (21) |
| MTPN | | transcription regulator | | 0.487 | 0.000168 | ANXA1,CASP7,CASP9,CCNA2,CEBPA,DDC,EDN1,ENG,FAS,FOS,JUN,S100A10,TGFB2 | 370 (11) |
| LGALS1 | | other | | 0.481 | 0.0281 | CXCL16,CXCL2,GLI1,GPCPD1,HBEGF,IFNGR2,INSIG1,LGALS3 | |
| HSD17B4 | | enzyme | | 0.478 | 0.000147 | ACAA1,ACSL1,GLI1,GLI2,PDK4,PEX11A,PTGS2 | 481 (7) |
| Growth hormone | | group | | 0.474 | 3.42E-09 | BPD,CLU,COL18A1,CXCL1,CXCL2,DYRK2,EGR1,FOS,GDF15,IGF2,IGFBP2,IGFBP6,IRF1,JUN,NDRG2,NEDD9,NR1D1,PDK4,PLAAT3,PPARG,PPP1R15A,S100A4,SCARB1,SGK1,TE | 540 (23) |
| gemcitabine | | chemical drug | | 0.472 | 0.000042 | CASP9,CDC25A,CPT1A,CXCL8,CXCR4,CYP3A5,EGR1,FAS,FOSL1,MDK,NT5E,PTGS2,SLC28A3 | 451 (24) |
| AGT | | growth factor | | 0.471 | 6.16E-16 | 1,GLI2,GRK5,HBEGF,HMGCR,HSPA1A/HSPA1B,HSPA1L,HSPB1,HSPG2,IDI1,IGFBP2,IRS2,ITPR2,JUN,KITLG,LATS2,LCN2,LGALS3,LIF,LIPG,LOXL4,LPP,LRP8,LSS,LTBP3,LTBP4, | 635 (23) |
| IRF2 | | transcription regulator | | 0.47 | 6.47E-10 | A13,B2M,CEACAM1,CFTR,CLDN2,CTSS,DST,DUSP5,EIF2AK2,HGD,HLA-B,IFI35,IRF1,IRF7,KLF4,LCN2,MMP7,PSMB10,PSMB8,PSME1,PSME2,PTGS2,RARRES1,TAP2,TAPBP,TRI | 293 (18) |
| daunorubicin | | chemical drug | | 0.464 | 0.000188 | ABCA2,ABCC1,CFTR,DMTF1,MCL1,PFKM,TAP2,UQCRFS1 | 428 (17) |
| 3,4,5,3',4'-pentachlorobiphenyl | | chemical toxicant | | 0.461 | 0.0232 | BMP4,CAPN9,CD44,CYP1A1,GADD45A,HSPA1A/HSPA1B,JUN,NR0B2,TIPARP | |
| mir-130 | | microRNA | | 0.454 | 0.0333 | CD68,HOXA5,KLF4,LCN2,PTGS2 | |
| BMP2 | | growth factor | | 0.453 | 0.00345 | AOC1,BMP4,DKK1,ETS1,FGFR2,FOS,FOSB,GADD45B,ID2,JUN,KLF4,KLF9,NFYA,PCSK6,PPARG,PTGS2,RUNX2,SOSTDC1,TGM2,TOB1,VDR | 544 (25) |
| AGER | | transmembrane receptor | | 0.45 | 0.0281 | CXCL2,CXCL3,CXCL8,EGR1,FAS,PTGS2,TJP1,TXNIP | |
| YY1 | | transcription regulator | | 0.449 | 0.000475 | P4,CASP7,CCNA2,CDKN2B,CFTR,CLU,CXCR4,CYP24A1,DNAJB4,DNMT3A,EDN1,EGR1,FAS,FOS,GRIN2D,HMGCR,ID2,IL1R2,INCENP,KLF4,MAT2A,NFKBIZ,PPP1R15A,RND3,SC | 506 (23) |
| ibrutinib | | chemical drug | | 0.447 | 0.00656 | CD44,CDK6,FAS,MCL1,UGCG | 319 (17) |
| THRA | | ligand-dependent nuclear receptor | | 0.447 | 0.00653 | BMP4,CD44,CEBPA,CIRBP,CPT1A,CXCL2,CXCL3,ENG,FOS,IRF1,JUN,KLF9,MMP7,PPARG | 463 (26) |
| SELP | | transmembrane receptor | | 0.447 | 0.0212 | CLK1,CXCL2,CXCL8,CXCR4,IL1R2 | |
| SFN | | other | | 0.447 | 0.0107 | BCL6,PDK4,PPP1R1B,SGK1,TJP1 | |
| hymecromone | | chemical drug | | 0.447 | 0.000763 | CASP9,CAV1,CD44,CXCL8,CXCR4,FAS,MCL1 | 433 (20) |
| romidepsin | | biologic drug | | 0.44 | 0.0079 | CDKN2B,CXCL8,DUSP1,EGR1,FOS,GSN,JUN,MCL1,PTGS2,RHOB | 458 (24) |
| lenalidomide | | chemical drug | | 0.437 | 0.0000803 | V1,CAV2,CD68,CEBPA,DEPP1,DKK1,DVL1,ELF3,ERN1,HIPK2,IFIH1,IFIT3,KLF4,LIMA1,MYB,NFKBIZ,NMI,PPIF,PROS1,RCAN3,REL,RNF213,ROR1,S100A16,SIPA1L2,SMAD3,WNT | 465 (13) |
| FAS | -0.581 | transmembrane receptor | | 0.431 | 0.00000025 | CXCL2,CXCL3,CXCL8,EFNA1,EGR1,F2RL1,F5,FAS,FOS,FOSB,GRK5,IL15RA,IQGAP2,IRF1,JUN,JUND,LGALS3,MAP4K4,MBOAT7,PDLIM7,PLD1,PLD2,PPFIBP2,PPP1R15A,RASA1 | 421 (17) |
| isobutylmethylxanthine | | chemical toxicant | | 0.429 | 0.0000244 | 1,ATP5MC1,CAV1,CAV2,CD44,CEBPA,CEBPD,CRABP2,CXCR4,DGAT2,DHRS3,EGR1,ELOVL6,FOS,GNPAT,IFIT3,IRF7,JUN,KLF4,LDHA,PDK4,PLAAT3,PPARG,SCARB1,SGK1,ZC | 415 (19) |
| PIM1 | 0.696 | kinase | | 0.426 | 0.00193 | BBC3,CAV2,CD44,CDC25A,EPHA2,FOSL1,ID2,PIM1,PROS1 | 461 (18) |
| tetrachlorodibenzodioxin | | chemical toxicant | | 0.421 | 0.00000435 | YP2S1,EDN1,FAS,FOS,GLI2,HDAC9,HSPA1A/HSPA1B,IGF2,IRF1,JCAD,JUN,JUND,KCNE3,KRT7,MMP15,MYB,NEDD9,NR0B2,PJA1,PRKCA,PTGS2,S100A4,SAT1,SERPINH1,SER | 545 (22) |
| SRA1 | | transcription regulator | | 0.418 | 0.00295 | CAV1,PPARG,SLC2A12,TBL1X,TGFB2 | 331 (13) |
| MED1 | | transcription regulator | | 0.415 | 0.000104 | ACAA1,AOC1,AURKA,BCL3,BCL6,CEBPA,CYP1A1,CYP24A1,CYP2B6,CYP3A5,GADD45A,ID2,KLF4,LIF,MCL1,PDK4,PEX11A,PIM1,PPARG,VDR | 456 (21) |
| PTK2 | | kinase | | 0.415 | 0.0247 | BIRC3,CCN1,CSPG4,FOS,LPP,MMP7,TJP1 | |
| WNT1 | | cytokine | | 0.406 | 0.0000266 | AXIN2,BMP4,CALB2,CEBPA,DKK1,EDN1,EGR1,FOS,HNF4A,IGF2,JUN,PPARG,PTGS2,RARG,RHOU,RUNX2,SERPINE2,STRA6,TJP1 | 498 (24) |
| gossypol | | chemical drug | | 0.404 | 7.02E-08 | ATF3,CXCL1,CXCL8,MCL1,NCOA3,PMAIP1,SAT1 | 398 (11) |
| AFP | | transporter | | 0.404 | 0.00144 | EGR1,FOS,TNFRSF12A,TSC22D1 | |
| BCL2 | | transporter | | 0.396 | 0.00714 | ABCC1,BBC3,CXCL1,CXCL8,FAS,FOS,GADD45A,IGFBP6,MCAM,MCL1,TGFB2,VDAC1 | 487 (21) |
| 1,25-dihydroxyvitamin D | | chemical drug | | 0.396 | 0.0158 | CCNG2,CXCL8,CYP24A1,EGR1,IGF2,REN,VDR | |
| nicotinic acid | | chemical - endogenous mammalian | | 0.391 | 0.0299 | ANGPT1,CD68,CEACAM1,ELOVL6,SCARB1 | |
| MMP14 | | peptidase | | 0.391 | 0.00485 | ATF3,BCAM,CSPG4,DAPK1,PTGS2,SMARCA1 | 259 (7) |
| SRSF3 | | other | | 0.391 | 0.000387 | CD44,CXCL8,E2F7,HIPK2,JUN | 359 (12) |
| carboplatin | | chemical drug | | 0.389 | 0.0003 | ABCC2,CD44,CLU,ELF3,LAMC2,LCN2,LGALS3,NT5E,PMAIP1,TNFRSF12A | 197 (7) |
| beta-carotene | | chemical - endogenous mammalian | | 0.389 | 0.00000504 | ACSS2,APOL6,ARRDC3,DGAT2,ELOVL6,HSPA1A/HSPA1B,ME1,NR1D1,PPARG,PTGS2,SCARB1 | 370 (13) |
| GFI1 | | transcription regulator | | 0.385 | 0.000000301 | BBC3,BCL3,CARD6,CEBPA,CEP95,CXCL8,CXCR4,EIF2AK2,ETS1,F2R,ID2,IRF1,JUN,MAFF,MMP7,NT5E,REL,SERPINA1,SERPINA3,SGK1,SMAD3,TFF3,TNK2,VDR | 453 (17) |
| paclitaxel | | chemical drug | | 0.382 | 3.69E-12 | CL2,CXCL8,CYP1A1,CYP3A5,DUSP5,EGR1,EMP1,EPHA2,EREG,ETV1,FAS,FOSL1,GADD45B,GDF15,HBEGF,JUN,JUND,KDM6B,MACF1,MCL1,MUC2,NT5E,PHLDA1,PTGS2,REN, | 510 (20) |
| CDKN1A | | kinase | | 0.382 | 0.00734 | RKA,BBC3,CCN1,CCNA2,CD44,CDC25A,CDKN2B,CXCL2,CXCL3,DUSP1,E2F7,FAS,GADD45A,GNAI1,HJURP,HSPA1A/HSPA1B,IFITM3,LGALS3,LGALS3BP,LIMA1,MUC2,STMN1,T | 453 (17) |
| NLRP12 | | other | | 0.381 | 7.59E-08 | BCL3,CXCL2,CXCL8,CXCR4,HLA-B,HLA-C,HLA-E,HLA-J,IL1RN,JUN,MAP3K14,PSMB8 | 449 (14) |
| CDH11 | | other | | 0.378 | 0.000319 | LAMA5,MMP15,MMP7,PIK3R1,SERPINA1,TJP1,TNK2 | 290 (7) |
| ARHGAP31 | | other | | 0.378 | 0.000928 | GADD45B,GDF15,ID2,ITGB6,JUN,SERTAD2,TGFB2 | |
| BACH1 | | transcription regulator | | 0.378 | 0.0105 | AIFM3,ATP5MC1,ATP5MC3,CXCR4,DUSP1,ME1,PDK4,PPARG | |
| PAK2 | | kinase | | 0.378 | 0.00221 | ASS1,BMP4,CKB,DUSP1,ETS1,GADD45A,HBEGF | 477 (18) |
| SAHM1 | | chemical reagent | | 0.378 | 0.00399 | ABLIM1,CD44,DDIT4,DOCK5,PPARG,RHOU,TBC1D4 | |
| CCN5 | | growth factor | | 0.377 | 0.00000118 | AXIN2,CD44,CEBPA,CLDN1,JUN,KLF4,LAMC2,PPARG,PROCR,SDC4,SMAD3,TFF3,TGFBR2 | 427 (17) |
| caffeine | | chemical drug | | 0.372 | 0.00188 | CAV1,CDC25A,CEBPA,CXCL8,EGR1,FOS,FOSB,PPARG,PRKCA,PTGS2 | 460 (21) |
| Hsp27 | | group | | 0.371 | 0.029 | CASP9,CXCL8,FBXO32,FOS,PPP1R1B,PTGS2 | |
| BRAF | | kinase | | 0.359 | 0.00022 | VR1,B3GNT5,BCAM,BIRC3,CASP9,CXCL8,EGR1,GCNT2,HSPG2,LAMC2,MCL1,MSLN,NPC1,NT5E,PSMB10,PSMB8,RND3,SERPINE2,SLC22A5,ST3GAL2,TSC22D1,TSPAN13,UG | 431 (22) |
| cyclophosphamide | | chemical drug | | 0.342 | 0.000000258 | ATF3,B2M,CD44,CLU,CYP2B6,ELF3,FAS,FOS,HLA-E,IL1R2,KITLG,LAMC2,LGALS3,PSMB8,SLC22A5,TNFRSF12A,TSPAN8,UGCG | 523 (21) |
| CDX1 | | transcription regulator | | 0.342 | 0.00358 | CLDN1,CLDN2,CLU,CTSE,DDC,FXYD3,HEPH,PTGS2 | |
| FGF10 | | growth factor | | 0.342 | 0.0174 | ANXA1,BMP4,FGFR2,LMO7,LSS,PTGS2,TSPAN8 | |
| conjugated linoleic acid | | chemical drug | | 0.342 | 0.000134 | CEBPA,CXCL8,CXCR4,DUSP1,FOS,GDF15,IL1RN,PPARG,PTGS2 | 476 (19) |
| flutamide | | chemical drug | | 0.34 | 0.00787 | CCN1,CXCL2,CYP1A1,CYP3A5,FOS,GPD1,KITLG,ME1,PSCA,REN,SERPINA5 | 385 (7) |
| butylated hydroxyanisol | | chemical toxicant | | 0.337 | 0.0165 | FOS,IRF1,IRF7,JUN,NOX1 | |
| EZH2 | | transcription regulator | | 0.337 | 0.0000411 | ISPLD2,CXCL1,CXCL2,CXCL8,CXCR4,DKK1,DUSP5,EFNA1,EGR1,EPC1,GATA6,GDF15,KLF4,LAMC2,LATS2,LCN2,MMP7,MYEOV,NCOA7,PARP12,PLAAT4,PPARG,PTGS2,RAP1 | 527 (23) |
| napabucasin | | chemical drug | | 0.333 | 0.00384 | ALCAM,ALDH1A1,B2M,CD44,CXCL8,ENG,FGFR2,KITLG,KLF4 | |
| MAPK11 | | kinase | | 0.314 | 0.0188 | CAV1,FAS,FOS,PTGS2,VDR | |
| fulvestrant | | chemical drug | | 0.313 | 4.53E-09 | NA2,CDC25A,CEBPD,CLK1,DDIT4,EGR1,F2R,FOS,GAB2,GADD45A,GJB2,ID2,IRF1,JUN,LRP8,MYB,MYO1B,NCOA3,NRP1,PMAIP1,PPARG,PTGS2,RNF126,SAPCD2,SEMA7A,SLC | 532 (24) |
| miR-124-3p (and other miRNAs w/seed AAGGCAC) | | mature microRNA | | 0.313 | 0.00103 | AK2,ARHGAP29,CAV1,CDC14B,CDK6,CEBPA,EGR1,ENDOD1,ERN1,GSN,ITPRID2,KANK1,KLHL24,NME4,PDLIM7,PGM1,RARG,SLC16A1,SLC22A5,SMCO4,SWAP70,TJP2,TRIM29 | |
| bucladesine | | chemical toxicant | | 0.313 | 4.47E-10 | 3A5,DUSP1,EGR1,ERRFI1,FAAH,FOS,FOSL1,FRRS1,FSTL3,GADD45B,GDF15,HMGCR,IGF2,IGFBP6,IRS2,JUN,KITLG,LIF,LPCAT1,MACROD1,MAT2A,MCAM,NOX1,PTGS2,PTPN2 | 515 (21) |
| BIRC3 | 1.042 | enzyme | | 0.311 | 0.00108 | BIRC3,CXCL1,CXCL8,MAP3K14,MCL1,REL | 439 (16) |
| IGF1 | | growth factor | | 0.31 | 1.82E-10 | O32,FOS,FOSB,GADD45A,GATA2,GPD1,HMGCR,HOXA5,ID2,IFITM3,IFNGR2,IGF2,IGFBP2,IGFBP6,IRS2,JDP2,JUN,KLF6,KLHL24,KLK6,LCN2,LDHA,MCL1,MDK,MYB,NOX1,PCK2,I | 598 (22) |
| bicalutamide | | chemical drug | | 0.308 | 0.000644 | CEBPA,CLU,DDIT4,DNMT3A,F5,FOS,SEC24D,SORD,SPINK1 | 389 (12) |
| ASPSCR1-TFE3 | | fusion gene/product | | 0.302 | 0.00787 | APOL2,CIB1,DEXI,DST,GDF15,HLA-E,LOC102724788/PRODH,MAFF,MYOF,SCARB1,TCN2 | |
| MAP3K14 | 0.338 | kinase | | 0.301 | 0.000148 | ADAM8,BIRC3,CD44,CXCL1,CXCL2,CXCL8,FAS,GADD45A,IL1R2,MUC2,PTGS2,RCAN1,REL | 408 (14) |
| Smad | | complex | | 0.299 | 0.00656 | CDKN2B,EGLN1,GLI2,ID2,MCL1 | |

Supplementary Table 4. Upstream regulator analysis based gene set enrichment analysis (GSEA) of differentially expressed genes (DEGs) after ATRA+PDT compared to PDT

| Upstream Regulator | Expr Log Ratio | Molecule Type | Predicted Activation State | Activation z-score | p-value of overlap | Target Molecules in Dataset | Mechanistic Network |
|---|---|---|---|---|---|---|---|
| TMPRSS2-ERG | | fusion gene/product | | 0.293 | 3.23E-09 | RDC4,BBC3,CALB2,CARD6,CHAC1,DDIT4,DHRS3,DNAJA4,EREG,GDF15,HCAR1,IFI44,IGFBP6,KLF9,MEGF6,MSLN,MYEOV,PCK2,PHLDA1,PIM1,PPARG,PSPH,ROR1,SAMD9,USF | |
| TRPV4 | | ion channel | | 0.283 | 0.0319 | AQP5,CXCL2,FAS,FOS | |
| mevalonic acid | | chemical - endogenous mammalian | | 0.279 | 0.00221 | CD55,CYP2B6,GNAI1,HMGCR,PTGS2,SCARB1,SMAD3 | 438 (15) |
| PHLPP1 | 0.446 | enzyme | | 0.277 | 0.00183 | CEBPD,CXCL2,CXCL3,MCL1,PRKCA | 471 (24) |
| BAP1 | | peptidase | | 0.277 | 0.00183 | ATF3,CCN1,CXCL2,LATS2,LIF | |
| AZGP1 | -0.672 | transporter | | 0.277 | 0.0078 | AZGP1,BMP4,EGR1,FBXO32,PNPLA2 | 216 (7) |
| KRAS | | enzyme | | 0.275 | 6.87E-25 | F2,IGFBP2,IQGAP2,IRF1,ITGB6,ITM2B,JUN,KLF6,LAMA5,LAMC2,LATS2,LCN2,LDHA,LGALS3,LIF,LIPG,LMO7,LRP8,LTBP4,MACC1,MAP1LC3B,MCAM,MMP7,MSLN,MUC20,NOX1, | 470 (25) |
| INS | | other | | 0.271 | 0.000135 | AMOTL2,CCN1,CPT1A,CRABP2,DUSP1,EDN1,EGR1,ERRFI1,FAS,FOS,FOSB,FOSL1,HMGCR,IGFBP2,IRS2,JUN,JUND,PPARG,PTGS2,UGCG | 457 (22) |
| S-nitrosoglutathione | | chemical toxicant | | 0.266 | 0.0158 | BIRC3,IRS2,LIF,PPARG,PTGS2,S100A10 | |
| 2-bromoethylamine | | chemical reagent | | 0.266 | 0.0000195 | ABCC2,ANXA1,CD44,CLU,ELF3,LAMC2,LCN2,LGALS3,S100A10,TNFRSF12A,TSPAN8 | |
| SPP1 | | cytokine | | 0.266 | 0.0000132 | ANGPT1,AURKA,CCL15,CD44,CDC25A,CXCL1,CXCL2,CXCL3,CXCL8,FGFR2,FOS,HMGCR,INSIG1,KRT20,LIF,MMP7,NDRG2,PIK3R1,PPARG,PTGS2,RUNX2,S100A4,SC5D,TGFBR2 | 427 (19) |
| IL2RG | | transmembrane receptor | | 0.263 | 0.0051 | BCL6,CXXC5,ETS1,FOS,IRF7,JUN,NEDD4L,TXK | 451 (23) |
| phosphate | | chemical - endogenous mammalian | | 0.262 | 0.00131 | ACAT1,AMBP,CYP24A1,DNMT3A,ECM1,EGR1,F2R,FGFR2,FOSL1,LDHA,RUNX2,SDF2L1,ZFHX3 | 197 (13) |
| allopurinol | | chemical drug | | 0.261 | 0.00000115 | ABCC2,ANXA1,BIRC3,CD44,CLU,ELF3,LAMC2,LCN2,LGALS3,PSMB10,PSMB8,PTGS2,RCN1,REN,S100A10,TNFRSF12A,TSPAN8,TXNIP | 463 (21) |
| OGA | | enzyme | | 0.258 | 9.67E-14 | 5,CXCL16,CXCL8,CYP2B6,DPEP1,DST,EPHB6,FAS,FLNB,GADD45A,GALNT6,GFOD1,GNE,GSN,HIPK2,HIVEP2,HMGCR,HPGD,IFIT3,IFNGR2,IGF2,JDP2,JUN,LIF,LRIG1,LSS,MCAM | 460 (19) |
| GATA4 | | transcription regulator | | 0.256 | 0.00019 | KN2B,CLDN1,CLDN2,CRABP2,CXCL2,CXCR4,EDN1,FOS,GATA6,GRHL3,HK1,IGFBP6,JUN,KRT7,LCN2,LDHA,NR5A2,OVOL1,PML,PPL,PTGS2,PTPRB,S100A4,SERPINA3,SERPIN | 179 (7) |
| PTAFR | | G-protein coupled receptor | | 0.249 | 0.0207 | CXCL2,CXCL8,MCAM,PTGS2 | |
| PTHLH | | other | | 0.248 | 0.00215 | ANGPT1,CDKN2B,CXCL8,ECM1,FOS,HSPA1A/HSPA1B,PTGS2,RGS2,RUNX2,SMPD3,VDR | 419 (24) |
| SP3 | | transcription regulator | | 0.247 | 0.000000079 | CAM1,CEBPA,DNMT3A,DUSP1,EGR1,EIF2AK2,EREG,F2R,FAS,FOS,FOSL1,GDF15,IGF2,JUN,KLF4,MAOB,MAT2A,MUC2,NR5A2,PADI1,PROCR,PTGS2,SCARB1,SERPINH1,SGK1, | 503 (18) |
| 1L-6-hydroxymethyl-chiro-inositol 2-(R)-2-O-methyl-3-O-octadecylcarbonate | | chemical - kinase inhibitor | | 0.246 | 0.000101 | CXCL16,GDF15,MT2A,PPARG,PTGS2 | 427 (17) |
| EHMT1 | | transcription regulator | | 0.243 | 0.0000249 | ACSL1,ALDH2,CEBPD,EFNA1,GFOD1,ITPRIP,LCN2,LOC102724788/PRODH,MSLN,PCSK6,PPL,PRR15L,RARRES1,SAT1,SLC7A7,STRA6,TJP3 | |
| clenbuterol | | chemical drug | | 0.243 | 0.00106 | CXCL1,FBXO32,IL1RN,SLC16A1,VDR | |
| TCF7L1 | | transcription regulator | | 0.243 | 0.00295 | ATF3,AXIN2,JUN,KLF4,NR5A2,PPARG | 321 (9) |
| levodopa | | chemical - endogenous mammalian | | 0.241 | 0.0000475 | HBEGF,IGFBP2,ITPKA,KLF6,KLF9,LAT2,LPP,MAT2A,MEIS2,MPP6,MSMO1,MXD1,MYO1B,MYORG,NEDD4L,NT5E,ONECUT2,PADI2,PGM2L1,PGPEP1,PIK3R1,PITPNM3,PLEKHA2, | 222 (12) |
| TSC22D3 | 0.927 | transcription regulator | | 0.239 | 0.000748 | ANXA1,BBC3,CEBPA,CXCL8,DUSP1,PPARG,PTGS2,RUNX2,SGK1 | 473 (25) |
| ZBTB17 | | transcription regulator | | 0.235 | 0.0475 | CCNA2,CDKN2B,EGR1,JUN,KDM8,PMAIP1,TNFAIP2,ZFP36 | |
| PARP1 | | enzyme | | 0.224 | 0.0392 | CASP4,CXCL1,CXCL3,FAS,FOS,GDF15,HRH1,JUN,LGALS3BP,LIF,PTGS2 | |
| MAPK7 | | kinase | | 0.22 | 3.02E-08 | ACSS2,CXCL2,CXCL3,CXCL8,DUSP1,EDN1,ELOVL6,IDI1,INSIG1,JUN,KLF4,LSS,MCL1,MSMO1,PTGS2,SQLE,STEAP4,TRAF4 | 426 (22) |
| ETS1 | 0.464 | transcription regulator | | 0.22 | 0.00000586 | BMP4,CAV1,CD44,CDK6,CYP24A1,EGR1,ETS1,FBXL19-AS1,HPGD,HSPA1A/HSPA1B,HSPA1L,HSPA6,ID2,INSIG1,ITGB6,MCAM,MCL1,MMP7,MYB,PML,REN,RUNX2,SP100,TBXAS | 508 (22) |
| C3AR1 | | G-protein coupled receptor | | 0.218 | 0.0264 | CASP4,CD55,CXCL8,DYRK2,TGFB2,TGFBR2 | |
| chlorpromazine | | chemical drug | | 0.218 | 0.00000393 | CYP2B6,DEPP1,FOS,HMGCR,IDI1,LSS,MSMO1,SC5D,SERPINA3,SQLE,TSC22D3 | |
| indirubin | | chemical drug | | 0.218 | 0.0101 | BIRC3,CCN1,CYP1A1,JUN | |
| miR-21-5p (and other miRNAs w/seed AGCUUAU) | | mature microRNA | | 0.213 | 0.0208 | CDC25A,CDK6,FAS,GLCCI1,PIK3R1,PRRG4,TGFB2,TGFBR2 | |
| HTT | | transcription regulator | | 0.211 | 0.0177 | 31,DNAJC3,DUSP5,EGR1,ETV1,FOS,FOSL1,GRIN2D,GRK5,GSN,HBEGF,HIVEP2,HMGCR,IGF2,JUN,KLF4,KLF9,LDHA,MCAM,MEIS2,NR1D2,PFKM,PPARG,PPP1R1B,PSMB8,PSMI | |
| miR-204-5p (and other miRNAs w/seed UCCCUUU) | | mature microRNA | | 0.21 | 0.0000403 | ACVR1,ATP2B1,HMGA2,ID2,PIK3R1,PPARG,SMAD3,SOX4,SPDEF,STX1A,TGFBR2,TRPS1 | |
| ORMDL3 | | other | | 0.2 | 0.0367 | ADAM8,CASP9,CXCL8,MAP1LC3B,OAS3 | |
| ROR1 | 0.776 | kinase | | 0.2 | 0.0188 | AMOTL2,CCN1,SGK1,STMN1,TJP1 | |
| daporinad | | chemical drug | | 0.194 | 0.000234 | ALDH1A1,CD44,FAM111B,LIF,LRIG1,RCAN1,VDR,ZNF488 | 319 (7) |
| NGFR | | transmembrane receptor | | 0.192 | 0.00655 | CCNA2,FGFR2,FOS,JUN,NR0B2,RHOB | 416 (19) |
| cyclopamine | | chemical reagent | | 0.187 | 0.00457 | ANGPT1,FAS,GLI1,GLI2,PTGS2,SMPD3,TJP1 | 332 (10) |
| ALDH1A2 | | enzyme | | 0.186 | 0.00414 | ALDH1A1,ALDH1A3,ATF3,BGN,CCN1,CD44 | 290 (7) |
| NR4A2 | | ligand-dependent nuclear receptor | | 0.184 | 0.0265 | CXCL8,DDC,ETV1,KITLG,PTGS2,SLC39A8,SMAD3,TSC22D3,TSFM | |
| CNR1 | | G-protein coupled receptor | | 0.18 | 0.0141 | BMP4,CA9,CEACAM1,DKK1,DYRK2,ERRFI1,FOS,MDK,PLD1,PLSCR1,PTGS2,SCARB1,SMURF1,TJP1 | |
| glucocorticoid | | chemical drug | | 0.17 | 0.0000777 | D,CLIP2,CLU,CXCL1,CXCL2,CXCL8,CXCR4,CYP24A1,CYP2B6,CYP3A5,DUSP1,EDN1,FAS,GABRP,GPR158,IL1R2,IL1RN,LAT2,MAOA,MT2A,PPARG,PTGS2,SERPINA3,SGK1,TGF | 404 (15) |
| MMP2 | | peptidase | | 0.168 | 0.00839 | CD44,COL18A1,HMGCR,IL1RN,INSIG1,IRF7,PDK4,SCARB1,TJP1 | 474 (19) |
| STK11 | | kinase | | 0.167 | 0.0000217 | TP5MC1,ATP5MC3,CAMKK1,CCNA2,CEBPA,CYP2B6,EREG,FGFR2,FOSL1,GADD45A,GADD45B,GALM,GLI1,KLF12,LDHA,MCAM,NEDD9,NR1D1,NT5E,PAPSS1,PCK2,PGM1,PPAI | 521 (22) |
| ERG | | transcription regulator | | 0.166 | 0.000253 | AXIN2,BIRC3,CCNA2,CXCL1,CXCL8,CXCR4,ETS1,F2R,FAM174B,FLNB,FOSB,FYN,HPGD,IRS2,LAMA5,PIM1,PLPP3,SLC24A1,SOX4,SVIL,TGFBR2,TSC22D3,WNT11 | 457 (18) |
| panobinostat | | chemical drug | | 0.155 | 0.013 | BBC3,CDK6,CEBPD,MCL1,PMAIP1,PPP1R1B,PTGS2 | |
| budesonide | | chemical drug | | 0.154 | 0.0000413 | CASP9,CXCL2,CXCL8,CYP3A5,FYN,GNA14,GNAI1,JUN,KLF9,NMI,PDLIM1,PRKCA,RAP1GAP,TSC22D3 | 280 (11) |
| BCL2L1 | | other | | 0.154 | 0.0138 | CD44,CXCL1,CXCL3,CXCL8,ECM1,FAS,MCL1,PRAP1 | |
| GW501516 | | chemical drug | | 0.154 | 0.00491 | ACSL1,CPT1A,CYP24A1,GPD1,IGFBP2,IGFBP6,KLK6,LIPG,PDK4,PPL,SNCG,SULT2B1 | 198 (9) |
| SLC51A | | transporter | | 0.152 | 0.00144 | ABCC2,CYP2B6,CYP3A5,NR0B2 | |
| PMP22 | | other | | 0.152 | 0.00651 | CSPG4,FLOT1,HMGCR,JUN | |
| LATS2 | 0.349 | kinase | | 0.152 | 0.0488 | ALDH1A3,BBC3,CCN1,CYP24A1,LATS2 | |
| ACSL5 | | enzyme | | 0.152 | 0.00818 | ADAMTS6,GPRC5A,MCAM,MDK | |
| DCAF1 | | kinase | | 0.152 | 0.0363 | PPIF,TGM2,TOB1,TXNIP | |
| let-7a-5p (and other miRNAs w/seed GAGGUAG) | | mature microRNA | | 0.146 | 0.00613 | AGO4,ATAD3B,CDC25A,CDK6,DOCK5,EDN1,HMGA2,KLK10,LOXL4,OTULINL,PTGS2,RHOB,S100A4,SIGMAR1,SNAP23,SPCS3,TGFBR2,UGT8 | |
| ABCA1 | | transporter | | 0.142 | 0.00213 | ANXA1,CPT1A,CXCL3,DGAT2,HMGCR,PPARG,PTGS2,SCARB1 | 530 (25) |
| mir-33 | | microRNA | | 0.139 | 0.000672 | CDK6,CEBPA,CPT1A,HMGA2,IRS2,PPARG | 79 (4) |
| CCNC | | other | | 0.131 | 0.041 | CEBPA,FAS,PNPLA2,PPARG | |
| ADRB | | group | | 0.129 | 0.00000591 | ATF3,BCL10,CASP4,CCNA2,CD68,CDK6,CXCL2,DDIT4,ENDOD1,FAS,FOS,GADD45A,GDF15,IFIT3,INSIG1,JUN,KLF4,LSS,MSMO1,PTGS2,SLCO4A1 | 496 (23) |
| PCDH11Y | | other | | 0.128 | 0.000393 | AXIN2,BMP4,CD44,FOSL1,JUN,PTGS2 | 289 (7) |
| mir-22 | | microRNA | | 0.128 | 0.0158 | CCN1,CLIP2,FOSL1,MALAT1,RGS2,TGFBR2 | |
| ATF3 | 1.18 | transcription regulator | | 0.128 | 0.000182 | ATF3,AURKA,BBC3,BIRC3,CDC25A,CHAC1,CXCL1,CXCL16,GSN,HSPB1,JUN,MT2A,NFKBIZ,PPP1R15A,SRI | 511 (20) |
| ATF6 | | transcription regulator | | 0.113 | 0.00118 | AURKA,CPT1A,DAPK1,DNAJC3,FOS,MAP1LC3B,MCL1,NR0B2,NUCB2,RCAN1 | 428 (17) |
| CNTF | | cytokine | | 0.105 | 0.00727 | CEBPD,CXCL8,EGR1,FOS,IRF1,LIF,PPP1R1B,SERPINA3,SLC16A1,ZFP36 | 492 (21) |
| ANXA2 | | other | | 0.104 | 0.00112 | CDK6,FAS,GADD45A,JUN,PTGS2,S100A10,SESN2 | 452 (20) |
| Calcineurin protein(s) | | complex | | 0.103 | 0.000814 | AQP5,CBLB,CXCL3,CXCL8,DUSP1,EGR1,FOS,PTGS2,RCAN1,RGS2,RNF128 | 436 (21) |
| CYP19A1 | | enzyme | | 0.103 | 0.0279 | B3GNT5,CD44,CLCF1,HLA-E,JUN,NR5A2,PPARG,S100A10,SCARB1,STEAP4 | |
| MLX | | transcription regulator | | 0.094 | 0.000293 | ARRDC4,ELOVL6,IGF2,IL1RN,LGALS3BP,TXNIP | |
| ETV5 | | transcription regulator | | 0.093 | 0.000000287 | ALCAM,AQP5,CAV1,CLDN1,CLIC5,CXCR4,KRT13,KRT7,KRT80,LCN2,LGALS3,LMO7,MYB,S100A14,SCARB1,TINAGL1,TJP1,TJP2,TJP3 | 153 (6) |
| SLC9A3R1 | | other | | 0.092 | 0.0207 | ABCC2,CFTR,S100A4,SCARB1 | |
| rifampin | | chemical drug | | 0.086 | 1.71E-09 | ,ALDH1A1,ATP2B1,AXIN2,CEACAM5,CEACAM6,CEP135,CEP70,CYP1A1,CYP24A1,CYP2B6,CYP3A5,ELOVL6,GATA6,MAOB,ME1,MYB,NR0B2,PADI2,PCK2,SLC39A10,TBC1D9,TM | 480 (18) |
| PAX3-FOXO1 | | fusion gene/product | | 0.084 | 0.00509 | XA1,ARHGDIB,AURKA,CCN1,CDKN2B,CEP41,CXCR4,DAPK1,DUSP1,FGFR2,FOSL1,GPRC5A,HSPA1A/HSPA1B,IGF2,IGFBP2,MCAM,MT1X,SDCCAG8,SLC16A1,SMAD3,SPHK2,SI | 394 (7) |
| RBPJ | | transcription regulator | | 0.083 | 0.00139 | AXIN2,CD44,DUSP1,ETS1,FOS,GNAI1,IGF2,JUN,KITLG,LIF,MDK,MIB2,MUC2,PRKAR2A,REN,SOX4,TGFB2 | 451 (22) |
| SOX11 | | transcription regulator | | 0.081 | 0.00141 | ADAM9,BMP4,BTNL9,CAV1,CD58,CXCR4,EGR1,FAS,IFIH1,KLK6,LGALS9,MYO1B,PAG1,RUNX2,SKAP2,SMAD3 | |
| cinnamaldehyde | | chemical toxicant | | 0.075 | 0.00000515 | CPT1A,CXCL8,DDIT4,EGR1,FGFR2,FOS,HSPA1A/HSPA1B,INSIG1,JUND,PDK4,PNPLA2,PPARG,PPP1R10,PTGS2,SLC2A12,TXNIP | 497 (20) |
| benzyl isothiocyanate | | chemical - endogenous non-mammalian | | 0.07 | 0.0488 | FOS,MCL1,PTGS2,ZBTB10,ZBTB4 | |
| tacrolimus | | chemical drug | | 0.069 | 0.000637 | ABLIM1,ANGPT1,CALB2,CASP7,CEBPA,CXCL2,CXCL8,CXCR4,DAPK1,EDN1,EGR1,ETV1,F5,FOS,FOSB,IL1R2,JUN,MARF1,NRP1,NT5E,PTGS2,RCN1,RUNX2,TJP1,TNFRSF12A | 444 (19) |
| lovastatin | | chemical drug | | 0.066 | 0.00218 | ATF3,CCNA2,COTL1,GNPAT,HMGCR,IRF1,KLF4,LIF,MCL1,MPP6,PTGS2,RHOB,RHOU,SDC4,SMAD3,TFPI | 493 (19) |
| RARA | | ligand-dependent nuclear receptor | | 0.065 | 9.22E-14 | E2F7,EGR1,EIF5A2,FOS,FUT4,GDF15,GLI1,GPR158,HLA-B,HLA-C,HLA-J,HOXA3,HOXA5,IFI44,IGFBP6,IRF1,JUN,KLF4,KRT20,LGALS3BP,MAOB,MUC20,MXD4,NCOA3,NEDD9,NRI | 444 (22) |
| brefeldin A | | chemical - endogenous non-mammalian | | 0.063 | 0.000185 | BBC3,CD44,CXCL3,GADD45B,HLA-B,HLA-C,KLF4,PTGS2,SESN2,SLC38A5 | 479 (22) |
| NANOG | | transcription regulator | | 0.061 | 0.00437 | AMER1,CDC25A,CDK6,EGR1,FOS,FOSB,GATA6,GLI1,HSPA1A/HSPA1B,IER5L,KLF4,NRIP1,TOB1,ZFHX3 | 401 (10) |
| dexamethasone | | chemical drug | | 0.06 | 3.66E-33 | DEPP1,DEXI,DGAT2,DNAJC3,DST,DUSP1,DUSP16,DUSP5,DYRK2,EDN1,EFNA1,EGR1,EHF,ELOVL6,EREG,ERRFI1,F2RL1,F5,FBXO32,FLNB,FOS,FOSB,FOSL1,FSTL3,GAB1,GAI | 559 (23) |
| CD28 | | transmembrane receptor | | 0.058 | 0.00148 | B,CDK6,CLK1,CXCL2,CXCL8,CXCR4,FOS,FYN,GLI1,HSPA1A/HSPA1B,IGFBP6,IL15RA,IRF1,JUN,LDHA,MSLN,PRDX4,PTGS2,REL,RNF128,SEC61B,SLC16A3,SRPRB,STEAP2,TA | 363 (17) |
| MAP3K7 | | kinase | | 0.057 | 0.000444 | BIRC3,CXCL2,CXCL8,IFI44,IFIH1,IFIT3,IRF1,IRF7,JUN,MUC2,PTGS2,RGS2 | 293 (18) |
| dextran sulfate | | chemical drug | | 0.051 | 0.000217 | A,BMP4,CCNA2,CCNF,COL18A1,CXCL2,CXCL3,CXCL8,E2F7,ENG,ERCC6L,FANCF,FIGNL1,HJURP,IFITM3,IRF7,NOX1,NRP1,PTGS2,REN,SOX12,STEAP3,STMN1,TGFB2,TGFBR | 464 (18) |
| beta-estradiol | | chemical - endogenous mammalian | | 0.035 | 8.45E-46 | ECE2,EDN1,EFNA1,EGR1,ELF3,EMP1,EREG,ERRFI1,ETS1,ETV1,F2R,F2RL1,F5,FAAH,FAM162A,FAM3D,FAS,FEZ2,FGFR2,FHL2,FLNB,FLOT1,FOS,FOSB,FOSL1,FRMD4B,FUT8,F | 636 (22) |
| CEBPA | -0.415 | transcription regulator | | 0.028 | 8.68E-11 | CXCR4,CYP2B6,CYP3A5,DGAT2,EEF1A2,FOS,GADD45A,GATA2,GATA6,GLI1,GRHL3,H1-10,HLA-B,HMGCR,HPGD,ID2,IL1RN,IRS2,JUN,KLF4,KRT7,LCN2,MT2A,NR5A2,NRP1,OVC | 516 (22) |
| cyclosporin A | | biologic drug | | 0.027 | 1.32E-09 | XCR4,CYP3A5,DDIT4,DEXI,EDN1,EGR1,ELF3,F5,FAS,FOS,FOSB,FYN,HMGCR,HNF4A,IQGAP2,ITGB6,ITPR2,JUN,LAMC2,LCN2,LGALS3,MARF1,MUC2,PRKCA,PSMB10,PSMB8,PT | 503 (19) |
| MEN1 | | transcription regulator | | 0.027 | 0.00846 | CDKN2B,FOS,GLI1,IGFBP2,IRS2,RUNX2,TGFBR2 | 426 (19) |
| cobalt chloride | | chemical reagent | | 0.021 | 0.0000147 | ABCB10,BBC3,CA9,CLU,COL18A1,CXCL8,CXCR4,CYP1A1,DDIT4,DUSP1,MCL1,PTGS2,SORL1,TJP1,VDAC1 | 430 (18) |
| TO-901317 | | chemical reagent | | 0.019 | 0.000017 | 2,ASS1,CASP4,CASP7,CPT1A,CYP3A5,DDC,ELOVL6,GPD1,HMGCS2,LGALS3,LIPG,MRPS2,NR0B2,NR1D1,NR5A2,PPARG,PSMB8,PSME1,PTGS2,PXMP2,SCARB1,SERPINA3,SIG | 497 (22) |
| miR-29b-3p (and other miRNAs w/seed AGCACCA) | | mature microRNA | | 0.004 | 0.00119 | CAV2,CDK6,DNMT3A,DUSP2,INSIG1,KLF4,LOXL4,MCL1,PIK3R1,TGFBR2,TUBB2A,ZFP36L1 | |
| RXRA | | ligand-dependent nuclear receptor | | 0.003 | 9.54E-11 | IA,CRABP2,CTSS,CYP24A1,CYP2B6,CYP3A5,ECM1,ENG,FGF19,FLRT3,FOS,GPD1,HBEGF,HMGCS2,HNF4A,IGFBP6,INSIG1,KLF9,LCN2,MAOB,NEDD9,NR0B2,NR1D1,NR5A2,PCI | 571 (27) |
| medroxyprogesterone acetate | | chemical drug | | 0 | 3.13E-09 | 068,CDK6,CDKN2B,CEBPA,CKB,CLDN1,CXCR4,CYP1A1,DKK1,ETS1,F2R,FAS,FLNB,FOSL1,FUT8,FYN,GLI1,GSN,HES6,ID2,IFI35,IGF2,IQGAP2,JUN,LGMN,LIF,MAOA,MAOB,MMP | 553 (18) |
| puromycin | | chemical - endogenous non-mammalian | | 0 | 0.0148 | ABCC1,CYP1A1,CYP2B6,CYP3A5 | |
| FKBP4 | | enzyme | | 0 | 0.00106 | CEACAM1,IL15RA,LIF,PDK4,PPARG | 218 (7) |
| EGLN1 | 0.381 | enzyme | | 0 | 0.00473 | AURKA,CDKN2B,CXCR4,EDN1,JAG2,PTGS2,SDC4,TGFB2,TGFBR2 | 460 (24) |
| N(G)-monomethyl-D-arginine | | chemical - endogenous mammalian | | 0 | 0.000503 | ACO1,CXCL3,FAS,FOS,GATA2,PMAIP1,PTGS2 | 395 (17) |
| S-nitroso-N-acetyl-DL-penicillamine | | chemical reagent | | 0 | 0.0268 | AQP5,CXCL8,FAS,HSPA1A/HSPA1B,PMAIP1,PTGS2,S100A10 | |
| CREB3L3 | | transcription regulator | | 0 | 0.041 | CYP2B6,IGFBP2,RUNX2,SMURF1 | |

Supplementary Table 4. Upstream regulator analysis based gene set enrichment analysis (GSEA) of differentially expressed genes (DEGs) after ATRA+PDT compared to PDT

| Upstream Regulator | Expr Log Ratio | Molecule Type | Predicted Activation State | Activation z-score | p-value of overlap | Target Molecules in Dataset | Mechanistic Network |
|---|---|---|---|---|---|---|---|
| 1810019D21Rik | | other | | 0 | 0.00782 | APOL6,ASS1,EDN1,F5,GPR87,NIPAL1,PADI1,TCEAL1 | |
| EIF4G2 | | translation regulator | | 0 | 0.000518 | EGLN1,IGF2,JUN,KLF4,PTGS2,SERPINE2 | |
| DMD | | other | | 0 | 0.0106 | BGN,CNKSR1,CTSS,FBXO32,GPD1,IGF2,LIF,MAOB,NDRG2,NQO2,PADI2,PDK4,PFKM,PLPP3,S100A10,SERPINA1,SLC9A7,STARD10,TBC1D4 | |
| miR-7a-5p (and other miRNAs w/seed GGAAGAC) | | mature microRNA | | 0 | 0.0207 | FOS,IRS2,KLF4,VDAC1 | |
| LPL | | enzyme | | 0 | 0.0158 | CPT1A,CXCL8,IRS2,PDK4,PNPLA2,PPARG,SCARB1 | |
| SS18 | | transcription regulator | | 0 | 0.00754 | FGFR2,GADD45B,IGF2,JAG2,RAB40B,SLC16A3 | |
| CBX5 | | transcription regulator | | 0 | 1.43E-18 | AM1,CEMIP,ERAP2,GASK1B,GRHL3,KRT20,KRT80,LCN2,MMP7,MUC13,MUC20,PLAAT4,PPARG,PPFIBP2,PRR15L,PSMB8,QPCT,RUNX2,SBSPON,SERPINA3,SLC7A7,SNTB1,SY | |
| HOXC8 | | transcription regulator | | 0 | 0.0188 | ASS1,IL1R2,MSLN,REN,SLC16A3 | |
| CEACAM1 | 0.309 | transporter | | 0 | 0.0126 | ALDH1A1,ANGPT1,CAPN9,COL18A1,CXCL8,CXCR4 | |
| NTRK1 | | kinase | | 0 | 0.00546 | EGR1,FOS,GADD45A,IGF2,TSC22D1 | 441 (22) |
| RORC | | ligand-dependent nuclear receptor | | 0 | 0.0116 | ATP1B1,CXCL2,CXCL3,CYP2B6,CYP3A5,DDC,ELOVL6,GADD45B,HMGCR,HPGD,KCNK5,LCN2,PROCR,REL,USP53 | |
| SRD5A1 | | enzyme | | 0 | 0.00108 | CLDN1,CLDN2,HMGCR,IRS2,LCN2,PNPLA2 | |
| KRIT1 | | other | | 0 | 0.00207 | CXCR4,JUN,KLF4,TGFB2 | |
| RPL22 | | translation regulator | | 0 | 0.00287 | ATF3,BBC3,DNAJB9,DNAJC3 | 324 (11) |
| DDX5 | | enzyme | | 0 | 0.0126 | ATP5MC1,ATP5MC3,BBC3,FOS,FOSL1,JUN,LCN2,S100A4 | |
| PCYT1A | | enzyme | | 0 | 0.0319 | CXCL2,GSN,KLF4,SIX4 | |
| AQP1 | | transporter | | 0 | 0.00651 | AK2,HSPA4L,HSPH1,LCN2 | |
| COLQ | | other | | 0 | 0.0193 | BGN,BMP4,ECM1,GPC4,HSPG2,LTBP3,LTBP4,SDC4 | |
| vanillin | | chemical - endogenous non-mammalian | | 0 | 0.000859 | DDIT4,FGFR2,HSPA1A/HSPA1B,INSIG1,JUND,PPP1R10 | |
| amlodipine | | chemical drug | | 0 | 0.000778 | EDN1,HMGCR,PTGS2,REN,S100A10 | 301 (7) |
| sea cucumber body wall meal | | chemical reagent | | 0 | 0.00507 | CPT1A,HMGCR,PPARG,SCARB1 | |
| putrescine | | chemical - endogenous mammalian | | 0 | 0.00207 | FOS,IGFBP2,JUN,MAT2A | 276 (11) |
| stearic acid | | chemical - endogenous mammalian | | 0 | 0.00295 | ATF3,CLU,EDN1,F2RL1,MCL1,PTGS2 | 434 (16) |
| MTORC1 | | complex | | 0 | 0.0443 | HMGCR,IDI1,INSIG1,KLF4,LDHA,PTGS2 | |
| atorvastatin | | chemical drug | | -0.007 | 1.14E-08 | /1,CCN1,CD55,CXCL8,CXCR4,CYP2B6,DKK1,DUSP1,EDN1,EGR1,ELF3,ENG,F2R,FGF19,FOS,FOSB,GADD45B,HMGCR,IDI1,JUN,LIPG,LSS,MSMO1,NOX1,NR0B2,PPARG,PTGS2, | 482 (18) |
| SNAI2 | | transcription regulator | | -0.014 | 0.00427 | BBC3,CCN1,CLDN1,CXCR4,DKK1,HPGD,RUNX2,TRPS1,VDR | 367 (14) |
| CD3 | | complex | | -0.015 | 0.00136 | PP4,DUSP2,EEF1A2,ETS1,FAS,FOS,FYN,GLI1,GPRC5A,HSPA1A/HSPA1B,IFI35,IGFBP6,IL15RA,IRF1,JUN,MSLN,NME4,PPARG,PRDX4,PTGS2,REL,RNF128,SCO1,SEC61B,SORL | 456 (18) |
| LEP | | growth factor | | -0.015 | 0.000000785 | 8,CEBPA,CPT1A,CXCL8,CYP24A1,CYP3A5,EDN1,EGR1,ELOVL6,FAAH,FAS,FGFR2,FOS,GADD45A,GNPAT,HGD,HMGCR,IGFBP2,IL1R2,IL1RN,IRS2,JUN,JUND,LIF,MAP1LC3B,M | 472 (24) |
| diethylstilbestrol | | chemical drug | | -0.016 | 0.000000218 | N8,CLU,CRABP2,CYP3A5,DNMT3A,EGR1,EMP1,EREG,FAS,FOS,GABRP,GADD45A,GATA6,GJB2,HBEGF,IGFBP2,IGFBP6,IL1R2,KLF4,KRT13,MAOA,MXD4,NDRG2,NPDC1,PIM1,P | 515 (22) |
| N-Ac-Leu-Leu-norleucinal | | chemical - protease inhibitor | | -0.017 | 0.00000139 | CASP9,CAV1,CDC25A,CEBPA,CXCL8,HLA-B,HLA-C,ITPR2,NEDD9,PRKCA,PTGS2,RARG,RUNX2,SCARB1,STMN1,TGFBR2 | 540 (23) |
| GATA6 | 0.363 | transcription regulator | | -0.029 | 0.0161 | ACO1,AQP5,BCAM,BMP4,CAV1,CRABP2,DPP4,EDN1,GATA6,GRHL3,HNF4A,KRT7,NR5A2,OVOL1,PPARG,PPL,SUCLA2,TGFB2 | |
| nickel chloride | | chemical toxicant | | -0.029 | 0.00387 | JUN,PHLPP1,PLK3,PTGS2 | 370 (12) |
| EPAS1 | | transcription regulator | | -0.034 | 0.0000312 | 2,BBC3,BIRC3,CA9,CAV1,CCN1,CEBPA,CEMIP,CKB,CLDN1,CXCL2,CXCR4,DGAT2,EDN1,FAM13A,FOS,GADD45B,HOXA5,IRS2,LDHA,MAFF,NT5E,PKIB,PPARG,PRKCA,TGFBR2, | 507 (25) |
| IL10 | | cytokine | | -0.036 | 0.0000635 | 16,CXCL2,CXCL3,CXCL8,CXCR4,DEPP1,DUSP1,EHF,FAAH,FAS,FOS,HDAC9,HPGD,IFNGR2,IL1R2,IL1RN,INCENP,IRF1,JUN,KITLG,LGMN,LIF,MCL1,MUC2,NEDD9,NT5E,PTGS2,F | 381 (16) |
| pilocarpine | | chemical drug | | -0.061 | 0.00295 | AQP5,EDN1,FOS,FOSB,LGALS3 | 390 (17) |
| salvin | | chemical toxicant | | -0.064 | 0.0176 | CXCL8,CYP2B6,EGR1,VDR | |
| PTGS1 | | enzyme | | -0.065 | 0.0196 | ANGPT1,ATF3,CFTR,EDN1,KLK6,PTGS2 | |
| D-glucose | | chemical - endogenous mammalian | | -0.066 | 0.000000526 | 1,EGR1,ELF3,ELOVL6,FAS,FBXO32,FOS,GDF15,GPD1,GSN,HK1,HMGCR,HNF4A,HSPB1,IGF2,IRS2,ITPR2,JUN,LDHA,LOC102724788/PRODH,MCAM,MCL1,NPNT,PCK2,PFKM,PG | 536 (19) |
| piceatannol | | chemical - endogenous non-mammalian | | -0.068 | 0.00656 | BCL6,EGLN1,FAS,PMAIP1,PTGS2 | 467 (22) |
| zVAD-FMK | | chemical - protease inhibitor | | -0.068 | 0.0106 | BBC3,BIRC3,CDK6,CXCL3,FOS,MAP1LC3B,MCL1 | |
| PRKAA2 | | kinase | | -0.069 | 0.00000673 | ARHGDIB,CAMKK1,CEBPA,CYP2B6,CYP3A5,EGR1,ERN1,FGFR2,GALM,IFI44,LDHA,NDRG2,PPARG,PPP1R3B,SLC16A1,SMAD3,SORL1,TGM2,ZFP36L1,ZYX | 482 (20) |
| mir-373 | | microRNA | | -0.069 | 0.00287 | CD44,CXCL8,LATS2,TGFBR2 | 279 (11) |
| CD44 | -0.402 | other | | -0.074 | 0.000000471 | DAM8,ANXA11,BCAM,BGN,BIRC3,CD44,CLU,COL18A1,CRABP2,ECM1,FAS,GADD45A,IFITM2,IL1R2,IL1RN,JUN,LGALS3,LIMS2,MCL1,MMP7,MYO1B,NPNT,PLPP3,SDC4,SERPINA | 470 (16) |
| SNAI1 | | transcription regulator | | -0.074 | 3.65E-08 | 3MP4,CCN1,CD55,CLDN1,CLDN12,CXCL8,DKK1,FLNB,FOS,FOSL1,GDF15,GLI1,GLI2,GSN,HNF4A,HPGD,LIF,LIMA1,PTGS2,RUNX2,SERPINH1,SLC28A3,SMAD3,SVIL,TGFBR2,VD | 508 (21) |
| puromycin aminonucleoside | | chemical reagent | | -0.074 | 0.00000642 | ABCC2,ANXA1,CD44,CLIC5,CLU,CYP2B6,CYP3A5,ELF3,LAMC2,LCN2,LGALS3,PSMB10,PSMB8,RCN1,S100A10,SCARB1,TNFRSF12A,TSPAN8 | |
| bexarotene | | chemical drug | | -0.083 | 0.0000293 | AKR1C1/AKR1C2,AKR1C3,ALCAM,AMOTL2,CCN1,DDIT4,DEPP1,DHRS3,DKK1,DUSP1,DUSP5,EPHA2,GUK1,IGFBP6,ITGB6,MAFF,MALAT1,PIK3R1,PPARG,PTGS2,TGFB2,TGM2, | 500 (24) |
| RUNX3 | | transcription regulator | | -0.087 | 0.00407 | ABCC1,ABCC2,BBC3,CCN1,CEBPA,CXCL8,DIP2C,GLI2,GRK5,MACF1,MID1,PTPN21,TGFB2,UGCG | 418 (15) |
| NFE2L2 | | transcription regulator | | -0.091 | 0.000843 | XCL3,CXCL8,DNAJC3,ENTPD5,ETV6,GNA14,GSTA4,HIPK2,IFIH1,IFNGR2,IL1R2,IL1RN,IMPDH1,MAFF,MAP1LC3B,ME1,MSMO1,NR0B2,NSMF,NUCB2,PPARG,PTGS2,PTPRB,SAT1 | 391 (17) |
| FSH | | complex | | -0.093 | 2.28E-09 | APK1,DHRS3,DNAJB9,DUSP1,EFNA1,EREG,FOS,FOSB,FOSL1,GATA6,GPRC5A,GRK5,IL1RN,JUN,KITLG,MAP1LC3B,MAP4K4,MCL1,MSMO1,MT1X,MT2A,NEO1,PCSK6,PMAIP1,F | 399 (23) |
| mevastatin | | chemical drug | | -0.105 | 0.00651 | BMP4,HMGCR,KLF4,PTGS2 | 250 (7) |
| TP63 | | transcription regulator | | -0.112 | 4.67E-14 | GFR2,FOS,FOSL1,GADD45A,GLB1L2,GNAI1,GRHL3,HBEGF,HMGA2,IGFBP2,IGFBP6,JAG2,JUN,KLF6,KRT20,KRT7,LOC102724788/PRODH,MAFF,MAP4K4,MYB,NT5E,PI3,PMAIP1, | 522 (21) |
| ROCK2 | | kinase | | -0.113 | 0.000156 | BCL6,CD44,CEBPA,CXCL8,FAS,PI3,PPARG,PPL,SCEL,TOB1 | 431 (18) |
| estrogen | | chemical drug | | -0.121 | 3.45E-09 | B,CLU,COL18A1,DPEP1,DUSP1,EGR1,FAS,FOS,GJB2,GRIN2D,HBEGF,HMGCR,HSPH1,IGF2,IGFBP2,IGFBP6,JUN,KLK10,LCN2,LIF,MYB,NCOA3,NRIP1,PIM1,PTGS2,SCARB1,SE | 568 (23) |
| K+ | | chemical - endogenous mammalian | | -0.126 | 0.029 | ATF3,EGR1,FOS,JUN,REN,SLC12A4 | |
| DIO2 | | enzyme | | -0.128 | 0.0472 | ALDH1A1,ATF3,B3GNT5,CIRBP,DDIT4,EGR1,FOS,FOSB,HSPA1A/HSPA1B,LIPG,SBNO2,SEMA7A,SLC23A3 | |
| roscovitine | | chemical drug | | -0.128 | 0.0264 | CD44,CDC25A,CEBPA,CYP2B6,MCL1,PPARG | |
| FBN1 | | other | | -0.13 | 0.00295 | CCN1,LTBP3,PPARG,RUNX2,TGFBR2 | 513 (16) |
| 1,1-bis(3'-indolyl)-1-(4-trifluoromethyl-phenyl)methane | | chemical reagent | | -0.132 | 0.000593 | ATF3,CAV1,CAV2,GDF15 | 209 (7) |
| NR0B2 | 1.085 | ligand-dependent nuclear receptor | | -0.152 | 0.0263 | CPT1A,CYP2B6,EGR1,HMGCR,HNF4A,NR0B2,PCK2,PDK4,PPARG,SCARB1 | |
| RASSF5 | | other | | -0.152 | 0.0461 | CIRBP,DNAJA1,MDK,SAT1 | |
| EIF2S1 | | translation regulator | | -0.152 | 0.000256 | ATP2A3,CA9,CHAC1,EIF2AK2,GADD45A,MAP1LC3B,MCL1,PHLDA1,PPP1R15A,SESN2 | |
| SPRY2 | | other | | -0.152 | 0.0368 | AXIN2,ETS1,ETV1,FOS,ID2 | |
| AXIN1 | | other | | -0.152 | 0.0176 | CEBPA,CSRNP1,FOS,PPARG | |
| ACVRL1 | | kinase | | -0.152 | 0.0016 | CXCL8,CXCR4,EFNA1,ENG,GDF15,ID2,ZYX | 309 (7) |
| mir-25 | | microRNA | | -0.156 | 0.0446 | BBC3,GADD45A,KITLG,MCL1,TOB1 | |
| FGF8 | | growth factor | | -0.156 | 0.00675 | BMP4,CMTM3,COL18A1,DKK1,ERRFI1,FGFR2,MDK,NDRG2,RUNX2,TRIP6,VDR | 285 (12) |
| GW9662 | | chemical reagent | | -0.16 | 0.000000398 | AGR2,CAV1,CEACAM1,CEACAM5,CEACAM6,CEBPA,EGLN1,FOS,GDF15,INSIG1,IRS2,KLF4,KLK6,KRT20,LCN2,LOC102724788/PRODH,NR1D1,PPARG,PTGS2 | 493 (22) |
| cytochalasin D | | chemical toxicant | | -0.16 | 0.0346 | AMOTL2,AQP5,CXCL8,FOS,MMP15,PTGS2 | |
| aldosterone | | chemical - endogenous mammalian | | -0.163 | 0.000174 | ATP1B1,CD68,DUSP1,EDN1,EGR1,FOS,IRS2,LCN2,LGALS3,NDRG2,NOX1,PPARG,PROCR,PSCA,PTGS2,RGS2,SGK1,TBC1D4,TSC22D3,WNK4 | 468 (22) |
| androgen | | chemical drug | | -0.17 | 0.0000011 | AGR2,AKR1C3,AXIN2,BMP4,CASP7,CD44,CXCR4,DNMT3A,ETS1,FGFR2,FOS,FOSL1,GSN,IGFBP2,JUN,PIK3R1,PIM1,PSCA,PTGS2,RUNX2,SDC4,SMAD3,SOX4,STEAP1,VIL1 | 369 (19) |
| STAT5a/b | | group | | -0.171 | 0.000445 | BCL6,CDC25A,CLU,DUSP1,DUSP2,EMP1,F2R,FOS,GATA2,GNAZ,ID2,IDI1,IRF1,MCL1,PIM1,PTGS2 | 417 (22) |
| pioglitazone | | chemical drug | | -0.172 | 0.0196 | ABCB10,ATP2A3,CAV1,CEBPD,CXCL8,CYP2B6,EGR1,FOS,FOSL1,HPGD,IL1RN,IRS2,KLF4,NOX1,PPARG,PTGS2,SCARB1 | |
| losartan potassium | | chemical drug | | -0.188 | 0.000717 | ANGPT1,CAV1,CXCL3,CXCL8,EDN1,EGR1,ENG,FOS,GRK5,ITPR2,PTGS2,REN,SMAD3,TGFB2,TGFBR2 | 438 (20) |
| LEF1 | | transcription regulator | | -0.196 | 0.00000354 | ADAMTS6,ARRDC4,AXIN2,CASP7,CASP9,CAV1,CD44,CXCR4,DKK1,ECM1,FAS,ID2,LRIG1,MMP7,MXD1,NCOA7,NTN4,PHLDA1,PTGS2,RUNX2,SGK1,SH3BGRL2,TSC22D1 | 461 (18) |
| SHC1 | | other | | -0.2 | 0.0484 | CXCL1,DKK1,EGR1,FOS,KRT7,PHLDA1,SDC4 | |
| FADD | | other | | -0.202 | 6.9E-09 | ASB13,BIRC3,CASP7,CDK6,CXCL2,CXCL8,CXCR4,EGR1,EIF2AK2,FAS,FOS,GADD45B,IFIH1,IRF7,JUN,KLF6,PIM1,PPP1R15A,PSMB8,TIPARP | 427 (17) |
| DICER1 | | enzyme | | -0.213 | 0.000652 | N1,CD44,CDKN2B,COL18A1,CXCL8,CXCR4,DEPP1,EGR1,ENG,FOS,GBP3,HMGCR,IGF2,JUN,LIPG,MCAM,MT2A,PELI2,PRKCA,PROCR,PROS1,SERPINA1,SLC4A4,TGFB2,TGFBR | 477 (22) |
| TAZ | | enzyme | | -0.213 | 4.25E-09 | AMOTL2,ANXA1,ARHGAP29,CAV1,CAV2,CCN1,CD44,DUSP1,EDN1,EGLN1,F2RL1,KLF6,LDHA,MYOF,PGM1,PMAIP1,PPP1R3B,PTGS2,RUNX2,SAMD4A,TBC1D2,WNT11 | 328 (12) |
| GCS-100 | | chemical drug | | -0.218 | 0.000343 | CDK6,LGALS3,MCL1,PMAIP1 | |
| dexmedetomidine | | chemical drug | | -0.218 | 0.000557 | CXCL8,DDIT4,FOS,FOSB,PTGS2 | 466 (22) |
| indole | | chemical - endogenous mammalian | | -0.218 | 0.00651 | CXCL8,CYP1A1,TJP1,TJP3 | |
| molybdenum disulfide | | chemical reagent | | -0.218 | 0.0000047 | KR1B10,AKR1C1/AKR1C2,AKR1C3,ALDH1B1,ANXA10,CDKN2B,CKB,CXCL8,DKK1,GPC4,IFI44,LAMA5,LIMS2,MEGF6,PHLDA1,RARRES1,RCAN1,STEAP1,TINAGL1,TP53I11,TUFT | |
| NR1H3 | | ligand-dependent nuclear receptor | | -0.219 | 0.00517 | ABCA2,ATF3,AURKA,CCNA2,CD55,CD68,CXCL16,CXCL2,E2F7,GLI1,HK1,HMGCR,IL1RN,IRF7,NR0B2,NR5A2,PPARG,PTGS2,REN,SCARB1,UBC | 434 (19) |
| TFAP2C | | transcription regulator | | -0.22 | 0.00388 | ALCAM,CD44,DYRK2,F2R,HK1,MT1X,MT2A,NRP1,RASA1,SEMA3B | |
| mir-10 | | microRNA | | -0.226 | 0.00943 | ALDH1A3,CCN1,CD44,GAB2,GLI1,GTPBP2,IGF2,KLF4,MCL1,SH2B3,TBC1D1,TGFBR2 | 477 (20) |
| ITGB2 | | transmembrane receptor | | -0.239 | 0.0117 | CD44,CXCL2,CXCL3,GADD45B,PTGS2,RUNX2,TGFBR2 | |
| ATG5 | | other | | -0.239 | 0.0484 | AGR2,CASP7,CYP2B6,IFIH1,RCAN1,SCARB1,TGM2 | |
| PLAGL1 | | transcription regulator | | -0.243 | 0.000557 | KRT20,LDHA,LOC102724788/PRODH,PPARG,SORD | |
| methotrexate | | chemical drug | | -0.246 | 0.00904 | ACAA1,ACO1,ACSS2,AK4,AMD1,ASS1,CASP4,CLDN2,CLN8,CPT1A,CTSS,CXCL8,DGAT2,DUSP1,FAS,GSN,HMGCS2,PLD2,PTGS2,TOB1,ZFP36L1 | 460 (18) |
| (+)-MK-801 | | chemical drug | | -0.247 | 0.015 | ABCC2,EGR1,FOS,FOSB,HSPA1A/HSPA1B,JUN,JUND,PTGS2,RUNX2 | |
| cerivastatin | | chemical drug | | -0.256 | 0.00128 | ASS1,CXCL8,CYP2B6,ETV6,HMGCR,HSPB1,IDI1,JUN,NFYA,PPARG,PTGS2 | 430 (23) |
| linoleic acid | | chemical - endogenous mammalian | | -0.258 | 0.0261 | CLU,CPT1A,DUSP1,EDN1,NEDD4L,PTGS2,REN,SULT2B1 | |
| pirinixic acid | | chemical toxicant | | -0.262 | 7.79E-09 | YP2B6,CYP3A5,DPP4,ELOVL6,FAS,GADD45A,GADD45B,GPD1,HLA-E,HMGCR,HMGCS2,HNF4A,HSPB1,ID2,IFITM3,IL1RN,INSIG1,JAG2,LCN2,LGALS3,LGALS4,MCL1,NPC1,NR1D | 478 (23) |
| MYOC | | other | | -0.277 | 0.00119 | ANXA1,CARD10,CD44,DDIT4,EPHA2,FBXO32,JDP2,RAB27B,RUNX2,SESN2,SOSTDC1,SP100 | 390 (7) |
| alpha-tocopherol | | chemical drug | | -0.277 | 0.0078 | CXCL3,HSPA1A/HSPA1B,JUND,PMAIP1,PPARG | 491 (20) |
| SERPINE1 | | other | | -0.277 | 0.0406 | CASP9,CXCL2,CXCL8,FOS,TGFBR2 | |
| MAPT | | other | | -0.283 | 0.00282 | EIF4A2,EIF5A2,FOS,FRRS1,FYN,GNAZ,HK1,HSPA1A/HSPA1B,HSPA4L,HSPH1,IFI44,IFIT3,IFITM3,IGFBP2,IL1RN,IRF7,JUN,LGALS3BP,PARP14,PLXNB2,PPP1R1B,RNF213,SLC16 | 461 (20) |
| mono-(2-ethylhexyl)phthalate | | chemical toxicant | | -0.283 | 0.00117 | ACO1,ACSL1,CEBPA,CYP1A1,DDIT4,DGAT2,FAM98A,FAS,FOS,GPD1,HIPK2,IRS2,NOX1,PDK4,PPARG,SC5D,UQCRFS1,UQCRHL | 430 (24) |
| sirolimus | | chemical drug | | -0.287 | 0.0000463 | ,CEBPA,CXCR4,DDIT4,EGR1,EIF2AK2,FAS,FOS,GPD1,HLA-B,HMGCS2,ID2,IDI1,IGF2,IRF7,IRS2,JUN,LCN2,LDHA,LGALS3,MAP1LC3B,MCL1,MUC2,MYB,PGM1,PIK3R1,PIM1,PML | 552 (22) |
| PAX3 | | transcription regulator | | -0.293 | 0.0000377 | ANXA1,ARHGDIB,ASS1,BMP4,CLU,CXCR4,DPEP1,DUSP2,F2RL1,HSPG2,ID2,IFITM2,IGFBP2,MCAM,NEDD9,PSME1,RHOB,SOX4,TCN2,TGFB2,TIA1,TJP2,TSC22D1,URB2 | 373 (7) |
| XDH | | enzyme | | -0.294 | 0.00863 | CEBPA,CXCL3,CXCL8,CXCR4,DUSP1,PPARG | 470 (22) |
| BCL6 | 0.41 | transcription regulator | | -0.294 | 0.0142 | ALCAM,BCL3,BCL6,CD44,CDKN2AIP,CGAS,CXCL3,FAS,GADD45A,GLI1,GLI2,HDAC9,HERC6,IRF1,ITPRIP,KLF6,LPP,MYB,PFKM,SLC39A8 | |

Supplementary Table 4. Upstream regulator analysis based gene set enrichment analysis (GSEA) of differentially expressed genes (DEGs) after ATRA+PDT compared to PDT

| Upstream Regulator | Expr Log Ratio | Molecule Type | Predicted Activation State | Activation z-score | p-value of overlap | Target Molecules in Dataset | Mechanistic Network |
|---|---|---|---|---|---|---|---|
| lysophosphatidylcholine | | chemical - other | | -0.296 | 0.0000488 | CXCL1,CXCL8,EGR1,HBEGF,HMGCR,INSIG1,JUN,LSS,MSMO1,PPARG,RGS2,SCARB1,SQLE | 418 (19) |
| Gsk3 | | group | | -0.296 | 0.00176 | BBC3,CEBPA,CXCL8,FOS,FOSB,GDF15,HMGCR,JUN,MYB,PPARG,PTGS2 | 497 (22) |
| CD38 | | enzyme | | -0.299 | 0.000000309 | ATP1B1,BCL6,CD44,EGLN1,FAM162A,IL1R2,LGALS3,LMO7,MYADM,NABP1,NT5E,PDLIM1,PGM1,PIM1,PKIB,PRDX4,RAP1GAP,RBPMS,S100A4,SCEL,SDF2L1,SLC16A3,SLC39A8, | |
| Y 27632 | | chemical drug | | -0.306 | 0.00115 | AMOTL2,BBC3,BMP4,CCN1,CD44,CDKN2B,CEBPA,FOS,PPARG,PTGS2,TJP1 | 508 (26) |
| berberine | | chemical drug | | -0.31 | 0.0015 | BIRC3,CASP7,CASP9,CPT1A,CXCL8,CYP1A1,CYP2B6,CYP3A5,HMGCR,MCL1,PPARG,PTGS2,SAT1,SCARB1 | 496 (20) |
| BMP7 | | growth factor | | -0.31 | 0.0000254 | ACVR1,BMP4,CAV2,CD44,CEBPA,CEBPD,CXCL8,DKK1,EDN1,EREG,ID2,IGF2,IGFBP6,IL1RN,KLF4,PCSK6,PIK3R1,PPARG,RGS2,RUNX2,S100A4,SOSTDC1,WNT11 | 481 (23) |
| mir-148 | | microRNA | | -0.312 | 0.041 | ALCAM,EGR1,MMP15,TGFBR2 | |
| CFTR | -0.375 | ion channel | | -0.323 | 0.000215 | ACAA1,CFTR,CXCL3,CXCL8,CYP2B6,CYP3A5,FAAH,FAS,HPGD,IDI1,NFKBIZ,PTGS2,SQLE | 386 (16) |
| S100A9 | | other | | -0.325 | 0.00026 | AKAP12,ALDH1A1,ATF3,BACE2,BCL3,CXCL3,CXCL8,DDC,DDIT4,EFNA1,FAS,FOSL1,GJB2,GPRC5A,LOC102724788/PRODH,PHLDA1,PLK3,SEMA3B,SGK1,TXNIP | 489 (22) |
| STAT5A | | transcription regulator | | -0.328 | 2.97E-18 | S,FOSB,GADD45A,GASK1B,GRHL3,ID2,IFITM2,IRF1,KRT20,KRT80,LCN2,MCL1,MMP7,MUC13,MUC20,PIM1,PLAAT4,PPARG,PPFIBP2,PRR15L,PSMB8,QPCT,RAB27B,RUNX2,SB | 493 (21) |
| Hif1 | | complex | | -0.329 | 0.0109 | CA9,CCNG2,CXCR4,DDIT4,EDN1,LDHA,SLC16A1,SLC16A3,TFF3 | |
| SOX7 | | transcription regulator | | -0.333 | 0.0015 | CRABP2,EFNA1,ENG,GRHL3,HSPG2,KRT7,OVOL1,PPL,PRKCA,SOX4 | |
| FLT1 | | kinase | | -0.333 | 0.0202 | DEPP1,FOS,HK1,HLA-B,MAT2A,NRP1,SNAP23,TGFBR2,VDAC1 | |
| CBFB | | transcription regulator | | -0.342 | 0.0302 | CEBPA,ITGB6,KCNK5,LGALS3BP,LGMN,NRP1,PPARG,RUNX2,TGFBR2 | |
| hemin | | chemical - endogenous mammalian | | -0.342 | 0.0457 | ACO1,AKR1C1/AKR1C2,CD68,CXCL8,HDAC9,JUN,NFYA,PROCR,PTGS2 | |
| RASSF1 | | other | | -0.345 | 7.38E-12 | ALDH1A1,CASP4,CAV1,CAV2,CCNA2,CLU,CRABP2,CXCR4,DKK1,EGR1,EMP1,FOS,FRMD4B,GDF15,GSN,HBEGF,KLF4,LIF,MDK,MTUS1,PROS1,PTGS2,TGM2,TUBB2A | 446 (18) |
| tempol | | chemical drug | | -0.346 | 0.00984 | BMP4,CXCL2,CXCL8,NOX1,PTGS2,SGK1 | 415 (24) |
| Insulin | | group | | -0.35 | 0.000000656 | T4,DGAT2,DNAJB4,DUSP1,EDN1,EGR1,ELOVL6,ERRFI1,FBXO32,FER,FOS,FOSB,HMGCR,HNF4A,IFIT3,IGF2,IGFBP2,INSIG1,IRF7,IRS2,JUN,KLF4,LDHA,LGALS3BP,MAOA,MAO | 378 (21) |
| FOXA1 | | transcription regulator | | -0.356 | 0.00352 | AGR2,ANXA1,CD58,CFTR,COL18A1,CXCL8,DDC,DDIT4,HK1,HNF4A,HNF4G,MUC2,NRIP1,S100A2,SPDEF,TFF3,TNFRSF12A | 357 (11) |
| TGFBR2 | -0.513 | kinase | | -0.357 | 0.0000218 | N,CD44,CDKN2B,CLU,CXCL2,DYRK2,GADD45B,GJB2,GLI1,GLI2,HPGD,IL1R2,JUN,MUC2,MYOF,NRP1,PTGS2,RUNX2,SLC12A4,SMAD3,TBXAS1,TGFB2,TGFBR2,TNFRSF12A,TX | 470 (18) |
| ERBB2 | | kinase | | -0.359 | 4.93E-17 | CR,HSPA1A/HSPA1B,HSPB1,ID2,IDI1,IGF2,IGFBP6,ITGB6,ITPR2,JAG2,JUN,KLF4,KRT7,LAMC2,LCN2,LDHA,LGALS3,LPCAT1,LSS,LTBP3,MACF1,MAOA,MCL1,MMP7,MSLN,NED | 464 (22) |
| FOSB | 0.826 | transcription regulator | | -0.37 | 0.0000566 | CEBPA,FOS,FOSB,FOSL1,HPGD,MT2A,PPARG,RUNX2,SMAD3 | 347 (15) |
| ciglitazone | | chemical drug | | -0.372 | 0.000872 | CAV1,CAV2,CXCL1,CXCL8,GPD1,GSN,LCN2,MXD1,NOX1,PPARG,PTGS2,SULT2B1 | 427 (22) |
| PGR | | ligand-dependent nuclear receptor | | -0.373 | 1.22E-16 | ST,EDN1,F2RL1,FOS,GPR160,GPRC5A,HBEGF,HPGD,IFI35,IFI44,IFIT3,IRF7,IRS2,KLF4,KLF9,MAT2A,MBOAT7,MTSS1,NEDD9,NPC1,NQO2,OAS3,PDLIM1,PHF21A,PPARG,PPIF, | 566 (24) |
| IgG | | complex | | -0.375 | 9.03E-11 | ),CRABP2,CXCL8,DUSP1,EDN1,EGR1,EMP1,FOSB,GPRC5A,GUK1,HLA-E,HSPB1,IFIH1,IFITM3,IL1RN,JUND,KLK6,KRT13,LCN2,LGALS3,PIM1,PPIF,PPL,PPP1R15A,PTGS2,RND3, | 507 (22) |
| HDL-cholesterol | | complex | | -0.378 | 0.00189 | CHAC1,EGR1,IDI1,KLHL24,MSMO1,SLC30A1,SQLE | |
| PPARD | | ligand-dependent nuclear receptor | | -0.382 | 0.000848 | A1,ALDH2,ASS1,CD68,CPT1A,CYP2B6,GPD1,HMGCS2,IGFBP2,IGFBP6,KLK6,LDHA,LGALS4,LIPG,PCSK6,PDK4,PNPLA2,PPARG,PTGS2,RUNX2,SAT1,SORD,TFF3,TNFRSF12A, | 363 (19) |
| gefitinib | | chemical drug | | -0.387 | 0.000476 | ALDH1A1,AURKA,CCNF,CEBPD,CXCL2,CXCL8,EFNA1,FGFR2,FOSL1,GADD45B,GLI1,HSPB1,IL1R2,MAFF,MAP4K4,MMP15,PPARG,PTGS2,RHOB,S100A14 | 459 (23) |
| KN-62 | | chemical - kinase inhibitor | | -0.396 | 0.00000232 | AQP5,ATF3,CXCL8,CYP3A5,EGR1,HNF4A,PPARG,PPP1R15A,PTGS2 | 481 (21) |
| ADCYAP1 | | other | | -0.401 | 0.0000488 | ,BCAM,CCNA2,CEBPD,CLCF1,CXCL2,CXCL3,DNMT3A,EGR1,EPHA2,FGFR2,FOS,FRRS1,GLI1,GRK5,HK1,IRF1,MCAM,PPP1R3B,RHOB,S100A10,SCARB1,SERPINA3,SERPINE2,T | 561 (22) |
| NPPA | | other | | -0.401 | 0.0165 | DUSP1,EDN1,EGR1,MCL1,PTGS2 | |
| celecoxib | | chemical drug | | -0.405 | 0.00353 | CD68,CYP3A5,DNAJB9,FAS,FOS,GADD45A,GDF15,MCL1,MUC2,PPARG,PTGS2,ST3GAL4,TGFBR2,TRAF4 | 423 (18) |
| MYRF | | transcription regulator | | -0.427 | 0.00636 | AKR1C3,AMOTL2,CEACAM1,GDF15,LOXL4,PARP10,PDK4,RASSF6,RHOU | |
| NCOR1 | | transcription regulator | | -0.436 | 0.000000206 | AZGP1,BCL3,CCNA2,CEBPA,CYP3A5,DUSP1,FXYD3,HK1,ME1,MT1X,NR1D1,PDE9A,PDK4,PPARG,PTGS2,SORD,SPINK1,STEAP2,STK39 | 492 (20) |
| miR-291a-3p (and other miRNAs w/seed AAGUGCU) | | mature microRNA | | -0.438 | 0.0483 | ADAM9,ANKRD13B,CD44,DKK1,GBP3,LATS2,TBC1D2,TGFBR2,TRPS1 | |
| interferon alfacon-1 | | biologic drug | | -0.447 | 0.0268 | CXCL8,HIPK2,IL1R2,MAFF,PARP14 | |
| xanthohumol | | chemical drug | | -0.447 | 0.00863 | ANXA1,BIRC3,DGAT2,FOS,GAB2,RUNX2 | 193 (7) |
| SETD2 | | enzyme | | -0.447 | 0.0268 | CAV1,CCN1,ENG,JUN,RHOB | |
| HOXD10 | | transcription regulator | | -0.447 | 0.00092 | CD44,CEACAM1,DAPP1,HBEGF,MAOB,RAB40B,REN,RHOB,TFPI,TJP1 | |
| EED | | transcription regulator | | -0.447 | 0.00457 | GATA6,IGFBP2,KLF4,MMP7,PTGS2,RUNX2,SLC26A6 | 371 (17) |
| BGN | -1.301 | other | | -0.447 | 0.00919 | AXIN2,CXCL3,CXCL8,RUNX2,SMAD3 | 481 (19) |
| MED13 | | transcription regulator | | -0.447 | 0.024 | ELOVL6,GPD1,HMGCR,KITLG,LIF,PPARG | |
| SGK1 | -0.361 | kinase | | -0.447 | 0.0422 | ASIC1,CDK6,CEBPA,CFTR,HBEGF,LIF,PPARG | |
| amiodarone | | chemical drug | | -0.447 | 0.0016 | AK4,CXCL8,CYP1A1,LSS,ME1,NDRG2,PPARG | 232 (7) |
| SB 290157 | | chemical reagent | | -0.447 | 0.00546 | BMP4,CASP4,CXCL2,CXCL3,PTGS2 | 466 (17) |
| 2-methoxyestradiol | | chemical - endogenous mammalian | | -0.452 | 0.00592 | ATF3,CAV1,CXCR4,LOC102724788/PRODH,MCL1,PTGS2,SERPINE2 | 506 (24) |
| GW3965 | | chemical reagent | | -0.453 | 0.00243 | CAV1,CCNA2,CXCL8,DGAT2,DUSP1,GNAI1,IL1RN,NR1D1,NR5A2,PDK4,PKIB,PPP1R15A,PTGS2,SC5D | 502 (25) |
| genistein | | chemical drug | | -0.456 | 0.000000126 | L3,CXCL8,CYP1A1,DKK1,DUSP1,EREG,FOS,GDF15,GJB2,GLI1,HMGCR,HPGD,HSPA1A/HSPA1B,ID2,IRF7,JUN,JUND,LGMN,MAOA,MUC2,MYB,NOX1,NPC1,PIK3R1,PIM1,PKIB,P | 591 (20) |
| ACOX1 | | enzyme | | -0.457 | 0.000000038 | ER2,ACSL1,CTSE,DPP4,GADD45A,IGFBP2,KLF6,LCN2,LGALS3,LGALS3BP,PDK4,PEX11A,PHLDA1,PLSCR1,PXMP2,QPCT,RGS2,SERPINA1,SGK1,SLC16A1,SLC4A4,TM4SF4,U | 180 (7) |
| spironolactone | | chemical drug | | -0.458 | 0.0283 | ABCC2,BMP4,CXCR4,CYP3A5,EDN1,LGALS3,PPARG,PTGS2,REN | |
| EIF4EBP1 | | translation regulator | | -0.462 | 0.00000607 | BBC3,CEBPA,CEBPD,EGR1,IL1RN,IRF7,PHLPP1,PNPLA2,PPARG,PTGS2 | 400 (12) |
| indole-3-carbinol | | chemical drug | | -0.462 | 0.0268 | ANXA1,CDK6,CYP3A5,FOS,GDF15 | |
| tributyltin | | chemical reagent | | -0.463 | 0.000043 | ACSL1,ASS1,ATP5MC1,CEBPA,DGAT2,ELOVL6,F2RL1,GPRC5A,IFIT3,IRF7,KRT80,LDHA,PDK4,PPARG,PPL,SLCO4A1,SNTB1,WNK4 | 387 (12) |
| tanespimycin | | chemical drug | | -0.473 | 0.00152 | BIRC3,CAV1,CDK6,CDKN2B,CXCL8,DNAJB1,FAS,HMGCR,HRH1,HSPA1A/HSPA1B,JUN,LDHA,MCL1,POLR2L,SERPINH1 | 503 (22) |
| resveratrol | | chemical drug | | -0.475 | 0.00000678 | 1A,CXCL2,CXCL3,CXCL8,CYP1A1,DAPK1,EDN1,EGLN1,EGR1,FAS,FOS,FOSL1,GDF15,GLI1,HMGCR,IGF2,JUN,JUND,KRT20,MALAT1,MCL1,NOX1,NRIP1,PDK4,PPARG,PPP1R15 | 463 (21) |
| GSTO1 | | enzyme | | -0.48 | 0.000000424 | CELSR1,CHAC1,FUT8,LAMA5,LMO7,MTUS1,NPDC1,OAS3,PKIB,SAMD9,SAT1,SEMA3A,SLFN5,SOX4,ZFP36L1 | |
| FGFR2 | 0.626 | kinase | | -0.489 | 0.000812 | BMP4,DKK1,FAS,FGFR2,FLRT3,GASK1B,GATA6,ID2,IGF2,NOSTRIN,NR5A2,PLXNB2,RUNX2,SPINK1,TRPS1,TSPAN8 | 473 (12) |
| ursodeoxycholic acid | | chemical - endogenous mammalian | | -0.499 | 0.00427 | ABCC2,CYP1A1,CYP2B6,ELF3,GLI1,LGALS3,PTGS2,TGFB2,VDR | 496 (14) |
| mir-21 | | microRNA | | -0.502 | 0.013 | CARD6,CASP4,CCNA2,CDC25A,CDK6,CDKN2B,CLU,CXCL2,FAS,GBP3,GLCCI1,HMGA2,IRF1,NPAS2,OAS3,PRRG4,PSME2,RHOB,SSR1,STMN1,TBC1D1,TGFBR2,TRMT10C | |
| ADORA2A | | G-protein coupled receptor | | -0.508 | 0.00839 | ATP2B1,CKB,CXCL2,DGAT2,DUSP1,FOS,GLI1,IFITM3,PMPCA,PPP1R1B,PTGS2,TJP1,TSC22D3,TUBB2A | 363 (19) |
| SRC (family) | | group | | -0.518 | 0.00454 | CYP1A1,EGR1,MAOA,MCL1,NOX1,PTGS2,S100A4,TJP1 | 464 (19) |
| caffeic acid phenethyl ester | | chemical drug | | -0.532 | 0.00241 | ABCB10,BIRC3,CD44,CXCL3,CXCL8,EGR1,HSPB1,IGFBP6,KLF6,MUC2,PLD1,PTGS2,RHOB | 441 (18) |
| ADIPOQ | | other | | -0.538 | 0.0124 | ACSL1,BCL3,CEBPA,CXCL1,CXCL2,CXCL3,CXCL8,DUSP1,EGR1,FOS,GPD1,HBEGF,HMGCS2,HNF4A,PPARG,PTGS2,RUNX2 | |
| glucagon | | biologic drug | | -0.552 | 0.0017 | ESAM,FOS,IRS2,LIMA1,NDRG2,PPP1R1B,RAP1GAP,SEC61B,TJP2 | |
| DL-fructose | | chemical - endogenous mammalian | | -0.555 | 0.0241 | BBC3,ELOVL6,GPD1,TXNIP | |
| SIAH2 | | transcription regulator | | -0.562 | 0.00000888 | AKR1C3,ANGPT1,CYP24A1,EGLN1,FXYD3,MT1X,PDE9A,PLK3,STEAP2 | 422 (11) |
| CITED2 | | transcription regulator | | -0.562 | 0.0196 | CDKN2B,CXCL1,CXCL8,DUSP1,GATA6,TGM2 | |
| GW7647 | | chemical drug | | -0.57 | 0.000193 | ABCC1,CPT1A,IGFBP2,IGFBP6,NPC1,PDK4,SLC22A5 | 211 (11) |
| EIF4EBP2 | | translation regulator | | -0.572 | 0.00000823 | CEBPA,CEBPD,EGR1,IL1RN,IRF7,PNPLA2,PPARG,PTGS2 | 254 (7) |
| FGF21 | | growth factor | | -0.577 | 0.000468 | CPT1A,DGAT2,EGR1,ERN1,FGFR2,FOS,HMGCS2,IRS2,ME1,PDK4 | 503 (24) |
| colistin | | biologic drug | | -0.577 | 0.00714 | ALCAM,BMP4,DNAJB4,FOS,LGALS3,MYO1B,NPNT,NRIP1,SLCO4A1,SMAD3,SOX4,TJP1 | |
| BHLHA15 | | transcription regulator | | -0.577 | 0.00651 | ARRDC3,CLU,GJB2,RAB3D | |
| kanamycin A | | chemical drug | | -0.577 | 0.00822 | ALCAM,BMP4,DNAJB4,FOS,LGALS3,MYO1B,NPNT,NRIP1,SLCO4A1,SMAD3,SOX4,TJP1 | |
| metronidazole | | chemical drug | | -0.577 | 0.0115 | ALCAM,BMP4,DNAJB4,FOS,LGALS3,MYO1B,NPNT,NRIP1,SLCO4A1,SMAD3,SOX4,TJP1 | |
| IGF2BP1 | | translation regulator | | -0.577 | 0.00565 | AXIN2,CD44,GADD45A,GLI1,IGF2,SERPINH1 | |
| SOX9 | | transcription regulator | | -0.581 | 0.00171 | CEACAM5,CXCR4,FOS,MMP15,MUC2,PPARG,PRKCA,RUNX2,S100A4,SMPD3,SOSTDC1,SOX4,WNT11 | 478 (23) |
| ATG7 | | enzyme | | -0.584 | 0.0000868 | ACAT1,ACSL1,AGR2,CASP7,CD44,CXCL8,HK1,IFIH1,PDLIM1,RCAN1,S100A4,SCARB1,TGFBR2 | 444 (16) |
| obeticholic acid | | chemical drug | | -0.588 | 0.000399 | ABCC2,CD68,CEBPA,CXCL2,CXCL3,FGF19,NR0B2,PPARG,PTGS2 | 541 (18) |
| KLF15 | | transcription regulator | | -0.589 | 0.029 | ACAT1,ACSL1,DGAT2,PDK4,PPARG,PXMP2 | |
| PTPN1 | | phosphatase | | -0.592 | 0.00454 | CXCL2,CXCL3,DUSP1,DUSP2,IRS2,NOX1,PIK3R1,RASA1 | 435 (20) |
| CTNNB1 | | transcription regulator | | -0.6 | 1.46E-15 | AS,FOS,FOSL1,FSTL3,GADD45B,GALM,GATA2,GDF15,GJB2,GLI1,GLI2,HBEGF,HMGCR,HMGCS2,HNF4A,HOXA5,ID2,IDI1,IGF2,IGFBP2,JUN,KITLG,KRT7,LAMC2,LGALS3,LIF,MA | 561 (20) |
| CREBZF | | transcription regulator | | -0.6 | 0.00183 | DNAJB9,ERN1,HSPA1A/HSPA1B,INSIG1,PDK4 | |
| IL4 | | cytokine | | -0.604 | 1.14E-14 | ,HPGD,HRH1,HSPA1A/HSPA1B,IFI44,IFIH1,IFIT3,IFITM3,IFNGR2,IL15RA,IL1R2,IL1RN,IMPDH1,IRF1,IRF7,JUN,JUND,KDM6B,KITLG,KLF6,LGALS3,LGALS3BP,LIF,LIPG,LMO7,MAC | 509 (19) |
| IKZF3 | | transcription regulator | | -0.607 | 0.00000438 | CAV1,CAV2,CD68,DEPP1,DKK1,IFIT3,IRF7,KLF4,NFKBIZ,PROS1,RNF213,S100A16,SIPA1L2,SMAD3 | |
| estrogen receptor | | group | | -0.608 | 2.07E-11 | ,CXCL8,CXCR4,CYP1A1,CYP2B6,EDN1,EGR1,EREG,ETS1,FGFR2,FOS,HBEGF,HSPA1A/HSPA1B,JUN,KRT7,LAMC2,MMP15,NEDD9,NRIP1,NT5E,PDK4,PMAIP1,PPARG,RGS2,SA | 568 (19) |
| IRS1 | | enzyme | | -0.615 | 0.00023 | ATP2A3,CDKN2B,CEBPA,CEBPD,CRABP2,EGR1,FBXO32,FOS,HMGCR,ID2,IRS2,KLF4,NRP1,PIK3R1,PLPP3,PPARG,RHOU | 476 (21) |
| WBP2 | | transcription regulator | | -0.621 | 0.00000293 | ADGRF1,AXIN2,BMP4,CASP7,CSPG4,CXCL1,CXCL2,CXCL3,CXCL8,CYP24A1,EGR1,GLCCI1,JDP2,JUN,LIPG,MAPRE3,MBOAT1,PTGS2,SIGMAR1,SLC16A3,SOX4 | 500 (17) |
| LRP5 | | transmembrane receptor | | -0.625 | 0.00485 | ALDH1A1,AXIN2,HNF4A,IGF2,IRS2,SLC38A5 | 433 (15) |
| KLF6 | 0.654 | transcription regulator | | -0.628 | 0.00000139 | ATF3,BCL6,CXCL2,CXCL8,CYP1A1,ENG,GLI1,HNF4A,MCL1,MSLN,NR1D1,PMAIP1,PPARG,PTGS2,RARG,SERPINH1 | 413 (20) |
| PPARGC1A | | transcription regulator | | -0.628 | 0.00000016 | ,EGLN1,FBXO32,GDF15,HMGCS2,IGFBP6,IL1R2,IL1RN,INSIG1,IRS2,LDHA,MAOB,ME1,MMP15,MSMO1,NR5A2,PCK2,PCSK5,PDIA5,PDK4,PIK3R1,PMAIP1,PNPLA2,PTGS2,RUNX | 372 (18) |
| curcumin | | chemical drug | | -0.631 | 0.000000487 | CL1,CXCL3,CXCL8,CXCR4,CYP1A1,DAPK1,DNAJB1,EDN1,EGR1,ENTPD5,ETS1,FAS,FER,FOS,FOSB,FOSL1,FYN,HSPA1A/HSPA1B,JUN,JUND,KLF9,MCL1,NEO1,PIM1,PNPLA2, | 496 (22) |
| EFNA5 | | kinase | | -0.632 | 0.000156 | CRABP2,EREG,FOSL1,IRF1,IRF7,ITGB6,KRT13,KRT7,NT5E,PTGS2 | 253 (7) |
| niacinamide | | chemical - endogenous mammalian | | -0.632 | 0.0117 | CCNG2,CXCL8,IRF1,KLF4,MCL1,NFYA,PDK4,PMAIP1,PTGS2,RUNX2 | |
| TAL1 | | transcription regulator | | -0.636 | 0.000767 | AZGP1,BCL6,CCNG2,CDK6,CEP70,CMTR2,ETS1,GATA2,GSDMD,HLA-B,HLA-C,HLA-E,ID2,JUN,MSLN,MYB,NFKBIZ,NFYA,REL,SOX4,STK17A,TOB1,ZC3H12A | 58 (2) |
| TCF3 | | transcription regulator | | -0.638 | 0.000000487 | 6,CDKN2B,CGREF1,CTSE,FGFR2,GADD45A,GADD45B,GATA2,GNE,H1-0,ID2,JUN,JUND,KLF4,KLF6,MSMO1,NR0B2,PARP8,PLAC8,PML,PPARG,PRDX4,RARG,SDF2L1,SEC11C, | 270 (7) |
| verapamil | | chemical drug | | -0.64 | 0.0196 | CYP24A1,FOS,HMGCR,JUN,S100A10,TXNIP | |
| SGPP2 | | phosphatase | | -0.64 | 0.00135 | DNAJA1,DNAJA4,DNAJC22,DNAJC3,HSPA4L,HSPH1 | |
| TGFBR1 | | kinase | | -0.647 | 0.0302 | BGN,BMP4,EDN1,HNF4A,KLF4,MUC2,PDK4,WNT11,ZFP36 | |
| C1QA | | other | | -0.647 | 0.00295 | CD44,EGR1,F2R,LCN2,S100A10,SERPINA3 | 466 (16) |
| diosgenin | | chemical - endogenous non-mammalian | | -0.651 | 0.00546 | ABCC2,CYP3A5,EDN1,HMGCR,PTGS2 | 424 (14) |
| sucrose | | chemical - endogenous mammalian | | -0.666 | 0.0406 | ACSS2,ELOVL6,FOS,FOSB,HMGCR | |
| PTPN11 | | phosphatase | | -0.673 | 0.000114 | CD68,CEBPA,CXCL8,EGR1,FOS,GAB1,GATA2,JUN,MCL1,MXD4,PIM1,SPDEF,TP53I11 | 403 (26) |

Supplementary Table 4. Upstream regulator analysis based gene set enrichment analysis (GSEA) of differentially expressed genes (DEGs) after ATRA+PDT compared to PDT

| Upstream Regulator | Expr Log Ratio | Molecule Type | Predicted Activation State | Activation z-score | p-value of overlap | Target Molecules in Dataset | Mechanistic Network |
|---|---|---|---|---|---|---|---|
| BTK | | kinase | | -0.68 | 0.000412 | BBC3,CDC25A,CDK6,CEBPA,CXCL3,CXCL8,CXCR4,EMP1,ETS1,IFI35,IFIT3,IRF1,JUN,OAS3,PIM1 | 424 (19) |
| glutamine | | chemical - endogenous mammalian | | -0.682 | 0.00000429 | AKR1C1/AKR1C2,ASS1,CXCL1,CXCL8,DUSP1,FOS,GADD45A,HK1,HSPA1A/HSPA1B,IGF2,IGFBP2,JUN,KLF4,MCL1,PCK2,PTGS2,SLC16A1,WARS1 | 492 (24) |
| mir-9 | | microRNA | | -0.689 | 0.0319 | BCL6,CXCR4,ETS1,MEIS2 | |
| PNPLA2 | 0.343 | enzyme | | -0.701 | 0.00473 | ACSL1,CD68,CEBPA,CPT1A,DGAT2,PDK4,PNPLA2,PPARG,SLC16A1 | 393 (21) |
| KDM3B | | enzyme | | -0.707 | 0.00917 | AXIN2,DAPK1,DEGS2,FRMD4B,GATA2,HJURP,NEDD4L,SORL1,STMN1 | |
| UBE2I | | enzyme | | -0.707 | 0.0000566 | CEBPA,CEBPD,CPT1A,IRF1,NR0B2,PDK4,PML,PPARG,SGK1 | 469 (24) |
| EFNA2 | | kinase | | -0.707 | 0.00635 | CRABP2,FOSL1,FOXL1,IRF7,KRT13,KRT7,NEDD4L,NT5E | |
| selenomethylselenocysteine | | chemical - endogenous mammalian | | -0.707 | 0.00000561 | ABCC1,ATP1B3,CLU,CXCL3,CYP3A5,EGR1,JUN,PSME1 | 170 (7) |
| CLOCK | | transcription regulator | | -0.711 | 0.0000779 | EMIP,DNAJB9,ELOVL6,F2R,GADD45A,GADD45B,HES6,HMGCS2,HSPH1,LRATD2,MMP7,MSMO1,NPAS2,NR0B2,NR1D1,NR1D2,OAS3,PPIF,RAP2B,SERINC5,ST6GALNAC4,TMED3 | |
| FOS | 0.547 | transcription regulator | | -0.72 | 4.51E-13 | A-B,HPGD,IGFBP6,IRS2,JUN,KITLG,KLF6,KRT13,LGALS3,LGALS3BP,LGALS4,LRP8,LTBP3,MALAT1,MAOA,MAP4K4,MMP7,MSLN,MT2A,MXD1,NR5A2,NRIP1,PARD6B,PHLDA1,PI | 477 (20) |
| CLDN7 | | other | | -0.72 | 3.87E-11 | DIB,ATP5MC1,CD68,CEACAM5,CEACAM6,CXCL8,DKK1,F5,FOSL1,HLA-B,IFI44,LGALS3,LGR4,LIMA1,MT1X,MT2A,MUC13,NCOA7,PLAAT3,SCEL,SOSTDC1,SYTL2,TEAD2,TINAGL | |
| prednisolone | | chemical drug | | -0.724 | 0.0000116 | RC3,BTG1,CEBPD,CTSS,CXCR4,CYP1A1,DAPK1,DUSP1,EDN1,FAS,FBXO32,GAB1,GLI1,IRS2,KLF9,MAP1LC3B,MCL1,NQO2,PLAAT4,PTGS2,SGK1,SPHK2,TGM2,TMEM43,TSC22 | 442 (19) |
| T3-TR-RXR | | complex | | -0.728 | 0.0142 | AKR1C1/AKR1C2,AKR1C3,KLF9,ME1,SCARB1,SLC16A3 | |
| 8-bromoguanosine 3',5'-cyclic monophosphate | | chemical - kinase inhibitor | | -0.73 | 0.00754 | CAV1,CYP3A5,EDN1,FOS,S100A10,SLC12A4 | 457 (23) |
| salicylic acid | | chemical drug | | -0.742 | 0.0144 | BGN,CXCL8,GDF15,KITLG,MCL1,PTGS2,SCARB1 | |
| testosterone | | chemical - endogenous mammalian | | -0.744 | 7.14E-09 | ,DDIT4,DNMT3A,EGR1,FAS,FBXO32,FGFR2,FOS,FOSB,FOSL1,GADD45A,HMGCR,HMGCS2,HSPA1A/HSPA1B,IRS2,JUN,JUND,MAOA,MMP7,NR5A2,PIK3R1,PKIB,PPARG,PTGS2 | 615 (22) |
| propranolol | | chemical drug | | -0.749 | 0.0488 | CYP1A1,DUSP1,FOS,MAOA,PTGS2 | |
| ESR1 | | ligand-dependent nuclear receptor | | -0.751 | 1.16E-21 | K2,HLA-B,HLA-C,HLA-E,HLA-J,HMGA2,HMGCR,HMGCS2,HSPH1,IFI44,IFITM2,IFITM3,IGF2,IGFBP2,INCENP,IRF1,IRS2,JUN,JUND,KLHL24,KRT13,KRT7,LATS2,LCN2,LGALS3BP,LI | 567 (21) |
| PI3K (family) | | group | | -0.752 | 2.57E-08 | AKR1B10,AKR1C1/AKR1C2,BIRC3,CCNG2,CDKN2B,CLDN2,CXCL1,CXCL8,CXCR4,DDIT4,EGLN1,FBXO32,FOS,GLI1,HK1,MCL1,PTGS2,SCARB1,SOX4,TGM2,TJP1,TXNIP | 553 (29) |
| silibinin | | chemical drug | | -0.756 | 0.0243 | BBC3,CD44,CDK6,CXCL8,EGR1,KLF4,MCL1,PTGS2 | |
| actinomycin D | | biologic drug | | -0.756 | 2.46E-12 | ,CAV1,CCN1,CD55,CXCL1,CXCL8,CYP1A1,CYP24A1,DUSP1,ERRFI1,ETS1,FAS,FOS,GADD45A,HBEGF,HK1,HMGA2,ID2,IL1R2,IRF1,IRS2,ITPR2,JUN,MAOA,MCL1,MUC2,PMAIP1,I | 496 (18) |
| KDM1A | | enzyme | | -0.758 | 0.0000571 | N12,CTSS,CXCL16,DAPP1,DKK1,EIF2AK2,FRRS1,ID2,IFI44,IFIT3,IFITM3,IGFBP2,IL1RN,IQGAP2,IRF7,LATS2,LGALS3BP,MYO1B,ONECUT2,PARP14,PLXNB2,PTPRB,RARG,RHOU | 268 (7) |
| desmopressin | | biologic drug | | -0.761 | 0.00623 | ESAM,FOS,HSPA1A/HSPA1B,HSPH1,LIMA1,NDRG2,PPP1R1B,RAP1GAP,SEC61B,TJP2,VDAC1 | |
| bisindolylmaleimide iv | | chemical - kinase inhibitor | | -0.762 | 0.0279 | CXCL8,EGR1,FOS,MUC2 | |
| SB-431542 | | chemical reagent | | -0.767 | 0.000407 | BGN,BMP4,CD44,CYP1A1,FOS,GADD45B,GATA2,GJB2,ID2,IDI1,IL1R2,RUNX2,SQLE,TGFB2,TJP1,TNFRSF12A | 558 (22) |
| BRD4 | | kinase | | -0.768 | 0.0000808 | ABLIM1,ALDH1B1,BCL3,BTN3A2,CCNA2,CDC25A,CDKN2B,CEBPA,CXCL2,CXCR4,ETV6,FOS,ID2,IRF1,MCL1,MYB,PIM1,SLC38A5,SORD | 486 (19) |
| TWIST1 | | transcription regulator | | -0.775 | 0.000205 | ALCAM,ALDH1A1,CD44,CELSR1,CLU,CXCL8,ETS1,FAS,FGFR2,FOS,FOSL1,GLI1,GSN,NT5E,RUNX2,S100A4,SLC28A3,TGFB2,TGFBR2,ZYX | 469 (23) |
| RNF31 | | enzyme | | -0.777 | 0.000238 | AURKA,CXCL1,CXCL16,CXCL2,CXCL3,CXCL8,CXCR4,IL15RA,IL1R2,IL1RN,NR5A2,SCARB1 | 437 (13) |
| MYC | | transcription regulator | | -0.78 | 9.33E-16 | HIVEP2,HLA-B,HLA-E,HOXA5,HSPB1,HSPH1,ID2,IFI35,IFI44,IFIH1,IFIT3,IQGAP2,IRF7,ITM2B,JAG2,JUN,KDM6B,KLF4,KLF6,KLK6,KRT7,LDHA,LGMN,LIMA1,LOC102724788/PRODH | 570 (19) |
| FGF7 | | growth factor | | -0.787 | 0.00143 | APRT,AQP5,CD44,CEBPA,CEBPD,FGFR2,IRF1,LPCAT1,RUNX2,S100A10,SMAD3 | 488 (22) |
| chromium | | chemical drug | | -0.788 | 0.00404 | AJUBA,ALG2,CXCL8,CYP1A1,ELF3,ENTPD5,PPIF,TNFAIP2 | 194 (7) |
| NR1H4 | | ligand-dependent nuclear receptor | | -0.789 | 0.000103 | ABCC2,AMBP,CASP9,CCL15,CEBPA,CYP2B6,CYP3A5,EDN1,FAS,FGF19,HNF4A,KLF4,LCN2,LDHA,NDRG2,NR0B2,NR5A2,PGM1,PPARG,PTGS2,SCARB1,TXNIP | 525 (23) |
| SOCS3 | | phosphatase | | -0.792 | 0.0385 | ATF3,EGR1,FOS,IL1RN,IRF1,IRS2,JUN,MUC2,TFF3 | |
| ESR2 | | ligand-dependent nuclear receptor | | -0.793 | 3.05E-14 | RL1,FGF19,FOS,GADD45B,GYG1,HMGA2,HPGD,IFI35,IFI44,IFIH1,IFIT3,IRF7,KLF4,KRT20,LGALS3BP,LOXL4,MAOA,MDK,MSLN,MXD1,NDRG2,NEDD9,NFYA,PARP12,PARP14,PDK | 618 (21) |
| S100A8 | | other | | -0.798 | 0.000591 | AKAP12,ALDH1A1,ATF3,BCL3,CXCL2,CXCL3,CXCL8,DDC,DDIT4,EFNA1,FAS,FOSL1,GJB2,GPRC5A,LOC102724788/PRODH,PHLDA1,PLK3,SEMA3B,SGK1,TXNIP | 496 (20) |
| NR1H2 | | ligand-dependent nuclear receptor | | -0.798 | 0.00143 | ABCA2,CXCL16,CXCL2,ELOVL6,GLI1,HK1,HMGCR,IL1RN,LRP8,PNPLA2,PTGS2,REN,SCARB1 | 374 (22) |
| vitamin D | | chemical drug | | -0.811 | 0.0291 | CCN1,CYP1A1,CYP24A1,GPR158,REN,SGK1,VDR | |
| 12-(3-adamantan-1-yl-ureido) dodecanoic acid | | chemical reagent | | -0.816 | 0.00427 | BIRC3,CARD6,CASP4,CASP7,CASP9,FAS,MCL1,SPHK2,TRAF4 | |
| selumetinib | | chemical drug | | -0.816 | 0.00246 | ABCC1,FOS,FOSB,FOSL1,JUN,RUNX2 | 489 (24) |
| ZBTB20 | | transcription regulator | | -0.816 | 0.0409 | ACSS2,CYP2B6,CYP3A5,IGFBP2,NEDD9,PDK4 | |
| MEX3A | 0.444 | other | | -0.816 | 0.00984 | AURKA,CD44,KCNE3,NRP1,PDK4,PPARG | |
| NEUROG1 | | transcription regulator | | -0.816 | 0.0393 | ASS1,CEMIP,CXCL1,GASK1B,LRIG1,NEO1,S100A4 | |
| HBB | | transporter | | -0.816 | 0.0218 | CXCL2,CXCL3,CXCL8,GDF15,LIF,SCARB1 | |
| evodiamine | | chemical - endogenous non-mammalian | | -0.816 | 0.000672 | BIRC3,CASP9,CXCL8,CYP1A1,MCL1,PTGS2 | 403 (16) |
| ZBTB16 | | transcription regulator | | -0.819 | 0.00124 | ACER2,BCL6,CCNA2,CD44,CD55,DDC,DHRS3,EIF2AK2,F2R,F2RL1,GALNT6,HPGD,ID2,ME1,NFKBIZ,PIK3R1,PTGS2,RARG,REL,RUNX2,TSC22D1,TSC22D3 | |
| MDM2 | | transcription regulator | | -0.832 | 0.0248 | ATF3,BBC3,CCNA2,CXCL8,ETS1,HIPK2,IGFBP6,IL1RN,NR0B2 | |
| ritonavir | | chemical drug | | -0.849 | 1.19E-09 | CAM5,CEACAM6,CEP135,CEP70,CXCL8,CYP1A1,CYP2B6,CYP3A5,HMGCR,IDI1,KITLG,MAOB,ME1,MSMO1,MYB,PADI2,PPARG,SC5D,SLC39A10,SQLE,TMPRSS4,TRIM31,TSC22D | 474 (16) |
| TRIB3 | | kinase | | -0.853 | 0.00000468 | CDC25A,CEBPA,DDIT4,GDF15,PCK2,PMAIP1,PPARG,PSPH,RUNX2 | 498 (22) |
| ERBB3 | | kinase | | -0.863 | 0.0000447 | ADAM9,AMOTL2,BGN,CCN1,CCNG2,CLU,CXCR4,DUSP1,EGR1,HBEGF,HMGA2,IGF2,PTGS2,SERPINA3,SGK1,STMN1,TGM2 | 443 (19) |
| CALCA | | other | | -0.863 | 0.000212 | CD44,CXCL8,CXCR4,CYP24A1,EDN1,ESAM,FOS,LIMA1,NDRG2,PPP1R1B,RAP1GAP,RUNX2,SEC61B,TJP2 | 452 (24) |
| GLI2 | -1.025 | transcription regulator | | -0.871 | 0.0169 | BMP4,CLU,CXCL2,CXCL3,GLI1,GLI2,JAG2,PI3,RUNX2,UGCG,VDR | |
| HLX | | transcription regulator | | -0.873 | 0.000928 | CD44,CLDN1,CTSS,CXCL8,EGR1,GDF15,JUN | 314 (7) |
| mirdametinib | | chemical drug | | -0.882 | 0.0239 | CXCL8,DDIT4,ETV1,KLF4,MCL1 | |
| Go6983 | | chemical - kinase inhibitor | | -0.883 | 0.0000716 | CLDN1,FOS,PPARG,PRKCA,PTGS2,SCARB1 | 449 (21) |
| fenofibrate | | chemical drug | | -0.884 | 0.000787 | ,ACSL1,CASP4,CD68,CDC25A,CLU,CPT1A,CXCL8,DGAT2,EDN1,FBXO32,GADD45A,GYG1,ID2,JAG2,JUN,PCK2,PDK4,PNPLA2,PTGS2,PTPN21,SC5D,SCARB1,TXNIP,UGT1A10 (i | 462 (25) |
| MYOD1 | | transcription regulator | | -0.892 | 0.0211 | ASS1,BCL6,CYP2S1,DUSP1,EIF2AK2,FOSL1,FYN,GADD45A,HES6,HJURP,HSPA4L,ID2,IGF2,KLF6,LIF,PFKM,SHCBP1,TEAD2,TJP1,TRIM21 | |
| ARNTL | | transcription regulator | | -0.896 | 0.0484 | ANGPT1,CCNG2,CEBPA,ELOVL6,HMGCS2,NR1D1,TBC1D4 | |
| SMAD7 | | transcription regulator | | -0.898 | 0.00929 | ACVR1,BGN,CCNA2,CDKN2B,CGREF1,FAS,FSTL3,GADD45A,HBEGF,HDAC9,ID2,LTBP3,SMAD3,TGFB2,TGFBR2 | 433 (21) |
| ilomastat | | chemical drug | | -0.9 | 0.0241 | CXCL3,CXCL8,HBEGF,TJP1 | |
| EPHB4 | | kinase | | -0.905 | 0.0000512 | BMP4,CXCR4,FGFR2,GATA2,IGF2,KITLG,PDLIM7,RASA1,TGFB2,TGFBR2,WNT11 | |
| ciprofloxacin | | chemical drug | | -0.905 | 0.0203 | ACAA1,ADAM8,ALDH2,BMP4,CD68,DNAJC3,LGMN,MYOF,NRP1,PLXNB2,VDAC1 | |
| VIP | | other | | -0.91 | 0.0414 | CEBPD,CFTR,CLCF1,CXCL2,CXCL3,CXCL8,FOS,IRF1,PTGS2,TJP1 | |
| mir-135 | | microRNA | | -0.911 | 0.0188 | EDN1,KLF4,MCL1,MTSS1,RUNX2 | |
| Wnt | | group | | -0.927 | 0.024 | CD44,CEBPA,GLI1,ID2,PPARG,RUNX2 | |
| HEXIM1 | | transcription regulator | | -0.927 | 0.0319 | ANGPT1,ANXA1,BBC3,TGFB2 | |
| ERBB4 | | kinase | | -0.927 | 0.000345 | ADAM9,CASP4,DUSP1,EGR1,GADD45A,HBEGF,HMGA2,IGF2,JUN,PTGS2,SERPINA3,SERPINE2,TBXAS1 | 471 (20) |
| glucosamine | | chemical - endogenous mammalian | | -0.928 | 0.00309 | ASS1,CEBPA,CXCL1,CXCL3,CXCL8,GPD1,PTGS2,REN,TXNIP | 494 (24) |
| PRNP | | other | | -0.933 | 0.00143 | ADAM8,CASP9,CD44,DKK1,EGR1,FOSL1,HSPA1A/HSPA1B,HSPB1,ID2,IGF2,PLPP3 | 469 (22) |
| NCOA1 | | transcription regulator | | -0.941 | 0.0141 | CASP7,CEBPA,CYP1A1,CYP2B6,EGR1,FGFR2,FOS,GJB2,GYG1,IL1RN,NFKBIZ,NR0B2,PARD6B,PPARG | |
| clofibrate | | chemical drug | | -0.942 | 0.0000927 | ACAA1,CEACAM1,CPT1A,CXCL8,CYP1A1,DPP4,FAS,GADD45A,HMGCR,HMGCS2,IL1RN,MYB,PEX11A,PTGS2,SULT2B1 | 340 (21) |
| Alpha catenin | | group | | -0.952 | 0.00000002 | ADAM8,AXIN2,BCL3,BGN,BIRC3,CXCL2,DKK1,ENG,EREG,IGF2,IGFBP2,IGFBP6,IRF1,NFKBIZ,PTGS2,SGK1,STEAP2,STEAP4,TGM2,TNFAIP2,TNFRSF12A,TSPAN8,ZYX | 410 (16) |
| ERN1 | 0.515 | kinase | | -0.952 | 0.000000137 | EBPD,CXCL2,DGAT2,DNAJB9,DNAJC3,ETS1,FAS,ITGB6,JUN,MAP1LC3B,MCL1,NCOA3,PDIA5,PPARG,PROS1,SDF2L1,SEC24D,SEC61A1,SLC25A28,TINAGL1,TMED3,TXNDC5,V | 419 (18) |
| canrenoate potassium | | chemical drug | | -0.958 | 0.00818 | CD68,EDN1,LGALS3,PTGS2 | |
| NS-398 | | chemical reagent | | -0.96 | 0.0113 | ANGPT1,CXCL8,CXCR4,F2R,GATA6,MCL1,MUC2,PPARG,PSMB8,PTGS2,RUNX2 | |
| miR-141-3p (and other miRNAs w/seed AACACUG) | | mature microRNA | | -0.963 | 0.0302 | AKR1C1/AKR1C2,CDK6,GADD45A,GRB7,MSLN,PMAIP1,RNF128,TGFB2 | |
| MTTP | | transporter | | -0.97 | 0.00112 | CPT1A,CXCL3,ERN1,HMGCR,JUN,PPARG,RUNX2 | 182 (7) |
| WWTR1 | | transcription regulator | | -0.971 | 0.00000641 | CCN1,CCNA2,CD44,CXCL1,CXCL2,CXCL3,CXCL8,EMP1,ID2,LAMA5,LATS2,RUNX2,UCA1 | |
| TG | | other | | -0.975 | 0.0165 | ANGPT1,CEBPA,CPT1A,IL1R2,PDK4 | |
| CSHL1 | | growth factor | | -0.976 | 0.0105 | BCL6,CYP2B6,CYP3A5,FOS,HNF4A,IRF1,PIK3R1,SERPINH1 | |
| Sch-23390 | | chemical drug | | -0.984 | 0.000672 | EGR1,FAS,FOS,FOSB,FOSL1,PLD2 | 419 (23) |
| HDAC1 | | transcription regulator | | -0.986 | 0.0000314 | IN2,BBC3,BCL6,CCNA2,CD44,CDC25A,CEACAM1,CEACAM6,CEBPA,CXCL1,CXCL8,DUSP1,EGR1,FAS,FOS,GLI1,LCN2,PMAIP1,PPARG,PTGS2,RHOB,RUNX2,SGK1,STX1A,TGFE | 473 (20) |
| pentobarbital | | chemical drug | | -0.987 | 0.00387 | CYP2B6,FOS,JUN,JUND | 478 (20) |
| PI3K (complex) | | complex | | -0.99 | 5.26E-12 | ,DDIT4,DKK1,DUSP1,ETS1,FBXO32,FOS,FOSL1,GATA2,GLI1,GLI2,HBEGF,HK1,HMGCR,IGF2,IGFBP2,IL1RN,JUN,LCN2,MCAM,MCL1,MT2A,NOX1,PIM1,PPP1R1B,PTGS2,RGS2,R | 488 (21) |
| PLAG1 | | transcription regulator | | -0.997 | 0.0051 | BTN3A2,CNKSR1,IGF2,QSOX1,S100A2,STX1A,TGM2,TSPAN4 | |
| pregna-4,17-diene-3,16-dione | | chemical - endogenous non-mammalian | | -1 | 0.0241 | CEBPA,MCL1,PPARG,PTGS2 | |
| AHI1 | | other | | -1 | 0.0363 | CCNG2,CMTM7,IL1RN,REPS2 | |
| KDM6A | | enzyme | | -1 | 0.0126 | ANKS4B,CSPG4,DKK1,HNF4A,TM4SF4,UPK3B | |
| GAS2L3 | | other | | -1 | 0.00651 | F2R,FAS,LIF,SAT1 | |
| fontolizumab | | biologic drug | | -1 | 0.00818 | CXCL1,FAS,IFI35,IL15RA | |
| TRG | | other | | -1 | 0.000593 | CAV1,CAV2,CXCL1,NT5E | |
| SEL1L | | other | | -1 | 0.0363 | COL17A1,DNAJC3,DOCK5,KANK1,NTN4,PRKCA,TGFBR2,UQCRFS1,ZFHX3 | |
| ASXL1 | | transcription regulator | | -1 | 0.0158 | CLP1,FAM162A,GPCPD1,HOXA5,PELI2,PPARG | |
| MYT1 | | transcription regulator | | -1 | 0.00818 | ARHGDIB,DAPP1,PTPRB,STEAP3 | |
| NKX2-2-AS1 | | other | | -1 | 0.041 | AK4,FAM162A,FAM210A,PDK4 | |
| LMNA | | other | | -1 | 0.0005 | C4,CCNG2,CEBPA,EDN1,ERN1,FUT4,GATA6,GLIS3,IFI44,IFIH1,IRF1,JUN,JUND,MAFF,MCAM,MMP15,MXD4,NPC1,PLD2,PPARG,RESF1,SEMA7A,SKAP2,SLC22A23,ST6GALNAC4, | 459 (17) |
| EFNA1 | -0.373 | other | | -1 | 0.00012 | CLDN2,CRABP2,EPHA2,FOSL1,GATA6,HERC6,IRF7,ITGB6,KRT13,NEDD4L,NT5E,RASA1 | 255 (7) |
| Gcg | | other | | -1 | 0.00193 | ALDH1B1,ASS1,ATP1B1,FOS,ME1,METTL7B,PEX11A,RARRES1,RNASE4 | |
| EMD | | other | | -1 | 0.0123 | EGR1,HOXA5,IRF1,PPL | |
| FABP1 | | transporter | | -1 | 0.0279 | FAAH,FGF19,PNPLA2,PPARG | |
| Z-DEVD-FMK | | chemical - protease inhibitor | | -1 | 0.00144 | CXCL2,CXCL3,MCL1,PLD1 | 419 (14) |
| daidzein | | chemical drug | | -1 | 0.00729 | AKR1C3,CAV1,CYP1A1,FOSB,HMGCR,ID2,IRF7,PIK3R1,PKIB,PTGS2,ZNF212 | 576 (23) |
| biochanin A | | chemical toxicant | | -1 | 0.0148 | CXCL8,CYP1A1,PTGS2,RUNX2 | |

Supplementary Table 4. Upstream regulator analysis based gene set enrichment analysis (GSEA) of differentially expressed genes (DEGs) after ATRA+PDT compared to PDT

| Upstream Regulator | Expr Log Ratio | Molecule Type | Predicted Activation State | Activation z-score | p-value of overlap | Target Molecules in Dataset | Mechanistic Network |
|---|---|---|---|---|---|---|---|
| chelerythrine | | chemical drug | | -1 | 0.0279 | CXCL3,MCL1,NOX1,PIM1 | |
| Immunoglobulin | | complex | | -1.001 | 2.96E-09 | M210A,FAS,FGFR2,FLNB,FOS,FOSB,GALM,GLI1,HLA-E,HSPA6,IFIT3,IFNGR2,IL1R2,IL1RN,IRF1,IRF7,KLF4,LCN2,LGALS3,LGALS3BP,LOC102724788/PRODH,LTBP3,MAFF,MAP3I | 477 (19) |
| metformin | | chemical drug | | -1.035 | 0.000977 | ATF3,BCL3,CAV1,CXCL8,CYP2B6,CYP3A5,DDIT4,EGLN1,ERN1,FOS,GADD45B,GPD1,IFI44,IL1RN,INSIG1,NR0B2,PARP14,PFKM,PPP1R3B,PTGS2,RUNX2,SCARB1 | 411 (22) |
| PPARA | | ligand-dependent nuclear receptor | | -1.037 | 0.000000104 | SE,CYP1A1,CYP2B6,EDN1,ELOVL6,FOS,GPD1,HLA-E,HMGCR,HMGCS2,HNF4A,IDI1,IFITM3,IGFBP2,IGFBP6,IL1RN,INSIG1,LGALS4,LIPG,LSS,MAT2A,MSMO1,NR1D1,PDK4,PEX1 | 433 (21) |
| apigenin | | chemical - endogenous non-mammalian | | -1.039 | 0.0217 | CDK6,CXCL3,CXCL8,CYP1A1,DGAT2,DPP4,DVL1,FOS,PTGS2 | |
| haloperidol | | chemical drug | | -1.052 | 0.00377 | EGR1,FOS,FOSB,HMGCR,HSPG2,PPP1R1B,PRKAR2A,PRKCA,RASA1,SC5D,SIGMAR1 | 394 (23) |
| HIC1 | | transcription regulator | | -1.055 | 0.0291 | CYP24A1,EPHA2,FHL2,LCN2,LIF,LRP8,MCAM | |
| IRF4 | | transcription regulator | | -1.061 | 0.00216 | B2M,BCL6,BIRC3,CD68,CDK6,CTSS,CXCL3,CXCR4,IL15RA,IL1RN,IRF1,IRF7,PLSCR1,PRKCA,PROCR,RHOB,RUNX2,TRIM21,ZFP36L1 | 467 (17) |
| UPF2 | | other | | -1.066 | 0.000012 | CLU,KANK1,NEO1,NRP1,PTGS2,PTPN21,RCAN1,TSPAN13,TSPAN8 | |
| FLZ | | chemical drug | | -1.067 | 0.00207 | DDIT4,FAS,PMAIP1,PTGS2 | 364 (12) |
| harmine | | chemical - endogenous non-mammalian | | -1.067 | 0.00287 | CYP1A1,FOS,PTGS2,RUNX2 | 466 (12) |
| SLC16A2 | | transporter | | -1.067 | 0.0368 | CIRBP,KLF9,SEMA7A,SLC16A1,SLC16A3 | |
| Mt1 | | other | | -1.067 | 0.0241 | ATF3,CEBPA,FOS,JUN | |
| Mt2 | | other | | -1.067 | 0.0148 | ATF3,CEBPA,FOS,JUN | |
| farglitazar | | chemical drug | | -1.067 | 0.00095 | EDN1,REN,SGK1,SULT2B1 | 210 (7) |
| calyculin A | | chemical toxicant | | -1.067 | 0.0363 | EGR1,MCL1,PRKCA,TGM2 | |
| choline | | chemical - endogenous mammalian | | -1.067 | 0.0268 | BBC3,DNMT3A,ERRFI1,IGF2,PTGS2 | |
| hydroxypropyl-beta-cyclodextrin | | chemical drug | | -1.067 | 0.0461 | CD68,HMGCR,NPC1,SCARB1 | |
| TAF4 | | transcription regulator | | -1.069 | 0.00232 | BCL6,CLDN1,EHF,ETS1,GJB2,HBEGF,ID2,IL1RN,KLF4,MAFF,MAP4K4,MXD1 | |
| paricalcitol | | chemical drug | | -1.072 | 0.0406 | F2RL1,PTGS2,REN,TFPI,VDR | |
| TFAP2A | | transcription regulator | | -1.081 | 0.0000401 | ALCAM,ANXA1,BIRC3,CCNDBP1,CEBPA,CXCL1,CXCL2,EREG,F2R,IGF2,KLF4,MT2A,PAG1,PLAAT3,PPARG,RAB27B | 449 (21) |
| sulforafan | | chemical drug | | -1.089 | 0.00203 | ABCC2,AKR1B10,AKR1C1/AKR1C2,AKR7A3,ALDH1A1,CD44,CXCL8,DDC,ENTPD5,FAS,FOS,GATA6,GSTA4,HIPK2,PML,PTGS2 | 506 (22) |
| KDM2B | -0.33 | enzyme | | -1.091 | 0.0368 | CDKN2B,CRABP2,FOS,GATA6,LAMA5 | |
| KMT5B | | enzyme | | -1.091 | 0.000178 | DUSP1,MT2A,SGK1,TSC22D3 | |
| FASN | | enzyme | | -1.093 | 0.00636 | AJUBA,AMOTL2,CAV1,CXCL8,DDIT4,DGAT2,ELOVL6,PNPLA2,PTGS2 | 412 (21) |
| CDK19 | | kinase | | -1.095 | 1.29E-15 | CL1,CXCL2,CXCL8,DDIT4,EPHA2,FAS,GADD45A,HBEGF,HSPA4L,JAG2,JUN,LIF,LSS,MXD1,MXD4,PLK3,PPP1R15A,PROCR,RAP2B,SAT1,SEC61A1,TCN2,TRAF4,TSC22D1,TXNIP | |
| CYP27A1 | | enzyme | | -1.103 | 0.0207 | ABCC2,CYP2B6,CYP3A5,HMGCR | |
| SS18-SSX2 | | fusion gene/product | | -1.103 | 0.00144 | AXIN2,CDC25A,DKK1,MCL1 | |
| parthenolide | | chemical drug | | -1.118 | 0.00259 | AXIN2,CXCL2,CXCL3,CXCL8,LIF,PDK4,PTGS2 | 425 (18) |
| eplerenone | | chemical drug | | -1.119 | 0.0268 | CD68,EDN1,LGALS3,PPARG,PTGS2 | |
| miR-26a-5p (and other miRNAs w/seed UCAAGUA) | | mature microRNA | | -1.121 | 0.000503 | CDK6,CHORDC1,CXCL3,EPHA2,HPGD,PTGS2,TGFBR2 | |
| sorafenib | | chemical drug | | -1.123 | 0.0126 | CDK6,CXCL8,ERN1,MCL1,PIM1,PPP1R15A | |
| CDX2 | | transcription regulator | | -1.127 | 5.77E-08 | ,AOC1,AXIN2,CAV1,CDHR5,CLDN1,CLDN2,CLU,CTSE,DDC,ETS1,FXYD3,GLI2,HEPH,HNF4A,HOXA3,HOXA5,JUN,KLF4,LAMC2,MTUS1,MUC2,PARD6B,SLC26A6,UGT1A10 (include | |
| PP1 | | chemical - kinase inhibitor | | -1.129 | 0.000134 | ANGPT1,CXCL16,CXCL8,CXCR4,FOS,IL1RN,JUN,PTGS2,TJP1 | 471 (24) |
| manidipine | | chemical drug | | -1.131 | 0.000178 | CXCL8,FOS,HMGCR,JUN | 273 (12) |
| BDKRB2 | | G-protein coupled receptor | | -1.131 | 0.00507 | CD44,EDN1,PTGS2,REN | 511 (27) |
| ZNF106 | | other | | -1.134 | 0.00521 | HSPB1,IFITM3,LGALS3,NCMAP,SERPINA3,TNFRSF12A,UGT8 | |
| lysophosphatidylinositol | | chemical - endogenous mammalian | | -1.134 | 0.00846 | CFTR,CPT1A,CXCL3,CXCL8,INSIG1,LDHA,PFKM | 163 (7) |
| trichloroethylene | | chemical toxicant | | -1.134 | 0.0227 | CDK6,EGR1,ID2,PLD1,RHOB,SLC16A1,TXNIP | |
| GPS2 | | transcription regulator | | -1.134 | 0.0366 | CDK6,CXCL8,DHRS3,HIVEP2,RHOB,SGK1,TBC1D2 | |
| benzene | | chemical toxicant | | -1.134 | 0.0227 | CDK6,EGR1,ID2,PLD1,RHOB,SLC16A1,TXNIP | |
| SATB1 | | transcription regulator | | -1.138 | 0.000559 | CEACAM1,DNMT3A,ETS1,F5,FEZ2,FOSB,GADD45B,GDF15,IL1R2,IRF7,KITLG,PTGS2,SGK1,SIPA1L2,SMAD3,TRPS1,TSC22D3,TUBA4A,WARS1 | |
| HNF1A | | transcription regulator | | -1.143 | 0.0000212 | DC,DPP4,ELF3,FAM107B,FAS,FUT8,FXYD3,GOLT1A,HGD,HHLA2,HMGCR,HNF4A,HNF4G,ITM2B,KDM2B,LINC01559,LRRC31,MT1X,MTMR11,NR0B2,NR1D1,NR5A2,RNASE4,RUN) | 514 (17) |
| AGTR1 | | G-protein coupled receptor | | -1.151 | 0.000731 | DUSP1,EDN1,FOS,JUN,PTGS2,REN,SGK1,TXNIP | 349 (22) |
| NR2F2 | | ligand-dependent nuclear receptor | | -1.154 | 0.00984 | ALDH2,ANGPT1,GATA6,HNF4A,NR5A2,NRP1 | 247 (10) |
| NPC2 | | transporter | | -1.154 | 0.0125 | AKR1C3,CREB5,MAP1LC3B,PPARG,PTGS2 | |
| honokiol | | chemical - endogenous non-mammalian | | -1.154 | 0.0212 | CDK6,FOS,MCL1,PPARG,PTGS2 | |
| SOCS1 | | other | | -1.157 | 0.00392 | CD44,CXCL2,CXCL8,DUSP1,FAS,FOS,IFI44,IFIH1,IFIT3,IRF1,IRF7,IRS2,JUN,PTGS2,TGM2 | 345 (17) |
| NR4A1 | | ligand-dependent nuclear receptor | | -1.157 | 0.00017 | ATF3,ATP5MC3,AXIN2,CEBPA,CRABP2,DAPK1,EHF,GALM,GPD1,HCAR1,KITLG,MAT2A,MMP7,NT5E,NUCB2,PCK2,PDK4,PFKM,PGM1,PPARG,SAT1,SMAD3,STARD10,SUCLA2,T | 425 (22) |
| LRP6 | | transmembrane receptor | | -1.165 | 0.00754 | ALDH1A1,AXIN2,CAV1,CD68,DKK1,FOS,MMP7 | 437 (21) |
| dimethyl itaconate | | chemical reagent | | -1.172 | 0.000319 | ATF3,ERN1,IFIT3,IFITM3,IRF1,LCN2,NFKBIZ | |
| ESRRA | | transcription regulator | | -1.172 | 0.000303 | ACSL1,CEBPA,CKB,GRB7,GSDMB,HK1,HMGCR,IDI1,KRT13,KRT20,LDHA,MAOB,NR0B2,NR1D1,NRIP1,PCK2,PDK4,PPARG,PPP1R1B,RUNX2,WNT11 | 419 (17) |
| ANXA1 | 0.312 | enzyme | | -1.176 | 0.00592 | ANXA1,COL18A1,CXCL2,CXCL8,CXCR4,PTGS2,TSC22D3 | 369 (20) |
| KCNK9 | | ion channel | | -1.195 | 0.00347 | CTSE,GADD45B,LCN2,REN,SLC16A3,SNCG,TM4SF4 | |
| FOSL1 | 0.522 | transcription regulator | | -1.197 | 0.000071 | CCN1,CD44,CXCL8,EDN1,EGR1,FOS,FOSB,FOSL1,HPGD,JUN,MT2A,NRP1,RUNX2,SERPINE2 | 467 (20) |
| 5-N-ethylcarboxamido adenosine | | chemical reagent | | -1.207 | 0.0000501 | BAG3,BCL3,CDKN2B,CEBPD,CXCL8,DUSP16,EGR1,GADD45A,IFIT3,KLF6,NRP1,NT5E,PROCR,RGS2,TXNIP,UBC,VDR | 465 (20) |
| DACH1 | | transcription regulator | | -1.213 | 0.000134 | BMP4,CDC25A,CXCL3,CXCL8,EGR1,FOS,JUN,KLF4,PTGS2 | 360 (12) |
| CDK5 | | kinase | | -1.214 | 0.0212 | CASP9,CXCL2,JUN,PMAIP1,PPP1R1B | |
| L-histidine | | chemical - endogenous mammalian | | -1.219 | 0.00207 | ATF3,DDIT4,EGR1,PPP1R15A | 417 (16) |
| IL13 | | cytokine | | -1.256 | 3.59E-09 | CXCL3,CXCL8,CXCR4,EGR1,FAM162A,FAS,FLOT1,FXYD3,GSN,HK1,HRH1,IFNGR2,IL1R2,IL1RN,JUN,KITLG,KLF6,KLK6,MAOA,MCL1,MSMO1,MT1X,MTSS1,MUC2,PAPSS1,PHLD | 457 (19) |
| Esrra | | transcription regulator | | -1.265 | 0.0115 | ACAT1,ACO1,ATP5MC1,ATP5MC3,CPT1A,GALM,PCK2,PDK4,PGM1,PPP1R1B,PTGS2,REN | |
| FBXW7 | | enzyme | | -1.27 | 0.0026 | CEBPD,DGAT2,DNAJA1,HMGCR,HSPB1,HSPH1,JUN,PPARG,RHOB,SQLE | 434 (18) |
| miR-125b-5p (and other miRNAs w/seed CCCUGAG) | | mature microRNA | | -1.272 | 0.0000824 | AJUBA,CASP7,CBX7,CCNA2,CD44,CDC25A,CDK6,CYP1A1,GPR160,ID2,IL1RN,JUN,SH2B3,TSPAN8,VSIR | 363 (13) |
| STAT6 | | transcription regulator | | -1.274 | 7.37E-13 | ,GBP3,GNA14,HIPK2,HMGCR,HMGCS2,HRH1,IFI44,IFIH1,IFIT3,IFITM3,IL1RN,IMPDH1,IRF1,IRF7,IRS2,KDM6B,LGALS3BP,LIF,LMO7,MYB,NCOA3,NEDD9,OAS3,PDK4,PKIB,PLD1,F | 525 (19) |
| mir-155 | | microRNA | | -1.287 | 0.00894 | BCL6,CD68,CEBPA,CLDN1,CXCL8,DNAJB1,DUSP5,EDN1,IFIT3,IRF7,MYB,PPARG,PTGS2 | 361 (20) |
| SPARC | | other | | -1.291 | 0.000339 | AKR1C3,CEBPD,CLDN1,CYP1A1,DVL1,GPD1,LOXL4,LRP8,NIPAL1,PAG1,SLC22A5,TMC5,TRIM25,WNK4,ZHX2 | |
| L-triiodothyronine | | chemical - endogenous mammalian | | -1.299 | 6.07E-09 | A,DNMT3A,DUSP1,DUSP5,DVL1,EDN1,EFNA1,EGR1,ENG,FAS,FOS,FUOM,GDF15,GLI2,GNA14,GPCPD1,ID2,IGFBP6,KLF9,LCN2,LGALS3,LGALS4,ME1,MSMO1,MYOM3,NR1D1,N | 445 (23) |
| oleic acid | | chemical - endogenous mammalian | | -1.302 | 0.00000368 | ACSL1,ATF3,BBC3,CD68,CLU,CXCL8,DGAT2,DUSP1,EDN1,EGR1,FOS,GADD45A,GADD45B,GDF15,HMGCR,LCN2,PDK4,PNPLA2,PTGS2,REN | 446 (20) |
| SYK | | kinase | | -1.304 | 0.000125 | AKAP12,BIRC3,CD44,CFTR,CXCL8,CXCR4,EDN1,FOSL1,GADD45A,GLI1,IRF1,PIM1,RASSF6,TSC22D1,TSC22D3,USP53 | 503 (21) |
| calphostin C | | chemical - kinase inhibitor | | -1.318 | 0.0000223 | AXIN2,CDC25A,CLU,CXCL8,DKK1,DUSP1,ETS1,FOS,MAOB,MUC2,PPARG,PTGS2,SCARB1 | 457 (24) |
| PTGER4 | | G-protein coupled receptor | | -1.32 | 3.65E-08 | DC3,CCNG2,CD44,CDK6,CXCL8,CXCR4,EDN1,EGR1,GAB1,GLIS3,HERC6,IFI35,IFIH1,IRF1,IRF7,MAFF,NOX1,PARP14,PTGS2,RNF213,RNF24,RUNX2,SLFN5,TBC1D4,TRIM21,ZFP | 428 (21) |
| APOE | | transporter | | -1.321 | 3.43E-13 | A1,CPT1A,CTSS,CXCL3,CYP3A5,ECM1,EDN1,EGR1,F2R,F2RL1,FOS,HMGCR,HSPA1A/HSPA1B,HSPG2,IGFBP6,IL1RN,JUN,KLF4,LCN2,LIMS2,LIPG,LRP8,MYO1B,NPNT,NQO2,PI | 511 (22) |
| PPARGC1B | | transcription regulator | | -1.331 | 0.0243 | CKB,FBXO32,GRB7,HMGCR,LSS,PDK4,RUNX2,SQLE | |
| LDLR | | transporter | | -1.331 | 0.0000337 | AURKA,AXIN2,BGN,CCNA2,CD55,CD68,CEBPA,DUSP1,E2F7,EGR1,ELOVL6,GALM,HMGCR,HSPG2,IL1RN,IRF7,LCN2,PCK2,PKIB,PPARG,PPP1R15A,PSME2,RUNX2,SC5D,SCAR | 475 (18) |
| pyrrolidine dithiocarbamate | | chemical reagent | | -1.333 | 0.0000281 | ANXA1,ASS1,BIRC3,CFTR,CXCL1,CXCL2,CXCL3,CXCL8,CXCR4,IRF1,IRF7,MMP7,NOX1,PLD1,PPARG,PSMB8,PTGS2 | 411 (17) |
| 4-phenylbutyric acid | | chemical - endogenous mammalian | | -1.335 | 0.0053 | ATP2A3,CASP9,CXCL8,DNAJB9,ERN1,GSN,MXD1,PADI2,PEX11A,PTGS2,SERPINA1 | 497 (21) |
| valproic acid | | chemical drug | | -1.336 | 1.85E-09 | ,EDN1,ELOVL6,ERN1,ETV6,FAS,GADD45B,HMGCS2,ID2,IDI1,IGSF1,JUN,JUND,LIMA1,LRP8,MCL1,MRPS2,MSMO1,MYOF,NFYA,NR0B2,OSBPL5,P3H2,PKIB,PNPLA2,PPARG,PSM | 593 (23) |
| EP400 | | other | | -1.342 | 0.0158 | CCNA2,CCNF,CDC25A,E2F7,INCENP,PPARG | |
| NFIL3 | | transcription regulator | | -1.342 | 0.0125 | FAS,GADD45A,GADD45B,ID2,PTGS2 | |
| STS | | enzyme | | -1.342 | 0.000622 | CD68,CEBPD,EEF1A2,HSPA1A/HSPA1B,IGFBP2,PNPLA2,PPARG | |
| DUSP5 | 0.488 | phosphatase | | -1.342 | 0.00234 | ALDH1A3,DUSP2,EGR1,ID2,ZFP36 | |
| JAK inhibitor I | | chemical - kinase inhibitor | | -1.342 | 0.0107 | CXCL1,CXCL2,CXCL3,PTGS2,SGK1 | |
| FUS-DDIT3 | | fusion gene/product | | -1.342 | 0.00655 | CXCL8,IGFBP6,LCN2,NPAS2,STX1A,TSC22D3 | |
| immethridine | | chemical reagent | | -1.342 | 0.0078 | CD44,CTSS,CXCL3,LCN2,SERPINA3 | |
| SP2509 | | chemical reagent | | -1.344 | 0.000723 | ,CDK6,CRABP2,DHRS3,DNMT3A,EIF2AK2,ELOVL6,FLRT3,FRYL,GATA6,GLI2,HMGA2,IGSF9,INPPL1,KLF4,MAP4K4,MICALL2,MYO1B,NPNT,PPP2R5B,SGK1,SMAD3,SOX12,SRI,S | |
| HNF4A | -0.821 | transcription regulator | | -1.346 | 1.32E-11 | 5,GJB2,GPR160,GRHL3,GRIN2D,GSN,GSTA4,GYG1,HBEGF,HGD,HHLA2,HLA-B,HLA-C,HNF4A,HNF4G,HSPA4L,HSPH1,IFITM2,INCENP,ITPRID2,JUN,KDM8,KRT7,LCN2,LDHA,LG | 517 (22) |
| triptolide | | chemical drug | | -1.355 | 3.29E-08 | AKAP12,BBC3,CAV1,CDKN2B,CXCL1,CXCL8,DPP4,DUSP1,EFNA1,EMP1,ENG,F2RL1,FAS,HBEGF,HPGD,IGFBP2,JUN,KITLG,MCAM,MCL1,MMP7,PMAIP1,PTGS2,SOX4,TJP1 | 489 (23) |
| PTGER2 | | G-protein coupled receptor | | -1.373 | 0.013 | AURKA,CCNA2,CKAP2L,CXCL8,CXCR4,ECM1,EGR1,IL1R2,PIM1,PTGS2,TBL3 | |
| TAB1 | | enzyme | | -1.385 | 0.00015 | BIRC3,CXCL8,IFIH1,IRF7,JUN,MCL1,PTGS2,RGS2 | 448 (23) |
| CBL | | transcription regulator | | -1.387 | 0.019 | CDKN2B,CFTR,CXCL2,CXCL3,FGFR2,FOS,RUNX2 | |
| PD 153035 | | chemical drug | | -1.399 | 0.000393 | CXCL8,FOS,KRT13,KRT20,MCL1,PTGS2 | 433 (20) |
| TMBIM6 | | other | | -1.412 | 0.0123 | DNAJB9,ERN1,MAP1LC3B,SEC61A1 | |
| EFNA4 | | kinase | | -1.414 | 0.000865 | CRABP2,FOSL1,IRF1,IRF7,ITGB6,KRT13,KRT7,NT5E | |
| RHO | | G-protein coupled receptor | | -1.414 | 0.00309 | B2M,CASP7,CD44,CEBPD,CTSS,FOS,IMPDH1,NT5E,SGK1 | 319 (7) |
| EFNA3 | | kinase | | -1.414 | 0.00102 | CRABP2,FOSL1,IRF1,IRF7,ITGB6,KRT13,KRT7,NT5E | |
| methylmercury | | chemical toxicant | | -1.414 | 0.015 | CDK6,CYP1A1,EGR1,ID2,PLD1,RHOB,SLC16A1,TXNIP | |
| epigallocatechin-gallate | | chemical drug | | -1.417 | 0.0000268 | 1,CASP9,CCN1,CEBPA,COL18A1,CXCL1,CXCL3,CXCL8,EDN1,EGR1,FAS,FGF19,FOS,FOSB,FOSL1,GADD45A,GADD45B,GBP3,GJB2,GLI1,IRF1,JUN,JUND,PRKCA,PTGS2,REL,SI | 453 (22) |
| rosiglitazone | | chemical drug | | -1.429 | 2.99E-11 | 3,EDN1,EGLN1,ELOVL6,ETS1,FOS,FOSL1,GPD1,GYG1,HCAR1,HMGCS2,HPGD,IFIT3,IGF2,IGFBP2,IGFBP6,IL1RN,INSIG1,IRF1,IRF7,IRS2,KLF4,KRT20,LCN2,LDHA,MAT2A,NOX1, | 378 (20) |
| HDAC4 | | transcription regulator | | -1.45 | 0.00342 | ATF3,FOS,FOSL1,GADD45A,HDAC9,HMGCR,IFITM3,JUN,LDHA,PRKCA,PTGS2,RGS2,RUNX2,TNFRSF12A,TSPAN13 | 433 (19) |
| baicalein | | chemical drug | | -1.455 | 0.0196 | CXCL8,CYP1A1,FOS,JUN,MCL1,PTGS2 | |
| miR-24-3p (and other miRNAs w/seed GGCUCAG) | | mature microRNA | | -1.461 | 0.0212 | BBC3,CCNA2,DUSP1,PMAIP1,SMAD3 | |
| heme | | chemical - endogenous mammalian | | -1.463 | 0.00782 | CD55,CLDN1,CXCL2,CXCL3,LCN2,LIF,SCARB1,TJP1 | 463 (18) |

Supplementary Table 4. Upstream regulator analysis based gene set enrichment analysis (GSEA) of differentially expressed genes (DEGs) after ATRA+PDT compared to PDT

| Upstream Regulator | Expr Log Ratio | Molecule Type | Predicted Activation State | Activation z-score | p-value of overlap | Target Molecules in Dataset | Mechanistic Network |
|---|---|---|---|---|---|---|---|
| miR-203a-3p (and other miRNAs w/seed UGAAAUG) | | mature microRNA | | -1.471 | 0.0107 | CDK6,F2RL1,KLF4,PRKCA,RUNX2 | |
| tyrphostin AG 1478 | | chemical - kinase inhibitor | | -1.493 | 0.00288 | CAV1,CXCL8,CXCR4,EDN1,FOS,HBEGF,MCL1,PTGS2,RUNX2,S100A10 | 417 (19) |
| pitavastatin | | chemical drug | | -1.495 | 0.000672 | HMGCR,IDI1,LSS,PTGS2,SCARB1,SQLE | 248 (10) |
| STEAP3 | -0.4 | transporter | | -1.508 | 0.0000728 | ACER2,CD44,ERRFI1,GCNT2,NEDD9,PIK3R1,S100A10,TGFB2,TGM2,TNFRSF12A,ZYX | |
| GLI1 | -0.756 | transcription regulator | | -1.518 | 5.11E-08 | 4,CEP70,CLU,DKK1,FAS,FOXL1,GATA6,GLI1,GLI2,HOXA5,ID2,IGF2,IGFBP6,IL1R2,INSIG1,JAG2,KLF4,NDRG2,NPDC1,NQO2,PGM2L1,PIM1,REG4,RHOB,RUNX2,SH2B3,TSC22D1 | 489 (19) |
| RPTOR | | other | | -1.521 | 0.00437 | BCL3,CDK6,EGLN1,HMGCR,IDI1,IFI44,JUN,LDHA,LSS,MSMO1,PPP1R3B,RUNX2,SC5D,SQLE | 485 (18) |
| Ro41-5253 | | chemical reagent | | -1.528 | 0.000672 | CRABP2,IRF1,PPARG,RARG,SMPD3,TGFB2 | 305 (11) |
| PLA2G10 | | enzyme | | -1.536 | 0.0232 | BMP4,DHRS3,FAS,KDM6B,KRT13,NABP1,PHLDA1,PPP1R10,PTGS2,TRAF4 | |
| luteolin | | chemical drug | | -1.546 | 0.0208 | CDK6,CYP1A1,FAS,FOS,HNF4A,JUN,PTGS2 | |
| diphenyleneiodonium | | chemical reagent | | -1.55 | 0.00473 | ATF3,CXCL2,CXCL3,CXCL8,DUSP2,EGR1,FOS,NOX1,PTGS2 | 433 (19) |
| tazemetostat | | chemical drug | | -1.553 | 0.000872 | K6,CRABP2,DHRS3,DNMT3A,EIF2AK2,ELOVL6,FLRT3,FRYL,GATA6,GLI2,HMGA2,IGSF9,INPPL1,KLF4,MAP4K4,MICALL2,MYO1B,NPNT,PPP2R5B,SGK1,SMAD3,SOX12,SRI,STEA | |
| mir-30 | | microRNA | | -1.566 | 0.00047 | ARID5B,BBC3,GADD45A,KLF9,NEDD4L,RUNX2,SH2B3,STK39,UGT8 | |
| 15-deoxy-delta-12,14 -PGJ 2 | | chemical - endogenous mammalian | | -1.576 | 3.27E-08 | CAV2,CEBPD,CXCL2,CXCL8,DNAJB1,EGR1,EHF,FOS,GADD45A,GDF15,GSN,HSPB1,IRF1,JUN,KLF4,MAP1LC3B,MCL1,MXD1,MYB,NOX1,PLAAT4,PPARG,PTGS2,REL,RUNX2,SC | 427 (17) |
| miR-133a-3p (and other miRNAs w/seed UUGGUCC) | | mature microRNA | | -1.586 | 0.0125 | CXCL3,KRT7,MALAT1,MCL1,RUNX2 | |
| SMARCA5 | | transcription regulator | | -1.604 | 0.00468 | ALDH1B1,ASS1,B2M,BCL6,CD44,CELSR1,EPHA2,FAS,IFITM2,IRF7,PHLDA1,PLAC8,PLXND1,UNC93B1 | 199 (7) |
| N-nitro-L-arginine methyl ester | | chemical drug | | -1.614 | 0.00143 | ACO1,ATF3,BGN,CD68,CXCL2,CXCL3,CXCL8,FAS,HMGCR,INSIG1,PDK4,PTGS2,REN | 407 (20) |
| LDB1 | | transcription regulator | | -1.616 | 0.0000359 | 1,ANXA1,AXIN2,BBC3,CASP4,CGAS,CLDN2,COL18A1,DPP4,ENG,F2R,GSN,HOXA5,IFIH1,IRS2,MDK,NT5E,PAG1,PATZ1,PLSCR1,SERPINE2,SLC16A1,SMAD3,TGFB2,TRIM25,TXI | |
| LMO2 | | transcription regulator | | -1.616 | 0.0000384 | 1,ANXA1,AXIN2,BBC3,CASP4,CGAS,CLDN2,COL18A1,DPP4,ENG,F2R,GSN,HOXA5,IFIH1,IRS2,MDK,NT5E,PAG1,PATZ1,PLSCR1,SERPINE2,SLC16A1,SMAD3,TGFB2,TRIM25,TXI | |
| EWSR1-FLI1 | | fusion gene/product | | -1.623 | 0.0162 | ARID5B,CAV1,CCN1,EDN1,HPGD,NFKBIZ,SLFN5,TGFBR2,TRNP1 | |
| metribolone | | chemical reagent | | -1.626 | 2.65E-15 | R1,ERRFI1,ETV1,FHL2,FOS,GDF15,GTPBP2,HES6,HK1,HMGCR,HMGCS2,HSPH1,IDI1,IFITM2,IRS2,LDHA,LRIG1,MAOA,ME1,MMP7,MRPL34,NEDD4L,NQO2,PROS1,PYGB,REG4,S | 552 (21) |
| simvastatin | | chemical drug | | -1.626 | 0.0000191 | 1,ANXA1,CCN1,CD55,COTL1,CXCL2,CXCL3,CXCL8,CYP2B6,EDN1,F2R,FOS,HK1,HMGCR,IRF1,KLF4,LCN2,MAP1LC3B,MPP6,NRIP1,PTGS2,RHOB,RHOU,SCARB1,SDC4,ST6GAL | 304 (19) |
| ARHGDIG | | other | | -1.633 | 0.000153 | ALCAM,ANXA1,EDN1,KANK1,PADI1,RGS2 | |
| ACSS2 | -0.621 | enzyme | | -1.633 | 0.000518 | ACAA1,DGAT2,HMGCR,LSS,PPARG,SCARB1 | 338 (13) |
| (-)-norephedrine | | chemical drug | | -1.633 | 0.0000467 | HMGCR,IDI1,LSS,MSMO1,NPC1,SQLE | 195 (8) |
| dihydrotestosterone | | chemical - endogenous mammalian | | -1.635 | 1.92E-09 | ,GPD1,GSN,HMGCR,HOXA5,HPGD,HSPA1A/HSPA1B,HSPB1,ID2,IGFBP2,IQGAP2,IRS2,JUN,KITLG,LAMA5,LDHA,MAFF,MCAM,ME1,MMP7,NEDD4L,PDE9A,PHF21A,PHLDA1,PMA | 530 (23) |
| MYCN | | transcription regulator | | -1.646 | 0.00403 | 2,ABCB10,ABCC1,ALDH1A1,AMOTL2,ATXN2,B2M,CAV1,CCNG2,CLU,COL18A1,DNMT3A,DUSP2,FGFR2,GATA6,ID2,LDHA,MAT2A,PMAIP1,POLR2L,S100A10,SLC16A1,SORD,TGM | 454 (13) |
| MAP2K5 | | kinase | | -1.646 | 0.000000935 | ACSS2,CXCL8,CYP24A1,ELOVL6,FOS,HMGCR,IDI1,INSIG1,JUN,LSS,MSMO1,PTGS2,SQLE | 379 (21) |
| BMP10 | | growth factor | | -1.648 | 0.00019 | ALDH1A3,ANGPT1,CD55,ECM1,ENG,FGFR2,FOS,GLI1,HPGD,ID2,LCN2,PHLDA1,SERPINA3,SERPINE2 | 257 (7) |
| TSC2 | | other | | -1.652 | 1.53E-08 | ,ANXA1,ATF3,AXIN2,BCL3,CASP9,CD68,EGLN1,EGR1,FOS,GSN,HMGCS2,HSPB1,IFI44,IFITM3,IRS2,LGALS3,MAOB,MCL1,MUC2,PDK4,PGM2L1,PHLPP1,PKIB,PPP1R3B,PRKCA, | 500 (23) |
| 26s Proteasome | | complex | | -1.661 | 0.000478 | BAG3,BCL3,BIRC3,CCNA2,CD68,CDC25A,CXCL3,CXCL8,EREG,FOSL1,HBEGF,IFI35,MAP1LC3B,PRKCA,PTGS2,TGM2,TRAF4,TXNIP | 476 (21) |
| RPSA | | translation regulator | | -1.664 | 0.00295 | CXCL3,DUSP1,DUSP2,GBP3,PTGS2,REL | 505 (20) |
| N-[N-(3,5-difluorophenacetyl-L-Ala)]-S-phenylglycine t-butyl ester | | chemical - protease inhibitor | | -1.664 | 0.0443 | CXCL1,CXCL2,CXCR4,REL,RND3,S100A4 | |
| apicidin | | chemical - endogenous non-mammalian | | -1.664 | 0.000518 | ATP2A3,CXCL8,FAS,GSN,ROR1,TGM2 | |
| SIN3A | | transcription regulator | | -1.673 | 0.00309 | BCL6,CCNG2,CYP1A1,FBXO32,GADD45B,KLF6,PTGS2,TGFBR2,TXNIP | 412 (14) |
| GAPDH | | enzyme | | -1.673 | 0.00189 | CXCL1,DUSP1,HLA-C,IFITM2,OAS3,TXNIP,WARS1 | |
| IL10RA | | transmembrane receptor | | -1.677 | 9.18E-10 | R,CTSE,CYP2S1,DDIT4,DPEP1,ECM1,EDN1,EHF,ENTPD5,ETV1,FAS,GNE,HPGD,IL15RA,IL1RN,IRF1,IRF7,KCNE3,KITLG,LCN2,MAOB,MYB,NDRG2,NR5A2,P3H2,PLAAT3,PROCR, | 273 (12) |
| docosahexaenoic acid | | chemical drug | | -1.681 | 0.000924 | ACSS2,CD44,CPT1A,CXCL8,CXCR4,DNAJB9,ELOVL6,ERN1,FAS,FOS,HPGD,PIK3R1,PNPLA2,PTGS2,SERPINH1,SORL1,TGFB2,TGFBR2,UGT1A10 (includes others) | 485 (22) |
| H-7 | | chemical - kinase inhibitor | | -1.683 | 0.000731 | CLCF1,CYP1A1,EGR1,FAS,FOS,PIM1,SMAD3,VDR | 482 (20) |
| NKX2-1 | | transcription regulator | | -1.698 | 0.00000467 | GR2,AQP5,BMP4,CDKN2B,CEBPA,CLU,DPP4,ESAM,FAM3D,FXYD3,GLCCI1,IGFBP6,KITLG,LCN2,MTUS1,MYB,PARD6B,QSOX1,RNASE4,ROR1,SEC14L2,TFF3,TGFB2,UGT8,WNT | |
| CD300LF | | other | | -1.709 | 0.000249 | ACSS2,CPT1A,CXCL8,GPD1,HK1,IL1RN,PCK2 | 257 (7) |
| miR-34a-5p (and other miRNAs w/seed GGCAGUG) | | mature microRNA | | -1.727 | 0.000812 | ATF3,AXIN2,CD55,CDC25A,CDK6,FAM221A,GSDMB,HID1,IFITM2,KLF4,LGALS3BP,MYB,PLAAT4,PPARG,TRPS1,TSPAN13 | 240 (7) |
| Sb202190 | | chemical - kinase inhibitor | | -1.73 | 0.0137 | CD55,CFTR,CXCL8,CYP1A1,DKK1,EGR1,FGFR2,FOS,PHLDA1,PTGS2,RHOB,SGK1,VDR | |
| rotenone | | chemical toxicant | | -1.746 | 0.0366 | BIRC3,CXCL2,DNAJB9,FOS,PTGS2,SLC16A1,TXNIP | |
| MYB | 0.316 | transcription regulator | | -1.749 | 0.0000075 | ATP2B1,AURKA,AXIN2,BIRC3,CDKN2B,CXCL16,CXCL2,CXCR4,ESAM,FUT8,IGF2,JUN,KITLG,KLF4,LCN2,LIF,MAT2A,METTL7B,MTSS1,MYB,NEDD9,PTGS2,SDC4,ZC3H12A | 419 (18) |
| glutathione | | chemical - endogenous mammalian | | -1.768 | 0.0484 | ABCC1,ASS1,ATF3,CA9,PTGS2,ZBTB10,ZBTB4 | |
| raloxifene | | chemical drug | | -1.774 | 8.36E-08 | 3,BMP4,CLK1,CLU,DDIT4,DUSP1,ENG,F2R,FLNB,GPRC5A,HSPA1A/HSPA1B,JUN,KLF6,KRT7,MXD4,MYO1B,NCOA3,PLK3,PTGS2,RAP1GAP,SLC22A5,SMAD3,SNCG,TGFB2,TGFE | 476 (23) |
| SS18-SSX1 | | fusion gene/product | | -1.783 | 0.00000242 | AXIN2,BBC3,CDC25A,DKK1,PMAIP1,SHCBP1 | 313 (7) |
| iron | | chemical - endogenous mammalian | | -1.801 | 0.0174 | CEBPA,GADD45A,GADD45B,LCN2,NFYA,PPP1R15A,SLC39A8 | |
| troglitazone | | chemical drug | | -1.802 | 3.85E-13 | P1A1,CYP2B6,CYP3A5,DGAT2,DHRS3,EGR1,ENTPD5,ETS1,FOS,FOSL1,GADD45A,GDF15,GPD1,HIPK2,JUN,KLF4,KRT13,KRT20,LOC102724788/PRODH,MAOB,MCL1,ME1,MYB, | 491 (20) |
| staurosporine | | chemical - kinase inhibitor | | -1.818 | 0.0000954 | BBC3,CFTR,CXCL8,CYP1A1,FAS,FOS,FOSL1,GATA2,IGF2,MCL1,NEDD9,PIM1,PTGS2,RCAN1,SERPINA3,SMAD3,VDR | 514 (23) |
| wortmannin | | chemical - kinase inhibitor | | -1.821 | 0.00724 | ATF3,CD44,CDKN2B,CEBPD,CKB,CXCL16,CXCL8,EGR1,F2R,FOS,GPD1,IRF1,IRS2,JUN,MCL1,MT2A,NT5E,PTGS2,SCARB1 | 497 (22) |
| HMOX1 | | enzyme | | -1.824 | 0.00141 | ANGPT1,CDK6,CXCL1,CXCL2,CXCL3,CXCL8,CYP1A1,ENG,FOS,IL1RN,MAP1LC3B,PROCR,PTGS2,TGFB2,ZFP36 | 454 (20) |
| BCL3 | 0.773 | transcription regulator | | -1.827 | 0.00092 | CXCL2,CXCL3,CXCL8,DUSP5,EGR1,FOS,ID2,IRF1,JUN,PDK4 | 431 (18) |
| FOXA2 | | transcription regulator | | -1.831 | 0.00864 | AGR2,CD55,CEBPA,CPT1A,CXCR4,DDC,DDIT4,HES6,HMGCS2,HNF4A,HPGD,MTUS1,MUC2,NR5A2,PIM1,PPARG,PROS1,RARG,SERPINA1,SPINK1,TBXAS1 | 418 (16) |
| mir-15 | | microRNA | | -1.838 | 0.0517 | CDK6,HSPA1A/HSPA1B,LATS2,MCL1,MYB,PIM1,PTGS2 | |
| lithium chloride | | chemical drug | | -1.841 | 0.000143 | ASS1,AXIN2,BCL3,BMP4,CD44,DKK1,FOS,GDF15,HMGCR,IDI1,KRT20,LSS,MT2A,MYB,PLD1,PPARG,PTGS2 | 436 (22) |
| Sn50 peptide | | chemical toxicant | | -1.845 | 0.00384 | BIRC3,CXCL2,CXCL8,FAS,IFNGR2,LIPG,MMP7,PPARG,PTGS2 | 413 (18) |
| mir-26 | | microRNA | | -1.854 | 0.000193 | CDK6,CHORDC1,CXCL3,HPGD,MCL1,PTGS2,STK39 | 198 (7) |
| KLF2 | | transcription regulator | | -1.857 | 1.99E-08 | CL3,BMP4,CAV1,CD44,CD55,CDKN2B,CEBPA,CXCL2,CXCL8,CXCR4,ECM1,EDN1,EFNA1,F2RL1,FUT4,GADD45A,IGF2,KITLG,LGALS9,MT2A,PLPP3,PPARG,PROS1,PTGS2,RUNX | 410 (16) |
| CST5 | | other | | -1.863 | 0.0000168 | 1/AKR1C2,ARHGAP29,CAV1,CD44,CEMIP,CEP170,CHD2,DNAJB4,DOCK5,EPN3,FER,GPRC5A,IGSF9,IQGAP2,ITPRID2,JMJD1C,NPAS2,NRP1,NT5E,PMAIP1,PPL,PYGB,RUNX2,S | |
| SREBF1 | | transcription regulator | | -1.887 | 1.38E-08 | LOVL6,FAS,FBXO32,HMGCR,HNF4A,HSPA1A/HSPA1B,IDI1,IL1R2,INSIG1,IRS2,LGALS3,LSS,MDK,MSMO1,NPC1,NR0B2,PCK2,PMPCA,PPARG,PTGS2,SC5D,SCARB1,SERPINA1, | 301 (16) |
| G protein alpha | | group | | -1.89 | 0.0943 | ANKS4B,CHPF,FOS,GNAZ,LYSMD4,QPCT,SPCS3,TMED3 | |
| NLRX1 | | other | | -1.89 | 0.0106 | BCL10,BIRC3,CARD10,IRF1,MAP3K14,PIK3R1,REL | |
| ATP7B | | transporter | | -1.89 | 0.003 | ELOVL6,HMGCR,IDI1,LSS,MSMO1,SHCBP1,SQLE | |
| PPP3R1 | | phosphatase | | -1.89 | 0.034 | AQP5,BAG3,BCL3,CLIC5,GBP3,H1-0,PDE9A | |
| TCF4 | | transcription regulator | | -1.892 | 0.00000175 | K6,CDKN2B,CSRNP1,CTSE,FAS,GBP3,GNE,GRB7,H1-0,ID2,IGF2,MCL1,NEDD9,PARP8,PLAC8,PML,PPIF,PRDX4,RUNX2,SDF2L1,SEC11C,SEC24D,SERINC5,SP140L,TAPBPL,TIA | 273 (7) |
| SFTPA1 | | transporter | | -1.912 | 0.000000216 | AGR3,AKR1C1/AKR1C2,CCN1,CXCL2,CXCL3,ECM1,EGR1,ERN1,FOS,FOSB,GABRP,HBEGF,HSPA6,LRIG1,PDK4,PHLDA1,PLAAT4 | 445 (19) |
| ZNF217 | | transcription regulator | | -1.912 | 0.00115 | AGR2,CDKN2B,CREB5,GPRC5A,MYO1B,RCAN1,SEC14L2,SEMA3A,SH3RF2,STRA6,ZHX2 | |
| THZ2 | | chemical - kinase inhibitor | | -1.912 | 0.000111 | FHL2,GLI2,HRH1,LIF,MCL1,RUNX2,SMAD3 | |
| mir-29 | | microRNA | | -1.93 | 0.00411 | CDK6,CLDN1,DKK1,DNMT3A,KLF4,MCL1,PIK3R1,RUNX2,SCARB1,TRAF4,ZFP36L1 | 371 (11) |
| Go 6976 | | chemical - kinase inhibitor | | -1.932 | 0.00427 | CD55,CXCL2,CXCL3,CXCL8,ETS1,FOS,PLD1,PRKCA,PTGS2 | 452 (23) |
| resolvin D1 | | chemical - endogenous mammalian | | -1.933 | 0.00134 | BCL6,CXCL2,CXCL3,CXCL8,CXCR4,JUN,PTGS2 | 424 (19) |
| UCN-01 | | chemical drug | | -1.934 | 0.00656 | CDC25A,CXCL8,MCL1,RHOB,XPA | 470 (21) |
| amino acids | | chemical - endogenous mammalian | | -1.936 | 0.0363 | FOS,FOSB,IGF2,JUN | |
| SUMO2 | | enzyme | | -1.937 | 0.000865 | CAV1,CAV2,CEACAM1,CXCL16,NR0B2,PIK3R1,RNASE4,TSC22D3 | 367 (7) |
| PAX7 | | transcription regulator | | -1.938 | 0.000814 | ASS1,CXCR4,F2RL1,FAM107B,ID2,MSLN,NPNT,PPARG,RNF128,SEMA3E,TOB1 | |
| fingolimod | | chemical drug | | -1.941 | 0.566 | CXCL2,FOSB,FOSL1,RUNX2 | |
| fludarabine | | chemical drug | | -1.949 | 0.0571 | CXCL2,FAS,IRF1,MCL1 | |
| RBM5 | | other | | -1.953 | 0.00000116 | ADAM9,ANXA1,ATP5MC3,BIRC3,IFITM3,INSIG1,MCL1,MYO1B,NCOA3,PIM1,SERPINH1,TSPAN6 | |
| BDNF | | growth factor | | -1.959 | 0.0000429 | R4,DUSP1,EGR1,ELOVL6,FLNB,FOS,FOSB,GNAI1,GPC4,HMGCR,HSPA4L,JUN,LCN2,MACROD1,MALAT1,PLXNB2,PPP1R1B,PTPN9,S100A10,SCARB1,SEMA3E,SERPINH1,SH3I | 469 (21) |
| dichlororibofuranosylbenzimidazole | | chemical toxicant | | -1.961 | 0.0123 | CXCL8,CYP1A1,CYP24A1,CYP3A5 | |
| PRDM16 | | transcription regulator | | -1.963 | 0.098 | GBP3,IFI44,IRF7,OAS3 | |
| SERCA | | group | | -1.964 | 0.0000795 | EGR1,FOSL1,JUN,ZFP36 | 420 (15) |
| fisetin | | chemical toxicant | | -1.964 | 0.149 | BIRC3,FOS,MCL1,PTGS2 | |
| taurine | | chemical - endogenous mammalian | | -1.964 | 0.122 | ABCC2,CEBPA,CPT1A,PTGS2 | |
| SIRT1 | | transcription regulator | | -1.967 | 0.000000285 | GADD45A,GBP3,GLI1,GLI2,HJURP,HMGCR,HOXA5,IFI44,IFIT3,IFITM3,IRF7,KDM2B,KIF13B,LGALS3,LGALS3BP,MAOA,MAT2A,MCL1,MGAT1,NEDD4L,NPNT,NR1D1,PARP12,PARI | 428 (21) |
| MIR101 | | group | | -1.969 | 0.00651 | DUSP1,MCL1,PTGS2,STMN1 | |
| naringenin | | chemical - endogenous non-mammalian | | -1.969 | 0.0368 | CPT1A,CXCL1,CXCL8,CYP1A1,PTGS2 | |
| PD184352 | | chemical drug | | -1.969 | 0.0241 | CXCL3,EGR1,FOS,MCL1 | |
| HDAC5 | | transcription regulator | | -1.97 | 0.0079 | ACSL1,AURKA,CPT1A,DAPK1,HMGCR,HMGCS2,JUN,KLF6,PPARG,RUNX2 | 504 (22) |
| TNIP1 | | other | | -1.972 | 0.14 | CXCL2,CXCL3,HLA-B,LCN2 | |
| EZR | | other | | -1.973 | 0.0268 | ATF3,DDIT4,FAS,IGFBP2,PTGS2 | |
| TRAF3 | | enzyme | | -1.977 | 0.165 | BIRC3,CXCL2,CXCL3,FAS,MAP3K14,MCL1,PPFIBP2 | |
| LY294002 | | chemical - kinase inhibitor | | -1.979 | 2.63E-15 | CL3,CXCL8,DUSP1,EGR1,EREG,ETS1,F2R,FAS,FER,FEZ2,FOS,FOSB,FOSL1,GDF15,GPD1,GSTA4,HBEGF,HMGCR,IDI1,IGF2,IGFBP2,IL1R2,IL1RN,IRF1,IRS2,JUN,LAMC2,LCN2, | 510 (20) |
| ARRB2 | | other | | -1.98 | 0.00863 | CXCL2,CXCL3,EGR1,GRK5,NFKBIZ,PTGS2 | 399 (17) |
| tosylphenylalanyl chloromethyl ketone | | chemical - protease inhibitor | | -1.981 | 0.0319 | BIRC3,CXCL2,CXCL8,PTGS2 | |
| SGPL1 | | enzyme | | -1.982 | 0.0363 | ABCC1,PPARG,PTGS2,SEC61A1 | |
| DNMT1 | | enzyme | | -1.982 | 0.472 | CYP24A1,IGF2,LRP8,PPARG,PTGS2 | |
| IL37 | | cytokine | | -1.982 | 0.0279 | CXCL2,CXCL3,CXCL8,IL1RN | |
| SL 327 | | chemical - protease inhibitor | | -1.982 | 0.0461 | BBC3,FOS,FOSB,NEDD4L | |
| fucoidin | | chemical reagent | | -1.982 | 0.0571 | CLDN1,CXCL8,IRF7,MCL1 | |
| mir-34 | | microRNA | | -1.984 | 0.000814 | BIRC3,CDC25A,CDK6,CXCL3,EMP1,FOSL1,MYB,PIP5K1A,PPP1R10,RUNX2,SEMA4B | 250 (10) |

Supplementary Table 4. Upstream regulator analysis based gene set enrichment analysis (GSEA) of differentially expressed genes (DEGs) after ATRA+PDT compared to PDT

| Upstream Regulator | Expr Log Ratio | Molecule Type | Predicted Activation State | Activation z-score | p-value of overlap | Target Molecules in Dataset | Mechanistic Network |
|---|---|---|---|---|---|---|---|
| N-acetyl-L-cysteine | | chemical drug | | -1.985 | 0.000163 | 3,BIRC3,CASP4,CASP9,CAV1,CCN1,CFTR,CXCL3,CXCL8,CXCR4,DUSP1,EDN1,EGR1,FAS,FBXO32,FOS,HBEGF,JUN,MAP1LC3B,NFKBIZ,PTGS2,RHOB,RUNX2,SLC16A1,TGFBR2 | 468 (20) |
| R-WIN 55,212 | | chemical reagent | | -1.986 | 0.0571 | CASP7,CASP9,CDK6,CXCL8 | |
| levothyroxine | | chemical - endogenous mammalian | | -1.987 | 0.306 | BBC3,FOS,GDF15,IGFBP6,PCSK6 | |
| NR1D1 | 0.662 | ligand-dependent nuclear receptor | | -1.987 | 0.122 | EGR1,FOS,ME1,NR1D1 | |
| STAT5B | | transcription regulator | | -1.993 | 0.00000474 | RD13B,BCL6,BIRC3,CASP4,CASP9,CDK6,CYP2B6,CYP3A5,DAPK1,DDC,DGAT2,FAS,FOS,GADD45A,GADD45B,GPD1,IRF1,MCL1,ME1,MTMR11,NIPAL1,NT5E,PCSK5,PDK4,PIM1,I | 484 (20) |
| anakinra | | biologic drug | Inhibited | -2 | 0.0165 | ABCC2,CXCL1,EDN1,PIM1,PTGS2 | |
| CBX7 | 0.635 | other | Inhibited | -2 | 0.0123 | CLDN1,CTSS,CXCL8,GDF15 | |
| ZBED2 | | other | Inhibited | -2 | 0.000593 | GATA6,IRF7,OAS3,PIM1 | 153 (7) |
| SPDEF | -1.01 | transcription regulator | Inhibited | -2 | 0.249 | BIRC3,CXCL8,PRKCA,SMAD3,STMN1 | |
| PPP1R15B | | phosphatase | Inhibited | -2 | 0.00144 | CHAC1,PPARG,PPP1R15A,SESN2 | |
| LRPAP1 | | other | Inhibited | -2 | 0.0319 | CXCL8,PTGS2,SERPINA3,SORL1 | |
| PPP2R5C | | other | Inhibited | -2 | 0.00818 | BCL3,BIRC3,CXCL8,REL | 354 (10) |
| S100A6 | | transporter | Inhibited | -2 | 0.0514 | ATF3,AURKA,CXCL2,CXCL8 | |
| JAG2 | -0.547 | growth factor | Inhibited | -2 | 0.122 | BCL6,CXCL1,CXCL3,IL1RN | |
| PAEP | | other | Inhibited | -2 | 0.0123 | CXCL1,CXCL16,EDN1,HBEGF | |
| NBEAL2 | | other | Inhibited | -2 | 0.00818 | CXCL16,CXCL2,CXCL3,GATA2 | 396 (7) |
| ADAMTS12 | | peptidase | Inhibited | -2 | 0.0363 | ALDH1A3,LCN2,MID1,TNFAIP2 | |
| tranilast | | chemical drug | Inhibited | -2 | 0.0176 | CXCL1,CXCL8,NRP1,TXNIP | |
| glutamyl-Se-methylselenocysteine | | chemical - endogenous mammalian | Inhibited | -2 | 0.00818 | CLU,EGR1,JUN,PSME1 | |
| AG490 | | chemical - kinase inhibitor | Inhibited | -2.004 | 0.0000213 | ACAT1,AKR1C3,BIRC3,CD68,CFTR,CXCL2,CXCL8,EDN1,FOS,HPGD,IGF2,IRF1,MCL1,PIM1,PTGS2,S100A10 | 484 (23) |
| miR-30c-5p (and other miRNAs w/seed GUAAACA) | | mature microRNA | Inhibited | -2.005 | 0.0000251 | ACVR1,BCL6,IRS2,JUN,KRT7,LOXL4,MAP4K4,MAT2A,NT5E,PGM1,PPARG,RAB27B,RUNX2,SH2B3,SLC12A4,STX1A,TGFBR2,TMED3,TNFAIP2,TRPS1 | |
| INSR | | kinase | Inhibited | -2.01 | 4.57E-10 | CXCR4,DGAT2,DUSP1,EDN1,EGR1,EREG,FOS,GADD45A,GATA6,GRB7,HBEGF,HMGCR,HPGD,IDI1,IFITM2,IGF2,INSIG1,KLF9,LAT2,LSS,MAOA,MAOB,MSMO1,MUC2,PDK4,PHLD | 489 (26) |
| miR-199a-5p (and other miRNAs w/seed CCAGUGU) | | mature microRNA | Inhibited | -2.032 | 0.0000655 | APOL1,BCAM,BGN,CELSR1,ETS1,GPRC5A,KLF4,LAMC2,LCN2,MMP7,NEDD9,PTGS2,TGFBR2 | |
| Nr1h | | group | Inhibited | -2.06 | 0.00000872 | ALDH1A1,ATP1B1,BCL3,CAV1,CXCL1,CXCL8,EDN1,ELOVL6,F2R,GLI1,IFIH1,LAT2,NFKBIZ,NPC1,NR5A2,PML,PTGS2,REN,SBNO2,SCARB1,SP110,SYT7 | 535 (22) |
| THRB | | ligand-dependent nuclear receptor | Inhibited | -2.111 | 0.0000856 | 3,CDK6,CEBPA,CFTR,CPT1A,CXCL2,CXCL3,CXCR4,DDC,DKK1,ENG,FOS,FOSL1,GSN,IGFBP2,JUN,KLF9,MACC1,ME1,MMP7,NR5A2,PDK4,PPARG,PTGS2,RUNX2,STRA6,UQCR | 459 (22) |
| MEOX2 | | transcription regulator | Inhibited | -2.121 | 0.000514 | ANGPT1,CXCL1,CXCL2,CXCL3,CXCL8,EDN1,EFNA1,HBEGF | 309 (7) |
| EPCAM | | other | Inhibited | -2.121 | 0.0000313 | ATF3,EGR1,FHL2,FOS,FOSL1,HMGA2,JUN,RUNX2 | 505 (21) |
| miR-182-5p (and other miRNAs w/seed UUGGCAA) | | mature microRNA | Inhibited | -2.137 | 0.000254 | CDK6,DDX60L,HERC6,MALAT1,MTSS1,NMI,RARG,SAMD9,SERPINE2,SLFN5,SP110 | |
| Ro31-8220 | | chemical - kinase inhibitor | Inhibited | -2.174 | 0.00139 | CD55,CLDN1,CXCL8,DUSP1,EGR1,FSTL3,PLD1,PTGS2 | 440 (20) |
| miR-27a-3p (and other miRNAs w/seed UCACAGU) | | mature microRNA | Inhibited | -2.183 | 0.0532 | BBC3,PMAIP1,PPARG,SMAD3,ZBTB10 | |
| RCAN1 | 0.7 | other | Inhibited | -2.183 | 0.00135 | CCNA2,CCNF,CXCL8,FOS,PTGS2,RCAN1 | 469 (22) |
| STK40 | | kinase | Inhibited | -2.186 | 0.0323 | CEBPA,CXCL1,CXCL8,FUT4,JUN,PPARG,PTGS2,WNT11 | |
| ethylene glycol tetraacetic acid | | chemical reagent | Inhibited | -2.194 | 0.0368 | BIRC3,CXCL8,EGR1,FOS,RCAN1 | |
| NRF1 | | transcription regulator | Inhibited | -2.2 | 0.122 | ATP1B1,CLDN1,CXCL8,IL1RN,TJP1,VDAC1 | |
| FBXO32 | 0.306 | enzyme | Inhibited | -2.2 | 0.0534 | ALDH1A3,AXIN2,CXCL2,CXCL3,DNAJB9,DNAJC3,GADD45A,PTGS2 | |
| HSPB1 | 0.966 | other | Inhibited | -2.2 | 0.00919 | BBC3,CCNA2,CXCL2,CXCL3,CXCL8 | 462 (21) |
| PS-1145 | | chemical - kinase inhibitor | Inhibited | -2.2 | 0.00183 | CXCL1,CXCL8,IRF1,PTGS2,SERPINA3 | 405 (16) |
| Bay 11-7082 | | chemical - kinase inhibitor | Inhibited | -2.201 | 0.00176 | CD44,CXCL1,CXCL2,CXCL3,CXCL8,IL15RA,MCL1,MMP7,PIM1,PTGS2,S100A4 | 407 (16) |
| mifepristone | | chemical drug | Inhibited | -2.206 | 0.000000838 | 1,CEBPD,CFTR,CLU,COL18A1,CYP3A5,DDIT4,DUSP1,EGR1,EHF,FAS,FOS,HBEGF,HPGD,IGF2,IRS2,JUN,KLF4,LAMA5,LAMC2,MYB,NCOA3,NQO2,PADI2,PTGS2,RAP1GAP,RGS2 | 545 (21) |
| MAF | | transcription regulator | Inhibited | -2.213 | 0.116 | ACSS2,HMGCR,IDI1,LSS,NT5E,SQLE | |
| GADD45A | 0.716 | other | Inhibited | -2.213 | 0.00367 | CXCL2,CXCL3,FOSB,JUN,MMP7 | 488 (18) |
| USP18 | | peptidase | Inhibited | -2.219 | 0.0165 | BBC3,IFIH1,IFITM3,IRF1,IRF7 | |
| DLK1 | | other | Inhibited | -2.232 | 0.00367 | GLI1,IRF1,IRF7,PPARG,REL | 433 (13) |
| ACKR2 | | G-protein coupled receptor | Inhibited | -2.236 | 0.0579 | EIF2AK2,IFI44,IFIT3,IRF7,OAS3 | |
| SIRT2 | | transcription regulator | Inhibited | -2.236 | 0.0078 | HMGCR,IDI1,LSS,SC5D,SQLE | |
| DBI | | other | Inhibited | -2.236 | 0.00000979 | ACSS2,ELOVL6,HMGCR,LSS,PPARG,SQLE | 94 (6) |
| PNPT1 | | enzyme | Inhibited | -2.236 | 0.000131 | CXCL8,EIF2AK2,IFI44,IFIH1,IFIT3,IRF7,LGALS3BP,PARP12,PARP14,RNF213 | 391 (12) |
| SU6656 | | chemical toxicant | Inhibited | -2.236 | 0.0268 | ATF3,EGR1,PHLDA1,PTGS2,ZFP36L1 | |
| PRDM1 | | transcription regulator | Inhibited | -2.271 | 5.49E-08 | ,CDK6,CLDN1,CRISPLD2,CXCL2,CXCL3,CXCR4,DDIT4,DUSP16,ECM1,F5,FOS,FOSL1,ID2,IRF7,MCL1,ME1,MT2A,MTMR11,PCSK6,PLAAT4,PLAC8,PMAIP1,PSMB10,PSMB8,SERP | 471 (19) |
| FOXM1 | | transcription regulator | Inhibited | -2.277 | 0.00189 | ANXA1,AURKA,AXIN2,CAV1,CCNA2,CCNF,CDC25A,FOS,FUT4,IL1RN,JUN,LDHA,MMP7,STMN1,TGFBR2,TJP1 | 462 (19) |
| U0126 | | chemical - kinase inhibitor | Inhibited | -2.278 | 4.03E-21 | EIF2AK2,ELOVL6,EMP1,EPHA2,EREG,ERRFI1,ETS1,ETV1,FGFR2,FOS,FOSB,FOSL1,GADD45B,GDF15,GLI1,GSN,GSTA4,HBEGF,HMGCR,HPGD,IDI1,IL1R2,IRF1,JDP2,JUN,JUND | 488 (19) |
| mir-8 | | microRNA | Inhibited | -2.284 | 6.68E-08 | 3,CCNG2,CDC25A,CDK6,CLDN1,CLDN12,DNAJC3,ERRFI1,ETS1,FGFR2,GRB7,GSE1,HOXA5,ITGB6,KITLG,KLF4,LATS2,MSLN,MYOF,PIP5K1A,PMAIP1,RNF128,SERPINE2,TGFB2 | 376 (7) |
| PSMB11 | | peptidase | Inhibited | -2.326 | 0.00283 | CAV1,CLDN2,CREB5,CTSS,CXCL3,GADD45A,GADD45B,HSPA1A/HSPA1B,LGALS3,NTN4,PTGS2,RHOB,TGFB2,TNFRSF12A | |
| TFRC | | transporter | Inhibited | -2.335 | 0.000026 | ATF3,BBC3,CCNA2,CHAC1,DDIT4,FOS,GADD45A,ID2,JUN,LDHA,ME1,PPARG,PPP1R15A,TGFB2,TNFRSF12A | 311 (7) |
| elaidic acid | | chemical - endogenous mammalian | Inhibited | -2.35 | 0.0000443 | ACSL1,ALDH1A1,CEBPA,DGAT2,DHRS3,ELOVL6,GNE,HMGCR,IDI1,LCN2,LPP,LSS,SC5D,SQLE,TFPI | |
| miR-155-5p (miRNAs w/seed UAAUGCU) | | mature microRNA | Inhibited | -2.361 | 0.00126 | BRPF3,CCN1,CLDN1,CXCL1,CXCL2,CXCL3,CXCL8,DNAJB1,ETS1,MYB,NT5E,PIK3R1,PLXND1,PMAIP1,PPL,PTGS2,RAB27B,SLC30A1 | 383 (18) |
| DUSP1 | 0.918 | phosphatase | Inhibited | -2.362 | 0.000000306 | APOL6,CXCL1,CXCL2,CXCL3,CXCL8,DKK1,DUSP1,FOSL1,GDF15,GJB2,IFIT3,IL1RN,IRF1,JUN,MXD1,NRP1,PEX11A,PIK3R1,PTGS2,TGFB2,ZFP36 | 437 (18) |
| ETV6-RUNX1 | | fusion gene/product | Inhibited | -2.372 | 0.0000166 | C42EP2,CXXC5,DDIT4,ENG,ID2,IFI35,IFIH1,IFIT3,IL15RA,IRF1,IRF7,JUND,KLF6,LAT2,NEDD4L,NMI,PAG1,PIM1,PLXNB2,PMAIP1,PML,PPP1R15A,S100A4,SH2B3,STMN1,SWAP70, | 155 (5) |
| BAPTA-AM | | chemical reagent | Inhibited | -2.376 | 0.000644 | AQP5,ATF3,CXCL8,CYP1A1,CYP3A5,EGR1,FOS,JUN,PTGS2 | 410 (19) |
| XBP1 | | transcription regulator | Inhibited | -2.392 | 0.000533 | CL2,CXCL8,DNAJB9,DNAJC3,ETS1,FAS,GOLPH3L,KLF4,MAP1LC3B,NCOA3,NUCB2,PDIA5,SDF2L1,SEC11C,SEC24D,SEC61A1,SEC61B,SEC61G,SERPINA1,SPCS3,SRPRB,SSR1 | 390 (16) |
| ZMPSTE24 | | peptidase | Inhibited | -2.418 | 0.0218 | ATF3,CXCL2,CXCL3,DDIT4,GADD45A,GADD45B | |
| SUMO3 | | other | Inhibited | -2.425 | 0.00984 | CAV1,CAV2,CEACAM1,CXCL16,PML,RNASE4 | |
| lapatinib | | chemical drug | Inhibited | -2.425 | 0.000393 | EREG,FOS,FOSB,FOSL1,HBEGF,JUN | 516 (23) |
| isoquercitrin | | chemical drug | Inhibited | -2.433 | 0.0000467 | HMGCR,HMGCS2,INSIG1,MSMO1,SC5D,SQLE | 72 (4) |
| SAFB2 | | other | Inhibited | -2.449 | 0.000859 | B2M,BBC3,FOS,HLA-C,HLA-E,JUN | |
| ELL2 | | transcription regulator | Inhibited | -2.449 | 0.0638 | DNAJB9,DNAJC3,FUT8,REXO2,SEC24D,SEC61A1 | |
| MNT | | transcription regulator | Inhibited | -2.449 | 0.000214 | BCL6,CCNG2,FBXO32,GADD45B,KLF6,TXNIP | |
| CISH | | other | Inhibited | -2.449 | 0.154 | CXCL3,DUSP2,EDN1,PPP1R15A,PTGS2,SORL1 | |
| miR-16-5p (and other miRNAs w/seed AGCAGCA) | | mature microRNA | Inhibited | -2.454 | 0.00000967 | DC25A,CDK6,CHORDC1,CLDN12,DMTF1,DNAJB4,GOLPH3L,HERC6,HSPA1A/HSPA1B,JUN,KITLG,MCL1,MYB,PPIF,PTGS2,RAB30,SERPINE2,SKAP2,SLC16A3,SLC38A5,SRPRB,T | |
| filgrastim | | biologic drug | Inhibited | -2.458 | 2.64E-12 | 2,F5,GADD45A,GATA6,GBP3,HIPK2,HPGD,IFI44,IFIT3,IL1R2,JUN,KLF6,KLF9,LDHA,LGALS3,MAFF,MYB,NHS,OAS3,PARP14,PATZ1,PDLIM1,PMAIP1,PRKAR2A,PRRG4,PSMB10,P | |
| PD98059 | | chemical - kinase inhibitor | Inhibited | -2.469 | 3.46E-23 | P1,ENG,EREG,ERRFI1,ETS1,F2R,FAM13A,FAS,FOS,FOSB,FOSL1,GADD45A,HBEGF,HMGA2,HPGD,IGF2,IL1RN,INSIG1,JUN,JUND,LAMA5,LDHA,LIPG,MAT2A,MCL1,MMP7,MUC2,I | 508 (17) |
| AIRE | | transcription regulator | Inhibited | -2.53 | 0.000628 | AKR1B10,AMBP,BMP4,EIF2AK2,HERC6,IFI44,IGF2,KLK6,MGAT1,NMI,PARP14,TFF3 | |
| MACROH2A1 | | other | Inhibited | -2.53 | 0.00215 | ALDH1A3,CDKN2B,EDN1,EGR1,ETS1,FGD3,FSTL3,GADD45A,IRF7,PPARG,TJP2 | 305 (13) |
| MAPK1 | | kinase | Inhibited | -2.563 | 5.07E-19 | B,GFOD1,GRB7,HLA-B,HLA-C,HLA-E,IFI35,IFI44,IFIH1,IFIT3,IFITM3,IGFBP6,IL1RN,IRF7,IRS2,ITPR2,JUN,JUND,LGALS3,LGALS3BP,LIF,MAOB,MCL1,NMI,NRP1,NT5E,OAS3,PARP | 430 (24) |
| mir-210 | | microRNA | Inhibited | -2.566 | 0.184 | EPB41L1,HOXA3,KLF6,SERTAD2,SMCHD1,STMN1,TP53I11 | |
| ZFP36 | 0.298 | transcription regulator | Inhibited | -2.595 | 0.0000389 | BIRC3,CTSS,CXCL2,CXCL3,FOS,FOSL1,IRF1,JUN,LATS2,LCN2,MXD1,PIM1,PTGS2,UQCRFS1,ZFP36 | 422 (16) |
| IKZF1 | | transcription regulator | Inhibited | -2.601 | 0.0000184 | CAV1,CAV2,CD68,CDK6,CXCL2,DEPP1,DKK1,DTX3L,DUSP5,EMP1,IFIT3,KITLG,KLF4,LGMN,MYO1B,NFKBIZ,PROS1,RNF19A,RNF213,S100A16,SERPINH1,SIPA1L2,SMAD3,TBC1 | 407 (8) |
| 2-amino-5-phosphonovaleric acid | | chemical - other | Inhibited | -2.607 | 0.00189 | BBC3,CAMKK1,CCDC28A,DUSP1,EGR1,FOS,FOSB,GADD45A,GADD45B,HDAC9,JUN,OTULINL,PHLDA1,PLK3,PTGS2,RGS2 | 438 (20) |
| infliximab | | biologic drug | Inhibited | -2.63 | 0.0422 | CXCL1,CXCL8,DDIT4,EDN1,MT1X,MYO1B,PTGS2 | |
| SH3TC2 | | other | Inhibited | -2.646 | 0.000402 | ACSL1,EGR1,HMGCR,IDI1,LSS,SC5D,SQLE | |
| SP600125 | | chemical - kinase inhibitor | Inhibited | -2.66 | 3.06E-08 | SPG4,CXCL1,CXCL16,CXCL2,CXCL3,CXCL8,CYP1A1,DDIT4,DKK1,DUSP1,EGR1,FAS,FLRT3,FOS,FOSB,GADD45A,HNF4A,JUN,LAMA5,MCL1,NEDD9,NOX1,NR0B2,PARP8,PCK2,F | 453 (19) |
| PTPRR | | phosphatase | Inhibited | -2.668 | 0.0218 | DHRS3,DNMT3A,ETS1,FOS,FOSL1,IFITM2,JUN,PLXNB2,PROCR,ROR1 | |
| ARID1A | | transcription regulator | Inhibited | -2.713 | 0.0000039 | ANXA1,AXIN2,BMP4,CEACAM1,CEACAM6,CXCL1,CXCL2,CXCL8,EREG,F2R,IRS2,UCA1 | 484 (20) |
| PKD1 | | ion channel | Inhibited | -2.758 | 0.0000237 | C,ALDH1A1,AMOTL2,ATP1B1,CCN1,CLCF1,CXCL16,CXCL2,DPEP1,FOSB,GLI2,HBEGF,HDAC9,HGD,MID1,NEDD9,NIPAL1,PML,PRAP1,QPCT,RASSF6,RUNX2,SMURF1,TBC1D4,Z | 301 (7) |
| PPARG | 0.347 | ligand-dependent nuclear receptor | Inhibited | -2.762 | 1.03E-11 | 1,GATA2,GDF15,GPD1,HCAR1,HIVEP2,HMGCR,HMGCS2,IGFBP6,INSIG1,IRS2,JUN,KLF4,KLF6,KLK6,KRT20,LOC102724788/PRODH,ME1,MUC13,NOX1,NR1D1,PCK2,PDK4,PEX | 505 (26) |
| herbimycin | | chemical - kinase inhibitor | Inhibited | -2.763 | 0.00213 | ATP2A3,CDK6,CXCL8,CXCR4,DUSP1,F2R,FOS,PTGS2 | 475 (22) |
| SAFB | | other | Inhibited | -2.813 | 0.0051 | B2M,BBC3,CXCL8,FOS,HLA-C,HLA-E,JUN,NEDD9 | 512 (19) |
| H89 | | chemical - kinase inhibitor | Inhibited | -2.882 | 0.00541 | ANXA1,AQP5,BMP4,CD55,CLCF1,CXCL8,DUSP1,EGR1,EREG,FOS,ITPR2,PLD1,PTGS2,REN,SIGMAR1,SMAD3 | 466 (22) |
| COL18A1 | -0.478 | other | Inhibited | -2.882 | 0.000026 | CCNDBP1,DDIT4,EDN1,EFNA1,EGLN1,ETS1,F2R,F2RL1,FOS,JUN,LDHA,MCL1,NRP1,PTGS2,VDAC1 | 483 (23) |
| bisindolylmaleimide I | | chemical - kinase inhibitor | Inhibited | -2.919 | 0.00000206 | BIRC3,CD44,CD55,CLCF1,CLDN1,CLU,CXCL8,EGR1,FOS,GDF15,JUND,MAT2A,MMP7,PLD1,PRKCA,PTGS2,RCAN1,TJP1,TJP2 | 461 (20) |
| mir-181 | | microRNA | Inhibited | -2.95 | 0.00473 | ATP2B1,CXCL8,GATA6,KLF6,MCL1,PTGS2,RUNX2,SH2B3,TGM2 | 363 (12) |
| MAX | | transcription regulator | Inhibited | -2.985 | 0.00132 | BCL6,CCNG2,CDC25A,CDKN2B,DUSP2,EDN1,FBXO32,GADD45A,GADD45B,ID2,KLF6,TXNIP | 225 (3) |
| thalidomide | | chemical drug | Inhibited | -2.985 | 0.0000133 | ANGPT1,BIRC3,CAV1,CXCL8,DKK1,FOS,GADD45B,GDF15,NRP1,PIM1 | 483 (21) |
| SP110 | 0.533 | transcription regulator | Inhibited | -2.985 | 0.00000144 | ATF3,BCL3,BMP4,CCN1,CDKN2B,CLU,ETS1,F2R,FER,IFIH1,IFIT3,IFITM3,MAOA,MCL1,OAS3,PLSCR1,PPARG,PRKCA,SERPINA1,SOX4,TIA1,TXNIP | |
| GMNN | | transcription regulator | Inhibited | -3 | 0.0202 | ACVR1,ANGPT1,DUSP1,DUSP5,EPHA2,GBP3,NRP1,TEAD2,TGFB2 | |
| SOX1 | | transcription regulator | Inhibited | -3 | 0.00917 | ACVR1,ANGPT1,DUSP1,DUSP5,EPHA2,GBP3,NRP1,TEAD2,TGFB2 | |
| SOX3 | | transcription regulator | Inhibited | -3 | 0.0162 | ACVR1,ANGPT1,DUSP1,DUSP5,EPHA2,GBP3,NRP1,TEAD2,TGFB2 | |
| SOX2 | | transcription regulator | Inhibited | -3.039 | 8.37E-13 | S,FUT4,FYN,GAB1,GAB2,GADD45B,GATA2,GATA6,GBP3,GLI2,GRHL3,GSN,ID2,IER5L,INPPL1,ITPR2,JUN,KANK1,KITLG,KLF4,KLF6,KRT7,LIMA1,LIMS2,NAAA,NR5A2,NRP1,OVOI | 462 (13) |
| Irgm1 | | other | Inhibited | -3.178 | 4.02E-08 | APOBEC3B,B2M,CCNA2,CGAS,CXCL16,FAS,GSDMD,HERC6,ID2,IFI35,IFI44,IFIH1,IFIT3,IFITM3,IRF1,IRF7,OAS3,TRIM21,TRIM25,TRIM56 | 293 (14) |
| SB203580 | | chemical - kinase inhibitor | Inhibited | -3.183 | 3.28E-08 | CXCL3,CXCL8,CYP1A1,DKK1,DNAJC3,DUSP1,EFNA1,EGR1,EHF,EIF2AK2,EREG,FAS,FOS,FOSB,GADD45A,GSTA4,IFI35,IFIT3,IL1RN,IRF1,IRF7,JUN,JUND,MAT2A,MUC2,MXD1,P | 508 (20) |
| rottlerin | | chemical - kinase inhibitor | Inhibited | -3.183 | 0.0000268 | APOBEC3B,BIRC3,CDK6,CLDN1,CXCL3,CXCL8,JUN,PLD1,PLSCR1,PTGS2,RGS2,TJP1,TJP2,ZFP36 | 498 (23) |
| IRGM | | enzyme | Inhibited | -3.268 | 0.0000291 | B2M,CGAS,DDX60L,HLA-B,IFI35,IFI44,IFIH1,IFIT3,IFITM3,PSMB8,SP140L | |

Supplementary Table 4. Upstream regulator analysis based gene set enrichment analysis (GSEA) of differentially expressed genes (DEGs) after ATRA+PDT compared to PDT

| Upstream Regulator | Expr Log Ratio | Molecule Type | Predicted Activation State | Activation z-score | p-value of overlap | Target Molecules in Dataset | Mechanistic Network |
|---|---|---|---|---|---|---|---|
| SREBF2 | | transcription regulator | Inhibited | -3.376 | 0.000000216 | ACSL1,ACSS2,BMP4,CXCL1,CXCL8,ELOVL6,HES6,HMGCR,IDI1,INSIG1,IRS2,LSS,MSMO1,PPARG,SC5D,SQLE,TFF3 | 491 (21) |
| Hdac | | group | Inhibited | -3.414 | 4.46E-08 | L6,CCNG2,CRABP2,CXCL8,CXCR4,CYP24A1,EDN1,EGR1,FBXO32,FOS,FOSL1,GADD45B,GLI2,HPGD,JUN,KLF6,KLF9,MT1E,MXD1,NPNT,NRP1,ONECUT2,SMPD3,STX1A,TGFBR | 458 (19) |
| NKX2-3 | | transcription regulator | Inhibited | -3.499 | 6.72E-13 | EDN1,EIF2AK2,F2RL1,GASK1B,GDF15,HLA-B,HLA-C,KLF4,LGMN,MMP7,PARP10,PARP12,PARP14,PLD1,PLSCR1,PMAIP1,PSMB8,PTGS2,RNF213,SAMD9,SEMA3A,SERPINE2,SH | |
| SCAP | | other | Inhibited | -3.541 | 0.000000653 | ACSL1,ACSS2,ATF3,CYP2B6,ELOVL6,HES6,HMGCR,HMGCS2,IDI1,INSIG1,LGALS3,LSS,MSMO1,PTGS2,SC5D,SQLE,TFF3 | 340 (11) |
| TRIM24 | | transcription regulator | Inhibited | -3.628 | 0.00000946 | CYP24A1,FLRT3,GBP3,HERC6,IFI35,IFI44,IFIH1,IFIT3,IFNGR2,IRF1,IRF7,LGALS3,LGALS3BP,NMI,PARP12,PLAC8,PSMB10,PSMB8 | 314 (14) |
| mibolerone | | chemical drug | Inhibited | -3.638 | 0.000157 | ACSL1,ALCAM,AMD1,AQP5,CEACAM1,CLU,DHRS3,DNAJA4,HLA-E,INCENP,IRF7,JUN,LSS,MAT2A,SLC22A5,SQLE,STMN1,TMED3 | |
| RC3H1 | | enzyme | Inhibited | -3.769 | 1.11E-09 | APOBEC3B,CD44,CLDN1,CXCL2,EDN1,FOSL1,HSPB1,IFI44,IFIT3,IFITM3,IRF1,MT2A,NRP1,OAS3,PLSCR1,SHFL,TRIM21,TRIM25,TRIM56 | 460 (12) |
| IL1RN | 0.525 | cytokine | Inhibited | -4.188 | 5.89E-12 | I,CD44,CHAC1,CTSS,CXCL1,CXCL3,CXCL8,CYP3A5,ERAP2,GSDMD,HDAC9,HERC6,IFI44,IFIH1,IFIT3,IRF1,IRF7,KLF6,LGALS9,LIF,NR0B2,OAS3,PMAIP1,PML,PTGS2,RARRES1,S | 471 (17) |
| phosphatidylinositol | | chemical - endogenous mammalian | | | 0.0428 | ACAT1,CXCL8 | |
| sphingosylphosphocholine | | chemical - endogenous mammalian | | | 0.00789 | FOS,JUN,TGM2 | 469 (18) |
| 1,2-dioctanoyl-sn-glycerol | | chemical reagent | | | 0.0217 | JUN,VDR | |
| octanoic acid | | chemical - endogenous mammalian | | | 0.0285 | CEBPA,DGAT2,PPARG | |
| 16,16-dimethylprostaglandin E2 | | chemical - endogenous mammalian | | | 0.00789 | CD44,FOS,NOX1 | 300 (12) |
| (+)-fluprostenol | | chemical drug | | | 0.0217 | NOX1,PPARG | |
| epoprostenol | | chemical - endogenous mammalian | | | 0.0408 | EGR1,FOS,PTGS2 | |
| 12(S)-hydroxyeicosatetraenoic acid | | chemical - endogenous non-mammalian | | | 0.00546 | PAG1,PPARG,PTGS2 | 445 (18) |
| ATP-gamma-S | | chemical reagent | | | 0.000384 | BTG1,CD55,CXCL2,CXCL8,CXCR4,EREG,IL15RA,IL1R2,NRP1,PTGS2,WARS1 | 524 (21) |
| zidovudine | | chemical drug | | | 0.0241 | BCL10,BCL3,CPT1A,CXCL8 | |
| acyclovir | | chemical drug | | | 0.049 | DUSP2 | |
| propylthiouracil | | chemical drug | | | 0.0169 | AKR1C3,CXCL2,GSN,IGF2,JUN,LOC102724788/PRODH,NRP1,NSMF,PHLDA1,QPCT,SPINK1 | |
| 3-aminoisobutanoate | | chemical - endogenous mammalian | | | 0.049 | CPT1A | |
| L-cysteine | | chemical - endogenous mammalian | | | 0.00789 | CXCL8,FOS,PCK2 | 424 (18) |
| L-methionine | | chemical - endogenous mammalian | | | 0.0208 | CLU,CXCL2,GADD45B,MAT2A,PCK2,RHOB,SPHK2 | |
| sorbitol | | chemical - endogenous mammalian | | | 0.00387 | CCN1,ERRFI1,SGK1,SORD | 171 (7) |
| glycerol | | chemical - endogenous mammalian | | | 0.0408 | GLI1,PPARG,SCARB1 | |
| pelargonidin | | chemical - endogenous non-mammalian | | | 0.049 | CYP1A1 | |
| amphotericin B | | chemical drug | | | 0.0343 | CXCL8,IL1RN,LCN2 | |
| ginsenoside Rb1 | | chemical - endogenous non-mammalian | | | 0.0285 | CXCL8,FOS,HBEGF | |
| saikosaponin A | | chemical reagent | | | 0.000343 | BMP4,CDKN2B,FOS,JUN | 422 (13) |
| epothilone B | | chemical drug | | | 0.0408 | FAS,MCL1,TBC1D1 | |
| troleandomycin | | chemical drug | | | 0.049 | CYP3A5 | |
| 3,7,12-trihydroxycoprostane | | chemical - endogenous mammalian | | | 0.049 | CYP3A5 | |
| 6beta-hydroxyestradiol-17beta | | chemical - endogenous non-mammalian | | | 0.049 | CYP2B6 | |
| estrone | | chemical - endogenous mammalian | | | 0.0343 | CYP2B6,IRS2,PIK3R1 | |
| taurolithocholate-3-sulfate | | chemical - endogenous mammalian | | | 0.049 | ABCC2 | |
| glycochenodeoxycholate | | chemical - endogenous mammalian | | | 0.0285 | BIRC3,HPGD,NR0B2 | |
| Srebp | | group | | | 0.0123 | ACSL1,ACSS2,FOS,NPC1 | |
| taurochenodeoxycholate | | chemical - endogenous mammalian | | | 0.00697 | HPGD,NR0B2 | |
| cholesterol sulfate | | chemical - endogenous mammalian | | | 0.00789 | FOSL1,JUND,NR1D1 | 241 (11) |
| CP-55940 | | chemical reagent | | | 0.0368 | CXCL8,GADD45A,GRK5,JUN,UBC | |
| fluoride | | chemical - endogenous mammalian | | | 0.00651 | FAS,FOS,GADD45A,RUNX2 | 234 (13) |
| dopamine D1 receptor | | group | | | 0.0428 | FOS,FOSB | |
| TEAD | | group | | | 0.0109 | CAV2,CCN1,CD44 | |
| Histone H2b | | group | | | 0.0428 | IRF1,PPP1R3B | |
| asialo GM1 ganglioside | | chemical - endogenous mammalian | | | 0.0217 | CXCL8,MUC2 | |
| thromboxane A2 | | chemical - endogenous mammalian | | | 0.049 | PTGS2 | |
| 2'-adenylic acid | | chemical - endogenous mammalian | | | 0.049 | TXNIP | |
| dibutyryl cGMP | | chemical reagent | | | 0.0217 | AQP5,PTGS2 | |
| Rp-8-CPT-cAMPS | | chemical - kinase inhibitor | | | 0.049 | SCARB1 | |
| ADP | | chemical - endogenous mammalian | | | 0.0343 | CXCL8,FOS,REN | |
| desoxycorticosterone | | chemical - endogenous mammalian | | | 0.0478 | CXCL2,FOS,PTGS2 | |
| monooleylphosphatidic acid | | chemical - endogenous mammalian | | | 0.00697 | EREG,PLPP3 | |
| pCPT-cAMP | | chemical - kinase inhibitor | | | 0.0478 | EGR1,FOS,JUN | |
| clobetasol propionate | | chemical drug | | | 0.0185 | ABCC1,CYP3A5,FAS | |
| beclomethasone dipropionate | | chemical drug | | | 0.049 | CYP3A5 | |
| bevacizumab | | biologic drug | | | 0.0478 | ANGPT1,CXCR4,NRP1 | |
| aflatoxin B1 | | chemical - endogenous non-mammalian | | | 0.0109 | CYP1A1,CYP3A5,IRS2 | |
| GON4L | | transcription regulator | | | 0.0285 | CCNA2,CCNF,MYB | |
| ADRA1 | | group | | | 0.0217 | EGR1,FOS | |
| dienogest | | chemical drug | | | 0.0217 | CXCL8,PTGS2 | |
| pivalyloxymethyl butyrate | | chemical drug | | | 0.049 | ATP2A3 | |
| (6)-gingerol | | chemical - endogenous non-mammalian | | | 0.0428 | CDK6,PTGS2 | |
| cyanuric acid | | chemical reagent | | | 0.049 | CXCL8 | |
| eicosa-11Z, 14Z-dienoic acid | | chemical - endogenous mammalian | | | 0.049 | PNPLA2 | |
| styrene | | chemical toxicant | | | 0.0217 | EGR1,HSPA1L | |
| peptide YY 3-36 | | biologic drug | | | 0.049 | FOS | |
| ATXN8OS | | other | | | 0.0217 | ANGPT1,TGFBR2 | |
| MEG3 | | other | | | 0.0285 | GDF15,JUN,PHLPP1 | |
| decabromobiphenyl ether | | chemical toxicant | | | 0.0135 | CYP2B6,CYP3A5 | |
| trans-cinnamaldehyde | | chemical drug | | | 0.0241 | CXCL8,EGR1,FAS,PTGS2 | |
| pyrvinium | | chemical drug | | | 0.0316 | AXIN2,ID2 | |
| 2-amino-5-azotoluene | | chemical toxicant | | | 0.00546 | ABCC2,CYP2B6,CYP3A5 | |
| hydroxamic acid | | chemical - other | | | 0.049 | CXCR4 | |
| JINK1/2 | | group | | | 0.041 | AXIN2,FAS,PTGS2,RHOB | |
| Sos | | group | | | 1.62E-08 | ,CRABP2,ELOVL6,FOS,IRS2,KLF6,KRT13,LGALS3BP,LGALS4,LRP8,LTBP3,MAOA,MAP4K4,MXD1,NRIP1,PHLDA1,PKN1,PLD1,PTGS2,RAP1GAP,SEMA3B,SEMA3E,SMURF1,STM | 515 (23) |
| Pdgfr | | group | | | 0.0232 | EGR1,EREG,HBEGF | |
| LTB4R/LTB4R2 | | group | | | 0.049 | CXCL8 | |
| Gαq | | group | | | 0.049 | PTGS2 | |
| NDPK | | group | | | 0.049 | CXCL8 | |
| Rsk | | group | | | 0.0135 | CXCL2,CXCL8 | |
| Foxo | | group | | | 0.00106 | ABCC2,CDKN2B,FBXO32,GADD45A,PMAIP1 | 388 (7) |
| CaMKII | | complex | | | 0.0207 | BIRC3,CD44,FOS,IRF1 | |
| Actin | | group | | | 0.0285 | ABCC1,CD44,LPP | |
| C4BP | | complex | | | 0.00546 | HMGCR,IDI1,MSMO1 | |
| RNA polymerase II | | complex | | | 0.0000425 | CXCL8,CXCR4,CYP1A1,CYP24A1,DKK1,DUSP1,FOS,FOSL1,GADD45A,GADD45B,GDF15,HNF4A,HSPA1A/HSPA1B,IL1RN,JUN,KITLG,NCOA3,NR5A2,PPARG,PRR15L,PTGS2,RND | 461 (13) |
| tyrosine kinase | | group | | | 0.0135 | CD44,CYP1A1 | |
| beta-glucuronidase | | group | | | 0.049 | PTGS2 | |
| nicotinic acetylcholine receptor | | complex | | | 0.0428 | FOS,JUN | |
| 15-LOX | | group | | | 0.00697 | MAOA,PPARG | |
| MKK3/6 | | group | | | 0.049 | CXCL8 | |
| Histone h3 | | group | | | 0.00000515 | PA,CSPG4,CXCL8,CYP3A5,DKK1,DUSP16,DUSP2,EFNA1,EGR1,FGFR2,FOS,GATA2,GATA6,GLI1,HNF4A,HOXA5,ID2,IFITM3,IGFBP2,IRF7,JUN,MEIS2,MTUS1,PMAIP1,PPARG,PR | 502 (21) |
| Fgf | | group | | | 0.0212 | FOS,JUN,LPCAT1,PTGS2,RUNX2 | |
| Gli | | group | | | 0.0123 | GLI1,JAG2,RUNX2,S100A4 | |
| MSK1/2 | | group | | | 0.0024 | DUSP1,FOS | 344 (7) |
| adenosine deaminase | | group | | | 0.049 | CXCL3 | |
| Erm | | group | | | 0.0217 | CAV1,CD44 | |
| ETS | | group | | | 0.0207 | EGR1,ENG,FOS,HPGD | |
| growth factor receptor | | group | | | 0.049 | FOS | |
| N-cor | | group | | | 0.00143 | ACSL1,AXIN2,CDK6,CTSS,CXCL2,DHRS3,ELOVL6,HBEGF,HIVEP2,PTGS2,RHOB,SGK1,TBC1D2 | 451 (13) |
| Ctnna | | group | | | 0.0217 | AXIN2,CXCR4 | |
| desmethoxyyangonin | | chemical - endogenous non-mammalian | | | 0.049 | CYP3A5 | |
| dihydrokawain | | chemical - endogenous non-mammalian | | | 0.049 | CYP3A5 | |

Supplementary Table 4. Upstream regulator analysis based gene set enrichment analysis (GSEA) of differentially expressed genes (DEGs) after ATRA+PDT compared to PDT

| Upstream Regulator | Expr Log Ratio | Molecule Type | Predicted Activation State | Activation z-score | p-value of overlap | Target Molecules in Dataset | Mechanistic Network |
|---|---|---|---|---|---|---|---|
| Stat1 dimer | | complex | | | 0.00354 | EIF2AK2,IRF1,PIM1 | 153 (7) |
| Stat1-Stat2 | | complex | | | 0.0109 | EIF2AK2,IRF1,IRF7 | |
| Smad2/3-Smad4 | | complex | | | 0.0478 | CDKN2B,HMGA2,IRF7 | |
| Integrin | | complex | | | 0.0478 | FOS,JUN,TGFB2 | |
| Bcl9-Cbp/p300-Ctnnb1-Lef/Tcf | | complex | | | 0.00789 | AXIN2,CD44,MMP7 | |
| rifamycin SV | | chemical drug | | | 0.049 | BCL6 | |
| 2,2',4,4',5,5'-hexabromodiphenyl ether | | chemical reagent | | | 0.049 | CYP1A1 | |
| endrin | | chemical toxicant | | | 0.0024 | CEBPA,PPARG | 197 (7) |
| 5,7-dimethoxyflavone | | chemical - endogenous mammalian | | | 0.049 | CYP1A1 | |
| DE 71 | | chemical reagent | | | 0.049 | CYP1A1 | |
| 3',5'-methoxyflavone | | chemical reagent | | | 0.049 | PTGS2 | |
| perfluorooctanoic acid | | chemical reagent | | | 0.0408 | CXCL8,DNMT3A,HMGCR | |
| stevioside | | chemical - endogenous non-mammalian | | | 0.0135 | PPARG,PTGS2 | |
| miocamycin | | chemical drug | | | 0.049 | MUC2 | |
| aurapten | | chemical - endogenous non-mammalian | | | 0.00109 | CPT1A,CXCL8,PTGS2 | 170 (7) |
| Hottip | | other | | | 0.0217 | SMAD3,TGFBR2 | |
| JUN/JUNB/JUND | | group | | | 0.041 | CCN1,FOS,FOSL1,JUN | |
| Rxr | | group | | | 0.000139 | ABCB10,ABCC2,CPT1A,CTSS,CYP24A1,CYP2B6,CYP3A5,GADD45A,HBEGF,INSIG1,KDM6B,NR0B2,NR5A2,PDK4,PPARG,PTGS2 | 456 (23) |
| RAR-RXR | | complex | | | 0.0217 | IRF1,TGM2 | |
| tylophorine | | chemical drug | | | 0.0135 | JUN,PTGS2 | |
| MLN8054 | | chemical drug | | | 0.00697 | BBC3,PMAIP1 | |
| perhexiline | | chemical drug | | | 0.0232 | DEPP1,LSS,SERPINA3 | |
| chlorcyclizine | | chemical drug | | | 0.0185 | DEPP1,LSS,SERPINA3 | |
| rh-endostatin | | chemical drug | | | 0.00354 | FAS,JUN,MAP1LC3B | |
| Cytochrome bc1 | | complex | | | 0.049 | MAOA | |
| CDKN2B-AS1 | | other | | | 0.0285 | CDKN2B,CXCL8,PTGS2 | |
| zinc oxide | | chemical drug | | | 0.0316 | MT1E,MT1X | |
| vanillyl-N-nonylamide | | chemical drug | | | 0.0021 | ATF3,CCNG2,GADD45A | |
| sucralose | | chemical reagent | | | 0.0135 | CEBPA,PPARG | |
| loxoprofen | | chemical drug | | | 0.049 | MUC2 | |
| tacedinaline | | chemical drug | | | 0.0021 | ELF3,FOS,IGF2 | |
| obatoclax | | chemical drug | | | 0.0021 | ATF3,MCL1,PMAIP1 | |
| PRC2 | | complex | | | 0.024 | CEACAM1,CEACAM6,CXCL1,CXCL8,GATA6,MMP7 | |
| G-Actin | | group | | | 0.049 | JUN | |
| thyroid hormone receptor | | group | | | 0.0363 | CPT1A,FOS,KLF9,NR5A2 | |
| 3-iodothyronamine | | chemical - endogenous mammalian | | | 0.049 | IRS2 | |
| 25-hydroxyvitamin D | | chemical drug | | | 0.0024 | CYP24A1,VDR | |
| bendamustine | | chemical drug | | | 0.00789 | AURKA,MCL1,PMAIP1 | 367 (7) |
| PKR dimer | | complex | | | 0.049 | ATF3 | |
| MAP2K4/7 | | group | | | 0.049 | PTGS2 | |
| Retinoic acid-RAR-RXR | | complex | | | 0.00546 | CRABP2,DUSP1,TGFB2 | |
| bovine testicular hyaluronidase | | biologic drug | | | 0.049 | CD44 | |
| dihydromethysticin | | chemical - endogenous non-mammalian | | | 0.049 | CYP3A5 | |
| NF279 | | chemical reagent | | | 0.049 | CXCL8 | |
| muraglitazar | | chemical drug | | | 0.00697 | EDN1,REN | 302 (7) |
| ZM 323881 | | chemical - kinase inhibitor | | | 0.049 | RCAN1 | |
| ALKBH5 | | enzyme | | | 0.0232 | CAV1,CLU,DNAJA1,FYN,IGF2,JAG2,PRDX4,RGS2,SERPINA5 | |
| LRRC26 | | ion channel | | | 0.0316 | CXCL1,CXCL8 | |
| GSK 189254 | | chemical drug | | | 0.049 | FOS | |
| MBNL3 | | other | | | 0.0024 | ABLIM1,CD44 | |
| INTS11 | | other | | | 0.00697 | EGR1,FOSB | |
| trypsin | | group | | | 0.0232 | CXCL8,FOS,PTGS2 | |
| (-)-OSU 6162 | | chemical drug | | | 0.049 | FOS | |
| ASB9 | | transcription regulator | | | 0.049 | CKB | |
| RNF187 | | enzyme | | | 0.0217 | HBEGF,JUN | |
| IGF2BP2 | | translation regulator | | | 0.0135 | ELOVL6,LIMS2 | |
| CYP2R1 | | enzyme | | | 0.049 | VDR | |
| FICD | | enzyme | | | 0.000452 | DNAJB1,HSPA1A/HSPA1B,HSPB1 | |
| Pad2 | | other | | | 0.049 | PTGS2 | |
| SLC22A2 | | transporter | | | 0.0217 | CPT1A,SLC22A5 | |
| LPIN1 | | phosphatase | | | 0.0144 | CEBPA,CPT1A,HMGCR,PPARG,PTGS2 | |
| patulin | | chemical toxicant | | | 0.0428 | ATF3,CYP1A1 | |
| LGALS12 | | other | | | 0.0217 | CEBPA,PPARG | |
| RNF217-AS1 | | other | | | 0.049 | INTU | |
| pridopidine | | chemical drug | | | 0.049 | FOS | |
| LINC01139 | | other | | | 0.0109 | EGR1,LDHA,SLC16A3 | |
| PDCD10 | | other | | | 0.0428 | KLF4,PROCR | |
| SSC5D | | transmembrane receptor | | | 0.049 | CXCL8 | |
| TMC8 | | other | | | 0.049 | CIB1 | |
| SLC27A4 | | transporter | | | 0.0135 | CEBPA,PPARG | |
| RHOJ | | enzyme | | | 0.00183 | AKAP12,ANXA1,CAV1,GSN,RND3 | |
| IL1RL2 | | transmembrane receptor | | | 0.0408 | CXCL1,CXCL2,CXCL8 | |
| TFAP2D | | transcription regulator | | | 0.049 | MT2A | |
| BSCL2 | | other | | | 0.0393 | ACSL1,AKR1C3,CEBPA,GPD1,PNPLA2,PPARG,PTGS2 | |
| NPC1 | 0.332 | transporter | | | 0.000345 | CAV1,CAV2,CD68,EGR1,HMGCR,IDI1,KLF4,LGALS3,LIPG,LSS,MMP15,NPC1,PTGS2 | 413 (17) |
| MARCHF3 | | other | | | 0.0217 | CXCL1,CXCL8 | |
| FBXO11 | | enzyme | | | 0.049 | BCL6 | |
| FBXO25 | | enzyme | | | 0.0024 | EGR1,FOS | 127 (5) |
| ZNF652 | | other | | | 0.0428 | TGFB2,TGFBR2 | |
| APOL1 | 0.986 | transporter | | | 0.0144 | CXCL2,IFNGR2,MUC13 | |
| TRIM29 | -0.497 | transcription regulator | | | 0.0428 | CD44,PMAIP1 | |
| CD3-TCR | | complex | | | 0.049 | TOB1 | |
| SRF-ELK1 | | complex | | | 0.049 | FOS | |
| Gpcr | | group | | | 0.0217 | FOS,JUN | |
| KREMEN1 | | other | | | 0.049 | AXIN2 | |
| BCAS2 | | other | | | 0.0217 | BBC3,PMAIP1 | |
| H2BW2 | | other | | | 0.049 | DNMT3A | |
| PCOLCE2 | | other | | | 0.049 | SCARB1 | |
| SLC15A4 | | transporter | | | 0.049 | IRF7 | |
| ATOH7 | | other | | | 0.0217 | FGF19,SOX4 | |
| ZC3H14 | | other | | | 0.0478 | IL15RA,NMI,TSC22D1 | |
| SLC51B | | transporter | | | 0.049 | NR0B2 | |
| ADTRP | | enzyme | | | 0.049 | TFPI | |
| RSPO3 | | kinase | | | 0.00789 | AXIN2,GLI1,PTGS2 | 236 (7) |
| STEAP1 | 0.932 | transporter | | | 0.0024 | CEBPA,PPARG | |
| PPIP5K1 | | phosphatase | | | 0.049 | RUNX2 | |
| RNASEH2A | | enzyme | | | 0.00000303 | IFI35,IFI44,IFIH1,IFIT3,IFITM3,IFNGR2,IRF1,IRF7,NMI,PTGS2,TRIM21 | |
| PRDM5 | | transcription regulator | | | 0.00108 | APOBEC3B,CEMIP,GADD45B,ID2,MYB,NCOA7,OVOL1,PML,RUNX2,S100A2,STK32C,TINAGL1 | |
| PRG3 | | other | | | 0.049 | CXCL8 | |
| PRKAG3 | | other | | | 0.000519 | ACVR1,AMD1,ATP1B1,CCNDBP1,CD55,EGR1,HIVEP2,KDM2B,KLF4,MAT2A,NEO1,NRIP1,RHOU,RNF128,SNCG,SORD,TSPAN8,UGT1A10 (includes others),ZFP36 | |
| HSPBP1 | | other | | | 0.0135 | HSPA1A/HSPA1B,HSPA1L | |
| FNDC3B | | other | | | 0.049 | RUNX2 | |

Supplementary Table 4. Upstream regulator analysis based gene set enrichment analysis (GSEA) of differentially expressed genes (DEGs) after ATRA+PDT compared to PDT

| Upstream Regulator | Expr Log Ratio | Molecule Type | Predicted Activation State | Activation z-score | p-value of overlap | Target Molecules in Dataset | Mechanistic Network |
|---|---|---|---|---|---|---|---|
| RND2 | | enzyme | | | 0.049 | RHOB | |
| FCN1 | | other | | | 0.049 | CXCL8 | |
| PPP2R2D | | other | | | 0.049 | PPARG | |
| HCAR1 | 0.404 | G-protein coupled receptor | | | 0.0185 | EDN1,SLC16A1,SLC16A3 | |
| POLR3G | | enzyme | | | 0.0232 | AURKA,KLF6,S100A4 | |
| procyanidin B2 | | chemical - endogenous non-mammalian | | | 0.049 | PTGS2 | |
| BUD23 | | enzyme | | | 0.0217 | CXCL8,TSC22D3 | |
| TMEM184A | | other | | | 0.049 | DUSP1 | |
| B3GNT6 | | enzyme | | | 0.049 | MUC2 | |
| UBN1 | | transcription regulator | | | 0.049 | CCNA2 | |
| LMAN1 | | other | | | 0.049 | F5 | |
| IKBIP | | other | | | 0.0135 | CXCL2,NFKBIZ | |
| USP9Y | | peptidase | | | 0.049 | MUC2 | |
| C1QL4 | | other | | | 0.0024 | CEBPA,PPARG | |
| IQCB1 | | other | | | 0.049 | BBC3 | |
| CLDN9 | | other | | | 0.049 | TJP1 | |
| ZMIZ2 | | transcription regulator | | | 0.00109 | AXIN2,CD44,JUN | |
| Histone h4 | | group | | | 0.00103 | AKAP12,CDC25A,CXCL3,CXCL8,CYP24A1,DKK1,DUSP16,FOS,IRF1,KIF13B,NEDD4L,OAS3,PMAIP1,PPARG,PTGS2,RUNX2,TGFBR2 | 478 (20) |
| CUZD1 | | other | | | 0.0217 | EREG,ID2 | |
| TRIM17 | | enzyme | | | 0.049 | MCL1 | |
| ZNF580 | | transcription regulator | | | 0.049 | CXCL8 | |
| CCDC22 | | other | | | 0.049 | BIRC3 | |
| MBOAT7 | 0.307 | enzyme | | | 0.0185 | ACSS2,ELOVL6,HMGCR | |
| SELENOS | | other | | | 0.0343 | CAV1,CEBPA,PPARG | |
| CIRBP | 0.347 | translation regulator | | | 0.0135 | CXCL2,CXCL3 | |
| IFNA17 | | cytokine | | | 0.0285 | EIF2AK2,IFIH1,PLAAT4 | |
| potassium channel | | group | | | 0.00697 | FOS,JUN | |
| sGC | | complex | | | 0.0217 | EGR1,FOS | |
| TCF/LEF | | group | | | 0.0343 | JUN,PTGS2,RUNX2 | |
| GPR39 | | G-protein coupled receptor | | | 0.0316 | CLU,PNPLA2 | |
| NAA30 | | enzyme | | | 0.00697 | FAS,PMAIP1 | |
| GALNT4 | | enzyme | | | 0.0428 | SMAD3,TJP1 | |
| EIF3M | | other | | | 0.0316 | CDC25A,MT2A | |
| ARID2 | | transcription regulator | | | 0.0217 | BMP4,FGFR2 | |
| METTL8 | | enzyme | | | 0.0024 | CEBPA,PPARG | |
| MDGA2 | | other | | | 0.0232 | CASP9,FAS,SESN2 | |
| CACNA1C | | ion channel | | | 0.00546 | CAV1,PTGS2,S100A10 | 163 (7) |
| PIWIL4 | | other | | | 0.0316 | FGFR2,TGFBR2 | |
| MLLT11 | | other | | | 0.049 | CD44 | |
| RGMB | | other | | | 0.0021 | CXCL2,CXCL3,LCN2 | |
| BATF2 | | other | | | 0.0024 | CCN1,JUN | 296 (7) |
| BCO1 | | enzyme | | | 0.00287 | ARRDC3,NR1D1,PPARG,SCARB1 | |
| MAPKBP1 | | other | | | 0.049 | CXCL8 | |
| ZG16B | | other | | | 0.049 | CXCR4 | |
| KCTD10 | | ion channel | | | 0.049 | RHOB | |
| Dcpp1 (includes others) | | other | | | 0.049 | OTUD7B | |
| NSD1 | | transcription regulator | | | 0.0343 | BMP4,KLF6,ZFP36L1 | |
| harman | | chemical - endogenous non-mammalian | | | 0.049 | CYP1A1 | |
| USP47 | | peptidase | | | 0.049 | CDC25A | |
| CPB1 | | peptidase | | | 0.049 | CASP4 | |
| FGF14 | | growth factor | | | 0.049 | FOS | |
| CERK | | kinase | | | 0.0368 | CLCF1,CXCL1,CXCL2,CXCL8,CXCR4 | |
| QRFP | | other | | | 0.00354 | ACSL1,CEBPA,PPARG | |
| CDC73 | | other | | | 0.0078 | GATA2,HMGCS2,IGF2,IRF1,LGALS3 | 290 (7) |
| CAMSAP2 | | other | | | 0.049 | TJP1 | |
| DSC2 | | other | | | 0.049 | CD44 | |
| TRPM3 | | ion channel | | | 0.049 | EGR1 | |
| FAM3B | | cytokine | | | 0.0241 | CASP4,CAV1,LCN2,LDHA | |
| KLHL21 | | other | | | 0.00697 | DUSP1,NFKBIZ | |
| MLXIP | | transcription regulator | | | 0.0024 | ARRDC4,TXNIP | |
| STEAP2 | 0.777 | enzyme | | | 0.0024 | CEBPA,PPARG | |
| LRRC4 | | other | | | 0.049 | STMN1 | |
| CCDC8 | | other | | | 0.049 | BBC3 | |
| NSD2 | | enzyme | | | 0.000426 | BACE2,CDC25A,CXCR4,FOS,GADD45A,IRF7,JUN,PRKCA | 249 (7) |
| FANK1 | | transcription regulator | | | 0.049 | JUN | |
| SLC4A7 | | transporter | | | 0.049 | SLC16A1 | |
| CLEC12A | | other | | | 0.0316 | IFIT3,IRF7 | |
| SBDS | | other | | | 0.000345 | AKR1C1/AKR1C2,AKR1C3,CHAC1,DNAJB1,EGR1,FBXO32,FOS,GADD45B,HSPA6,MAFF,MAOA,MT2A,PPP1R15A | |
| TCIM | | other | | | 0.00354 | CXCL1,CXCL8,PTGS2 | 398 (14) |
| LTBP4 | -0.661 | growth factor | | | 0.00546 | BMP4,ID2,TGFB2 | 441 (11) |
| Gata6os | | other | | | 0.049 | GATA6 | |
| Hedgehog | | group | | | 0.0106 | CCNA2,FOXL1,GATA6,HOXA5,IGF2,KLF4,VIL1 | |
| Sapk | | group | | | 0.000452 | FBXO32,FOS,GADD45A | 254 (10) |
| GPX6 | | enzyme | | | 0.049 | PPP1R1B | |
| MBNL2 | | other | | | 0.049 | ABLIM1 | |
| GPR68 | | G-protein coupled receptor | | | 0.0217 | DUSP1,PTGS2 | |
| nickel sulfide | | chemical reagent | | | 0.049 | PTGS2 | |
| rimantadine | | chemical drug | | | 0.049 | CXCL8 | |
| Eif2 | | complex | | | 0.0135 | EIF2AK2,PTGS2 | |
| sphingomyelinase | | group | | | 0.0428 | JUN,PTGS2 | |
| P-TEFb | | complex | | | 0.0135 | BBC3,CXCL8 | |
| black raspberry extract | | chemical drug | | | 0.00943 | ADAMTS6,AKR1B10,AMBP,CD44,CLDN1,DUSP5,KRT20,MCL1,PIP5K1A,PLK3,SCEL,SLC30A1 | |
| FAS-AS1 | | other | | | 0.049 | FAS | |
| Myosin2 | | complex | | | 0.049 | PPARG | |
| Pro-inflammatory Cytokine | | group | | | 0.00699 | ABCC2,ASS1,CD55,CLU,LCN2,LGALS9,LIPG,PI3,SDC4 | |
| proanthocyanidin derivative | | chemical - other | | | 0.0135 | CYP1A1,JUN | |
| Tpl2 kinase inhibitor | | chemical - kinase inhibitor | | | 0.049 | PTGS2 | |
| IL7R | | transmembrane receptor | | | 0.00344 | C1orf116,CXCL8,DAPK1,DPP4,FHL2,MCL1,MMRN2,PCSK6,PIM1,SKAP1,TJP3 | |
| TAF9 | | transcription regulator | | | 0.0217 | CXCL8,IRF1 | |
| cis-vaccenic acid | | chemical - endogenous mammalian | | | 0.049 | PNPLA2 | |
| iguratimod | | chemical drug | | | 0.0428 | CXCL2,CXCL3 | |
| IDH2 | | enzyme | | | 0.00387 | CEBPA,EGR1,PPARG,TGFB2 | 248 (7) |
| hispidulin | | chemical drug | | | 0.049 | EGR1 | |
| urotensin II | | biologic drug | | | 0.0428 | CXCL8,SMAD3 | |
| NNZ 2566 | | biologic drug | | | 0.049 | ATF3 | |
| neurotropin | | chemical drug | | | 0.049 | SEMA3A | |
| vasopressins | | biologic drug | | | 0.049 | FOS | |
| firtecan pegol | | chemical drug | | | 0.0135 | CA9,CXCR4 | |
| pycnogenols | | chemical drug | | | 0.00697 | CXCL8,PTGS2 | |
| HCV 796 | | chemical drug | | | 0.049 | MAP1LC3B | |
| mediator | | complex | | | 0.049 | FOS | |
| 7S NGF | | complex | | | 0.0217 | CLU,FOS | |

Supplementary Table 4. Upstream regulator analysis based gene set enrichment analysis (GSEA) of differentially expressed genes (DEGs) after ATRA+PDT compared to PDT

| Upstream Regulator | Expr Log Ratio | Molecule Type | Predicted Activation State | Activation z-score | p-value of overlap | Target Molecules in Dataset | Mechanistic Network |
|---|---|---|---|---|---|---|---|
| Ppp2c | | group | | | 0.0232 | CDKN2B,DUSP1,PIM1 | |
| p85 (pik3r) | | group | | | 0.0478 | MCL1,PPARG,PTGS2 | |
| Mucin | | group | | | 0.0217 | CXCL8,PTGS2 | |
| Pik3r | | group | | | 0.0316 | BBC3,CXCL8 | |
| Rho gdi | | group | | | 0.049 | RHOB | |
| LAMTOR5 | | other | | | 0.0232 | CXCL8,LDHA,S100A4 | |
| BCOR | | transcription regulator | | | 0.0207 | ARID5B,ETV6,GLI1,GLI2 | |
| PMF1/PMF1-BGLAP | | transcription regulator | | | 0.049 | SAT1 | |
| ASCL2 | | transcription regulator | | | 0.0428 | CXCR4,EGR1 | |
| FLII | | other | | | 0.0135 | CLDN1,CXCL8 | |
| MEIS2 | 0.552 | transcription regulator | | | 0.0316 | CDKN2B,ETV1 | |
| PF-4523655 | | biologic drug | | | 0.049 | DDIT4 | |
| PDGFD | | growth factor | | | 0.0428 | CXCR4,PTGS2 | |
| cucurbitacin E | | chemical - endogenous non-mammalian | | | 0.049 | PTGS2 | |
| bitertanol | | chemical drug | | | 0.0135 | CYP1A1,CYP3A5 | |
| ARHGEF17 | | other | | | 0.049 | FOS | |
| interferon beta-1a | | biologic drug | | | 0.0011 | ATP5MC3,CXCR4,DNAJA1,DNAJB1,GADD45B,IFI35,IL1RN,IRF1,IRF7,NMI,PSMB8,PSME1,SERPINA1,TBXAS1,UQCRFS1 | 219 (8) |
| idelalisib | | chemical drug | | | 0.0428 | MCL1,UGCG | |
| IER3 | | other | | | 0.0428 | BIRC3,MCL1 | |
| RAB11FIP3 | | other | | | 0.0135 | FOS,JUN | |
| ANTXR1 | | transmembrane receptor | | | 0.049 | HSPA1A/HSPA1B | |
| pidilizumab | | biologic drug | | | 0.049 | CXCR4 | |
| Atf | | group | | | 0.0217 | JUN,PTGS2 | |
| DIO1 | | enzyme | | | 0.0135 | SLC16A1,SLC16A3 | |
| NCOR2 | | transcription regulator | | | 0.0017 | AXIN2,BIRC3,CEBPA,CTSS,CXCL8,FOS,HBEGF,JUN,PTGS2 | 506 (25) |
| MEDI-547 | | biologic drug | | | 0.049 | EPHA2 | |
| PF-4691502 | | chemical drug | | | 0.0135 | BBC3,PMAIP1 | |
| GSK0660 | | chemical reagent | | | 0.0428 | FBXO32,PDK4 | |
| NELFB | | other | | | 0.0185 | ATF3,GADD45A,GADD45B | |
| cercosporin | | chemical - endogenous non-mammalian | | | 0.0109 | AXIN2,CDC25A,DKK1 | |
| IGF2R | | transmembrane receptor | | | 0.0109 | IGF2,NEDD9,NPC1 | |
| APOA4 | | transporter | | | 0.0285 | DGAT2,FOS,SCARB1 | |
| TMC6 | | transporter | | | 0.049 | CIB1 | |
| GTF2H2 | | transcription regulator | | | 0.049 | KLF9 | |
| SYTL4 | | transporter | | | 0.049 | STX1A | |
| PAK4 | | kinase | | | 0.0185 | NOX1,PPARG,TJP1 | |
| NOD1 | | other | | | 0.0123 | CARD6,CXCL2,CXCL8,DUSP1 | |
| RAP1GAP | -0.574 | other | | | 0.0428 | CDK6,FOS | |
| GPLD1 | | enzyme | | | 0.049 | LGALS3BP | |
| MBNL1 | | other | | | 0.0185 | ABLIM1,SOX4,TGFB2 | |
| GNA14 | -0.765 | enzyme | | | 0.000000306 | AKAP12,ATF3,BMP4,CSPG4,CXCL3,CXCL8,DUSP5,GADD45A,HBEGF,IL1RN,KLF6,LIF,PHLDA1,PTGS2,SGK1,ZFP36L1 | 309 (14) |
| REV1 | | enzyme | | | 0.00354 | BBC3,PMAIP1,SESN2 | 367 (7) |
| ITGA5 | | transmembrane receptor | | | 0.0317 | CD44,FOS,IGF2,IGFBP2,JUN,TGFBR2 | |
| CD244 | | transmembrane receptor | | | 0.0109 | CXCL8,EGR1,FOS | |
| EDN2 | | growth factor | | | 0.049 | PTGS2 | |
| TLX1 | | transcription regulator | | | 0.0461 | GATA2,MCL1,MYB,NMI | |
| YWHAQ | | other | | | 0.00507 | CASP7,CASP9,ETV1,MMP7 | |
| CTF1 | | cytokine | | | 0.00863 | CEBPD,FOS,IRF1,PNPLA2,PPARG,SERPINA3 | 506 (19) |
| OPRK1 | | G-protein coupled receptor | | | 0.0021 | FOS,NRP1,PLXND1 | 297 (7) |
| AHRR | | transcription regulator | | | 0.049 | CYP1A1 | |
| FAF1 | | other | | | 0.00109 | AXIN2,DKK1,RUNX2 | 289 (7) |
| KIFAP3 | | other | | | 0.049 | FOS | |
| TFDP1 | | transcription regulator | | | 0.0299 | CASP7,CASP9,FGFR2,MYB,STMN1 | |
| RAB27A | | enzyme | | | 0.049 | CD44 | |
| AGR2 | -0.77 | other | | | 0.0428 | EGR1,FOS | |
| LGALS8 | | other | | | 0.0109 | CXCL1,CXCL3,CXCL8 | |
| ZBTB33 | | transcription regulator | | | 0.000557 | BBC3,BCL6,FAS,MMP7,S100A4 | |
| PPY | | other | | | 0.049 | FOS | |
| PROK2 | | other | | | 0.0316 | CDC25A,FOS | |
| IDH1 | | enzyme | | | 0.013 | EGR1,GLI1,PPARG,PPP1R15A,RUNX2,SERPINH1,TGFB2 | |
| POU2F1 | | transcription regulator | | | 0.00262 | ACSL1,BBC3,BCL6,CD55,CXCL8,CYP1A1,CYP2B6,DDIT4,FRRS1,GADD45A,JUND,PDE9A,REL,SESN2 | 412 (12) |
| RXRB | | ligand-dependent nuclear receptor | | | 0.00288 | ACSL1,ALDH1A1,CTSS,CYP24A1,ECM1,FLRT3,HBEGF,IGFBP6,LCN2,S100A4 | 353 (13) |
| LINC01370 | | other | | | 0.049 | GLIS3 | |
| LINC01503 | | other | | | 0.049 | FOSL1 | |
| entolimod | | biologic drug | | | 0.0316 | CXCL2,CXCL3 | |
| WBP11 | | phosphatase | | | 0.049 | TUBGCP6 | |
| GPHA2 | | other | | | 0.049 | FOS | |
| SYMPK | | other | | | 0.0024 | CLDN2,TJP1 | |
| dodecylbenzenesulfonic acid | | chemical reagent | | | 0.049 | PI3 | |
| cyproconazole | | chemical drug | | | 0.0024 | CYP2B6,GADD45B | |
| recombinant interferon alpha | | biologic drug | | | 0.049 | RNF213 | |
| SNX9 | | transporter | | | 0.049 | ADAM9 | |
| OSMR | | transmembrane receptor | | | 0.00345 | BCL3,BIRC3,CASP4,CXCL2,FOS,PIM1,SBNO2,SERPINA1,ZFP36 | 394 (18) |
| KSR2 | | kinase | | | 0.049 | CXCL8 | |
| CIC | | transcription regulator | | | 0.0185 | ETV1,FOSL1,MAFF | |
| NMB | | other | | | 0.0024 | FOS,JUN | 444 (22) |
| MPZ | | other | | | 0.00000241 | DNAJC3,HMGCR,ID2,JUN,PPP1R15A,SDF2L1,SOX4,SQLE | |
| NFYB | | transcription regulator | | | 0.0139 | AMD1,APOBEC3B,CAV1,CDC25A,CHPT1,DPP4,FAS,FGFR2,FRYL,GADD45A,GPR160,KDM8,PAQR5,PITPNM3,RHOB,RUNX2,S100A4,SCARB1,SLC25A28,SLC39A8,TGFBR2,TJP1 | |
| CDK5R1 | | kinase | | | 0.0123 | CXCL8,FOS,PMAIP1,PPP1R1B | |
| LY96 | | transmembrane receptor | | | 0.0408 | CXCL2,CXCL3,CXCL8 | |
| LBP | | transporter | | | 0.0478 | CXCL3,CXCL8,IL1RN | |
| DAB2IP | | other | | | 0.0109 | CD44,CLU,EGR1 | |
| GNE | -0.386 | kinase | | | 0.0428 | CHAC1,JDP2 | |
| PIAS2 | | transcription regulator | | | 0.0185 | CDKN2B,FOS,IRF1 | |
| DAB2 | | other | | | 0.0428 | FOS,HMGCR | |
| PHF19 | | other | | | 0.0135 | FGF19,HOXA5 | |
| LOC102724788/PRODH | -1.858 | enzyme | | | 0.0217 | LOC102724788/PRODH,PTGS2 | |
| CHRM3 | | G-protein coupled receptor | | | 0.0144 | CCN1,EGR1,FOS | |
| SMC3 | | other | | | 0.0343 | CCNG2,IRS2,SOX4 | |
| PNN | | other | | | 0.0343 | GDF15,JUN,MMP7 | |
| MDM4 | | transcription regulator | | | 0.0478 | BBC3,HIPK2,LGALS3 | |
| Lysosomal Protease | | group | | | 0.049 | MAP1LC3B | |
| miR-338-5p (miRNAs w/seed ACAAUAU) | | mature microRNA | | | 0.049 | SPINK1 | |
| mir-338 | | microRNA | | | 0.0217 | RUNX2,SPINK1 | |
| miR-516a-5p (miRNAs w/seed UCUCGAG) | | mature microRNA | | | 0.049 | KLK10 | |
| mir-506 | | microRNA | | | 0.0428 | CXCL8,SPDEF | |
| mir-488 | | microRNA | | | 0.0217 | ADAM9,CXCL8 | |
| miR-27b-5p (miRNAs w/seed GAGCUUA) | | mature microRNA | | | 0.049 | ATP2B1 | |
| mir-196 | | microRNA | | | 0.0343 | ANXA1,FOS,HMGA2 | |
| miR-293-3p (miRNAs w/seed GUGCCGC) | | mature microRNA | | | 0.049 | DKK1 | |
| miR-467a-5p (and other miRNAs w/seed AAGUGCC) | | mature microRNA | | | 0.049 | DKK1 | |
| miR-129-5p (and other miRNAs w/seed UUUUUGC) | | mature microRNA | | | 0.0343 | DNMT3A,ETV6,SOX4 | |

Supplementary Table 4. Upstream regulator analysis based gene set enrichment analysis (GSEA) of differentially expressed genes (DEGs) after ATRA+PDT compared to PDT

| Upstream Regulator | Expr Log Ratio | Molecule Type | Predicted Activation State | Activation z-score | p-value of overlap | Target Molecules in Dataset | Mechanistic Network |
|---|---|---|---|---|---|---|---|
| mir-129 | | microRNA | | | 0.049 | CXCL8 | |
| mir-345 | | microRNA | | | 0.049 | ABCC1 | |
| miR-193a-3p (and other miRNAs w/seed ACUGGCC) | | mature microRNA | | | 0.0144 | ETS1,MCL1,RPS6KB2 | |
| mir-489 | | microRNA | | | 0.0217 | ADAM9,MMP7 | |
| miR-135a-5p (and other miRNAs w/seed AUGGCUU) | | mature microRNA | | | 0.0285 | PPARG,RUNX2,TRPS1 | |
| miR-125a-3p (miRNAs w/seed CAGGUGA) | | mature microRNA | | | 0.049 | FYN | |
| miR-605-5p (miRNAs w/seed AAAUCCC) | | mature microRNA | | | 0.049 | SEC24D | |
| miR-142-5p (and other miRNAs w/seed AUAAAGU) | | mature microRNA | | | 0.049 | TGFBR2 | |
| miR-23a-5p (and other miRNAs w/seed GGGUUCC) | | mature microRNA | | | 0.049 | LOC102724788/PRODH | |
| miR-186-5p (miRNAs w/seed AAAGAAU) | | mature microRNA | | | 0.0144 | AURKA,HMGA2,TGFBR2 | |
| miR-328-3p (and other miRNAs w/seed UGGCCCU) | | mature microRNA | | | 0.0135 | CD44,CEBPA | |
| mir-199 | | microRNA | | | 0.0408 | DNMT3A,EDN1,EGLN1 | |
| miR-690 (miRNAs w/seed AAGGCUA) | | mature microRNA | | | 0.049 | CEBPA | |
| Mir690 | | microRNA | | | 0.049 | CEBPA | |
| miR-130a-3p (and other miRNAs w/seed AGUGCAA) | | mature microRNA | | | 0.0343 | CXCL2,CXCL3,HOXA5 | |
| mir-205 | | microRNA | | | 0.0478 | AXIN2,INPPL1,RUNX2 | |
| miR-197-3p (and other miRNAs w/seed UCACCAC) | | mature microRNA | | | 0.0428 | ACVR1,TGFBR2 | |
| mir-663 | | microRNA | | | 0.0024 | CXCR4,HSPG2 | |
| mir-1301 | | microRNA | | | 0.049 | KLF6 | |
| mir-887 | | microRNA | | | 0.049 | PLD2 | |
| MED15 | 0.292 | transcription regulator | | | 0.00789 | BBC3,LIMA1,S100A2 | |
| MIR3619 | | microRNA | | | 0.049 | PLD2 | |
| KREMEN2 | | other | | | 0.049 | AXIN2 | |
| PPP1R13B | | phosphatase | | | 0.0217 | BBC3,LATS2 | |
| TRIM28 | | transcription regulator | | | 0.0452 | CASP7,CXCR4,ETS1,MYEOV,PMAIP1,RCAN1,S100A4 | |
| RGS19 | | enzyme | | | 0.00546 | AXIN2,BMP4,DKK1 | |
| MLK | | group | | | 0.049 | MMP7 | |
| NUCB2 | -0.489 | other | | | 0.0316 | FOS,PTGS2 | |
| RIPK4 | | kinase | | | 0.0217 | GRHL3,OVOL1 | |
| HOXB9 | | transcription regulator | | | 0.0101 | CXCL8,EREG,REN,TGFB2 | |
| WSB1 | | other | | | 0.0217 | CA9,HIPK2 | |
| ABHD5 | | enzyme | | | 0.0316 | PDK4,PNPLA2 | |
| ANXA4 | | other | | | 0.049 | CXCL8 | |
| Irs3 | | other | | | 0.0217 | EGR1,IRS2 | |
| HBP1 | | transcription regulator | | | 0.0232 | H1-0,PIM1,PTGS2 | |
| TIAL1 | | transcription regulator | | | 0.0217 | MAP1LC3B,TIA1 | |
| NPVF | | other | | | 0.049 | FOS | |
| KRT14 | | other | | | 0.00984 | CXCL1,CXCL8,GJB2,KLK10,KLK6,MMP7 | 299 (7) |
| RCE1 | | peptidase | | | 0.0488 | B2M,GPC4,HBEGF,RCN1,TGFB2 | |
| HDAC3 | | transcription regulator | | | 0.0192 | BIRC3,CD55,CXCL2,CXCL8,ELOVL6,FOS,LGALS9,ME1,PHLPP1,PTGS2 | |
| miR-211-3p (miRNAs w/seed CAGGGAC) | | mature microRNA | | | 0.049 | CDK6 | |
| MTA2 | | transcription regulator | | | 0.0363 | GATA2,ID2,JUN,SQLE | |
| IL1R2 | -0.516 | transmembrane receptor | | | 0.0144 | CXCL2,CXCL3,CXCL8 | |
| F10 | | peptidase | | | 0.0285 | CCN1,CXCL8,EGR1 | |
| ATN1 | | transcription regulator | | | 0.0145 | ATP1B1,ATP1B3,BCL6,CCNDBP1,ETV1,GRK5,GSN,HBEGF,MCAM,PSME1,SOX4,STMN1,TUBA4A | |
| DRD1 | | G-protein coupled receptor | | | 0.00207 | EGR1,FOS,FOSB,S100A10 | 441 (20) |
| TLE1 | | transcription regulator | | | 0.0000174 | ATF3,DUSP1,EGR1,FOS,MCL1,MEIS2 | 313 (7) |
| Scd2 | | enzyme | | | 0.049 | PPARG | |
| BORCS8-MEF2B | | transcription regulator | | | 0.049 | JUN | |
| HCRT | | other | | | 0.0109 | EGR1,FOS,KITLG | |
| GJA5 | | transporter | | | 0.049 | REN | |
| RBX1 | | enzyme | | | 0.0428 | MCL1,RHOB | |
| UTS2 | | other | | | 0.0217 | FOS,HPGD | |
| eprenetapopt | | chemical drug | | | 0.0368 | BBC3,CARD10,GADD45B,MCL1,PMAIP1 | |
| pituitary adenylate cyclase-activating polypeptide | | biologic drug | | | 0.00102 | ACER2,AKR1B10,AXIN2,EGR1,ERRFI1,FOSL1,FRRS1,MACROD1 | |
| SLC30A9 | | transporter | | | 0.049 | CYP1A1 | |
| CAMK2 | | group | | | 0.049 | BIRC3 | |
| IL4I1 | | enzyme | | | 0.00789 | CXCL1,CXCL8,TOB1 | 211 (7) |
| DLC1 | | other | | | 0.00354 | CDK6,CDKN2B,S100A10 | |
| UGT2B4 | | enzyme | | | 0.049 | NR0B2 | |
| Nisch | | other | | | 0.0316 | FAS,FOS | |
| SIGLEC7 | | transmembrane receptor | | | 0.049 | PTGS2 | |
| SIGLEC9 | | other | | | 0.049 | PTGS2 | |
| ZNF613 | | transcription regulator | | | 0.049 | ID2 | |
| COL4A2 | | other | | | 0.049 | FAS | |
| LY6D | | other | | | 0.049 | TSTA3 | |
| COX4I1 | | enzyme | | | 0.0285 | PDK4,TGFB2,TSPAN8 | |
| FOXC1 | | transcription regulator | | | 0.0279 | CXCR4,MMP7,RUNX2,TSC22D1 | |
| GNB2 | | enzyme | | | 0.00863 | CXCL3,DUSP1,FOSB,IL1R2,JAG2,TSC22D3 | |
| ELK4 | | transcription regulator | | | 0.00287 | EGR1,FOS,FOSL1,NCOA7 | 250 (12) |
| PPM1B | | phosphatase | | | 0.0428 | CEBPD,HBEGF | |
| BCAR1 | | enzyme | | | 0.0232 | CDKN2B,EGR1,MMP7 | |
| XIAP | | enzyme | | | 0.0363 | CDK6,CXCL1,CXCL8,PTGS2 | |
| ITGA1 | | other | | | 0.0279 | CAV1,MMP7,PPARG,PTGS2 | |
| SUPT16H | | transcription regulator | | | 0.0363 | DUSP5,EGR1,HSPA1A/HSPA1B,ID2 | |
| DDB2 | | other | | | 0.0185 | BBC3,EIF2AK2,SEMA3A | |
| GCLC | | enzyme | | | 0.000452 | FOS,JUN,SLC39A8 | 231 (7) |
| SSPN | | other | | | 0.0316 | CEACAM5,PTGS2 | |
| RNGTT | | phosphatase | | | 0.049 | SCARB1 | |
| MAPKAP1 | | other | | | 0.0408 | CXCR4,DAPK1,SLFN5 | |
| RAD51AP1 | | other | | | 0.0316 | CD44,KLF4 | |
| FABP2 | | transporter | | | 0.0232 | CPT1A,HMGCR,PPARG | |
| INSIG2 | | other | | | 0.0343 | ELOVL6,HMGCR,INSIG1 | |
| GPI | | enzyme | | | 0.0478 | CAV1,CXCL8,STEAP4 | |
| LIMA1 | 0.456 | other | | | 0.0144 | ID2,MMP7,VDR | |
| PIAS4 | | transcription regulator | | | 0.0343 | CXCL2,IRF1,PDK4 | |
| PIKFYVE | | kinase | | | 0.0144 | ALDH1A1,ATF3,MAP1LC3B | |
| SLC22A1 | | transporter | | | 0.0428 | CPT1A,SLC22A5 | |
| RPS6KA4 | | kinase | | | 0.0185 | DUSP1,FOS,JUN | |
| WNK4 | -0.875 | kinase | | | 0.0217 | CFTR,WNK4 | |
| ADRA1B | | G-protein coupled receptor | | | 0.029 | CEBPD,DUSP1,EGR1,FOS,JUN,LIF | |
| ADRA1A | | G-protein coupled receptor | | | 0.00316 | CEBPD,CYP1A1,DUSP1,EGR1,FOS,JUN,LIF,MMP7 | 461 (23) |
| KCNE3 | 1.117 | ion channel | | | 0.000104 | AKR1C3,CDHR5,CSRNP1,EPN3,FOSB,HMGCS2,HSPA1A/HSPA1B,KCNK5,KLF4,PHLDA1,PLK3,PTGS2 | |
| ADRA1D | | G-protein coupled receptor | | | 0.029 | CEBPD,DUSP1,EGR1,FOS,JUN,LIF | |
| ONECUT1 | | transcription regulator | | | 0.00156 | AMBP,CDC25A,CEBPA,CEBPD,CLDN2,CXCL1,CXCL3,EHF,FAS,GSTA4,HMGCR,HNF4A,HSPA1A/HSPA1B,HSPH1,KDM8,LAMC2,MYOF,NR0B2,PMAIP1,RARG,RBM48,SERPINA1,S | 141 (5) |
| TNPO2 | | transporter | | | 0.049 | CXCL8 | |
| PFKFB2 | | kinase | | | 0.049 | PDK4 | |
| Zfp54 | | other | | | 0.049 | PPARG | |
| GATAD2B | | transcription regulator | | | 0.0024 | CXCL8,PTGS2 | |
| MEOX1 | | transcription regulator | | | 0.0109 | BMP4,GLI1,GLI2 | |
| TIA1 | 0.395 | other | | | 0.0185 | CCNA2,MAP1LC3B,PTGS2 | |
| TYMP | | growth factor | | | 0.049 | CXCL8 | |

Supplementary Table 4. Upstream regulator analysis based gene set enrichment analysis (GSEA) of differentially expressed genes (DEGs) after ATRA+PDT compared to PDT

| Upstream Regulator | Expr Log Ratio | Molecule Type | Predicted Activation State | Activation z-score | p-value of overlap | Target Molecules in Dataset | Mechanistic Network |
|---|---|---|---|---|---|---|---|
| PTPRE | | phosphatase | | | 0.0285 | CXCL2,CXCL3,TSC22D1 | |
| TPSD1 | | peptidase | | | 0.0428 | CXCL8,PTGS2 | |
| SLC1A2 | | transporter | | | 0.049 | FOS | |
| RAC2 | | enzyme | | | 0.00358 | ANXA1,FHL2,FOS,LDHA,MACF1,RUNX2,TSC22D1,ZYX | |
| IAPP | | other | | | 0.00234 | BACE2,FOS,HMGCR,IDI1,MSMO1 | 303 (7) |
| KIDINS220 | | transcription regulator | | | 0.0217 | EGR1,FOS | |
| GPHB5 | | other | | | 0.049 | FOS | |
| anatabine | | chemical - endogenous non-mammalian | | | 0.0135 | IL1R2,PTGS2 | |
| SCN7A | | ion channel | | | 0.049 | FOS | |
| RNF7 | | enzyme | | | 0.049 | JUN | |
| AURKAIP1 | | enzyme | | | 0.049 | AURKA | |
| RDX | | other | | | 0.049 | ABCC2 | |
| IGFBP4 | | other | | | 0.0316 | IGFBP6,MCAM | |
| CKS1B | | kinase | | | 0.0428 | CCNA2,CXCL8 | |
| TERC | | other | | | 0.00134 | AURKA,CLU,EGR1,H1-0,HMGCR,NR1D1,TUBB2A | |
| CD3E | | transmembrane receptor | | | 0.015 | BCL10,CBLB,FAS,FOS,JUN,MYB,PIK3R1,PTGS2 | |
| PROK1 | | growth factor | | | 0.000178 | CXCL8,MCL1,PTGS2,RCAN1 | 447 (24) |
| FOLR1 | | transporter | | | 0.00729 | ACVR1,ARHGDIB,BIRC3,CAV1,MDK,MPP6,PPARG,S100A4,SHCBP1,SORD,ST3GAL4 | |
| PPP2R2A | | phosphatase | | | 0.0217 | SAT1,SERPINE2 | |
| NAPB | | transporter | | | 0.049 | CYP1A1 | |
| FANCC | | other | | | 0.000438 | ATP1B1,AXIN2,DKK1,FHL2,GDF15,MMP7,MYOF,PTGS2,RND3,SKAP2,TBC1D9 | 400 (12) |
| PIP5K1C | | kinase | | | 0.0428 | ERRFI1,PIP5K1A | |
| N4BP1 | | other | | | 0.0109 | HLA-B,JUN,MAP3K14 | |
| ANKS1A | | other | | | 0.049 | FOS | |
| bis(4-hydroxyphenyl)sulfone | | chemical reagent | | | 0.0316 | CEBPA,PPARG | |
| 5-methoxytryptophan | | chemical - endogenous mammalian | | | 0.049 | PTGS2 | |
| venetoclax | | chemical drug | | | 0.049 | MCL1 | |
| 20-hydroxyprostaglandin E2 | | chemical - endogenous mammalian | | | 0.049 | PPARG | |
| MYCT1 | | other | | | 0.0217 | EGLN1,MXD4 | |
| BCL6B | | transcription regulator | | | 0.0109 | CASP7,CASP9,S100A4 | |
| CCL22 | | cytokine | | | 0.0217 | CXCL2,CXCL3 | |
| HNRNPA2B1 | | other | | | 0.0000102 | ARSJ,CEMIP,CLDN1,GDF15,GSN,HIPK2,IGFBP2,ITPRID2,MEIS2,MUC13,PGM2L1,PITPNC1,PKIB,ROR1,SEMA3B,SERPINA1,SRI,TGFB2,TRIM31,USH1C,VIL1 | |
| AP2A1 | | transporter | | | 0.0135 | CEBPA,MT2A | |
| NKX3-1 | | transcription regulator | | | 0.00115 | CASP7,ETS1,FOS,PIK3R1,PIM1,SDC4,SOX4,STEAP1,VIL1 | 252 (11) |
| LPAR2 | | G-protein coupled receptor | | | 0.0428 | CXCL8,PTGS2 | |
| DLST | | enzyme | | | 0.00697 | JUN,PIK3R1 | |
| SND1 | | enzyme | | | 0.0000554 | EMP1,EREG,ETV1,PTGS2,QPCT,SLCO4A1,SMURF1,SOX4 | |
| IGBP1 | | phosphatase | | | 0.0109 | CLDN1,IRF1,TJP1 | |
| CD160 | | transmembrane receptor | | | 0.00789 | FOS,JUN,MCL1 | 236 (7) |
| LUC7L3 | | other | | | 0.049 | PTGS2 | |
| TOR2A | | other | | | 0.00697 | ACAT1,FOS | |
| CNOT7 | | transcription regulator | | | 7.93E-10 | B2M,CEACAM6,CLDN1,HERC6,IFI35,LGALS3BP,OAS3,PARP12,PLSCR1,PSMB8,SP110,TAP2,TFF3 | 200 (7) |
| CARD14 | | other | | | 0.0316 | BCL10,CXCL2 | |
| IFNGR2 | 0.403 | transmembrane receptor | | | 0.00697 | GBP3,IRF1 | 290 (10) |
| GFRA2 | | transmembrane receptor | | | 0.00697 | EGR1,FOSB | 165 (7) |
| HSPB8 | | kinase | | | 0.0185 | BMP4,CXCL8,MAP1LC3B | |
| TRIM33 | | transcription regulator | | | 0.0316 | GATA2,TGFBR2 | |
| NMU | | other | | | 0.0319 | FOSB,IL1R2,MYB,ZFP36L1 | |
| CITED1 | | transcription regulator | | | 0.0217 | CYP24A1,IGF2 | |
| OXTR | | G-protein coupled receptor | | | 0.0428 | FOS,RUNX2 | |
| GPD1 | -1.167 | enzyme | | | 0.0167 | ADGRF1,BCL6,CGREF1,CYP2B6,ELOVL6,GADD45A,GPCPD1,RARRES1,SERPINA3,SERPINE2 | |
| Cyp2c70 | | enzyme | | | 0.0232 | CYP3A5,HMGCR,HNF4A | |
| NFKBIB | | transcription regulator | | | 0.00454 | CARD6,CASP7,CASP9,CXCL1,CXCL2,CXCL8,DAPK1,EGR1 | 396 (15) |
| CFH | | other | | | 0.0316 | CXCL2,PPARG | |
| WBP1 | | other | | | 0.049 | MCL1 | |
| ZNF350 | | transcription regulator | | | 0.00697 | GADD45A,HMGA2 | 341 (7) |
| Zfp55 | | other | | | 0.049 | PPARG | |
| BMX | | kinase | | | 0.0285 | CXCL8,FOS,HSPA1A/HSPA1B | |
| ANGPTL3 | | growth factor | | | 0.0109 | ACSS2,ELOVL6,HSPG2 | |
| COMMD1 | | transporter | | | 0.0319 | BIRC3,DDIT4,EGLN1,LDHA | |
| KPNB1 | | transporter | | | 0.00697 | BBC3,PMAIP1 | |
| MAP3K10 | | kinase | | | 0.00354 | CXCL8,GLI1,GLI2 | |
| SLC27A1 | | transporter | | | 0.00697 | CEBPA,PPARG | |
| HOXB7 | | transcription regulator | | | 0.00095 | ANGPT1,CXCL1,CXCL8,REN | 229 (11) |
| GNB1 | | enzyme | | | 0.00259 | CXCL3,DUSP1,F2R,FOSB,IL1R2,JAG2,TSC22D3 | 277 (9) |
| IL4R | | transmembrane receptor | | | 0.0478 | ACSL1,ADAM8,FAS,FXYD3,PPARG,RUNX2 | |
| TFF1 | | other | | | 0.049 | PTGS2 | |
| TMSB10/TMSB4X | | other | | | 0.00109 | CXCL8,FAS,MMP7 | |
| RAD9A | | enzyme | | | 0.0428 | CDC25A,KLF12 | |
| RGD1560225 | | other | | | 0.000452 | EDN1,ITPR2,PTGS2 | |
| IGF2BP3 | | translation regulator | | | 0.00697 | CD44,IGF2 | 202 (7) |
| STK38 | | kinase | | | 0.0316 | CXCL2,CXCL3 | |
| DMTF1 | 0.372 | transcription regulator | | | 0.0428 | BBC3,DMTF1 | |
| MLLT3 | | transcription regulator | | | 0.0428 | EDN1,SGK1 | |
| F2RL3 | | G-protein coupled receptor | | | 0.0316 | EGR1,FOS | |
| GPS1 | | other | | | 0.0316 | AXIN2,FOS | |
| MECOM | | transcription regulator | | | 0.0165 | CEBPA,GATA2,ITPR2,MAP3K14,PML | |
| FABP5 | | transporter | | | 0.0478 | CXCL8,IFIT3,PPARG | |
| IL18RAP | | transmembrane receptor | | | 0.0428 | CXCL8,PTGS2 | |
| IL2RB | | transmembrane receptor | | | 0.0408 | ETS1,FOS,JUN | |
| BBC3 | 0.317 | other | | | 0.0217 | BBC3,STMN1 | |
| Pla2g2a | | other | | | 0.049 | PTGS2 | |
| PLA2G2A | | enzyme | | | 0.00354 | CD44,CXCL8,PTGS2 | 468 (20) |
| SHOX2 | | transcription regulator | | | 0.0316 | FGFR2,RUNX2 | |
| SIN3B | | transcription regulator | | | 0.0000423 | BCL6,CCNG2,FBXO32,GADD45B,KLF6,LDHA,TXNIP | 338 (7) |
| SRY | | transcription regulator | | | 0.0232 | BBC3,FOSL1,MAOA | |
| TNKS | | enzyme | | | 0.0428 | AXIN2,MCL1 | |
| VIPR1 | | G-protein coupled receptor | | | 0.0408 | CXCL2,CXCL3,IRF1 | |
| WAC | | other | | | 0.0316 | PDK4,PPARG | |
| MED13L | | other | | | 0.0428 | KITLG,LIF | |
| EPGN | | growth factor | | | 0.049 | FOS | |
| C5AR2 | | G-protein coupled receptor | | | 0.0101 | CXCL2,CXCL3,CXCL8,EIF2AK2 | |
| IFT20 | | other | | | 0.0135 | GLI1,GLI2 | |
| PLAGL2 | | transcription regulator | | | 0.0135 | IGF2,LDHA | |
| SP2 | | transcription regulator | | | 0.00507 | CEACAM1,FOS,MAT2A,SERPINH1 | |
| TAX1BP3 | | transcription regulator | | | 0.049 | FOS | |
| ADCYAP1R1 | | G-protein coupled receptor | | | 0.0428 | DUSP5,FOS | |
| LMNB1 | | other | | | 0.00454 | B2M,GPC4,HBEGF,HMGCR,MSMO1,RCN1,SQLE,TGFB2 | |
| NOXA1 | | other | | | 0.049 | NOX1 | |
| CCR4 | | G-protein coupled receptor | | | 0.0428 | CXCL3,FOS | |
| PDPN | | other | | | 0.049 | CD44 | |
| CRABP1 | | transporter | | | 0.049 | CRABP2 | |

Supplementary Table 4. Upstream regulator analysis based gene set enrichment analysis (GSEA) of differentially expressed genes (DEGs) after ATRA+PDT compared to PDT

| Upstream Regulator | Expr Log Ratio | Molecule Type | Predicted Activation State | Activation z-score | p-value of overlap | Target Molecules in Dataset | Mechanistic Network |
|---|---|---|---|---|---|---|---|
| FOXH1 | | transcription regulator | | | 0.0135 | ALDH1A1,ALDH1A3 | |
| MZF1 | | transcription regulator | | | 0.0232 | MYB,PADI1,PRKCA | |
| glaucocalyxin A | | chemical - endogenous non-mammalian | | | 0.0316 | LCN2,PTGS2 | |
| HOXB6 | | transcription regulator | | | 0.049 | REN | |
| YWHAB | | other | | | 0.0428 | BCAM,DUSP1 | |
| PPP3CA | | phosphatase | | | 0.0188 | CPT1A,MAOB,PLD1,PLPP3,PTGS2,RCAN1,RUNX2,S100A4,SDC4 | |
| PEA15 | | transporter | | | 0.0109 | JUN,PLD1,PLD2 | |
| UTS2R | | G-protein coupled receptor | | | 0.00789 | ACAT1,PPARG,SCARB1 | |
| RNF17 | | other | | | 0.0316 | EGLN1,SERPINA3 | |
| PON1 | | phosphatase | | | 0.0232 | CD68,PPARG,SCARB1 | |
| THEM4 | | enzyme | | | 0.049 | PTGS2 | |
| HIVEP3 | | transcription regulator | | | 0.0428 | ALG2,S100A4 | |
| DSPP | | other | | | 0.0316 | BGN,RUNX2 | |
| TAPBP | 0.646 | transporter | | | 0.00546 | HLA-B,HLA-E,TAP2 | |
| PELI2 | 1.015 | enzyme | | | 0.0217 | CXCL2,CXCL8 | |
| SDC1 | | enzyme | | | 0.0232 | CXCL2,CXCL3,TGM2 | |
| Tnxa-ps1 | | other | | | 0.049 | DUSP1 | |
| TFAP2B | | transcription regulator | | | 0.0478 | EDN1,IGF2,MT2A | |
| ENTPD1 | | enzyme | | | 0.0144 | CXCL2,CXCL3,CXCL8 | |
| RBP1 | | transporter | | | 0.0232 | ALDH1A1,PPARG,STRA6 | |
| BAG3 | 0.487 | other | | | 0.0185 | CDKN2B,MAP1LC3B,MCL1 | |
| IRAK2 | | kinase | | | 0.0408 | CXCL2,CXCL8,MAT2A | |
| CD99 | | other | | | 0.00789 | CAV1,FOSB,JUND | 394 (7) |
| ARHGDIB | 1.094 | enzyme | | | 0.049 | PTGS2 | |
| PTGER1 | | G-protein coupled receptor | | | 0.0232 | CXCL8,FOS,PTGS2 | |
| MAML1 | | transcription regulator | | | 0.0408 | CXCL8,EGR1,RND3 | |
| BCL10 | 0.695 | transcription regulator | | | 0.0408 | CD44,CXCL2,CXCL8 | |
| PEX5L | | ion channel | | | 0.0232 | HMGCR,IDI1,LSS | |
| CNR2 | | G-protein coupled receptor | | | 0.0363 | CXCL8,EGR1,GRK5,PTGS2 | |
| PBX1 | | transcription regulator | | | 0.0239 | AMD1,CDKN2B,ETV1,HOXA3,REN | |
| M6PR | | transporter | | | 0.049 | NPC1 | |
| CCX771 | | chemical reagent | | | 0.00546 | CD44,CXCR4,TJP1 | |
| TAF6 | | transcription regulator | | | 0.0461 | ATF3,GADD45A,IRF1,JUN | |
| ROR2 | | kinase | | | 0.000394 | AURKA,AXIN2,CDC25A,CDKN2B,CEBPA,GALM,MAP1LC3B,MYB,PPARG,SLC17A9,TEAD2,TRAF4 | |
| RARRES1 | 2.587 | other | | | 0.049 | GRK5 | |
| XPA | -0.522 | other | | | 0.0316 | FOS,PTGS2 | |
| 16:0 ceramide-1-phosphate | | chemical - endogenous mammalian | | | 0.049 | CXCL8 | |
| RAB7A | | enzyme | | | 0.0316 | EGR1,FOS | |
| ST14 | | peptidase | | | 0.0285 | CLDN2,CXCL2,CXCL8 | |
| cytarabine/idarubicin | | chemical drug | | | 0.049 | CEBPA | |
| RAB5B | | enzyme | | | 0.049 | FOS | |
| PAK1 | | kinase | | | 0.0268 | FBXO32,FOSL1,PFKM,SCARB1,TFPI | |
| ITIH4 | | other | | | 0.049 | VDR | |
| CRY1 | | enzyme | | | 0.0109 | CEBPA,FOS,SGK1 | |
| TOPBP1 | | other | | | 0.0343 | BBC3,CDKN2B,GADD45A | |
| HDC | | enzyme | | | 0.0478 | CYP24A1,FOS,PTGS2 | |
| CLEC11A | | growth factor | | | 0.019 | ANXA1,FHL2,FOS,LDHA,RUNX2,TSC22D1,ZYX | |
| GNPAT | 0.499 | enzyme | | | 0.0135 | UGCG,UGT8 | |
| CD164 | | other | | | 0.049 | CXCR4 | |
| cytarabine/filgrastim/fludarabine phosphate | | biologic drug | | | 0.049 | FOS | |
| ETS1-AS1 | | other | | | 0.049 | ETS1 | |
| TCL1A | | transcription regulator | | | 0.000386 | CD68,CTSE,CTSS,CXCL3,DNMT3A,FOS,GDF15,IL1R2,LCN2,MCL1,SEMA7A,STEAP4,TNFAIP2 | 447 (17) |
| HSF2 | | transcription regulator | | | 0.0333 | CLU,HSPA1A/HSPA1B,HSPB1,HSPH1,JUN | |
| ERC1 | | other | | | 0.00697 | CXCL8,PTGS2 | 317 (9) |
| CCDC80 | | other | | | 0.00546 | AXIN2,CEBPA,PPARG | 361 (12) |
| ERO1A | | enzyme | | | 0.0135 | CXCL2,CXCL3 | |
| NFKBIE | | transcription regulator | | | 0.0109 | CXCL3,CXCL8,LIF | |
| tofogliflozin | | chemical drug | | | 0.0217 | CPT1A,DGAT2 | |
| PRC1 | | other | | | 0.049 | PTGS2 | |
| GREM1 | | other | | | 0.0232 | AXIN2,RUNX2,WNT11 | |
| TFF3 | -4.091 | other | | | 0.0241 | CD55,CLDN1,FOS,TJP1 | |
| Gm35986 | | other | | | 0.0316 | HMGCR,IDI1 | |
| farnesyl transferase | | complex | | | 0.00546 | HBEGF,KLF6,MMP7 | 299 (7) |
| FLT3 | | kinase | | | 0.0218 | CDC25A,IFIT3,IRF7,MCL1,NOX1,PIM1 | |
| ESRRB | | transcription regulator | | | 0.041 | MCL1,NR0B2,PCSK6,PDIA5 | |
| PLAA | | other | | | 0.0217 | CLU,PTGS2 | |
| DYSF | | other | | | 0.0000268 | ANXA1,B2M,CD55,CD68,CTSS,DNAJB1,ERRFI1,HSPA1A/HSPA1B,HSPB1,LGALS3,PROS1,S100A4,SERPINA3,TCEAL9 | |
| tetramethylpyrazine | | chemical - endogenous non-mammalian | | | 0.049 | CXCR4 | |
| MST1R | | kinase | | | 0.0123 | CXCL3,DVL1,FOS,JUN | |
| ADCY3 | | enzyme | | | 0.049 | NRP1 | |
| SERPIND1 | | other | | | 0.0316 | EGR1,F2R | |
| IFNLR1 | | transmembrane receptor | | | 0.0000623 | ATF3,CXCL2,CXCL3,IFI44,IFIH1,IFIT3,IRF7,LIF,PTGS2,TNFAIP2 | 313 (13) |
| glyceraldehyde-BSA | | chemical reagent | | | 0.0428 | CEBPA,PPARG | |
| methylglyoxal-BSA | | chemical reagent | | | 0.0428 | CEBPA,PPARG | |
| HESX1 | | transcription regulator | | | 0.00354 | AXIN2,GATA2,MYB | |
| KIT | | transmembrane receptor | | | 0.0015 | CD44,CDKN2B,CXCL3,CXCR4,ETV1,IFIT3,IL15RA,IRF1,KITLG,LGALS3,LIF,MCL1,RAB27B,TJP1 | 522 (20) |
| GTx-560 | | chemical reagent | | | 0.049 | PROS1 | |
| EXOSC10 | | kinase | | | 0.049 | FOS | |
| EHMT2 | | transcription regulator | | | 0.0174 | BCAS3,MAP1LC3B,NCOA3,PKIB,PMAIP1,PPL,PTGS2 | |
| WNT10B | | other | | | 0.0144 | CEBPA,PPARG,RUNX2 | |
| CDKL2 | | kinase | | | 0.049 | CD44 | |
| GNAQ | | enzyme | | | 8.05E-09 | AK4,ATF3,BMP4,CD44,CSPG4,CXCL3,CXCL8,DUSP5,EDN1,EGR1,F2RL1,FOS,GADD45A,IL1RN,INSIG1,NDRG2,PHLDA1,PLSCR1,PTGS2,RHOB,SGK1,SLCO4A1,TGFB2,ZFP36L1 | 481 (19) |
| RING1 | | transcription regulator | | | 0.00697 | FOS,JUN | 278 (6) |
| EEF1A1 | | translation regulator | | | 0.00697 | CCNA2,S100A4 | |
| CD300C | | transmembrane receptor | | | 0.0316 | CXCL2,CXCL8 | |
| CAV2 | -0.516 | other | | | 0.049 | CAV1 | |
| SLC25A13 | | transporter | | | 0.0179 | ADGRF1,BCL6,CGREF1,CYP2B6,ELOVL6,GADD45A,GPCPD1,RARRES1,SERPINA3,SERPINE2 | |
| SRRM1 | | other | | | 0.049 | CD44 | |
| CHRM2 | | G-protein coupled receptor | | | 0.049 | EGR1 | |
| ACP1 | | phosphatase | | | 0.0135 | EPHA2,FOS | |
| NEK2 | | kinase | | | 0.0135 | FAS,GADD45A | |
| Mt3 | | other | | | 0.0135 | CEBPD,MT2A | |
| BEX2 | | other | | | 0.0185 | BBC3,ITGB6,SWAP70 | |
| H4C3 | | other | | | 0.0428 | FOSB,TGFBR2 | |
| CDC42 | | enzyme | | | 0.0188 | ATF3,FOS,JUN,SCARB1,TGM2 | |
| Cyp2a12/Cyp2a22 | | enzyme | | | 0.0135 | CYP3A5,HNF4A | |
| Flt1 | | kinase | | | 0.049 | ATF3 | |
| GNMT | | enzyme | | | 0.00354 | CYP1A1,NPC1,SCARB1 | 301 (7) |
| HSPA1A/HSPA1B | 0.551 | enzyme | | | 0.0232 | CXCL2,CXCL8,FOS | |
| Ifna4 | | other | | | 0.0316 | IRF7,TRIM38 | |
| Irs4 | | other | | | 0.0024 | EGR1,IRS2 | 405 (17) |
| PPP1R15A | 0.574 | other | | | 0.00295 | CA9,CEBPA,GADD45A,MAP1LC3B,MCL1 | |

Supplementary Table 4. Upstream regulator analysis based gene set enrichment analysis (GSEA) of differentially expressed genes (DEGs) after ATRA+PDT compared to PDT

| Upstream Regulator | Expr Log Ratio | Molecule Type | Predicted Activation State | Activation z-score | p-value of overlap | Target Molecules in Dataset | Mechanistic Network |
|---|---|---|---|---|---|---|---|
| Cxcl3 | | cytokine | | | 0.00109 | CEBPA,CEBPD,PPARG | |
| TCF7 | | transcription regulator | | | 0.00754 | AXIN2,BCL6,CEBPA,CEBPD,FOS,HNF4A,PPARG | 242 (9) |
| PAXIP1 | | other | | | 0.0316 | CSPG4,DKK1 | |
| Paxip1 | | other | | | 0.0021 | ANXA1,CEACAM1,REL | |
| TWSG1 | | other | | | 0.0217 | BCL6,ID2 | |
| OBP2B | | transporter | | | 0.0316 | CPT1A,PPARG | |
| HMGB2 | | transcription regulator | | | 0.0078 | CXCL1,CXCL2,CXCL8,PHLDA1,RUNX2 | 375 (9) |
| NKRF | | transcription regulator | | | 0.0428 | CXCL8,PTGS2 | |
| H1-4 | | other | | | 0.0217 | GLI1,GLI2 | |
| H1f4 | | other | | | 0.00109 | AXIN2,GLI1,GLI2 | |
| RFX5 | | transcription regulator | | | 0.0343 | B2M,GCNT2,HLA-B | |
| S100A7 | | other | | | 0.0144 | CA9,CXCL1,CXCL8 | |
| PPP2R5B | 0.609 | phosphatase | | | 0.0217 | SAT1,SERPINE2 | |
| GCLM | | enzyme | | | 0.0217 | FOS,JUN | |
| GNRH2 | | other | | | 0.00789 | FOS,FOSB,JUN | 498 (22) |
| MPT0E028 | | chemical drug | | | 0.049 | MCL1 | |
| HDAC8 | | transcription regulator | | | 0.0109 | BBC3,HOXA5,STX1A | |
| PEX14 | | transcription regulator | | | 0.049 | PTGS2 | |
| DEK | | transcription regulator | | | 0.00789 | BIRC3,CXCL8,MCL1 | 309 (7) |
| ID4 | | transcription regulator | | | 0.00387 | BBC3,CDKN2B,CEBPA,PPARG | 248 (7) |
| ZYX | 0.622 | other | | | 0.0316 | FOS,HIPK2 | |
| GNA15 | | enzyme | | | 0.000000123 | AKAP12,BMP4,CASP4,CAV2,CEBPD,CXCL2,CXCL3,CXCL8,FHL2,FOSL1,GADD45A,HLA-E,IL1RN,LGALS3,MAT2A,NOX1,PDLIM1,PTGS2,RGS2,SERPINA3,ZFP36L1 | 413 (16) |
| spliceostatin A | | chemical reagent | | | 0.049 | MCL1 | |
| oligomycin A | | chemical - endogenous non-mammalian | | | 0.049 | TXNIP | |
| Rp-cAMP triethylamine | | chemical reagent | | | 0.049 | CXCL3 | |
| CML advanced glycation end products | | chemical reagent | | | 0.049 | EGR1 | |
| SR11256 | | chemical reagent | | | 0.049 | EDN1 | |
| cAMP-dependent protein kinase inhibitor | | chemical drug | | | 0.049 | CYP3A5 | |
| HADHB | | enzyme | | | 0.049 | REN | |
| SOD1 | | enzyme | | | 0.00563 | ACO1,ASS1,B2M,BAG3,BMP4,CASP4,CKB,CLU,FAS,FGF19,FOS,GUK1,HSPB1,JUN,MAOB,MCL1,PSMB8,PTGS2,PTPN21,RUNX2,SNCG,STMN1,TGFBR2 | 483 (21) |
| 3,4-(methylenedioxy)cinnamic acid | | chemical reagent | | | 0.049 | MCL1 | |
| CC-122 | | chemical drug | | | 0.0217 | IFIT3,IRF7 | |
| polyinosine-polycytidylic acid/polyethylenimine formulation | | chemical reagent | | | 0.0217 | IFIH1,PMAIP1 | |
| COG112 peptide | | chemical reagent | | | 0.0316 | CXCL2,CXCL3 | |
| STF 083010 | | chemical reagent | | | 0.049 | NCOA3 | |
| Immunoglobulin Lambda Light Chain | | group | | | 0.0428 | ERN1,PMAIP1 | |
| RB4 | | chemical reagent | | | 0.049 | MCL1 | |
| ciliobrevin A | | chemical reagent | | | 0.0024 | GLI1,GLI2 | |
| RB3 | | chemical reagent | | | 0.049 | MCL1 | |
| HNRNPAB | | enzyme | | | 0.0185 | PTGS2,S100A4,TJP1 | |
| J11-Cl | | chemical reagent | | | 0.0144 | CXCL2,CXCL3,PTGS2 | |
| KMUP-1 | | chemical reagent | | | 0.0144 | HMGCR,PPARG,SCARB1 | |
| CH-223191 | | chemical reagent | | | 0.0408 | CYP1A1,NT5E,PTGS2 | |
| MLS000389544 | | chemical reagent | | | 0.0024 | PCK2,PDK4 | |
| NCGC00184710 | | chemical reagent | | | 0.049 | PDK4 | |
| finerenone | | chemical drug | | | 0.0217 | LCN2,LGALS3 | |
| arsenic trichloride | | chemical reagent | | | 0.049 | GADD45A | |
| SYK023 | | chemical reagent | | | 0.0135 | CDC25A,CDKN2B | |
| meayamycin B | | chemical reagent | | | 0.049 | MCL1 | |
| 1-(2-hydroxy-5-methylphenyl)-3-phenyl-1,3-propanedione | | chemical reagent | | | 0.0135 | CEBPD,PPARG | |
| 24-methylenecycloartanyl ferulate | | chemical reagent | | | 0.049 | PPARG | |
| BCL201 | | chemical drug | | | 0.049 | MCL1 | |
| KRT7-AS | | other | | | 0.049 | KRT7 | |
| acriflavine | | chemical toxicant | | | 0.0144 | CA9,NT5E,PTPRB | |
| beractant | | chemical drug | | | 0.049 | CXCL8 | |
| sulfamethoxazole/trimethoprim | | chemical drug | | | 0.0217 | CXCL8,IRF7 | |
| asbestos | | chemical toxicant | | | 0.00789 | CXCL8,FOSL1,JUN | |
| hydroxyl radical | | chemical toxicant | | | 0.0024 | CAV1,KLF4 | |
| AP20187 | | chemical reagent | | | 0.0428 | EGR1,FOS | |
| furosemide | | chemical drug | | | 0.0217 | PTGS2,REN | |
| enzastaurin | | chemical drug | | | 0.0316 | CXCL8,TJP1 | |
| halofuginone | | chemical drug | | | 0.0000041 | AKAP12,AOC1,BGN,CD44,CD55,CKB,CLIP2,CTSS,EGR1,EMP1,F2R,GYG1,HK1,IFITM2,LGALS3BP,PCSK6,PLPP3,SLC39A8,ST3GAL2,STEAP3,TGFBR2,TGM2 | |
| NBI 27914 | | chemical drug | | | 0.049 | FOS | |
| nicorandil | | chemical drug | | | 0.0428 | ATF3,JUN | |
| diltiazem | | chemical drug | | | 0.0109 | EDN1,FOS,JUN | |
| benzoquinone | | chemical - endogenous mammalian | | | 0.0428 | APOL1,IRF1 | |
| probenecid | | chemical drug | | | 0.049 | ABCC1 | |
| thenoyltrifluoroacetone | | chemical reagent | | | 0.0428 | BIRC3,CXCL8 | |
| carboxyamido-triazole | | chemical drug | | | 0.0135 | BAG3,FOS | |
| LY379196 | | chemical - kinase inhibitor | | | 0.0217 | CD55,EGR1 | |
| cantharidin | | chemical drug | | | 0.0232 | BAG3,MCL1,SERPINH1 | |
| SB202474 | | chemical - kinase inhibitor | | | 0.049 | CYP1A1 | |
| EGTA acetoxymethyl ester | | chemical reagent | | | 0.0316 | CXCL8,PTGS2 | |
| 2-aminoethoxydiphenylborane | | chemical reagent | | | 0.0144 | CYP1A1,ITPR2,PTGS2 | |
| S-methylisothiopseudouronium | | chemical reagent | | | 0.049 | PTGS2 | |
| phenylacetate | | chemical - endogenous mammalian | | | 0.0135 | CAV1,CAV2 | |
| itraconazole | | chemical drug | | | 0.00546 | AGR2,CYP1A1,GLI1 | |
| ifosfamide | | chemical drug | | | 0.0316 | CYP2B6,SLC22A5 | |
| N-(3-oxododecanoyl)-homoserine lactone | | chemical reagent | | | 0.0285 | CXCL2,CXCL8,PTGS2 | |
| dichlorodiphenyl dichloroethylene | | chemical toxicant | | | 0.00697 | CYP2B6,CYP3A5 | 441 (12) |
| trabectedin | | chemical drug | | | 0.00295 | CCNG2,GADD45A,H1-10,LGMN,PPARG,SAT1 | |
| guanfacine | | chemical drug | | | 0.049 | FOS | |
| phenylpropanoic acid | | chemical - endogenous mammalian | | | 0.049 | ATP2A3 | |
| amantadine | | chemical drug | | | 0.00697 | DDC,FOS | |
| capsazepine | | chemical toxicant | | | 0.00789 | CXCL8,FOS,PTGS2 | 430 (21) |
| AR-A014418 | | chemical reagent | | | 0.0428 | CD44,GDF15 | |
| raltitrexed | | chemical drug | | | 0.00789 | FAS,FXYD3,SAT1 | |
| atipamezole | | chemical drug | | | 0.0135 | EGR1,FOS | |
| antimony potassium tartrate | | chemical toxicant | | | 0.049 | ABCC2 | |
| coomassie brilliant blue | | chemical drug | | | 0.0285 | LIF,PTGS2,TJP1 | |
| XCT790 | | chemical reagent | | | 0.0316 | MAOB,PDK4 | |
| pyridoxine | | chemical - endogenous mammalian | | | 0.0343 | DKK1,JUN,PTGS2 | |
| bisphenol A diglycidyl ether | | chemical reagent | | | 0.0185 | EGLN1,TGFBR2,VDR | |
| alpha-ketoisocaproic acid | | chemical - endogenous mammalian | | | 0.0316 | ATP2B1,ITPR2 | |
| clioquinol | | chemical drug | | | 0.0144 | FOS,JUN,KDM6B | |
| nimesulide | | chemical drug | | | 0.0478 | CYP1A1,EREG,PTGS2 | |
| cimetidine | | chemical drug | | | 0.0217 | FAS,GDF15 | |
| SR 144528 | | chemical reagent | | | 0.0185 | CXCL8,EGR1,IL1RN | |
| bupivacaine | | chemical drug | | | 0.0109 | CASP9,FOSL1,JUND | |
| 4-coumaric acid | | chemical - endogenous mammalian | | | 0.0428 | CCNA2,PTGS2 | |
| ferric nitrilotriacetate | | chemical toxicant | | | 0.0316 | KLF4,PTGS2 | |
| 2,5-dihydroxymethylcinnamate | | chemical - kinase inhibitor | | | 0.049 | EDN1 | |

Supplementary Table 4. Upstream regulator analysis based gene set enrichment analysis (GSEA) of differentially expressed genes (DEGs) after ATRA+PDT compared to PDT

| Upstream Regulator | Expr Log Ratio | Molecule Type | Predicted Activation State | Activation z-score | p-value of overlap | Target Molecules in Dataset | Mechanistic Network |
|---|---|---|---|---|---|---|---|
| LY311727 | | chemical reagent | | | 0.049 | PTGS2 | |
| BN 50730 | | chemical reagent | | | 0.049 | PTGS2 | |
| saccharin | | chemical reagent | | | 0.00697 | CEBPA,PPARG | 344 (7) |
| sterigmatocystin | | chemical toxicant | | | 0.049 | CYP1A1 | |
| hoechst 33342 | | chemical reagent | | | 0.0135 | EPC1,FOS | |
| emricasan | | chemical drug | | | 0.0316 | CXCL2,CXCL3 | |
| tecalcet | | chemical reagent | | | 0.049 | EDN1 | |
| BW-755C | | chemical drug | | | 0.049 | JUND | |
| ondansetron | | chemical drug | | | 0.049 | FOS | |
| phenidone | | chemical drug | | | 0.049 | REN | |
| cyclothiazide | | chemical drug | | | 0.049 | EGR1 | |
| flurothyl | | chemical drug | | | 0.049 | FOS | |
| carbonyl cyanide p-(trifluoromethoxy)phenylhydrazone | | chemical reagent | | | 0.0285 | CHAC1,PCK2,PSPH | |
| pustulan | | chemical reagent | | | 0.049 | CXCL8 | |
| trichloroacetic acid | | chemical drug | | | 0.049 | MCL1 | |
| L 733060 | | chemical reagent | | | 0.049 | FOS | |
| lansoprazole | | chemical drug | | | 0.0135 | CXCL8,PTGS2 | |
| 1-(1-glycero)dodeca-1,3,5,7,9-pentaene | | chemical toxicant | | | 0.049 | PTGS2 | |
| volinanserin | | chemical drug | | | 0.00354 | EGR1,FOS,REN | 231 (7) |
| phenyl-N-tert-butylnitrone | | chemical reagent | | | 0.0478 | CXCL8,HSPA1A/HSPA1B,PTGS2 | |
| warfarin | | chemical drug | | | 0.0428 | PROCR,PROS1 | |
| n-nitrosomethylbenzylamine | | chemical toxicant | | | 0.0172 | ADAMTS6,AKR1B10,AMBP,CD44,CLDN1,DUSP5,KRT20,MCL1,PIP5K1A,PLK3,PTGS2,SCEL,SLC30A1 | |
| L-threo-safingol | | chemical drug | | | 0.049 | CXCL8 | |
| CHS-828 | | chemical drug | | | 0.00546 | ALDH1A1,CD44,LRIG1 | |
| selfotel | | chemical drug | | | 0.049 | FOS | |
| liarozole | | chemical drug | | | 0.0135 | PI3,VDR | |
| alpha-methylhydrocinnamic acid | | chemical - endogenous mammalian | | | 0.0135 | MCL1,MYB | |
| tetrabenazine | | chemical drug | | | 0.049 | PPP1R1B | |
| manumycin A | | chemical reagent | | | 0.0144 | CXCL8,PLD1,PTGS2 | |
| ibotenic acid | | chemical toxicant | | | 0.0135 | FOS,JUN | |
| methylselenic acid | | chemical reagent | | | 8.19E-09 | [illegible]4,CASP9,CCNA2,CD44,CDC14B,CDC25A,CLU,DNAJB9,DNAJC3,ELF3,GADD45A,GATA2,GDF15,H1-0,ID2,IRF1,JUN,KLF4,KRT20,MCL1,MXD1,NEDD4L,NMI,PATZ1,PRKCA,RHOB,[illegible] | 566 (22) |
| butylated hydroxytoluene | | chemical toxicant | | | 0.00109 | FOS,JUN,PTGS2 | 326 (13) |
| (R)-5-(2-azetidnylmethoxy)-2-chloropyridine | | chemical drug | | | 0.049 | FOS | |
| 6-cyano-7-nitroquinoxaline-2,3-dione | | chemical reagent | | | 0.0021 | EGR1,FOS,JUN | 186 (7) |
| ethacrynic acid | | chemical drug | | | 0.0217 | AKR1C1/AKR1C2,MCL1 | |
| RWJ 67657 | | chemical - kinase inhibitor | | | 0.0428 | CXCL8,PTGS2 | |
| rofecoxib | | chemical drug | | | 0.0478 | PTGS2,REN,TBC1D9 | |
| dibenzoylmethane | | chemical toxicant | | | 0.0316 | CYP1A1,PTGS2 | |
| zimelidine | | chemical drug | | | 0.0428 | DEPP1,SERPINA3 | |
| D609 | | chemical reagent | | | 0.0478 | CXCL8,JUN,PTGS2 | |
| bergamottin | | chemical - endogenous non-mammalian | | | 0.0428 | CYP1A1,CYP2B6 | |
| clomipramine | | chemical drug | | | 0.0109 | DEPP1,LSS,SERPINA3 | |
| barbiturate | | chemical drug | | | 0.0135 | CYP2B6,CYP3A5 | |
| BTG3-AS1 | | other | | | 0.049 | ATF3 | |
| clofarabine | | chemical drug | | | 0.049 | CDC25A | |
| histone deacetylase | | complex | | | 0.0446 | CCN1,CXCR4,CYP24A1,ID2,NDRG2 | |
| Hif | | complex | | | 0.0408 | CKB,CXCL8,LDHA | |
| D-sphingosine | | chemical - endogenous mammalian | | | 0.0185 | CLDN1,CXCL2,PTGS2 | |
| phytosphingosine | | chemical - endogenous mammalian | | | 0.049 | CLDN1 | |
| apomine | | chemical drug | | | 0.00697 | HMGCR,NR0B2 | |
| PK11007 | | chemical reagent | | | 0.0316 | BBC3,PMAIP1 | |
| aspergillus beta-glucan | | chemical - endogenous non-mammalian | | | 0.049 | CXCL8 | |
| pneumocystis beta-glucan | | chemical - endogenous non-mammalian | | | 0.049 | CXCL8 | |
| cortistatin A | | chemical reagent | | | 0.0109 | CEBPA,ETV6,IRF1 | |
| sevelamer | | chemical drug | | | 0.0316 | FGF19,NR0B2 | |
| diazepam | | chemical drug | | | 0.0428 | CXCL8,FOS | |
| flunitrazepam | | chemical drug | | | 0.049 | CXCL8 | |
| oxazepam | | chemical drug | | | 0.0024 | GADD45B,TSC22D1 | |
| thiabendazole | | chemical drug | | | 0.049 | CYP1A1 | |
| methylnitrosourea | | chemical toxicant | | | 0.00134 | ATP1B3,AURKA,CLU,CXCL3,EGR1,JUN,PSME1 | 467 (18) |
| arachidonyl-2-chloroethylamide | | chemical reagent | | | 0.049 | EGR1 | |
| glyburide | | chemical drug | | | 0.0478 | FOS,PPARG,SCARB1 | |
| naphthalene-1-2-dione | | chemical toxicant | | | 0.049 | CYP1A1 | |
| trans-7,8-dihydroxy-7,8-dihydrobenzo(a)pyrene | | chemical toxicant | | | 0.049 | CYP1A1 | |
| anthracene | | chemical toxicant | | | 0.049 | CYP1A1 | |
| tolrestat | | chemical drug | | | 0.049 | CXCL8 | |
| MUC2 | | other | | | 0.049 | CXCL8 | |
| dieldrin | | chemical toxicant | | | 0.0316 | CYP2B6,CYP3A5 | |
| DDT | | chemical toxicant | | | 0.0428 | CYP2B6,CYP3A5 | |
| naftifine | | chemical drug | | | 0.049 | SQLE | |
| 2-amino-3-methylimidazo(4,5-f)quinoline | | chemical toxicant | | | 0.049 | CYP1A1 | |
| benzo(a)pyrene-7,8-dione | | chemical toxicant | | | 0.0024 | AKR1C1/AKR1C2,CYP1A1 | |
| Ro 31-7549 | | chemical - kinase inhibitor | | | 0.049 | PTGS2 | |
| ketorolac | | chemical drug | | | 0.00789 | CXCL8,PTGS2,REN | 411 (20) |
| ethyl linoleate | | chemical - endogenous mammalian | | | 0.0135 | PTGS2,SC5D | |
| Ki16425 | | chemical reagent | | | 0.0428 | PLPP3,PTGS2 | |
| ONO-8711 | | chemical drug | | | 0.049 | FOS | |
| nitrendipine | | chemical drug | | | 0.000452 | CYP2B6,CYP3A5,HMGCR | |
| acetic acid | | chemical - endogenous mammalian | | | 0.0109 | EGR1,FOS,JUN | |
| isobutyric acid | | chemical - endogenous non-mammalian | | | 0.049 | ATP2A3 | |
| catechol | | chemical - endogenous mammalian | | | 0.0316 | FOS,JUN | |
| mesalamine | | chemical drug | | | 0.0109 | HPGD,PPARG,PTGS2 | |
| eugenol | | chemical - endogenous non-mammalian | | | 0.0428 | CXCL8,PTGS2 | |
| theaflavin monogallate B | | chemical - endogenous non-mammalian | | | 0.049 | PTGS2 | |
| propyl gallate | | chemical toxicant | | | 0.0316 | CXCL8,PTGS2 | |
| epicatechin gallate | | chemical drug | | | 0.0241 | CXCL8,GDF15,PTGS2,TP53I11 | |
| taprostene | | chemical reagent | | | 0.00697 | DUSP1,TSC22D3 | |
| AP-1 decoy | | chemical reagent | | | 0.049 | DUSP1 | |
| lavendustin A | | chemical - kinase inhibitor | | | 0.0024 | CYP1A1,EDN1 | |
| vanadyl sulfate | | chemical reagent | | | 0.0217 | IL1R2,IL1RN | |
| potassium iodide | | chemical drug | | | 0.049 | CXCL8 | |
| MitoBloCK-6 | | chemical drug | | | 0.00697 | PCK2,PSPH | |
| mercuric chloride | | chemical toxicant | | | 0.0299 | GADD45A,HBEGF,HSPA1A/HSPA1B,HSPB1,JUN | |
| spermine nitric oxide complex | | chemical toxicant | | | 0.0232 | ACO1,CXCL8,S100A10 | |
| chlorophyll a | | chemical - endogenous non-mammalian | | | 0.0135 | FAS,PPARG | |
| zinc protoporphyrin IX | | chemical - endogenous mammalian | | | 0.0343 | ANGPT1,FGFR2,PTGS2 | |
| senexin B | | chemical drug | | | 0.000117 | CXCL1,CXCL2,CXCL8 | 117 (5) |
| RUNX1-RUNX1T1 | | fusion gene/product | | | 0.000892 | CEBPA,CYP2S1,GRK5,ID2,IFITM3,KCNE3,KLHL24,MT2A,MTSS1,RND3,SH2B3,SORL1,SOX4 | 315 (11) |
| NUP98-KDM5A | | fusion gene/product | | | 0.0343 | CDK6,MCL1,MYB | |
| FUS-ERG | | fusion gene/product | | | 0.049 | PIM1 | |
| EWSR1-ERG | | fusion gene/product | | | 0.049 | PIM1 | |
| NUP98-NSD1 | | fusion gene/product | | | 0.0343 | CDK6,MCL1,MYB | |

Supplementary Table 4. Upstream regulator analysis based gene set enrichment analysis (GSEA) of differentially expressed genes (DEGs) after ATRA+PDT compared to PDT

| Upstream Regulator | Expr Log Ratio | Molecule Type | Predicted Activation State | Activation z-score | p-value of overlap | Target Molecules in Dataset | Mechanistic Network |
|---|---|---|---|---|---|---|---|
| PML-RARA | | fusion gene/product | | | 0.0299 | EGR1,FOS,JUN,PSMB10,PSMB8 | |
| retroinverso ERG inhibitory peptide 2 | | chemical reagent | | | 0.0428 | ARHGDIB,CXCR4 | |
| retroinverso ERG inhibitory peptide 1 | | chemical reagent | | | 0.0428 | ARHGDIB,CXCR4 | |
| miR-124 inhibitor | | chemical reagent | | | 0.049 | MCL1 | |
| anti-miR-32 inhibitor | | chemical reagent | | | 0.049 | KLF4 | |
| WZB117 | | chemical reagent | | | 0.0135 | CXCL3,PI3 | |
| cathelicidin-WA | | chemical - endogenous mammalian | | | 0.00109 | MUC2,TFF3,TJP1 | |
| dexamethasone/INS/isobutylmethylxanthine | | chemical reagent | | | 0.049 | CCNA2 | |
| BIM5078 | | chemical reagent | | | 0.049 | SULT2B1 | |
| pubchem compound 25247557 | | chemical reagent | | | 0.0135 | PDK4,PEX11A | |
| REC2923 | | chemical reagent | | | 0.00697 | LDHA,SLC16A1 | |
| tazarotene | | chemical drug | | | 0.0217 | PLAAT4,RARRES1 | |
| retinaldehyde | | chemical - endogenous mammalian | | | 0.0428 | CD44,PTGS2 | |
| thioridazine | | chemical drug | | | 0.0123 | DEPP1,LSS,MCL1,SERPINA3 | |
| CB-PIC | | chemical reagent | | | 0.0217 | ABCB10,CASP9 | |
| ethylenediaminetetraacetic acid | | chemical drug | | | 0.0232 | CXCL8,EGR1,MCAM | |
| spermidine | | chemical - endogenous mammalian | | | 0.00651 | AMD1,FOS,IGFBP2,SAT1 | |
| spermine | | chemical - endogenous mammalian | | | 0.0285 | IGFBP2,RUNX2,SAT1 | |
| meclizine | | chemical drug | | | 0.0285 | ABCC2,CYP2B6,CYP3A5 | |
| CTOP | | chemical reagent | | | 0.049 | FOS | |
| apamin | | chemical toxicant | | | 0.0316 | FOS,JUN | |
| BMS200261 | | chemical reagent | | | 0.049 | CD55 | |
| JMF3086 | | chemical reagent | | | 0.00789 | CXCL2,CXCL3,PTGS2 | 451 (16) |
| astressin 2B | | biologic drug | | | 0.0232 | CXCL2,FOS,HPGD | |
| Z-IETD-FMK | | chemical - protease inhibitor | | | 0.0316 | CXCL8,PLD1 | |
| N-t-Boc-Phe-D-Leu-Phe-D-Leu-Phe | | chemical reagent | | | 0.049 | PTGS2 | |
| N-carbobenzyloxy-leucine-leucine-norvalinal | | chemical - protease inhibitor | | | 0.0343 | CXCL8,MCL1,RARG | |
| BQ-788 | | biologic drug | | | 0.0109 | CXCL2,EDN1,MCAM | |
| myristoylated Protein Kinase C peptide inhibitor | | chemical - kinase inhibitor | | | 0.0135 | CD55,PPARG | |
| NCB-0846 | | chemical - kinase inhibitor | | | 0.049 | JUN | |
| Ac-DEVD-CHO | | chemical reagent | | | 0.049 | FOS | |
| aclarubicin | | chemical drug | | | 0.0217 | MCL1,PTGS2 | |
| idarubicin | | chemical drug | | | 0.0123 | BBC3,GADD45A,MCL1,SESN2 | |
| ibutamoren | | chemical drug | | | 0.049 | FOS | |
| trandolapril | | chemical drug | | | 0.049 | REN | |
| bisindolylmaleimide II | | chemical - kinase inhibitor | | | 0.0135 | JUN,PPARG | |
| linalool | | chemical - endogenous non-mammalian | | | 0.0176 | ACSL1,HMGCR,PCK2,SQLE | |
| geranylgeraniol | | chemical - endogenous mammalian | | | 0.0428 | HMGCR,MCL1 | |
| ETC-642 | | chemical reagent | | | 0.049 | HMGCR | |
| 18-alpha-glycyrrhetinic acid | | chemical - endogenous non-mammalian | | | 0.0316 | PPARG,TJP1 | |
| hyperforin | | chemical drug | | | 0.0343 | ABCC2,CYP24A1,CYP2B6 | |
| myricetin | | chemical - endogenous non-mammalian | | | 0.00651 | CYP1A1,EGLN1,PTGS2,SLC16A1 | 329 (7) |
| kaempferol | | chemical toxicant | | | 0.0319 | CYP1A1,JUN,PTGS2,RUNX2 | |
| cytisine | | chemical drug | | | 0.049 | FOSB | |
| naltrindole | | chemical drug | | | 0.0285 | IGF2,IGFBP2,UBC | |
| atropine | | chemical drug | | | 0.0428 | FOS,PTGS2 | |
| SN-38 | | chemical drug | | | 0.00655 | BIRC3,CASP9,CLU,FAS,GDF15,MCL1 | 395 (11) |
| yohimbine | | chemical drug | | | 0.0135 | EGR1,FOS | |
| NHB2005 | | chemical reagent | | | 0.0316 | CASP9,ERN1 | |
| tetrahydropalmatine | | chemical - endogenous non-mammalian | | | 0.0024 | CXCL8,PTGS2 | |
| noscapine | | chemical drug | | | 0.0285 | BIRC3,MCL1,PTGS2 | |
| omacetaxine mepesuccinate | | chemical drug | | | 0.0217 | CASP9,MCL1 | |
| arecoline | | chemical drug | | | 0.0207 | F2R,MAOA,S100A4,TJP1 | |
| 2-methoxy-5-(2',3',4'-trimethoxyphenyl)-2,4,6-cycloheptatrien-1-one | | chemical reagent | | | 0.049 | JUN | |
| harmaline | | chemical drug | | | 0.049 | CYP1A1 | |
| D-lysergic acid diethylamide | | chemical drug | | | 0.0024 | DDC,SGK1 | |
| LY 354740 | | chemical drug | | | 0.049 | FOS | |
| droxidopa | | chemical drug | | | 0.049 | FOS | |
| S(-)-carbidopa | | chemical drug | | | 0.049 | FOS | |
| tyrphostin AG 1296 | | chemical - kinase inhibitor | | | 0.0144 | CEACAM6,JUN,PTGS2 | |
| phosphatidic acid | | chemical - endogenous mammalian | | | 0.0428 | IGF2,PTGS2 | |
| isovaleric acid | | chemical - endogenous mammalian | | | 0.049 | ATP2A3 | |
| oleoylethanolamide | | chemical - endogenous mammalian | | | 0.00789 | CPT1A,FOS,FOSB | 449 (25) |
| tridecanoic acid | | chemical - endogenous mammalian | | | 0.0109 | DUSP5,KLF4,MXD1 | |
| 10-hydroxydecanoic acid | | chemical - endogenous non-mammalian | | | 0.049 | IRF1 | |
| n-6 docosapentaenoic acid | | chemical - other | | | 0.049 | PTGS2 | |
| fumagillin | | chemical - endogenous non-mammalian | | | 0.049 | CAV1 | |
| long-chain alcohol | | chemical - endogenous non-mammalian | | | 0.049 | PTGS2 | |
| 12-hydroxyeicosatetraenoic acid | | chemical - endogenous mammalian | | | 0.0428 | NOX1,PTGS2 | |
| 3'-adenylic acid | | chemical - endogenous mammalian | | | 0.049 | TXNIP | |
| stavudine | | chemical drug | | | 0.00697 | CPT1A,PPARG | |
| gambogic acid | | chemical - endogenous non-mammalian | | | 0.0232 | BIRC3,EGLN1,PTGS2 | |
| poractant alfa | | chemical drug | | | 0.049 | CXCL8 | |
| tosyllysine chloromethyl ketone | | chemical - protease inhibitor | | | 0.00546 | CYP1A1,IRF1,PTGS2 | 374 (14) |
| OSU8-1 | | chemical - protease inhibitor | | | 0.049 | HBEGF | |
| 5-hydroxytryptophan | | chemical - endogenous mammalian | | | 0.0217 | FOS,PTGS2 | |
| sarcosine | | chemical - endogenous mammalian | | | 0.049 | FOS | |
| 2-amino-3-phosphonopropionic acid | | chemical - endogenous mammalian | | | 0.00697 | EGR1,FOS | 300 (7) |
| quisqualic acid | | chemical - endogenous non-mammalian | | | 0.0316 | EGR1,FOS | |
| pectin | | chemical drug | | | 0.0144 | CD68,LGALS3,SMAD3 | |
| allose | | chemical - endogenous mammalian | | | 0.049 | TXNIP | |
| galactose | | chemical - endogenous mammalian | | | 0.0343 | GPD1,VDAC1,VDR | |
| mannose | | chemical - endogenous mammalian | | | 0.0408 | FOS,GPD1,MCL1 | |
| 3-deoxyglucosone | | chemical - endogenous mammalian | | | 0.049 | HBEGF | |
| growth factor | | group | | | 0.00387 | CLU,DUSP1,FOS,PTGS2 | 473 (18) |
| lactulose | | chemical - endogenous mammalian | | | 0.049 | CXCL8 | |
| allopregnanolone | | chemical - endogenous mammalian | | | 0.0144 | FOS,LGALS3,PTGS2 | |
| estramustine | | chemical drug | | | 0.049 | CYP1A1 | |
| sterol | | chemical - endogenous mammalian | | | 0.0461 | ACSL1,ACSS2,HMGCR,SCARB1 | |
| steroid hormone | | chemical - other | | | 0.0135 | CREBRF,CXCR4 | |
| oxysterol | | chemical - endogenous mammalian | | | 0.0478 | CAV1,CXCL8,HMGCR | |
| epiallopregnanolone | | chemical - endogenous mammalian | | | 0.00697 | LGALS3,PTGS2 | |
| kb-NB 142-70 | | chemical - kinase inhibitor | | | 0.00109 | DNAJB9,DNAJC3,ERN1 | |
| vinpocetine | | chemical drug | | | 0.0428 | CXCL3,CXCL8 | |
| pyocyanin | | chemical reagent | | | 0.0135 | CXCL8,NOX1 | |
| Fe3+ | | chemical - endogenous mammalian | | | 0.049 | HMGCR | |
| Cd2+ | | chemical toxicant | | | 0.0285 | CXCL3,FOS,PTGS2 | |
| Cr6+ | | chemical reagent | | | 0.0217 | MT2A,SLC30A1 | |
| V2+ | | chemical toxicant | | | 0.049 | CXCL8 | |
| D-alpha-hydroxyglutarate | | chemical - endogenous mammalian | | | 0.0135 | GLI1,RUNX2 | |
| CIL56 | | chemical reagent | | | 0.0135 | ALDH1A1,CDK6 | |
| RO5256390 | | chemical reagent | | | 0.049 | IRS2 | |

Supplementary Table 4. Upstream regulator analysis based gene set enrichment analysis (GSEA) of differentially expressed genes (DEGs) after ATRA+PDT compared to PDT

| Upstream Regulator | Expr Log Ratio | Molecule Type | Predicted Activation State | Activation z-score | p-value of overlap | Target Molecules in Dataset | Mechanistic Network |
|---|---|---|---|---|---|---|---|
| PC-SPES | | chemical drug | | | 0.000073 | ANXA1,B2M,CCNA2,CLU,DNAJB1,HLA-B,HLA-C,IMPDH1,IRF1,PPP1R15A,QSOX1,SCARB1,TIA1,TUBA4A,TUBB2A | |
| TCF | | group | | | 4.33E-10 | [illegible]LDH1A1,ARL4A,BMP4,CD44,CEACAM1,CYP24A1,DPEP1,ECM1,EGR1,FOS,LGALS3,MMP7,PCSK6,PTGS2,QPCT,RCN1,SERPINA1,SERPINA3,SERPINA5,TSC22D1,TSPAN8,UGC[illegible] | 448 (18) |
